# Functional innovation in the evolution of the calcium-dependent system of the eukaryotic endoplasmic reticulum

**DOI:** 10.1101/716472

**Authors:** Daniel E. Schäffer, Lakshminarayan M. Iyer, A. Maxwell Burroughs, L. Aravind

**Author notes:** **Correspondence:** L. Aravind. School of Computer Science, Carnegie Mellon University, Pittsburgh, PA, USA.

## Abstract

The origin of eukaryotes was marked by the emergence of several novel subcellular systems. One such is the calcium (Ca^2+^)-stores system of the endoplasmic reticulum, which profoundly influences diverse aspects of cellular function including signal transduction, motility, division, and biomineralization. We use comparative genomics and sensitive sequence and structure analyses to investigate the evolution of this system. Our findings reconstruct the core form of the Ca^2+^- stores system in the last eukaryotic common ancestor as having at least 15 proteins that constituted a basic system for facilitating both Ca^2+^ flux across endomembranes and Ca^2+^-dependent signaling. We present evidence that the key EF-hand Ca^2+^-binding components had their origins in a likely bacterial symbiont other than the mitochondrial progenitor, whereas the protein phosphatase subunit of the ancestral calcineurin complex was likely inherited from the asgardarchaeal progenitor of the stem eukaryote. This further points to the potential origin of the eukaryotes in a Ca^2+^-rich biomineralized environment such as stromatolites. We further show that throughout eukaryotic evolution there were several acquisitions from bacteria of key components of the Ca^2+^-stores system, even though no prokaryotic lineage possesses a comparable system. Further, using quantitative measures derived from comparative genomics we show that there were several rounds of lineage-specific gene expansions, innovations of novel gene families, and gene losses correlated with biological innovation such as the biomineralized molluscan shells, coccolithophores, and animal motility. The burst of innovation of new genes in animals included the wolframin protein associated with Wolfram syndrome in humans. We show for the first time that it contains previously unidentified Sel1, EF-hand, and OB-fold domains, which might have key roles in its biochemistry.

## 1 Introduction

The emergence of a conserved endomembrane system marks the seminal transition in cell structure that differentiates eukaryotes from their prokaryotic progenitors (Jekely, 2007). This event saw the emergence of a diversity of eukaryotic systems and organelles such as the nucleus, the endoplasmic reticulum (ER), vesicular trafficking, and several, novel signaling systems that are uniquely associated with this sub-cellular environment. A major eukaryotic innovation in this regard is the intracellular ER-dependent calcium (Ca^2+^)-stores system that regulates the cytosolic concentration of Ca^2+^ (Ashby and Tepikin, 2001). Although Ca^2+^ ions are maintained at a 4000 to 10,000-fold higher concentration in the endoplasmic reticulum (ER) lumen as compared to the cytoplasm (Woo et al., 2018), upon appropriate stimulus they are released into the cytoplasm by Ca^2+^-release channels such as the inositol trisphosphate receptors (IP3Rs) and the ryanodine receptors (RyRs). The process is then reversed and Ca^2+^ is pumped back into the ER by the ATP-dependent SERCA (sarcoplasmic/endoplasmic reticulum calcium ATPase) pumps, members of the P-type ATPase superfamily (Ashby and Tepikin, 2001; Altshuler et al., 2012). In addition to the above core components that mediate the flux of Ca^2+^ from and into the ER-dependent stores, several other proteins have been linked to the regulation of this process and transmission of Ca^2+^-dependent signals, including: 1) chaperones such as calreticulin, calnexin, and calsequestrin in the ER lumen (Kozlov et al., 2010); 2) diverse EF-hand proteins such as calmodulin (and its relatives), calcineurin B, and sorcin that bind Ca^2+^ and regulate the response to the Ca^2+^ flux (Denessiouk et al., 2014); 3) channel proteins such as the voltage-gated calcium channels (VGCC) and trimeric intracellular cation channels (TRIC) that influence the flow of Ca^2+^ (Lanner et al., 2010; Zhou et al., 2014); and 4) protein kinases (calcium/calmodulin-dependent kinases (CaMKs)) and protein phosphatases (calcineurin A) that mediate the Ca^2+^-dependent signaling response (Berridge, 2012). Together, the Ca^2+^-stores system and intracellular Ca^2+^-dependent signaling apparatus regulate a variety of cellular functions required for eukaryotic life, such as transcription, cellular motility, cell growth, stress response, and cell division (Clapham, 2007; Berridge, 2012; Krebs et al.).

Comparative evolutionary analyses of the proteins in the ER Ca^2+^-stores and-signaling system have revealed that some components were either present in the last eukaryotic common ancestor (LECA) (e.g. Calmodulin and SERCA) or derived early in the evolution of the eukaryotes (e.g. IP3R) (Nolan et al., 1994; Moreno and Docampo, 2003; Reiner et al., 2003; Prole and Taylor, 2011; Plattner and Verkhratsky, 2013; Verkhratsky and Parpura, 2014; Perez-Gordones et al., 2015). Other proteins show a patchier distribution in lineages outside of metazoans (e.g. calreticulin and calnexin) (Moreno and Docampo, 2003; Banerjee et al., 2007), or were reconstructed to have been derived in lineages closely related to the metazoans (e.g. RyR, which diverged from the ancestor of IP3R at the base of filozoans) (Alzayady et al., 2015). A substantial number of the components that have been studied in this system are primarily found in the metazoans, with no identifiable homologs outside of metazoa (Cai et al., 2015). Most studies have focused on animal proteins of these systems, highlighting the general lack of knowledge regarding the regulation of ER Ca^2+^-stores and the potential diversity in the regulatory systems present in other eukaryotes. To our knowledge, a systematic assessment of the evolutionary origins of the entire Ca^2+^-stores system and its regulatory components, as currently understood, has yet to be attempted.

Given our long-term interest in the origin and evolution of the eukaryotic subcellular systems, we conducted a comprehensive analysis of the core and regulatory components of the Ca^2+^-stores system, analyzing their known and predicted interactions and inferring the evolutionary depth of various components. We show that an ancient core of at least 15 protein families was already in place at the stem of the eukaryotic lineage. Of these, a subset of proteins is of recognizable bacterial ancestry, although there is no evidence of a bacterial Ca^2+^-stores system resembling those in eukaryotes. We also show that gene loss and lineage-specific expansions of these components shaped the system in different eukaryotic lineages, and sometimes corresponds to recognized adaptive features unique to particular organisms or lineages. Further, we conducted a systematic domain analysis of the proteins in the system, uncovering three novel unreported domains in the enigmatic wolframin protein. These provide further testable hypotheses on the functions of wolframin in the context of the Ca^2+^-stores system and in protecting cells against the response stresses that impinge on the ER.

## 2 Methods

### 2.1 Sequence analysis

Iterative sequence profile searches were performed using the PSI-BLAST program (RRID: SCR_001010) (Altschul et al., 1997) against a curated database of 236 eukaryotic proteomes retrieved from the National Center for Biotechnology Information (NCBI), with search parameters varying based on the query sequence (see Supplementary Figure S1) and composition. For building a curated dataset of eukaryotic proteomes, completely sequenced eukaryotic genomes were culled from Refseq and Genbank, and representative genomes (with a preference for reference genomes) were chosen from different phyletic groups using the eukaryotic phylogenetic tree as guide. If more than one genome was available for a species, we typically chose the one listed as a reference genome, or one that had the best assembly, or one with the most complete proteome set. The list of genomes is given in the supplementary data of the supplementary material. The program HHpred (RRID:SCR_010276) (Soding, 2005; Alva et al., 2016) was used for profile-profile comparisons. The BLASTCLUST program^1^ (RRID: SCR_016641) was used to cluster protein sequences based on BLAST similarity scores. Support for inclusion of a protein in an orthologous cluster involved reciprocal BLAST searches, conservation of domain architectures, and, when required, construction of phylogenetic trees with FastTree 2.1.3 (RRID: SCR_015501) (Price et al., 2010) with default parameters. The trees were visualized using FigTree^2^ (RRID: SCR_008515). Selected taxonomic absences were further investigated with targeted BLASTP (RRID: SCR_001010) (Altschul et al., 1990) and TBLASTN (RRID: SCR_011822) (Gertz et al., 2006) searches against NCBI’s non-redundant (nr) and nucleotide (nt) databases (Benson et al., 2013), respectively. Multiple sequence alignments were constructed using the MUSCLE (Edgar, 2004) and GISMO (Neuwald and Altschul, 2016) programs with default parameters. Alignments were manually adjusted using BLAST high-score pair (hsp) results as guides. Secondary structure predictions were performed with the Jpred 4 program (RRID: SCR_016504) with default settings (Drozdetskiy et al., 2015). EMBOSS (RRID: SCR_008493) pepwheel^3^ was used to generate renderings of amino acid positions on the circumference of an α-helix.

### 2.2 Protein network construction

Protein-protein interactions were extracted from published data sources, updating any outdated gene/protein names and making substantial efforts to disambiguate between paralogs (Supplementary Data). High-throughput/predicted protein-protein interactions were extracted from the FunCoup database (Ogris et al., 2018). Networks were visualized using the R-language implementations of the iGraph and qGraph packages. For network rendering, the Fruchterman-Reingold force-directed algorithm was used (Fruchterman and Reingold, 1991).

### 2.3 Comparative genome analyses

Quantitative analysis of phyletic patterns and paralog counts for different proteins/families were first obtained using a combination of sequence similarity searches as outlined above. We filtered the counts to exclude multiple identical sequences annotated with the same gene name, using the latest genome assemblies for each of these organisms as available in NCBI GenBank (RRID: SCR_002760) as anchors. Proteomes with identifiable quality issues were removed from downstream analyses (e.g. genomes containing sequences with ambiguous strain assignment), leaving counts for 216 organisms. These counts and their phyletic patterns defined two sets of vectors, namely the distribution by organism for a given protein and the complement of proteins for a given organism. These vectors for the protein families and organisms were used to compute the inter-protein or inter-organism Canberra distances (Lance and Williams, 1966), which is best suited for such integer data of the form of presences and absences. The Canberra distance between two vectors 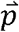 and 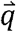 is defined as:

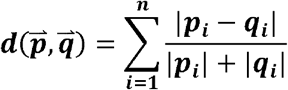

These distances were used to cluster the protein families and organisms through agglomerative hierarchical clustering using Ward’s method (Kaufman and Rousseeuw, 1990). The results were rendered as dendrograms. Ward’s method takes the distance between two clusters, A and B to be the amount by which the sum of squares from the center of the cluster will increase when they are merged. Ward’s method then tries to keep this growth as small as possible. It tends to merge smaller clusters that are at the same distance from each other as larger ones, a behavior useful in lumping “stragglers” in terms of both organisms and proteins with correlated phyletic patterns.

The protein complement vectors for organisms were also used to perform principal component analysis to detect spatial clustering upon reducing dimensionality. The variables were scaled to have unit variance for this analysis. Similarly, a linear discriminant analysis was performed on these vectors using representatives of the major eukaryotic evolutionary lineages (see Supplementary Table S1 for list) as the prior groups for classification. This was then used on our complete phyletic pattern data for classification of the organisms based on their protein complements.

Organism polydomain scores were calculated as follows: if ***c***(***o,p***) counts the number of paralogs of some protein domain family ***p*** in some organism ***o, P*** is the set of all proteins studied, and ***O*** is the set of all organisms studied, then the polydomain score for an organism ***o*** ∈ ***O*** is defined as:

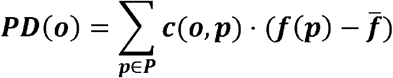

where 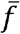 is the mean of *f*(*p*) for all proteins *p* ∈ *P*, and *f*(*p*) is defined as:

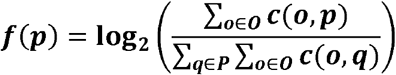

Computations and visualizations were performed using the R language.

## 3 Results

### 3.1 Protein-protein interaction network for the ER-dependent calcium stores regulatory/signaling apparatus

Metazoan Ca^2+^ stores have been extensively studied, leading to the identification of several proteins directly or indirectly involved in calcium transport or signaling and regulation of these processes. In order to apprehend the global structure of this system, we used published literature and the FunCoup database to derive a network of protein-protein interactions (PPIs) containing human genes that have roles in Ca^2+^ stores, centering the network on the three families of ER Ca^2+^ channels (SERCA, IP3R, RyR) (see Methods). The resulting network totaled 173 protein nodes and 761 interaction edges (Figure 1A, Supplementary Data). The distribution of the number of connections per node (degree distribution) in this network displays an inverse (rectangular hyperbolic) relationship (R^2^ = 0.88; Figure 1C). This is a notable departure from typical PPI networks which tend to show power-law degree distributions (Bader and Hogue, 2002; Rodrigues et al., 2011). To better understand this pattern, we studied its most densely connected subnetworks by searching for cliques, where every node is connected to every other node in the subnetwork. The largest cliques in this network have ten nodes. As the degree distribution graph shows an inflection around degree 6, we merged all cliques of size 6 or greater resulting in a subnetwork of 46 nodes comprising close to 50% of the edges of the overall network (Figure 1B). This suggests that the inverse relationship of the degree distribution is a consequence of the presence of a core of several highly connected nodes (6 or more edges), which is in contrast to other system-specific PPI networks showing a power-law degree distribution, as in the case of the ubiquitin network (Venancio et al., 2009).

**Figure 1.**
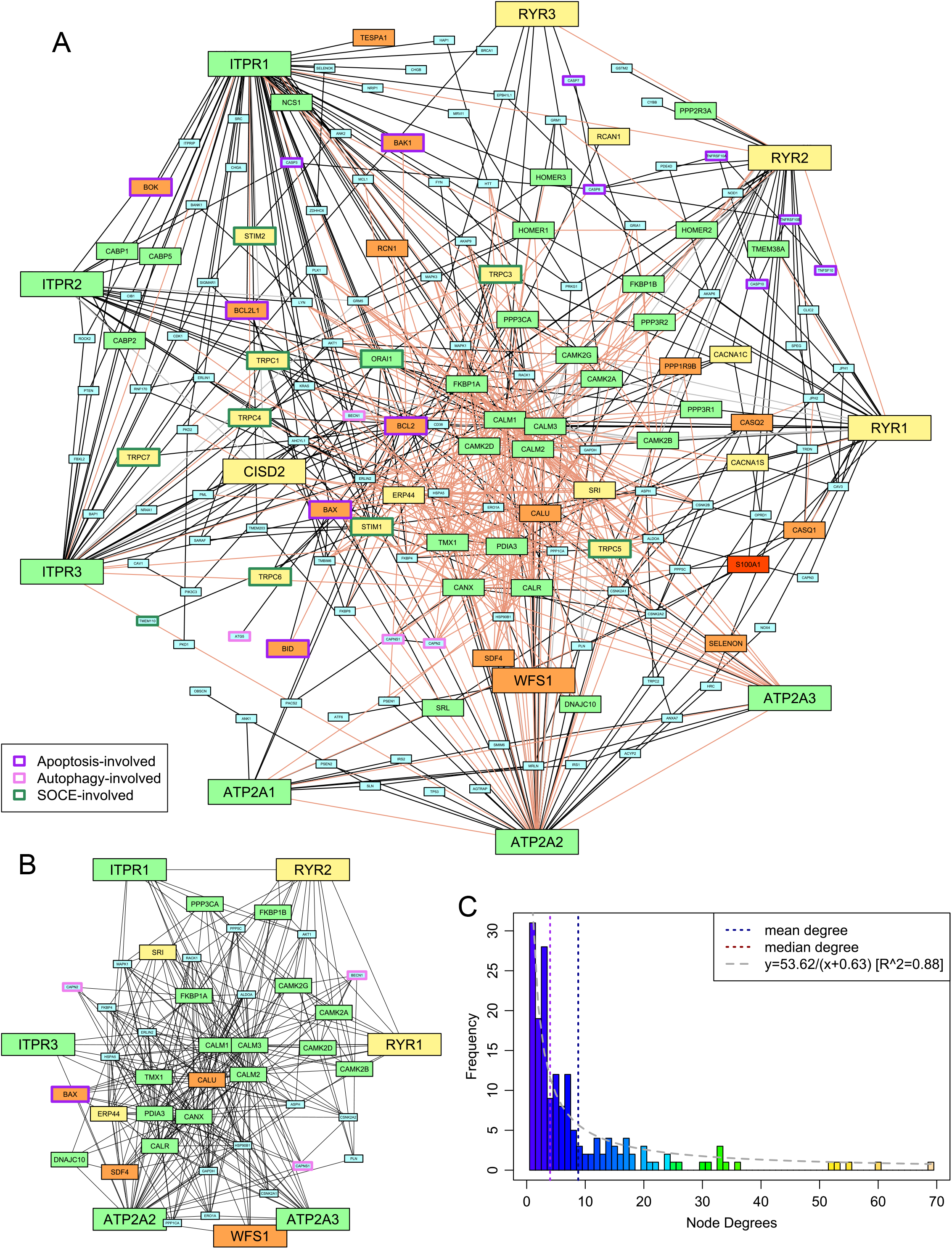
Protein-protein interaction network of the ER-dependent Ca^2+^-stores and-signaling system. (**A**) The network. Edges are classified as follows: interactions reported in the literature in which each partner gene either has no other paralog or has been specifically identified (solid black), reported interactions where the specific paralog for at least one of the two genes is unclear (dashed gray), and interactions found in FunCoup 4.0 with P ≤ 0.9 (light pink). Medium-and large-sized nodes represent the proteins whose phyletic distribution was selected for detailed analysis and are colored based on phylogeny as follows: pan-eukaryotic, green; metazoans and close relatives, yellow; metazoan-specific, orange; chordate-specific, red. Nodes labeled by HUGO nomenclature to capture paralog-specific interactions. The borders of nodes involved in selected intracellular systems are colored based on the key given in the lower left. (**B**) The highly-connected subnetwork. Edge coloring has no significance; nodes are colored and sized as in (A). (**C**) A histogram of the degrees of the nodes of the network shown in (A). A curve of best fit is shown in gray.

Analysis of the proteins in this highly connected sub-network suggests that the 46 proteins can be broadly classified into 5 groups: 1) the channels and ATP-dependent pumps which constitute the core Ca^2+^ transport system for ER stores; 2) EF-hand domain proteins such as calmodulin and sorcin that bind Ca^2+^ and consequently interact with and regulate the biochemistry of numerous other proteins; 3) proteins involved in folding and stability of other proteins, such as chaperones, redox proteins, and protein disulfide isomerases; 4) components of the protein phosphorylation response that is downstream of Ca^2+^ flux into the cytoplasm; and 5) proteins linking the network to other major functional systems, such as Bcl2, which is involved in the apoptosis response in animals, and beclin-1, which is involved in autophagy.

### 3.2 Phyletic distribution of key proteins and implications for the evolution of the ER Ca^2+^-stores and-signaling pathway

This densely connected sub-network invites questions about its evolutionary origin, particularly given that the wet-lab results that inform the connections are predominantly drawn from mammalian studies (see Methods). To understand better the emergence of this sub-network and the conservation of its nodes across eukaryotes, we systematically analyzed their phyletic patterns (Figure 2A, Supplementary Figure S1 and Data). A list of the 34 proteins and protein families studied, as well as their domain architectures, is in Supplementary Table S2. Further, Ward clustering analysis of the core components based on their phyletic patterns (see Methods) revealed the presence of 5 distinct clusters (Figure 2C). These clusters appear to have an evolutionary basis with distinct clusters accommodating proteins that could be inferred as having been in the LECA (e.g. cluster 1) and those that emerged in a metazoan-specific expansion of the ER Ca^2+^-stores system (e.g. clusters 4 and 5).

**Figure 2.**
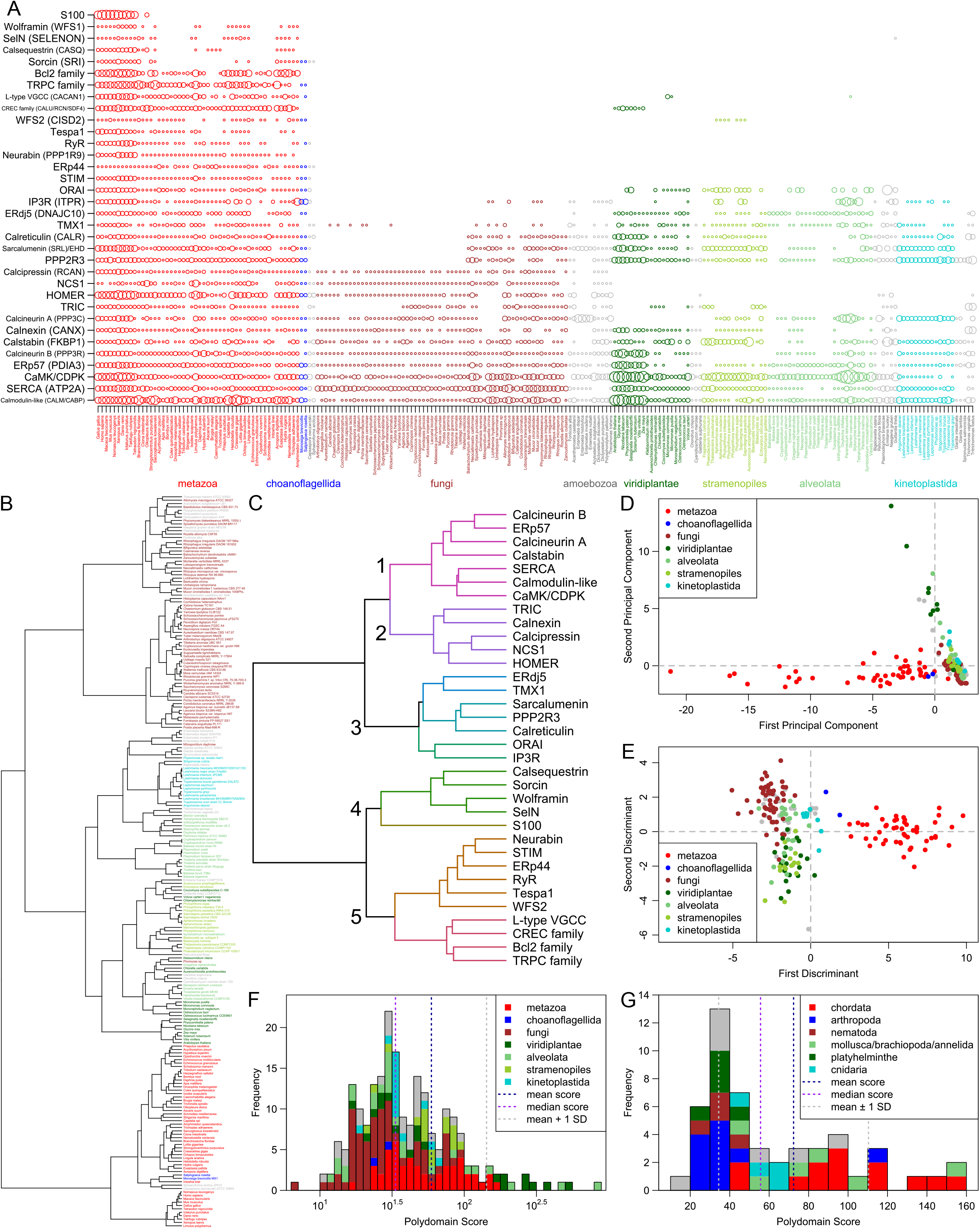
Comparative genome analyses. (**A**) A visualization of the number of paralogs of the 34 key proteins/protein families (vertical axis) across 216 eukaryotes (horizontal axis). Each circle represents the number of paralogs of one protein/family in one organism; the radius of each circle is scaled by the hyperbolic arcsine of the represented number of paralogs. The proteins and protein families shown can be broadly divided into two categories: those that are largely pan-eukaryotic (calmodulin to ORAI) and those that originate in the metazoans or their close relatives (STIM to S100). Eukaryotic clades are color-coded and labeled below the plot. Where needed, protein/protein family names are supplemented in parenthesis with human gene names from Figure 1, where node labels distinguish between paralogs. (**B**) A dendrogram of the 216 eukaryotes based on the counts of their paralogs of the 34 proteins/families. Tip labels are colored by clade using the same system as in panel A. (**C**) A dendrogram of the 34 proteins/families based on the counts of their paralogs. Five major clusters are numbered to their left. Branch coloring corresponds to major clusters. (**D-E**) Plots of the first two (**D**) principal components and (**E**) discriminants of each of the 216 eukaryotes resulting from principal component and linear discriminant analyses on the paralog count dataset. Color keys linking colors to high-level clades are given in the upper right and lower right, respectively, of the plots. (**F-G**) Histograms of the polydomain scores of (**F**) all 216 eukaryotes, shown on a base-10 log scale, and (**G**) 50 metazoans, shown on a linear scale. Contributions of some clades to each bar of the histograms are shown through coloring given by the keys in the upper right of the histograms.

#### 3.2.1 The core LECA complement of the Ca^2+^-stores system

At least 15 proteins of the ER Ca^2+^-stores and signaling system are found across all or most eukaryotic lineages, suggesting that they were present in the LECA. These proteins include key components of the dense sub-network, such as 1) the SERCA Ca^2+^ pumps; 2) the Ca^2+^-binding EF-hand proteins like calmodulin; 3) chaperones involved in protein folding that are mostly found in the ER and that sometimes act as either Ca^2+^ binding proteins (e.g. calreticulin, calnexin) or as regulators of other components of the Ca^2+^-stores and-signaling system (e.g. ERp57/PDIA3, calstabin, TMX1, 2+ ERdj5/DNAJC10); and 4) core enzymes of the Ca^2+^-dependent phosphorylation-based signaling system including the CaMKs and the calcineurin A phosphatase. This set of components is likely to have comprised the minimal ER-Ca^2+^-stores and -signaling system in the LECA and suggests that there were already sub-systems in place to mediate: 1) the dynamic transport of Ca^2+^ ions across the emerging eukaryotic intracellular membrane system and 2) the transmission of signals affecting a wide-range of subcellular processes based on the sensing of Ca^2+^ ions. 3) A potential stress response system that channels Ca-dependent signals to regulation of protein folding (Krebs et al.).

Notably, the IP3R channels are absent in lineages that are often considered the basal-most eukaryotes, namely the parabasalids and diplomonads; however, they are present in some other early-branching eukaryotes such as kinetoplastids (Cavalier-Smith et al.). Their absence in certain extant eukaryotes (Figure 2A) suggests that they can be dispensable, or that their role can be taken up by other channels in eukaryotes that lack them. A comparable phyletic pattern is also seen for certain other key components of the densely connected sub-network, namely the ERdj 5 and calnexin chaperones.

#### 3.2.2 Components with clearly identifiable prokaryotic origins

Deeper sequence-based homology searches revealed that at least three protein families of the ER Ca^2+^-stores system have a clearly-identifiable bacterial provenance, namely the P-type ATPase pump SERCA, sarcalumenin, and calmodulin and related EF-hand proteins. Phylogenetic analyses suggest that the P-type ATPases SERCA and plasma membrane calcium-transporting ATPase (PMCA) were both present in the LECA. They are most closely-related to bacterial P-type ATPases (Plattner and Verkhratsky, 2015), which commonly associate with transporters (e.g. Na^+^-Ca^2+^ antiporters), ion exchangers (Na^+^-H^+^ exchangers), permeases, and other distinct P-type ATPases in conserved gene-neighborhoods (Supplementary Figure S2) suggesting a role in maintenance of ionic homeostasis even in bacteria.

The GTPases EHD and sarcalumenin, whose GTPase domains belong to the dynamin family (Leipe et al., 2002), also show clear bacterial origins based on their phyletic patterns and phylogenetic affinities. The closest bacterial homologs possess a pair of transmembrane helices C-terminal to the GTPase domain, suggesting a possible role in membrane dynamics (Figure 3B, Supplementary Data). Phylogenetic trees show that although they are related to the dynamins, the progenitor of the eukaryotic sarcalumenin and EHD was acquired independently of the dynamins via a separate transfer from a proteobacterial lineage early in eukaryotic evolution (Figure 3A) (Leipe et al., 2002). This gave rise to the EH-domain-containing EHD clade of GTPases, which are involved in regulating vesicular trafficking and membrane/Golgi reorganization. A further secondary transfer, likely from the kinetoplastid lineage to the metazoans, gave rise to sarcalumenin proper, which has characterized roles in the Ca^2+^-stores system (Figure 3A). This raises the possibility that in other eukaryotes, EHD performs additional roles in the Ca^2+^-stores system overlapping with metazoan sarcalumenin.

**Figure 3.**
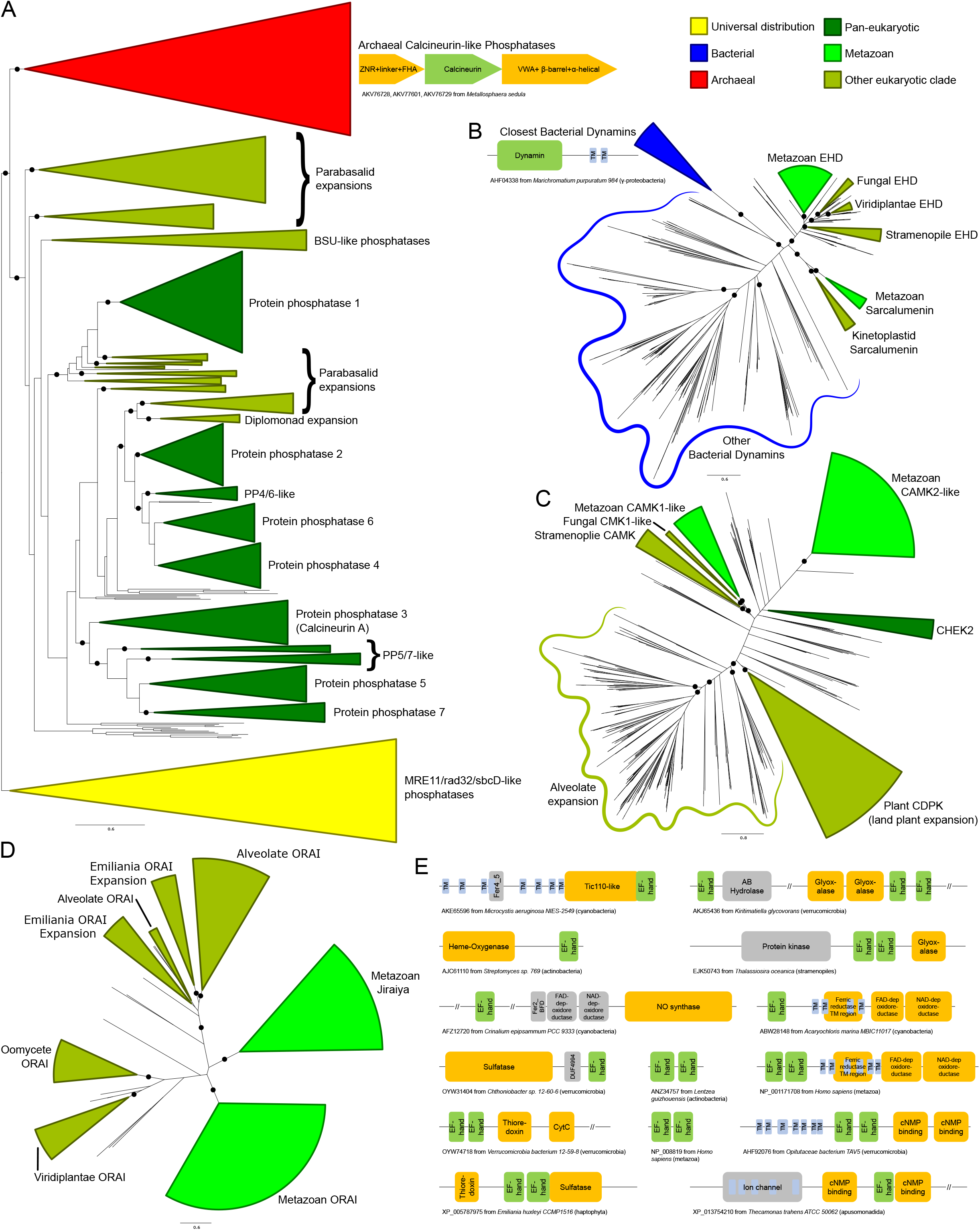
(**A-D**) Stylized phylogenetic trees showing (**A**) sarcalumenin, EHD, and related bacterial dynamins; (**B**) a partial tree of the calcineurin-like superfamily, containing the classical calcineurin A phosphatases with their immediate eukaryotic relatives and newly-recognized archaeal orthologs, along with the related MRE11/rad32/sbcD-like phosphoesterases as an outgroup; (**C**) the expansions of the calcium/calmodulin-dependent kinase in eukaryotes; and (**D**) ORAI and Jiraiya. Collapsed groups are colored as follows: universal distribution, yellow; bacterial, blue; archaeal, red; pan-eukaryotic, dark green; metazoan, light green; restricted to non-metazoan eukaryotes, blue-green. Branches with bootstrap support of greater than 85% are marked with a black circle. Genome contextual associations pertinent to a given clade are provided within context of trees. Conserved gene neighborhoods are depicted as boxed arrows and protein domain architectures as boxes linked in the same polypeptide. The trees are also provided in Newick format in the Supplementary Data. (**E**) Domain architectures of proteins containing calmodulin-like EF-hands. Each EF-hand is a dyad of EF repeats. Long (>200 residue) regions without an annotated domain are collapsed using “//”.

Our analysis revealed that the bacterial calmodulin-like EF-hands show a great diversity of domain-architectural associations. Versions closest to the eukaryotic ones are found in actinobacteria, cyanobacteria, proteobacteria, and verrucomicrobiae. These prokaryotic calmodulin-like EF-hand domains are found fused to a variety of other domains (Figure 3E, Supplementary Data), such as a 7-transmembrane domain (7TM) (cyanobacteria and to a lesser extent in actinobacteria), the prokaryotic Tic110-like α-helical domain (cyanobacteria), heme-oxygenases (actinobacteria), the nitric oxide synthase and NADPH oxidase with ferredoxin and nucleotide-binding domains (cyanobacteria and δ-proteobacteria), as well as cNMP-binding, thioredoxin, cytochrome, and sulfatase domains (verrucomicrobiae). Comparable architectural associations with several of these domains, such as the fusions to the sulfatases and the redox-regulator domains thioredoxin, cytochromes, glyoxylases, and NADPH oxidases, are also observed in eukaryotes. However, our analysis showed that these fusions appear to be independently derived in the two superkingdoms. These diverse associations suggest that even in bacteria the calmodulin-like EF-hands function in the context of membrane-associated signaling and redox reactions, possibly regulated in a Ca^2+^-flux-dependent manner (Zhou et al., 2006; Dominguez et al., 2015). We infer that a version of these was transferred to the stem eukaryote, probably from the cyanobacteria or the actinobacteria, and had already triplicated by the LECA, giving rise to the ancestral versions of calmodulin and calcineurin B, which function as part of the Ca^2+^-stores system, and the centrins, which were recruited for a eukaryote-specific role in cell division is association with the centrosome (Dantas et al., 2012).

Among components with indirect regulatory roles in the ER Ca^2+^-dependent system is the peptidyl-prolyl cis-trans isomerase (PPIase) calstabin (FKBP1A/B in humans), which is a member of the FKBP family of PPIase chaperones. Phyletic pattern analysis indicates that calstabin was present in the LECA (Figure 2A). Eukaryotic PPIases are more similar to bacterial PPIases (Supplementary Data); hence, the ancestral eukaryotic PPIase was likely acquired from the alphaproteobacterial mitochondrial progenitor. This is also supported by the evidence from extant pathogenic/symbiotic bacteria wherein the bacterial FKBP-like PPIases play a role in establishing associations with eukaryotic hosts (Unal and Steinert, 2014). In the stem eukaryotes, the ancestral PPIase acquired from the bacterial source underwent a large radiation resulting in diverse PPIases that probably went hand-in-hand with the eukaryotic expansion of low-complexity proteins with potential substrate prolines. Thus, eukaryotes acquired a wide range of substrate proteins in several eukaryote-specific pathways and function in several cellular compartments (Trandinh et al., 1992). Calstabin is one of the paralogs that arose as part of this radiation and appears to have been dedicated to the ER Ca^2+^-stores system.

Beyond the above-mentioned, several other components inferred to be part of the LECA complement of the ER Ca^2+^-stores system are likely of bacterial origin. However, they do not have obvious bacterial orthologs and might have diverged considerably from their bacterial precursors in the stem eukaryote itself. These include the TRIC-like channels (Silverio and Saier, 2011), the CaMKs, and the thioredoxin domains found in ERp57, ERdj5, and TMX1.

In contrast to the several components of bacterial provenance, the large eukaryotic assemblage of calcineurin-like protein phosphatases, which includes the Ca^2+^-stores regulator calcineurin A, are specifically related to an archaeal clade to the exclusion of all other members of the superfamily (Figure 3A, Supplementary Data). Notably, these close relatives are present in several Asgardarchaea, suggesting the eukaryotes may have directly inherited the ancestral version of these phosphoesterases from their archaeal progenitor (Zaremba-Niedzwiedzka et al., 2017). Strikingly, the archaeal calcineurin-like phosphatases occur in a conserved operon (Figure 3A) with genes for two others proteins, one combining a zinc ribbon fused to a phosphopeptide-recognition FHA domain and the second with a vWA domain fused to a β-barrel-like domain. This supports a similar role for these archaeal calcineurin-like phosphatases to their eukaryotic counterparts in transducing a signal through dephosphorylation of a protein-substrate.

Thus, the core, ancestrally-conserved components of the ER-dependent stores system predominantly descend from the bacteria, although at least one component was inherited from the archaea. While the roles for some of these domains in possible bacterial Ca^2+^-dependent systems are apparent, there is no evidence that these versions function in a coordinated fashion in any single bacterial species or clade. Further, it is also clear that there was likely more than one bacterial source for the proteins: components such as SERCA and calstabin, whose closest relatives are proteobacterial, are likely to have been acquired from the mitochondrial ancestor, whereas calmodulin is likely to have been derived from a cyanobacterium or actinobacterium. Thus, the ER-dependent Ca^2+^-stores network was assembled in the stem eukaryote from diverse components drawn from different prokaryotic lineages. This assembly of the system in eukaryotes is strikingly illustrated by the case of the calcineurin complex. Here, the protein phosphatase component has a clear-cut origin from the archaeal precursor of the eukaryotes, whereas the Ca^2+^-binding calcineurin B component descends from a bacterial source. It was the combination of these proteins with very distinct ancestries that allowed the emergence of a Ca^2+^-signaling system.

#### 3.2.3 Lineage-specific expansions and gene loss shape the Ca2+ response across eukaryotes

In order to obtain some insights regarding the major developments in the evolution of the eukaryotic Ca^2+^-stores system, we systematically assembled protein complement vectors for each of the organisms in the curated proteome dataset (see Methods). After computing the pairwise Canberra distance between these vectors, we performed clustering using the Ward algorithm (see Methods). The resulting clusters recapitulate aspects of eukaryote phylogeny, with animals, fungi, plants, kinetoplastids and apicomplexans forming distinct monotypic clusters (Figure 2B). These observations together indicate that most major eukaryotic lineages have likely evolved specific component around the core Ca^2+^-stores system inherited from the LECA that distinguish them from related lineages. We then used principal component and linear discriminant analysis (see Methods) on these vectors to obtain a global quantitative view of the diversification of the Ca^2+^-stores system. Plotting the first two principal components/discriminants reveals discernable spatial separation of the metazoans from their nearest sister group, the fungi (Figure 2D-E). Further, diverse photosynthetic eukaryotes tended to be spatially colocalized. These observations suggest that certain accretions to the core Ca^2+^-stores system probably occurred alongside unique functional developments such as motility and multicellularity (metazoa) and autotrophy (photosynthetic lineages). Hence, investigations on individual lineages are likely to unearth novel, lineage-specific regulatory devices, which potentially reflect adaptations to their respective environments and provide hints about the biological contexts in which they were found.

We further investigated these potential lineage-specific developments by defining a measure, the polydomain score (see Methods), which captures the overall amplification of the Ca^2+^-stores system of an organism in terms of the contribution of the different constituent protein families of the system (Figure 2F). While PCA and LDA help identify the overall tendencies among organisms in terms of the system under consideration, the polydomain score is better-suited for identifying specific unusual features in terms of individual protein families/domains in particular organisms or clades. This score allowed us to capture some of the key developments in the reconstructed Ca^2+^-stores system of particular organisms and lineages. Notably, this led to the identification some of the distinctions between Ca^2+^-regulatory systems that may reflect differential strategies for the incorporation of calcium into exo/endo-skeletal structures in eukaryotes.

Comparison of polydomain scores within metazoa (Figure 2G) showed a striking reduction of the Ca^2+^-stores system in arthropods as well as in some molluscs and other marine lineages, consistent with their distinct grouping in the PCA/LDA plots. This appears to be generally related to the lower dependence on biomineralized structures in these organisms – arthropods utilize chitin as the primary exoskeletal material as opposed to calcareous structures, and the molluscan lineages showing this pattern have lost their calcified shells (e.g. octopi). Similarly, fungi, which use chitin as a central cell wall component, show lower polydomain scores relative to sister eukaryotic lineages. Conversely, we noted elevated polydomain scores in molluscs with calcareous shells. Likewise, ciliates, which have a evolved a distinct extension to the ancestral eukaryotic Ca^2+^-stores system in the form of the alveoli (Plattner, 2017), also have elevated polydomain scores. Ca^2+^-based signaling has been shown to be important for defensive trichocyst exocytosis, ciliary action, and other pathways in ciliates (Plattner, 2015). These observations suggest a linkage between strategies for structural and signaling usage of calcium and the regulation of Ca^2+^ distribution between intracellular compartments.

In qualitative terms, the distinct spatial positioning of different eukaryotic lineages in the PCA/LDA plots and the difference in their polydomain scores could be explained on the basis of lineage-specific expansions (LSEs), gene losses, and domain architectural diversity between lineages (Lespinet et al., 2002). We identified specific gains and losses based on inferred ancestral protein complements (Supplementary Figure S1). One of the most frequent proteins displaying LSEs is the CaMK: expansions have occurred independently in animals, plants, oomycetes and alveolates (Figure 3C, Supplementary Data). Ciliates in particular show a dramatic expansion of the family with over one hundred copies in *Paramecium* and *Stentor*. Other proteins that show LSEs in specific lineages (Supplementary Data) include calmodulin (in metazoans, certain fungal lineages, and plants), calstabin (independently in different stramenopile lineages and haptophytes), PP2R3-like proteins (in kinetoplastids and Trichomonas), calcineurin A (in ciliates and *Entamoeba),* calnexins (in *Trichomonas* and diatoms), ORAI (in *Emiliania,* Figure 3D), and SERCA (in certain fungi). The metazoans also display LSEs for several protein families that appear to have specifically emerged in metazoa (see below).

Some of these LSEs have clear biological correlates that were also suggested by the polydomain scores. For example, LSEs of the calmodulin family in the shelled molluscs (20-46 copies) might correspond to or be involved in the regulation of Ca^2+^ concentration during the biomineralization processes involved in the formation of calcareous shells. Concomitant with this expansion, in shelled molluscs several transporters, including P-type ATPases, voltage-gated channels, and ion exchangers, have been adapted for the transportation of Ca^2+^ ions for shell formation (Sillanpaa et al., 2018). Similarly, the development of the distinct set of Ca^2+^ stores in the form of the alveoli, which play several key ciliate-specific roles (Plattner, 2015; 2017), might explain the expansions in these organisms. Thus, the very large LSEs of the CaMK and calcineurin A-like phosphoesterase likely reflect the many diverse pathways into which Ca^2+^-based signaling is incorporated in these organisms. Additionally, experimental evidence shows that some pan-eukaryotic members of the ER Ca^2+^ stores system have acquired additional roles in ciliates; for example, SERCA-like Ca^2+^-pumps are also in the membranes of *Paramecium* alveolar calcium stores (Plattner et al., 1999).

ORAI family LSEs in *Emiliania* might correspond to the need to regulate Ca^2+^ transport for mineralizing the calcareous coccoliths (Yin et al., 2018). Further, calcium for coccolith formation may also supplied in part by an ER-membrane-localized Ca^2+^/H^+^ exchanger, as well as by increased activity of SERCA and a plasma membrane Ca^2+^/H^+^ exchanger (Mackinder et al., 2011). Here, as in ciliates, these expansions might have accompanied the emergence of a distinct Ca^2+^-stores system associated with the nuclear envelope (an extension of the ER), which is close in proximity to the envelope of the coccolith and the site of biomineralization (Brownlee et al., 2015).

In contrast, we observed extensive gene losses of proteins including ERdj5, sarcalumenin, IP3R, and ORAI across several or all fungi and losses of at least 10 conserved proteins in *Entamoeba* (Figure 2A). However, experimental studies have shown that both fungi and *Entamoeba* have Ca^2+^ transients and associated signaling systems (Makioka et al., 2002; Kim et al., 2012); hence, despite these losses, the evidence favors these organisms retaining at least a limited Ca^2+^-store-dependent signaling network. The CREC (calumenin, reticulocalbin, and Cab45) family displays an unusual phyletic pattern of particular note, being found only in metazoans and land plants (Figure 2A). Barring the unusual possibility of lateral transfer between these two lineages, this would imply extensive loss of these proteins across most other major eukaryotic lineages. Within metazoa, several proteins display unusual loss patterns, including sorcin, calsequestrin, and SelN (Figure 2A).

Notably, the land plants and their sister group the green algae have entirely lost ancient core Ca^2+^- dependent signaling proteins such as calcineurin A and HOMER. The IP3R channels are retained by the chlorophytes but lost entirely by the land plants, while the ORAI channels are present in the basal land plants *Physcomitrella* and *Selaginella* but are not observed in crown land plants (Edel et al., 2017). Land plants have also lost the TRIC channels, but the basal streptophyte *Chara braunii* retains a copy. The differential retention and expansion of various ancestral Ca^2+^ components (Figure 2A) in the land plants might be seen as adaptations to a sessile lifestyle no longer requiring Ca^2+^-signaling components associated with active cell or organismal motility. However, the paradoxical LSEs of the calmodulin-like and EF-hand-fused CaMK-like proteins (CDPK family) (Edel et al., 2017) in land plants relative to their chlorophyte sister-group (Figure 2A, 3C). These might have emerged as part of Ca^2+^ sequestration and signaling mechanisms relevant in the Ca^2+^-poor freshwater ecosystems, wherein the land plants originated from algal progenitors (Delwiche and Cooper, 2015).

Little in the way of domain architectural diversity is observed in the core conserved proteins of the animal Ca^2+^-stores system. A notable exception is seen in the CaMKs: metazoan CaMKs show a higher architectural complexity via their fusion to several distinct globular domains. These domains act as adaptors in interactions with Ca^2+^-binding regulatory domains, effectively broadening the total range of signaling pathways in which the CaMKs participate in metazoa (Wang et al., 2015). In contrast, land plants display lower architectural complexity with CaMK orthologs directly fused to Ca^2+^-binding domains (Klimecka and Muszynska, 2007).

#### 3.2.4 Accretion of novel Ca2+ signaling pathway components in the Metazoa

Case by case examination revealed that the distinct position of the metazoans in the PCA/LDA plots relative to other eukaryotes (Figure 2D-E) is due to a major accretion of novel signaling and flux-related Ca^2+^-stores components at the base of the metazoan lineage (Figure 2A). Our analyses suggest three distinct origins of these proteins: 1) several emerged as paralogs of domains already present in the LECA and functioning in the context of Ca^2+^-signaling, including the EF-hand domains found in the STIM1/2, calumenin, SelN, S100, sorcin, and NCS1 proteins, the thioredoxin domain found in the ERp44 and calsequestrin proteins, and the RyR channel proteins. 2) Proteins or component domains which are involved in a broader range of functions but have been recruited to roles in Ca^2+^-stores regulation specifically in the animal lineages, such as the SAM domain in the STIM1/2 proteins, the PDZ domain in the neurabin protein, and the TRPC channels. 3) Proteins containing domains which appear to be either novel metazoan innovations, such as the KRAP domain of the Tespa1 protein, or whose origins have yet to be traced, such as VGCC and wolframin. Further, the origin of certain metazoan-specific proteins such as phospholamban, which inhibits the SERCA ATPase, remain difficult to trace because of their small size and highly-biased composition as membrane proteins. Inspection of the neighbors of these proteins in the constructed network suggests that in almost all instances, these proteins were added to the Ca^2+^ signaling network via interactions with one or more of the ancient components of the system (Figure 1A).

This sudden accretion of Ca^2+^-stores components likely coincided with emergence of well-studied aspects of differentiated metazoan tissues, such as muscle contraction and neurotransmitter release (Zucchi and Ronca-Testoni, 1997; Clapham, 2007). Other additions to the network likely arose via interface with other pathways like apoptosis and autophagy in the context of ER stress response (Smaili et al., 2013). Notable in this regard is the metazoan-specific Bcl2 family of membrane-associated proteins associated with regulation of apoptosis. They appear to have been derived via rapid divergence in metazoans from pore-forming toxin domains of ultimately bacterial provenance (Peng et al., 2009), which are found in pathogenic bacteria and fungi (Aravind et al., 2012).

Still other proteins were recruited to the system as part of newly-emergent regulatory subnetworks, such as the Store-Operated Calcium Entry (SOCE) pathway for re-filling the ER from extracellular Ca^2+^ stores (Prakriya and Lewis, 2015; Ong et al., 2016). Of the proteins identified in this pathway, the ORAI Ca^2+^ channels are present in the early-branching kinetoplastid lineage but were lost in several later-branching lineages (Figure 2A), while other components like the ORAI-regulating STIM1/2 proteins and the TRPC channels emerged around the metazoan accretion event, suggesting the core SOCE pathway came together at or near the base of the animal lineage. In course of this analysis, we identified a common origin for the ORAI and Jiraiya/TMEM221 ER channels, the latter of which are characterized in BMP signaling (Aramaki et al., 2010). The Jiraiya channels are observed in animals but are absent in earlier-branching eukaryotic lineages, suggesting they emerged from a duplication of an ORAI channel early in metazoan evolution (Figure 3D, Supplementary Data). Jiraiya channel domains lack the Ca^2+^ binding residues seen in ORAI channels, suggesting they are unlikely to directly bind Ca^2+^. However, it is possible that Jiraiya/TMEM221 physically associates with components of the Ca^2+^-stores system to regulate them.

### 3.3 Domain architectural anatomy and functional analysis of the wolframin protein

As noted above, one of the uniquely metazoan proteins in the Ca^2+^-stores system is wolframin, a transmembrane protein localized to the ER membrane (Hildebrand et al., 2008; Rigoli et al., 2011; Qian et al., 2015) (Figure 2A, Supplementary Data). Wolframin, along with the structurally-unrelated and more widely phyletically distributed (Figure 2A) Wolfram syndrome 2 (WFS2) protein, is implicated in Wolfram syndrome (Inoue et al., 1998; Strom et al., 1998; Amr et al., 2007; Urano, 2016). Experimental studies have attributed biological roles to unannotated regions upstream and downstream of the TM region in wolframin, and our analyses revealed four uncharacterized globular regions therein (Figure 4A). The cytosolic N-terminal region was identified through iterative database searches (see Methods) as containing Sel1-like repeats (SLRs; query: NP_005996, hit: OYV16035, iteration 2, e-value: 5×10^-16^), which are α-helical superstructure forming repeats structurally comparable to the tetratricopeptide repeats (TPRs) (Ponting et al., 1999; Karpenahalli et al., 2007). Profile-profile searches (see Methods) unified the remaining two globular regions respectively with various EF-hand domains (e.g., query: XP_011608878, hit: 1SNL_A, p-value 2.3×10^-4^) and OB fold-containing domains (e.g. query: XP_017330582, hit: 2FXQ_A, p-value: 1.3×10^-4^).

**Figure 4.**
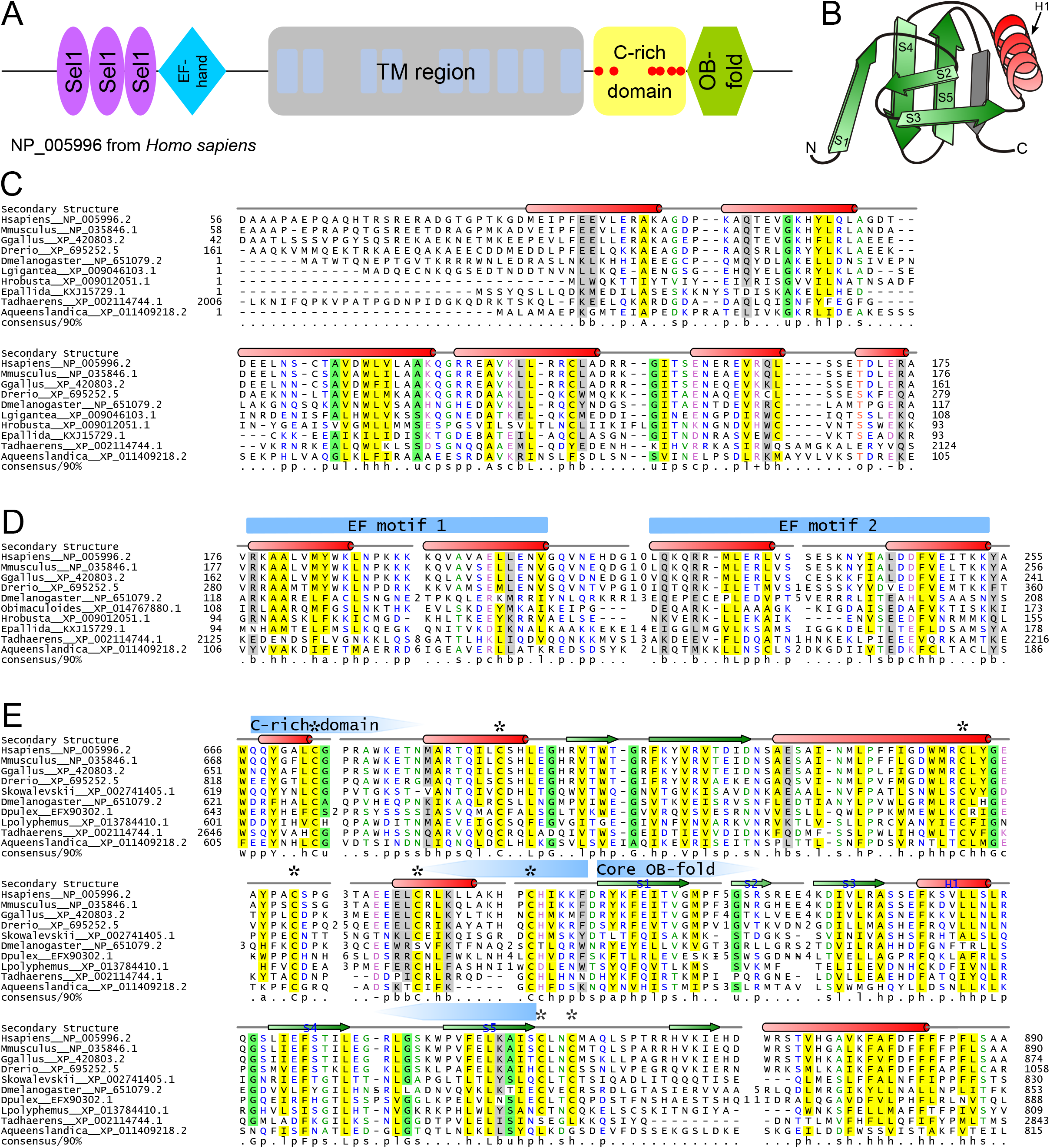
Structural and sequence overview of wolframin. (A) Domain architecture of wolframin. (B) Topology diagram of the wolframin OB-fold. The labeled secondary structure elements correspond with the labeled secondary structure elements and coloring in the OB-fold alignment in part E. The β-strand shaded in gray is a possible sixth strand stacking with the core OB-fold domain barrel. (C-E) Multiple sequence alignments of the (C) Sel1-like repeats, (**D**) EF-hand, (**E**) and cysteine-rich and OB-fold domains of wolframin. Sequences are labeled to the left of the alignments by organism abbreviation (see Supplementary Table S3) and NCBI GenBank accession number. Secondary structure is provided above the alignments; green arrows represent strands and red cylinders represent helices. The two EF motifs are marked with blue arrows above the secondary structure line. The boundaries of the core OB-fold and the cysteine-rich region (E) are marked with pentagonal arrows above the secondary structure line and pointing towards the center of the domain. The six secondary structure elements that comprise the OB-fold are labeled with blue text in the secondary structure line. Conserved cysteines are marked with an asterisk. A 90% consensus line is provided below the alignments; the coloring and abbreviations used are: h (hydrophobic), l (aliphatic), and a (aromatic) are shown on a yellow background; o (alcohol) is shown in salmon font; p (polar) is shown in blue font; + (positively charged), - (negatively charged), and c (charged) are shown in pink font; s (small) is shown in green font; u (tiny) is shown on a green background; b (big) is shaded gray.

The wolframin SLR possess several conserved residues seen in the classical SLRs (Figure 4C, Supplementary Table S4)(Mittl and Schneider-Brachert, 2007), and additionally display a truncated first loop when compared to known SLRs (Figure 4C). The SLRs of wolframin appear most closely-related to bacterial versions, suggesting a possible horizontal transfer at the base of animals (Ponting et al., 1999); however, the short length and rapid divergence of such repeats complicates definitive ascertainment of such evolutionary relationships. SLRs and related α/α repeats are often involved in coordinating interactions within protein complexes (Karpenahalli et al., 2007; Mittl and Schneider-Brachert, 2007) and have been characterized specifically in ER stress and misfolded protein degradation responses (Jeong et al., 2016) and in mediating interactions of membrane-associated protein complexes (Mittl and Schneider-Brachert, 2007). As the wolframin SLRs roughly correspond to an experimentally-determined calmodulin-binding region (Yurimoto et al., 2009), they might specifically mediate that protein interaction.

The EF-hand region of wolframin contains the two characteristic copies of the bihelical repeat that form the basic EF-hand unit. However, it lacks the well-characterized Ca^2+^-binding DxDxDG motif or any comparable residue conservation (Figure 4D) (Gifford et al., 2007; Denessiouk et al., 2014). It also features striking loop length variability in between the two helix-loop-helix motifs (Figure 4D). Such “inactive” EF-hand units typically dimerize with other EF-hand proteins (Kawasaki et al., 1998; Gifford et al., 2007), suggesting that wolframin could be self-dimerizing or that its EF-hand could interact at the ER membrane with calmodulin and/or a distinct EF-hand protein.

The C-terminal OB fold domain (Figure 4B,E) lacks the conserved polar residues typical of nucleic acid-binding OB-fold domains (Watson et al., 2007; Guardino et al., 2009). Further, the localization of this region to the ER lumen suggests that it is unlikely to be involved in nucleic acid binding (Arcus, 2002; Flynn and Zou, 2010; Krishna et al., 2010). Alternatively, this OB-fold domain could mediate protein-protein interactions as has been observed for other members of the fold (Flynn and Zou, 2010) and is also consistent with previous experimental studies implicating this region of wolframin in binding the pre-folded form of ATP1B1 (Zatyka et al., 2008). Strikingly, in the region N-terminal to the OB fold domain we observe a further globular region containing a set of six absolutely-conserved cysteines (Figure 4E). This region is not unifiable with any known domains and could conceivably represent an extension to the core OB fold domain. These conserved cysteine residues could contribute to disulfide-bond-mediated cross-linking, a well-studied regulatory mechanism of Ca^2+^-stores regulation (Ushioda et al., 2016). Additionally, C-terminal to the OB-fold, wolframin has a hydrophobic helix that might be involved in intra-or inter-molecular interactions (Figure 4E).

The wolframin TM region, located in the central region of the protein (Figure 4A), consists of nine transmembrane helices (Hofmann et al., 2003; Rigoli et al., 2011; Qian et al., 2015). Despite extensive studies on the TM region, its precise role in affecting Ca^2+^ flow across the ER membrane remains the subject of some debate (Osman et al., 2003; Aloi et al., 2012; Zatyka et al., 2015; Cagalinec et al., 2016). Inspection of a multiple sequence alignment of the wolframin transmembrane region (Supplementary Figure S3) revealed a concentration of polar residues which are spatially alignable in helices 4 and 5 (Supplementary Figure S4). This is reminiscent of membrane associated polar residue configurations seen in proteins that allow transmembrane flux of ions. Hence, it would be of interest to investigate if these residues might play a role in ion transport by wolframin.

## 4 Discussion

### 4.1 Evolutionary and functional considerations

#### 4.1.1 Early and later landmarks in the evolution of the ER Ca^2+^-stores system

The ER Ca^2+^-stores system displays several parallels in its evolutionary history to other endomembrane-dependent systems such as the nuclear membrane and vesicular trafficking systems (Mans et al., 2004; Jekely, 2008). Like in the case of these systems, dedicated ER Ca^2+^-stores systems are absent in the prokaryotes, despite the presence of Ca^2+^ transients and Ca^2+^-dependent signaling pathways in them (Dominguez et al., 2015). However, several of the more ancient individual components of eukaryotic Ca^2+^-stores systems are of clear-cut prokaryotic origin. Notably, we show here that not all of these proteins originated from the proto-mitochondrion; notably, calmodulin is of likely cyanobacterial or actinobacterial provenance and the calcineurin-like phosphoesterases originally descended from the archaea. The former observation adds to accumulating evidence of LECA acquiring bacterial contributions from non-α-proteobacterial lineages. It is therefore possible that LECA had a more extensive set of associated symbionts than what was fixed as the mitochondrion in eukaryotic evolution (Huang and Gogarten; Burroughs et al., 2017; Verma et al., 2018).

The cyanobacterial/actinobacterial origin of calmodulin and the role for the closely-related and early-branching centrin family of EF-hand proteins in microtubule dynamics during cell division suggest that the LECA already had a strong Ca^2+^ dependency. This raises the possibility that the prokaryotic ancestors of the eukaryotes might have existed in a calcium-rich environment such as the biomineralized structures (e.g. stromatolites) formed by cyanobacteria (Bosak et al., 2013). This is consistent with the diversity of domain architectures for calmodulin-like proteins with ramifications into various functional systems in the cyanobacteria that we reported here. This diversified pool of Ca^2+^-binding domains could have contributed raw materials needed during the initial emergence of Ca^2+^ flux-based signaling and regulation across endomembranes in eukaryotes. The newly emerged intracellular Ca^2+^ gradients were likely fixed by the myriad advantages it bestowed in the stem eukaryotes, including increased signaling capacity and a regulatory mechanism for processes like growth/proliferation, secretion, and motility.

Figure 5 shows a summary of key acquisitions and losses of Ca^2+^-stores proteins during eukaryotic evolution, placed onto a simplified model of eukaryotic evolution. In inferring ancestral states, and thereby gains and losses, we followed the (relatively) certain contours of the eukaryotic tree (see also Supplementary Figure S1). For example, we assumed that the root of eukaryotes lies in the excavates and that the SAR (stramenopile/alveolate/rhizarian) group is monophyletic. In general, we strove to make the least number of assumptions possible; therefore, many of our estimates – for example of the number of Ca^2+^-stores proteins in the LECA – are lower bounds and conservative. By these criteria, we infer that the ancestral eukaryotic ER-dependent Ca^2+^-stores system likely consisted of a combination of the SERCA pump, a cation channel, EF-hand-containing proteins, and phosphorylation enzymes (Figure 5). Early in eukaryotic evolution, chaperone domains associated with protein folding and thioredoxin fold domains were added to the system, likely recruited from roles as general regulatory domains, some of which could also bind Ca^2+^ (Figure 5). Association with thioredoxin fold domains, involved in disulfide-bond isomerization, is of note as this points to early emergence of a link between redox-dependent folding of cysteine-rich proteins and Ca^2+^ concentrations. The striking presence of cysteine-rich domains associated with cyanobacterial calmodulin homologs (Figure 3E; DES, LMI, and LA unpublished observations) suggests that such a connection might have emerged even before the origin of the eukaryotic Ca^2+^-stores system. These functional links might have persisted until later in eukaryotic evolution as hinted by the cysteine-rich domain present in wolframin. This led to the basic system as reconstructed in the LECA (Figure 2A).

**Figure 5.**
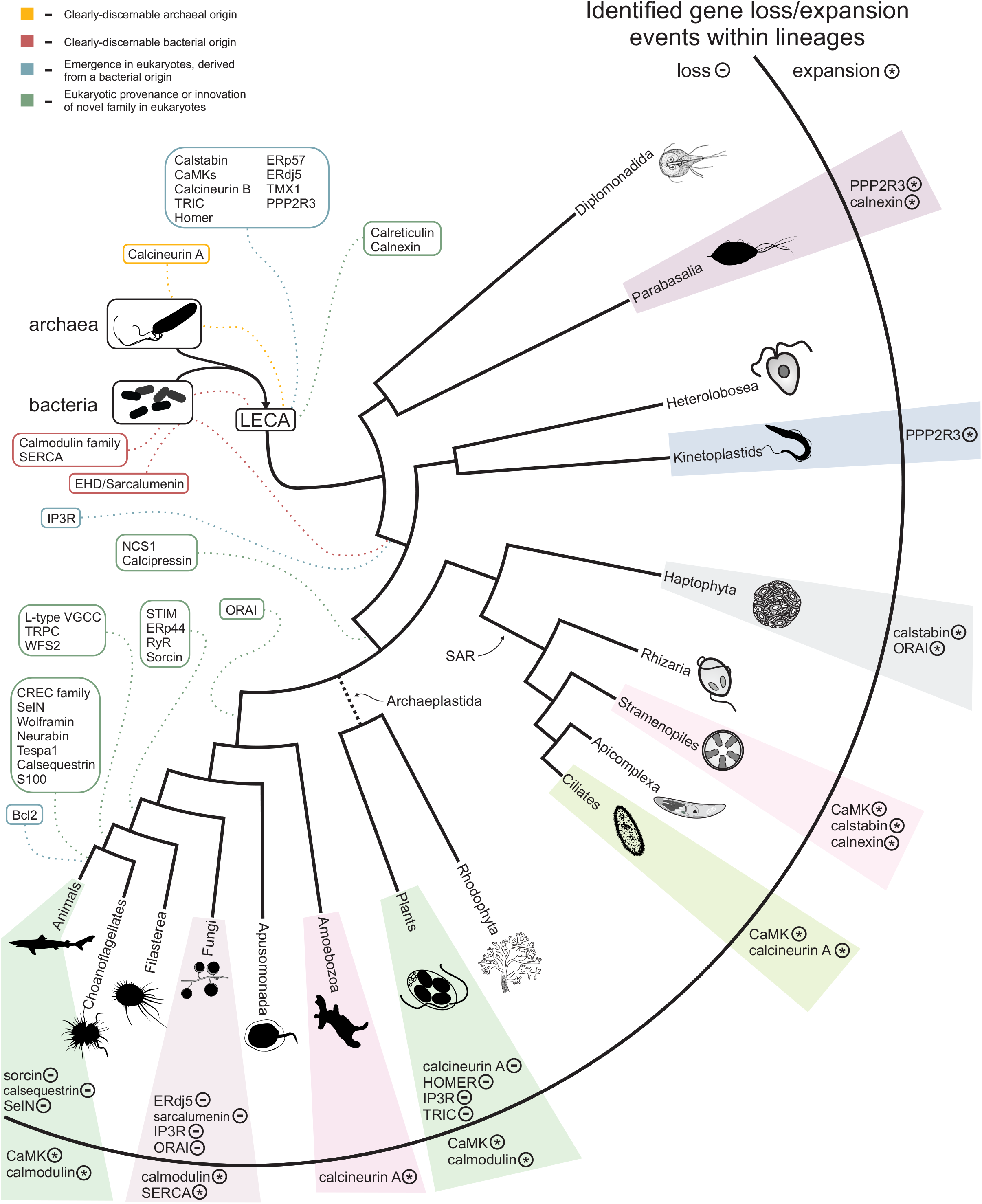
Summary of the evolution of the Ca^2+^ stores system. The temporal timing of the emergence of the core components of the system are overlaid on a consensus phylogenetic tree depicting eukaryotic evolution. Colors indicate component provenance, as labeled in the key provided in upper-left corner. Components with identified losses or expansions within a particular lineage are listed to the right, with ‘-’ and ‘*’ labels denoting loss and expansion of a particular component, respectively. A detailed breakdown of losses/expansions within the labeled lineages are provided in Figure 2 and Supplementary Figure S1.

Waves of additional accretion events added components to the Ca^2+^-stores system at distinct points in eukaryotic evolution, often appearing to correlate with adaptations to distinct lifestyles, such as the evolution of motile multicellular forms, loss of motility (the crown plant lineage), or the evolution of calcium-rich biomineralized skeletons and shells (Figures 2A, 5, Supplementary Figure S1). Strikingly, we appear to observe some correlation between the loss-and-gain patterns of the regulatory components of Ca^2+^-stores systems and the degree of structural utilization of calcium. For example, a relative dearth of these components is observed in the arthropod and fungal lineage, which use chitin-based structural components as opposed to the calcium-based biomineralized skeletons and shells of vertebrates and certain molluscs (Figure 2F-G). “Because the data in this study necessarily relies on experimental findings primarily from animals, it is best suited to characterizing the system from the viewpoint of metazoa. However, variations in presence/absence across lineages and LSEs provide insight into the dynamic evolution of Ca^2+^ stores regulation and suggests there are further complexities to explore in more poorly-characterized eukaryotes/

#### 4.1.2 Wolframin domain architecture and interactors

Even within animal Ca^2+^-stores systems, several proteins with important regulatory roles remain poorly-understood in terms of their functional mechanisms. Wolframin is such a protein whose domain composition has eluded researchers for over two decades (Inoue et al., 1998; Strom et al., 1998; Osman et al., 2003). Assignment of domains at the N-and C-termini of the central TM region (Figure 4A), as well as the positioning of wolframin in the assembled interaction network, (Figure 1A) (Zatyka et al., 2015) supports a role in coordinating interactions on both sides of the ER membrane. These are likely to take the form of PPIs with Ca^2+^-binding proteins or through disulfide bond interactions (see above).

However, outside of a possible bacterial origin for the SLR region (see above, Figure 4C), the precise evolutionary origins of the remaining domains comprising wolframin remain mostly unclear. It appears likely these domains were derived from paralogs of existing EF-hand and OB fold domains, both of which had already undergone extensive domain radiations in the eukaryotes prior to the emergence of the animals, and then assembled and recruited to a Ca^2+^-stores regulatory role at the ER (Lespinet et al., 2002). Their rapid divergence, evident by the lack of recognizable relationships to known families with their respective folds, could have resulted from the extraordinary selective pressures occurring with the major burst of evolutionary changes during the emergence of the metazoan lineage.

Despite observations that genic disruptions contribute to similar though not identical phenotypes (Urano, 2016), WFS1 and WFS2 are structurally unrelated. Our interaction network analysis further failed to uncover any shared interactors (Figure 1A), although both have been linked in the past to calpain activity (Lu et al., 2014), mitochondrial dysfunction (Chang et al., 2012; Cagalinec et al., 2016), and apoptosis (Yamada et al., 2006; Chang et al., 2010). Recent research on WFS2 has particularly focused on its role in calcium stores regulation at the intersection of the ER and mitochondrial membranes (Rouzier et al., 2017). We believe that our identification of the constituent domains of wolframin reported herein might help clarify its function better through target deletion and mutagenesis of these domains.

### 4.2 Conclusions

Reconstruction of the evolution of the eukaryotic Ca^2+^-stores regulatory system points to a core of domains inherited from distinct prokaryotic sources conserved across most eukaryotes. Lineage-specific differentiation of the system across eukaryotes is driven by complexities stemming from both the loss and/or expansion of the core complement of domains by the addition of components via LSEs or recruitment of domains of diverse provenance. We analyze in depth one such striking example of the latter, namely wolframin, which has been previously implicated in human disease.

The evolution of wolframin provides a model for how regulatory components of the Ca^2+^-stores system emerged: through the combining of existing mediators of Ca^2+^ signaling, like the EF-hand domains, with other domains originally not found in the system. Such transitions often happened at the base of lineages that subsequently underwent substantial diversification. We hope the findings presented here open novel avenues for the ongoing research on the regulation of calcium stores across eukaryotes, including providing new handles for understanding the functional mechanisms of wolframin and its dysregulation in Wolfram syndrome.

## 5 Conflict of Interest

The authors declare that the research was conducted in the absence of any commercial or financial relationships that could be construed as a potential conflict of interest.

## 6 Author Contributions

Conceptualization, DS, AMB, LA; formal analysis, DS, AMB, LMI; analytical tools, DS, LA; project administration, AMB, LMI, LA; visualization, DS; writing—original draft DS; AMB.; writing—review and editing, AMB, LMI, and LA

## 7 Funding

DES, LMI., AMB., and LA are supported by the Intramural Research Program of the NIH, National Library of Medicine.

## 8 Acknowledgments

This is a short text to acknowledge the contributions of specific colleagues, institutions, or agencies that aided the efforts of the authors.

## 9 Data Availability Statement

All datasets analyzed for this study are included in the manuscript and the supplementary files.

## Footnotes

1 ftp://ftp.ncbi.nih.gov/blast/documents/blastclust.html

2 http://tree.bio.ed.ac.uk/software/figtree

3 http://www.bioinformatics.nl/cgi-bin/emboss/pepwheel

