## Supplementary Material for "Functional innovation in the evolution of the calcium-dependent system of the eukaryotic endoplasmic reticulum"

Daniel E. Schäffer[1,2], Lakshminarayan M. Iyer[1], Alexander M. Burroughs[1], L Aravind*[1]

1-National Center for Biotechnology Information, National Library of Medicine, National Institutes of Health, Bethesda, MD 20894, USA

2-Montgomery Blair High School, Science, Mathematics, and Computer Science Magnet Program, Silver Spring, MD 20901, USA

*-corresponding authors

Table of Contents


I. Supplementary Figure Legends
II. Supplementary Table S1.
III. Supplementary Table S2.
IV. Supplementary Table S3.
V. Supplementary Table S4.
VI. Supplementary Figure S1.
VII. Supplementary Figure S2.
VIII. Supplementary Figure S3.
IX. Supplementary Figure S4.
X. Supplementary Data.

#### I. Supplementary Figure Legends

**Supplementary Figure S1.** Summary of evolutionary analysis of representative proteins involved in calcium signaling, including wolframin. The figure shows a matrix showing the presence (P / green) or absence (A / red) of homologs of 34 ER-calcium-signaling-related proteins in 41 extant taxonomic groups. Additionally, inferred presence/absence patters are shown for five key ancestral states, denoted in **bold**. Each row represents one protein or protein family, and each column represents one eukaryotic taxonomic group, sorted left-to-right by approximate time of divergence from the most recent ancestor shared with Vertebrata. 'Ties' among the columns are broken arbitrarily with some effort made to put related groups close to each other. The rightmost (i.e. most basal) "P" for each protein is bolded. The proteins are sorted top to bottom in approximate order of evolutionary occurrence, as determined by the most basal group in which a homolog of that protein was identified. The "Accession Used" column gives the RefSeq accession of the specific sequence used in the searches used to generate that row (see Methods).

**Supplementary Figure S2.** Six examples of conserved gene neighborhoods showing the associations between bacterial P-type ATPases and other proteins involved in ion homeostasis.

**Supplementary Figure S3.** Multiple-sequence alignment of the transmembrane region of wolframin. The transmembrane helices (as reported in Qian et al., 2015) are indicated by red cylinders above the alignment. Sequences are identified on the left by organism abbreviation (see Supplementary Table S3) and NCBI GenBank accession number, separated by two underscores. The alignment is colored as follows: h (hydrophobic), l (aliphatic), and a (aromatic) are shown on a yellow background; o (alcohol) is shown in salmon font; p (polar) is shown in blue font; + (positively charged), − (negatively charged), and c (charged) are shown in pink font; s (small) is shown in green font; u (tiny) is shown on a green background; b (big) is shaded gray. Columns in which the same amino acid is in at least 9 out of the 10 sequences are denoted with that amino acid's one-letter abbreviation in the consensus row and are colored based on a category to which that amino acid belongs.

**Supplementary Figure S4.** Renderings ('pepwheels') of amino acid positions on the circumference of wolframin's transmembrane helices. Each subfigure corresponds to one of wolframin's nine TM helices; the positions in human wolframin are indicated above the figure. The first amino acid is at the top, and the amino acids move clockwise at a rate of 18 amino acids per five rotations. Hydrophilic amino acids are in red.

**II. Supplementary Table S1**

| Major Eukaryotic Lineages |
|---|
| alveolata |
| amoebozoa |
| apusozoa |
| choanoflagellida |
| cristidiscoidea |
| cryptophyta |
| diplomonadida |
| euglenozoa |
| filasterea |
| fungi |
| haptophyceae |
| heterolobosea |
| ichthyosporea |
| metazoa |
| parabasalia |
| rhizaria |
| rhodophyta |
| stramenopiles |
| viridiplantae |

##### III. Supplementary Table S2

| Protein / protein family | Domain annotation |
|---|---|
| Calmodulin family | EF-hand_7+EF-hand_7 |
| ERp57 (PDIA3) | Thioredoxin+Thioredoxin_6+Thioredoxin+Thioredoxin_8 |
| SERCA | Cation_ATPase_N+TM+TM+E1-E2_ATPase+TM+TM+Cation_ATPase+Hydrolase+Hydrolase_3+TM+Cation_ATPase_C |
| CaMK / CDPK | Pkinase^ |
| Calstabin | FKBP_c |
| PPP2R3 | EF-hand_7* |
| Calcineurin A | Metallophos |
| Calcineurin B | EF-hand_7+EF-hand_7+EF-hand_6+EF-hand_4 |
| TMX1 | Thioredoxin+TM |
| Calreticulin | Calreticulin |
| Calnexin | Calreticulin+TM |
| TRIC | TRIC |
| Homer (WH1 dom.) | EVH1^ |
| ERdj5 | DnaJ+Thioredoxin+Thioredoxin+Thioredoxin+Thioredoxin |
| Sarcalumenin | EHD_N+Dynamin_N |
| IP3R | Ins145_P3_rec+MIR+RYDR_ITPR+RYDR_ITPR+RIH_assoc+TM+TM+TM+TM+TM+TM |
| ORAI | Orai-1 |
| NCS1 | EF-hand_7+EF-hand_7+EF-hand_7+EF-hand_6 |
| Calcipressin | Calcipressin |
| STIM | EF-hand+SAM+TM+SOAR |
| ERp44 | Thioredoxin+Thioredoxin_6 |
| RyR | Ins145_P3_rec+MIR+RYDR_ITPR+SPRY+RyR+RyR+SPRY+SPRY+RYDR_ITPR+RyR+RyR+RIH_assoc+EF-hand_8+TM+TM+TM+TM+TM+TM |
| Sorcin | EF-hand_8+EF-hand_7 |
| L-type VGCC (pore) | TM+TM+TM+TM+TM+TM+TM+TM+TM+TM+TM+TM+TM+TM+TM+TM+TM+TM+TM+TM+TM+TM+TM+TM+GPHH+Ca_chan_IQ+CAC1F_C |
| TRPC | Ank_2+TRP_2+TM+TM+TM+TM+TM+TM |

| | |
|---|---|
| WFS2 (CISD2) | MitoNEET_N+zf-CDGSH |
| CREC family | EF-hand_7+EF-hand_7+EF-hand_7 |
| Bcl2 | BH4+Bcl-2+TM† |
| SelN | EF-hand_6 |
| Wolframin | Sel1+Sel1+Sel1+wEF-hand+TM+TM+TM+TM+TM+TM+TM+TM+TM+wOB‡ |
| Neurabin | PDZ* |
| Tespa1 (KRAP dom.) | KRAP_IP3R_bind |
| Calsequestrin | Calsequestrin |
| S100 | S_100+EF-hand_7 |

^: Other domains vary

*: Other domains vary and are not very globular

†: Domain architecture varies between family members

‡: The domains wEP-hand and wOB refer to the wolframin-specific version of the EF-hand and OB-fold domains reported in this paper

**IV. Supplementary Table S3**

| Alignment Abbreviation | Scientific name | Common name | Clade |
|---|---|---|---|
| Hsapiens | *Homo sapiens* | Human | *Mammalia* |
| Mmusculus | *Mus musculus* | House mouse | *Mammalia* |
| Ggallus | *Gallus gallus* | Chicken | *Aves* |
| Drerio | *Danio rerio* | Zebrafish | *Actinopterygii* |
| Skowalevskii | *Saccoglossus kowalevskii* | Acorn worm | *Hemichordata* |
| Dmelanogaster | *Drosophila melanogaster* | Fruit fly | *Hexapoda* |
| Tcastaneum | *Tribolium castaneum* | Red flour beetle | *Hexapoda* |
| Dpulex | Daphnia pulex | Water flea | *Crustacea* |
| Lpolyphemus | *Limulus polyphemus* | Atlantic horseshoe crab | *Chelicerata* |
| Lgigantea | *Lottia gigantea* | Owl limpet | *Mollusca* |
| Obimaculodies | *Octopus bimaculodies* | California two-spot octopus | *Mollusca* |
| Hrobusta | *Helobdella robusta* | Leech | *Annelida* |
| Celegans | *Caenorhabditis elegans* | | *Nematoda* |
| Epallida | *Exaiptasia pallida* | Brown glass anemone | *Cnidaria* |
| Tadhaerens | *Trichoplax adhaerens* | | *Placozoa* |
| Aqueesnlandica | *Amphimedon queenslandica* | Sponge | *Porifera* |
| Srosetta | *Salpingoeca rosetta* | | *Choanoflagellatea* |
| Smoellendorffii | *Selaginella moellendorffii* | Spikemoss | *Streptophyta* |
| Vcarteri | *Volvox carteri f. nagariensis* | | *Chlorophyta* |
| Tpseudonana | *Thalassiosira pseudonana CCMP1335* | | *Bacillariophyta* |
| Pinfestans | *Phytophthora infestans T30-4* | Late potato blight | *Oomycetes* |
| Tgondii | *Toxoplasma gondii ME49* | | *Apicomplexa* |
| Slemnae | *Stylonychia lemnae* | | *Ciliophora* |
| Ehuxleyi | *Emiliania huxleyi CCMP1516* | | *Haptophyta* |

##### V. Supplementary Table S4

| Human wolframin repeat 1 | | Human wolframin repeat 2 | | Human wolframin repeat 3 | | Canonical SLR^ | |
|---|---|---|---|---|---|---|---|
| Position | Residue | Position | Residue | Position | Residue | Position | Residue |
| 65 | A | 102 | A | 141 | A | 3 | A |
| 69 | R | 106 | V | 145 | L | 7 | L |
| 70 | S | 107 | G | 146 | R | 8 | G |
| 73 | R | 110 | Y | 149 | L | 11 | Y |
| 76 | G | 113 | L | 154 | G | 14 | G |
| 83 | D | 123 | S | 158 | E | 28 | D |
| 87 | P | 126 | A | 162 | E | 32 | A |
| 94 | R | 133 | A | 169 | E | 39 | A |
| 95 | A | 134 | A | 170 | T | 40 | A |
| 98 | G | 137 | G | 173 | E | 43 | G |

^: From Mittl & Schneider-Brachert, 2007

### VI. Supplementary Figure S1

| Name | Accession Used | metazoa>chordata>vertebrata | metazoa>chordata>tunicata | metazoa>chordata>cephalochordata | metazoa>echinodermata | metazoa>hemichordata | metazoa>arthropoda>hexapoda | metazoa>arthropoda>chelicerata | metazoa>arthropoda>crustacea | metazoa>nematoda | metazoa>priapulida | metazoa>annelida | metazoa>mollusca | metazoa>brachiopoda | metazoa>platyhelminthe | metazoa>cnidaria | metazoa>placozoa | metazoa>porifera | **Metazoan progenitor** | choanoflagellida | filasterea | **Holozoan progenitor** | fungi>dikarya | fungi>chytridiomycota | fungi>mucoromycotina | fungi>kickxellomycotina | **Fungal progenitor** | cristidiscoidea | apusozoa>apusomonadidae | amoebozoa>acanthamoebidae | amoebozoa>entamoebidae | amoebozoa>mycetozoa>dictyostelida | **Unnikont progenitor** | viridiplantae>streptophyta | viridiplantae>chlorophyta | rhodophyta | stramenopiles>bacillariophyta | stramenopiles>oomycetes | alveolata>apicomplexa | alveolata>ciliophora | rhizaria | **SAR progenitor** | haptophyceae | heterolobosea | euglenozoa>kinetoplastida | diplomonadida | parabasalia | **LECA** |
|---|---|---|---|---|---|---|---|---|---|---|---|---|---|---|---|---|---|---|---|---|---|---|---|---|---|---|---|---|---|---|---|---|---|---|---|---|---|---|---|---|---|---|---|---|---|---|---|---|
| Calmodulin family | NP_008819.1 | P | P | P | P | P | P | P | P | P | P | P | P | P | P | P | P | P | P | P | P | P | P | P | P | P | P | P | P | P | P | P | P | P | P | P | P | P | P | P | P | P | P | P | P | P | P | P |
| ERp57 (PDIA3) | NP_005304.3 | P | P | P | P | P | P | P | P | P | P | P | P | P | P | P | P | P | P | P | P | P | P | P | P | P | P | P | P | P | P | P | P | P | P | P | A | P | P | P | P | P | P | P | P | P | P | P |
| SERCA | NP_733765.1 | P | P | P | P | P | P | P | P | P | P | P | P | P | P | P | P | P | P | P | P | P | P | P | P | P | P | P | P | P | P | A | P | P | P | P | P | P | P | P | P | P | P | P | P | A | P | P |
| CaMK / CDPK | NP_001308498.1 | P | P | P | P | P | P | P | P | P | P | P | P | P | P | P | P | P | P | P | P | P | P | P | P | P | P | A | P | P | P | P | P | P | P | P | P | P | P | P | P | P | P | P | P | A | P | P |
| Calstabin | NP_463460.1 | P | P | P | P | P | P | P | P | P | P | P | P | P | P | P | P | P | P | P | P | P | P | P | P | P | P | A | P | P | A | P | P | P | P | P | P | P | P | P | P | P | P | P | P | P | P | P |
| PPP2R3 | NP_002709.2 | P | P | P | P | P | P | P | P | P | P | P | P | P | P | P | P | P | P | P | A | P | A | P | P | P | P | P | P | P | A | P | P | P | P | P | P | P | P | P | P | P | P | P | P | P | P | P |
| Calcineurin A | NP_000935.1 | P | P | P | P | P | P | P | P | P | P | P | P | P | P | P | P | P | P | P | P | P | P | P | P | P | P | P | P | P | P | P | P | A | A | A | A | P | P | P | P | P | A | P | P | P | P | P |
| Calcineurin B | NP_000936.1 | P | P | P | P | P | P | P | P | P | P | P | P | P | P | P | P | P | P | P | P | P | P | P | P | P | P | P | P | P | P | P | P | P | P | P | A | A | P | P | P | P | P | P | P | P | P | P |
| TMX1 | NP_110382.3 | P | P | P | P | P | P | P | P | P | P | P | P | P | P | P | P | P | P | A | P | P | P | P | P | A | P | P | A | P | A | P | P | P | P | P | P | P | P | P | P | P | P | P | P | A | P | P |
| Calreticulin | NP_004334.1 | P | P | P | P | P | P | P | P | P | P | P | P | P | P | P | P | P | P | P | P | P | A | P | P | P | P | P | A | P | P | P | P | P | P | A | P | P | A | P | P | P | P | P | P | A | P | P |
| Calnexin | NP_001737.1 | P | P | P | P | P | P | P | P | P | P | P | P | P | P | P | P | P | P | P | A | P | P | P | A | P | P | A | P | A | A | P | P | P | P | P | A | P | P | A | P | P | P | P | A | A | P | P |
| TRIC | NP_076979.1 | P | P | P | P | P | P | P | P | P | P | P | P | P | P | P | P | P | P | A | P | P | P | P | P | P | P | P | P | P | A | P | P | A | P | P | P | A | A | A | P | P | P | P | A | A | P | P |
| Homer (WH1 dom.) | NP_004263.1 | P | P | P | P | P | P | P | P | P | P | P | P | P | P | P | P | P | P | P | P | P | P | P | P | A | P | P | P | P | A | P | P | A | A | P | A | A | A | A | P | P | A | P | A | A | P | P |
| ERdj5 | NP_061854.1 | P | P | P | P | P | P | P | P | P | A | P | P | P | P | P | P | A | P | P | A | P | A | A | A | A | A | A | A | P | A | A | P | P | P | A | A | P | P | P | P | P | P | A | A | A | P | P |
| Sarcalumenin | NP_001092284.1 | P | P | P | P | P | P | P | P | P | P | P | P | P | P | P | P | P | P | P | P | P | A | P | P | P | P | P | P | P | P | P | P | P | P | P | P | P | P | A | P | P | P | A | P | A | A | A |
| IP3R | NP_001161744.1 | P | P | P | P | P | P | P | P | P | P | P | P | P | P | P | P | P | P | P | P | P | A | P | P | A | P | P | P | P | A | P | P | A | P | A | A | P | A | P | P | P | P | P | P | A | A | A |
| ORAI | NP_116179.2 | P | P | P | P | P | P | P | P | P | P | P | P | P | P | P | P | P | P | P | P | P | A | A | A | A | A | A | A | A | A | A | P | P | P | A | P | P | P | P | A | P | P | A | A | A | A | A |
| NCS1 | NP_055101.2 | P | P | P | P | P | P | P | P | P | P | P | P | P | P | P | P | P | P | P | P | P | P | P | P | P | P | A | P | P | A | P | P | A | A | A | A | A | A | A | A | A | A | A | A | A | A | A |
| Calcipressin | NP_001272320.2 | P | P | P | P | P | P | P | P | P | P | P | P | P | P | P | P | P | P | P | P | P | P | P | P | P | P | A | A | P | A | P | P | A | A | A | A | A | A | A | A | A | A | A | A | A | A | A |
| STIM | NP_001264890.1 | P | P | P | P | P | P | P | P | P | P | P | P | P | P | P | P | P | P | P | P | P | A | A | A | A | A | A | A | A | A | A | A | A | A | A | A | A | A | A | A | A | A | A | A | A | A | A |
| ERp44 | NP_055866.1 | P | P | P | P | P | P | P | P | P | P | P | P | P | P | P | P | P | P | P | P | P | A | A | A | A | A | A | A | A | A | A | A | A | A | A | A | A | A | A | A | A | A | A | A | A | A | A |
| RyR | NP_000531.2 | P | P | P | P | P | P | P | P | P | P | P | P | P | P | A | P | P | P | P | P | P | A | A | A | A | A | A | A | A | A | A | A | A | A | A | A | A | A | A | A | A | A | A | A | A | A | A |
| Sorcin | NP_003121.1 | P | A | P | P | P | A | A | A | A | A | A | P | P | A | P | P | P | P | P | P | P | A | A | A | A | A | A | A | A | A | A | A | A | A | A | A | A | A | A | A | A | A | A | A | A | A | A |
| L-type VGCC (pore) | NP_000060.2 | P | P | P | P | P | P | P | P | P | P | P | P | P | P | P | P | P | P | P | A | A | A | A | A | A | A | A | A | A | A | A | A | A | A | A | A | A | A | A | A | A | A | A | A | A | A | A |
| TRPC | NP_003297.1 | P | P | P | P | P | P | P | P | P | P | P | P | P | P | P | A | A | P | P | A | A | A | A | A | A | A | A | A | A | A | A | A | A | A | A | A | A | A | A | A | A | A | A | A | A | A | A |
| WFS2 (CISD2) | NP_001008389.1 | P | P | P | P | P | P | A | P | A | P | P | P | P | P | P | P | P | P | P | A | A | A | A | A | A | A | A | A | A | A | A | A | A | A | A | A | P | A | A | A | A | A | A | A | A | A | A |
| CREC family | NP_001186601.1 | P | P | P | P | P | P | P | P | P | P | P | P | P | P | P | P | P | P | A | A | A | A | A | A | A | A | A | A | A | A | A | A | P | A | A | A | A | A | A | A | A | A | A | A | A | A | A |
| Bcl2 | NP_000624.2 | P | P | P | P | P | P | P | P | P | P | P | P | P | P | P | P | P | P | A | A | A | A | A | A | A | A | A | A | A | A | A | A | A | A | A | A | A | A | A | A | A | A | A | A | A | A | A |
| SelN | NP_065184.2 | P | P | P | P | P | A | P | A | A | P | P | P | P | P | P | A | P | P | A | A | A | A | A | A | A | A | A | A | A | A | A | A | A | A | A | A | A | A | A | A | A | A | A | A | A | A | A |
| Wolframin | NP_005996.2 | P | P | A | A | P | P | P | P | P | A | P | P | P | A | P | P | P | P | A | A | A | A | A | A | A | A | A | A | A | A | A | A | A | A | A | A | A | A | A | A | A | A | A | A | A | A | A |
| Neurabin | NP_115984.3 | P | P | P | P | P | P | P | P | P | P | P | P | P | P | P | P | A | A | A | A | A | A | A | A | A | A | A | A | A | A | A | A | A | A | A | A | A | A | A | A | A | A | A | A | A | A | A |
| Tespa1 (KRAP dom.) | NP_001129502.1 | P | P | P | P | P | P | P | P | A | P | P | P | P | A | P | P | A | A | A | A | A | A | A | A | A | A | A | A | A | A | A | A | A | A | A | A | A | A | A | A | A | A | A | A | A | A | A |
| Calsequestrin | NP_001222.3 | P | P | P | A | P | A | P | A | P | A | A | A | A | A | P | P | A | A | A | A | A | A | A | A | A | A | A | A | A | A | A | A | A | A | A | A | A | A | A | A | A | A | A | A | A | A | A |
| S100 | NP_006262.1 | P | P | A | A | A | A | A | A | A | A | A | A | A | A | A | A | A | A | A | A | A | A | A | A | A | A | A | A | A | A | A | A | A | A | A | A | A | A | A | A | A | A | A | A | A | A | A |

### VII. Supplementary Figure S2

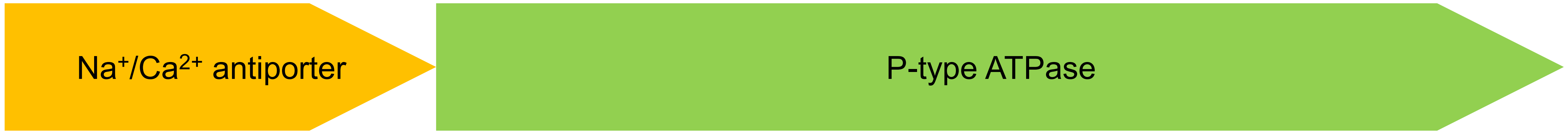


ABQ30334, ABQ30333 from *Acidiphilium cryptum JF-5* (α-proteobacteria)

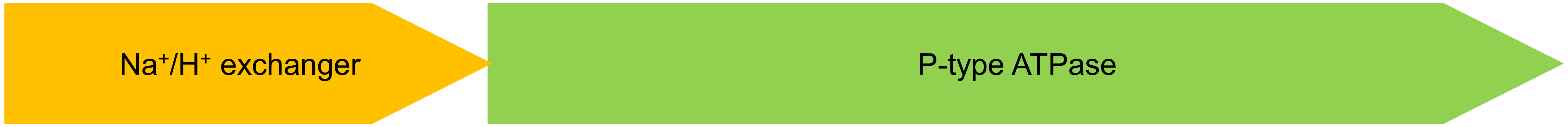


ADN08417, ADN08416 from *Sulfurimonas autotrophica DSM 16294* (ε-proteobacteria)

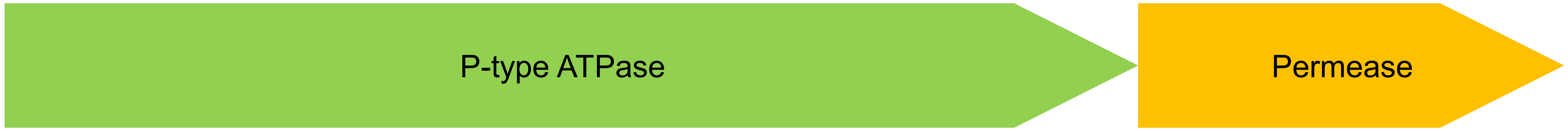


AFL87427, AFL87428 from *Terriglobus roseus DSM 18391* (acidobacteria)

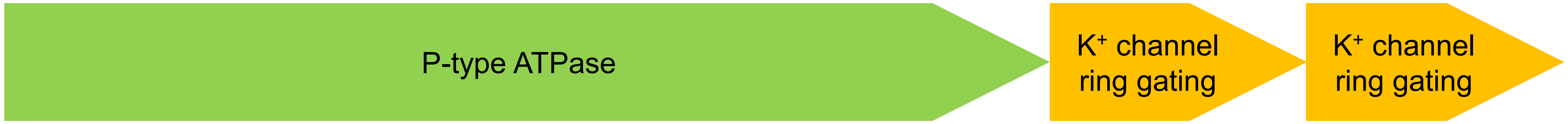


AEY94042, AEY94041, AEY94040 from *Streptomyces hygroscopicus subsp. jinggangensis 5008* (actinobacteria)

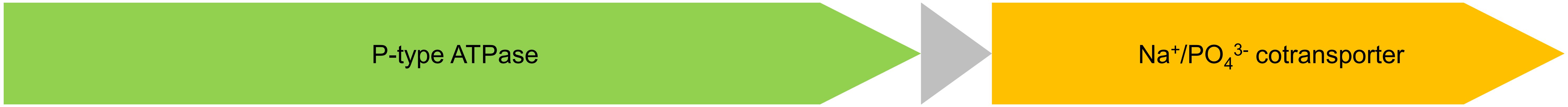


APC18922, APC18921, APC18920 from *Pseudomonas frederiksbergensis* (γ-proteobacteria)

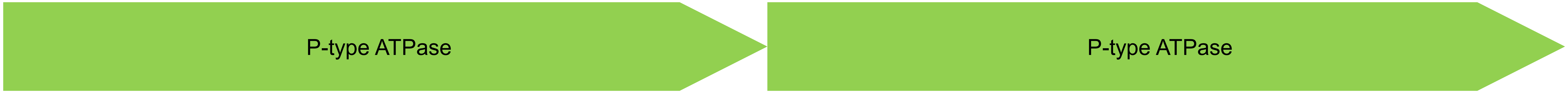


AEN97664, AEN97663 from *Roseburia hominis A2-183* (firmicutes)

### VIII. Supplementary Figure S3

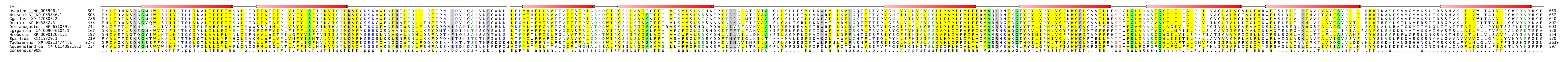

### IX. Supplementary Figure S4

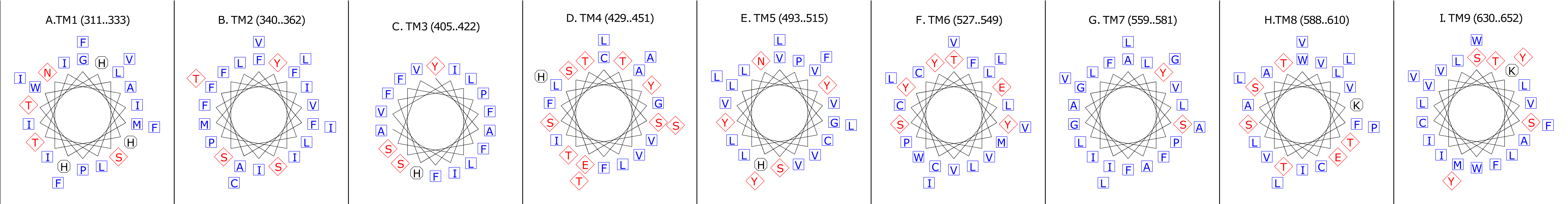

SUPPLEMENTARY DATA

### Functional innovation in the evolution of the calcium-dependent system of the eukaryotic endoplasmic reticulum

Daniel E. Schäffer1,2, Lakshminarayan M. Iyer1, Alexander M. Burroughs1, L Aravind1

1 - National Center for Biotechnology Information, National Library of Medicine, National Institutes of Health
2 - Montgomery Blair High School, Science, Mathematics, and Computer Science Magnet Program

ABSTRACT

The origin of eukaryotes was marked by the emergence of several novel subcellular systems. One such is the calcium (Ca2+)-stores system of the endoplasmic reticulum, which profoundly influences diverse aspects of cellular function including signal transduction, motility, division, and biomineralization. We use comparative genomics and sensitive sequence and structure analyses to investigate the evolution of this system. Our findings reconstruct the core form of the Ca2+- stores system in the last eukaryotic common ancestor as having at least 15 proteins that constituted a basic system for facilitating both Ca2+ flux across endomembranes and Ca2+-dependent signaling. We present evidence that the key EF-hand Ca2+-binding components had their origins in a likely bacterial symbiont other than the mitochondrial progenitor, whereas the protein phosphatase subunit of the ancestral calcineurin complex was likely inherited from the asgardarchaeal progenitor of the stem eukaryote. This further points to the potential origin of the eukaryotes in a Ca2+-rich biomineralized environment such as stromatolites. We further show that throughout eukaryotic evolution there were several acquisitions from bacteria of key components of the Ca2+-stores system, even though no prokaryotic lineage possesses a comparable system. Further, using quantitative measures derived from comparative genomics we show that there were several rounds of lineage-specific gene expansions, innovations of novel gene families, and gene losses correlated with biological innovation such as the biomineralized molluscan shells, coccolithophores, and animal motility. The burst of innovation of new genes in animals included the wolframin protein associated with Wolfram syndrome in humans. We show for the first time that it contains previously unidentified Sel1, EF-hand, and OB-fold domains, which might have key roles in its biochemistry.

TABLE OF CONTENTS

I. Edges in the protein-protein interaction network of ER-dependent Ca2+ storage and signaling
II. Accession numbers for homologs of proteins and protein families key to calcium storage and signaling, with their phylogenies
  A. Calmodulin family
  B. ERp57 (PDIA3)
  C. SERCA
  D. CaMK / CDPK
  E. Calstabin
  F. PPP2R3
  G. Calcineurin A
  H. Calcineurin B
  I. TMX1
  J. Calreticulin
  K. Calnexin
  L. TRIC
  M. HOMER (WH1 dom.)
  N. ERdj5
  O. Sarcalumenin/EHD
  P. IP3R
  Q. ORAI
  R. NCS1
  S. Calcipressin
  T. STIM
  U. ERp44
  V. RyR
  W. Sorcin
  X. L-type VGCC (pore)
  Y. TRPC
  Z. WFS2 (CISD2)
  AA. CREC family
  AB. Bcl2
  AC. SelN
  AD. Wolframin
  AE. Neurabin
  AF. Tespa1 (KRAP dom.)
  AG. Calsequestrin
  AH. S100
III. Accession numbers for prokaryotic relatives of proteins and protein families key to calcium storage and signaling, with their phylogenies
  A. Accession numbers for selected calmodulin-like proteins with domain fusions
    i. Fusions to Tic110-like domains
    ii. Fusions to Heme-Oxygenase domains
    iii Fusions to NO synthase domains
    iv. Fusions to NADPH oxidase domains
    v. Fusions to cNMP-binding domains
    vi. Fusions to thioredoxin and/or sulfatase domains
    vii. Fusions to glyoxalase domains
  B. Accession numbers for selected bacterial PPIases
IV. Newick-format trees
  A. Sarcalumenin, EHD, and bacterial relatives thereof
  B. Calcineurin-like phosphatases, archaeal relatives thereof, and MRE11/rad32/sbcD-like phosphatases
  C. Calcium/calmodulin-dependent kinases
  D. ORAI and Jiraiya
V. Alignments
  A. Alignment of representative Jiraya and ORAI family members
VI. List of completely sequenced curated eukaryotic genomes used for the analysis.

#### I. Edges in the protein-protein interaction network of ER-dependent Ca2+ storage and signaling

Edges which were included in the protein-protein interaction network presented in the paper are provided below in a tab-seperated format, where the first and second columns are two interacting proteins (represented by their encoding gene), the third column is the interaction type, and the fourth and subsequent columns (if any) provide the source of the interaction. There are three possible interaction types: 1 (interactions reported in the literature in which each partner gene either has no other paralog or has been specifically identified), 3 (reported interactions where the specific paralog for at least one of the two genes is unclear), and 4 (interactions found in FunCoup 4.0 with P = 0.9). Sources are in the form of PubMed IDs of papers that describe the interactions for types 1 and 3, and "FC P" for type 4 interactions, where P is the FunCoup P-score.

| Gene 1 | Gene 2 | Type | Source | Second source |
|---|---|---|---|---|
| ACYP2 | ATP2A2 | 1 | PMID16105684 | |
| AGTRAP | ATP2A2 | 1 | PMID27015675 | |
| AHCYL1 | CAMK2A | 1 | PMID19133301 | PMID25922519 |
| AHCYL1 | CISD2 | 1 | PMID19133301 | |
| AHCYL1 | ITPR1 | 1 | PMID19133301 | |
| AHCYL1 | ITPR2 | 1 | PMID17429043 | |
| AHCYL1 | ITPR3 | 1 | PMID17429043 | |
| AHCYL1 | PPP1CA | 1 | PMID19133301 | |
| AHCYL1 | SRL | 1 | PMID10398759 | |
| AKAP6 | PDE4D | 1 | PMID19345240 | |
| AKAP6 | RYR1 | 1 | PMID12709444 | |
| AKAP6 | RYR2 | 1 | PMID19345240 | |
| AKAP9 | ITPR1 | 1 | PMID19133301 | |
| AKAP9 | PPP1CA | 1 | PMID19133301 | PMID14982933 |
| AKT1 | CAMK2G | 1 | PMID26803057 | |
| AKT1 | ITPR1 | 1 | PMID19133301 | |
| AKT1 | ITPR2 | 1 | PMID19133301 | |
| AKT1 | ITPR3 | 1 | PMID19133301 | |
| AKT1 | PML | 1 | PMID21030605 | |
| ALDOA | CAPN3 | 1 | PMID18676612 | |
| ALDOA | RYR1 | 1 | PMID18676612 | |
| ANK1 | ATP2A1 | 1 | PMID28487373 | |
| ANK1 | OBSCN | 1 | PMID19002483 | |
| ANK1 | SLN | 1 | PMID28487373 | |
| ANK2 | ITPR1 | 1 | PMID17429043 | |
| ANK2 | SIGMAR1 | 1 | PMID17429043 | |
| ANXA7 | SRI | 1 | PMID9268363 | PMID10767422 |
| ASPH | CASQ1 | 1 | PMID20961976 | |
| ASPH | CASQ2 | 1 | PMID20961976 | |
| ASPH | ORAI1 | 1 | PMID22586105 | |
| ASPH | RYR1 | 1 | PMID19398037 | |
| ASPH | RYR2 | 1 | PMID15041652 | |
| ASPH | STIM1 | 1 | PMID22586105 | |
| ASPH | TRDN | 1 | PMID20961976 | |
| ASPH | TRPC2 | 1 | PMID16153633 | |
| ASPH | TRPC5 | 1 | PMID16153633 | |
| ATF6 | WFS1 | 1 | PMID20160352 | |
| ATG5 | BCL2 | 1 | PMID19043451 | |
| ATP2A1 | BCL2 | 1 | PMID16105864 | |
| ATP2A1 | IRS1 | 1 | PMID16105684 | |
| ATP2A1 | IRS2 | 1 | PMID16105684 | |
| ATP2A1 | MRLN | 1 | PMID26816378 | |
| ATP2A1 | SLN | 1 | PMID28487373 | |
| ATP2A1 | SMIM6 | 1 | PMID27923914 | |
| ATP2A1 | SRL | 1 | PMID16105684 | |
| ATP2A1 | STIM1 | 1 | PMID24077737 | |
| ATP2A2 | BCL2 | 1 | PMID16105684 | PMID26772784 |
| ATP2A2 | CALR | 1 | PMID16105684 | |
| ATP2A2 | CALU | 1 | PMID20356454 | PMID18562801 |
| ATP2A2 | CAMK2A | 1 | PMID16105684 | PMID25515219 |
| ATP2A2 | CAMK2B | 1 | PMID16105684 | PMID25515219 |
| ATP2A2 | CANX | 1 | PMID16105684 | PMID21234650 |
| ATP2A2 | DNAJC10 | 1 | PMID28614305 | PMID27694578 |
| ATP2A2 | HRC | 1 | PMID22952658 | |
| ATP2A2 | IRS1 | 1 | PMID16105684 | |
| ATP2A2 | IRS2 | 1 | PMID16105684 | |
| ATP2A2 | MRLN | 1 | PMID26816378 | |
| ATP2A2 | NOX4 | 1 | PMID24386930 | |
| ATP2A2 | OPRD1 | 1 | PMID21234650 | |
| ATP2A2 | PLN | 1 | PMID27923914 | |
| ATP2A2 | PRKG1 | 1 | PMID16105684 | |
| ATP2A2 | PSEN1 | 1 | PMID18591429 | |
| ATP2A2 | PSEN2 | 1 | PMID18591429 | |
| ATP2A2 | S100A1 | 1 | PMID16105684 | |
| ATP2A2 | SDF4 | 1 | PMID25772237 | |
| ATP2A2 | SELENON | 1 | PMID28506443 | PMID25452428 |
| ATP2A2 | SLN | 1 | PMID27923914 | |
| ATP2A2 | SMIM6 | 1 | PMID27923914 | |
| ATP2A2 | STIM1 | 1 | PMID25772237 | |
| ATP2A2 | TMEM203 | 1 | PMID25996873 | |
| ATP2A2 | TMX1 | 1 | PMID28619231 | |
| ATP2A2 | TP53 | 1 | PMID26772784 | PMID25624484 |
| ATP2A2 | WFS1 | 1 | PMID25274773 | |
| ATP2A3 | BCL2 | 1 | PMID27639965 | |
| ATP2A3 | IRS1 | 1 | PMID16105684 | |
| ATP2A3 | MRLN | 1 | PMID26316378 | |
| ATP2A3 | SMIM6 | 1 | PMID27923914 | |
| ATP2A3 | STIM1 | 1 | PMID18439569 | |
| BAK1 | ITPR1 | 1 | PMID15613488 | |
| BANK1 | ITPR1 | 1 | PMID19133301 | |
| BANK1 | ITPR2 | 1 | PMID19133301 | |
| BANK1 | LYN | 1 | PMID19133301 | |
| BAP1 | ITPR3 | 1 | PMID28614305 | PMID28619231 |
| BAX | BID | 1 | PMID19043451 | |
| BAX | ITPR1 | 1 | PMID15613488 | |
| BAX | TMBIM6 | 1 | PMID21718971 | |
| BCL2 | BECN1 | 1 | PMID28725634 | PMID11309306 |
| BCL2 | CISD2 | 1 | PMID22617439 | |
| BCL2 | FKBP8 | 1 | PMID19043451 | PMID12510191 |
| BCL2 | ITPR1 | 1 | PMID15613488 | PMID19133301, PMID22617439 |
| BCL2 | ITPR2 | 1 | PMID19133301 | PMID19752026, PMID22617439 |
| BCL2 | ITPR3 | 1 | PMID19133301 | PMID22617439 |
| BCL2 | NR4A1 | 1 | PMID19043451 | |
| BCL2 | PPP1CA | 1 | PMID19133301 | |
| BCL2 | RYR1 | 1 | PMID24762814 | |
| BCL2 | RYR2 | 1 | PMID24762814 | |
| BCL2 | RYR3 | 1 | PMID25957473 | PMID24762814 |
| BCL2 | TMBIM6 | 1 | PMID21718971 | |
| BCL2L1 | FKBP8 | 1 | PMID12510191 | |
| BCL2L1 | ITPR1 | 1 | PMID19133301 | |
| BCL2L1 | ITPR2 | 1 | PMID19133301 | |
| BCL2L1 | ITPR3 | 1 | PMID19133301 | |
| BCL2L1 | RYR3 | 1 | PMID25872771 | |
| BCL2L1 | TMBIM6 | 1 | PMID21718971 | |
| BECN1 | CAV1 | 1 | PMID28542134 | |
| BECN1 | ITPR1 | 1 | PMID17256008 | |
| BECN1 | ITPR3 | 1 | PMID17256008 | |
| BECN1 | PIK3C3 | 1 | PMID28725634 | PMID28542134 |
| BOK | ITPR1 | 1 | PMID26772784 | |
| BOK | ITPR2 | 1 | PMID26772784 | |
| BRCA1 | ITPR1 | 1 | PMID26772784 | |
| CABP1 | ITPR1 | 1 | PMID17429043 | PMID14685260 |
| CABP1 | ITPR2 | 1 | PMID17429043 | |
| CABP1 | ITPR3 | 1 | PMID17429043 | |
| CABP2 | ITPR1 | 1 | PMID12032348 | |
| CABP2 | ITPR2 | 1 | PMID12032348 | |
| CABP2 | ITPR3 | 1 | PMID12032348 | |
| CABP5 | ITPR1 | 1 | PMID12032348 | |
| CABP5 | ITPR2 | 1 | PMID12032348 | |
| CABP5 | ITPR3 | 1 | PMID12032348 | |
| CACNA1C | CALM1 | 1 | PMID16299511 | |
| CACNA1C | CALM2 | 1 | PMID16299511 | |
| CACNA1C | CALM3 | 1 | PMID16299511 | |
| CACNA1C | OPRD1 | 1 | PMID25552382 | |
| CACNA1C | RYR2 | 1 | PMID20961976 | |
| CACNA1S | CAV3 | 1 | PMID22020936 | |
| CACNA1S | JPH1 | 1 | PMID22020936 | |
| CACNA1S | JPH2 | 1 | PMID22020936 | |
| CACNA1S | RYR1 | 1 | PMID22020936 | PMID20961976 |
| CALM1 | ITPR1 | 1 | PMID19133301 | |
| CALM1 | ITPR2 | 1 | PMID19133301 | |
| CALM1 | ITPR3 | 1 | PMID19133301 | |
| CALM1 | ORAI1 | 1 | PMID23109337 | |
| CALM1 | PPP3CA | 1 | PMID12809556 | PMID21115349 |
| CALM1 | RYR1 | 1 | PMID20961976 | |
| CALM1 | RYR2 | 1 | PMID20961976 | |
| CALM1 | RYR3 | 1 | PMID20961976 | |
| CALM1 | WFS1 | 1 | PMID19292454 | |
| CALM2 | ORAI1 | 1 | PMID23109337 | |
| CALM2 | PPP3CA | 1 | PMID21115349 | |
| CALM2 | RYR1 | 1 | PMID20961976 | |
| CALM2 | WFS1 | 1 | PMID19292454 | |
| CALM3 | ORAI1 | 1 | PMID23109337 | |
| CALM3 | PPP3CA | 1 | PMID21115349 | |
| CALM3 | RYR1 | 1 | PMID20961976 | |
| CALM3 | WFS1 | 1 | PMID19292454 | |
| CALR | PDIA3 | 1 | PMID16105684 | PMID28619231 |
| CALU | RYR1 | 1 | PMID16527250 | |
| CALU | RYR2 | 1 | PMID20356454 | |
| CAMK2A | FYN | 1 | PMID28212797 | |
| CAMK2A | RYR3 | 1 | PMID23825405 | |
| CAMK2D | FYN | 1 | PMID28212797 | |
| CAMK2D | ITPR2 | 1 | PMID19133301 | PMID23019322 |
| CAMK2D | PLN | 1 | PMID28209766 | |
| CAMK2D | RYR2 | 1 | PMID28209766 | PMID20961976 |
| CAMK2D | SRI | 1 | PMID17707398 | |
| CANX | CSNK2A1 | 1 | PMID10994779 | |
| CANX | CSNK2A2 | 1 | PMID10994779 | |
| CANX | PACS2 | 1 | PMID18417615 | |
| CANX | PDIA3 | 1 | PMID16105684 | |
| CAPN2 | CAPNS1 | 1 | PMID25422446 | |
| CAPN2 | CISD2 | 1 | PMID25422446 | |
| CAPN3 | RYR1 | 1 | PMID18676612 | |
| CAPNS1 | CISD2 | 1 | PMID25422446 | |
| CASP10 | TNFRSF10A | 1 | PMID9325248 | |
| CASP10 | TNFRSF10B | 1 | PMID9325248 | |
| CASP3 | CASP8 | 1 | PMID28049841 | |
| CASP3 | ITPR1 | 1 | PMID17429043 | PMID11013462 |
| CASP3 | ITPR2 | 1 | PMID17429043 | PMID11013462 |
| CASP7 | CASP8 | 1 | PMID28049841 | |
| CASP8 | RYR2 | 1 | PMID21788490 | |
| CASP8 | TNFRSF10A | 1 | PMID9325248 | |
| CASP8 | TNFRSF10B | 1 | PMID9325248 | |
| CASQ1 | CSNK2A1 | 1 | PMID10398759 | PMID1985907 |
| CASQ1 | CSNK2A2 | 1 | PMID10398759 | PMID1985907 |
| CASQ1 | RYR1 | 1 | PMID20961976 | |
| CASQ1 | TRDN | 1 | PMID20961976 | |
| CASQ2 | CSNK2A1 | 1 | PMID10398759 | PMID1985907 |
| CASQ2 | CSNK2A2 | 1 | PMID10398759 | PMID1985907 |
| CASQ2 | PDIA3 | 1 | PMID10398759 | |
| CASQ2 | RYR2 | 1 | PMID20961976 | |
| CASQ2 | TRDN | 1 | PMID20961976 | |
| CAV1 | PIK3C3 | 1 | PMID28542134 | |
| CAV3 | JPH1 | 1 | PMID22020936 | |
| CAV3 | JPH2 | 1 | PMID22020936 | |
| CAV3 | RYR1 | 1 | PMID22020936 | |
| CD38 | RYR1 | 1 | PMID16751979 | |
| CD38 | TRPC1 | 1 | PMID16751979 | |
| CD38 | TRPC3 | 1 | PMID16751979 | |
| CD38 | TRPC4 | 1 | PMID16751979 | |
| CD38 | TRPC5 | 1 | PMID16751979 | |
| CD38 | TRPC6 | 1 | PMID16751979 | |
| CD38 | TRPC7 | 1 | PMID16751979 | |
| CDK1 | ITPR1 | 1 | PMID19133301 | |
| CDK1 | ITPR3 | 1 | PMID19133301 | |
| CHGA | ITPR1 | 1 | PMID17429043 | |
| CHGB | ITPR1 | 1 | PMID17429043 | |
| CIB1 | ITPR1 | 1 | PMID17429043 | |
| CIB1 | ITPR2 | 1 | PMID17429043 | |
| CIB1 | ITPR3 | 1 | PMID17429043 | |
| CLIC2 | RYR1 | 1 | PMID19356589 | |
| CSNK2A1 | CSNK2A2 | 1 | PMID10398759 | |
| CSNK2A1 | CSNK2B | 1 | PMID10398759 | |
| CSNK2A1 | SRL | 1 | PMID10398759 | |
| CSNK2A2 | CSNK2B | 1 | PMID10398759 | |
| CSNK2A2 | SRL | 1 | PMID10398759 | |
| CYBB | RYR2 | 1 | PMID24386930 | |
| EPB41L1 | GRIA1 | 1 | PMID17429043 | |
| EPB41L1 | ITPR1 | 1 | PMID17429043 | |
| ERLIN1 | ERLIN2 | 1 | PMID27378757 | |
| ERLIN1 | ITPR1 | 1 | PMID27378757 | PMID19240031 |
| ERLIN1 | ITPR2 | 1 | PMID27378757 | |
| ERLIN1 | ITPR3 | 1 | PMID27378757 | |
| ERLIN1 | RNF170 | 1 | PMID27378757 | |
| ERLIN2 | ITPR1 | 1 | PMID27378757 | |
| ERLIN2 | ITPR2 | 1 | PMID27378757 | |
| ERLIN2 | ITPR3 | 1 | PMID27378757 | |
| ERLIN2 | RNF170 | 1 | PMID27378757 | |
| ERO1A | ERP44 | 1 | PMID28619231 | |
| ERO1A | ITPR1 | 1 | PMID28619231 | PMID19752026 |
| FBXL2 | ITPR3 | 1 | PMID28614300 | |
| FKBP1A | ITPR1 | 1 | PMID17429043 | |
| FKBP1A | ITPR2 | 1 | PMID17429043 | |
| FKBP1A | ITPR3 | 1 | PMID17429043 | |
| FKBP1A | PPP3CA | 1 | PMID9346894 | |
| FKBP1A | RYR1 | 1 | PMID24509862 | |
| FKBP1A | RYR2 | 1 | PMID24509862 | |
| FKBP1A | RYR3 | 1 | PMID24509862 | |
| FKBP1A | TRPC3 | 1 | PMID23228564 | |
| FKBP1A | TRPC6 | 1 | PMID23228564 | |
| FKBP1B | RYR1 | 1 | PMID20961976 | |
| FKBP1B | RYR2 | 1 | PMID20961976 | |
| FKBP1B | RYR3 | 1 | PMID20961976 | |
| FKBP4 | ORAI1 | 1 | PMID23228564 | |
| FKBP4 | TRPC1 | 1 | PMID23228564 | |
| FKBP4 | TRPC4 | 1 | PMID23228564 | |
| FKBP4 | TRPC5 | 1 | PMID23228564 | |
| FKBP8 | PPP3CA | 1 | PMID15757646 | |
| FYN | ITPR1 | 1 | PMID19133301 | |
| GAPDH | ITPR1 | 1 | PMID17429043 | |
| GRM1 | HOMER1 | 1 | PMID17429043 | PMID12072556 |
| GRM1 | HOMER3 | 1 | PMID17429043 | PMID12072556 |
| GRM5 | HOMER1 | 1 | PMID17429043 | PMID12072556 |
| GRM5 | HOMER3 | 1 | PMID17429043 | PMID12072556 |
| GSTM2 | RYR2 | 1 | PMID18308613 | |
| HAP1 | HTT | 1 | PMID17429043 | |
| HAP1 | ITPR1 | 1 | PMID12877301 | PMID17429043 |
| HOMER1 | ITPR1 | 1 | PMID16226241 | PMID12072556 |
| HOMER1 | RYR1 | 1 | PMID16226241 | PMID 12223488 |
| HOMER1 | TRPC1 | 1 | PMID17429043 | PMID14505576 |
| HOMER2 | RYR1 | 1 | PMID16226241 | PMID12223488 |
| HOMER3 | ITPR1 | 1 | PMID16226241 | PMID12072556 |
| HOMER3 | RYR1 | 1 | PMID16226241 | PMID12223488 |
| HRC | TRDN | 1 | PMID22952658 | |
| HSP90B1 | WFS1 | 1 | PMID19125190 | |
| HSPA5 | ITPR1 | 1 | PMID28619231 | PMID21145001 |
| HSPA5 | ITPR3 | 1 | PMID28619231 | |
| HSPA5 | SIGMAR1 | 1 | PMID28619231 | |
| HTT | ITPR1 | 1 | PMID12877301 | PMID17429043 |
| ITPR1 | ITPRIP | 1 | PMID16990268 | |
| ITPR1 | KRAS | 1 | PMID26772784 | |
| ITPR1 | LYN | 1 | PMID19133301 | |
| ITPR1 | MAPK1 | 1 | PMID19133301 | |
| ITPR1 | MAPK3 | 1 | PMID19133301 | |
| ITPR1 | MCL1 | 1 | PMID26772784 | |
| ITPR1 | MRVI1 | 1 | PMID19133301 | |
| ITPR1 | NCS1 | 1 | PMID21104311 | PMID16691292 |
| ITPR1 | ORAI1 | 1 | PMID25506690 | |
| ITPR1 | PKD2 | 1 | PMID16223735 | |
| ITPR1 | PLK1 | 1 | PMID19133301 | |
| ITPR1 | PRKG1 | 1 | PMID19133301 | PMID10724174 |
| ITPR1 | PTEN | 1 | PMID26772784 | |
| ITPR1 | RACK1 | 1 | PMID25506690 | |
| ITPR1 | RCN1 | 1 | PMID28319095 | |
| ITPR1 | NRIP1 | 1 | PMID25066731 | |
| ITPR1 | RNF170 | 1 | PMID27378757 | PMID21610068 |
| ITPR1 | ROCK2 | 1 | PMID19133301 | |
| ITPR1 | SELENOK | 1 | PMID25368151 | |
| ITPR1 | SIGMAR1 | 1 | PMID28619231 | |
| ITPR1 | SRC | 1 | PMID19133301 | |
| ITPR1 | STIM1 | 1 | PMID25506690 | |
| ITPR1 | STIM2 | 1 | PMID25506690 | |
| ITPR1 | TESPA1 | 1 | PMID28598420 | |
| ITPR1 | TMEM203 | 1 | PMID25996873 | |
| ITPR1 | TRPC3 | 1 | PMID25506690 | |
| ITPR1 | ZDHHC6 | 1 | PMID25368151 | |
| ITPR2 | LYN | 1 | PMID19133301 | |
| ITPR2 | RNF170 | 1 | PMID27378757 | PMID21610068 |
| ITPR2 | ROCK2 | 1 | PMID19133301 | |
| ITPR2 | TRPC1 | 1 | PMID23228564 | |
| ITPR3 | KRAS | 1 | PMID26772784 | |
| ITPR3 | PLK1 | 1 | PMID19133301 | |
| ITPR3 | PML | 1 | PMID26772784 | PMID28515000 |
| ITPR3 | PTEN | 1 | PMID26772784 | |
| ITPR3 | RNF170 | 1 | PMID27378757 | PMID21610068 |
| ITPR3 | ROCK2 | 1 | PMID19133301 | |
| ITPR3 | SIGMAR1 | 1 | PMID28619231 | PMID17981125 |
| JPH1 | RYR1 | 1 | PMID22020936 | |
| JPH2 | RYR1 | 1 | PMID22020936 | |
| JPH2 | RYR2 | 1 | PMID22020936 | |
| JPH2 | SPEG | 1 | PMID27729468 | |
| MRLN | PLN | 1 | PMID26816378 | |
| MRLN | SLN | 1 | PMID26816378 | |
| MRVI1 | PRKG1 | 1 | PMID19133301 | |
| NOD1 | RYR2 | 1 | PMID28126160 | |
| NOX4 | RYR1 | 1 | PMID24386930 | |
| OPRD1 | RYR2 | 1 | PMID25552382 | |
| ORAI1 | RACK1 | 1 | PMID25506690 | |
| ORAI1 | SARAF | 1 | PMID29223474 | |
| ORAI1 | STIM1 | 1 | PMID25506690 | |
| ORAI1 | STIM2 | 1 | PMID25506690 | PMID20807283 |
| ORAI1 | TRPC1 | 1 | PMID23228564 | |
| ORAI1 | TRPC3 | 1 | PMID25506690 | |
| ORAI1 | TRPC4 | 1 | PMID23228564 | |
| ORAI1 | TRPC6 | 1 | PMID23228564 | |
| PDE4D | RYR2 | 1 | PMID19345240 | |
| PDIA3 | STIM1 | 1 | PMID21941299 | PMID28619231 |
| PKD1 | PKD2 | 1 | PMID16223735 | |
| PLN | S100A1 | 1 | PMID12804600 | |
| PPP1R9B | RYR2 | 1 | PMID19345240 | |
| PPP2R3A | RYR2 | 1 | PMID19345240 | |
| PPP3CA | PPP3R1 | 1 | PMID21115349 | |
| PPP3CA | PPP3R2 | 1 | PMID21115349 | |
| PPP3CA | RCAN1 | 1 | PMID12809556 | |
| PPP5C | S100A1 | 1 | PMID27600583 | |
| PRKG1 | RYR2 | 1 | PMID20961976 | |
| RACK1 | STIM1 | 1 | PMID25506690 | |
| RACK1 | TRPC3 | 1 | PMID25506690 | |
| RYR1 | S100A1 | 1 | PMID18650434 | |
| RYR1 | SELENON | 1 | PMID28506443 | |
| RYR1 | TMEM38A | 1 | PMID28387457 | |
| RYR1 | TRPC3 | 1 | PMID16751979 | |
| RYR1 | TRPC5 | 1 | PMID16751979 | |
| RYR2 | S100A1 | 1 | PMID18650434 | |
| RYR2 | SPEG | 1 | PMID27729468 | |
| RYR2 | SRI | 1 | PMID17707398 | |
| RYR2 | TMEM38A | 1 | PMID24526676 | |
| RYR2 | TRDN | 1 | PMID15041652 | |
| SARAF | STIM1 | 1 | PMID29223474 | |
| SARAF | TRPC1 | 1 | PMID29223474 | |
| SELENOK | ZDHHC6 | 1 | PMID25368151 | |
| STIM1 | STIM2 | 1 | PMID25506690 | |
| STIM1 | TMEM110 | 1 | PMID26419802 | |
| STIM1 | TMEM203 | 1 | PMID25996873 | |
| STIM1 | TRPC1 | 1 | PMID25772237 | |
| STIM1 | TRPC3 | 1 | PMID25506690 | |
| TNFRSF10A | TNFSF10 | 1 | PMID9325248 | |
| TNFRSF10B | TNFSF10 | 1 | PMID9325248 | |
| ATP2A2 | CAMK2D | 3 | PMID16105684 | |
| ATP2A2 | CAMK2G | 3 | PMID16105684 | |
| BECN1 | CAMK2A | 3 | PMID28725634 | |
| BECN1 | CAMK2B | 3 | PMID28725634 | |
| BECN1 | CAMK2D | 3 | PMID28725634 | |
| BECN1 | CAMK2G | 3 | PMID28725634 | |
| CALM1 | CAMK2A | 3 | PMID20961976 | |
| CALM1 | CAMK2B | 3 | PMID20961976 | |
| CALM1 | CAMK2D | 3 | PMID20961976 | |
| CALM1 | CAMK2G | 3 | PMID20961976 | |
| CALM2 | CAMK2A | 3 | PMID20961976 | |
| CALM2 | CAMK2B | 3 | PMID20961976 | |
| CALM2 | CAMK2D | 3 | PMID20961976 | |
| CALM2 | CAMK2G | 3 | PMID20961976 | |
| CALM3 | CAMK2A | 3 | PMID20961976 | |
| CALM3 | CAMK2B | 3 | PMID20961976 | |
| CALM3 | CAMK2D | 3 | PMID20961976 | |
| CALM3 | CAMK2G | 3 | PMID20961976 | |
| CAMK2A | PLN | 3 | PMID20961976 | |
| CAMK2A | RYR1 | 3 | PMID20961976 | |
| CAMK2A | RYR2 | 3 | PMID20961976 | |
| CAMK2B | PLN | 3 | PMID20961976 | |
| CAMK2B | RYR1 | 3 | PMID20961976 | |
| CAMK2B | RYR2 | 3 | PMID20961976 | |
| CAMK2D | PLN | 3 | PMID20961976 | |
| CAMK2D | RYR1 | 3 | PMID20961976 | |
| CAMK2G | PLN | 3 | PMID20961976 | |
| CAMK2G | RYR1 | 3 | PMID20961976 | |
| CAMK2G | RYR2 | 3 | PMID20961976 | |
| CISD2 | ITPR1 | 3 | PMID22617439 | |
| CISD2 | ITPR2 | 3 | PMID22617439 | |
| CISD2 | ITPR3 | 3 | PMID22617439 | |
| GRM5 | ITPR2 | 3 | PMID19133301 | |
| GRM5 | ITPR3 | 3 | PMID19133301 | |
| AHCYL1 | CALU | 4 | FC0.92 | |
| AHCYL1 | ERP44 | 4 | FC0.957 | |
| AKAP6 | CALM1 | 4 | FC0.979 | |
| AKAP6 | CALM3 | 4 | FC0.921 | |
| AKAP9 | CALM1 | 4 | FC0.983 | |
| AKAP9 | CALM2 | 4 | FC0.983 | |
| AKAP9 | CALM3 | 4 | FC0.982 | |
| AKT1 | CALM1 | 4 | FC0.957 | |
| AKT1 | CALM2 | 4 | FC0.963 | |
| AKT1 | CALM3 | 4 | FC0.957 | |
| AKT1 | CAMK2A | 4 | FC0.972 | |
| AKT1 | PPP3CA | 4 | FC0.987 | |
| ALDOA | ATP2A2 | 4 | FC0.998 | |
| ALDOA | CALM1 | 4 | FC1.0 | |
| ALDOA | CALM2 | 4 | FC1.0 | |
| ALDOA | CALM3 | 4 | FC1.0 | |
| ALDOA | CALR | 4 | FC1.0 | |
| ALDOA | CALU | 4 | FC0.999 | |
| ALDOA | CANX | 4 | FC0.997 | |
| ALDOA | FKBP1A | 4 | FC1.0 | |
| ALDOA | FKBP1B | 4 | FC0.966 | |
| ALDOA | PDIA3 | 4 | FC1.0 | |
| ALDOA | SRI | 4 | FC0.998 | |
| ALDOA | TMX1 | 4 | FC0.999 | |
| ANK2 | HOMER1 | 4 | FC0.927 | |
| ASPH | ATP2A2 | 4 | FC0.993 | |
| ASPH | CALM1 | 4 | FC0.955 | |
| ASPH | CALM2 | 4 | FC0.945 | |
| ASPH | CALM3 | 4 | FC0.97 | |
| ASPH | CALU | 4 | FC0.999 | |
| ASPH | CANX | 4 | FC0.993 | |
| ASPH | ERP44 | 4 | FC0.999 | |
| ASPH | TMX1 | 4 | FC0.992 | |
| ATP2A1 | ATP2A2 | 4 | FC0.994 | |
| ATP2A1 | ATP2A3 | 4 | FC0.991 | |
| ATP2A1 | HSP90B1 | 4 | FC0.963 | |
| ATP2A1 | PLN | 4 | FC0.918 | |
| ATP2A2 | ATP2A3 | 4 | FC1.0 | |
| ATP2A2 | BAX | 4 | FC0.997 | |
| ATP2A2 | CALM1 | 4 | FC1.0 | |
| ATP2A2 | CALM2 | 4 | FC1.0 | |
| ATP2A2 | CALM3 | 4 | FC1.0 | |
| ATP2A2 | CAPNS1 | 4 | FC0.995 | |
| ATP2A2 | CSNK2A1 | 4 | FC0.929 | |
| ATP2A2 | ERLIN2 | 4 | FC0.986 | |
| ATP2A2 | ERO1A | 4 | FC1.0 | |
| ATP2A2 | ERP44 | 4 | FC0.999 | |
| ATP2A2 | FKBP8 | 4 | FC0.911 | |
| ATP2A2 | GAPDH | 4 | FC1.0 | |
| ATP2A2 | HSP90B1 | 4 | FC1.0 | |
| ATP2A2 | HSPA5 | 4 | FC1.0 | |
| ATP2A2 | ITPR3 | 4 | FC0.9 | |
| ATP2A2 | PDIA3 | 4 | FC0.999 | |
| ATP2A2 | PPP1CA | 4 | FC0.998 | |
| ATP2A2 | RACK1 | 4 | FC0.999 | |
| ATP2A3 | CALM1 | 4 | FC0.988 | |
| ATP2A3 | CALM2 | 4 | FC0.988 | |
| ATP2A3 | CALM3 | 4 | FC0.988 | |
| ATP2A3 | CALR | 4 | FC0.999 | |
| ATP2A3 | CALU | 4 | FC0.954 | |
| ATP2A3 | CANX | 4 | FC0.986 | |
| ATP2A3 | ERLIN2 | 4 | FC0.904 | |
| ATP2A3 | ERO1A | 4 | FC1.0 | |
| ATP2A3 | GAPDH | 4 | FC0.994 | |
| ATP2A3 | HSP90B1 | 4 | FC1.0 | |
| ATP2A3 | HSPA5 | 4 | FC0.984 | |
| ATP2A3 | PDIA3 | 4 | FC0.924 | |
| ATP2A3 | TMX1 | 4 | FC0.996 | |
| BAK1 | BCL2 | 4 | FC0.999 | |
| BAX | BCL2 | 4 | FC1.0 | |
| BAX | CALM1 | 4 | FC1.0 | |
| BAX | CALM2 | 4 | FC1.0 | |
| BAX | CALM3 | 4 | FC1.0 | |
| BAX | CANX | 4 | FC1.0 | |
| BAX | ERP44 | 4 | FC0.992 | |
| BAX | PDIA3 | 4 | FC0.907 | |
| BAX | TMX1 | 4 | FC0.999 | |
| BCL2 | BCL2L1 | 4 | FC1.0 | |
| BCL2 | BID | 4 | FC0.928 | |
| BCL2 | CASP8 | 4 | FC0.927 | |
| BCL2 | MAPK1 | 4 | FC0.969 | |
| BCL2 | MCL1 | 4 | FC0.996 | |
| BCL2 | PSEN1 | 4 | FC0.927 | |
| BECN1 | CALM1 | 4 | FC0.999 | |
| BECN1 | CALM2 | 4 | FC0.999 | |
| BECN1 | CALM3 | 4 | FC0.998 | |
| BECN1 | CALR | 4 | FC0.99 | |
| CACNA1S | CALM1 | 4 | FC0.942 | |
| CACNA1S | CALM2 | 4 | FC0.915 | |
| CACNA1S | CALM3 | 4 | FC0.942 | |
| CALM1 | CALM2 | 4 | FC0.918 | |
| CALM1 | CALM3 | 4 | FC0.992 | |
| CALM1 | CALR | 4 | FC1.0 | |
| CALM1 | CALU | 4 | FC1.0 | |
| CALM1 | CANX | 4 | FC1.0 | |
| CALM1 | CAPNS1 | 4 | FC0.999 | |
| CALM1 | CSNK2A1 | 4 | FC1.0 | |
| CALM1 | CSNK2A2 | 4 | FC0.924 | |
| CALM1 | CSNK2B | 4 | FC1.0 | |
| CALM1 | ERLIN2 | 4 | FC0.998 | |
| CALM1 | ERO1A | 4 | FC1.0 | |
| CALM1 | ERP44 | 4 | FC0.95 | |
| CALM1 | FKBP1A | 4 | FC1.0 | |
| CALM1 | FKBP1B | 4 | FC0.989 | |
| CALM1 | FKBP4 | 4 | FC1.0 | |
| CALM1 | GAPDH | 4 | FC1.0 | |
| CALM1 | GRM5 | 4 | FC0.998 | |
| CALM1 | HOMER1 | 4 | FC0.93 | |
| CALM1 | HSP90B1 | 4 | FC1.0 | |
| CALM1 | HSPA5 | 4 | FC1.0 | |
| CALM1 | HTT | 4 | FC0.998 | |
| CALM1 | KRAS | 4 | FC1.0 | |
| CALM1 | LYN | 4 | FC0.985 | |
| CALM1 | MAPK1 | 4 | FC1.0 | |
| CALM1 | MAPK3 | 4 | FC0.998 | |
| CALM1 | PDIA3 | 4 | FC1.0 | |
| CALM1 | PLK1 | 4 | FC0.99 | |
| CALM1 | PPP1CA | 4 | FC1.0 | |
| CALM1 | PPP3R1 | 4 | FC1.0 | |
| CALM1 | PPP3R2 | 4 | FC0.985 | |
| CALM1 | PPP5C | 4 | FC0.999 | |
| CALM1 | RACK1 | 4 | FC1.0 | |
| CALM1 | SDF4 | 4 | FC0.975 | |
| CALM1 | SRI | 4 | FC0.996 | |
| CALM1 | TMBIM6 | 4 | FC0.986 | |
| CALM1 | TMX1 | 4 | FC0.984 | |
| CALM2 | CALR | 4 | FC1.0 | |
| CALM2 | CALU | 4 | FC1.0 | |
| CALM2 | CANX | 4 | FC1.0 | |
| CALM2 | CAPNS1 | 4 | FC0.999 | |
| CALM2 | CSNK2A1 | 4 | FC1.0 | |

| | | | |
|---|---|---|---|
| CALM2 | CSNK2A2 | 4 | FC0.922 |
| CALM2 | CSNK2B | 4 | FC1.0 |
| CALM2 | ERLIN2 | 4 | FC0.997 |
| CALM2 | ERO1A | 4 | FC1.0 |
| CALM2 | FKBP1A | 4 | FC1.0 |
| CALM2 | FKBP1B | 4 | FC0.99 |
| CALM2 | FKBP4 | 4 | FC1.0 |
| CALM2 | GAPDH | 4 | FC1.0 |
| CALM2 | GRM5 | 4 | FC0.99 |
| CALM2 | HSP90B1 | 4 | FC1.0 |
| CALM2 | HSPA5 | 4 | FC1.0 |
| CALM2 | HTT | 4 | FC0.998 |
| CALM2 | ITPR1 | 4 | FC0.982 |
| CALM2 | KRAS | 4 | FC0.999 |
| CALM2 | LYN | 4 | FC0.986 |
| CALM2 | MAPK1 | 4 | FC1.0 |
| CALM2 | MAPK3 | 4 | FC0.999 |
| CALM2 | PDIA3 | 4 | FC1.0 |
| CALM2 | PLK1 | 4 | FC0.99 |
| CALM2 | PPP1CA | 4 | FC1.0 |
| CALM2 | PPP3R1 | 4 | FC1.0 |
| CALM2 | PPP3R2 | 4 | FC0.985 |
| CALM2 | PPP5C | 4 | FC0.998 |
| CALM2 | RACK1 | 4 | FC1.0 |
| CALM2 | RYR2 | 4 | FC1.0 |
| CALM2 | SDF4 | 4 | FC0.971 |
| CALM2 | SRI | 4 | FC0.998 |
| CALM2 | TMBIM6 | 4 | FC0.976 |
| CALM2 | TMX1 | 4 | FC0.989 |
| CALM3 | CALR | 4 | FC1.0 |
| CALM3 | CALU | 4 | FC1.0 |
| CALM3 | CANX | 4 | FC1.0 |
| CALM3 | CAPNS1 | 4 | FC0.999 |
| CALM3 | CSNK2A1 | 4 | FC1.0 |
| CALM3 | CSNK2A2 | 4 | FC0.922 |
| CALM3 | CSNK2B | 4 | FC1.0 |
| CALM3 | ERLIN2 | 4 | FC0.997 |
| CALM3 | ERO1A | 4 | FC1.0 |
| CALM3 | FKBP1A | 4 | FC1.0 |
| CALM3 | FKBP1B | 4 | FC0.989 |
| CALM3 | FKBP4 | 4 | FC1.0 |
| CALM3 | GAPDH | 4 | FC1.0 |
| CALM3 | GRM5 | 4 | FC0.998 |
| CALM3 | HOMER1 | 4 | FC0.906 |
| CALM3 | HSP90B1 | 4 | FC1.0 |
| CALM3 | HSPA5 | 4 | FC1.0 |
| CALM3 | HTT | 4 | FC0.998 |
| CALM3 | ITPR1 | 4 | FC0.907 |
| CALM3 | KRAS | 4 | FC1.0 |
| CALM3 | LYN | 4 | FC0.986 |
| CALM3 | MAPK1 | 4 | FC1.0 |
| CALM3 | MAPK3 | 4 | FC0.999 |
| CALM3 | PDIA3 | 4 | FC1.0 |
| CALM3 | PLK1 | 4 | FC0.991 |
| CALM3 | PPP1CA | 4 | FC1.0 |
| CALM3 | PPP3R1 | 4 | FC1.0 |
| CALM3 | PPP3R2 | 4 | FC0.985 |
| CALM3 | PPP5C | 4 | FC0.999 |
| CALM3 | RACK1 | 4 | FC1.0 |
| CALM3 | RYR2 | 4 | FC1.0 |
| CALM3 | SDF4 | 4 | FC0.975 |
| CALM3 | SRI | 4 | FC0.997 |
| CALM3 | TMBIM6 | 4 | FC0.959 |
| CALM3 | TMX1 | 4 | FC0.986 |
| CALR | CALU | 4 | FC1.0 |
| CALR | CANX | 4 | FC1.0 |
| CALR | CAPN2 | 4 | FC0.996 |
| CALR | ERLIN2 | 4 | FC0.992 |
| CALR | ERO1A | 4 | FC1.0 |
| CALR | ERP44 | 4 | FC1.0 |
| CALR | FKBP1A | 4 | FC1.0 |
| CALR | FKBP4 | 4 | FC1.0 |
| CALR | GAPDH | 4 | FC1.0 |
| CALR | HSP90B1 | 4 | FC1.0 |
| CALR | HSPA5 | 4 | FC1.0 |
| CALR | MAPK1 | 4 | FC0.942 |
| CALR | PPP1CA | 4 | FC1.0 |
| CALR | PSEN1 | 4 | FC0.984 |
| CALR | RACK1 | 4 | FC1.0 |
| CALR | SDF4 | 4 | FC0.999 |
| CALR | TMX1 | 4 | FC1.0 |
| CALU | CANX | 4 | FC1.0 |
| CALU | CAPN2 | 4 | FC0.982 |
| CALU | CISD2 | 4 | FC0.942 |
| CALU | CSNK2A1 | 4 | FC0.996 |
| CALU | CSNK2A2 | 4 | FC0.998 |
| CALU | DNAJC10 | 4 | FC0.999 |
| CALU | ERLIN1 | 4 | FC0.906 |
| CALU | ERLIN2 | 4 | FC1.0 |
| CALU | ERO1A | 4 | FC1.0 |
| CALU | ERP44 | 4 | FC1.0 |
| CALU | FKBP1A | 4 | FC0.966 |
| CALU | FKBP4 | 4 | FC0.996 |
| CALU | GAPDH | 4 | FC0.997 |
| CALU | HSP90B1 | 4 | FC1.0 |
| CALU | HSPA5 | 4 | FC1.0 |
| CALU | MAPK1 | 4 | FC0.987 |
| CALU | PDIA3 | 4 | FC1.0 |
| CALU | PML | 4 | FC0.963 |
| CALU | PPP5C | 4 | FC1.0 |
| CALU | SDF4 | 4 | FC1.0 |
| CALU | SRI | 4 | FC0.937 |
| CALU | TMX1 | 4 | FC1.0 |
| CAMK2A | CAMK2B | 4 | FC1.0 |
| CAMK2A | CAMK2D | 4 | FC1.0 |
| CAMK2A | CAMK2G | 4 | FC1.0 |
| CAMK2A | GRIA1 | 4 | FC0.996 |
| CAMK2A | PPP3CA | 4 | FC0.96 |
| CAMK2B | CAMK2D | 4 | FC1.0 |
| CAMK2B | CAMK2G | 4 | FC0.999 |
| CAMK2B | GRIA1 | 4 | FC0.978 |
| CAMK2D | CAMK2G | 4 | FC1.0 |
| CAMK2D | FKBP4 | 4 | FC0.922 |
| CAMK2D | HSPA5 | 4 | FC0.977 |
| CAMK2D | ITPR3 | 4 | FC0.985 |
| CAMK2D | MAPK1 | 4 | FC0.993 |
| CAMK2D | PPP1CA | 4 | FC0.985 |
| CAMK2D | PPP3CA | 4 | FC1.0 |
| CAMK2G | GRIA1 | 4 | FC0.998 |
| CAMK2G | ITPR1 | 4 | FC0.907 |
| CAMK2G | PPP3CA | 4 | FC0.94 |
| CAMK2G | RACK1 | 4 | FC0.908 |
| CANX | CAPN2 | 4 | FC0.96 |
| CANX | CAPNS1 | 4 | FC0.997 |
| CANX | DNAJC10 | 4 | FC0.988 |
| CANX | ERLIN2 | 4 | FC1.0 |
| CANX | ERO1A | 4 | FC1.0 |
| CANX | ERP44 | 4 | FC1.0 |
| CANX | FKBP1A | 4 | FC0.988 |
| CANX | FKBP4 | 4 | FC1.0 |
| CANX | FKBP8 | 4 | FC0.982 |
| CANX | GAPDH | 4 | FC1.0 |
| CANX | HSP90B1 | 4 | FC1.0 |
| CANX | HSPA5 | 4 | FC1.0 |
| CANX | ITPR3 | 4 | FC0.993 |
| CANX | MAPK1 | 4 | FC0.997 |
| CANX | PKD2 | 4 | FC0.997 |
| CANX | PPP1CA | 4 | FC0.917 |
| CANX | PPP5C | 4 | FC0.921 |
| CANX | RACK1 | 4 | FC0.999 |
| CANX | SDF4 | 4 | FC0.998 |
| CANX | TMBIM6 | 4 | FC0.998 |
| CANX | TMX1 | 4 | FC0.999 |
| CANX | WFS1 | 4 | FC0.942 |
| CAPN2 | FKBP1A | 4 | FC0.93 |
| CAPN2 | PDIA3 | 4 | FC0.995 |
| CAPN2 | SRI | 4 | FC0.954 |
| CAPNS1 | PDIA3 | 4 | FC0.908 |
| CASQ1 | CAV3 | 4 | FC0.96 |
| CASQ1 | JPH2 | 4 | FC0.921 |
| CASQ2 | JPH2 | 4 | FC0.959 |
| CAV1 | ITPR3 | 4 | FC1.0 |
| CDK1 | ERP44 | 4 | FC1.0 |
| CISD2 | ERP44 | 4 | FC0.952 |
| CISD2 | PDIA3 | 4 | FC0.961 |
| CISD2 | TMX1 | 4 | FC0.942 |
| CSNK2A1 | PDIA3 | 4 | FC0.997 |
| CSNK2A1 | PPP1R9B | 4 | FC0.97 |
| CSNK2A1 | PPP3CA | 4 | FC0.927 |
| CSNK2A1 | SRI | 4 | FC0.999 |
| CSNK2B | PDIA3 | 4 | FC0.935 |
| CSNK2B | PPP1R9B | 4 | FC0.943 |
| CSNK2B | SRI | 4 | FC0.982 |
| DNAJC10 | HSPA5 | 4 | FC1.0 |
| DNAJC10 | SDF4 | 4 | FC0.905 |
| ERLIN2 | ERP44 | 4 | FC0.984 |
| ERLIN2 | FKBP1A | 4 | FC0.963 |
| ERLIN2 | PDIA3 | 4 | FC0.999 |
| ERLIN2 | SRI | 4 | FC0.999 |
| ERLIN2 | TMX1 | 4 | FC0.999 |
| ERO1A | PDIA3 | 4 | FC1.0 |
| ERO1A | TMX1 | 4 | FC1.0 |
| ERP44 | FKBP1A | 4 | FC0.98 |
| ERP44 | FKBP4 | 4 | FC0.984 |
| ERP44 | FKBP8 | 4 | FC0.99 |
| ERP44 | HSP90B1 | 4 | FC1.0 |
| ERP44 | HSPA5 | 4 | FC1.0 |
| ERP44 | ITPR1 | 4 | FC0.985 |
| ERP44 | MAPK1 | 4 | FC0.975 |
| ERP44 | PDIA3 | 4 | FC1.0 |
| ERP44 | PPP1CA | 4 | FC1.0 |
| ERP44 | SDF4 | 4 | FC0.998 |
| ERP44 | SRI | 4 | FC0.973 |
| ERP44 | TMX1 | 4 | FC0.992 |
| FKBP1A | FKBP1B | 4 | FC0.976 |
| FKBP1A | FKBP4 | 4 | FC1.0 |
| FKBP1A | GAPDH | 4 | FC0.999 |
| FKBP1A | HSP90B1 | 4 | FC1.0 |
| FKBP1A | HSPA5 | 4 | FC0.999 |
| FKBP1A | MAPK1 | 4 | FC1.0 |
| FKBP1A | MAPK3 | 4 | FC0.995 |
| FKBP1A | PDIA3 | 4 | FC0.999 |
| FKBP1A | PPP5C | 4 | FC0.928 |
| FKBP1A | RACK1 | 4 | FC0.997 |
| FKBP1A | SRI | 4 | FC0.907 |
| FKBP1B | GAPDH | 4 | FC0.933 |
| FKBP4 | PDIA3 | 4 | FC0.997 |
| FKBP4 | SRI | 4 | FC0.999 |
| FKBP4 | STIM1 | 4 | FC0.936 |
| FKBP4 | TMX1 | 4 | FC0.977 |
| GAPDH | PDIA3 | 4 | FC1.0 |
| GAPDH | SRI | 4 | FC0.985 |
| GAPDH | TMX1 | 4 | FC0.998 |
| GRM1 | HOMER2 | 4 | FC0.992 |
| HOMER1 | HOMER2 | 4 | FC1.0 |
| HOMER1 | HOMER3 | 4 | FC0.997 |
| HOMER1 | PPP3CA | 4 | FC0.992 |
| HSP90B1 | PDIA3 | 4 | FC1.0 |
| HSP90B1 | RCN1 | 4 | FC1.0 |
| HSP90B1 | SDF4 | 4 | FC1.0 |
| HSP90B1 | SRI | 4 | FC0.956 |
| HSP90B1 | TMX1 | 4 | FC1.0 |
| HSPA5 | PDIA3 | 4 | FC1.0 |
| HSPA5 | PPP3CA | 4 | FC0.999 |
| HSPA5 | RCN1 | 4 | FC1.0 |
| HSPA5 | SDF4 | 4 | FC1.0 |
| HSPA5 | SRI | 4 | FC1.0 |
| HSPA5 | TMX1 | 4 | FC0.994 |
| ITPR1 | ITPR3 | 4 | FC1.0 |
| ITPR1 | PPP3CA | 4 | FC1.0 |
| ITPR1 | RYR2 | 4 | FC0.987 |
| ITPR1 | TRPC4 | 4 | FC0.966 |
| ITPR3 | TMX1 | 4 | FC0.998 |
| ITPR3 | TRPC4 | 4 | FC0.958 |
| JPH1 | TMEM38A | 4 | FC0.904 |
| KRAS | PDIA3 | 4 | FC0.947 |
| LYN | TMX1 | 4 | FC0.945 |
| MAPK1 | PDIA3 | 4 | FC1.0 |
| MAPK1 | PPP1R9B | 4 | FC1.0 |
| MAPK1 | PPP3CA | 4 | FC0.981 |
| MAPK1 | SRI | 4 | FC0.999 |
| MAPK1 | TMX1 | 4 | FC0.923 |
| MAPK3 | PDIA3 | 4 | FC0.991 |
| PDIA3 | PPP1CA | 4 | FC0.984 |
| PDIA3 | RACK1 | 4 | FC1.0 |
| PDIA3 | TMX1 | 4 | FC1.0 |
| PKD2 | TRPC1 | 4 | FC0.936 |
| PPP1CA | PPP1R9B | 4 | FC1.0 |
| PPP1CA | SRI | 4 | FC1.0 |
| PPP1R9B | PPP3CA | 4 | FC0.994 |
| RACK1 | RYR2 | 4 | FC0.933 |
| RACK1 | TMX1 | 4 | FC0.915 |
| RYR1 | RYR2 | 4 | FC0.999 |
| RYR1 | TRDN | 4 | FC0.992 |
| RYR2 | RYR3 | 4 | FC0.999 |
| SDF4 | TMX1 | 4 | FC0.985 |
| TMX1 | ZDHHC6 | 4 | FC0.908 |
| TRPC1 | TRPC3 | 4 | FC0.919 |
| TRPC4 | TRPC5 | 4 | FC0.968 |

II. Accession numbers for homologs of proteins and protein families key to calcium storage and signaling, with their phylogenies

Accession numbers which were included in the phylogenetic analyses presented in the paper are provided below in clusters which correspond to the 34 proteins and protein families studied. Accession numbers are further annotated by the following columns in order from left to right: length, GenBank gene name, phylogeny, genus, species, and GenBank defline. Some accessions (in the format Aaaa##########) correspond to translationed gene predictions done by our group for genomes without published protein predictions.

A. Calmodulin family

| | | | | | | |
|---|---|---|---|---|---|---|
| NP_001031523.1 | 289 | ARABIDOPSIS,THALIANA,CALMODULIN,LIKE,4,ATCAL4,calmodulin-like,12,CALMODULIN-RELATED,PROTEIN,CML12,T3K9.13,T3K9_13,TOUCH,3 | eukaryota>viridiplantae | Arabidopsis | thaliana | Calcium-binding EF hand family protein [Arabidopsis thaliana] |
| NP_001318400.1 | 289 | ARABIDOPSIS,THALIANA,CALMODULIN,LIKE,4,ATCAL4,calmodulin-like,12,CALMODULIN-RELATED,PROTEIN,CML12,T3K9.13,T3K9_13,TOUCH,3 | eukaryota>viridiplantae | Arabidopsis | thaliana | Calcium-binding EF hand family protein [Arabidopsis thaliana] |
| NP_001324806.1 | 289 | ARABIDOPSIS,THALIANA,CALMODULIN,LIKE,4,ATCAL4,calmodulin-like,12,CALMODULIN-RELATED,PROTEIN,CML12,T3K9.13,T3K9_13,TOUCH,3 | eukaryota>viridiplantae | Arabidopsis | thaliana | Calcium-binding EF hand family protein [Arabidopsis thaliana] |
| XP_015838420.1 | 276 | LOC659938 | eukaryota>metazoa>hexapoda | Tribolium | castaneum | PREDICTED: calmodulin-beta isoform X1 [Tribolium castaneum] |
| XP_624589.2 | 276 | LOC552209 | eukaryota>metazoa>hexapoda | Apis | mellifera | PREDICTED: calmodulin-beta-like isoform X1 [Apis mellifera] |
| NP_660201.1 | 275 | CRSD,CSNB2B | eukaryota>metazoa>chordata>vertebrata | Homo | sapiens | calcium-binding protein 4 isoform a [Homo sapiens] |
| XP_003273984.1 | 275 | CABP4 | eukaryota>metazoa>chordata>vertebrata | Nomascus | leucogenys | PREDICTED: calcium-binding protein 4 [Nomascus leucogenys] |
| XP_005577127.1 | 275 | CABP4 | eukaryota>metazoa>chordata>vertebrata | Macaca | fascicularis | PREDICTED: calcium-binding protein 4 isoform X2 [Macaca fascicularis] |
| XP_011150174.1 | 274 | LOC105189634 | eukaryota>metazoa>hexapoda | Harpegnathos | saltator | PREDICTED: calmodulin-beta isoform X1 [Harpegnathos saltator] |
| OQV24127.1 | 274 | BV898_02079 | eukaryota>metazoa | Hypsibius | dujardini | putative Calmodulin [Hypsibius dujardini] |
| XP_016771317.1 | 268 | LOC552209 | eukaryota>metazoa>hexapoda | Apis | mellifera | PREDICTED: calmodulin-A-like isoform X2 [Apis mellifera] |
| XP_002602906.1 | 266 | BRAFLDRAFT_98094 | eukaryota>metazoa>chordata | Branchiostoma | floridae | hypothetical protein BRAFLDRAFT_98094 [Branchiostoma floridae] |
| KXJ07556.1 | 265 | AC249_AIPGENE20456 | eukaryota>metazoa>cnidaria | Exaiptasia | pallida | Calmodulin [Exaiptasia pallida] |
| XP_015766476.1 | 265 | LOC107345284 | eukaryota>metazoa>cnidaria | Acropora | digitifera | PREDICTED: calmodulin-like [Acropora digitifera] |
| KXJ21637.1 | 263 | AC249_AIPGENE18020 | eukaryota>metazoa>cnidaria | Exaiptasia | pallida | Calmodulin [Exaiptasia pallida] |
| XP_006558596.1 | 262 | LOC552209 | eukaryota>metazoa>hexapoda | Apis | mellifera | PREDICTED: neo-calmodulin-like isoform X3 [Apis mellifera] |
| XP_008197239.1 | 262 | LOC659938 | eukaryota>metazoa>hexapoda | Tribolium | castaneum | PREDICTED: neo-calmodulin isoform X2 [Tribolium castaneum] |
| XP_016875727.1 | 262 | CALBRAIN,HCALB_BR | eukaryota>metazoa>chordata>vertebrata | Homo | sapiens | PREDICTED: calcium-binding protein 1 isoform X4 [Homo sapiens] |
| XP_008197240.1 | 260 | LOC659938 | eukaryota>metazoa>hexapoda | Tribolium | castaneum | PREDICTED: neo-calmodulin isoform X3 [Tribolium castaneum] |
| XP_011150175.1 | 260 | LOC105189634 | eukaryota>metazoa>hexapoda | Harpegnathos | saltator | PREDICTED: calmodulin-A isoform X2 [Harpegnathos saltator] |
| XP_016771318.1 | 259 | LOC552209 | eukaryota>metazoa>hexapoda | Apis | mellifera | PREDICTED: neo-calmodulin-like isoform X4 [Apis mellifera] |
| XP_001948572.2 | 259 | LOC100162151 | eukaryota>metazoa>hexapoda | Acyrthosiphon | pisum | PREDICTED: neo-calmodulin-like isoform X1 [Acyrthosiphon pisum] |
| XP_015141709.1 | 258 | CABP2 | eukaryota>metazoa>chordata>vertebrata | Gallus | gallus | PREDICTED: calcium-binding protein 2 isoform X1 [Gallus gallus] |
| XP_006531829.1 | 256 | Cabp2 | eukaryota>metazoa>chordata>vertebrata | Mus | musculus | PREDICTED: calcium-binding protein 2 isoform X4 [Mus musculus] |
| XP_006531830.1 | 256 | Cabp2 | eukaryota>metazoa>chordata>vertebrata | Mus | musculus | PREDICTED: calcium-binding protein 2 isoform X4 [Mus musculus] |
| XP_012356868.1 | 256 | CALM2 | eukaryota>metazoa>chordata>vertebrata | Nomascus | leucogenys | PREDICTED: calmodulin isoform X1 [Nomascus leucogenys] |
| XP_003074444.1 | 255 | Ot01g03840 | eukaryota>viridiplantae>chlorophyta | Ostreococcus | tauri | calmodulin mutant SYNCAM9 (ISS) [Ostreococcus tauri] |
| XP_004927631.1 | 255 | LOC101742890 | eukaryota>metazoa>hexapoda | Bombyx | mori | neo-calmodulin [Bombyx mori] |
| XP_006558597.1 | 254 | LOC552209 | eukaryota>metazoa>hexapoda | Apis | mellifera | PREDICTED: neo-calmodulin-like isoform X5 [Apis mellifera] |
| XP_011608941.1 | 253 | LOC101063225 | eukaryota>metazoa>chordata>vertebrata>actinopterygii | Takifugu | rubripes | PREDICTED: calcium-binding protein 2-like [Takifugu rubripes] |
| KXJ25153.1 | 252 | AC249_AIPGENE22463 | eukaryota>metazoa>cnidaria | Exaiptasia | pallida | Calmodulin [Exaiptasia pallida] |
| XP_018099550.1 | 247 | LOC108707103 | eukaryota>metazoa>chordata>vertebrata | Xenopus | laevis | PREDICTED: calcium-binding protein 1-like isoform X2 [Xenopus laevis] |
| XP_971297.2 | 246 | LOC659938 | eukaryota>metazoa>hexapoda | Tribolium | castaneum | PREDICTED: neo-calmodulin isoform X4 [Tribolium castaneum] |
| XP_006558598.1 | 245 | LOC552209 | eukaryota>metazoa>hexapoda | Apis | mellifera | PREDICTED: neo-calmodulin-like isoform X6 [Apis mellifera] |
| XP_016657504.1 | 245 | LOC100162151 | eukaryota>metazoa>hexapoda | Acyrthosiphon | pisum | PREDICTED: neo-calmodulin-like isoform X2 [Acyrthosiphon pisum] |
| XP_011150176.1 | 244 | LOC105189634 | eukaryota>metazoa>hexapoda | Harpegnathos | saltator | PREDICTED: neo-calmodulin isoform X3 [Harpegnathos saltator] |
| XP_002161262.1 | 243 | LOC100215966 | eukaryota>metazoa>cnidaria | Hydra | vulgaris | PREDICTED: calmodulin-A-like [Hydra vulgaris] |
| Smar1000005545 | 243 | Smar1000005545 | eukaryota>metazoa>arthropoda>myriapoda | Strigamia | maritima | >Smar1000005545 |
| XP_002168034.1 | 242 | LOC100203928 | eukaryota>metazoa>cnidaria | Hydra | vulgaris | PREDICTED: calmodulin-like [Hydra vulgaris] |
| XP_002503099.1 | 242 | MICPUN_108387 | eukaryota>viridiplantae>chlorophyta | Micromonas | commoda | predicted protein [Micromonas commoda] |
| CAF93636.1 | 242 | GSTEN:00009382:G:001 | eukaryota>metazoa>chordata>vertebrata>actinopterygii | Tetraodon | nigroviridis | unnamed protein product, partial [Tetraodon nigroviridis] |
| XP_016440228.1 | 241 | LOC107766031 | eukaryota>viridiplantae | Nicotiana | tabacum | PREDICTED: calmodulin-like protein 3 [Nicotiana tabacum] |
| XP_016463804.1 | 241 | LOC107786814 | eukaryota>viridiplantae | Nicotiana | tabacum | PREDICTED: calmodulin-like protein 3 [Nicotiana tabacum] |
| XP_003379560.1 | 241 | Tsp_03217 | eukaryota>metazoa>nematoda | Trichinella | spiralis | calmodulin-1 [Trichinella spiralis] |
| XP_016450019.1 | 240 | LOC107774876 | eukaryota>viridiplantae | Nicotiana | tabacum | PREDICTED: calmodulin-like protein 3 [Nicotiana tabacum] |
| XP_016479889.1 | 240 | LOC107801119 | eukaryota>viridiplantae | Nicotiana | tabacum | PREDICTED: calmodulin-like protein 3 [Nicotiana tabacum] |
| XP_017211029.1 | 240 | zgc:112300 | eukaryota>metazoa>chordata>vertebrata>actinopterygii | Danio | rerio | PREDICTED: calcium-binding protein 2 isoform X1 [Danio rerio] |
| XP_003372739.1 | 239 | Tsp_11726 | eukaryota>metazoa>nematoda | Trichinella | spiralis | EF hand domain containing protein [Trichinella spiralis] |
| XP_017328917.1 | 238 | LOC108268443 | eukaryota>metazoa>chordata>vertebrata>actinopterygii | Ictalurus | punctatus | PREDICTED: calcium-binding protein 2-like isoform X1 [Ictalurus punctatus] |
| XP_017338502.1 | 238 | LOC108273649 | eukaryota>metazoa>chordata>vertebrata>actinopterygii | Ictalurus | punctatus | PREDICTED: calcium-binding protein 5-like [Ictalurus punctatus] |
| XP_006345220.1 | 237 | LOC102595634 | eukaryota>viridiplantae | Solanum | tuberosum | PREDICTED: calmodulin-like protein 3 [Solanum tuberosum] |
| XP_003064500.1 | 237 | MICPUCDRAFT_54225 | eukaryota>viridiplantae>chlorophyta | Micromonas | pusilla CCMP1545 | predicted protein [Micromonas pusilla CCMP1545] |
| NP_001025439.2 | 236 | zgc:112300 | eukaryota>metazoa>chordata>vertebrata>actinopterygii | Danio | rerio | calcium-binding protein 2 [Danio rerio] |
| XP_005160309.1 | 236 | zgc:112300 | eukaryota>metazoa>chordata>vertebrata>actinopterygii | Danio | rerio | PREDICTED: calcium-binding protein 2 isoform X2 [Danio rerio] |
| NP_001255489.1 | 236 | cal-4 | eukaryota>metazoa>nematoda | Caenorhabditis | elegans | CALmodulin related genes [Caenorhabditis elegans] |
| XP_003539163.2 | 236 | LOC100780797 | eukaryota>viridiplantae | Glycine | max | PREDICTED: probable calcium-binding protein CML17 [Glycine max] |
| XP_011401016.1 | 235 | F751_0028 | eukaryota>viridiplantae>chlorophyta | Auxenochlorella | protothecoides | Calmodulin [Auxenochlorella protothecoides] |
| OQV22258.1 | 235 | BV898_03761 | eukaryota>metazoa | Hypsibius | dujardini | putative Calmodulin [Hypsibius dujardini] |
| NP_001167949.1 | 234 | GRMZM2G072052 | eukaryota>viridiplantae | Zea | mays | uncharacterized protein LOC100381663 [Zea mays] |
| NP_500421.2 | 234 | cal-3 | eukaryota>metazoa>nematoda | Caenorhabditis | elegans | CALmodulin related genes [Caenorhabditis elegans] |
| XP_006358302.1 | 234 | LOC102585648 | eukaryota>viridiplantae | Solanum | tuberosum | PREDICTED: calmodulin-like protein 3 [Solanum tuberosum] |
| XP_015296417.1 | 234 | CALM3 | eukaryota>metazoa>chordata>vertebrata | Macaca | fascicularis | PREDICTED: calmodulin isoform X1 [Macaca fascicularis] |
| XP_003641327.1 | 234 | CABP4 | eukaryota>metazoa>chordata>vertebrata | Gallus | gallus | PREDICTED: calcium-binding protein 2 [Gallus gallus] |
| XP_009291662.1 | 233 | zgc:112300 | eukaryota>metazoa>chordata>vertebrata>actinopterygii | Danio | rerio | calcium-binding protein 2 isoform X1 [Danio rerio] |
| XP_017318617.1 | 232 | cabp2 | eukaryota>metazoa>chordata>vertebrata>actinopterygii | Ictalurus | punctatus | PREDICTED: calcium-binding protein 2 isoform X1 [Ictalurus punctatus] |
| XP_016515675.1 | 231 | LOC107832358 | eukaryota>viridiplantae | Nicotiana | tabacum | PREDICTED: calmodulin-like protein 1 [Nicotiana tabacum] |
| XP_017318618.1 | 231 | cabp2 | eukaryota>metazoa>chordata>vertebrata>actinopterygii | Ictalurus | punctatus | PREDICTED: calcium-binding protein 2 isoform X2 [Ictalurus punctatus] |
| XP_011610696.1 | 230 | LOC101066819 | eukaryota>metazoa>chordata>vertebrata>actinopterygii | Takifugu | rubripes | PREDICTED: calcium-binding protein 2-like [Takifugu rubripes] |
| XP_017318619.1 | 230 | cabp2 | eukaryota>metazoa>chordata>vertebrata>actinopterygii | Ictalurus | punctatus | PREDICTED: calcium-binding protein 2 isoform X3 [Ictalurus punctatus] |
| XP_016487971.1 | 230 | LOC107808003 | eukaryota>viridiplantae | Nicotiana | tabacum | PREDICTED: probable calcium-binding protein CML25 [Nicotiana tabacum] |
| NP_001240157.1 | 229 | LOC100810081 | [illegible] | [illegible] | [illegible] | [illegible] |
| XP_003549743.1 | 229 | LOC100803032 | [illegible] | [illegible] | [illegible] | [illegible] |
| CAF96460.1 | 229 | GSTEN:00013499:G:001 | eukaryota>metazoa>chordata>vertebrata>actinopterygii | Tetraodon | nigroviridis | unnamed protein product [Tetraodon nigroviridis] |
| XP_009037109.1 | 229 | AURANDRAFT_26147 | eukaryota>stramenopiles | Aureococcus | anophagefferens | hypothetical protein AURANDRAFT_26147 [Aureococcus anophagefferens] |
| XP_003372845.1 | 228 | Tsp_10406 | eukaryota>metazoa>nematoda | Trichinella | spiralis | calmodulin [Trichinella spiralis] |
| XP_006823622.1 | 228 | LOC102802917 | eukaryota>metazoa>hemichordata | Saccoglossus | kowalevskii | PREDICTED: calmodulin-B-like [Saccoglossus kowalevskii] |
| NP_038907.1 | 227 | caldendrin | eukaryota>metazoa>chordata>vertebrata | Mus | musculus | calcium-binding protein 1 isoform 2 [Mus musculus] |
| NP_112482.1 | 227 | CALBRAIN,HCALB_BR | eukaryota>metazoa>chordata>vertebrata | Homo | sapiens | calcium-binding protein 1 isoform 1 [Homo sapiens] |
| XP_003280334.1 | 227 | CABP1 | eukaryota>metazoa>chordata>vertebrata | Nomascus | leucogenys | PREDICTED: calcium-binding protein 1 isoform X1 [Nomascus leucogenys] |
| XP_005572469.1 | 227 | CABP1 | eukaryota>metazoa>chordata>vertebrata | Macaca | fascicularis | PREDICTED: calcium-binding protein 1 isoform X2 [Macaca fascicularis] |
| CDS36359.1 | 227 | EmuJ_000344500 | eukaryota>metazoa | Echinococcus | multilocularis | calmodulin [Echinococcus multilocularis] |
| ETO13322.1 | 227 | RFI_24053 | eukaryota>rhizaria | Reticulomyxa | filosa | calmodulin 2-like protein [Reticulomyxa filosa] |
| NP_001305425.1 | 226 | DFNB93 | eukaryota>metazoa>chordata>vertebrata | Homo | sapiens | calcium-binding protein 2 isoform 3 [Homo sapiens] |
| XP_012351438.1 | 226 | CABP2 | eukaryota>metazoa>chordata>vertebrata | Nomascus | leucogenys | PREDICTED: calcium-binding protein 2 isoform X1 [Nomascus leucogenys] |
| XP_015289856.1 | 226 | CABP2 | eukaryota>metazoa>chordata>vertebrata | Macaca | fascicularis | PREDICTED: calcium-binding protein 2 isoform X1 [Macaca fascicularis] |
| NP_001147205.1 | 226 | LOC100280813 | eukaryota>viridiplantae | Zea | mays | calmodulin [Zea mays] |
| XP_001233509.2 | 224 | CABP1 | eukaryota>metazoa>chordata>vertebrata | Gallus | gallus | PREDICTED: calcium-binding protein 1 isoform X5 [Gallus gallus] |
| XP_016612824.1 | 224 | SPPG_00489 | eukaryota>fungi>chytridiomycota | Spizellomyces | punctatus DAOM BR117 | hypothetical protein SPPG_00489 [Spizellomyces punctatus DAOM BR117] |
| XP_019925473.1 | 223 | LOC105334726 | eukaryota>metazoa>mollusca | Crassostrea | gigas | PREDICTED: calmodulin-beta isoform X1 [Crassostrea gigas] |
| XP_019925474.1 | 223 | LOC105334726 | eukaryota>metazoa>mollusca | Crassostrea | gigas | PREDICTED: calmodulin isoform X2 [Crassostrea gigas] |
| XP_015141710.1 | 222 | CABP2 | eukaryota>metazoa>chordata>vertebrata | Gallus | gallus | PREDICTED: calcium-binding protein 2 isoform X2 [Gallus gallus] |
| NP_038906.2 | 221 | Cabp2 | eukaryota>metazoa>chordata>vertebrata | Mus | musculus | calcium-binding protein 2 isoform 1 [Mus musculus] |
| XP_001897398.1 | 221 | Bm1_29780 | eukaryota>metazoa>nematoda | Brugia | malayi | EF hand family protein [Brugia malayi] |
| NP_057450.2 | 220 | DFNB93 | eukaryota>metazoa>chordata>vertebrata | Homo | sapiens | calcium-binding protein 2 isoform 1 [Homo sapiens] |
| XP_003273991.1 | 220 | CABP2 | eukaryota>metazoa>chordata>vertebrata | Nomascus | leucogenys | PREDICTED: calcium-binding protein 2 isoform X2 [Nomascus leucogenys] |
| XP_005577112.1 | 220 | CABP2 | eukaryota>metazoa>chordata>vertebrata | Macaca | fascicularis | PREDICTED: calcium-binding protein 2 isoform X2 [Macaca fascicularis] |
| ELR18329.1 | 220 | ACA1_371900 | eukaryota>amoebozoa>acanthamoebidae | Acanthamoeba | castellanii str. Neff | EF hand domain containing protein [Acanthamoeba castellanii str. Neff] |
| XP_002285404.1 | 220 | LOC100245811 | eukaryota>viridiplantae | Vitis | vinifera | PREDICTED: calmodulin-like protein 3 [Vitis vinifera] |
| XP_016460667.1 | 219 | LOC107784110 | eukaryota>viridiplantae | Nicotiana | tabacum | PREDICTED: probable calcium-binding protein CML36 [Nicotiana tabacum] |
| XP_016462443.1 | 219 | LOC107785616 | eukaryota>viridiplantae | Nicotiana | tabacum | PREDICTED: probable calcium-binding protein CML36 [Nicotiana tabacum] |
| XP_006347296.1 | 219 | LOC102598823 | eukaryota>viridiplantae | Solanum | tuberosum | PREDICTED: calmodulin-like protein 3 [Solanum tuberosum] |
| XP_003554140.1 | 218 | LOC100815559 | eukaryota>viridiplantae | Glycine | max | PREDICTED: probable calcium-binding protein CML36 [Glycine max] |
| XP_017343770.1 | 217 | LOC108276532 | eukaryota>metazoa>chordata>vertebrata>actinopterygii | Ictalurus | punctatus | PREDICTED: calcium-binding protein 1-like isoform X2 [Ictalurus punctatus] |
| XP_011604896.1 | 216 | LOC101066197 | eukaryota>metazoa>chordata>vertebrata>actinopterygii | Takifugu | rubripes | PREDICTED: calcium-binding protein 2-like [Takifugu rubripes] |
| XP_017322763.1 | 216 | LOC108265140 | eukaryota>metazoa>chordata>vertebrata>actinopterygii | Ictalurus | punctatus | PREDICTED: calcium-binding protein 2-like isoform X2 [Ictalurus punctatus] |
| ERG84951.1 | 216 | ASU_04582 | eukaryota>metazoa>nematoda | Ascaris | suum | putative calcium-binding protein [Ascaris suum] |
| XP_003524630.2 | 216 | LOC100786354 | eukaryota>viridiplantae | Glycine | max | PREDICTED: calmodulin-like protein 1 [Glycine max] |
| NP_181672.1 | 216 | F13H10.4,F13H10_4 | eukaryota>viridiplantae | Arabidopsis | thaliana | Calcium-binding EF-hand family protein [Arabidopsis thaliana] |
| NP_001264540.1 | 215 | CABP2 | eukaryota>metazoa>chordata>vertebrata | Gallus | gallus | calcium-binding protein 2 [Gallus gallus] |
| NP_565996.1 | 215 | F14B2.33,multicopy,suppressors,of,snf4,deficiency,in,yeast,3 | eukaryota>viridiplantae | Arabidopsis | thaliana | Calcium-binding EF-hand family protein [Arabidopsis thaliana] |
| XP_003373971.1 | 215 | Tsp_08698 | eukaryota>metazoa>nematoda | Trichinella | spiralis | calmodulin [Trichinella spiralis] |
| XP_013426777.1 | 215 | M436DRAFT_82463 | eukaryota>fungi>ascomycota | Aureobasidium | namibiae CBS 147.97 | EF-hand [Aureobasidium namibiae CBS 147.97] |
| Caps1000024397 | 215 | Caps1000024397 | eukaryota>metazoa>annelida | Capitella | spI | >Caps1000024397 |
| XP_017316965.1 | 214 | LOC108260839 | eukaryota>metazoa>chordata>vertebrata>actinopterygii | Ictalurus | punctatus | PREDICTED: calcium-binding protein 2-like [Ictalurus punctatus] |
| XP_002266359.1 | 214 | LOC100254364 | eukaryota>viridiplantae | Vitis | vinifera | PREDICTED: calmodulin-like protein 3 [Vitis vinifera] |
| NP_001326523.1 | 214 | AtCAM7,calmodulin,7,CALMODULIN,7,Z-box,Binding,Factor,3,ZBF3 | eukaryota>viridiplantae | Arabidopsis | thaliana | calmodulin 7 [Arabidopsis thaliana] |
| KNE55575.1 | 213 | AMAG_01466 | eukaryota>fungi | Allomyces | macrogynus ATCC 38327 | calmodulin [Allomyces macrogynus ATCC 38327] |
| XP_003874723.1 | 213 | LMXM_36_3675 | eukaryota>euglenozoa>kinetoplastida | Leishmania | mexicana MHOM/GT/2001/U1103 | putative calmodulin [Leishmania mexicana MHOM/GT/2001/U1103] |
| XP_780862.3 | 213 | LOC575365 | eukaryota>metazoa>echinodermata | Strongylocentrotus | purpuratus | PREDICTED: calmodulin [Strongylocentrotus purpuratus] |
| XP_013400665.1 | 212 | LOC106166584 | eukaryota>metazoa | Lingula | anatina | PREDICTED: neo-calmodulin-like isoform X1 [Lingula anatina] |
| Smar1000001531 | 212 | Smar1000001531 | eukaryota>metazoa>arthropoda>myriapoda | Strigamia | maritima | >Smar1000001531 |
| Smed1000023664 | 212 | Smed1000023664 | eukaryota>metazoa>platyhelminthe | Schmidtea | mediterranea | >Smed1000023664 |
| XP_010663735.1 | 212 | LOC100262849 | eukaryota>viridiplantae | Vitis | vinifera | PREDICTED: putative calcium-binding protein CML19 [Vitis vinifera] |
| NP_001152651.1 | 211 | LOC100286292 | eukaryota>viridiplantae | Zea | mays | calmodulin [Zea mays] |
| XP_006351283.1 | 211 | LOC102603406 | eukaryota>viridiplantae | Solanum | tuberosum | PREDICTED: probable calcium-binding protein CML36 [Solanum tuberosum] |
| XP_011427010.1 | 210 | LOC105328000 | eukaryota>metazoa>mollusca | Crassostrea | gigas | PREDICTED: calmodulin-A isoform X1 [Crassostrea gigas] |
| XP_016493926.1 | 209 | LOC107813208 | eukaryota>viridiplantae | Nicotiana | tabacum | PREDICTED: probable calcium-binding protein CML36 [Nicotiana tabacum] |
| XP_016499615.1 | 209 | LOC107818184 | eukaryota>viridiplantae | Nicotiana | tabacum | PREDICTED: probable calcium-binding protein CML36 [Nicotiana tabacum] |
| XP_011455014.1 | 209 | LOC105347567 | eukaryota>metazoa>mollusca | Crassostrea | gigas | PREDICTED: neo-calmodulin [Crassostrea gigas] |
| NP_187630.1 | 209 | AT3G10190 | eukaryota>viridiplantae | Arabidopsis | thaliana | Calcium-binding EF-hand family protein [Arabidopsis thaliana] |
| NP_001255490.1 | 208 | cal-4 | eukaryota>metazoa>nematoda | Caenorhabditis | elegans | CALmodulin related genes [Caenorhabditis elegans] |
| Lhya1000005584 | 208 | Lhya1000005584 | eukaryota>fungi>mucoromycotina | Lichtheimia | hyalospora | >Lhya1000005584 |
| XP_808089.1 | 207 | Tc00.1047053507483.30 | eukaryota>euglenozoa>kinetoplastida | Trypanosoma | cruzi strain CL Brener | calmodulin [Trypanosoma cruzi strain CL Brener] |
| Caps1000022106 | 207 | Caps1000022106 | eukaryota>metazoa>annelida | Capitella | spI | >Caps1000022106 |
| XP_017318620.1 | 207 | cabp2 | eukaryota>metazoa>chordata>vertebrata>actinopterygii | Ictalurus | punctatus | PREDICTED: calcium-binding protein 2 isoform X4 [Ictalurus punctatus] |
| XP_018082625.1 | 207 | LOC108697221 | eukaryota>metazoa>chordata>vertebrata | Xenopus | laevis | PREDICTED: calcium-binding protein 2-like [Xenopus laevis] |
| NP_001256433.1 | 206 | E02A10.3 | eukaryota>metazoa>nematoda | Caenorhabditis | elegans | Uncharacterized protein CELE_E02A10.3 [Caenorhabditis elegans] |
| XP_011441728.2 | 206 | LOC105338356 | eukaryota>metazoa>mollusca | Crassostrea | gigas | PREDICTED: neo-calmodulin-like [Crassostrea gigas] |
| XP_012894039.1 | 206 | GSBLH_T00000385001 | eukaryota>stramenopiles | Blastocystis | hominis | uncharacterized protein [Blastocystis hominis] |
| XP_018117078.1 | 206 | LOC108716018 | eukaryota>metazoa>chordata>vertebrata | Xenopus | laevis | PREDICTED: calcium-binding protein 1 isoform X2 [Xenopus laevis] |
| XP_006351257.1 | 205 | LOC102595230 | eukaryota>viridiplantae | Solanum | tuberosum | PREDICTED: probable calcium-binding protein CML36 [Solanum tuberosum] |
| XP_003961445.1 | 205 | LOC101077996 | eukaryota>metazoa>chordata>vertebrata>actinopterygii | Takifugu | rubripes | PREDICTED: calcium-binding protein 5-like [Takifugu rubripes] |
| NP_190646.1 | 205 | calmodulin-like,41 | eukaryota>viridiplantae | Arabidopsis | thaliana | calmodulin-like 41 [Arabidopsis thaliana] |
| XP_010645228.1 | 204 | LOC100267033 | eukaryota>viridiplantae | Vitis | vinifera | PREDICTED: calmodulin-like protein 3 [Vitis vinifera] |
| XP_011427011.1 | 204 | LOC105328000 | eukaryota>metazoa>mollusca | Crassostrea | gigas | PREDICTED: calmodulin-A isoform X2 [Crassostrea gigas] |
| XP_002279679.1 | 204 | LOC100267822 | eukaryota>viridiplantae | Vitis | vinifera | PREDICTED: probable calcium-binding protein CML23 [Vitis vinifera] |
| XP_005244786.2 | 203 | CAGLP | eukaryota>metazoa>chordata>vertebrata | Homo | sapiens | PREDICTED: calmodulin-like protein 6 isoform X1 [Homo sapiens] |
| NP_001153725.1 | 203 | Cabp2 | eukaryota>metazoa>chordata>vertebrata | Mus | musculus | calcium-binding protein 2 isoform 3 [Mus musculus] |
| XP_006531832.1 | 203 | Cabp2 | eukaryota>metazoa>chordata>vertebrata | Mus | musculus | PREDICTED: calcium-binding protein 2 isoform X5 [Mus musculus] |
| NP_001168222.1 | 203 | GRMZM2G031329 | eukaryota>viridiplantae | Zea | mays | uncharacterized protein LOC100381981 [Zea mays] |
| ETO17697.1 | 203 | RFI_19618 | eukaryota>rhizaria | Reticulomyxa | filosa | calmodulin [Reticulomyxa filosa] |
| XP_014630165.1 | 203 | LOC100813924 | eukaryota>viridiplantae | Glycine | max | PREDICTED: probable calcium-binding protein CML15 [Glycine max] |
| XP_015768205.1 | 202 | LOC107346864 | eukaryota>metazoa>cnidaria | Acropora | digitifera | PREDICTED: calmodulin-like [Acropora digitifera] |
| XP_015768206.1 | 202 | LOC107346865 | eukaryota>metazoa>cnidaria | Acropora | digitifera | PREDICTED: calmodulin-like [Acropora digitifera] |
| NP_001256434.1 | 202 | E02A10.3 | eukaryota>metazoa>nematoda | Caenorhabditis | elegans | Uncharacterized protein CELE_E02A10.3 [Caenorhabditis elegans] |
| NP_001256435.1 | 202 | E02A10.3 | eukaryota>metazoa>nematoda | Caenorhabditis | elegans | Uncharacterized protein CELE_E02A10.3 [Caenorhabditis elegans] |
| XP_016875728.1 | 202 | CALBRAIN,HCALB_BR | [illegible] | [illegible] | sapiens | PREDICTED: calcium-binding protein 1 isoform X5 [Homo sapiens] |
| XP_014771763.1 | 201 | LOC106870257 | [illegible] | [illegible] | [illegible] | [illegible] |
| NP_001150089.1 | 200 | GRMZM2G349655 | [illegible] | [illegible] | [illegible] | [illegible] |
| XP_014149168.1 | 200 | SARC_12204 | [illegible] | [illegible] | [illegible] | [illegible] |
| NP_001318399.1 | 199 | ARABIDOPSIS,THALIANA,CALMODULIN,LIKE,4,ATCAL4,calmodulin-like[illegible] | [illegible] | [illegible] | [illegible] | [illegible] |
| NP_001150915.1 | 199 | GRMZM2G097900,si606063b10 | [illegible] | [illegible] | [illegible] | [illegible] |
| OQV20899.1 | 199 | BV898_05239 | [illegible] | [illegible] | [illegible] | [illegible] |
| EPB82295.1 | 198 | HMPREF1544_10975 | [illegible] | [illegible] | [illegible] | [illegible] |
| XP_002590348.1 | 198 | BRAFLDRAFT_76605 | [illegible] | [illegible] | [illegible] | [illegible] |
| XP_016483638.1 | 197 | LOC107804288 | [illegible] | [illegible] | [illegible] | [illegible] |
| NP_001292553.1 | 197 | caM,CAMII,LQT15,PHKD,PHKD2 | [illegible] | [illegible] | [illegible] | [illegible] |
| NP_001300863.1 | 197 | AI256814,AI327027,AI461935,AL024000,Calm,CaM,Cam1 | [illegible] | [illegible] | [illegible] | [illegible] |
| XP_003061114.1 | 197 | MICPUCDRAFT_34833 | [illegible] | [illegible] | [illegible] | [illegible] |
| XP_013400674.1 | 197 | LOC106166584 | [illegible] | [illegible] | [illegible] | [illegible] |
| XP_013760455.1 | 197 | AMSG_03124 | [illegible] | [illegible] | [illegible] | [illegible] |
| NP_219501.2 | 196 | NY-BR-20 | [illegible] | [illegible] | [illegible] | [illegible] |
| XP_003267191.1 | 196 | CALML4 | [illegible] | [illegible] | [illegible] | [illegible] |
| XP_006531833.1 | 196 | Cabp2 | [illegible] | [illegible] | [illegible] | [illegible] |
| XP_011613778.1 | 196 | cabp1 | [illegible] | [illegible] | [illegible] | [illegible] |
| XP_005559954.1 | 195 | CALML4 | [illegible] | [illegible] | [illegible] | [illegible] |
| ETO18056.1 | 195 | RFI_19237 | [illegible] | [illegible] | [illegible] | [illegible] |
| NP_191503.1 | 195 | AT3G59440 | [illegible] | [illegible] | [illegible] | [illegible] |
| XP_008673745.1 | 195 | LOC103649813 | [illegible] | [illegible] | [illegible] | [illegible] |
| ETO16277.1 | 194 | RFI_21077 | [illegible] | [illegible] | [illegible] | [illegible] |
| Caps1000006906 | 194 | Caps1000006906 | [illegible] | [illegible] | [illegible] | [illegible] |
| Lhya1000002576 | 194 | Lhya1000002576 | [illegible] | [illegible] | [illegible] | [illegible] |
| Lhya1000005583 | 194 | Lhya1000005583 | [illegible] | [illegible] | [illegible] | [illegible] |
| XP_006365377.1 | 194 | LOC102601367 | [illegible] | [illegible] | [illegible] | [illegible] |
| Bcir1000004971 | 193 | Bcir1000004971 | [illegible] | [illegible] | [illegible] | [illegible] |
| XP_015170234.1 | 193 | LOC102591062 | [illegible] | [illegible] | [illegible] | [illegible] |
| XP_009020056.1 | 193 | HELRODRAFT_161938 | [illegible] | [illegible] | [illegible] | [illegible] |
| NP_524902.2 | 193 | 63F-1,Ca[2+],binding,protein,CBP12,CG15855,Dmel\CG15855,E63-[illegible] | [illegible] | [illegible] | [illegible] | [illegible] |
| Caps1000002977 | 192 | Caps1000002977 | [illegible] | [illegible] | [illegible] | [illegible] |
| XP_006339442.1 | 192 | LOC102582450 | [illegible] | [illegible] | [illegible] | [illegible] |
| XP_016511236.1 | 191 | LOC107828446 | [illegible] | [illegible] | [illegible] | [illegible] |
| NP_181642.1 | 191 | T3K9.14,T3K9_14 | [illegible] | [illegible] | [illegible] | [illegible] |
| NP_193810.1 | 191 | calmodulin,like,42,F21C20.130,F21C20_130 | [illegible] | [illegible] | [illegible] | [illegible] |
| XP_014787760.1 | 191 | LOC106881770 | [illegible] | [illegible] | [illegible] | [illegible] |
| XP_808994.1 | 191 | Tc00.1047053508951.50 | [illegible] | [illegible] | [illegible] | [illegible] |
| XP_016482875.1 | 190 | LOC107803626 | [illegible] | [illegible] | [illegible] | [illegible] |
| XP_003375537.1 | 190 | Tsp_08523 | [illegible] | [illegible] | [illegible] | [illegible] |
| XP_003550069.1 | 190 | LOC100802167 | [illegible] | [illegible] | [illegible] | [illegible] |
| XP_011441748.1 | 190 | LOC105338363 | [illegible] | [illegible] | [illegible] | [illegible] |
| XP_013779887.1 | 190 | LOC106464303 | [illegible] | [illegible] | [illegible] | [illegible] |
| XP_018286143.1 | 190 | PHYBLDRAFT_173596 | [illegible] | [illegible] | [illegible] | [illegible] |
| Uram1000006768 | 190 | Uram1000006768 | [illegible] | [illegible] | [illegible] | [illegible] |
| EPB87027.1 | 189 | HMPREF1544_06144 | [illegible] | [illegible] | [illegible] | [illegible] |
| Bcir1000000008 | 189 | Bcir1000000008 | [illegible] | [illegible] | [illegible] | [illegible] |
| XP_016487304.1 | 189 | LOC107807439 | [illegible] | [illegible] | [illegible] | [illegible] |
| EIE91510.1 | 188 | RO3G_16221 | [illegible] | [illegible] | [illegible] | [illegible] |
| NP_001141189.1 | 188 | GRMZM2G309327 | [illegible] | [illegible] | [illegible] | [illegible] |
| NP_001168304.1 | 188 | GRMZM2G312661 | [illegible] | [illegible] | [illegible] | [illegible] |
| XP_004932140.1 | 188 | LOC101738501 | [illegible] | [illegible] | [illegible] | [illegible] |
| XP_004932141.1 | 188 | LOC101738501 | [illegible] | [illegible] | [illegible] | [illegible] |
| XP_015299308.1 | 188 | CALML6 | [illegible] | [illegible] | [illegible] | [illegible] |
| ETO13499.1 | 188 | RFI_23867 | [illegible] | [illegible] | [illegible] | [illegible] |
| KNE58371.1 | 188 | AMAG_03954 | [illegible] | [illegible] | [illegible] | [illegible] |
| XP_006815692.1 | 188 | LOC100371565 | [illegible] | [illegible] | [illegible] | [illegible] |
| XP_014770346.1 | 188 | LOC106869213 | [illegible] | [illegible] | [illegible] | [illegible] |
| XP_017313791.1 | 188 | calml4 | [illegible] | [illegible] | [illegible] | [illegible] |
| XP_018736275.1 | 188 | CMD1 | [illegible] | [illegible] | [illegible] | [illegible] |
| XP_017340988.1 | 188 | LOC108274992 | [illegible] | [illegible] | [illegible] | [illegible] |
| XP_003524362.1 | 188 | LOC100813092 | [illegible] | [illegible] | [illegible] | [illegible] |
| NP_001241462.1 | 187 | LOC100790495 | [illegible] | [illegible] | [illegible] | [illegible] |
| XP_003537323.1 | 187 | LOC100787013 | [illegible] | [illegible] | [illegible] | [illegible] |
| NP_179170.1 | 187 | AtCML30,calmodulin-like,30 | [illegible] | [illegible] | [illegible] | [illegible] |
| XP_002611450.1 | 187 | BRAFLDRAFT_63918 | [illegible] | [illegible] | [illegible] | [illegible] |
| XP_013400682.1 | 187 | LOC106166584 | [illegible] | [illegible] | [illegible] | [illegible] |
| ORE07332.1 | 187 | BCV72DRAFT_110282 | [illegible] | [illegible] | [illegible] | [illegible] |
| EJY65763.1 | 186 | OXYTRI_14079 | [illegible] | [illegible] | [illegible] | [illegible] |
| XP_009496274.1 | 186 | H696_04116 | [illegible] | [illegible] | [illegible] | [illegible] |
| XP_816466.1 | 186 | Tc00.1047053510121.50 | [illegible] | [illegible] | [illegible] | [illegible] |
| NP_001005962.1 | 186 | cabp1,zgc:101878 | [illegible] | [illegible] | [illegible] | [illegible] |
| NP_724103.1 | 186 | BcDNA:AT22559,CG15158,Dmel\CG31802 | [illegible] | [illegible] | [illegible] | [illegible] |
| NP_173866.1 | 186 | F21J9.28 | [illegible] | [illegible] | [illegible] | [illegible] |
| XP_016503215.1 | 186 | LOC107821294 | [illegible] | [illegible] | [illegible] | [illegible] |
| XP_006346428.1 | 185 | LOC102596749 | [illegible] | [illegible] | [illegible] | [illegible] |
| XP_016468454.1 | 185 | LOC107790990 | [illegible] | [illegible] | [illegible] | [illegible] |
| XP_016470715.1 | 185 | LOC107792965 | [illegible] | [illegible] | [illegible] | [illegible] |
| NP_001236739.1 | 185 | LOC100527487 | [illegible] | [illegible] | [illegible] | [illegible] |
| NP_001235680.1 | 185 | LOC100499763 | [illegible] | [illegible] | [illegible] | [illegible] |
| NP_001276189.1 | 185 | LOC100803158 | [illegible] | [illegible] | [illegible] | [illegible] |
| XP_016507843.1 | 185 | LOC107825494 | [illegible] | [illegible] | [illegible] | [illegible] |
| Caps1000006465 | 185 | Caps1000006465 | [illegible] | [illegible] | [illegible] | [illegible] |
| NP_001041263.2 | 184 | K03A1.4 | [illegible] | [illegible] | [illegible] | [illegible] |
| NP_001183849.1 | 183 | LOC100502442 | [illegible] | [illegible] | [illegible] | [illegible] |
| XP_015299310.1 | 183 | CALML6 | [illegible] | [illegible] | [illegible] | [illegible] |
| XP_003533646.1 | 183 | LOC100807117 | [illegible] | [illegible] | [illegible] | [illegible] |
| XP_011608260.1 | 183 | calml4 | [illegible] | [illegible] | [illegible] | [illegible] |
| XP_016471425.1 | 183 | LOC107793563 | [illegible] | [illegible] | [illegible] | [illegible] |
| XP_018655557.1 | 183 | Smp_026560.1 | [illegible] | [illegible] | [illegible] | [illegible] |
| XP_013774956.1 | 183 | LOC106459837 | [illegible] | [illegible] | [illegible] | [illegible] |
| NP_001255491.1 | 182 | cal-4 | [illegible] | [illegible] | [illegible] | [illegible] |
| XP_012554753.1 | 182 | LOC100212129 | [illegible] | [illegible] | [illegible] | [illegible] |
| CAG07602.1 | 182 | GSTEN:00028321:G:001 | [illegible] | [illegible] | [illegible] | [illegible] |
| XP_001785090.1 | 182 | PHYPADRAFT_154846 | [illegible] | [illegible] | [illegible] | [illegible] |
| XP_002285223.1 | 182 | LOC100257203 | [illegible] | [illegible] | [illegible] | [illegible] |
| XP_006583713.1 | 182 | LOC100790109 | [illegible] | [illegible] | [illegible] | [illegible] |
| NP_850097.1 | 181 | 1357,ACAM-2,calmodulin,2,calmodulin,5,CAM2,T20P8.8 | [illegible] | [illegible] | [illegible] | [illegible] |
| XP_016455012.1 | 181 | LOC107779166 | [illegible] | [illegible] | [illegible] | [illegible] |
| XP_016493990.1 | 181 | LOC107813260 | [illegible] | [illegible] | [illegible] | [illegible] |
| NP_619650.2 | 181 | CAGLP | [illegible] | [illegible] | [illegible] | [illegible] |
| XP_001947193.2 | 181 | LOC100161899 | [illegible] | [illegible] | [illegible] | [illegible] |
| XP_008182669.1 | 181 | LOC100161899 | [illegible] | [illegible] | [illegible] | [illegible] |
| XP_012553386.1 | 181 | LOC101741854 | [illegible] | [illegible] | [illegible] | [illegible] |
| XP_012553387.1 | 181 | LOC101741854 | [illegible] | [illegible] | [illegible] | [illegible] |
| CDS36411.1 | 181 | EmuJ_000350400 | [illegible] | [illegible] | [illegible] | [illegible] |
| EJY69755.1 | 181 | OXYTRI_09506 | [illegible] | [illegible] | [illegible] | [illegible] |
| NP_199259.1 | 181 | calmodulin,like,43,MFC16.12,MFC16_12 | [illegible] | [illegible] | [illegible] | [illegible] |
| XP_011446785.1 | 181 | LOC105341779 | [illegible] | [illegible] | [illegible] | [illegible] |
| XP_013401701.1 | 181 | LOC106167460 | [illegible] | [illegible] | [illegible] | [illegible] |
| XP_016448278.1 | 181 | LOC107773369 | [illegible] | [illegible] | [illegible] | [illegible] |
| XP_018293892.1 | 181 | PHYBLDRAFT_180432 | [illegible] | [illegible] | [illegible] | [illegible] |
| NP_001189046.1 | 181 | 63F-1,Ca[2+],binding,protein,CBP12,CG15855,Dmel\CG15855,E63-[illegible] | [illegible] | [illegible] | [illegible] | [illegible] |
| XP_002974189.1 | 181 | SELMODRAFT_100711 | [illegible] | [illegible] | [illegible] | [illegible] |
| XP_003632230.1 | 180 | LOC100261610 | [illegible] | [illegible] | [illegible] | [illegible] |
| XP_006340365.2 | 180 | LOC102594384 | [illegible] | [illegible] | [illegible] | [illegible] |
| XP_016455013.1 | 180 | LOC107779166 | [illegible] | [illegible] | [illegible] | [illegible] |
| XP_016493991.1 | 180 | LOC107813260 | [illegible] | [illegible] | [illegible] | [illegible] |
| NP_001237156.1 | 180 | LOC100500636 | [illegible] | [illegible] | [illegible] | [illegible] |
| XP_003530955.1 | 180 | LOC100783572 | [illegible] | [illegible] | [illegible] | [illegible] |
| XP_014634193.1 | 180 | LOC100783572 | [illegible] | [illegible] | [illegible] | [illegible] |
| NP_001151914.1 | 180 | GRMZM2G146720 | [illegible] | [illegible] | [illegible] | [illegible] |
| XP_008668118.1 | 180 | LOC100285141 | [illegible] | [illegible] | [illegible] | [illegible] |
| XP_002972882.1 | 180 | SELMODRAFT_270984 | [illegible] | [illegible] | [illegible] | [illegible] |
| XP_011150177.1 | 180 | LOC105189634 | [illegible] | [illegible] | [illegible] | [illegible] |
| KXJ21616.1 | 180 | AC249_AIPGENE18012 | [illegible] | [illegible] | [illegible] | [illegible] |
| NP_001256428.1 | 180 | cal-1 | [illegible] | [illegible] | [illegible] | [illegible] |
| XP_016450675.1 | 179 | LOC107775463 | [illegible] | [illegible] | [illegible] | [illegible] |
| XP_009167655.1 | 179 | T265_04609 | [illegible] | [illegible] | [illegible] | [illegible] |
| XP_009172902.1 | 179 | T265_08749 | [illegible] | [illegible] | [illegible] | [illegible] |
| XP_016466291.1 | 179 | LOC107789034 | [illegible] | [illegible] | [illegible] | [illegible] |
| XP_016488038.1 | 179 | LOC107808069 | [illegible] | [illegible] | [illegible] | [illegible] |
| XP_016771319.1 | 179 | LOC552209 | [illegible] | [illegible] | [illegible] | [illegible] |
| XP_018095703.1 | 179 | LOC108703912 | [illegible] | [illegible] | [illegible] | [illegible] |
| XP_003528969.1 | 179 | LOC100815264 | [illegible] | [illegible] | [illegible] | [illegible] |
| XP_011414765.1 | 178 | LOC105319062 | [illegible] | [illegible] | [illegible] | [illegible] |
| XP_011415115.1 | 177 | LOC105319316 | [illegible] | [illegible] | [illegible] | [illegible] |
| XP_011415116.1 | 177 | LOC105319316 | [illegible] | [illegible] | [illegible] | [illegible] |
| XP_011415118.1 | 177 | LOC105319316 | [illegible] | [illegible] | [illegible] | [illegible] |
| XP_011415119.1 | 177 | LOC105319316 | [illegible] | [illegible] | [illegible] | [illegible] |
| XP_015838421.1 | 177 | LOC659938 | [illegible] | [illegible] | [illegible] | [illegible] |
| ERG85040.1 | 177 | ASU_04393 | [illegible] | [illegible] | [illegible] | [illegible] |
| XP_003084116.1 | 177 | Ot18g00850 | [illegible] | [illegible] | [illegible] | [illegible] |
| XP_004206928.1 | 177 | LOC100212129 | [illegible] | [illegible] | [illegible] | [illegible] |
| XP_008180058.1 | 177 | LOC100162151 | [illegible] | [illegible] | [illegible] | [illegible] |
| XP_009059542.1 | 177 | LOTGIDRAFT_192654 | [illegible] | [illegible] | [illegible] | [illegible] |
| XP_002988053.1 | 176 | SELMODRAFT_159200 | [illegible] | [illegible] | [illegible] | [illegible] |
| CDW77243.1 | 176 | Contig4287.g4591 | [illegible] | [illegible] | [illegible] | [illegible] |
| EFA85881.1 | 176 | calB | [illegible] | [illegible] | [illegible] | [illegible] |
| XP_006819033.1 | 176 | LOC102806008 | [illegible] | [illegible] | [illegible] | [illegible] |
| XP_014778343.1 | 176 | LOC106874943 | [illegible] | [illegible] | [illegible] | [illegible] |
| XP_001762647.1 | 176 | PHYPADRAFT_75995 | [illegible] | [illegible] | [illegible] | [illegible] |
| XP_001768532.1 | 176 | PHYPADRAFT_187280 | [illegible] | [illegible] | [illegible] | [illegible] |
| NP_001154755.1 | 175 | ACAM-1,calmodulin,1,K22F20.20,K22F20_20,TCH1,TOUCH,1 | [illegible] | [illegible] | [illegible] | [illegible] |
| XP_011414766.1 | 175 | LOC105319062 | [illegible] | [illegible] | [illegible] | [illegible] |
| XP_017334567.1 | 175 | calml6 | [illegible] | [illegible] | [illegible] | [illegible] |
| XP_001779680.1 | 175 | PHYPADRAFT_147571 | [illegible] | [illegible] | [illegible] | [illegible] |
| OQV19740.1 | 175 | BV898_06279 | [illegible] | [illegible] | [illegible] | [illegible] |
| CAG10181.1 | 174 | GSTEN:00031732:G:001 | [illegible] | [illegible] | [illegible] | [illegible] |
| XP_002165237.1 | 174 | LOC100212129 | [illegible] | [illegible] | [illegible] | [illegible] |
| Lhya1000006016 | 174 | Lhya1000006016 | [illegible] | [illegible] | [illegible] | [illegible] |
| XP_006367611.1 | 174 | LOC102603618 | [illegible] | [illegible] | [illegible] | [illegible] |
| XP_008680817.1 | 174 | LOC103655921 | [illegible] | [illegible] | [illegible] | [illegible] |
| OZJ02928.1 | 174 | BZG36_04121 | [illegible] | [illegible] | [illegible] | [illegible] |
| NP_188933.1 | 173 | AtCML11,calmodulin-like,11 | [illegible] | [illegible] | [illegible] | [illegible] |
| XP_005832678.1 | 173 | GUITHDRAFT_108572 | [illegible] | [illegible] | [illegible] | [illegible] |
| XP_007331737.1 | 173 | AGABI1DRAFT_77172 | [illegible] | [illegible] | [illegible] | [illegible] |
| XP_013762720.1 | 173 | AMSG_00442 | [illegible] | [illegible] | [illegible] | [illegible] |
| XP_001768907.1 | 173 | PHYPADRAFT_133902 | [illegible] | [illegible] | [illegible] | [illegible] |
| OZJ01474.1 | 173 | BZG36_05497 | [illegible] | [illegible] | [illegible] | [illegible] |
| NP_001147004.1 | 172 | LOC100280614 | [illegible] | [illegible] | [illegible] | [illegible] |
| XP_001622774.1 | 172 | NEMVEDRAFT_v1g220271 | [illegible] | [illegible] | [illegible] | [illegible] |
| XP_011399134.1 | 172 | F751_4731 | [illegible] | [illegible] | [illegible] | [illegible] |
| XP_013400688.1 | 172 | LOC106166584 | [illegible] | [illegible] | [illegible] | [illegible] |
| NP_495906.2 | 171 | cal-2 | [illegible] | [illegible] | [illegible] | [illegible] |
| XP_012554754.1 | 171 | LOC100212129 | [illegible] | [illegible] | [illegible] | [illegible] |
| XP_012759347.1 | 171 | SAMD00019534_004600 | [illegible] | [illegible] | [illegible] | [illegible] |
| XP_013400697.1 | 171 | LOC106166584 | [illegible] | [illegible] | [illegible] | [illegible] |
| NP_001287824.1 | 170 | CRSD,CSNB2B | [illegible] | [illegible] | [illegible] | [illegible] |
| NP_001287825.1 | 170 | CRSD,CSNB2B | [illegible] | [illegible] | [illegible] | [illegible] |
| XP_011543485.1 | 170 | CRSD,CSNB2B | [illegible] | [illegible] | [illegible] | [illegible] |
| XP_016873514.1 | 170 | CRSD,CSNB2B | [illegible] | [illegible] | [illegible] | [illegible] |
| NP_173259.1 | 170 | T10F20.22 | [illegible] | [illegible] | [illegible] | [illegible] |
| NP_849686.1 | 170 | T10F20.22 | [illegible] | [illegible] | [illegible] | [illegible] |
| XP_011449950.1 | 170 | LOC105344068 | [illegible] | [illegible] | [illegible] | [illegible] |
| XP_011449951.1 | 170 | LOC105344068 | [illegible] | [illegible] | [illegible] | [illegible] |
| KXJ14149.1 | 170 | AC249_AIPGENE7476 | [illegible] | [illegible] | [illegible] | [illegible] |

| | | | | | | |
|---|---|---|---|---|---|---|
| NP_001231973.1 | 170 | LOC494376 | eukaryota>metazoa>chordata | Ciona | intestinalis | [illegible] |
| XP_005843158.1 | 170 | CHLNCDRAFT_55421 | eukaryota>viridiplantae>chlorophyta | Chlorella | variabilis | [illegible] |
| XP_013236837.1 | 170 | DI09_71p40 | eukaryota>fungi>microsporidia | Mitosporidium | daphniae | [illegible] |
| XP_789104.4 | 170 | LOC584132 | eukaryota>metazoa>echinodermata | Strongylocentrotus | purpuratus | [illegible] |
| Caps1000019672 | 170 | Caps1000019672 | eukaryota>metazoa>annelida | Capitella | spI | [illegible] |
| XP_017322402.1 | 170 | cabp1 | eukaryota>metazoa>chordata>vertebrata>actinopterygii | Ictalurus | punctatus | [illegible] |
| NP_001146794.1 | 169 | calmodulin,GRMZM2G444621 | eukaryota>viridiplantae | Zea | mays | [illegible] |
| XP_002946868.1 | 169 | fap272 | eukaryota>viridiplantae>chlorophyta | Volvox | carteri f. nagariensis | [illegible] |
| XP_003543563.3 | 169 | LOC100813656 | eukaryota>viridiplantae | Glycine | max | [illegible] |
| XP_005786038.1 | 169 | CAM3 | eukaryota>haptophyceae | Emiliania | huxleyi CCMP1516 | [illegible] |
| XP_006678916.1 | 169 | BATDEDRAFT_19649 | eukaryota>fungi>chytridiomycota | Batrachochytrium | dendrobatidis JAM81 | [illegible] |
| XP_008648304.1 | 169 | ZMCALM1 | eukaryota>viridiplantae | Zea | mays | [illegible] |
| XP_014771770.1 | 169 | LOC106870257 | eukaryota>metazoa>mollusca | Octopus | bimaculoides | [illegible] |
| NP_001018568.1 | 169 | zgc:109977 | eukaryota>metazoa>chordata>vertebrata>actinopterygii | Danio | rerio | [illegible] |
| XP_001752070.1 | 169 | PHYPADRAFT_111790 | eukaryota>viridiplantae | Physcomitrella | patens | [illegible] |
| XP_006345065.1 | 169 | LOC102599351 | eukaryota>viridiplantae | Solanum | tuberosum | [illegible] |
| XP_011441749.1 | 168 | LOC105338364 | eukaryota>metazoa>mollusca | Crassostrea | gigas | [illegible] |
| XP_019927029.1 | 168 | LOC105338364 | eukaryota>metazoa>mollusca | Crassostrea | gigas | [illegible] |
| XP_019927030.1 | 168 | LOC105338364 | eukaryota>metazoa>mollusca | Crassostrea | gigas | [illegible] |
| ERG83959.1 | 168 | ASU_06137 | eukaryota>metazoa>nematoda | Ascaris | suum | [illegible] |
| GAQ86360.1 | 168 | KFL_002850040 | eukaryota>viridiplantae | Klebsormidium | nitens | [illegible] |
| XP_013400705.1 | 168 | LOC106166584 | eukaryota>metazoa | Lingula | anatina | [illegible] |
| XP_015755705.1 | 168 | LOC107335225 | eukaryota>metazoa>cnidaria | Acropora | digitifera | [illegible] |
| NP_001153724.1 | 168 | Cabp2 | eukaryota>metazoa>chordata>vertebrata | Mus | musculus | [illegible] |
| XP_003964546.1 | 168 | LOC101075297 | eukaryota>metazoa>chordata>vertebrata>actinopterygii | Takifugu | rubripes | [illegible] |
| XP_005589779.1 | 168 | CABP5 | eukaryota>metazoa>chordata>vertebrata | Macaca | fascicularis | [illegible] |
| XP_011617066.1 | 168 | cabp4 | eukaryota>metazoa>chordata>vertebrata>actinopterygii | Takifugu | rubripes | [illegible] |
| XP_002277499.1 | 168 | LOC100247858 | eukaryota>viridiplantae | Vitis | vinifera | [illegible] |
| NP_001002414.1 | 167 | Cabp1,zgc:92680 | eukaryota>metazoa>chordata>vertebrata>actinopterygii | Danio | rerio | [illegible] |
| NP_001297644.1 | 167 | caldendrin | eukaryota>metazoa>chordata>vertebrata | Mus | musculus | [illegible] |
| NP_004267.2 | 167 | CALBRAIN,HCALB_BR | eukaryota>metazoa>chordata>vertebrata | Homo | sapiens | [illegible] |
| XP_003280335.1 | 167 | CABP1 | eukaryota>metazoa>chordata>vertebrata | Nomascus | leucogenys | [illegible] |
| XP_004934447.1 | 167 | CABP1 | eukaryota>metazoa>chordata>vertebrata | Gallus | gallus | [illegible] |
| XP_017343771.1 | 167 | LOC108276532 | eukaryota>metazoa>chordata>vertebrata>actinopterygii | Ictalurus | punctatus | [illegible] |
| XP_001901898.1 | 167 | Bm1_52150 | eukaryota>metazoa>nematoda | Brugia | malayi | [illegible] |
| ETO09562.1 | 167 | RFI_27815 | eukaryota>rhizaria | Reticulomyxa | filosa | [illegible] |
| XP_001849785.1 | 167 | CpipJ_CPIJ007602 | eukaryota>metazoa>hexapoda | Culex | quinquefasciatus | [illegible] |
| XP_003726780.1 | 167 | LOC100891445 | eukaryota>metazoa>echinodermata | Strongylocentrotus | purpuratus | [illegible] |
| XP_011416018.1 | 167 | LOC105319972 | eukaryota>metazoa>mollusca | Crassostrea | gigas | [illegible] |
| XP_016657505.1 | 167 | LOC100162151 | eukaryota>metazoa>hexapoda | Acyrthosiphon | pisum | [illegible] |
| Caps1000024396 | 167 | Caps1000024396 | eukaryota>metazoa>annelida | Capitella | spI | [illegible] |
| XP_015160192.1 | 167 | LOC107058689 | eukaryota>viridiplantae | Solanum | tuberosum | [illegible] |
| XP_011603190.1 | 166 | LOC101065517 | eukaryota>metazoa>chordata>vertebrata>actinopterygii | Takifugu | rubripes | [illegible] |
| NP_174504.1 | 166 | F27G20.1 | eukaryota>viridiplantae | Arabidopsis | thaliana | [illegible] |
| ERG83873.1 | 166 | ASU_06315 | eukaryota>metazoa>nematoda | Ascaris | suum | [illegible] |
| KXJ22231.1 | 166 | calm | eukaryota>metazoa>cnidaria | Exaiptasia | pallida | [illegible] |
| XP_009295966.1 | 166 | calml4b | eukaryota>metazoa>chordata>vertebrata>actinopterygii | Danio | rerio | [illegible] |
| XP_015761991.1 | 166 | LOC107341081 | eukaryota>metazoa>cnidaria | Acropora | digitifera | [illegible] |
| CAF91408.1 | 165 | GSTEN:00006151:G:001 | eukaryota>metazoa>chordata>vertebrata>actinopterygii | Tetraodon | nigroviridis | [illegible] |
| NP_956992.1 | 165 | zgc:73060 | eukaryota>metazoa>chordata>vertebrata>actinopterygii | Danio | rerio | [illegible] |
| XP_005156057.1 | 165 | zgc:73060 | eukaryota>metazoa>chordata>vertebrata>actinopterygii | Danio | rerio | [illegible] |
| OAF65977.1 | 165 | A3Q56_06298 | eukaryota>metazoa | Intoshia | linei | [illegible] |
| XP_002733157.1 | 165 | LOC100377516 | eukaryota>metazoa>hemichordata | Saccoglossus | kowalevskii | [illegible] |
| XP_002946692.1 | 165 | camA | eukaryota>viridiplantae>chlorophyta | Volvox | carteri f. nagariensis | [illegible] |
| XP_009050133.1 | 165 | LOTGIDRAFT_142024 | eukaryota>metazoa>mollusca | Lottia | gigantea | [illegible] |
| XP_009171875.1 | 165 | T265_07946 | eukaryota>metazoa | Opisthorchis | viverrini | [illegible] |
| XP_969500.1 | 165 | LOC657988 | eukaryota>metazoa>hexapoda | Tribolium | castaneum | [illegible] |
| NP_186950.1 | 165 | F13E7.5,F13E7_5 | eukaryota>viridiplantae | Arabidopsis | thaliana | [illegible] |
| XP_001765534.1 | 165 | PHYPADRAFT_129576 | eukaryota>viridiplantae | Physcomitrella | patens | [illegible] |
| NP_001154754.1 | 164 | ACAM-1,calmodulin,1,K22F20.20,K22F20_20,TCH1,TOUCH,1 | eukaryota>viridiplantae | Arabidopsis | thaliana | [illegible] |
| XP_003279757.1 | 164 | CALML6 | eukaryota>metazoa>chordata>vertebrata | Nomascus | leucogenys | [illegible] |
| XP_011539143.1 | 164 | CAGLP | eukaryota>metazoa>chordata>vertebrata | Homo | sapiens | [illegible] |
| XP_006341535.1 | 164 | LOC102599328 | eukaryota>viridiplantae | Solanum | tuberosum | [illegible] |
| XP_016468045.1 | 164 | LOC107790609 | eukaryota>viridiplantae | Nicotiana | tabacum | [illegible] |

[illegible]

| | | | | | | |
|---|---|---|---|---|---|---|
| XP_002779137.1 | 149 | Pmar_PMAR021412 | eukaryota>alveolata | Perkinsus | marinus ATCC 50983 | calmodulin, putative [Perkinsus marinus ATCC 50983] |
| XP_002785606.1 | 149 | Pmar_PMAR022352 | eukaryota>alveolata | Perkinsus | marinus ATCC 50983 | calmodulin, putative [Perkinsus marinus ATCC 50983] |
| XP_002963446.1 | 149 | SELMODRAFT_141966 | eukaryota>viridiplantae | Selaginella | moellendorffii | hypothetical protein SELMODRAFT_141966 [Selaginella moellendorffii] |
| XP_002971581.1 | 149 | SELMODRAFT_172372 | eukaryota>viridiplantae | Selaginella | moellendorffii | hypothetical protein SELMODRAFT_172372 [Selaginella moellendorffii] |
| XP_002977832.1 | 149 | SELMODRAFT_107557 | eukaryota>viridiplantae | Selaginella | moellendorffii | hypothetical protein SELMODRAFT_107557 [Selaginella moellendorffii] |
| XP_002979493.1 | 149 | SELMODRAFT_233374 | eukaryota>viridiplantae | Selaginella | moellendorffii | hypothetical protein SELMODRAFT_233374 [Selaginella moellendorffii] |
| XP_002998616.1 | 149 | PITG_06514 | eukaryota>stramenopiles | Phytophthora | infestans T30-4 | calmodulin [Phytophthora infestans T30-4] |
| XP_003056131.1 | 149 | CAM | eukaryota>viridiplantae>chlorophyta | Micromonas | pusilla CCMP1545 | calmodulin [Micromonas pusilla CCMP1545] |
| XP_003257649.1 | 149 | CALML3 | eukaryota>metazoa>chordata>vertebrata | Nomascus | leucogenys | PREDICTED: calmodulin-like protein 3 [Nomascus leucogenys] |
| XP_003260924.1 | 149 | CALM1 | eukaryota>metazoa>chordata>vertebrata | Nomascus | leucogenys | PREDICTED: calmodulin isoform X3 [Nomascus leucogenys] |
| XP_003262870.1 | 149 | CALM2 | eukaryota>metazoa>chordata>vertebrata | Nomascus | leucogenys | PREDICTED: calmodulin isoform X3 [Nomascus leucogenys] |
| XP_003277701.2 | 149 | CALM3 | eukaryota>metazoa>chordata>vertebrata | Nomascus | leucogenys | PREDICTED: calmodulin isoform X1 [Nomascus leucogenys] |
| XP_003329587.1 | 149 | PGTG_11337 | eukaryota>fungi>basidiomycota | Puccinia | graminis f. sp. tritici CRL 75-36-700-3 | calmodulin [Puccinia graminis f. sp. tritici CRL 75-36-700-3] |
| XP_003519492.1 | 149 | LOC100793700 | eukaryota>viridiplantae | Glycine | max | PREDICTED: calmodulin [Glycine max] |
| XP_003545473.1 | 149 | LOC100791702 | eukaryota>viridiplantae | Glycine | max | PREDICTED: calmodulin [Glycine max] |
| XP_003726932.1 | 149 | LOC591003 | eukaryota>metazoa>echinodermata | Strongylocentrotus | purpuratus | PREDICTED: calmodulin isoform X2 [Strongylocentrotus purpuratus] |
| XP_003858776.1 | 149 | LDBPK_090970 | eukaryota>euglenozoa>kinetoplastida | Leishmania | donovani | calmodulin, putative [Leishmania donovani] |
| XP_003858777.1 | 149 | LDBPK_090980 | eukaryota>euglenozoa>kinetoplastida | Leishmania | donovani | calmodulin, putative [Leishmania donovani] |
| XP_003872774.1 | 149 | LMXM_09_0910 | eukaryota>euglenozoa>kinetoplastida | Leishmania | mexicana MHOM/GT/2001/U1103 | putative calmodulin [Leishmania mexicana MHOM/GT/2001/U1103] |
| XP_003872775.1 | 149 | LMXM_09_0920 | eukaryota>euglenozoa>kinetoplastida | Leishmania | mexicana MHOM/GT/2001/U1103 | putative calmodulin [Leishmania mexicana MHOM/GT/2001/U1103] |
| XP_003872776.1 | 149 | LMXM_09_0930 | eukaryota>euglenozoa>kinetoplastida | Leishmania | mexicana MHOM/GT/2001/U1103 | putative calmodulin [Leishmania mexicana MHOM/GT/2001/U1103] |
| XP_003886144.1 | 149 | NCLIV_065440 | eukaryota>alveolata>apicomplexa | Neospora | caninum Liverpool | hypothetical protein NCLIV_065440 [Neospora caninum Liverpool] |
| XP_003963666.1 | 149 | calm2 | eukaryota>metazoa>chordata>vertebrata>actinopterygii | Takifugu | rubripes | PREDICTED: calmodulin [Takifugu rubripes] |
| XP_003968409.1 | 149 | calm3 | eukaryota>metazoa>chordata>vertebrata>actinopterygii | Takifugu | rubripes | PREDICTED: calmodulin [Takifugu rubripes] |
| XP_003971505.1 | 149 | LOC101068488 | eukaryota>metazoa>chordata>vertebrata>actinopterygii | Takifugu | rubripes | PREDICTED: calmodulin [Takifugu rubripes] |
| XP_003976021.1 | 149 | LOC101064505 | eukaryota>metazoa>chordata>vertebrata>actinopterygii | Takifugu | rubripes | PREDICTED: calmodulin-alpha-like [Takifugu rubripes] |
| XP_004039912.1 | 149 | IMG5_006070 | eukaryota>alveolata>ciliophora | Ichthyophthirius | multifiliis | hypothetical protein IMG5_006070 [Ichthyophthirius multifiliis] |
| XP_004091507.1 | 149 | CALM2 | eukaryota>metazoa>chordata>vertebrata | Nomascus | leucogenys | PREDICTED: calmodulin isoform X3 [Nomascus leucogenys] |
| XP_004349444.1 | 149 | CAOG_02694 | eukaryota>ichthyosporea | Capsaspora | owczarzaki ATCC 30864 | calmodulin [Capsaspora owczarzaki ATCC 30864] |
| XP_004829358.1 | 149 | BEWA_025410 | eukaryota>alveolata>apicomplexa | Theileria | equi | calmodulin, putative [Theileria equi] |
| XP_004993230.1 | 149 | PTSG_06340 | eukaryota>choanoflagellida | Salpingoeca | rosetta | calmodulin [Salpingoeca rosetta] |
| XP_005562071.1 | 149 | CALM1 | eukaryota>metazoa>chordata>vertebrata | Macaca | fascicularis | PREDICTED: calmodulin [Macaca fascicularis] |
| XP_005564612.1 | 149 | CALML3 | eukaryota>metazoa>chordata>vertebrata | Macaca | fascicularis | PREDICTED: calmodulin-like protein 3 [Macaca fascicularis] |
| XP_005836584.1 | 149 | GUITHDRAFT_151413 | eukaryota>cryptophyta | Guillardia | theta CCMP2712 | hypothetical protein GUITHDRAFT_151413 [Guillardia theta CCMP2712] |
| XP_005839886.1 | 149 | GUITHDRAFT_150466 | eukaryota>cryptophyta | Guillardia | theta CCMP2712 | hypothetical protein GUITHDRAFT_150466 [Guillardia theta CCMP2712] |
| XP_005843378.1 | 149 | CHLNCDRAFT_59820 | eukaryota>viridiplantae>chlorophyta | Chlorella | variabilis | hypothetical protein CHLNCDRAFT_59820 [Chlorella variabilis] |
| XP_006339234.1 | 149 | LOC102579473 | eukaryota>viridiplantae | Solanum | tuberosum | PREDICTED: calmodulin-7 [Solanum tuberosum] |
| XP_006364752.1 | 149 | LOC102599353 | eukaryota>viridiplantae | Solanum | tuberosum | PREDICTED: calmodulin [Solanum tuberosum] |
| XP_006565317.1 | 149 | LOC551859 | eukaryota>metazoa>hexapoda | Apis | mellifera | PREDICTED: calmodulin [Apis mellifera] |
| XP_006959848.1 | 149 | WALSEDRAFT_70094 | eukaryota>fungi>basidiomycota | Wallemia | mellicola CBS 633.66 | EF-hand [Wallemia mellicola CBS 633.66] |
| XP_007331882.1 | 149 | AGABI1DRAFT_86553 | eukaryota>fungi>basidiomycota | Agaricus | bisporus var. burnettii JB137-S8 | hypothetical protein AGABI1DRAFT_86553 [Agaricus bisporus var. burnettii JB137-S8] |
| XP_008607073.1 | 149 | SDRG_03224 | eukaryota>stramenopiles | Saprolegnia | diclina VS20 | calmodulin [Saprolegnia diclina VS20] |
| XP_008655487.1 | 149 | LOC103634668 | eukaryota>viridiplantae | Zea | mays | calmodulin [Zea mays] |
| XP_008660204.1 | 149 | LOC103639205 | eukaryota>viridiplantae | Zea | mays | calmodulin [Zea mays] |
| XP_008883927.1 | 149 | HHA_249240 | eukaryota>alveolata>apicomplexa | Hammondia | hammondi | calmodulin, putative [Hammondia hammondi] |
| XP_008908788.1 | 149 | PPTG_13130 | eukaryota>stramenopiles | Phytophthora | parasitica INRA-310 | calmodulin [Phytophthora parasitica INRA-310] |
| XP_009033803.1 | 149 | AURANDRAFT_36419 | eukaryota>stramenopiles | Aureococcus | anophagefferens | hypothetical protein AURANDRAFT_36419 [Aureococcus anophagefferens] |
| XP_009033883.1 | 149 | AURANDRAFT_58710 | eukaryota>stramenopiles | Aureococcus | anophagefferens | hypothetical protein AURANDRAFT_58710 [Aureococcus anophagefferens] |
| XP_009054739.1 | 149 | LOTGIDRAFT_215397 | eukaryota>metazoa>mollusca | Lottia | gigantea | hypothetical protein LOTGIDRAFT_215397 [Lottia gigantea] |
| XP_009056063.1 | 149 | LOTGIDRAFT_203998 | eukaryota>metazoa>mollusca | Lottia | gigantea | hypothetical protein LOTGIDRAFT_203998 [Lottia gigantea] |
| XP_009056065.1 | 149 | LOTGIDRAFT_216228 | eukaryota>metazoa>mollusca | Lottia | gigantea | hypothetical protein LOTGIDRAFT_216228 [Lottia gigantea] |
| XP_009315838.1 | 149 | DQ04_17211000 | eukaryota>euglenozoa>kinetoplastida | Trypanosoma | grayi | calmodulin [Trypanosoma grayi] |
| XP_009539924.1 | 149 | PHYSODRAFT_292780 | eukaryota>stramenopiles | Phytophthora | sojae | hypothetical protein PHYSODRAFT_292780 [Phytophthora sojae] |
| XP_009690713.1 | 149 | TOT_020000669 | eukaryota>alveolata>apicomplexa | Theileria | orientalis strain Shintoku | calmodulin [Theileria orientalis strain Shintoku] |
| XP_010645766.1 | 149 | LOC100260795 | eukaryota>viridiplantae | Vitis | vinifera | PREDICTED: calmodulin [Vitis vinifera] |
| XP_010704329.1 | 149 | LPMP_090920 | eukaryota>euglenozoa>kinetoplastida | Leishmania | panamensis | calmodulin, putative [Leishmania panamensis] |
| XP_010704330.1 | 149 | LPMP_090930 | eukaryota>euglenozoa>kinetoplastida | Leishmania | panamensis | calmodulin, putative [Leishmania panamensis] |
| XP_011129786.1 | 149 | GNI_051490 | eukaryota>alveolata>apicomplexa | Gregarina | niphandrodes | calmodulin [Gregarina niphandrodes] |
| XP_011139526.1 | 149 | LOC105183241 | eukaryota>metazoa>hexapoda | Harpegnathos | saltator | PREDICTED: calmodulin isoform X1 [Harpegnathos saltator] |
| XP_011139527.1 | 149 | LOC105183241 | eukaryota>metazoa>hexapoda | Harpegnathos | saltator | PREDICTED: calmodulin isoform X2 [Harpegnathos saltator] |
| XP_011390388.1 | 149 | UMAG_03910 | eukaryota>fungi>basidiomycota | Ustilago | maydis 521 | putative calmodulin [Ustilago maydis 521] |
| XP_011416022.2 | 149 | LOC105319976 | eukaryota>metazoa>mollusca | Crassostrea | gigas | PREDICTED: neo-calmodulin-like [Crassostrea gigas] |
| XP_011427008.1 | 149 | LOC105327998 | eukaryota>metazoa>mollusca | Crassostrea | gigas | PREDICTED: calmodulin-A-like [Crassostrea gigas] |
| XP_011427009.1 | 149 | LOC105327999 | eukaryota>metazoa>mollusca | Crassostrea | gigas | PREDICTED: calmodulin-A-like [Crassostrea gigas] |
| XP_011436587.1 | 149 | LOC105334732 | eukaryota>metazoa>mollusca | Crassostrea | gigas | PREDICTED: calmodulin-A [Crassostrea gigas] |
| XP_011436588.1 | 149 | LOC105334733 | eukaryota>metazoa>mollusca | Crassostrea | gigas | PREDICTED: calmodulin isoform X2 [Crassostrea gigas] |
| XP_011436589.1 | 149 | LOC105334734 | eukaryota>metazoa>mollusca | Crassostrea | gigas | PREDICTED: calmodulin-like [Crassostrea gigas] |
| XP_011444268.1 | 149 | LOC105340108 | eukaryota>metazoa>mollusca | Crassostrea | gigas | PREDICTED: calmodulin [Crassostrea gigas] |
| XP_011675926.1 | 149 | LOC105443934 | eukaryota>metazoa>echinodermata | Strongylocentrotus | purpuratus | PREDICTED: calmodulin-A-like [Strongylocentrotus purpuratus] |
| XP_011675927.1 | 149 | LOC585084 | eukaryota>metazoa>echinodermata | Strongylocentrotus | purpuratus | PREDICTED: calmodulin-beta [Strongylocentrotus purpuratus] |
| XP_011780594.1 | 149 | TbgDal_XI14490 | eukaryota>euglenozoa>kinetoplastida | Trypanosoma | brucei gambiense DAL972 | calmodulin, putative [Trypanosoma brucei gambiense DAL972] |
| XP_011780595.1 | 149 | TbgDal_XI14500 | eukaryota>euglenozoa>kinetoplastida | Trypanosoma | brucei gambiense DAL972 | calmodulin, putative [Trypanosoma brucei gambiense DAL972] |
| XP_011780596.1 | 149 | TbgDal_XI14510 | eukaryota>euglenozoa>kinetoplastida | Trypanosoma | brucei gambiense DAL972 | calmodulin, putative [Trypanosoma brucei gambiense DAL972] |
| XP_011780597.1 | 149 | TbgDal_XI14520 | eukaryota>euglenozoa>kinetoplastida | Trypanosoma | brucei gambiense DAL972 | calmodulin, putative [Trypanosoma brucei gambiense DAL972] |
| XP_012052369.1 | 149 | CNAG_01557 | eukaryota>fungi>basidiomycota | Cryptococcus | neoformans var. grubii H99 | calmodulin [Cryptococcus neoformans var. grubii H99] |
| XP_012195381.1 | 149 | SPRG_01624 | eukaryota>stramenopiles | Saprolegnia | parasitica CBS 223.65 | hypothetical protein SPRG_01624 [Saprolegnia parasitica CBS 223.65] |
| XP_012769886.1 | 149 | BBBOND_0401920 | eukaryota>alveolata>apicomplexa | Babesia | bigemina | calmodulin, putative [Babesia bigemina] |
| XP_013233382.1 | 149 | ETH_00032320 | eukaryota>alveolata>apicomplexa | Eimeria | tenella | calmodulin, putative [Eimeria tenella] |
| XP_013381516.1 | 149 | LOC106152471 | eukaryota>metazoa | Lingula | anatina | PREDICTED: calmodulin isoform X2 [Lingula anatina] |
| XP_013381517.1 | 149 | LOC106152471 | eukaryota>metazoa | Lingula | anatina | PREDICTED: calmodulin isoform X3 [Lingula anatina] |
| XP_013756148.1 | 149 | AMSG_07687 | eukaryota>apusozoa>apusomonadidae | Thecamonas | trahens ATCC 50062 | calmodulin [Thecamonas trahens ATCC 50062] |
| XP_013780919.1 | 149 | LOC106465255 | eukaryota>metazoa | Limulus | polyphemus | calmodulin [Limulus polyphemus] |
| XP_013781372.1 | 149 | LOC106465681 | eukaryota>metazoa | Limulus | polyphemus | calmodulin [Limulus polyphemus] |
| XP_013781807.1 | 149 | LOC106466113 | eukaryota>metazoa | Limulus | polyphemus | calmodulin [Limulus polyphemus] |
| XP_013781942.1 | 149 | LOC106466237 | eukaryota>metazoa | Limulus | polyphemus | calmodulin [Limulus polyphemus] |
| XP_013785128.1 | 149 | LOC106469207 | eukaryota>metazoa | Limulus | polyphemus | calmodulin [Limulus polyphemus] |
| XP_013901830.1 | 149 | MNEG_5147 | eukaryota>viridiplantae>chlorophyta | Monoraphidium | neglectum | calmodulin [Monoraphidium neglectum] |
| XP_014153486.1 | 149 | SARC_08017 | eukaryota>ichthyosporea | Sphaeroforma | arctica JP610 | calmodulin [Sphaeroforma arctica JP610] |
| XP_014536691.1 | 149 | PDIP_25360 | eukaryota>fungi>ascomycota | Penicillium | digitatum Pd1 | Calmodulin [Penicillium digitatum Pd1] |
| XP_014673041.1 | 149 | LOC106813418 | eukaryota>metazoa | Priapulus | caudatus | PREDICTED: calmodulin [Priapulus caudatus] |
| XP_014786992.1 | 149 | LOC106881201 | eukaryota>metazoa>mollusca | Octopus | bimaculoides | PREDICTED: calmodulin isoform X1 [Octopus bimaculoides] |
| XP_014787000.1 | 149 | LOC106881201 | eukaryota>metazoa>mollusca | Octopus | bimaculoides | PREDICTED: calmodulin isoform X2 [Octopus bimaculoides] |
| XP_015656044.1 | 149 | ABB37_06970 | eukaryota>euglenozoa>kinetoplastida | Leptomonas | pyrrhocoris | putative calmodulin [Leptomonas pyrrhocoris] |
| XP_015656045.1 | 149 | ABB37_06971 | eukaryota>euglenozoa>kinetoplastida | Leptomonas | pyrrhocoris | Trypanosoma cruzi [Leptomonas pyrrhocoris] |
| XP_015764671.1 | 149 | LOC107343607 | eukaryota>metazoa>cnidaria | Acropora | digitifera | PREDICTED: calmodulin [Acropora digitifera] |
| XP_016435527.1 | 149 | LOC107761764 | eukaryota>viridiplantae | Nicotiana | tabacum | PREDICTED: calmodulin-7 [Nicotiana tabacum] |
| XP_016437737.1 | 149 | LOC107763754 | eukaryota>viridiplantae | Nicotiana | tabacum | PREDICTED: calmodulin-like [Nicotiana tabacum] |
| XP_016450742.1 | 149 | LOC107775488 | eukaryota>viridiplantae | Nicotiana | tabacum | PREDICTED: calmodulin-7 [Nicotiana tabacum] |
| XP_016455453.1 | 149 | LOC107779518 | eukaryota>viridiplantae | Nicotiana | tabacum | PREDICTED: calmodulin [Nicotiana tabacum] |
| XP_016458299.1 | 149 | LOC107781984 | eukaryota>viridiplantae | Nicotiana | tabacum | PREDICTED: calmodulin [Nicotiana tabacum] |
| XP_016467127.1 | 149 | LOC107789771 | eukaryota>viridiplantae | Nicotiana | tabacum | PREDICTED: calmodulin [Nicotiana tabacum] |
| XP_016478865.1 | 149 | LOC107800228 | eukaryota>viridiplantae | Nicotiana | tabacum | PREDICTED: calmodulin-7 [Nicotiana tabacum] |
| XP_016496183.1 | 149 | LOC107815174 | eukaryota>viridiplantae | Nicotiana | tabacum | PREDICTED: calmodulin-7 isoform X1 [Nicotiana tabacum] |
| XP_016501754.1 | 149 | LOC107820061 | eukaryota>viridiplantae | Nicotiana | tabacum | PREDICTED: calmodulin-7 [Nicotiana tabacum] |
| XP_016509951.1 | 149 | LOC107827355 | eukaryota>viridiplantae | Nicotiana | tabacum | PREDICTED: calmodulin-7 isoform X1 [Nicotiana tabacum] |
| XP_016609142.1 | 149 | SPPG_04193 | eukaryota>fungi>chytridiomycota | Spizellomyces | punctatus DAOM BR117 | calmodulin [Spizellomyces punctatus DAOM BR117] |
| XP_016611050.1 | 149 | SPPG_02082 | eukaryota>fungi>chytridiomycota | Spizellomyces | punctatus DAOM BR117 | calmodulin [Spizellomyces punctatus DAOM BR117] |
| XP_017312248.1 | 149 | LOC108258233 | eukaryota>metazoa>chordata>vertebrata>actinopterygii | Ictalurus | punctatus | PREDICTED: calmodulin [Ictalurus punctatus] |
| XP_017319419.1 | 149 | LOC108263280 | eukaryota>metazoa>chordata>vertebrata>actinopterygii | Ictalurus | punctatus | PREDICTED: calmodulin [Ictalurus punctatus] |
| XP_017339860.1 | 149 | calm2 | eukaryota>metazoa>chordata>vertebrata>actinopterygii | Ictalurus | punctatus | PREDICTED: calmodulin [Ictalurus punctatus] |
| XP_017346481.1 | 149 | calm3 | eukaryota>metazoa>chordata>vertebrata>actinopterygii | Ictalurus | punctatus | PREDICTED: calmodulin [Ictalurus punctatus] |
| XP_017348609.1 | 149 | LOC108279128 | eukaryota>metazoa>chordata>vertebrata>actinopterygii | Ictalurus | punctatus | PREDICTED: calmodulin [Ictalurus punctatus] |
| XP_018085904.1 | 149 | LOC108698710 | eukaryota>metazoa>chordata>vertebrata | Xenopus | laevis | PREDICTED: calmodulin [Xenopus laevis] |
| XP_018187267.1 | 149 | L228DRAFT_248440 | eukaryota>fungi>ascomycota | Xylona | heveae TC161 | EF-hand protein [Xylona heveae TC161] |
| XP_018291668.1 | 149 | Cam1 | eukaryota>fungi | Phycomyces | blakesleeanus NRRL 1555(-) | calmodulin [Phycomyces blakesleeanus NRRL 1555(-)] |
| XP_019016027.1 | 149 | PICMEDRAFT_12670 | eukaryota>fungi>ascomycota | Pichia | membranifaciens NRRL Y-2026 | hypothetical protein PICMEDRAFT_12670 [Pichia membranifaciens NRRL Y-2026] |
| XP_019077160.1 | 149 | LOC100232908 | eukaryota>viridiplantae | Vitis | vinifera | PREDICTED: calmodulin [Vitis vinifera] |
| XP_019330937.1 | 149 | CMD1 | eukaryota>fungi>ascomycota | Candida | albicans SC5314 | calmodulin [Candida albicans SC5314] |
| XP_727667.1 | 149 | PY17X_1012200 | eukaryota>alveolata>apicomplexa | Plasmodium | yoelii | calmodulin, putative [Plasmodium yoelii] |
| XP_765284.1 | 149 | TP02_0717 | eukaryota>alveolata>apicomplexa | Theileria | parva strain Muguga | calmodulin [Theileria parva strain Muguga] |
| XP_780925.4 | 149 | LOC575430 | eukaryota>metazoa>echinodermata | Strongylocentrotus | purpuratus | PREDICTED: calmodulin [Strongylocentrotus purpuratus] |
| XP_805243.1 | 149 | Tc00.1047053506391.10 | eukaryota>euglenozoa>kinetoplastida | Trypanosoma | cruzi strain CL Brener | calmodulin [Trypanosoma cruzi strain CL Brener] |
| XP_805244.1 | 149 | Tc00.1047053506391.20 | eukaryota>euglenozoa>kinetoplastida | Trypanosoma | cruzi strain CL Brener | calmodulin [Trypanosoma cruzi strain CL Brener] |
| XP_808090.1 | 149 | Tc00.1047053507483.50 | eukaryota>euglenozoa>kinetoplastida | Trypanosoma | cruzi strain CL Brener | calmodulin [Trypanosoma cruzi strain CL Brener] |
| XP_808093.1 | 149 | Tc00.1047053507483.39 | eukaryota>euglenozoa>kinetoplastida | Trypanosoma | cruzi strain CL Brener | calmodulin [Trypanosoma cruzi strain CL Brener] |
| XP_951999.1 | 149 | TA14735 | eukaryota>alveolata>apicomplexa | Theileria | annulata | calmodulin, putative [Theileria annulata] |
| XP_961276.2 | 149 | cmd | eukaryota>fungi>ascomycota | Neurospora | crassa OR74A | calmodulin A [Neurospora crassa OR74A] |
| XP_972156.1 | 149 | LOC660864 | eukaryota>metazoa>hexapoda | Tribolium | castaneum | PREDICTED: calmodulin [Tribolium castaneum] |
| Bcir1000006777 | 149 | Bcir1000006777 | eukaryota>fungi>mucoromycotina | Backusella | circina | >Bcir1000006777 |
| Lhya1000001900 | 149 | Lhya1000001900 | eukaryota>fungi>mucoromycotina | Lichtheimia | hyalospora | >Lhya1000001900 |
| OHS94714.1 | 149 | CMD1 | eukaryota>parabasalia | Tritrichomonas | foetus | Calmodulin [Tritrichomonas foetus] |
| OHT04308.1 | 149 | CMD1 | eukaryota>parabasalia | Tritrichomonas | foetus | Calmodulin [Tritrichomonas foetus] |
| XP_001323137.1 | 149 | TVAG_259990 | eukaryota>parabasalia | Trichomonas | vaginalis G3 | calmodulin [Trichomonas vaginalis G3] |
| XP_001325744.1 | 149 | TVAG_343500 | eukaryota>parabasalia | Trichomonas | vaginalis G3 | calmodulin [Trichomonas vaginalis G3] |
| XP_001328926.1 | 149 | TVAG_067080 | eukaryota>parabasalia | Trichomonas | vaginalis G3 | calmodulin [Trichomonas vaginalis G3] |
| NP_001235963.1 | 149 | LOC100500550 | eukaryota>viridiplantae | Glycine | max | uncharacterized protein LOC100500550 [Glycine max] |
| XP_009494169.1 | 149 | H696_01995 | [illegible] | Fonticula | alba | calmodulin [Fonticula alba] |
| XP_793110.3 | 149 | LOC588327 | eukaryota>metazoa>echinodermata | Strongylocentrotus | purpuratus | PREDICTED: calmodulin-alpha [Strongylocentrotus purpuratus] |
| OEU22277.1 | 149 | FRACYDRAFT_166325 | eukaryota>stramenopiles | Fragilariopsis | cylindrus CCMP1102 | calmodulin [Fragilariopsis cylindrus CCMP1102] |
| XP_002598167.1 | 149 | BRAFLDRAFT_114725 | eukaryota>metazoa>chordata | Branchiostoma | floridae | hypothetical protein BRAFLDRAFT_114725 [Branchiostoma floridae] |

[illegible]

| | | | | | | |
|---|---|---|---|---|---|---|
| XP_002285864.1 | 499 | LOC100259138 | eukaryota>viridiplantae | Vitis | vinifera | PREDICTED: protein disulfide-isomerase [Vitis vinifera] |
| XP_005837233.1 | 499 | GUITHDRAFT_85427 | eukaryota>cryptophyta | Guillardia | theta CCMP2712 | hypothetical protein GUITHDRAFT_85427 [Guillardia theta CCMP2712] |
| XP_014785522.1 | 499 | LOC106880182 | eukaryota>metazoa>mollusca | Octopus | bimaculoides | PREDICTED: protein disulfide-isomerase A3-like [Octopus bimaculoides] |
| ORY00427.1 | 499 | K493DRAFT_103679 | eukaryota>fungi | Basidiobolus | meristosporus CBS 931.73 | hypothetical protein K493DRAFT_103679 [Basidiobolus meristosporus CBS 931.73] |
| EIE76133.1 | 498 | RO3G_00837 | eukaryota>fungi | Rhizopus | delemar RA 99-880 | hypothetical protein RO3G_00837 [Rhizopus delemar RA 99-880] |
| KXN72927.1 | 498 | CONCODRAFT_15852 | eukaryota>fungi | Conidiobolus | coronatus NRRL 28638 | protein disulfide-isomerase [Conidiobolus coronatus NRRL 28638] |
| XP_001766340.1 | 498 | PHYPADRAFT_130646 | eukaryota>viridiplantae | Physcomitrella | patens | predicted protein [Physcomitrella patens] |
| XP_008190988.1 | 498 | LOC103312345 | eukaryota>metazoa>hexapoda | Tribolium | castaneum | PREDICTED: protein disulfide-isomerase [Tribolium castaneum] |
| XP_008877298.1 | 498 | H310_12113 | eukaryota>stramenopiles | Aphanomyces | invadans | protein disulfide-isomerase domain [Aphanomyces invadans] |
| XP_009833015.1 | 498 | H257_08690 | eukaryota>stramenopiles | Aphanomyces | astaci | protein disulfide-isomerase domain [Aphanomyces astaci] |
| Lhya1000002422 | 498 | Lhya1000002422 | eukaryota>fungi>mucoromycotina | Lichtheimia | hyalospora | >Lhya1000002422 |
| XP_001897232.1 | 497 | Bm1_28935 | eukaryota>metazoa>nematoda | Brugia | malayi | transglutaminase [Brugia malayi] |
| XP_006363890.1 | 497 | LOC102592089 | eukaryota>viridiplantae | Solanum | tuberosum | PREDICTED: protein disulfide-isomerase-like [Solanum tuberosum] |
| XP_011138070.1 | 497 | LOC105182389 | eukaryota>metazoa>hexapoda | Harpegnathos | saltator | PREDICTED: protein disulfide-isomerase [Harpegnathos saltator] |
| XP_013778350.1 | 497 | LOC106462929 | eukaryota>metazoa | Limulus | polyphemus | PREDICTED: protein disulfide-isomerase 2-like [Limulus polyphemus] |
| XP_015159154.1 | 497 | LOC102592089 | eukaryota>viridiplantae | Solanum | tuberosum | PREDICTED: protein disulfide-isomerase-like [Solanum tuberosum] |
| XP_015159155.1 | 497 | LOC102592089 | eukaryota>viridiplantae | Solanum | tuberosum | PREDICTED: protein disulfide-isomerase-like [Solanum tuberosum] |
| Bcir1000001554 | 497 | Bcir1000001554 | eukaryota>fungi>mucoromycotina | Backusella | circina | >Bcir1000001554 |
| XP_011778178.1 | 497 | TbgDal_X10050 | eukaryota>euglenozoa>kinetoplastida | Trypanosoma | brucei gambiense DAL972 | bloodstream-specific protein 2 precursor,putative [Trypanosoma brucei gambiense DAL972] |
| OWA54404.1 | 497 | BV898_18808 | eukaryota>metazoa | Hypsibius | dujardini | Protein disulfide-isomerase [Hypsibius dujardini] |
| EPB91883.1 | 496 | HMPREF1544_01177 | eukaryota>fungi | Mucor | circinelloides f. circinelloides 1006PhL | hypothetical protein HMPREF1544_01177 [Mucor circinelloides f. circinelloides 1006PhL] |
| NP_001287070.1 | 496 | CG6988,Dmel\CG6988,dpdi,Dpdi,EP(3)1022,pdi,PDI | eukaryota>metazoa>hexapoda | Drosophila | melanogaster | protein disulfide isomerase, isoform E [Drosophila melanogaster] |
| NP_524079.1 | 496 | CG6988,Dmel\CG6988,dpdi,Dpdi,EP(3)1022,pdi,PDI | eukaryota>metazoa>hexapoda | Drosophila | melanogaster | protein disulfide isomerase, isoform A [Drosophila melanogaster] |
| NP_999697.1 | 496 | ECaSt/PDI | eukaryota>metazoa>echinodermata | Strongylocentrotus | purpuratus | ER calcistorin precursor [Strongylocentrotus purpuratus] |
| XP_003545643.1 | 496 | LOC100788702 | eukaryota>viridiplantae | Glycine | max | PREDICTED: protein disulfide-isomerase-like isoform X1 [Glycine max] |
| XP_005647426.1 | 496 | COCSUDRAFT_16178 | eukaryota>viridiplantae>chlorophyta | Coccomyxa | subellipsoidea C-169 | protein disulfide isomerase [Coccomyxa subellipsoidea C-169] |
| XP_006593329.1 | 496 | LOC100305460 | eukaryota>viridiplantae | Glycine | max | PREDICTED: protein disulfide isomerase like protein isoform X1 [Glycine max] |
| XP_008616243.1 | 496 | SDRG_11966 | eukaryota>stramenopiles | Saprolegnia | diclina VS20 | hypothetical protein SDRG_11966 [Saprolegnia diclina VS20] |
| XP_009028587.1 | 496 | HELRODRAFT_194128 | eukaryota>metazoa>annelida | Helobdella | robusta | hypothetical protein HELRODRAFT_194128 [Helobdella robusta] |
| XP_012198778.1 | 496 | SPRG_04464 | eukaryota>stramenopiles | Saprolegnia | parasitica CBS 223.65 | hypothetical protein SPRG_04464 [Saprolegnia parasitica CBS 223.65] |
| XP_017330169.1 | 496 | pdia3 | eukaryota>metazoa>chordata>vertebrata>actinopterygii | Ictalurus | punctatus | PREDICTED: protein disulfide-isomerase A3 [Ictalurus punctatus] |
| CEM39955.1 | 495 | Vbra_19861 | eukaryota>alveolata | Vitrella | brassicaformis CCMP3155 | unnamed protein product [Vitrella brassicaformis CCMP3155] |
| XP_001761068.1 | 495 | PHYPADRAFT_123708 | eukaryota>viridiplantae | Physcomitrella | patens | predicted protein [Physcomitrella patens] |
| XP_003969188.1 | 495 | LOC101067334 | eukaryota>metazoa>chordata>vertebrata>actinopterygii | Takifugu | rubripes | PREDICTED: protein disulfide-isomerase A3-like [Takifugu rubripes] |
| XP_006593605.1 | 495 | LOC100789569 | eukaryota>viridiplantae | Glycine | max | PREDICTED: protein disulfide-isomerase-like [Glycine max] |
| XP_011445708.1 | 495 | LOC105341095 | eukaryota>metazoa>mollusca | Crassostrea | gigas | PREDICTED: protein disulfide-isomerase [Crassostrea gigas] |
| XP_016770060.1 | 495 | LOC551435 | eukaryota>metazoa>hexapoda | Apis | mellifera | PREDICTED: protein disulfide-isomerase [Apis mellifera] |
| XP_017337773.1 | 495 | LOC108273190 | eukaryota>metazoa>chordata>vertebrata>actinopterygii | Ictalurus | punctatus | PREDICTED: protein disulfide-isomerase A3-like [Ictalurus punctatus] |
| ORX80283.1 | 495 | K493DRAFT_320917 | eukaryota>fungi | Basidiobolus | meristosporus CBS 931.73 | protein disulfide isomerase [Basidiobolus meristosporus CBS 931.73] |
| OAD04586.1 | 495 | MUCCIDRAFT_177573 | eukaryota>fungi | Mucor | circinelloides f. lusitanicus CBS 277.49 | hypothetical protein MUCCIDRAFT_177573 [Mucor circinelloides f. lusitanicus CBS 277.49] |
| CBJ29913.1 | 494 | Pdi3 | eukaryota>stramenopiles | Ectocarpus | siliculosus | protein disulfide isomerase [Ectocarpus siliculosus] |
| NP_001037171.1 | 494 | Pdi | eukaryota>metazoa>hexapoda | Bombyx | mori | protein disulfide isomerase precursor [Bombyx mori] |
| NP_001186666.1 | 494 | Grp58,sb:cb825 | eukaryota>metazoa>chordata>vertebrata>actinopterygii | Danio | rerio | protein disulfide-isomerase A3 precursor [Danio rerio] |
| XP_003967256.1 | 494 | pdia3 | eukaryota>metazoa>chordata>vertebrata>actinopterygii | Takifugu | rubripes | PREDICTED: protein disulfide-isomerase A3 [Takifugu rubripes] |
| Smar1000001458 | 494 | Smar1000001458 | eukaryota>metazoa>arthropoda>myriapoda | Strigamia | maritima | >Smar1000001458 |
| NP_001003517.1 | 493 | zgc:100906 | eukaryota>metazoa>chordata>vertebrata>actinopterygii | Danio | rerio | protein disulfide-isomerase A3 precursor [Danio rerio] |
| NP_508778.1 | 493 | pdi-2 | eukaryota>metazoa>nematoda | Caenorhabditis | elegans | Protein disulfide-isomerase 2 [Caenorhabditis elegans] |
| XP_001784758.1 | 493 | PHYPADRAFT_154382 | eukaryota>viridiplantae | Physcomitrella | patens | predicted protein [Physcomitrella patens] |
| XP_001847538.1 | 493 | CpipJ_CPIJ005219 | eukaryota>metazoa>hexapoda | Culex | quinquefasciatus | disulfide isomerase [Culex quinquefasciatus] |
| XP_001864249.1 | 493 | CpipJ_CPIJ013559 | eukaryota>metazoa>hexapoda | Culex | quinquefasciatus | disulfide isomerase [Culex quinquefasciatus] |
| XP_002964632.1 | 493 | SELMODRAFT_142613 | eukaryota>viridiplantae | Selaginella | moellendorffii | hypothetical protein SELMODRAFT_142613 [Selaginella moellendorffii] |
| XP_011151095.1 | 493 | LOC105190195 | eukaryota>metazoa>hexapoda | Harpegnathos | saltator | PREDICTED: protein disulfide-isomerase A3 [Harpegnathos saltator] |
| XP_016605428.1 | 493 | SPPG_07316 | eukaryota>fungi>chytridiomycota | Spizellomyces | punctatus DAOM BR117 | protein disulfide isomerase [Spizellomyces punctatus DAOM BR117] |
| XP_017327509.1 | 493 | LOC108267688 | eukaryota>metazoa>chordata>vertebrata>actinopterygii | Ictalurus | punctatus | PREDICTED: protein disulfide-isomerase A3-like isoform X1 [Ictalurus punctatus] |
| Bcir1000015014 | 493 | Bcir1000015014 | eukaryota>fungi>mucoromycotina | Backusella | circina | >Bcir1000015014 |
| Uram1000005567 | 493 | Uram1000005567 | eukaryota>fungi>mucoromycotina | Umbelopsis | ramanniana | >Uram1000005567 |
| XP_002788655.1 | 493 | Pmar_PMAR010193 | eukaryota>alveolata | Perkinsus | marinus ATCC 50983 | protein disulfide isomerase, putative [Perkinsus marinus ATCC 50983] |
| NP_592871.1 | 492 | SPAC1F5.02 | eukaryota>fungi>ascomycota | Schizosaccharomyces | pombe | protein disulfide isomerase (predicted) [Schizosaccharomyces pombe] |
| XP_003380462.1 | 492 | Tsp_06198 | eukaryota>metazoa>nematoda | Trichinella | spiralis | protein disulfide-isomerase 2 [Trichinella spiralis] |
| XP_011453191.1 | 492 | LOC105346349 | eukaryota>metazoa>mollusca | Crassostrea | gigas | PREDICTED: protein disulfide-isomerase A3 [Crassostrea gigas] |
| XP_012053226.1 | 492 | CNAG_06240 | eukaryota>fungi>basidiomycota | Cryptococcus | neoformans var. grubii H99 | protein disulfide-isomerase [Cryptococcus neoformans var. grubii H99] |
| XP_013389396.1 | 492 | LOC106158083 | eukaryota>metazoa | Lingula | anatina | PREDICTED: LOW QUALITY PROTEIN: protein disulfide-isomerase A3-like [Lingula anatina] |
| XP_014571096.1 | 492 | L969DRAFT_84341 | eukaryota>fungi>basidiomycota | Mixia | osmundae IAM 14324 | hypothetical protein L969DRAFT_84341 [Mixia osmundae IAM 14324] |
| XP_018268022.1 | 492 | RHOBADRAFT_66973 | eukaryota>fungi>basidiomycota | Rhodotorula | graminis WP1 | hypothetical protein RHOBADRAFT_66973 [Rhodotorula graminis WP1] |
| XP_971685.1 | 492 | LOC660355 | eukaryota>metazoa>hexapoda | Tribolium | castaneum | PREDICTED: protein disulfide-isomerase A3 [Tribolium castaneum] |
| XP_002765336.1 | 492 | Pmar_PMAR016131 | eukaryota>alveolata | Perkinsus | marinus ATCC 50983 | protein disulfide isomerase, putative [Perkinsus marinus ATCC 50983] |
| XP_005711820.1 | 492 | CHC_T00009389001 | eukaryota>rhodophyta | Chondrus | crispus | Protein disulfide-isomerase, PDIA-4 Endoplasmic reticulum lumen [Chondrus crispus] |
| XP_021881607.1 | 492 | BCR41DRAFT_352944 | eukaryota>fungi | Lobosporangium | transversale | hypothetical protein BCR41DRAFT_352944 [Lobosporangium transversale] |
| CAG06136.1 | 491 | GSTEN:00026385:G:001 | eukaryota>metazoa>chordata>vertebrata>actinopterygii | Tetraodon | nigroviridis | unnamed protein product, partial [Tetraodon nigroviridis] |
| NP_001036997.1 | 491 | ERp57 | eukaryota>metazoa>hexapoda | Bombyx | mori | protein disulfide-isomerase like protein ERp57 precursor [Bombyx mori] |
| XP_002109713.1 | 491 | TRIADDRAFT_52862 | eukaryota>metazoa>placozoa | Trichoplax | adhaerens | hypothetical protein TRIADDRAFT_52862 [Trichoplax adhaerens] |
| XP_003384866.1 | 491 | LOC100638379 | eukaryota>metazoa | Amphimedon | queenslandica | PREDICTED: protein disulfide-isomerase A3-like [Amphimedon queenslandica] |
| XP_004037058.1 | 491 | IMG5_062500 | eukaryota>alveolata>ciliophora | Ichthyophthirius | multifiliis | hypothetical protein IMG5_062500 [Ichthyophthirius multifiliis] |
| XP_006959979.1 | 491 | WALSEDRAFT_61210 | eukaryota>fungi>basidiomycota | Wallemia | mellicola CBS 633.66 | protein disulfide isomerase [Wallemia mellicola CBS 633.66] |
| XP_012352692.1 | 491 | PDIA2 | eukaryota>metazoa>chordata>vertebrata | Nomascus | leucogenys | PREDICTED: protein disulfide-isomerase A2 [Nomascus leucogenys] |
| XP_016515375.1 | 491 | LOC107832078 | eukaryota>viridiplantae | Nicotiana | tabacum | PREDICTED: protein disulfide-isomerase-like [Nicotiana tabacum] |
| XP_970692.1 | 491 | LOC659274 | eukaryota>metazoa>hexapoda | Tribolium | castaneum | PREDICTED: protein disulfide-isomerase A3 [Tribolium castaneum] |
| Capa1000012415 | 491 | Capa1000012415 | eukaryota>metazoa>annelida | Capitella | sp1 | >Capa1000012415 |
| Pram1000013023 | 491 | Pram1000013023 | eukaryota>stramenopiles | Phytophthora | ramorum | >Pram1000013023 |
| Smed1000024692 | 491 | Smed1000024692 | eukaryota>metazoa>platyhelminthe | Schmidtea | mediterranea | >Smed1000024692 |
| XP_725916.1 | 491 | PY00638 | eukaryota>alveolata>apicomplexa | Plasmodium | yoelii yoelii 17XNL | protein disulfide isomerase, partial [Plasmodium yoelii yoelii 17XNL] |
| CAG01048.1 | 490 | GSTEN:00019677:G:001 | eukaryota>metazoa>chordata>vertebrata>actinopterygii | Tetraodon | nigroviridis | unnamed protein product, partial [Tetraodon nigroviridis] |
| KXJ25978.1 | 490 | PDIA3 | eukaryota>metazoa>cnidaria | Exaiptasia | pallida | Protein disulfide-isomerase A3 [Exaiptasia pallida] |
| XP_001950406.1 | 490 | LOC100164598 | eukaryota>metazoa>hexapoda | Acyrthosiphon | pisum | PREDICTED: protein disulfide-isomerase A3 [Acyrthosiphon pisum] |
| XP_002160150.2 | 490 | LOC100203161 | eukaryota>metazoa>cnidaria | Hydra | vulgaris | PREDICTED: protein disulfide-isomerase A3-like [Hydra vulgaris] |
| XP_006338174.1 | 490 | LOC102598658 | eukaryota>viridiplantae | Solanum | tuberosum | PREDICTED: protein disulfide-isomerase-like [Solanum tuberosum] |
| XP_009030127.1 | 490 | HELRODRAFT_70378 | eukaryota>metazoa>annelida | Helobdella | robusta | hypothetical protein HELRODRAFT_70378 [Helobdella robusta] |
| XP_623282.1 | 490 | LOC409911 | eukaryota>metazoa>hexapoda | Apis | mellifera | PREDICTED: protein disulfide-isomerase A3 [Apis mellifera] |
| XP_976697.1 | 490 | TTHERM_00549480 | eukaryota>alveolata>ciliophora | Tetrahymena | thermophila SB210 | protein disulfide isomerase [Tetrahymena thermophila SB210] |
| OHS95737.1 | 490 | TRFO_10394 | eukaryota>parabasalia | Tritrichomonas | foetus | hypothetical protein TRFO_10394 [Tritrichomonas foetus] |
| EFX70620.1 | 489 | DAPPUDRAFT_202253 | eukaryota>metazoa>crustacea | Daphnia | pulex | hypothetical protein DAPPUDRAFT_202253 [Daphnia pulex] |
| NP_725084.2 | 489 | anon-EST:fe3D9,CG8983,D-ERp60,Dmel\CG8983,Erp60,Ov6 | eukaryota>metazoa>hexapoda | Drosophila | melanogaster | ERp60 [Drosophila melanogaster] |
| XP_009056950.1 | 489 | LOTGIDRAFT_190816 | eukaryota>metazoa>mollusca | Lottia | gigantea | hypothetical protein LOTGIDRAFT_190816 [Lottia gigantea] |
| KFH72373.1 | 488 | MVEG_02664 | eukaryota>fungi | Mortierella | verticillata NRRL 6337 | prolyl 4-hydroxylase, beta polypeptide [Mortierella verticillata NRRL 6337] |
| NP_491995.1 | 488 | pdi-3 | eukaryota>metazoa>nematoda | Caenorhabditis | elegans | Protein disulfide-isomerase [Caenorhabditis elegans] |
| NP_998070.1 | 488 | zgc:77086 | eukaryota>metazoa>chordata>vertebrata>actinopterygii | Danio | rerio | zgc:77086 precursor [Danio rerio] |
| XP_006524935.1 | 488 | 1810041F13Rik,AI661267,Pdip,Pdipl | eukaryota>metazoa>chordata>vertebrata | Mus | musculus | PREDICTED: protein disulfide-isomerase A2 isoform X1 [Mus musculus] |
| XP_018281499.1 | 488 | CC85DRAFT_241448 | eukaryota>fungi>basidiomycota | Cutaneotrichosporon | oleaginosum | protein disulfide isomerase [Cutaneotrichosporon oleaginosum] |
| XP_002764929.1 | 488 | Pmar_PMAR007496 | eukaryota>alveolata | Perkinsus | marinus ATCC 50983 | protein disulfide isomerase, putative [Perkinsus marinus ATCC 50983] |
| EJY66972.1 | 487 | OXYTRI_12735 | eukaryota>alveolata>ciliophora | Oxytricha | trifallax | Disulfide isomerase (macronuclear) [Oxytricha trifallax] |
| EJY67614.1 | 487 | OXYTRI_01869 | eukaryota>alveolata>ciliophora | Oxytricha | trifallax | Disulfide isomerase (macronuclear) [Oxytricha trifallax] |
| ERG84937.1 | 487 | ASU_04589 | eukaryota>metazoa>nematoda | Ascaris | suum | protein disulfide-isomerase a3 [Ascaris suum] |
| NP_849696.1 | 487 | ARABIDOPSIS,THALIANA,PROTEIN,DISULFIDE,ISOMERASE,5,ATPDI5,ATPDIL1-1,F8K7.19,F8K7_19,PDI-like,1-1,PDI5,PROTEIN,DISULFIDE,ISOMERASE,5 | eukaryota>viridiplantae | Arabidopsis | thaliana | PDI-like 1-1 [Arabidopsis thaliana] |
| XP_001623734.1 | 487 | NEMVEDRAFT_v1g218982 | eukaryota>metazoa>cnidaria | Nematostella | vectensis | predicted protein [Nematostella vectensis] |
| XP_001899304.1 | 487 | Bm1_39250 | eukaryota>metazoa>nematoda | Brugia | malayi | protein disulphide isomerase [Brugia malayi] |
| XP_004365843.1 | 487 | CAOG_00972 | eukaryota>ichthyosporea | Capsaspora | owczarzaki ATCC 30864 | protein disulfide-isomerase ERp60 [Capsaspora owczarzaki ATCC 30864] |
| XP_014157833.1 | 487 | SARC_03846 | eukaryota>ichthyosporea | Sphaeroforma | arctica JP610 | hypothetical protein SARC_03846 [Sphaeroforma arctica JP610] |
| Smed1000021326 | 487 | Smed1000021326 | eukaryota>metazoa>platyhelminthe | Schmidtea | mediterranea | >Smed1000021326 |
| CDW72484.1 | 486 | Contig4677.g4996 | eukaryota>alveolata>ciliophora | Stylonychia | lemnae | disulfide isomerase [Stylonychia lemnae] |
| XP_002406442.1 | 486 | IscW_ISCW016161 | eukaryota>metazoa | Ixodes | scapularis | protein disulfide isomerase, putative [Ixodes scapularis] |
| XP_002595781.1 | 486 | BRAFLDRAFT_287617 | eukaryota>metazoa>chordata | Branchiostoma | floridae | hypothetical protein BRAFLDRAFT_287617 [Branchiostoma floridae] |
| XP_003373217.1 | 486 | Tsp_10062 | eukaryota>metazoa>nematoda | Trichinella | spiralis | protein disulfide-isomerase A3 [Trichinella spiralis] |
| XP_013776776.1 | 486 | LOC106461500 | eukaryota>metazoa | Limulus | polyphemus | protein disulfide-isomerase A3-like isoform X1 [Limulus polyphemus] |
| XP_014662378.1 | 486 | LOC106805339 | eukaryota>metazoa | Priapulus | caudatus | PREDICTED: protein disulfide-isomerase A3-like [Priapulus caudatus] |
| XP_013231631.1 | 486 | ETH_00006210 | eukaryota>alveolata>apicomplexa | Eimeria | tenella | protein disulfide isomerase, putative [Eimeria tenella] |
| OHS96937.1 | 486 | TRFO_02072 | eukaryota>parabasalia | Tritrichomonas | foetus | hypothetical protein TRFO_02072 [Tritrichomonas foetus] |
| NP_497746.1 | 485 | pdi-1 | eukaryota>metazoa>nematoda | Caenorhabditis | elegans | Protein disulfide-isomerase 1 [Caenorhabditis elegans] |
| XP_001023996.2 | 485 | TTHERM_00475140 | eukaryota>alveolata>ciliophora | Tetrahymena | thermophila SB210 | protein disulfide-isomerase [Tetrahymena thermophila SB210] |
| XP_002740183.1 | 485 | LOC100374196 | eukaryota>metazoa>hemichordata | Saccoglossus | kowalevskii | PREDICTED: protein disulfide-isomerase A3-like [Saccoglossus kowalevskii] |
| XP_004088301.1 | 485 | PDIA3 | eukaryota>metazoa>chordata>vertebrata | Nomascus | leucogenys | PREDICTED: protein disulfide-isomerase A3 isoform X2 [Nomascus leucogenys] |
| XP_009308609.1 | 485 | DQ04_01181060 | eukaryota>euglenozoa>kinetoplastida | Trypanosoma | grayi | protein disulfide isomerase [Trypanosoma grayi] |
| XP_001866126.1 | 484 | CpipJ_CPIJ015893 | eukaryota>metazoa>hexapoda | Culex | quinquefasciatus | disulfide isomerase [Culex quinquefasciatus] |
| XP_004991467.1 | 484 | PTSG_07668 | eukaryota>choanoflagellida | Salpingoeca | rosetta | disulfide isomerase [Salpingoeca rosetta] |
| XP_009170645.1 | 484 | T265_06975 | eukaryota>metazoa | Opisthorchis | viverrini | hypothetical protein T265_06975 [Opisthorchis viverrini] |
| XP_018651240.1 | 484 | Smp_079770.1 | eukaryota>metazoa | Schistosoma | mansoni | putative protein disulfide-isomerase ER-60 precursor (ERP60) [Schistosoma mansoni] |
| EFA77013.1 | 483 | PPL_09765 | eukaryota>amoebozoa>mycetozoa>dictyostelida | Polysphondylium | pallidum PN500 | protein disulfide isomerase [Polysphondylium pallidum PN500] |
| EUB56196.1 | 483 | EGR_08944 | eukaryota>metazoa | Echinococcus | granulosus | Protein disulfide-isomerase [Echinococcus granulosus] |
| XP_001437694.1 | 483 | GSPATT00038399001 | eukaryota>alveolata>ciliophora | Paramecium | tetraurelia strain d4-2 | hypothetical protein (macronuclear) [Paramecium tetraurelia strain d4-2] |
| XP_004032029.1 | 483 | IMG5_131860 | eukaryota>alveolata>ciliophora | Ichthyophthirius | multifiliis | prolyl 4- beta subunit precursor, putative [Ichthyophthirius multifiliis] |
| XP_011611082.1 | 483 | LOC101064129 | eukaryota>metazoa>chordata>vertebrata>actinopterygii | Takifugu | rubripes | PREDICTED: protein disulfide-isomerase-like isoform X2 [Takifugu rubripes] |
| XP_002808883.1 | 483 | PfPDI-8 | eukaryota>alveolata>apicomplexa | Plasmodium | falciparum 3D7 | protein disulfide isomerase [Plasmodium falciparum 3D7] |
| XP_015754696.1 | 482 | LOC107334277 | eukaryota>metazoa>cnidaria | Acropora | digitifera | PREDICTED: protein disulfide-isomerase A3-like [Acropora digitifera] |
| XP_018646261.1 | 482 | Smp_056760 | eukaryota>metazoa | Schistosoma | mansoni | protein disulfide-isomerase,putative [Schistosoma mansoni] |
| XP_001614884.1 | 482 | PVX_088960 | eukaryota>alveolata>apicomplexa | Plasmodium | vivax Sal-1 | protein disulfide isomerase [Plasmodium vivax Sal-1] |
| ELR12342.1 | 482 | ACA1_374090 | eukaryota>amoebozoa>acanthamoebidae | Acanthamoeba | castellanii str. Neff | protein disulfideisomerase domain containing protein [Acanthamoeba castellanii str. Neff] |
| XP_001628893.1 | 481 | NEMVEDRAFT_v1g118540 | eukaryota>metazoa>cnidaria | Nematostella | vectensis | predicted protein, partial [Nematostella vectensis] |
| XP_625352.1 | 481 | cgd6_120 | eukaryota>alveolata>apicomplexa | Cryptosporidium | parvum | Iowa II disulfide-isomerase, signal peptide plus ER retention motif ER protein [Cryptosporidium parvum Iowa II] |
| XP_015660348.1 | 481 | ABB37_04152 | eukaryota>euglenozoa>kinetoplastida | Leptomonas | pyrrhocoris | protein disulfide isomerase [Leptomonas pyrrhocoris] |
| XP_821173.1 | 481 | Tc00.1047053507611.370 | eukaryota>euglenozoa>kinetoplastida | Trypanosoma | cruzi strain CL Brener | protein disulfide isomerase [Trypanosoma cruzi strain CL Brener] |
| XP_001310020.1 | 481 | TVAG_086290 | eukaryota>parabasalia | Trichomonas | vaginalis G3 | hypothetical protein [Trichomonas vaginalis G3] |
| ORE06497.1 | 481 | BCV72DRAFT_123957 | eukaryota>fungi | Rhizopus | microsporus var. microsporus | thioredoxin-like protein [Rhizopus microsporus var. microsporus] |
| XP_012362681.1 | 480 | PDIA3 | eukaryota>metazoa>chordata>vertebrata | Nomascus | leucogenys | PREDICTED: protein disulfide-isomerase A3 isoform X3 [Nomascus leucogenys] |
| CBY21606.1 | 479 | GSOID_T00009378001 | eukaryota>metazoa>chordata | Oikopleura | dioica | unnamed protein product [Oikopleura dioica] |
| XP_002979092.1 | 479 | SELMODRAFT_110105 | eukaryota>viridiplantae | Selaginella | moellendorffii | hypothetical protein SELMODRAFT_110105 [Selaginella moellendorffii] |
| XP_002993987.1 | 479 | SELMODRAFT_163263 | eukaryota>viridiplantae | Selaginella | moellendorffii | hypothetical protein SELMODRAFT_163263 [Selaginella moellendorffii] |
| KPI89399.1 | 479 | ABL78_1525 | eukaryota>euglenozoa>kinetoplastida | Leptomonas | seymouri | protein disulfide isomerase [Leptomonas seymouri] |
| XP_013757691.1 | 478 | AMSG_05621 | eukaryota>apusozoa>apusomonadidae | Thecamonas | trahens ATCC 50062 | hypothetical protein AMSG_05621 [Thecamonas trahens ATCC 50062] |
| XP_014153070.1 | 478 | SARC_08432 | eukaryota>ichthyosporea | Sphaeroforma | arctica JP610 | hypothetical protein SARC_08432 [Sphaeroforma arctica JP610] |
| XP_002140148.1 | 478 | CMU_028090 | eukaryota>alveolata>apicomplexa | Cryptosporidium | muris RN66 | disulfide isomerase protein [Cryptosporidium muris RN66] |
| XP_002974021.1 | 477 | SELMODRAFT_52354 | eukaryota>viridiplantae | Selaginella | moellendorffii | hypothetical protein SELMODRAFT_52354, partial [Selaginella moellendorffii] |
| XP_001469404.1 | 477 | PDI-2 | eukaryota>euglenozoa>kinetoplastida | Leishmania | infantum JPCM5 | protein disulfide isomerase [Leishmania infantum JPCM5] |
| XP_001687289.1 | 477 | PDI-2 | eukaryota>euglenozoa>kinetoplastida | Leishmania | major strain Friedlin | protein disulfide isomerase [Leishmania major strain Friedlin] |
| XP_003865838.1 | 477 | LDBPK_367280 | eukaryota>euglenozoa>kinetoplastida | Leishmania | donovani | protein disulfide isomerase [Leishmania donovani] |
| XP_002126714.1 | 476 | LOC100179353 | eukaryota>metazoa>chordata | Ciona | intestinalis | PREDICTED: protein disulfide-isomerase A3 [Ciona intestinalis] |
| XP_018285422.1 | 476 | PHYBLDRAFT_19447 | eukaryota>fungi | Phycomyces | blakesleeanus NRRL 1555(-) | hypothetical protein PHYBLDRAFT_19447 [Phycomyces blakesleeanus NRRL 1555(-)] |
| Capa1000007768 | 476 | Capa1000007768 | eukaryota>metazoa>annelida | Capitella | sp1 | >Capa1000007768 |
| EPY37757.1 | 476 | AGDE_06177 | eukaryota>euglenozoa>kinetoplastida | Angomonas | deanei | protein disulfide-isomerase A6 [Angomonas deanei] |
| XP_004256756.1 | 476 | EIN_402890 | eukaryota>amoebozoa>entamoebidae | Entamoeba | invadens IP1 | protein disulfide isomerase, putative [Entamoeba invadens IP1] |
| ELR13716.1 | 476 | ACA1_305150 | eukaryota>amoebozoa>acanthamoebidae | Acanthamoeba | castellanii str. Neff | protein disulfide isomerase associated 4, putative, partial [Acanthamoeba castellanii str. Neff] |
| XP_002678133.1 | 476 | NAEGRDRAFT_57827 | eukaryota>heterolobosea | Naegleria | gruberi strain NEG-M | disulfide isomerase family protein [Naegleria gruberi strain NEG-M] |
| OMJ67705.1 | 475 | SteCoe_35059 | eukaryota>alveolata>ciliophora | Stentor | coeruleus | hypothetical protein SteCoe_35059 [Stentor coeruleus] |
| OMJ86534.1 | 474 | SteCoe_11951 | eukaryota>alveolata>ciliophora | Stentor | coeruleus | hypothetical protein SteCoe_11951 [Stentor coeruleus] |
| OMJ94810.1 | 474 | SteCoe_1994 | eukaryota>alveolata>ciliophora | Stentor | coeruleus | hypothetical protein SteCoe_1994 [Stentor coeruleus] |
| CBY14738.1 | 473 | GSOID_T00009810001 | eukaryota>metazoa>chordata | Oikopleura | dioica | unnamed protein product [Oikopleura dioica] |
| EIE84149.1 | 473 | RO3G_08859 | eukaryota>fungi | Rhizopus | delemar RA 99-880 | hypothetical protein RO3G_08859 [Rhizopus delemar RA 99-880] |
| OMJ67351.1 | 473 | SteCoe_35512 | eukaryota>alveolata>ciliophora | Stentor | coeruleus | hypothetical protein SteCoe_35512 [Stentor coeruleus] |
| OMJ73739.1 | 473 | SteCoe_27509 | eukaryota>alveolata>ciliophora | Stentor | coeruleus | hypothetical protein SteCoe_27509 [Stentor coeruleus] |
| XP_003875055.1 | 473 | LMXM_36_6940 | eukaryota>euglenozoa>kinetoplastida | Leishmania | mexicana MHOM/GT/2001/U1103 | protein disulfide isomerase [Leishmania mexicana MHOM/GT/2001/U1103] |
| OMJ90871.1 | 472 | SteCoe_6630 | eukaryota>alveolata>ciliophora | Stentor | coeruleus | hypothetical protein SteCoe_6630 [Stentor coeruleus] |
| XP_012896346.1 | 472 | GSBLH_T00002347001 | eukaryota>stramenopiles | Blastocystis | hominis | uncharacterized protein [Blastocystis hominis] |
| XP_011132043.1 | 472 | GNI_127040 | eukaryota>alveolata>apicomplexa | Gregarina | niphandrodes | disulfide isomerase [Gregarina niphandrodes] |
| XP_002371293.1 | 471 | TGME49_211680 | eukaryota>alveolata>apicomplexa | Toxoplasma | gondii ME49 | protein disulfide isomerase [Toxoplasma gondii ME49] |
| XP_003880708.1 | 471 | NCLIV_011410 | eukaryota>alveolata>apicomplexa | Neospora | caninum Liverpool | protein disulfide isomerase [Neospora caninum Liverpool] |
| XP_008885290.1 | 471 | HHA_211680 | eukaryota>alveolata>apicomplexa | Hammondia | hammondi | protein disulfide isomerase [Hammondia hammondi] |
| CCW70112.1 | 471 | GSHART1_T00007599001 | eukaryota>euglenozoa>kinetoplastida | Phytomonas | sp. isolate Hart1 | unnamed protein product [Phytomonas sp. isolate Hart1] |
| XP_001327862.1 | 471 | TVAG_209180 | eukaryota>parabasalia | Trichomonas | vaginalis G3 | hypothetical protein [Trichomonas vaginalis G3] |
| OMJ79678.1 | 470 | SteCoe_20242 | eukaryota>alveolata>ciliophora | Stentor | coeruleus | hypothetical protein SteCoe_20242 [Stentor coeruleus] |
| EPY22551.1 | 470 | STCU_08222 | eukaryota>euglenozoa>kinetoplastida | Strigomonas | culicis | protein disulfide-isomerase [Strigomonas culicis] |
| EPY28046.1 | 470 | STCU_05328 | eukaryota>euglenozoa>kinetoplastida | Strigomonas | culicis | protein disulfide-isomerase [Strigomonas culicis] |
| XP_001569341.1 | 470 | LBRM_35_7320 | eukaryota>euglenozoa>kinetoplastida | Leishmania | braziliensis MHOM/BR/75/M2904 | protein disulfide isomerase [Leishmania braziliensis MHOM/BR/75/M2904] |
| XP_010703642.1 | 470 | PDI-2 | eukaryota>euglenozoa>kinetoplastida | Leishmania | panamensis | protein disulfide isomerase [Leishmania panamensis] |
| XP_012647399.1 | 468 | BBM_III670 | eukaryota>alveolata>apicomplexa | Babesia | microti strain RI | uncharacterized protein [Babesia microti strain RI] |
| Crev1000000943 | 467 | Crev1000000943 | eukaryota>fungi>kickxellomycotina | Coemansia | reversa | >Crev1000000943 |
| XP_001747756.1 | 466 | MONBRDRAFT_27243 | eukaryota>choanoflagellida | Monosiga | brevicollis MX1 | hypothetical protein [Monosiga brevicollis MX1] |
| XP_002111036.1 | 465 | TRIADDRAFT_63251 | eukaryota>metazoa>placozoa | Trichoplax | adhaerens | hypothetical protein TRIADDRAFT_63251 [Trichoplax adhaerens] |
| EPZ33203.1 | 464 | O9G_001172 | eukaryota>fungi | Rozella | allomycis CSF55 | Thioredoxin-like fold domain-containing protein [Rozella allomycis CSF55] |
| XP_006505513.1 | 464 | AI987846,Cai,ERp-72,Erp72 | eukaryota>metazoa>chordata>vertebrata | Mus | musculus | PREDICTED: protein disulfide-isomerase A4 isoform X1 [Mus musculus] |
| CEL96760.1 | 464 | Vbra_12065 | eukaryota>alveolata | Vitrella | brassicaformis CCMP3155 | unnamed protein product [Vitrella brassicaformis CCMP3155] |
| XP_001609398.1 | 463 | 21.m03010 | eukaryota>alveolata>apicomplexa | Babesia | bovis T2Bo | protein disulfide-isomerase [Babesia bovis T2Bo] |
| XP_004832266.1 | 460 | BEWA_013730 | eukaryota>alveolata>apicomplexa | Theileria | equi | protein disulfide isomerase, putative [Theileria equi] |
| Bnat1000012902 | 458 | Bnat1000012902 | eukaryota>rhizaria | Bigelowiella | natans | >Bnat1000012902 |
| EPZ33204.1 | 456 | O9G_001173 | eukaryota>fungi | Rozella | allomycis CSF55 | Thioredoxin-like fold domain-containing protein [Rozella allomycis CSF55] |
| XP_003293872.1 | 456 | DICPUDRAFT_58708 | eukaryota>amoebozoa>mycetozoa>dictyostelida | Dictyostelium | purpureum | hypothetical protein DICPUDRAFT_58708 [Dictyostelium purpureum] |
| XP_017327519.1 | 455 | LOC108267688 | eukaryota>metazoa>chordata>vertebrata>actinopterygii | Ictalurus | punctatus | PREDICTED: protein disulfide-isomerase A3-like isoform X2 [Ictalurus punctatus] |
| XP_005559436.1 | 453 | PDIA3 | eukaryota>metazoa>chordata>vertebrata | Macaca | fascicularis | PREDICTED: protein disulfide-isomerase A3 isoform X2 [Macaca fascicularis] |
| XP_013243228.1 | 453 | K437DRAFT_120456 | eukaryota>fungi>basidiomycota | Tilletiaria | anomala UBC 951 | protein disulfide isomerase [Tilletiaria anomala UBC 951] |
| XP_004092689.1 | 451 | P4HB | eukaryota>metazoa>chordata>vertebrata | Nomascus | leucogenys | PREDICTED: protein disulfide-isomerase isoform X2 [Nomascus leucogenys] |
| XP_014622056.1 | 451 | LOC100788702 | eukaryota>viridiplantae | Glycine | max | PREDICTED: protein disulfide-isomerase-like isoform X2 [Glycine max] |
| XP_001318330.1 | 451 | TVAG_349600 | eukaryota>parabasalia | Trichomonas | vaginalis G3 | hypothetical protein [Trichomonas vaginalis G3] |
| ESU44748.1 | 449 | GSB_9413 | eukaryota>diplomonadida | Giardia | intestinalis | Protein disulfide isomerase [Giardia intestinalis] |
| XP_001707782.1 | 449 | GL50803_9413 | eukaryota>diplomonadida | Giardia | lamblia ATCC 50803 | Protein disulfide isomerase PDI2 [Giardia lamblia ATCC 50803] |
| OQV19186.1 | 442 | BV898_06823 | eukaryota>metazoa | Hypsibius | dujardini | Protein disulfide-isomerase A3 [Hypsibius dujardini] |
| Smar1000000668 | 441 | Smar1000000668 | eukaryota>metazoa>arthropoda>myriapoda | Strigamia | maritima | >Smar1000000668 |
| XP_002602037.1 | 440 | BRAFLDRAFT_59159 | eukaryota>metazoa>chordata | Branchiostoma | floridae | hypothetical protein BRAFLDRAFT_59159 [Branchiostoma floridae] |
| NP_872239.1 | 437 | pdi-2 | eukaryota>metazoa>nematoda | Caenorhabditis | elegans | Protein Disulfide Isomerase [Caenorhabditis elegans] |
| XP_001744423.1 | 436 | MONBRDRAFT_784 | eukaryota>choanoflagellida | Monosiga | brevicollis MX1 | hypothetical protein, partial [Monosiga brevicollis MX1] |
| XP_016479813.1 | 420 | LOC107801043 | eukaryota>viridiplantae | Nicotiana | tabacum | PREDICTED: protein disulfide-isomerase-like isoform X4 [Nicotiana tabacum] |
| XP_016514999.1 | 420 | LOC107831717 | eukaryota>viridiplantae | Nicotiana | tabacum | PREDICTED: protein disulfide-isomerase-like isoform X2 [Nicotiana tabacum] |
| ORX80531.1 | 416 | K493DRAFT_389936 | eukaryota>fungi | Basidiobolus | meristosporus CBS 931.73 | hypothetical protein K493DRAFT_389936 [Basidiobolus meristosporus CBS 931.73] |
| XP_011677933.1 | 414 | LOC105436389 | eukaryota>metazoa>echinodermata | Strongylocentrotus | purpuratus | PREDICTED: protein disulfide-isomerase 2-like [Strongylocentrotus purpuratus] |
| EST49530.1 | 372 | SS50377_10134 | eukaryota>diplomonadida | Spironucleus | salmonicida | Protein disulfide isomerase [Spironucleus salmonicida] |
| XP_001710228.1 | 354 | GL50803_103713 | eukaryota>diplomonadida | Giardia | lamblia ATCC 50803 | Protein disulfide isomerase PDI4 [Giardia lamblia ATCC 50803] |

C. SERCA

| | | | | | | |
|---|---|---|---|---|---|---|
| OMH79210.1 | 1617 | AX774_g7385 | eukaryota>fungi | Sancudomyces | culisetae | Calcium-transporting ATPase 1 [Sancudomyces culisetae] |
| XP_004363878.1 | 1448 | CAOG_03039 | eukaryota>ichthyosporea | Capsaspora | owczarzaki ATCC 30864 | Na+ P-type-ATPase [Capsaspora owczarzaki ATCC 30864] |
| XP_680933.1 | 1413 | AN7664.2 | eukaryota>fungi>ascomycota | Aspergillus | nidulans FGSC A4 | hypothetical protein AN7664.2 [Aspergillus nidulans FGSC A4] |
| EWM23830.1 | 1342 | Naga_100362g4 | eukaryota>stramenopiles | Nannochloropsis | gaditana | p-type had subfamily ic [Nannochloropsis gaditana] |
| KFH73317.1 | 1340 | MVEG_00534 | eukaryota>fungi | Mortierella | verticillata NRRL 6337 | hypothetical protein MVEG_00534 [Mortierella verticillata NRRL 6337] |
| XP_021883455.1 | 1281 | BCR41DRAFT_380461 | eukaryota>fungi | Lobosporangium | transversale | hypothetical protein BCR41DRAFT_380461 [Lobosporangium transversale] |
| EUB59093.1 | 1262 | EGR_06085 | eukaryota>metazoa | Echinococcus | granulosus | Sarcoplasmic/endoplasmic reticulum calcium ATPase 3 [Echinococcus granulosus] |
| OEU14943.1 | 1241 | FRACYDRAFT_188142 | eukaryota>stramenopiles | Fragilariopsis | cylindrus CCMP1102 | calcium ATPase [Fragilariopsis cylindrus CCMP1102] |
| XP_009497389.1 | 1234 | H696_05252 | eukaryota | Fonticula | alba | hypothetical protein H696_05252 [Fonticula alba] |
| XP_015664582.1 | 1226 | ABB37_00393 | eukaryota>euglenozoa>kinetoplastida | Leptomonas | pyrrhocoris | putative mitochondrial calcium motive p-type ATPase [Leptomonas pyrrhocoris] |
| XP_003879203.1 | 1225 | LMXM_34_2080 | eukaryota>euglenozoa>kinetoplastida | Leishmania | mexicana MHOM/GT/2001/U1103 | putative calcium motive p-type ATPase [Leishmania mexicana MHOM/GT/2001/U1103] |
| EPB85147.1 | 1218 | HMPREF1544_08119 | eukaryota>fungi | Mucor | circinelloides f. circinelloides 1006PhL | hypothetical protein HMPREF1544_08119 [Mucor circinelloides f. circinelloides 1006PhL] |
| KPI88812.1 | 1215 | ABL78_2071 | eukaryota>euglenozoa>kinetoplastida | Leptomonas | seymouri | putative Calcium motive p-type ATPase [Leptomonas seymouri] |
| CEM18264.1 | 1212 | Vbra_16176 | eukaryota>alveolata | Vitrella | brassicaformis CCMP3155 | unnamed protein product [Vitrella brassicaformis CCMP3155] |
| EPZ33470.1 | 1204 | O9G_001221 | eukaryota>fungi | Rozella | allomycis CSF55 | P-type ATPase, A domain-containing protein [Rozella allomycis CSF55] |
| EXX59744.1 | 1199 | RirG_186260 | eukaryota>fungi | Rhizophagus | irregularis DAOM 197198w | Ca(2+)/Mn(2+)-transporting P-type ATPase PMR1 [Rhizophagus irregularis DAOM 197198w] |
| CEP01863.1 | 1198 | PBRA_008806 | eukaryota>rhizaria>cercozoa>plasmodiophorida | Plasmodiophora | brassicae | hypothetical protein PBRA_008806 [Plasmodiophora brassicae] |
| ORE03465.1 | 1198 | BCV72DRAFT_315304 | eukaryota>fungi | Rhizopus | microsporus var. microsporus | calcium ATPase [Rhizopus microsporus var. microsporus] |
| XP_016609006.1 | 1194 | SPPG_04066 | eukaryota>fungi>chytridiomycota | Spizellomyces | punctatus DAOM BR117 | potassium/sodium efflux P-type ATPase, fungal-type [Spizellomyces punctatus DAOM BR117] |
| Uram1000006337 | 1190 | Uram1000006337 | eukaryota>fungi>mucoromycotina | Umbelopsis | ramanniana | >Uram1000006337 |
| ORY43529.1 | 1189 | LY90DRAFT_385125 | eukaryota>fungi>chytridiomycota | Neocallimastix | californiae | calcium ATPase [Neocallimastix californiae] |
| ORY06544.1 | 1189 | LY90DRAFT_392966 | eukaryota>fungi>chytridiomycota | Neocallimastix | californiae | calcium ATPase [Neocallimastix californiae] |
| CEM04548.1 | 1187 | Vbra_8653 | eukaryota>alveolata | Vitrella | brassicaformis CCMP3155 | unnamed protein product [Vitrella brassicaformis CCMP3155] |
| Uram1000001495 | 1183 | Uram1000001495 | eukaryota>fungi>mucoromycotina | Umbelopsis | ramanniana | >Uram1000001495 |
| XP_001228662.1 | 1182 | CHGG_02146 | eukaryota>fungi>ascomycota | Chaetomium | globosum CBS 148.51 | hypothetical protein CHGG_02146 [Chaetomium globosum CBS 148.51] |
| GAQ82796.1 | 1181 | KFL_001240270 | eukaryota>viridiplantae | Klebsormidium | nitens | Ca2+ transporting ATPase [Klebsormidium nitens] |
| EWM25701.1 | 1173 | Naga_100222g2 | eukaryota>stramenopiles | Nannochloropsis | gaditana | putative serca-type calcium atpase [Nannochloropsis gaditana] |
| KFH70534.1 | 1171 | MVEG_03384 | eukaryota>fungi | Mortierella | verticillata NRRL 6337 | hypothetical protein MVEG_03384 [Mortierella verticillata NRRL 6337] |
| XP_021869883.1 | 1170 | BD324DRAFT_631378 | eukaryota>fungi>basidiomycota | Kockovaella | imperatae | hypothetical protein BD324DRAFT_631378 [Kockovaella imperatae] |
| XP_001221054.1 | 1162 | CHGG_01833 | eukaryota>fungi>ascomycota | Chaetomium | globosum CBS 148.51 | hypothetical protein CHGG_01833 [Chaetomium globosum CBS 148.51] |
| XP_009167367.1 | 1151 | T265_13488 | eukaryota>metazoa | Opisthorchis | viverrini | hypothetical protein T265_13488, partial [Opisthorchis viverrini] |
| XP_018190425.1 | 1143 | L228DRAFT_243609 | eukaryota>fungi>ascomycota | Xylona | heveae TC161 | Na+-exporting ATPase [Xylona heveae TC161] |
| CAF98515.1 | 1141 | GSTEN:00016251:G:001 | eukaryota>metazoa>chordata>vertebrata>actinopterygii | Tetraodon | nigroviridis | unnamed protein product, partial [Tetraodon nigroviridis] |
| XP_004184491.1 | 1137 | EIN_081770 | eukaryota>amoebozoa>entamoebidae | Entamoeba | invadens IP1 | cation-transporting ATPase pacL, putative [Entamoeba invadens IP1] |
| XP_008855949.1 | 1137 | ENU1_046870 | eukaryota>amoebozoa>entamoebidae | Entamoeba | nuttalli P19 | calcium-transporting P-type ATPase, putative [Entamoeba nuttalli P19] |
| XP_001735630.1 | 1137 | EDI_097030 | eukaryota>amoebozoa>entamoebidae | Entamoeba | dispar SAW760 | cation-transporting ATPase pacL [Entamoeba dispar SAW760] |
| GAT95225.1 | 1137 | CL6EHI_027710 | eukaryota>amoebozoa>entamoebidae | Entamoeba | histolytica | calcium-transporting p-type ATPase putative [Entamoeba histolytica] |
| XP_002141105.1 | 1134 | CMU_014320 | eukaryota>alveolata>apicomplexa | Cryptosporidium | muris RN66 | calcium-translocating P-type ATPase, SERCA-type family protein [Cryptosporidium muris RN66] |
| Lhya1000007956 | 1134 | Lhya1000007956 | eukaryota>fungi>mucoromycotina | Lichtheimia | hyalospora | >Lhya1000007956 |
| KFH73316.1 | 1133 | MVEG_00534 | eukaryota>fungi | Mortierella | verticillata NRRL 6337 | hypothetical protein, variant [Mortierella verticillata NRRL 6337] |
| XP_001224140.1 | 1130 | CHGG_04926 | eukaryota>fungi>ascomycota | Chaetomium | globosum CBS 148.51 | hypothetical protein CHGG_04926 [Chaetomium globosum CBS 148.51] |
| XP_628574.1 | 1129 | cgd7_4190 | eukaryota>alveolata>apicomplexa | Cryptosporidium | parvum Iowa II | cation-transporting P-type ATPase with 11 or more transmembrane domains [Cryptosporidium parvum Iowa II] |
| KNE62172.1 | 1126 | AMAG_07418 | eukaryota>fungi | Allomyces | macrogynus ATCC 38327 | HAD ATPase, P-type, family IC [Allomyces macrogynus ATCC 38327] |
| XP_011386140.1 | 1125 | UMAG_00204 | eukaryota>fungi>basidiomycota | Ustilago | maydis 521 | hypothetical protein UMAG_00204 [Ustilago maydis 521] |
| XP_002953236.1 | 1123 | VOLCADRAFT_105844 | eukaryota>viridiplantae>chlorophyta | Volvox | carteri f. nagariensis | hypothetical protein VOLCADRAFT_105844 [Volvox carteri f. nagariensis] |
| XP_956354.1 | 1121 | ena-1 | eukaryota>fungi>ascomycota | Neurospora | crassa OR74A | calcium-transporting ATPase 3 [Neurospora crassa OR74A] |
| XP_005847889.1 | 1119 | CHLNCDRAFT_145247 | eukaryota>viridiplantae>chlorophyta | Chlorella | variabilis | hypothetical protein CHLNCDRAFT_145247 [Chlorella variabilis] |
| XP_012048517.1 | 1117 | CNAG_05135 | eukaryota>fungi>basidiomycota | Cryptococcus | neoformans var. grubii H99 | Ca2 -transporting ATPase [Cryptococcus neoformans var. grubii H99] |
| XP_011397223.1 | 1117 | F751_3053 | eukaryota>viridiplantae>chlorophyta | Auxenochlorella | protothecoides | calcium-transporting ATPase 1, endoplasmic reticulum-type [Auxenochlorella protothecoides] |
| XP_018278279.1 | 1115 | CC85DRAFT_328798 | eukaryota>fungi>basidiomycota | Cutaneotrichosporon | oleaginosum | potassium/sodium eff [Cutaneotrichosporon oleaginosum] |
| XP_011522183.1 | 1114 | SERCA3 | eukaryota>metazoa>chordata>vertebrata | Homo | sapiens | sarcoplasmic/endoplasmic reticulum calcium ATPase 3 isoform X1 [Homo sapiens] |
| XP_021869351.1 | 1112 | BD324DRAFT_634005 | eukaryota>fungi>basidiomycota | Kockovaella | imperatae | putative calcium-transporting ATPase 3 [Kockovaella imperatae] |
| XP_013241736.1 | 1111 | K437DRAFT_172527 | eukaryota>fungi>basidiomycota | Tilletiaria | anomala UBC 951 | calcium-transporting P [Tilletiaria anomala UBC 951] |
| XP_001469025.1 | 1109 | LINJ_35_2080 | eukaryota>euglenozoa>kinetoplastida | Leishmania | infantum JPCM5 | putative calcium motive p-type ATPase [Leishmania infantum JPCM5] |
| XP_003722601.1 | 1109 | LMJF_35_2080 | eukaryota>euglenozoa>kinetoplastida | Leishmania | major strain Friedlin | putative calcium motive p-type ATPase [Leishmania major strain Friedlin] |
| XP_003864789.1 | 1109 | LDBPK_352080 | eukaryota>euglenozoa>kinetoplastida | Leishmania | donovani | calcium motive p-type ATPase, putative [Leishmania donovani] |
| XP_962099.1 | 1109 | NCU07966 | eukaryota>fungi>ascomycota | Neurospora | crassa OR74A | calcium-transporting ATPase 3 [Neurospora crassa OR74A] |
| XP_007328195.1 | 1107 | AGABI1DRAFT_72076 | eukaryota>fungi>basidiomycota | Agaricus | bisporus var. burnettii JB137-S8 | hypothetical protein AGABI1DRAFT_72076 [Agaricus bisporus var. burnettii JB137-S8] |
| XP_013243622.1 | 1106 | K437DRAFT_256149 | eukaryota>fungi>basidiomycota | Tilletiaria | anomala UBC 951 | putative ENA2-plasma membrane P-type ATPase [Tilletiaria anomala UBC 951] |
| XP_011133287.1 | 1105 | GNI_170150 | eukaryota>alveolata>apicomplexa | Gregarina | niphandrodes | calcium-transporting ATPase 1, endoplasmic reticulum-type protein [Gregarina niphandrodes] |
| CAG02473.1 | 1105 | GSTEN:00021597:G:001 | eukaryota>metazoa>chordata>vertebrata>actinopterygii | Tetraodon | nigroviridis | unnamed protein product, partial [Tetraodon nigroviridis] |
| XP_001568308.1 | 1104 | LBRM_34_1990 | eukaryota>euglenozoa>kinetoplastida | Leishmania | braziliensis MHOM/BR/75/M2904 | putative calcium motive p-type ATPase [Leishmania braziliensis MHOM/BR/75/M2904] |
| XP_010702617.1 | 1104 | LPMP_341910 | eukaryota>euglenozoa>kinetoplastida | Leishmania | panamensis | calcium motive p-type ATPase, putative [Leishmania panamensis] |
| XP_005652084.1 | 1103 | COCSUDRAFT_26789 | eukaryota>viridiplantae>chlorophyta | Coccomyxa | subellipsoidea C-169 | endoplasmic reticulum-type calcium-transporting ATPase [Coccomyxa subellipsoidea C-169] |
| EPB92068.1 | 1100 | HMPREF1544_01132 | eukaryota>fungi | Mucor | circinelloides f. circinelloides 1006PhL | hypothetical protein HMPREF1544_01132 [Mucor circinelloides f. circinelloides 1006PhL] |
| XP_011388144.1 | 1098 | UMAG_15043 | eukaryota>fungi>basidiomycota | Ustilago | maydis 521 | putative plasma membrane P-type ATPase [Ustilago maydis 521] |
| KNE68426.1 | 1097 | AMAG_13080 | eukaryota>fungi | Allomyces | macrogynus ATCC 38327 | calcium-translocating P-type ATPase, SERCA-type [Allomyces macrogynus ATCC 38327] |
| XP_002367953.1 | 1093 | TGME49_230420 | eukaryota>alveolata>apicomplexa | Toxoplasma | gondii ME49 | sarco/endoplasmic reticulum Ca2+-ATPase [Toxoplasma gondii ME49] |
| XP_008889593.1 | 1093 | HHA_230420 | eukaryota>alveolata>apicomplexa | Hammondia | hammondi | sarco/endoplasmic reticulum Ca2+-ATPase [Hammondia hammondi] |
| XP_001541567.1 | 1093 | HCAG_03665 | eukaryota>fungi>ascomycota | Histoplasma | capsulatum NAm1 | calcium-transporting ATPase 1 [Histoplasma capsulatum NAm1] |
| XP_959558.3 | 1092 | ena-2 | eukaryota>fungi>ascomycota | Neurospora | crassa OR74A | Na or K P-type ATPase [Neurospora crassa OR74A] |
| ORZ31211.1 | 1092 | BCR44DRAFT_119611 | eukaryota>fungi | Catenaria | anguillulae PL171 | hypothetical protein BCR44DRAFT_119611 [Catenaria anguillulae PL171] |
| NP_010325.1 | 1091 | HOR6,PMR2 | eukaryota>fungi>ascomycota | Saccharomyces | cerevisiae S288C | Na(+)/Li(+)-exporting P-type ATPase ENA1 [Saccharomyces cerevisiae S288C] |
| NP_010324.3 | 1091 | ENA2 | eukaryota>fungi>ascomycota | Saccharomyces | cerevisiae S288C | Na(+)-exporting P-type ATPase ENA2 [Saccharomyces cerevisiae S288C] |
| NP_010323.3 | 1091 | ENA5 | eukaryota>fungi>ascomycota | Saccharomyces | cerevisiae S288C | putative Na(+)-exporting P-type ATPase ENA5 [Saccharomyces cerevisiae S288C] |
| XP_012046341.1 | 1091 | CNAG_00531 | eukaryota>fungi>basidiomycota | Cryptococcus | neoformans var. grubii H99 | potassium/sodium efflux P-type ATPase, fungal-type [Cryptococcus neoformans var. grubii H99] |
| XP_005707399.1 | 1089 | Gasu_18920 | eukaryota>rhodophyta | Galdieria | sulphuraria | calcium-transporting P-type ATPase [Galdieria sulphuraria] |
| XP_001219310.1 | 1088 | CHGG_00089 | eukaryota>fungi>ascomycota | Chaetomium | globosum CBS 148.51 | hypothetical protein CHGG_00089 [Chaetomium globosum CBS 148.51] |
| XP_017992933.1 | 1087 | Malapachy_2261 | eukaryota>fungi>basidiomycota | Malassezia | pachydermatis | ena2-plasma membrane p-type atpase [Malassezia pachydermatis] |
| XP_001832079.1 | 1086 | CC1G_07450 | eukaryota>fungi>basidiomycota | Coprinopsis | cinerea okayama7#130 | calcium-transporting ATPase 3 [Coprinopsis cinerea okayama7#130] |
| XP_006461327.1 | 1086 | AGABI2DRAFT_185605 | eukaryota>fungi>basidiomycota | Agaricus | bisporus var. bisporus H97 | hypothetical protein AGABI2DRAFT_185605 [Agaricus bisporus var. bisporus H97] |
| Pisp1000002294 | 1085 | Pisp1000002294 | eukaryota>fungi>neocallimastigomycota | Piromyces | sp | >Pisp1000002294 |
| XP_001837092.1 | 1084 | CC1G_00228 | eukaryota>fungi>basidiomycota | Coprinopsis | cinerea okayama7#130 | sodium transport ATPase [Coprinopsis cinerea okayama7#130] |
| Bcir1000009373 | 1084 | Bcir1000009373 | eukaryota>fungi>mucoromycotina | Backusella | circina | >Bcir1000009373 |
| XP_011522184.1 | 1084 | SERCA3 | eukaryota>metazoa>chordata>vertebrata | Homo | sapiens | sarcoplasmic/endoplasmic reticulum calcium ATPase 3 isoform X2 [Homo sapiens] |
| CEP01171.1 | 1083 | PBRA_008483 | eukaryota>rhizaria>cercozoa>plasmodiophorida | Plasmodiophora | brassicae | hypothetical protein PBRA_008483 [Plasmodiophora brassicae] |
| XP_454607.1 | 1082 | KLLA0_E14587g | eukaryota>fungi>ascomycota | Kluyveromyces | lactis | uncharacterized protein KLLA0_E14587g [Kluyveromyces lactis] |
| XP_456007.1 | 1082 | KLLA0_F20658g | eukaryota>fungi>ascomycota | Kluyveromyces | lactis | uncharacterized protein KLLA0_F20658g [Kluyveromyces lactis] |
| XP_019015284.1 | 1080 | PICMEDRAFT_18434 | eukaryota>fungi>ascomycota | Pichia | membranifaciens NRRL Y-2026 | hypothetical protein PICMEDRAFT_18434 [Pichia membranifaciens NRRL Y-2026] |
| XP_003883339.1 | 1079 | NCLIV_030940 | eukaryota>alveolata>apicomplexa | Neospora | caninum Liverpool | hypothetical protein NCLIV_030940 [Neospora caninum Liverpool] |
| XP_014537390.1 | 1079 | PDIP_16990 | eukaryota>fungi>ascomycota | Penicillium | digitatum Pd1 | Sodium P-type ATPase, putative [Penicillium digitatum Pd1] |
| XP_003062749.1 | 1079 | MICPUCDRAFT_36018 | eukaryota>viridiplantae>chlorophyta | Micromonas | pusilla CCMP1545 | p-type ATPase superfamily [Micromonas pusilla CCMP1545] |
| XP_013228780.1 | 1078 | ETH_00026655 | eukaryota>alveolata>apicomplexa | Eimeria | tenella | calcium-transporting ATPase, putative [Eimeria tenella] |
| XP_018738072.1 | 1078 | ENA5 | eukaryota>fungi>ascomycota | Sugiyamaella | lignohabitans | putative Na(+)-exporting P-type ATPase ENA5 [Sugiyamaella lignohabitans] |
| XP_499639.1 | 1078 | YALI0_A01023g | eukaryota>fungi>ascomycota | Yarrowia | lipolytica CLIB122 | YALI0A01023p [Yarrowia lipolytica CLIB122] |
| XP_017318576.1 | 1077 | LOC108262771 | eukaryota>metazoa>chordata>vertebrata>actinopterygii | Ictalurus | punctatus | PREDICTED: sarcoplasmic/endoplasmic reticulum calcium ATPase 2-like isoform X1 [Ictalurus punctatus] |
| XP_001830557.1 | 1076 | CC1G_06823 | eukaryota>fungi>basidiomycota | Coprinopsis | cinerea okayama7#130 | calcium-transporting ATPase [Coprinopsis cinerea okayama7#130] |
| XP_004829101.1 | 1075 | BEWA_022830 | eukaryota>alveolata>apicomplexa | Theileria | equi strain WA | p-type ATPase family member protein [Theileria equi strain WA] |
| XP_011123014.1 | 1075 | AOL_s00080g337 | eukaryota>fungi>ascomycota | Arthrobotrys | oligospora ATCC 24927 | hypothetical protein AOL_s00080g337 [Arthrobotrys oligospora ATCC 24927] |
| XP_005537020.1 | 1075 | CYME_CMM114C | eukaryota>rhodophyta | Cyanidioschyzon | merolae strain 10D | calcium-transporting ATPase [Cyanidioschyzon merolae strain 10D] |
| XP_001222926.1 | 1074 | CHGG_03712 | eukaryota>fungi>ascomycota | Chaetomium | globosum CBS 148.51 | hypothetical protein CHGG_03712 [Chaetomium globosum CBS 148.51] |
| XP_013432376.1 | 1073 | M436DRAFT_77535 | eukaryota>fungi>ascomycota | Aureobasidium | namibiae CBS 147.97 | Na(+)/Li(+)-exporting P-type ATPase [Aureobasidium namibiae CBS 147.97] |
| Lhya1000000800 | 1071 | Lhya1000000800 | eukaryota>fungi>mucoromycotina | Lichtheimia | hyalospora | >Lhya1000000800 |
| Chet1000007197 | 1070 | Chet1000007197 | eukaryota>fungi>ascomycota | Cochliobolus | heterostrophus | >Chet1000007197 |
| XP_019023864.1 | 1070 | SAICODRAFT_57537 | eukaryota>fungi>ascomycota | Saitoella | complicata NRRL Y-17804 | putative sodium P-type ATPase [Saitoella complicata NRRL Y-17804] |
| XP_005646370.1 | 1070 | COCSUDRAFT_66699 | eukaryota>viridiplantae>chlorophyta | Coccomyxa | subellipsoidea C-169 | calcium ATPase [Coccomyxa subellipsoidea C-169] |
| Chet1000000275 | 1069 | Chet1000000275 | eukaryota>fungi>ascomycota | Cochliobolus | heterostrophus | >Chet1000000275 |
| XP_011522186.1 | 1069 | SERCA3 | eukaryota>metazoa>chordata>vertebrata | Homo | sapiens | sarcoplasmic/endoplasmic reticulum calcium ATPase 3 isoform X3 [Homo sapiens] |
| XP_001700727.1 | 1069 | CHLREDRAFT_77047 | eukaryota>viridiplantae>chlorophyta | Chlamydomonas | reinhardtii | calcium-transporting ATPase, endoplasmic reticulum-type [Chlamydomonas reinhardtii] |
| EPB83742.1 | 1067 | HMPREF1544_09483 | eukaryota>fungi | Mucor | circinelloides f. circinelloides 1006PhL | hypothetical protein HMPREF1544_09483 [Mucor circinelloides f. circinelloides 1006PhL] |
| XP_719032.1 | 1067 | ENA2 | eukaryota>fungi>ascomycota | Candida | albicans SC5314 | Na(+)-exporting P-type ATPase [Candida albicans SC5314] |
| CDS42374.1 | 1067 | EmuJ_001008200 | eukaryota>metazoa | Echinococcus | multilocularis | calcium transporting atpase [Echinococcus multilocularis] |
| XP_664246.1 | 1066 | AN6642.2 | eukaryota>fungi>ascomycota | Aspergillus | nidulans FGSC A4 | hypothetical protein AN6642.2 [Aspergillus nidulans FGSC A4] |
| Lhya1000005955 | 1066 | Lhya1000005955 | eukaryota>fungi>mucoromycotina | Lichtheimia | hyalospora | >Lhya1000005955 |
| XP_001896249.1 | 1065 | Bm1_23960 | eukaryota>metazoa>nematoda | Brugia | malayi | Calcium-transporting ATPase sarcoplasmic/endoplasmic reticulum type [Brugia malayi] |
| XP_002958852.1 | 1065 | VOLCADRAFT_84661 | eukaryota>viridiplantae>chlorophyta | Volvox | carteri f. nagariensis | hypothetical protein VOLCADRAFT_84661 [Volvox carteri f. nagariensis] |
| XP_002277306.1 | 1063 | LOC100248326 | eukaryota>viridiplantae | Vitis | vinifera | PREDICTED: calcium-transporting ATPase 4, endoplasmic reticulum-type [Vitis vinifera] |
| XP_019077314.1 | 1063 | LOC100248326 | eukaryota>viridiplantae | Vitis | vinifera | PREDICTED: calcium-transporting ATPase 4, endoplasmic reticulum-type [Vitis vinifera] |
| XP_680733.1 | 1062 | AN7464.2 | eukaryota>fungi>ascomycota | Aspergillus | nidulans FGSC A4 | hypothetical protein AN7464.2 [Aspergillus nidulans FGSC A4] |
| XP_659232.1 | 1062 | AN1628.2 | eukaryota>fungi>ascomycota | Aspergillus | nidulans FGSC A4 | hypothetical protein AN1628.2 [Aspergillus nidulans FGSC A4] |
| XP_016880181.1 | 1062 | SERCA3 | eukaryota>metazoa>chordata>vertebrata | Homo | sapiens | sarcoplasmic/endoplasmic reticulum calcium ATPase 3 isoform X4 [Homo sapiens] |
| NP_001262220.1 | 1062 | anon-WO0118547.249,CG14449,CG32451,CG7651,Dmel\CG32451,SPoCK | eukaryota>metazoa>hexapoda | Drosophila | melanogaster | secretory pathway calcium atpase, isoform G [Drosophila melanogaster] |
| GAQ82464.1 | 1062 | KFL_001120080 | eukaryota>viridiplantae | Klebsormidium | nitens | Ca2+ transporting ATPase [Klebsormidium nitens] |
| XP_016457472.1 | 1062 | LOC107781299 | eukaryota>viridiplantae | Nicotiana | tabacum | PREDICTED: calcium-transporting ATPase 4, endoplasmic reticulum-type-like [Nicotiana tabacum] |
| XP_016486962.1 | 1062 | LOC107807149 | eukaryota>viridiplantae | Nicotiana | tabacum | PREDICTED: calcium-transporting ATPase 4, endoplasmic reticulum-type-like [Nicotiana tabacum] |
| XP_008660272.1 | 1062 | LOC103639290 | eukaryota>viridiplantae | Zea | mays | calcium-transporting ATPase 4, endoplasmic reticulum-type [Zea mays] |
| XP_013423194.1 | 1061 | M436DRAFT_85848 | eukaryota>fungi>ascomycota | Aureobasidium | namibiae CBS 147.97 | ENA P-type ATPase 2 [Aureobasidium namibiae CBS 147.97] |
| XP_011610806.1 | 1061 | LOC101071061 | eukaryota>metazoa>chordata>vertebrata>actinopterygii | Takifugu | rubripes | PREDICTED: sarcoplasmic/endoplasmic reticulum calcium ATPase 2-like [Takifugu rubripes] |
| NP_001321027.1 | 1061 | ATECA4,endomembrane-type,CA-ATPase,4,F24B9.24,F24B9_24 | eukaryota>viridiplantae | Arabidopsis | thaliana | endomembrane-type CA-ATPase 4 [Arabidopsis thaliana] |
| NP_172246.3 | 1061 | ATECA4,endomembrane-type,CA-ATPase,4,F24B9.24,F24B9_24 | eukaryota>viridiplantae | Arabidopsis | thaliana | endomembrane-type CA-ATPase 4 [Arabidopsis thaliana] |
| NP_172259.1 | 1061 | ACA3,ARABIDOPSIS,THALIANA,ER-TYPE,CA2+-ATPASE,1,ATECA1,ER-TYPE,CA2+-ATPASE,ER-type,Ca2+-ATPase,1,F24B9.9,F24B9_9 | eukaryota>viridiplantae | Arabidopsis | thaliana | ER-type Ca2+-ATPase 1 [Arabidopsis thaliana] |
| XP_006363343.1 | 1061 | LOC102605714 | eukaryota>viridiplantae | Solanum | tuberosum | PREDICTED: calcium-transporting ATPase 4, endoplasmic reticulum-type-like [Solanum tuberosum] |
| XP_002284552.1 | 1061 | LOC100243325 | eukaryota>viridiplantae | Vitis | vinifera | PREDICTED: calcium-transporting ATPase 4, endoplasmic reticulum-type [Vitis vinifera] |
| XP_010651081.1 | 1061 | LOC100243325 | eukaryota>viridiplantae | Vitis | vinifera | PREDICTED: calcium-transporting ATPase 4, endoplasmic reticulum-type [Vitis vinifera] |
| XP_014530613.1 | 1060 | PDIP_87250 | eukaryota>fungi>ascomycota | Penicillium | digitatum Pd1 | Calcium/mangenese P-type ATPase, putative [Penicillium digitatum Pd1] |
| XP_017318577.1 | 1060 | LOC108262771 | eukaryota>metazoa>chordata>vertebrata>actinopterygii | Ictalurus | punctatus | PREDICTED: sarcoplasmic/endoplasmic reticulum calcium ATPase 2-like isoform X2 [Ictalurus punctatus] |
| XP_003554341.1 | 1060 | LOC100788807 | eukaryota>viridiplantae | Glycine | max | PREDICTED: calcium-transporting ATPase 4, endoplasmic reticulum-type [Glycine max] |
| XP_003521357.1 | 1060 | LOC100812151 | eukaryota>viridiplantae | Glycine | max | PREDICTED: calcium-transporting ATPase 1, endoplasmic reticulum-type-like [Glycine max] |
| XP_001759652.1 | 1060 | PHYPADRAFT_179791 | eukaryota>viridiplantae | Physcomitrella | patens | predicted protein [Physcomitrella patens] |
| ORX90715.1 | 1060 | K493DRAFT_229839 | eukaryota>fungi | Basidiobolus | meristosporus CBS 931.73 | calcium ATPase [Basidiobolus meristosporus CBS 931.73] |
| XP_005781031.1 | 1059 | EMIHUDRAFT_522053 | eukaryota>haptophyceae | Emiliania | huxleyi CCMP1516 | hypothetical protein EMIHUDRAFT_522053 [Emiliania huxleyi CCMP1516] |
| NP_499386.1 | 1059 | sca-1 | eukaryota>metazoa>nematoda | Caenorhabditis | elegans | Calcium-transporting ATPase [Caenorhabditis elegans] |
| XP_003322729.1 | 1058 | PGTG_04266 | eukaryota>fungi>basidiomycota | Puccinia | graminis f. sp. tritici CRL 75-36-700-3 | Ca2+-transporting ATPase [Puccinia graminis f. sp. tritici CRL 75-36-700-3] |
| OEU12112.1 | 1058 | FRACYDRAFT_210916 | eukaryota>stramenopiles | Fragilariopsis | cylindrus CCMP1102 | calcium ATPase [Fragilariopsis cylindrus CCMP1102] |
| XP_001779936.1 | 1058 | Ppena2 | eukaryota>viridiplantae | Physcomitrella | patens | Na+ P-type-ATPase [Physcomitrella patens] |
| XP_014633223.1 | 1057 | LOC100800498 | eukaryota>viridiplantae | Glycine | max | PREDICTED: calcium-transporting ATPase, endoplasmic reticulum-type-like [Glycine max] |
| XP_014633222.1 | 1057 | LOC100800498 | eukaryota>viridiplantae | Glycine | max | PREDICTED: calcium-transporting ATPase, endoplasmic reticulum-type-like [Glycine max] |
| XP_003528778.1 | 1057 | LOC100800498 | eukaryota>viridiplantae | Glycine | max | PREDICTED: calcium-transporting ATPase, endoplasmic reticulum-type-like [Glycine max] |
| XP_003548255.1 | 1057 | LOC100792952 | eukaryota>viridiplantae | Glycine | max | PREDICTED: calcium-transporting ATPase 1, endoplasmic reticulum-type-like [Glycine max] |
| XP_006598889.1 | 1057 | LOC100792952 | eukaryota>viridiplantae | Glycine | max | PREDICTED: calcium-transporting ATPase 1, endoplasmic reticulum-type-like [Glycine max] |
| CDW82171.1 | 1056 | Contig9263.g9900 | eukaryota>alveolata>ciliophora | Stylonychia | lemnae | calcium-transporting atpase endoplasmic reticulum- [Stylonychia lemnae] |
| XP_009291732.1 | 1056 | si:dkey-28b4.8 | eukaryota>metazoa>chordata>vertebrata>actinopterygii | Danio | rerio | PREDICTED: sarcoplasmic/endoplasmic reticulum calcium ATPase 2 [Danio rerio] |
| XP_684227.1 | 1056 | si:dkey-28b4.8 | eukaryota>metazoa>chordata>vertebrata>actinopterygii | Danio | rerio | sarcoplasmic/endoplasmic reticulum calcium ATPase 2 [Danio rerio] |
| XP_018281950.1 | 1055 | CC85DRAFT_269298 | eukaryota>fungi>basidiomycota | Cutaneotrichosporon | oleaginosum | calcium-transporting P [Cutaneotrichosporon oleaginosum] |
| Crev1000005492 | 1055 | Crev1000005492 | eukaryota>fungi>kickxellomycotina | Coemansia | reversa | >Crev1000005492 |
| XP_001765945.1 | 1055 | PHYPADRAFT_212461 | eukaryota>viridiplantae | Physcomitrella | patens | predicted protein [Physcomitrella patens] |
| XP_504141.1 | 1054 | YALI0_E19338g | eukaryota>fungi>ascomycota | Yarrowia | lipolytica CLIB122 | YALI0E19338p [Yarrowia lipolytica CLIB122] |
| XP_014566252.1 | 1054 | L969DRAFT_96202 | eukaryota>fungi>basidiomycota | Mixia | osmundae IAM 14324 | hypothetical protein L969DRAFT_96202 [Mixia osmundae IAM 14324] |
| NP_001329712.1 | 1054 | ARABIDOPSIS,THALIANA,ER-TYPE,CA2+-ATPASE,2,ATECA2,A_TM018A10.4,A_TM018A10_4,ER-type,Ca2+-ATPase,2,T18A10.20,T18A10_20 | eukaryota>viridiplantae | Arabidopsis | thaliana | ER-type Ca2+-ATPase 2 [Arabidopsis thaliana] |
| NP_191999.1 | 1054 | ARABIDOPSIS,THALIANA,ER-TYPE,CA2+-ATPASE,2,ATECA2,A_TM018A10.4,A_TM018A10_4,ER-type,Ca2+-ATPase,2,T18A10.20,T18A10_20 | eukaryota>viridiplantae | Arabidopsis | thaliana | ER-type Ca2+-ATPase 2 [Arabidopsis thaliana] |
| XP_001889866.1 | 1053 | LACBIDRAFT_192284 | eukaryota>fungi>basidiomycota | Laccaria | bicolor S238N-H82 | Ca-transporting ATPase [Laccaria bicolor S238N-H82] |
| NP_777613.1 | 1052 | SERCA3 | eukaryota>metazoa>chordata>vertebrata | Homo | sapiens | sarcoplasmic/endoplasmic reticulum calcium ATPase 3 isoform e [Homo sapiens] |
| XP_008656544.1 | 1052 | LOC103635958 | eukaryota>viridiplantae | Zea | mays | PREDICTED: calcium-transporting ATPase 4, endoplasmic reticulum-type-like [Zea mays] |
| XP_002508654.1 | 1052 | MICPUN_97933 | eukaryota>viridiplantae>chlorophyta | Micromonas | commoda | p-type ATPase superfamily [Micromonas commoda] |
| XP_002175285.1 | 1051 | SJAG_04167 | eukaryota>fungi>ascomycota | Schizosaccharomyces | japonicus yFS275 | P-type ATPase [Schizosaccharomyces japonicus yFS275] |
| XP_010661979.1 | 1051 | LOC100253574 | eukaryota>viridiplantae | Vitis | vinifera | PREDICTED: calcium-transporting ATPase, endoplasmic reticulum-type [Vitis vinifera] |
| XP_002264585.2 | 1051 | LOC100253574 | eukaryota>viridiplantae | Vitis | vinifera | PREDICTED: calcium-transporting ATPase, endoplasmic reticulum-type [Vitis vinifera] |
| EJY74000.1 | 1050 | OXYTRI_04747 | eukaryota>alveolata>ciliophora | Oxytricha | trifallax | Calcium-transporting ATPase 1, endoplasmic reticulum-type, putative (macronuclear) [Oxytricha trifallax] |
| XP_011391135.1 | 1050 | UMAG_04461 | eukaryota>fungi>basidiomycota | Ustilago | maydis 521 | putative secretory pathway Ca++-transporting P-type ATPase [Ustilago maydis 521] |
| XP_016498837.1 | 1049 | LOC107817505 | eukaryota>viridiplantae | Nicotiana | tabacum | PREDICTED: calcium-transporting ATPase, endoplasmic reticulum-type-like [Nicotiana tabacum] |
| XP_016498836.1 | 1049 | LOC107817505 | eukaryota>viridiplantae | Nicotiana | tabacum | PREDICTED: calcium-transporting ATPase, endoplasmic reticulum-type-like [Nicotiana tabacum] |
| XP_016480058.1 | 1049 | LOC107801276 | eukaryota>viridiplantae | Nicotiana | tabacum | PREDICTED: calcium-transporting ATPase, endoplasmic reticulum-type [Nicotiana tabacum] |
| XP_016498838.1 | 1049 | LOC107817505 | eukaryota>viridiplantae | Nicotiana | tabacum | PREDICTED: calcium-transporting ATPase, endoplasmic reticulum-type-like [Nicotiana tabacum] |
| XP_016480053.1 | 1049 | LOC107801276 | eukaryota>viridiplantae | Nicotiana | tabacum | PREDICTED: calcium-transporting ATPase, endoplasmic reticulum-type [Nicotiana tabacum] |
| XP_016498840.1 | 1049 | LOC107817505 | eukaryota>viridiplantae | Nicotiana | tabacum | PREDICTED: calcium-transporting ATPase, endoplasmic reticulum-type-like [Nicotiana tabacum] |
| XP_016480055.1 | 1049 | LOC107801276 | eukaryota>viridiplantae | Nicotiana | tabacum | PREDICTED: calcium-transporting ATPase, endoplasmic reticulum-type [Nicotiana tabacum] |
| XP_016480057.1 | 1049 | LOC107801276 | eukaryota>viridiplantae | Nicotiana | tabacum | PREDICTED: calcium-transporting ATPase, endoplasmic reticulum-type [Nicotiana tabacum] |
| XP_016480054.1 | 1049 | LOC107801276 | eukaryota>viridiplantae | Nicotiana | tabacum | PREDICTED: calcium-transporting ATPase, endoplasmic reticulum-type [Nicotiana tabacum] |
| XP_016498839.1 | 1049 | LOC107817505 | eukaryota>viridiplantae | Nicotiana | tabacum | PREDICTED: calcium-transporting ATPase, endoplasmic reticulum-type-like [Nicotiana tabacum] |
| XP_016480056.1 | 1049 | LOC107801276 | eukaryota>viridiplantae | Nicotiana | tabacum | PREDICTED: calcium-transporting ATPase, endoplasmic reticulum-type [Nicotiana tabacum] |
| ABO93900.1 | 1049 | OSTLU_29095 | eukaryota>viridiplantae>chlorophyta | Ostreococcus | lucimarinus CCE9901 | P-ATPase family transporter: calcium ion [Ostreococcus lucimarinus CCE9901] |
| EPY22984.1 | 1048 | STCU_07975 | eukaryota>euglenozoa>kinetoplastida | Strigomonas | culicis | calcium motive p-type ATPase [Strigomonas culicis] |
| XP_018187906.1 | 1048 | L228DRAFT_230765 | eukaryota>fungi>ascomycota | Xylona | heveae TC161 | secretory pathway Ca2+-ATPase [Xylona heveae TC161] |
| XP_006339744.1 | 1048 | LOC102591952 | eukaryota>viridiplantae | Solanum | tuberosum | PREDICTED: calcium-transporting ATPase, endoplasmic reticulum-type [Solanum tuberosum] |
| XP_006339745.1 | 1048 | LOC102591952 | eukaryota>viridiplantae | Solanum | tuberosum | PREDICTED: calcium-transporting ATPase, endoplasmic reticulum-type [Solanum tuberosum] |
| XP_015167921.1 | 1048 | LOC102591952 | eukaryota>viridiplantae | Solanum | tuberosum | PREDICTED: calcium-transporting ATPase, endoplasmic reticulum-type [Solanum tuberosum] |
| XP_015167927.1 | 1048 | LOC102591952 | eukaryota>viridiplantae | Solanum | tuberosum | PREDICTED: calcium-transporting ATPase, endoplasmic reticulum-type [Solanum tuberosum] |
| XP_006339743.1 | 1048 | LOC102591952 | eukaryota>viridiplantae | Solanum | tuberosum | PREDICTED: calcium-transporting ATPase, endoplasmic reticulum-type [Solanum tuberosum] |
| XP_015167930.1 | 1048 | LOC102591952 | eukaryota>viridiplantae | Solanum | tuberosum | PREDICTED: calcium-transporting ATPase, endoplasmic reticulum-type [Solanum tuberosum] |
| XP_015167932.1 | 1048 | LOC102591952 | eukaryota>viridiplantae | Solanum | tuberosum | PREDICTED: calcium-transporting ATPase, endoplasmic reticulum-type [Solanum tuberosum] |
| XP_002968844.1 | 1047 | SELMODRAFT_170407 | eukaryota>viridiplantae | Selaginella | moellendorffii | hypothetical protein SELMODRAFT_170407 [Selaginella moellendorffii] |
| XP_002974128.1 | 1047 | SELMODRAFT_267711 | eukaryota>viridiplantae | Selaginella | moellendorffii | hypothetical protein SELMODRAFT_267711 [Selaginella moellendorffii] |
| XP_019039479.1 | 1046 | WICANDRAFT_62836 | eukaryota>fungi>ascomycota | Wickerhamomyces | anomalus NRRL Y-366-8 | hypothetical protein WICANDRAFT_62836 [Wickerhamomyces anomalus NRRL Y-366-8] |
| XP_005582600.1 | 1046 | ATP2A3 | eukaryota>metazoa>chordata>vertebrata | Macaca | fascicularis | PREDICTED: sarcoplasmic/endoplasmic reticulum calcium ATPase 3 isoform X1 [Macaca fascicularis] |
| XP_002903567.1 | 1046 | PITG_09282 | eukaryota>stramenopiles | Phytophthora | infestans T30-4 | calcium-transporting ATPase 1, endoplasmic reticulum-type, putative [Phytophthora infestans T30-4] |
| XP_009821873.1 | 1046 | H257_00743 | eukaryota>stramenopiles | Aphanomyces | astaci | hypothetical protein H257_00743 [Aphanomyces astaci] |
| XP_001542093.1 | 1045 | HCAG_02264 | eukaryota>fungi>ascomycota | Histoplasma | capsulatum NAm1 | sodium transport ATPase 5 [Histoplasma capsulatum NAm1] |
| XP_008862320.1 | 1045 | H310_01079 | eukaryota>stramenopiles | Aphanomyces | invadans | hypothetical protein H310_01079 [Aphanomyces invadans] |
| XP_008894556.1 | 1045 | PPTG_03625 | eukaryota>stramenopiles | Phytophthora | parasitica INRA-310 | hypothetical protein PPTG_03625 [Phytophthora parasitica INRA-310] |
| XP_002980453.1 | 1045 | SELMODRAFT_112465 | eukaryota>viridiplantae | Selaginella | moellendorffii | hypothetical protein SELMODRAFT_112465, partial [Selaginella moellendorffii] |
| NP_777614.1 | 1044 | SERCA3 | eukaryota>metazoa>chordata>vertebrata | Homo | sapiens | sarcoplasmic/endoplasmic reticulum calcium ATPase 3 isoform d [Homo sapiens] |
| NP_001103610.1 | 1044 | 9530097L16Rik,D5Wsu150e,mKIAA4195,SERCA2,Serca2a,SERCA2B | eukaryota>metazoa>chordata>vertebrata | Mus | musculus | sarcoplasmic/endoplasmic reticulum calcium ATPase 2 isoform b [Mus musculus] |
| OEU08503.1 | 1044 | FRACYDRAFT_264535 | eukaryota>stramenopiles | Fragilariopsis | cylindrus CCMP1102 | P-type ATPase [Fragilariopsis cylindrus CCMP1102] |
| XP_005821222.1 | 1043 | GUITHDRAFT_80735 | eukaryota>cryptophyta | Guillardia | theta CCMP2712 | hypothetical protein GUITHDRAFT_80735 [Guillardia theta CCMP2712] |
| Chet1000002268 | 1043 | Chet1000002268 | eukaryota>fungi>ascomycota | Cochliobolus | heterostrophus | >Chet1000002268 |
| NP_777615.1 | 1043 | SERCA3 | eukaryota>metazoa>chordata>vertebrata | Homo | sapiens | sarcoplasmic/endoplasmic reticulum calcium ATPase 3 isoform b [Homo sapiens] |
| XP_017323832.1 | 1043 | LOC108265716 | eukaryota>metazoa>chordata>vertebrata>actinopterygii | Ictalurus | punctatus | PREDICTED: sarcoplasmic/endoplasmic reticulum calcium ATPase 2-like isoform X1 [Ictalurus punctatus] |
| XP_009535831.1 | 1043 | PHYSODRAFT_564911 | eukaryota>stramenopiles | Phytophthora | sojae | hypothetical protein PHYSODRAFT_564911 [Phytophthora sojae] |
| XP_011122837.1 | 1042 | AOL_s00080g160 | eukaryota>fungi>ascomycota | Arthrobotrys | oligospora ATCC 24927 | hypothetical protein AOL_s00080g160 [Arthrobotrys oligospora ATCC 24927] |
| XP_011122415.1 | 1042 | AOL_s00079g238 | eukaryota>fungi>ascomycota | Arthrobotrys | oligospora ATCC 24927 | hypothetical protein AOL_s00079g238 [Arthrobotrys oligospora ATCC 24927] |
| NP_001258903.1 | 1042 | SERCA2 | eukaryota>metazoa>chordata>vertebrata | Gallus | gallus | sarcoplasmic/endoplasmic reticulum calcium ATPase 2 isoform 2 [Gallus gallus] |
| NP_990222.1 | 1042 | SERCA | eukaryota>metazoa>chordata>vertebrata | Gallus | gallus | sarcoplasmic/endoplasmic reticulum calcium ATPase 3 [Gallus gallus] |
| XP_011522187.1 | 1042 | SERCA3 | eukaryota>metazoa>chordata>vertebrata | Homo | sapiens | sarcoplasmic/endoplasmic reticulum calcium ATPase 3 isoform X5 [Homo sapiens] |
| NP_733765.1 | 1042 | ATP2B,DAR,DD,SERCA2 | eukaryota>metazoa>chordata>vertebrata | Homo | sapiens | sarcoplasmic/endoplasmic reticulum calcium ATPase 2 isoform b [Homo sapiens] |
| XP_005572290.1 | 1042 | ATP2A2 | eukaryota>metazoa>chordata>vertebrata | Macaca | fascicularis | PREDICTED: sarcoplasmic/endoplasmic reticulum calcium ATPase 2 isoform X1 [Macaca fascicularis] |
| XP_012365368.1 | 1042 | ATP2A2 | eukaryota>metazoa>chordata>vertebrata | Nomascus | leucogenys | PREDICTED: sarcoplasmic/endoplasmic reticulum calcium ATPase 2 isoform X1 [Nomascus leucogenys] |
| NP_001088218.1 | 1042 | LOC495046 | eukaryota>metazoa>chordata>vertebrata | Xenopus | laevis | uncharacterized protein LOC495046 [Xenopus laevis] |
| NP_001086443.1 | 1042 | atp2a,atp2a1,atp2a2,atp2b,dar,serca1,serca2 | eukaryota>metazoa>chordata>vertebrata | Xenopus | laevis | ATPase, Ca++ transporting, slow twitch 2 [Xenopus laevis] |
| XP_005166856.1 | 1042 | atp2a2,zgc:55380 | eukaryota>metazoa>chordata>vertebrata>actinopterygii | Danio | rerio | sarcoplasmic/endoplasmic reticulum calcium ATPase 2 isoform X1 [Danio rerio] |
| XP_005166855.1 | 1042 | atp2a2,zgc:55380 | eukaryota>metazoa>chordata>vertebrata>actinopterygii | Danio | rerio | sarcoplasmic/endoplasmic reticulum calcium ATPase 2 isoform X1 [Danio rerio] |
| XP_002974690.1 | 1042 | SELMODRAFT_102055 | eukaryota>viridiplantae | Selaginella | moellendorffii | hypothetical protein SELMODRAFT_102055 [Selaginella moellendorffii] |
| XP_001030432.2 | 1041 | TTHERM_01084370 | eukaryota>alveolata>ciliophora | Tetrahymena | thermophila SB210 | sarco/endoplasmic reticulum calcium-translocating P-type ATPase [Tetrahymena thermophila SB210] |
| XP_011777149.1 | 1041 | TbgDal_IX9590 | eukaryota>euglenozoa>kinetoplastida | Trypanosoma | brucei gambiense DAL972 | calcium motive p-type ATPase, putative [Trypanosoma brucei gambiense DAL972] |
| XP_016606375.1 | 1041 | SPPG_06728 | eukaryota>fungi>chytridiomycota | Spizellomyces | punctatus DAOM BR117 | potassium/sodium efflux P-type ATPase, fungal-type [Spizellomyces punctatus DAOM BR117] |
| Bcir1000012323 | 1041 | Bcir1000012323 | eukaryota>fungi>mucoromycotina | Backusella | circina | >Bcir1000012323 |
| XP_017347928.1 | 1041 | LOC108278817 | eukaryota>metazoa>chordata>vertebrata>actinopterygii | Ictalurus | punctatus | PREDICTED: sarcoplasmic/endoplasmic reticulum calcium ATPase 2 [Ictalurus punctatus] |
| XP_005161253.1 | 1041 | fb51e04,fb80c02,SERCA2,si:dkey-83d9.6,wu:fb51e04,wu:fb80c02,zK83D9.6 | eukaryota>metazoa>chordata>vertebrata>actinopterygii | Danio | rerio | ATPase, Ca++ transporting, cardiac muscle, slow twitch 2b isoform X1 [Danio rerio] |
| XP_002962638.1 | 1041 | SELMODRAFT_78895 | eukaryota>viridiplantae | Selaginella | moellendorffii | hypothetical protein SELMODRAFT_78895, partial [Selaginella moellendorffii] |
| XP_004040046.1 | 1040 | IMG5_002750 | eukaryota>alveolata>ciliophora | Ichthyophthirius | multifiliis | hypothetical protein IMG5_002750 [Ichthyophthirius multifiliis] |
| KNE56773.1 | 1040 | AMAG_02549 | eukaryota>fungi | Allomyces | macrogynus ATCC 38327 | potassium/sodium efflux P-type ATPase, fungal-type [Allomyces macrogynus ATCC 38327] |
| XP_005783170.1 | 1040 | ATPCa | eukaryota>haptophyceae | Emiliania | huxleyi CCMP1516 | Ca2+ ATPase [Emiliania huxleyi CCMP1516] |
| XP_817442.1 | 1039 | Tc00.1047053510769.120 | eukaryota>euglenozoa>kinetoplastida | Trypanosoma | cruzi strain CL Brener | calcium motive p-type ATPase [Trypanosoma cruzi strain CL Brener] |
| XP_817886.1 | 1039 | Tc00.1047053506241.70 | eukaryota>euglenozoa>kinetoplastida | Trypanosoma | cruzi strain CL Brener | calcium motive p-type ATPase [Trypanosoma cruzi strain CL Brener] |
| XP_018274164.1 | 1039 | RHOBADRAFT_32754 | eukaryota>fungi>basidiomycota | Rhodotorula | graminis WP1 | hypothetical protein RHOBADRAFT_32754 [Rhodotorula graminis WP1] |
| XP_001765694.1 | 1039 | PHYPADRAFT_184915 | eukaryota>viridiplantae | Physcomitrella | patens | predicted protein [Physcomitrella patens] |
| XP_002963405.1 | 1039 | SELMODRAFT_166180 | eukaryota>viridiplantae | Selaginella | moellendorffii | hypothetical protein SELMODRAFT_166180 [Selaginella moellendorffii] |
| NP_058025.2 | 1038 | Serca3,SERCA3b | eukaryota>metazoa>chordata>vertebrata | Mus | musculus | sarcoplasmic/endoplasmic reticulum calcium ATPase 3 isoform b [Mus musculus] |
| XP_003974774.1 | 1038 | atp2a2 | eukaryota>metazoa>chordata>vertebrata>actinopterygii | Takifugu | rubripes | PREDICTED: sarcoplasmic/endoplasmic reticulum calcium ATPase 2 [Takifugu rubripes] |
| XP_012766333.1 | 1037 | BBBOND_0104560 | eukaryota>alveolata>apicomplexa | Babesia | bigemina | calcium ATPase SERCA-like, putative [Babesia bigemina] |
| XP_001453966.1 | 1037 | GSPATT00020240001 | eukaryota>alveolata>ciliophora | Paramecium | tetraurelia strain d4-2 | hypothetical protein (macronuclear) [Paramecium tetraurelia strain d4-2] |
| NP_595246.1 | 1037 | cta3 | eukaryota>fungi>ascomycota | Schizosaccharomyces | pombe | P-type calcium transport ATPase Cta3 [Schizosaccharomyces pombe] |
| XP_005582601.1 | 1037 | ATP2A3 | eukaryota>metazoa>chordata>vertebrata | Macaca | fascicularis | PREDICTED: sarcoplasmic/endoplasmic reticulum calcium ATPase 3 isoform X2 [Macaca fascicularis] |
| XP_001778955.1 | 1037 | PHYPADRAFT_223041 | eukaryota>viridiplantae | Physcomitrella | patens | predicted protein [Physcomitrella patens] |
| XP_014526596.1 | 1036 | JH06_4380 | eukaryota>stramenopiles | Blastocystis | sp. subtype 4 | Ca2+ ATPase [Blastocystis sp. subtype 4] |
| Uram1000002146 | 1035 | Uram1000002146 | eukaryota>fungi>mucoromycotina | Umbelopsis | ramanniana | >Uram1000002146 |
| NP_001025448.1 | 1035 | fb51e04,fb80c02,SERCA2,si:dkey-83d9.6,wu:fb51e04,wu:fb80c02,zK83D9.6 | eukaryota>metazoa>chordata>vertebrata>actinopterygii | Danio | rerio | ATPase sarcoplasmic/endoplasmic reticulum Ca2+ transporting 2b [Danio rerio] |
| KNE72470.1 | 1034 | AMAG_16513 | eukaryota>fungi | Allomyces | macrogynus ATCC 38327 | potassium/sodium efflux P-type ATPase, fungal-type [Allomyces macrogynus ATCC 38327] |
| NP_730742.1 | 1034 | anon-WO0118547.249,CG14449,CG32451,CG7651,Dmel\CG32451,SPoCK | eukaryota>metazoa>hexapoda | Drosophila | melanogaster | secretory pathway calcium atpase, isoform B [Drosophila melanogaster] |
| XP_008609540.1 | 1034 | SDRG_05595 | eukaryota>stramenopiles | Saprolegnia | diclina VS20 | Ca2+-transporting ATPase [Saprolegnia diclina VS20] |
| XP_012196455.1 | 1034 | SPRG_02491 | eukaryota>stramenopiles | Saprolegnia | parasitica CBS 223.65 | calcium-translocating P-type ATPase [Saprolegnia parasitica CBS 223.65] |
| XP_006960882.1 | 1033 | PMR-1 | eukaryota>fungi>basidiomycota | Wallemia | mellicola CBS 633.66 | calcium-transporting P [Wallemia mellicola CBS 633.66] |
| XP_006675121.1 | 1033 | BATDEDRAFT_15466 | eukaryota>fungi>chytridiomycota | Batrachochytrium | dendrobatidis JAM81 | hypothetical protein BATDEDRAFT_15466 [Batrachochytrium dendrobatidis JAM81] |
| XP_018105186.1 | 1033 | LOC108709662 | eukaryota>metazoa>chordata>vertebrata | Xenopus | laevis | PREDICTED: sarcoplasmic/endoplasmic reticulum calcium ATPase 3-like [Xenopus laevis] |
| NP_001088563.1 | 1033 | atp2a3,serca3 | eukaryota>metazoa>chordata>vertebrata | Xenopus | laevis | ATPase sarcoplasmic/endoplasmic reticulum Ca2+ transporting 3 L homeolog [Xenopus laevis] |
| Bnat1000012666 | 1033 | Bnat1000012666 | eukaryota>rhizaria | Bigelowiella | natans | >Bnat1000012666 |
| XP_009038985.1 | 1033 | AURANDRAFT_29439 | eukaryota>stramenopiles | Aureococcus | anophagefferens | hypothetical protein AURANDRAFT_29439 [Aureococcus anophagefferens] |
| ORE10299.1 | 1033 | BCV72DRAFT_200094 | eukaryota>fungi | Rhizopus | microsporus var. microsporus | potassium/sodium efflux P-type ATPase [Rhizopus microsporus var. microsporus] |
| XP_005787448.1 | 1032 | ECA1 | eukaryota>haptophyceae | Emiliania | huxleyi CCMP1516 | Ca2+ ATPase [Emiliania huxleyi CCMP1516] |
| XP_008185966.1 | 1032 | LOC100167769 | eukaryota>metazoa>hexapoda | Acyrthosiphon | pisum | PREDICTED: calcium-transporting ATPase sarcoplasmic/endoplasmic reticulum type isoform X1 [Acyrthosiphon pisum] |
| XP_008185967.1 | 1032 | LOC100167769 | eukaryota>metazoa>hexapoda | Acyrthosiphon | pisum | PREDICTED: calcium-transporting ATPase sarcoplasmic/endoplasmic reticulum type isoform X1 [Acyrthosiphon pisum] |
| EJK45298.1 | 1032 | THAOC_36093 | eukaryota>stramenopiles | Thalassiosira | oceanica | hypothetical protein THAOC_36093 [Thalassiosira oceanica] |
| EIE86803.1 | 1031 | RO3G_11514 | eukaryota>fungi | Rhizopus | delemar RA 99-880 | potassium/sodium efflux P-type ATPase [Rhizopus delemar RA 99-880] |
| XP_013424406.1 | 1031 | M436DRAFT_75407 | eukaryota>fungi>ascomycota | Aureobasidium | namibiae CBS 147.97 | calcium transporter ATPase [Aureobasidium namibiae CBS 147.97] |
| XP_009039668.1 | 1030 | AURANDRAFT_10071 | eukaryota>stramenopiles | Aureococcus | anophagefferens | hypothetical protein AURANDRAFT_10071, partial [Aureococcus anophagefferens] |
| XP_002507319.1 | 1030 | MICPUN_78119 | eukaryota>viridiplantae>chlorophyta | Micromonas | commoda | p-type ATPase superfamily [Micromonas commoda] |
| NP_777618.1 | 1029 | SERCA3 | eukaryota>metazoa>chordata>vertebrata | Homo | sapiens | sarcoplasmic/endoplasmic reticulum calcium ATPase 3 isoform c [Homo sapiens] |
| NP_777616.1 | 1029 | SERCA3 | eukaryota>metazoa>chordata>vertebrata | Homo | sapiens | sarcoplasmic/endoplasmic reticulum calcium ATPase 3 isoform c [Homo sapiens] |
| XP_009051621.1 | 1029 | LOTGIDRAFT_208914 | eukaryota>metazoa>mollusca | Lottia | gigantea | hypothetical protein LOTGIDRAFT_208914 [Lottia gigantea] |
| XP_001610089.1 | 1028 | 18.m06474 | eukaryota>alveolata>apicomplexa | Babesia | bovis T2Bo | calcium ATPase SERCA-like [Babesia bovis T2Bo] |
| EPB86926.1 | 1028 | HMPREF1544_06249 | eukaryota>fungi | Mucor | circinelloides f. circinelloides 1006PhL | hypothetical protein HMPREF1544_06249 [Mucor circinelloides f. circinelloides 1006PhL] |
| XP_011522190.1 | 1028 | SERCA3 | eukaryota>metazoa>chordata>vertebrata | Homo | sapiens | sarcoplasmic/endoplasmic reticulum calcium ATPase 3 isoform X6 [Homo sapiens] |
| XP_002183209.1 | 1028 | PHATRDRAFT_228 | eukaryota>stramenopiles | Phaeodactylum | tricornutum CCAP 1055/1 | probable serca-type calcium ATPase, partial [Phaeodactylum tricornutum CCAP 1055/1] |
| XP_001447148.1 | 1026 | GSPATT00014681001 | eukaryota>alveolata>ciliophora | Paramecium | tetraurelia strain d4-2 | hypothetical protein (macronuclear) [Paramecium tetraurelia strain d4-2] |
| NP_001156809.1 | 1026 | Serca3,SERCA3b | eukaryota>metazoa>chordata>vertebrata | Mus | musculus | sarcoplasmic/endoplasmic reticulum calcium ATPase 3 isoform c [Mus musculus] |
| XP_010703651.1 | 1025 | LPMP_040010 | eukaryota>euglenozoa>kinetoplastida | Leishmania | panamensis | calcium-translocating P-type ATPase [Leishmania panamensis] |
| XP_001561691.1 | 1025 | LBRM_04_0030 | eukaryota>euglenozoa>kinetoplastida | Leishmania | braziliensis MHOM/BR/75/M2904 | calcium-translocating P-type ATPase [Leishmania braziliensis MHOM/BR/75/M2904] |
| XP_964218.1 | 1025 | pmr-1 | eukaryota>fungi>ascomycota | Neurospora | crassa OR74A | calcium-transporting ATPase type 2C member 1 [Neurospora crassa OR74A] |
| CDI98584.1 | 1025 | EmuJ_000244800 | eukaryota>metazoa | Echinococcus | multilocularis | sarco/endoplasmic reticulum calcium ATPase [Echinococcus multilocularis] |
| NP_001157948.1 | 1025 | LOC100302733 | eukaryota>metazoa>hexapoda | Bombyx | mori | sarco/endoplasmic reticulum calcium ATPase [Bombyx mori] |
| ERG85907.1 | 1025 | ASU_03077 | eukaryota>metazoa>nematoda | Ascaris | suum | calcium-transporting atpase sarcoplasmic endoplasmic reticulum type [Ascaris suum] |
| CBJ48791.1 | 1025 | Esi_0047_0101 | eukaryota>stramenopiles | Ectocarpus | siliculosus | conserved unknown protein [Ectocarpus siliculosus] |
| ABO95354.1 | 1025 | OSTLU_14777 | eukaryota>viridiplantae>chlorophyta | Ostreococcus | lucimarinus CCE9901 | P-ATPase family transporter: calcium ion [Ostreococcus lucimarinus CCE9901] |
| XP_013245771.1 | 1024 | K437DRAFT_219607 | eukaryota>fungi>basidiomycota | Tilletiaria | anomala UBC 951 | potassium/sodium eff [Tilletiaria anomala UBC 951] |
| Capa1000000589 | 1024 | Capa1000000589 | eukaryota>metazoa>annelida | Capitella | sp1 | >Capa1000000589 |
| XP_011522191.1 | 1024 | SERCA3 | eukaryota>metazoa>chordata>vertebrata | Homo | sapiens | sarcoplasmic/endoplasmic reticulum calcium ATPase 3 isoform X7 [Homo sapiens] |
| EFX65740.1 | 1024 | DAPPUDRAFT_65262 | eukaryota>metazoa>crustacea | Daphnia | pulex | hypothetical protein DAPPUDRAFT_65262 [Daphnia pulex] |
| XP_013755992.1 | 1023 | AMSG_07502 | eukaryota>apusozoa>apusomonadidae | Thecamonas | trahens ATCC 50062 | ATPase [Thecamonas trahens ATCC 50062] |
| XP_888512.1 | 1023 | LMJF_04_0010 | eukaryota>euglenozoa>kinetoplastida | Leishmania | major strain Friedlin | calcium-translocating P-type ATPase [Leishmania major strain Friedlin] |
| XP_003858064.1 | 1023 | LDBPK_040010 | eukaryota>euglenozoa>kinetoplastida | Leishmania | donovani | calcium-translocating P-type ATPase [Leishmania donovani] |
| XP_001462838.2 | 1023 | LINJ_04_0010 | eukaryota>euglenozoa>kinetoplastida | Leishmania | infantum JPCM5 | calcium-translocating P-type ATPase [Leishmania infantum JPCM5] |
| EPB85748.1 | 1023 | HMPREF1544_07499 | eukaryota>fungi | Mucor | circinelloides f. circinelloides 1006PhL | Ca2+ transporting ATPase, sarcoplasmic/endoplasmic reticulum [Mucor circinelloides f. circinelloides 1006PhL] |
| EXX62354.1 | 1023 | RirG_162570 | eukaryota>fungi | Rhizophagus | irregularis DAOM 197198w | Na(+)-exporting P-type ATPase ENA2 [Rhizophagus irregularis DAOM 197198w] |
| XP_012895122.1 | 1023 | GSBLH_T00001287001 | eukaryota>stramenopiles | Blastocystis | hominis | uncharacterized protein [Blastocystis hominis] |
| XP_016607069.1 | 1022 | SPPG_05978 | eukaryota>fungi>chytridiomycota | Spizellomyces | punctatus DAOM BR117 | calcium-translocating P-type ATPase, SERCA-type [Spizellomyces punctatus DAOM BR117] |
| XP_018648452.1 | 1022 | Smp_007260.1 | eukaryota>metazoa | Schistosoma | mansoni | putative atpase [Schistosoma mansoni] |
| NP_001032719.1 | 1022 | SERCA | eukaryota>metazoa>echinodermata | Strongylocentrotus | purpuratus | sarco/endoplasmic reticulum calcium transporting ATPase [Strongylocentrotus purpuratus] |
| XP_005757745.1 | 1020 | EMIHUDRAFT_62350 | eukaryota>haptophyceae | Emiliania | huxleyi CCMP1516 | hypothetical protein EMIHUDRAFT_62350 [Emiliania huxleyi CCMP1516] |
| OAF65999.1 | 1020 | A3Q56_06285 | eukaryota>metazoa | Intoshia | linei | hypothetical protein A3Q56_06285 [Intoshia linei] |
| XP_013421498.1 | 1020 | LOC106181601 | eukaryota>metazoa | Lingula | anatina | calcium-transporting ATPase sarcoplasmic/endoplasmic reticulum type isoform X1 [Lingula anatina] |
| XP_013421499.1 | 1020 | LOC106181601 | eukaryota>metazoa | Lingula | anatina | calcium-transporting ATPase sarcoplasmic/endoplasmic reticulum type isoform X2 [Lingula anatina] |
| XP_013421500.1 | 1020 | LOC106181601 | eukaryota>metazoa | Lingula | anatina | calcium-transporting ATPase sarcoplasmic/endoplasmic reticulum type isoform X3 [Lingula anatina] |
| XP_013421502.1 | 1020 | LOC106181601 | eukaryota>metazoa | Lingula | anatina | calcium-transporting ATPase sarcoplasmic/endoplasmic reticulum type isoform X1 [Lingula anatina] |
| XP_013775740.1 | 1020 | LOC106460569 | eukaryota>metazoa | Limulus | polyphemus | calcium-transporting ATPase sarcoplasmic/endoplasmic reticulum type-like isoform X1 [Limulus polyphemus] |
| XP_016880182.1 | 1020 | SERCA3 | eukaryota>metazoa>chordata>vertebrata | Homo | sapiens | sarcoplasmic/endoplasmic reticulum calcium ATPase 3 isoform X8 [Homo sapiens] |
| XP_018115259.1 | 1020 | atp2c2 | eukaryota>metazoa>chordata>vertebrata | Xenopus | laevis | PREDICTED: ATPase, Ca++ transporting, type 2C, member 2 S homeolog isoform X1 [Xenopus laevis] |
| XP_011601308.1 | 1020 | LOC101069346 | eukaryota>metazoa>chordata>vertebrata>actinopterygii | Takifugu | rubripes | PREDICTED: LOW QUALITY PROTEIN: sarcoplasmic/endoplasmic reticulum calcium ATPase 1-like [Takifugu rubripes] |
| XP_019697792.1 | 1020 | LOC105184991 | eukaryota>metazoa>hexapoda | Harpegnathos | saltator | calcium-transporting ATPase sarcoplasmic/endoplasmic reticulum type isoform X1 [Harpegnathos saltator] |
| XP_016770979.1 | 1020 | LOC410371 | eukaryota>metazoa>hexapoda | Apis | mellifera | PREDICTED: calcium-transporting ATPase sarcoplasmic/endoplasmic reticulum type isoform X1 [Apis mellifera] |
| XP_011142454.1 | 1020 | LOC105184991 | eukaryota>metazoa>hexapoda | Harpegnathos | saltator | calcium-transporting ATPase sarcoplasmic/endoplasmic reticulum type isoform X1 [Harpegnathos saltator] |
| XP_011142453.1 | 1020 | LOC105184991 | eukaryota>metazoa>hexapoda | Harpegnathos | saltator | calcium-transporting ATPase sarcoplasmic/endoplasmic reticulum type isoform X1 [Harpegnathos saltator] |
| XP_006565283.1 | 1020 | LOC410371 | eukaryota>metazoa>hexapoda | Apis | mellifera | PREDICTED: calcium-transporting ATPase sarcoplasmic/endoplasmic reticulum type isoform X1 [Apis mellifera] |
| XP_006565282.1 | 1020 | LOC410371 | eukaryota>metazoa>hexapoda | Apis | mellifera | PREDICTED: calcium-transporting ATPase sarcoplasmic/endoplasmic reticulum type isoform X1 [Apis mellifera] |
| NP_726387.1 | 1020 | 17136664,A4UZU0_DROME,ATC1_DROME,Ca-ATPase,Ca-p,Ca-P60,Ca-P60A,CA-P60A,CaP60A,Ca[2+]ATPase,CG3725,Dmel\CG3725,dserca,dSERCA,kum,Kum,l(2)k08308ab,l(2)k09025,l(2)SH0211,serca,Serca,SERCA2,Serca2a | eukaryota>metazoa>hexapoda | Drosophila | melanogaster | Sarco/endoplasmic reticulum Ca(2+)-ATPase, isoform H [Drosophila melanogaster] |
| NP_726381.1 | 1020 | 17136664,A4UZU0_DROME,ATC1_DROME,Ca-ATPase,Ca-p,Ca-P60,Ca-P60A,CA-P60A,CaP60A,Ca[2+]ATPase,CG3725,Dmel\CG3725,dserca,dSERCA,kum,Kum,l(2)k08308ab,l(2)k09025,l(2)SH0211,serca,Serca,SERCA2,Serca2a | eukaryota>metazoa>hexapoda | Drosophila | melanogaster | Sarco/endoplasmic reticulum Ca(2+)-ATPase, isoform B [Drosophila melanogaster] |
| NP_726386.1 | 1020 | 17136664,A4UZU0_DROME,ATC1_DROME,Ca-ATPase,Ca-p,Ca-P60,Ca-P60A,CA-P60A,CaP60A,Ca[2+]ATPase,CG3725,Dmel\CG3725,dserca,dSERCA,kum,Kum,l(2)k08308ab,l(2)k09025,l(2)SH0211,serca,Serca,SERCA2,Serca2a | eukaryota>metazoa>hexapoda | Drosophila | melanogaster | Sarco/endoplasmic reticulum Ca(2+)-ATPase, isoform G [Drosophila melanogaster] |
| NP_726382.1 | 1020 | 17136664,A4UZU0_DROME,ATC1_DROME,Ca-ATPase,Ca-p,Ca-P60,Ca-P60A,CA-P60A,CaP60A,Ca[2+]ATPase,CG3725,Dmel\CG3725,dserca,dSERCA,kum,Kum,l(2)k08308ab,l(2)k09025,l(2)SH0211,serca,Serca,SERCA2,Serca2a | eukaryota>metazoa>hexapoda | Drosophila | melanogaster | Sarco/endoplasmic reticulum Ca(2+)-ATPase, isoform C [Drosophila melanogaster] |
| NP_726383.1 | 1020 | 17136664,A4UZU0_DROME,ATC1_DROME,Ca-ATPase,Ca-p,Ca-P60,Ca-P60A,CA-P60A,CaP60A,Ca[2+]ATPase,CG3725,Dmel\CG3725,dserca,dSERCA,kum,Kum,l(2)k08308ab,l(2)k09025,l(2)SH0211,serca,Serca,SERCA2,Serca2a | eukaryota>metazoa>hexapoda | Drosophila | melanogaster | Sarco/endoplasmic reticulum Ca(2+)-ATPase, isoform D [Drosophila melanogaster] |
| NP_726384.1 | 1020 | 17136664,A4UZU0_DROME,ATC1_DROME,Ca-ATPase,Ca-p,Ca-P60,Ca-P60A,CA-P60A,CaP60A,Ca[2+]ATPase,CG3725,Dmel\CG3725,dserca,dSERCA,kum,Kum,l(2)k08308ab,l(2)k09025,l(2)SH0211,serca,Serca,SERCA2,Serca2a | eukaryota>metazoa>hexapoda | Drosophila | melanogaster | Sarco/endoplasmic reticulum Ca(2+)-ATPase, isoform E [Drosophila melanogaster] |
| XP_011142452.1 | 1020 | LOC105184991 | eukaryota>metazoa>hexapoda | Harpegnathos | saltator | calcium-transporting ATPase sarcoplasmic/endoplasmic reticulum type isoform X1 [Harpegnathos saltator] |
| NP_726385.1 | 1020 | 17136664,A4UZU0_DROME,ATC1_DROME,Ca-ATPase,Ca-p,Ca-P60,Ca-P60A,CA-P60A,CaP60A,Ca[2+]ATPase,CG3725,Dmel\CG3725,dserca,dSERCA,kum,Kum,l(2)k08308ab,l(2)k09025,l(2)SH0211,serca,Serca,SERCA2,Serca2a | eukaryota>metazoa>hexapoda | Drosophila | melanogaster | Sarco/endoplasmic reticulum Ca(2+)-ATPase, isoform F [Drosophila melanogaster] |
| XP_006565281.1 | 1020 | LOC410371 | eukaryota>metazoa>hexapoda | Apis | mellifera | PREDICTED: calcium-transporting ATPase sarcoplasmic/endoplasmic reticulum type isoform X1 [Apis mellifera] |
| XP_966783.1 | 1019 | LOC658328 | eukaryota>metazoa>hexapoda | Tribolium | castaneum | PREDICTED: calcium-transporting ATPase sarcoplasmic/endoplasmic reticulum type isoform X1 [Tribolium castaneum] |
| XP_015839081.1 | 1019 | LOC658328 | eukaryota>metazoa>hexapoda | Tribolium | castaneum | PREDICTED: calcium-transporting ATPase sarcoplasmic/endoplasmic reticulum type isoform X1 [Tribolium castaneum] |
| XP_008198220.1 | 1019 | LOC658328 | eukaryota>metazoa>hexapoda | Tribolium | castaneum | PREDICTED: calcium-transporting ATPase sarcoplasmic/endoplasmic reticulum type isoform X1 [Tribolium castaneum] |
| XP_008198221.1 | 1019 | LOC658328 | eukaryota>metazoa>hexapoda | Tribolium | castaneum | PREDICTED: calcium-transporting ATPase sarcoplasmic/endoplasmic reticulum type isoform X1 [Tribolium castaneum] |
| ORZ30791.1 | 1019 | BCR44DRAFT_1443840 | eukaryota>fungi | Catenaria | anguillulae PL171 | putative calcium ion P-type ATPase [Catenaria anguillulae PL171] |
| XP_021870418.1 | 1019 | BD324DRAFT_580559 | eukaryota>fungi>basidiomycota | Kockovaella | imperatae | putative calcium-transporting ATPase sarcoplasmic/endoplasmic reticulum type [Kockovaella imperatae] |
| XP_006956013.1 | 1018 | cta3 | eukaryota>fungi>basidiomycota | Wallemia | mellicola CBS 633.66 | potassium/sodium eff [Wallemia mellicola CBS 633.66] |
| XP_018113448.1 | 1018 | LOC108714077 | eukaryota>metazoa>chordata>vertebrata | Xenopus | laevis | PREDICTED: calcium-transporting ATPase type 2C member 2-like [Xenopus laevis] |
| XP_018115260.1 | 1018 | atp2c2 | eukaryota>metazoa>chordata>vertebrata | Xenopus | laevis | PREDICTED: ATPase, Ca++ transporting, type 2C, member 2 S homeolog isoform X2 [Xenopus laevis] |
| XP_016770980.1 | 1018 | LOC410371 | eukaryota>metazoa>hexapoda | Apis | mellifera | PREDICTED: calcium-transporting ATPase sarcoplasmic/endoplasmic reticulum type isoform X2 [Apis mellifera] |
| XP_021875769.1 | 1018 | BCR41DRAFT_364239 | eukaryota>fungi | Lobosporangium | transversale | hypothetical protein BCR41DRAFT_364239 [Lobosporangium transversale] |
| OZ06563.1 | 1018 | BZG36_00488 | eukaryota>fungi | Bifiguratus | adelaidae | hypothetical protein BZG36_00488 [Bifiguratus adelaidae] |
| XP_018270006.1 | 1017 | RHOBADRAFT_28354 | eukaryota>fungi>basidiomycota | Rhodotorula | graminis WP1 | hypothetical protein RHOBADRAFT_28354 [Rhodotorula graminis WP1] |
| XP_009025276.1 | 1017 | HELRODRAFT_185062 | eukaryota>metazoa>annelida | Helobdella | robusta | hypothetical protein HELRODRAFT_185062 [Helobdella robusta] |
| XP_012895489.1 | 1017 | GSBLH_T00001611001 | eukaryota>stramenopiles | Blastocystis | hominis | uncharacterized protein [Blastocystis hominis] |
| EPB85791.1 | 1016 | HMPREF1544_07460 | eukaryota>fungi | Mucor | circinelloides f. circinelloides 1006PhL | hypothetical protein HMPREF1544_07460 [Mucor circinelloides f. circinelloides 1006PhL] |
| XP_019041886.1 | 1016 | WICANDRAFT_99077 | eukaryota>fungi>ascomycota | Wickerhamomyces | anomalus NRRL Y-366-8 | hypothetical protein WICANDRAFT_99077 [Wickerhamomyces anomalus NRRL Y-366-8] |
| XP_001540332.1 | 1016 | HCAG_04172 | eukaryota>fungi>ascomycota | Histoplasma | capsulatum NAm1 | sarcoplasmic/endoplasmic reticulum calcium ATPase 1 [Histoplasma capsulatum NAm1] |
| XP_019922302.1 | 1016 | LOC105327139 | eukaryota>metazoa>mollusca | Crassostrea | gigas | PREDICTED: calcium-transporting ATPase sarcoplasmic/endoplasmic reticulum type [Crassostrea gigas] |
| CCW66645.1 | 1015 | GSHART1_T00000934001 | eukaryota>euglenozoa>kinetoplastida | Phytomonas | sp. isolate Hart1 | unnamed protein product [Phytomonas sp. isolate Hart1] |
| XP_013421503.1 | 1015 | LOC106181601 | eukaryota>metazoa | Lingula | anatina | calcium-transporting ATPase sarcoplasmic/endoplasmic reticulum type isoform X4 [Lingula anatina] |
| XP_013421504.1 | 1015 | LOC106181601 | eukaryota>metazoa | Lingula | anatina | calcium-transporting ATPase sarcoplasmic/endoplasmic reticulum type isoform X4 [Lingula anatina] |
| XP_002286890.1 | 1015 | THAPSDRAFT_204 | eukaryota>stramenopiles | Thalassiosira | pseudonana CCMP1335 | cation transport ATPase [Thalassiosira pseudonana CCMP1335] |
| XP_003055417.1 | 1015 | MICPUCDRAFT_50020 | eukaryota>viridiplantae>chlorophyta | Micromonas | pusilla CCMP1545 | p-type ATPase superfamily [Micromonas pusilla CCMP1545] |
| XP_009015784.1 | 1014 | HELRODRAFT_106126 | eukaryota>metazoa>annelida | Helobdella | robusta | hypothetical protein HELRODRAFT_106126 [Helobdella robusta] |
| XP_003871768.1 | 1013 | LMXM_04_0010 | eukaryota>euglenozoa>kinetoplastida | Leishmania | mexicana MHOM/GT/2001/U1103 | calcium-translocating P-type ATPase [Leishmania mexicana MHOM/GT/2001/U1103] |
| XP_004344407.2 | 1013 | CAOG_06786 | eukaryota>ichthyosporea | Capsaspora | owczarzaki ATCC 30864 | sarco/endoplasmic reticulum calcium transporting ATPase [Capsaspora owczarzaki ATCC 30864] |
| CEO97902.1 | 1013 | PBRA_006016 | eukaryota>rhizaria>cercozoa>plasmodiophorida | Plasmodiophora | brassicae | hypothetical protein PBRA_006016 [Plasmodiophora brassicae] |
| XP_003078384.1 | 1013 | Ot03g04740 | eukaryota>viridiplantae>chlorophyta | Ostreococcus | tauri | Ca2+-ATPase (ISS) [Ostreococcus tauri] |
| XP_013421505.1 | 1012 | LOC106181601 | eukaryota>metazoa | Lingula | anatina | calcium-transporting ATPase sarcoplasmic/endoplasmic reticulum type isoform X5 [Lingula anatina] |
| ORX97427.1 | 1012 | K493DRAFT_314088 | eukaryota>fungi | Basidiobolus | meristosporus CBS 931.73 | Ca-transporting ATPase [Basidiobolus meristosporus CBS 931.73] |
| XP_021879152.1 | 1012 | BCR41DRAFT_423988 | eukaryota>fungi | Lobosporangium | transversale | Ca-transporting ATPase [Lobosporangium transversale] |
| XP_011773626.1 | 1011 | TbgDal_V4780 | eukaryota>euglenozoa>kinetoplastida | Trypanosoma | brucei gambiense DAL972 | calcium pump, putative [Trypanosoma brucei gambiense DAL972] |
| GAQ88260.1 | 1011 | KFL_004130060 | eukaryota>viridiplantae | Klebsormidium | nitens | Ca2+ transporting ATPase [Klebsormidium nitens] |
| XP_002985331.1 | 1011 | SELMODRAFT_122175 | eukaryota>viridiplantae | Selaginella | moellendorffii | hypothetical protein SELMODRAFT_122175 [Selaginella moellendorffii] |
| XP_021880735.1 | 1011 | BCR41DRAFT_316570 | eukaryota>fungi | Lobosporangium | transversale | putative Na P-type ATPase [Lobosporangium transversale] |
| XP_015651596.1 | 1010 | ABB37_10062 | eukaryota>euglenozoa>kinetoplastida | Leptomonas | pyrrhocoris | putative mitochondrial calcium-translocating P-type ATPase [Leptomonas pyrrhocoris] |
| KPI84659.1 | 1010 | ABL78_6300 | eukaryota>euglenozoa>kinetoplastida | Leptomonas | seymouri | calcium-translocating P-type ATPase [Leptomonas seymouri] |
| KFH62632.1 | 1009 | MVEG_12024 | eukaryota>fungi | Mortierella | verticillata NRRL 6337 | Ca2+ transporting ATPase, sarcoplasmic/endoplasmic reticulum [Mortierella verticillata NRRL 6337] |
| XP_018282490.1 | 1009 | CC85DRAFT_325254 | eukaryota>fungi>basidiomycota | Cutaneotrichosporon | oleaginosum | calcium ATPase [Cutaneotrichosporon oleaginosum] |
| XP_011388455.1 | 1009 | UMAG_02081 | eukaryota>fungi>basidiomycota | Ustilago | maydis 521 | endoplasmic reticulum calcium transporter [Ustilago maydis 521] |
| XP_002987187.1 | 1009 | SELMODRAFT_158488 | eukaryota>viridiplantae | Selaginella | moellendorffii | hypothetical protein SELMODRAFT_158488 [Selaginella moellendorffii] |
| XP_014566126.1 | 1008 | L969DRAFT_96071 | eukaryota>fungi>basidiomycota | Mixia | osmundae IAM 14324 | hypothetical protein L969DRAFT_96071 [Mixia osmundae IAM 14324] |
| Bcir1000000233 | 1008 | Bcir1000000233 | eukaryota>fungi>mucoromycotina | Backusella | circina | >Bcir1000000233 |
| EUB64292.1 | 1008 | EGR_00836 | eukaryota>metazoa | Echinococcus | granulosus | Calcium-transporting ATPase sarcoplasmic/endoplasmic reticulum type [Echinococcus granulosus] |
| XP_019850897.1 | 1008 | LOC100634808 | eukaryota>metazoa | Amphimedon | queenslandica | PREDICTED: calcium-transporting ATPase sarcoplasmic/endoplasmic reticulum type-like [Amphimedon queenslandica] |
| CAG02658.1 | 1008 | GSTEN:00021814:G:001 | eukaryota>metazoa>chordata>vertebrata>actinopterygii | Tetraodon | nigroviridis | unnamed protein product [Tetraodon nigroviridis] |
| XP_005715614.1 | 1008 | CHC_T00008559001 | eukaryota>rhodophyta | Chondrus | crispus | calcium-transporting ATPase, endoplasmic reticulum type [Chondrus crispus] |
| XP_009312974.1 | 1007 | DQ04_06441000 | eukaryota>euglenozoa>kinetoplastida | Trypanosoma | grayi | calcium-transporting P-type ATPase [Trypanosoma grayi] |
| Chet1000002786 | 1007 | Chet1000002786 | eukaryota>fungi>ascomycota | Cochliobolus | heterostrophus | >Chet1000002786 |
| Lhya1000005952 | 1007 | Lhya1000005952 | eukaryota>fungi>mucoromycotina | Lichtheimia | hyalospora | >Lhya1000005952 |
| CBY21930.1 | 1007 | GSOID_T00011465001 | eukaryota>metazoa>chordata | Oikopleura | dioica | unnamed protein product [Oikopleura dioica] |
| XP_814228.1 | 1006 | Tc00.1047053509777.70 | eukaryota>euglenozoa>kinetoplastida | Trypanosoma | cruzi strain CL Brener | calcium-translocating P-type ATPase [Trypanosoma cruzi strain CL Brener] |
| XP_014531013.1 | 1006 | PDIP_68560 | eukaryota>fungi>ascomycota | Penicillium | digitatum Pd1 | Endoplasmic reticulum calcium ATPase, putative [Penicillium digitatum Pd1] |
| XP_012053868.1 | 1006 | CNAG_05592 | eukaryota>fungi>basidiomycota | Cryptococcus | neoformans var. grubii H99 | potassium/sodium efflux P-type ATPase, fungal-type [Cryptococcus neoformans var. grubii H99] |
| XP_002177149.1 | 1006 | PHATRDRAFT_53964 | eukaryota>stramenopiles | Phaeodactylum | tricornutum CCAP 1055/1 | probable calcium ATPase [Phaeodactylum tricornutum CCAP 1055/1] |
| ORZ31935.1 | 1006 | BCR44DRAFT_1480249 | eukaryota>fungi | Catenaria | anguillulae PL171 | hypothetical protein BCR44DRAFT_1480249 [Catenaria anguillulae PL171] |
| KFH74078.1 | 1005 | MVEG_01291 | eukaryota>fungi | Mortierella | verticillata NRRL 6337 | hypothetical protein MVEG_01291 [Mortierella verticillata NRRL 6337] |
| XP_009013683.1 | 1005 | HELRODRAFT_156619 | eukaryota>metazoa>annelida | Helobdella | robusta | hypothetical protein HELRODRAFT_156619 [Helobdella robusta] |
| XP_017308246.1 | 1005 | atp2a1 | eukaryota>metazoa>chordata>vertebrata>actinopterygii | Ictalurus | punctatus | PREDICTED: LOW QUALITY PROTEIN: sarcoplasmic/endoplasmic reticulum calcium ATPase 1 [Ictalurus punctatus] |
| XP_017315524.1 | 1005 | atp2a3 | eukaryota>metazoa>chordata>vertebrata>actinopterygii | Ictalurus | punctatus | PREDICTED: sarcoplasmic/endoplasmic reticulum calcium ATPase 3 [Ictalurus punctatus] |
| XP_697108.5 | 1005 | atp2a3 | eukaryota>metazoa>chordata>vertebrata>actinopterygii | Danio | rerio | sarcoplasmic/endoplasmic reticulum calcium ATPase 3 isoform X1 [Danio rerio] |
| XP_001639528.1 | 1005 | NEMVEDRAFT_v1g160437 | eukaryota>metazoa>cnidaria | Nematostella | vectensis | predicted protein [Nematostella vectensis] |
| XP_019697793.1 | 1005 | LOC105184991 | eukaryota>metazoa>hexapoda | Harpegnathos | saltator | calcium-transporting ATPase sarcoplasmic/endoplasmic reticulum type isoform X2 [Harpegnathos saltator] |
| XP_016662301.1 | 1005 | LOC100167769 | eukaryota>metazoa>hexapoda | Acyrthosiphon | pisum | PREDICTED: calcium-transporting ATPase sarcoplasmic/endoplasmic reticulum type isoform X2 [Acyrthosiphon pisum] |
| XP_005539258.1 | 1005 | CYME_CMT241C | eukaryota>rhodophyta | Cyanidioschyzon | merolae strain 10D | calcium-transporting ATPase, endoplasmic reticulum type [Cyanidioschyzon merolae strain 10D] |
| XP_018282216.1 | 1004 | CC85DRAFT_282349 | eukaryota>fungi>basidiomycota | Cutaneotrichosporon | oleaginosum | putative calcium-transporting ATPase 3 [Cutaneotrichosporon oleaginosum] |
| Bcir1000006932 | 1004 | Bcir1000006932 | eukaryota>fungi>mucoromycotina | Backusella | circina | >Bcir1000006932 |
| XP_018648453.1 | 1004 | Smp_007260.1 | eukaryota>metazoa | Schistosoma | mansoni | putative atpase [Schistosoma mansoni] |
| XP_013775741.1 | 1004 | LOC106460569 | eukaryota>metazoa | Limulus | polyphemus | calcium-transporting ATPase sarcoplasmic/endoplasmic reticulum type-like isoform X2 [Limulus polyphemus] |

| | | | | | | |
|---|---|---|---|---|---|---|
| NP_499385.3 | 1004 | sca-1 | eukaryota>metazoa>nematoda | Caenorhabditis | elegans | Calcium-transporting ATPase [Caenorhabditis elegans] |
| XP_001696690.1 | 1004 | CHLREDRAFT_104709 | eukaryota>viridiplantae>chlorophyta | Chlamydomonas | reinhardtii | cation transporting ATPase, partial [Chlamydomonas reinhardtii] |
| ORX91705.1 | 1004 | K493DRAFT_264186 | eukaryota>fungi | Basidiobolus | meristosporus CBS 931.73 | Ca-transporting ATPase [Basidiobolus meristosporus CBS 931.73] |
| XP_004992033.1 | 1003 | PTSG_06645 | eukaryota>choanoflagellida | Salpingoeca | rosetta | ATPase [Salpingoeca rosetta] |
| XP_005821433.1 | 1003 | GUITHDRAFT_158817 | eukaryota>cryptophyta | Guillardia | theta CCMP2712 | hypothetical protein GUITHDRAFT_158817 [Guillardia theta CCMP2712] |
| KFH69757.1 | 1003 | MVEG_04563 | eukaryota>fungi | Mortierella | verticillata NRRL 6337 | hypothetical protein MVEG_04563 [Mortierella verticillata NRRL 6337] |
| Crev1000001104 | 1003 | Crev1000001104 | eukaryota>fungi>kickxellomycotina | Coemansia | reversa | >Crev1000001104 |
| XP_002122235.2 | 1003 | LOC100175957 | eukaryota>metazoa>chordata | Ciona | intestinalis | PREDICTED: sarcoplasmic/endoplasmic reticulum calcium ATPase 1-like [Ciona intestinalis] |
| XP_009861042.2 | 1003 | LOC100182988 | eukaryota>metazoa>chordata | Ciona | intestinalis | PREDICTED: sarcoplasmic/endoplasmic reticulum calcium ATPase 1-like isoform X1 [Ciona intestinalis] |
| XP_009861043.2 | 1003 | LOC100182988 | eukaryota>metazoa>chordata | Ciona | intestinalis | PREDICTED: sarcoplasmic/endoplasmic reticulum calcium ATPase 2-like isoform X2 [Ciona intestinalis] |
| XP_003970855.1 | 1003 | atp2a3 | eukaryota>metazoa>chordata>vertebrata>actinopterygii | Takifugu | rubripes | PREDICTED: sarcoplasmic/endoplasmic reticulum calcium ATPase 3 [Takifugu rubripes] |
| XP_016770981.1 | 1003 | LOC410371 | eukaryota>metazoa>hexapoda | Apis | mellifera | PREDICTED: calcium-transporting ATPase sarcoplasmic/endoplasmic reticulum type isoform X3 [Apis mellifera] |
| ORX95125.1 | 1003 | K493DRAFT_283100 | eukaryota>fungi | Basidiobolus | meristosporus CBS 931.73 | potassium/sodium efflux P-type ATPase [Basidiobolus meristosporus CBS 931.73] |
| XP_009496868.1 | 1002 | H696_04730 | eukaryota | Fonticula | alba | Ca2+ transporting ATPase, sarcoplasmic/endoplasmic reticulum [Fonticula alba] |
| XP_004226981.1 | 1002 | LOC100180638 | eukaryota>metazoa>chordata | Ciona | intestinalis | PREDICTED: sarcoplasmic/endoplasmic reticulum calcium ATPase 1-like isoform X1 [Ciona intestinalis] |
| XP_011663710.1 | 1002 | SERCA | eukaryota>metazoa>echinodermata | Strongylocentrotus | purpuratus | PREDICTED: sarco/endoplasmic reticulum calcium transporting ATPase isoform X1 [Strongylocentrotus purpuratus] |
| NP_476832.1 | 1002 | 17136664,A4UZU0_DROME,ATC1_DROME,Ca-ATPase,Ca-p,Ca-P60,Ca-P60A,CA-P60A,CaP60A,Ca[2+]ATPase,CG3725,Dmel\CG3725,dserca,dSERCA,kum,Kum,l(2)k08308ab,l(2)k09025,l(2)SH0211,serca,Serca,SERCA2,Serca2a | eukaryota>metazoa>hexapoda | Drosophila | melanogaster | Sarco/endoplasmic reticulum Ca(2+)-ATPase, isoform A [Drosophila melanogaster] |
| XP_016770982.1 | 1002 | LOC410371 | eukaryota>metazoa>hexapoda | Apis | mellifera | PREDICTED: calcium-transporting ATPase sarcoplasmic/endoplasmic reticulum type isoform X4 [Apis mellifera] |
| XP_019697794.1 | 1002 | LOC105184991 | eukaryota>metazoa>hexapoda | Harpegnathos | saltator | calcium-transporting ATPase sarcoplasmic/endoplasmic reticulum type isoform X3 [Harpegnathos saltator] |
| XP_018296061.1 | 1001 | PHYBLDRAFT_185726 | eukaryota>fungi | Phycomyces | blakesleeanus NRRL 1555(-) | hypothetical protein PHYBLDRAFT_185726 [Phycomyces blakesleeanus NRRL 1555(-)] |
| XP_013243079.1 | 1001 | K437DRAFT_256669 | eukaryota>fungi>basidiomycota | Tilletiaria | anomala UBC 951 | Ca-transporting ATPase [Tilletiaria anomala UBC 951] |
| NP_775293.1 | 1001 | ATP2A,SERCA1 | eukaryota>metazoa>chordata>vertebrata | Homo | sapiens | sarcoplasmic/endoplasmic reticulum calcium ATPase 1 isoform b [Homo sapiens] |
| XP_015297988.1 | 1001 | ATP2A1 | eukaryota>metazoa>chordata>vertebrata | Macaca | fascicularis | PREDICTED: sarcoplasmic/endoplasmic reticulum calcium ATPase 1 [Macaca fascicularis] |
| XP_006507331.1 | 1001 | SERCA1 | eukaryota>metazoa>chordata>vertebrata | Mus | musculus | PREDICTED: sarcoplasmic/endoplasmic reticulum calcium ATPase 1 isoform X1 [Mus musculus] |
| XP_012360942.1 | 1001 | ATP2A1 | eukaryota>metazoa>chordata>vertebrata | Nomascus | leucogenys | PREDICTED: LOW QUALITY PROTEIN: sarcoplasmic/endoplasmic reticulum calcium ATPase 1 [Nomascus leucogenys] |
| XP_015839082.1 | 1001 | LOC658328 | eukaryota>metazoa>hexapoda | Tribolium | castaneum | PREDICTED: calcium-transporting ATPase sarcoplasmic/endoplasmic reticulum type isoform X2 [Tribolium castaneum] |
| XP_002294029.1 | 1001 | THAPSDRAFT_37662 | eukaryota>stramenopiles | Thalassiosira | pseudonana CCMP1335 | cation transporting ATPase [Thalassiosira pseudonana CCMP1335] |
| XP_003524018.1 | 1001 | LOC100783509 | eukaryota>viridiplantae | Glycine | max | PREDICTED: calcium-transporting ATPase 3, endoplasmic reticulum-type isoform X1 [Glycine max] |
| XP_001696782.1 | 1001 | CHLREDRAFT_130931 | eukaryota>viridiplantae>chlorophyta | Chlamydomonas | reinhardtii | calcium transporting ATPase [Chlamydomonas reinhardtii] |
| XP_012647248.1 | 1000 | BBM_I00885 | eukaryota>alveolata>apicomplexa | Babesia | microti strain RI | uncharacterized protein [Babesia microti strain RI] |
| OMH84339.1 | 1000 | AX774_g2139 | eukaryota>fungi | Zancudomyces | culisetae | Calcium-transporting ATPase 3 [Zancudomyces culisetae] |
| XP_007326281.1 | 1000 | AGABI1DRAFT_69139 | eukaryota>fungi>basidiomycota | Agaricus | bisporus var. burnettii JB137-S8 | hypothetical protein AGABI1DRAFT_69139 [Agaricus bisporus var. burnettii JB137-S8] |
| XP_009168054.1 | 1000 | T265_04922 | eukaryota>metazoa | Opisthorchis | viverrini | hypothetical protein T265_04922 [Opisthorchis viverrini] |
| XP_002127180.1 | 1000 | LOC100180619 | eukaryota>metazoa>chordata | Ciona | intestinalis | PREDICTED: sarcoplasmic/endoplasmic reticulum calcium ATPase 1-like [Ciona intestinalis] |
| XP_002119917.1 | 1000 | LOC100180638 | eukaryota>metazoa>chordata | Ciona | intestinalis | PREDICTED: sarcoplasmic/endoplasmic reticulum calcium ATPase 1-like isoform X2 [Ciona intestinalis] |
| XP_015839084.1 | 1000 | LOC658328 | eukaryota>metazoa>hexapoda | Tribolium | castaneum | PREDICTED: calcium-transporting ATPase sarcoplasmic/endoplasmic reticulum type isoform X3 [Tribolium castaneum] |
| XP_012895050.1 | 1000 | GSBLH_T00001230001 | eukaryota>stramenopiles | Blastocystis | hominis | uncharacterized protein [Blastocystis hominis] |
| XP_001778460.1 | 1000 | PHYPADRAFT_222630 | eukaryota>viridiplantae | Physcomitrella | patens | predicted protein [Physcomitrella patens] |
| XP_006347865.1 | 1000 | LOC102595434 | eukaryota>viridiplantae | Solanum | tuberosum | PREDICTED: calcium-transporting ATPase 3, endoplasmic reticulum-type isoform X1 [Solanum tuberosum] |
| XP_010664492.1 | 1000 | LOC100266003 | eukaryota>viridiplantae | Vitis | vinifera | PREDICTED: calcium-transporting ATPase 3, endoplasmic reticulum-type isoform X1 [Vitis vinifera] |
| XP_006458484.1 | 1000 | AGABI2DRAFT_183521 | eukaryota>fungi>basidiomycota | Agaricus | bisporus var. bisporus H97 | Ca-transporting ATPase [Agaricus bisporus var. bisporus H97] |
| XP_013425005.1 | 999 | M436DRAFT_84359 | eukaryota>fungi>ascomycota | Aureobasidium | namibiae CBS 147.97 | calcium ATPase [Aureobasidium namibiae CBS 147.97] |
| Lhya1000006398 | 999 | Lhya1000006398 | eukaryota>fungi>mucoromycotina | Lichtheimia | hyalospora | >Lhya1000006398 |
| XP_015151158.1 | 999 | SERCA | eukaryota>metazoa>chordata>vertebrata | Gallus | gallus | PREDICTED: sarcoplasmic/endoplasmic reticulum calcium ATPase 3 isoform X1 [Gallus gallus] |
| XP_011536704.1 | 999 | ATP2B,DAR,DD,SERCA2 | eukaryota>metazoa>chordata>vertebrata | Homo | sapiens | sarcoplasmic/endoplasmic reticulum calcium ATPase 2 isoform X1 [Homo sapiens] |
| XP_005253945.1 | 999 | ATP2B,DAR,DD,SERCA2 | eukaryota>metazoa>chordata>vertebrata | Homo | sapiens | sarcoplasmic/endoplasmic reticulum calcium ATPase 2 isoform X1 [Homo sapiens] |
| NP_005164.2 | 999 | SERCA3 | eukaryota>metazoa>chordata>vertebrata | Homo | sapiens | sarcoplasmic/endoplasmic reticulum calcium ATPase 3 isoform a [Homo sapiens] |
| XP_015287004.1 | 999 | ATP2A2 | eukaryota>metazoa>chordata>vertebrata | Macaca | fascicularis | PREDICTED: sarcoplasmic/endoplasmic reticulum calcium ATPase 2 isoform X2 [Macaca fascicularis] |
| XP_005572291.1 | 999 | ATP2A2 | eukaryota>metazoa>chordata>vertebrata | Macaca | fascicularis | PREDICTED: sarcoplasmic/endoplasmic reticulum calcium ATPase 2 isoform X2 [Macaca fascicularis] |
| NP_001156808.1 | 999 | Serca3,SERCA3b | eukaryota>metazoa>chordata>vertebrata | Mus | musculus | sarcoplasmic/endoplasmic reticulum calcium ATPase 3 isoform a [Mus musculus] |
| NP_001286796.1 | 999 | 17136664,A4UZU0_DROME,ATC1_DROME,Ca-ATPase,Ca-p,Ca-P60,Ca-P60A,CA-P60A,CaP60A,Ca[2+]ATPase,CG3725,Dmel\CG3725,dserca,dSERCA,kum,Kum,l(2)k08308ab,l(2)k09025,l(2)SH0211,serca,Serca,SERCA2,Serca2a | eukaryota>metazoa>hexapoda | Drosophila | melanogaster | Sarco/endoplasmic reticulum Ca(2+)-ATPase, isoform I [Drosophila melanogaster] |
| OZJ04612.1 | 999 | BZG36_02032 | eukaryota>fungi | Bifiguratus | adelaidae | hypothetical protein BZG36_02032 [Bifiguratus adelaidae] |
| ORY72275.1 | 999 | LY90DRAFT_379315 | eukaryota>fungi>chytridiomycota | Neocallimastix | californiae | Ca-transporting ATPase [Neocallimastix californiae] |
| XP_001744818.1 | 998 | MONBRDRAFT_7291 | eukaryota>choanoflagellida | Monosiga | brevicollis MX1 | hypothetical protein [Monosiga brevicollis MX1] |
| KXN70998.1 | 998 | CONCODRAFT_6371 | eukaryota>fungi | Conidiobolus | coronatus NRRL 28638 | calcium-transporting ATPase [Conidiobolus coronatus NRRL 28638] |
| XP_011121099.1 | 998 | AOL_s00076g244 | eukaryota>fungi>ascomycota | Arthrobotrys | oligospora ATCC 24927 | hypothetical protein AOL_s00076g244 [Arthrobotrys oligospora ATCC 24927] |
| XP_013421506.1 | 998 | LOC106181601 | eukaryota>metazoa | Lingula | anatina | calcium-transporting ATPase sarcoplasmic/endoplasmic reticulum type isoform X6 [Lingula anatina] |
| [illegible] | [illegible] | [illegible] | [illegible] | [illegible] | [illegible] | [illegible] |

| | | | | | | |
|---|---|---|---|---|---|---|
| XP_005156418.1 | 627 | camk2ga,zgc:109908 | eukaryota>metazoa>chordata>vertebrata>actinopterygii | Danio | rerio | calcium/calmodulin-dependent protein kinase type II subunit gamma isoform X1 [Danio rerio] |
| XP_006514532.1 | 627 | Camk2b | eukaryota>metazoa>chordata>vertebrata | Mus | musculus | PREDICTED: calcium/calmodulin-dependent protein kinase type II subunit beta isoform X6 [Mus musculus] |
| XP_017306963.1 | 627 | LOC108255499 | eukaryota>metazoa>chordata>vertebrata>actinopterygii | Ictalurus | punctatus | PREDICTED: calcium/calmodulin-dependent protein kinase type II delta chain-like isoform X15 [Ictalurus punctatus] |
| XP_008664763.1 | 627 | LOC103643377 | eukaryota>viridiplantae | Zea | mays | CDPK-related kinase 3 [Zea mays] |
| XP_005156961.1 | 626 | camk2g2 | eukaryota>metazoa>chordata>vertebrata>actinopterygii | Danio | rerio | calcium/calmodulin-dependent protein kinase type II subunit gamma isoform X1 [Danio rerio] |
| XP_006514533.1 | 626 | Camk2b | eukaryota>metazoa>chordata>vertebrata | Mus | musculus | PREDICTED: calcium/calmodulin-dependent protein kinase type II subunit beta isoform X7 [Mus musculus] |
| XP_006715844.1 | 626 | CAM2,CAMK2,CAMKB | eukaryota>metazoa>chordata>vertebrata | Homo | sapiens | PREDICTED: calcium/calmodulin-dependent protein kinase type II subunit beta isoform X27 [Homo sapiens] |
| XP_002903060.1 | 626 | PITG_09039 | eukaryota>stramenopiles | Phytophthora | infestans T30-4 | calcium/calmodulin-dependent protein kinase, putative [Phytophthora infestans T30-4] |
| XP_011121085.1 | 626 | AOL_s00076g230 | eukaryota>fungi>ascomycota | Arthrobotrys | oligospora ATCC 24927 | hypothetical protein AOL_s00076g230 [Arthrobotrys oligospora ATCC 24927] |
| XP_015833937.1 | 625 | LOC655266 | eukaryota>metazoa>hexapoda | Tribolium | castaneum | PREDICTED: calcium/calmodulin-dependent protein kinase type II alpha chain isoform X1 [Tribolium castaneum] |
| NP_001105740.1 | 625 | GRMZM2G115518,MCK1 | eukaryota>viridiplantae | Zea | mays | calcium dependent protein kinase 1 [Zea mays] |
| XP_012757847.1 | 625 | SAMD00019534_018020 | eukaryota>amoebozoa>mycetozoa>dictyosteliida | Acytostelium | subglobosum LB1 | hypothetical protein SAMD00019534_018020 [Acytostelium subglobosum LB1] |
| XP_005156419.1 | 624 | camk2ga,zgc:109908 | eukaryota>metazoa>chordata>vertebrata>actinopterygii | Danio | rerio | calcium/calmodulin-dependent protein kinase type II subunit gamma isoform X2 [Danio rerio] |
| XP_017338896.1 | 624 | LOC108273829 | eukaryota>metazoa>chordata>vertebrata>actinopterygii | Ictalurus | punctatus | PREDICTED: calcium/calmodulin-dependent protein kinase type II subunit gamma isoform X1 [Ictalurus punctatus] |
| CBJ27073.1 | 624 | CDPK2 | eukaryota>stramenopiles | Ectocarpus | siliculosus | conserved unknown protein [Ectocarpus siliculosus] |
| XP_009305633.1 | 623 | camk2g2 | eukaryota>metazoa>chordata>vertebrata>actinopterygii | Danio | rerio | calcium/calmodulin-dependent protein kinase type II subunit gamma isoform X2 [Danio rerio] |
| XP_015833938.1 | 623 | LOC655266 | eukaryota>metazoa>hexapoda | Tribolium | castaneum | PREDICTED: calcium/calmodulin-dependent protein kinase type II alpha chain isoform X2 [Tribolium castaneum] |
| XP_018106263.1 | 623 | cam2,camk2,camk2b,camkb | eukaryota>metazoa>chordata>vertebrata | Xenopus | laevis | PREDICTED: calcium/calmodulin-dependent protein kinase (CaM kinase) II beta L homeolog isoform X1 [Xenopus laevis] |
| XP_018106264.1 | 623 | cam2,camk2,camk2b,camkb | eukaryota>metazoa>chordata>vertebrata | Xenopus | laevis | PREDICTED: calcium/calmodulin-dependent protein kinase (CaM kinase) II beta L homeolog isoform X2 [Xenopus laevis] |
| XP_018109671.1 | 623 | camk2,camk2a,camk2b | eukaryota>metazoa>chordata>vertebrata | Xenopus | laevis | PREDICTED: calcium/calmodulin-dependent protein kinase II beta isoform X1 [Xenopus laevis] |
| XP_018109672.1 | 623 | camk2,camk2a,camk2b | eukaryota>metazoa>chordata>vertebrata | Xenopus | laevis | PREDICTED: calcium/calmodulin-dependent protein kinase II beta isoform X2 [Xenopus laevis] |
| EPY26434.1 | 623 | AGDE_11328 | eukaryota>euglenozoa>kinetoplastida | Angomonas | deanei | protein kinase [Angomonas deanei] |
| XP_012656331.1 | 623 | TTHERM_000841299 | eukaryota>alveolata>ciliophora | Tetrahymena | thermophila SB210 | calmodulin-domain kinase [Tetrahymena thermophila SB210] |
| XP_018080932.1 | 622 | LOC108696257 | eukaryota>metazoa>chordata>vertebrata | Xenopus | laevis | PREDICTED: calcium/calmodulin-dependent protein kinase type II subunit gamma-like isoform X3 [Xenopus laevis] |
| XP_016496330.1 | 622 | LOC107815283 | eukaryota>viridiplantae | Nicotiana | tabacum | PREDICTED: calcium-dependent protein kinase 26-like [Nicotiana tabacum] |
| XP_016510658.1 | 622 | LOC107827935 | eukaryota>viridiplantae | Nicotiana | tabacum | PREDICTED: calcium-dependent protein kinase 26-like [Nicotiana tabacum] |
| XP_003884375.1 | 622 | NCLIV_047750 | eukaryota>alveolata>apicomplexa | Neospora | caninum Liverpool | hypothetical protein NCLIV_047750 [Neospora caninum Liverpool] |
| XP_001388076.1 | 622 | cgd4_3330 | eukaryota>alveolata>apicomplexa | Cryptosporidium | parvum Iowa II | hypothetical protein, partial [Cryptosporidium parvum Iowa II] |
| XP_012759150.1 | 622 | SAMD00019534_002620 | eukaryota>amoebozoa>mycetozoa>dictyosteliida | Acytostelium | subglobosum LB1 | hypothetical protein SAMD00019534_002620 [Acytostelium subglobosum LB1] |
| XP_017338897.1 | 621 | LOC108273829 | eukaryota>metazoa>chordata>vertebrata>actinopterygii | Ictalurus | punctatus | PREDICTED: calcium/calmodulin-dependent protein kinase type II subunit gamma isoform X2 [Ictalurus punctatus] |
| CDW74871.1 | 621 | Contig12712.g13564 | eukaryota>alveolata>ciliophora | Stylonychia | lemnae | protein kinase domain containing protein [Stylonychia lemnae] |
| XP_018080933.1 | 620 | LOC108696257 | eukaryota>metazoa>chordata>vertebrata | Xenopus | laevis | PREDICTED: calcium/calmodulin-dependent protein kinase type II subunit gamma-like isoform X4 [Xenopus laevis] |
| CEM28668.1 | 619 | Vbra_17763 | eukaryota>alveolata | Vitrella | brassicaformis CCMP3155 | unnamed protein product [Vitrella brassicaformis CCMP3155] |
| XP_006560571.1 | 617 | Camkii | eukaryota>metazoa>hexapoda | Apis | mellifera | PREDICTED: calcium/calmodulin-dependent protein kinase II isoform X4 [Apis mellifera] |
| XP_015833939.1 | 617 | LOC655266 | eukaryota>metazoa>hexapoda | Tribolium | castaneum | PREDICTED: calcium/calmodulin-dependent protein kinase type II alpha chain isoform X3 [Tribolium castaneum] |
| XP_018082303.1 | 617 | camk2,camk2g | eukaryota>metazoa>chordata>vertebrata | Xenopus | laevis | PREDICTED: calcium/calmodulin-dependent protein kinase (CaM kinase) II gamma S homeolog isoform X1 [Xenopus laevis] |
| XP_002774607.1 | 616 | Pmar_PMAR006234 | eukaryota>alveolata | Perkinsus | marinus ATCC 50983 | calcium-dependent protein kinase, putative [Perkinsus marinus ATCC 50983] |
| XP_006560572.1 | 615 | Camkii | eukaryota>metazoa>hexapoda | Apis | mellifera | PREDICTED: calcium/calmodulin-dependent protein kinase II isoform X5 [Apis mellifera] |
| XP_006560573.1 | 615 | Camkii | eukaryota>metazoa>hexapoda | Apis | mellifera | PREDICTED: calcium/calmodulin-dependent protein kinase II isoform X6 [Apis mellifera] |
| XP_015833940.1 | 615 | LOC655266 | eukaryota>metazoa>hexapoda | Tribolium | castaneum | PREDICTED: calcium/calmodulin-dependent protein kinase type II alpha chain isoform X4 [Tribolium castaneum] |
| XP_019081311.1 | 615 | LOC100250518 | eukaryota>viridiplantae | Vitis | vinifera | PREDICTED: CDPK-related kinase 3 isoform X2 [Vitis vinifera] |
| XP_016872205.1 | 615 | CAMK,CAMK-II,CAMKG | eukaryota>metazoa>chordata>vertebrata | Homo | sapiens | PREDICTED: calcium/calmodulin-dependent protein kinase type II subunit gamma isoform X1 [Homo sapiens] |
| EJY84673.1 | 615 | OXYTRI_17480 | eukaryota>alveolata>ciliophora | Oxytricha | trifallax | Protein kinase domain containing protein (macronuclear) [Oxytricha trifallax] |
| XP_001691570.1 | 613 | CDPK1 | eukaryota>viridiplantae>chlorophyta | Chlamydomonas | reinhardtii | calcium-dependent protein kinase 1 [Chlamydomonas reinhardtii] |
| CBN78721.1 | 613 | PK | eukaryota>stramenopiles | Ectocarpus | siliculosus | calmodulin-dependent protein kinase I [Ectocarpus siliculosus] |
| XP_009299603.1 | 612 | camk2b,CAMK2Gb,fj90d04,si:dkey-174n20.2,wu:fj90d04,zgc:165373 | eukaryota>metazoa>chordata>vertebrata>actinopterygii | Danio | rerio | calcium/calmodulin-dependent protein kinase type II subunit beta isoform X20 [Danio rerio] |
| XP_002949744.1 | 612 | VOLCADRAFT_74309 | eukaryota>viridiplantae>chlorophyta | Volvox | carteri f. nagariensis | hypothetical protein VOLCADRAFT_74309 [Volvox carteri f. nagariensis] |
| XP_012759910.1 | 612 | SAMD00019534_035640 | eukaryota>amoebozoa>mycetozoa>dictyosteliida | Acytostelium | subglobosum LB1 | hypothetical protein SAMD00019534_035640 [Acytostelium subglobosum LB1] |
| XP_005156962.1 | 611 | camk2g2 | eukaryota>metazoa>chordata>vertebrata>actinopterygii | Danio | rerio | calcium/calmodulin-dependent protein kinase type II subunit gamma isoform X3 [Danio rerio] |
| CEM12751.1 | 611 | Vbra_15452 | eukaryota>alveolata | Vitrella | brassicaformis CCMP3155 | unnamed protein product [Vitrella brassicaformis CCMP3155] |
| NP_196107.1 | 610 | ATCPK1,CALCIUM,DEPENDENT,PROTEIN,KINASE,calcium,dependent,protein,kinase,1 | eukaryota>viridiplantae | Arabidopsis | thaliana | calcium dependent protein kinase 1 [Arabidopsis thaliana] |
| XP_016482898.1 | 610 | LOC107803647 | eukaryota>viridiplantae | Nicotiana | tabacum | PREDICTED: calcium-dependent protein kinase 26-like [Nicotiana tabacum] |
| XP_006338215.1 | 610 | LOC102588824 | eukaryota>viridiplantae | Solanum | tuberosum | PREDICTED: CDPK-related kinase 5 [Solanum tuberosum] |
| XP_017306964.1 | 610 | LOC108255499 | eukaryota>metazoa>chordata>vertebrata>actinopterygii | Ictalurus | punctatus | PREDICTED: calcium/calmodulin-dependent protein kinase type II subunit gamma-like isoform X16 [Ictalurus punctatus] |
| XP_008674183.1 | 609 | LOC103650380 | eukaryota>viridiplantae | Zea | mays | calcium-dependent protein kinase 27 [Zea mays] |
| XP_009305634.1 | 608 | camk2g2 | eukaryota>metazoa>chordata>vertebrata>actinopterygii | Danio | rerio | calcium/calmodulin-dependent protein kinase type II subunit gamma isoform X5 [Danio rerio] |
| XP_015143790.1 | 608 | CAMK2G | eukaryota>metazoa>chordata>vertebrata | Gallus | gallus | PREDICTED: calcium/calmodulin-dependent protein kinase type II subunit gamma isoform X1 [Gallus gallus] |
| [illegible] | [illegible] | [illegible] | [illegible] | [illegible] | [illegible] | [illegible] |

| | | | | | | |
|---|---|---|---|---|---|---|
| XP_002769463.1 | 535 | Pmar PMAR008962 | eukaryota>alveolata | Perkinsus | marinus ATCC 50983 | calcium-dependent protein kinase 1, putative [Perkinsus marinus ATCC 50983] |
| XP_009304937.1 | 534 | camk2d,camk2da,wu:fq17a12,zgc:92792 | eukaryota>metazoa>chordata>vertebrata>actinopterygii | Danio | rerio | calcium/calmodulin-dependent protein kinase type II delta 2 chain isoform X5 [Danio rerio] |
| NP_001132098.1 | 534 | LOC100193514 | eukaryota>viridiplantae | Zea | mays | uncharacterized protein LOC100193514 [Zea mays] |
| XP_002276630.2 | 534 | LOC100251670 | eukaryota>viridiplantae | Vitis | vinifera | PREDICTED: calcium-dependent protein kinase 34 [Vitis vinifera] |
| XP_003521182.1 | 534 | LOC100808399 | eukaryota>viridiplantae | Glycine | max | PREDICTED: calcium-dependent protein kinase 32-like [Glycine max] |
| XP_016477399.1 | 534 | LOC107798882 | eukaryota>viridiplantae | Nicotiana | tabacum | PREDICTED: calcium-dependent protein kinase 21-like [Nicotiana tabacum] |
| XP_016479757.1 | 534 | LOC107801007 | eukaryota>viridiplantae | Nicotiana | tabacum | PREDICTED: calcium-dependent protein kinase 8-like [Nicotiana tabacum] |
| NP_195331.2 | 534 | calcium-dependent,protein,kinase,18,T19K4.200,T19K4_200 | eukaryota>viridiplantae | Arabidopsis | thaliana | calcium-dependent protein kinase 18 [Arabidopsis thaliana] |
| NP_001023297.1 | 533 | unc-43 | eukaryota>metazoa>nematoda | Caenorhabditis | elegans | Calcium/calmodulin-dependent protein kinase type II [Caenorhabditis elegans] |
| NP_001308498.1 | 533 | CAMKD | eukaryota>metazoa>chordata>vertebrata | Homo | sapiens | calcium/calmodulin-dependent protein kinase type II subunit delta isoform 9 [Homo sapiens] |
| NP_001308500.1 | 533 | CAMKD | eukaryota>metazoa>chordata>vertebrata | Homo | sapiens | calcium/calmodulin-dependent protein kinase type II subunit delta isoform 11 [Homo sapiens] |
| XP_003269380.1 | 533 | CAMK2D | eukaryota>metazoa>chordata>vertebrata | Nomascus | leucogenys | PREDICTED: calcium/calmodulin-dependent protein kinase type II subunit delta isoform X1 [Nomascus leucogenys] |
| XP_005565529.1 | 533 | CAMK2G | eukaryota>metazoa>chordata>vertebrata | Macaca | fascicularis | PREDICTED: calcium/calmodulin-dependent protein kinase type II subunit gamma isoform X12 [Macaca fascicularis] |
| XP_006500879.1 | 533 | 2810011D23Rik,8030469K03Rik,CaMK,II,[d]-CaMKII | eukaryota>metazoa>chordata>vertebrata | Mus | musculus | PREDICTED: calcium/calmodulin-dependent protein kinase type II subunit delta isoform X1 [Mus musculus] |
| XP_006500880.1 | 533 | 2810011D23Rik,8030469K03Rik,CaMK,II,[d]-CaMKII | eukaryota>metazoa>chordata>vertebrata | Mus | musculus | PREDICTED: calcium/calmodulin-dependent protein kinase type II subunit delta isoform X2 [Mus musculus] |
| XP_006518560.1 | 533 | Camkg | eukaryota>metazoa>chordata>vertebrata | Mus | musculus | PREDICTED: calcium/calmodulin-dependent protein kinase type II subunit gamma isoform X10 [Mus musculus] |
| XP_009304938.1 | 533 | camk2d,camk2da,wu:fq17a12,zgc:92792 | eukaryota>metazoa>chordata>vertebrata>actinopterygii | Danio | rerio | calcium/calmodulin-dependent protein kinase type II delta 2 chain isoform X6 [Danio rerio] |
| XP_013779505.1 | 533 | LOC106463959 | eukaryota>metazoa | Limulus | polyphemus | calcium/calmodulin-dependent protein kinase type II alpha chain isoform X2 [Limulus polyphemus] |
| XP_015305988.1 | 533 | CAMK2D | eukaryota>metazoa>chordata>vertebrata | Macaca | fascicularis | PREDICTED: calcium/calmodulin-dependent protein kinase type II subunit delta isoform X1 [Macaca fascicularis] |
| XP_015305989.1 | 533 | CAMK2D | eukaryota>metazoa>chordata>vertebrata | Macaca | fascicularis | PREDICTED: calcium/calmodulin-dependent protein kinase type II subunit delta isoform X2 [Macaca fascicularis] |
| GAQ88622.1 | 533 | KFL_004450090 | eukaryota>viridiplantae | Klebsormidium | nitens | Serine Threonine protein kinase [Klebsormidium nitens] |
| NP_001190672.1 | 533 | ATCPK23,calcium-dependent,protein,kinase,23,T4B21.15,T4B21_15 | eukaryota>viridiplantae | Arabidopsis | thaliana | calcium-dependent protein kinase 23 [Arabidopsis thaliana] |
| NP_197446.1 | 533 | calcium-dependent,protein,kinase,19,CALCIUM-DEPENDENT,PROTEIN,KINASE,19,CPK8,F7K24.200,F7K24_200 | eukaryota>viridiplantae | Arabidopsis | thaliana | calcium-dependent protein kinase 19 [Arabidopsis thaliana] |
| NP_850853.1 | 533 | calcium-dependent,protein,kinase,19,CALCIUM-DEPENDENT,PROTEIN,KINASE,19,CPK8,F7K24.200,F7K24_200 | eukaryota>viridiplantae | Arabidopsis | thaliana | calcium-dependent protein kinase 19 [Arabidopsis thaliana] |
| XP_003529159.1 | 533 | LOC100783922 | eukaryota>viridiplantae | Glycine | max | PREDICTED: calcium-dependent protein kinase 13 [Glycine max] |
| XP_003540704.1 | 533 | LOC100801716 | eukaryota>viridiplantae | Glycine | max | PREDICTED: calcium-dependent protein kinase 24-like [Glycine max] |
| XP_006366539.1 | 533 | LOC102594770 | eukaryota>viridiplantae | Solanum | tuberosum | PREDICTED: calcium-dependent protein kinase 8-like [Solanum tuberosum] |
| XP_001434254.1 | 533 | GSPATT00035988001 | eukaryota>alveolata>ciliophora | Paramecium | tetraurelia strain d4-2 | hypothetical protein (macronuclear) [Paramecium tetraurelia strain d4-2] |
| XP_001438645.1 | 533 | GSPATT00000961001 | eukaryota>alveolata>ciliophora | Paramecium | tetraurelia strain d4-2 | hypothetical protein (macronuclear) [Paramecium tetraurelia strain d4-2] |
| XP_001439733.1 | 533 | GSPATT00008778001 | eukaryota>alveolata>ciliophora | Paramecium | tetraurelia strain d4-2 | hypothetical protein (macronuclear) [Paramecium tetraurelia strain d4-2] |
| XP_016872218.1 | 533 | CAMK,CAMK-II,CAMKG | eukaryota>metazoa>chordata>vertebrata | Homo | sapiens | PREDICTED: calcium/calmodulin-dependent protein kinase type II subunit gamma isoform X25 [Homo sapiens] |
| XP_005263310.1 | 532 | CAMKD | eukaryota>metazoa>chordata>vertebrata | Homo | sapiens | PREDICTED: calcium/calmodulin-dependent protein kinase type II subunit delta isoform X1 [Homo sapiens] |
| XP_005263311.1 | 532 | CAMKD | eukaryota>metazoa>chordata>vertebrata | Homo | sapiens | PREDICTED: calcium/calmodulin-dependent protein kinase type II subunit delta isoform X2 [Homo sapiens] |
| XP_006500881.1 | 532 | 2810011D23Rik,8030469K03Rik,CaMK,II,[d]-CaMKII | eukaryota>metazoa>chordata>vertebrata | Mus | musculus | PREDICTED: calcium/calmodulin-dependent protein kinase type II subunit delta isoform X3 [Mus musculus] |
| XP_015305990.1 | 532 | CAMK2D | eukaryota>metazoa>chordata>vertebrata | Macaca | fascicularis | PREDICTED: calcium/calmodulin-dependent protein kinase type II subunit delta isoform X3 [Macaca fascicularis] |
| XP_015305991.1 | 532 | CAMK2D | eukaryota>metazoa>chordata>vertebrata | Macaca | fascicularis | PREDICTED: calcium/calmodulin-dependent protein kinase type II subunit delta isoform X4 [Macaca fascicularis] |
| XP_017174904.1 | 532 | 2810011D23Rik,8030469K03Rik,CaMK,II,[d]-CaMKII | eukaryota>metazoa>chordata>vertebrata | Mus | musculus | PREDICTED: calcium/calmodulin-dependent protein kinase type II subunit delta isoform X4 [Mus musculus] |
| XP_017306968.1 | 532 | LOC108255499 | eukaryota>metazoa>chordata>vertebrata>actinopterygii | Ictalurus | punctatus | PREDICTED: calcium/calmodulin-dependent protein kinase type II subunit beta-like isoform X20 [Ictalurus punctatus] |
| NP_001275322.1 | 532 | CDPK | eukaryota>viridiplantae | Solanum | tuberosum | calcium-dependent protein kinase [Solanum tuberosum] |
| XP_002983821.1 | 532 | CPK3-2 | eukaryota>viridiplantae | Selaginella | moellendorffii | calcium dependent protein kinase 3 [Selaginella moellendorffii] |
| XP_002989102.1 | 532 | CPK3-1 | eukaryota>viridiplantae | Selaginella | moellendorffii | calcium dependent protein kinase [Selaginella moellendorffii] |
| XP_006351684.1 | 532 | LOC102602736 | eukaryota>viridiplantae | Solanum | tuberosum | PREDICTED: calcium-dependent protein kinase 7-like [Solanum tuberosum] |
| XP_015166078.1 | 532 | LOC102602736 | eukaryota>viridiplantae | Solanum | tuberosum | PREDICTED: calcium-dependent protein kinase 7-like [Solanum tuberosum] |
| XP_016470886.1 | 532 | LOC107793116 | eukaryota>viridiplantae | Nicotiana | tabacum | PREDICTED: calcium-dependent protein kinase 34-like isoform X1 [Nicotiana tabacum] |
| XP_019077317.1 | 532 | LOC100258857 | eukaryota>viridiplantae | Vitis | vinifera | PREDICTED: calcium-dependent protein kinase 32 isoform X1 [Vitis vinifera] |
| XP_001434671.1 | 532 | GSPATT00006580001 | eukaryota>alveolata>ciliophora | Paramecium | tetraurelia strain d4-2 | hypothetical protein (macronuclear) [Paramecium tetraurelia strain d4-2] |
| XP_001438861.1 | 532 | GSPATT00008011001 | eukaryota>alveolata>ciliophora | Paramecium | tetraurelia strain d4-2 | hypothetical protein (macronuclear) [Paramecium tetraurelia strain d4-2] |
| XP_019075935.1 | 532 | LOC100267353 | eukaryota>viridiplantae | Vitis | vinifera | PREDICTED: calcium-dependent protein kinase 8 isoform X2 [Vitis vinifera] |
| NP_001162833.1 | 531 | Ca2+/calmodulin-dependent,protein,kinase,calcium/calmodulin-dependent,protein,kinase,II,Cam,CaM,CaM,II,CaM,KII,CaM,kinase,CaMK,II,CaMK-II,camkII,camKII,Camkii,CamKII,CA [illegible] Fb,l(4)16,l(4)ar | eukaryota>metazoa>hexapoda | Drosophila | melanogaster | Calcium/calmodulin-dependent protein kinase II, isoform J [Drosophila melanogaster] |
| NP_001258322.1 | 531 | camk2b,CAMK2Gb,fj90d04,si:dkey-174n20.2,wu:fj90d04,zgc:165373 | eukaryota>metazoa>chordata>vertebrata>actinopterygii | Danio | rerio | calcium/calmodulin-dependent protein kinase type II subunit beta [Danio rerio] |
| XP_016767053.1 | 531 | Camkii | eukaryota>metazoa>hexapoda | Apis | mellifera | PREDICTED: calcium/calmodulin-dependent protein kinase II isoform X18 [Apis mellifera] |
| XP_018654510.1 | 531 | Smp_011660.1 | eukaryota>metazoa | Schistosoma | mansoni | protein kinase [Schistosoma mansoni] |
| NP_001105306.1 | 531 | GRMZM2G168706 | eukaryota>viridiplantae | Zea | mays | calcium-dependent protein kinase [Zea mays] |
| NP_001319867.1 | 531 | AtCPK21,calcium-dependent,protein,kinase,21,T4B21.13,T4B21_13 | eukaryota>viridiplantae | Arabidopsis | thaliana | calcium-dependent protein kinase 21 [Arabidopsis thaliana] |
| NP_192381.1 | 531 | AtCPK21,calcium-dependent,protein,kinase,21,T4B21.13,T4B21_13 | eukaryota>viridiplantae | Arabidopsis | thaliana | calcium-dependent protein kinase 21 [Arabidopsis thaliana] |
| XP_002976252.1 | 531 | CPK13-1 | eukaryota>viridiplantae | Selaginella | moellendorffii | calcium dependent protein kinase 7 [Selaginella moellendorffii] |
| XP_002987149.1 | 531 | CPK13-2 | eukaryota>viridiplantae | Selaginella | moellendorffii | calcium dependent protein kinase [Selaginella moellendorffii] |
| [illegible] | [illegible] | [illegible] | [illegible] | [illegible] | [illegible] | [illegible] |

| | | | | | | |
|---|---|---|---|---|---|---|
| OMJ88622.1 | 496 | SteCoe_9402 | eukaryota>alveolata>ciliophora | Stentor | coeruleus | hypothetical protein SteCoe_9402 [Stentor coeruleus] |
| XP_001452952.1 | 496 | GSPATT00019272001 | eukaryota>alveolata>ciliophora | Paramecium | tetraurelia strain d4-2 | hypothetical protein (macronuclear) [Paramecium tetraurelia strain d4-2] |
| XP_001012485.2 | 496 | TTHERM_01052880 | eukaryota>alveolata>ciliophora | Tetrahymena | thermophila SB210 | calmodulin-domain kinase [Tetrahymena thermophila SB210] |
| XP_005646351.1 | 496 | COCSUDRAFT_29803 | eukaryota>viridiplantae>chlorophyta | Coccomyxa | subellipsoidea C-169 | Pkinase-domain-containing protein [Coccomyxa subellipsoidea C-169] |
| NP_001034228.1 | 495 | Camkg | eukaryota>metazoa>chordata>vertebrata | Mus | musculus | calcium/calmodulin-dependent protein kinase type II subunit gamma isoform 3 [Mus musculus] |
| NP_001213.2 | 495 | CAMK,CAMK-II,CAMKG | eukaryota>metazoa>chordata>vertebrata | Homo | sapiens | calcium/calmodulin-dependent protein kinase type II subunit gamma isoform 4 [Homo sapiens] |
| XP_003978131.2 | 495 | LOC101071995 | eukaryota>metazoa>chordata>vertebrata>actinopterygii | Takifugu | rubripes | PREDICTED: calcium/calmodulin-dependent protein kinase type II delta chain-like [Takifugu rubripes] |
| XP_005565537.1 | 495 | CAMK2G | eukaryota>metazoa>chordata>vertebrata | Macaca | fascicularis | PREDICTED: calcium/calmodulin-dependent protein kinase type II subunit gamma isoform X18 [Macaca fascicularis] |
| XP_006560593.1 | 495 | Camkii | eukaryota>metazoa>hexapoda | Apis | mellifera | PREDICTED: calcium/calmodulin-dependent protein kinase II isoform X30 [Apis mellifera] |
| XP_015833952.1 | 495 | LOC655266 | eukaryota>metazoa>hexapoda | Tribolium | castaneum | PREDICTED: calcium/calmodulin-dependent protein kinase type II alpha chain isoform X15 [Tribolium castaneum] |
| NP_174807.1 | 495 | ATCDPK2,ATCPK11,calcium-dependent,protein,kinase,2,CALCIUM-DEPENDENT,PROTEIN,KINASE,2,CPK11,F15O4.8 | eukaryota>viridiplantae | Arabidopsis | thaliana | calcium-dependent protein kinase 2 [Arabidopsis thaliana] |
| XP_001699889.1 | 495 | CHLREDRAFT_126628 | eukaryota>viridiplantae>chlorophyta | Chlamydomonas | reinhardtii | predicted protein, partial [Chlamydomonas reinhardtii] |
| EJY79908.1 | 495 | OXYTRI_22813 | eukaryota>alveolata>ciliophora | Oxytricha | trifallax | Protein kinase domain containing protein (macronuclear) [Oxytricha trifallax] |
| OMJ68897.1 | 495 | SteCoe_33514 | eukaryota>alveolata>ciliophora | Stentor | coeruleus | hypothetical protein SteCoe_33514 [Stentor coeruleus] |
| OMJ88179.1 | 495 | SteCoe_9912 | eukaryota>alveolata>ciliophora | Stentor | coeruleus | hypothetical protein SteCoe_9912 [Stentor coeruleus] |
| XP_002777779.1 | 495 | Pmar_PMAR008714 | eukaryota>alveolata | Perkinsus | marinus ATCC 50983 | Calcium-dependent protein kinase, putative [Perkinsus marinus ATCC 50983] |
| XP_002779826.1 | 495 | Pmar_PMAR009835 | eukaryota>alveolata | Perkinsus | marinus ATCC 50983 | Calcium-dependent protein kinase, putative [Perkinsus marinus ATCC 50983] |
| XP_002781466.1 | 495 | Pmar_PMAR029698 | eukaryota>alveolata | Perkinsus | marinus ATCC 50983 | Calcium-dependent protein kinase, putative [Perkinsus marinus ATCC 50983] |
| OMJ85303.1 | 495 | SteCoe_13397 | eukaryota>alveolata>ciliophora | Stentor | coeruleus | hypothetical protein SteCoe_13397 [Stentor coeruleus] |
| OMJ90109.1 | 495 | SteCoe_7597 | eukaryota>alveolata>ciliophora | Stentor | coeruleus | hypothetical protein SteCoe_7597 [Stentor coeruleus] |
| OMJ74649.1 | 495 | SteCoe_26394 | eukaryota>alveolata>ciliophora | Stentor | coeruleus | hypothetical protein SteCoe_26394 [Stentor coeruleus] |
| OMJ79795.1 | 495 | SteCoe_20095 | eukaryota>alveolata>ciliophora | Stentor | coeruleus | hypothetical protein SteCoe_20095 [Stentor coeruleus] |
| OMJ85801.1 | 495 | SteCoe_12794 | eukaryota>alveolata>ciliophora | Stentor | coeruleus | hypothetical protein SteCoe_12794 [Stentor coeruleus] |
| OMJ68694.1 | 495 | SteCoe_33780 | eukaryota>alveolata>ciliophora | Stentor | coeruleus | hypothetical protein SteCoe_33780 [Stentor coeruleus] |
| XP_002615124.1 | 495 | CLUG_05139 | eukaryota>fungi>ascomycota | Clavispora | lusitaniae ATCC 42720 | hypothetical protein CLUG_05139 [Clavispora lusitaniae ATCC 42720] |
| XP_009037159.1 | 495 | AURANDRAFT_26376 | eukaryota>stramenopiles | Aureococcus | anophagefferens | hypothetical protein AURANDRAFT_26376 [Aureococcus anophagefferens] |
| XP_005249921.1 | 494 | CAM2,CAMK2,CAMKB | eukaryota>metazoa>chordata>vertebrata | Homo | sapiens | PREDICTED: calcium/calmodulin-dependent protein kinase type II subunit beta isoform X20 [Homo sapiens] |
| XP_011142947.1 | 494 | LOC105185279 | eukaryota>metazoa>hexapoda | Harpegnathos | saltator | PREDICTED: calcium/calmodulin-dependent protein kinase type II alpha chain isoform X13 [Harpegnathos saltator] |
| XP_015131771.1 | 494 | CAMK2D | eukaryota>metazoa>chordata>vertebrata | Gallus | gallus | PREDICTED: calcium/calmodulin-dependent protein kinase type II delta chain isoform X7 [Gallus gallus] |
| XP_015143807.1 | 494 | CAMK2G | eukaryota>metazoa>chordata>vertebrata | Gallus | gallus | PREDICTED: calcium/calmodulin-dependent protein kinase type II subunit gamma isoform X16 [Gallus gallus] |
| XP_016864164.1 | 494 | CAMKD | eukaryota>metazoa>chordata>vertebrata | Homo | sapiens | PREDICTED: calcium/calmodulin-dependent protein kinase type II subunit delta isoform X9 [Homo sapiens] |
| XP_016864165.1 | 494 | CAMKD | eukaryota>metazoa>chordata>vertebrata | Homo | sapiens | PREDICTED: calcium/calmodulin-dependent protein kinase type II subunit delta isoform X9 [Homo sapiens] |
| XP_018080949.1 | 494 | LOC108696257 | eukaryota>metazoa>chordata>vertebrata | Xenopus | laevis | PREDICTED: calcium/calmodulin-dependent protein kinase type II subunit gamma-like isoform X19 [Xenopus laevis] |
| XP_018082324.1 | 494 | camk2,camk2g | eukaryota>metazoa>chordata>vertebrata | Xenopus | laevis | PREDICTED: calcium/calmodulin-dependent protein kinase (CaM kinase) II gamma S homeolog isoform X21 [Xenopus laevis] |
| XP_001781693.1 | 494 | cpk7 | eukaryota>viridiplantae | Physcomitrella | patens | cpk7 calcium-dependent protein kinase [Physcomitrella patens] |
| OMJ79435.1 | 494 | SteCoe_20536 | eukaryota>alveolata>ciliophora | Stentor | coeruleus | hypothetical protein SteCoe_20536 [Stentor coeruleus] |
| OMJ67272.1 | 494 | SteCoe_35610 | eukaryota>alveolata>ciliophora | Stentor | coeruleus | hypothetical protein SteCoe_35610 [Stentor coeruleus] |
| OMJ73324.1 | 494 | SteCoe_28008 | eukaryota>alveolata>ciliophora | Stentor | coeruleus | hypothetical protein SteCoe_28008 [Stentor coeruleus] |
| EJY85924.1 | 494 | OXYTRI_16088 | eukaryota>alveolata>ciliophora | Oxytricha | trifallax | Protein kinase domain containing protein (macronuclear) [Oxytricha trifallax] |
| CBJ27498.1 | 494 | CAMK | eukaryota>stramenopiles | Ectocarpus | siliculosus | Ca2+/calmodulin-dependent protein kinase, EF-Hand protein superfamily [Ectocarpus siliculosus] |
| XP_005768816.1 | 494 | EMIHUDRAFT_451551 | eukaryota>haptophyceae | Emiliania | huxleyi CCMP1516 | hypothetical protein EMIHUDRAFT_451551 [Emiliania huxleyi CCMP1516] |
| Bnat1000019719 | 494 | Bnat1000019719 | eukaryota>rhizaria | Bigelowiella | natans | >Bnat1000019719 |
| NP_001002542.1 | 493 | camk2d,camk2da,wu:fq17a12,zgc:92792 | eukaryota>metazoa>chordata>vertebrata>actinopterygii | Danio | rerio | calcium/calmodulin-dependent protein kinase type II delta 2 chain [Danio rerio] |
| XP_005156428.1 | 493 | camk2ga,zgc:109908 | eukaryota>metazoa>chordata>vertebrata>actinopterygii | Danio | rerio | calcium/calmodulin-dependent protein kinase type II subunit gamma isoform X15 [Danio rerio] |
| XP_009174101.1 | 493 | T265_09682 | eukaryota>metazoa | Opisthorchis | viverrini | hypothetical protein T265_09682 [Opisthorchis viverrini] |
| XP_015131772.1 | 493 | CAMK2D | eukaryota>metazoa>chordata>vertebrata | Gallus | gallus | PREDICTED: calcium/calmodulin-dependent protein kinase type II delta chain isoform X8 [Gallus gallus] |
| XP_015131773.1 | 493 | CAMK2D | eukaryota>metazoa>chordata>vertebrata | Gallus | gallus | PREDICTED: calcium/calmodulin-dependent protein kinase type II delta chain isoform X9 [Gallus gallus] |
| XP_016868154.1 | 493 | CAM2,CAMK2,CAMKB | eukaryota>metazoa>chordata>vertebrata | Homo | sapiens | PREDICTED: calcium/calmodulin-dependent protein kinase type II subunit beta isoform X21 [Homo sapiens] |
| XP_017327127.1 | 493 | LOC108267517 | eukaryota>metazoa>chordata>vertebrata>actinopterygii | Ictalurus | punctatus | PREDICTED: calcium/calmodulin-dependent protein kinase type II delta 1 chain isoform X5 [Ictalurus punctatus] |
| XP_017327128.1 | 493 | LOC108267517 | eukaryota>metazoa>chordata>vertebrata>actinopterygii | Ictalurus | punctatus | PREDICTED: calcium/calmodulin-dependent protein kinase type II delta 1 chain isoform X6 [Ictalurus punctatus] |
| XP_017327132.1 | 493 | LOC108267517 | eukaryota>metazoa>chordata>vertebrata>actinopterygii | Ictalurus | punctatus | PREDICTED: calcium/calmodulin-dependent protein kinase type II delta 1 chain isoform X10 [Ictalurus punctatus] |
| XP_017338908.1 | 493 | LOC108273829 | eukaryota>metazoa>chordata>vertebrata>actinopterygii | Ictalurus | punctatus | PREDICTED: calcium/calmodulin-dependent protein kinase type II subunit gamma isoform X13 [Ictalurus punctatus] |
| XP_001029735.2 | 493 | TTHERM_01333220 | eukaryota>alveolata>ciliophora | Tetrahymena | thermophila SB210 | calmodulin-domain kinase [Tetrahymena thermophila SB210] |
| XP_001442992.1 | 493 | GSPATT00011227001 | eukaryota>alveolata>ciliophora | Paramecium | tetraurelia strain d4-2 | hypothetical protein (macronuclear) [Paramecium tetraurelia strain d4-2] |
| EJY88048.1 | 493 | OXYTRI_21209 | eukaryota>alveolata>ciliophora | Oxytricha | trifallax | Protein kinase domain containing protein (macronuclear) [Oxytricha trifallax] |
| OMJ89974.1 | 493 | SteCoe_7762 | eukaryota>alveolata>ciliophora | Stentor | coeruleus | hypothetical protein SteCoe_7762 [Stentor coeruleus] |
| OMJ86407.1 | 493 | SteCoe_12063 | eukaryota>alveolata>ciliophora | Stentor | coeruleus | hypothetical protein SteCoe_12063 [Stentor coeruleus] |
| [illegible] | [illegible] | [illegible] | [illegible] | [illegible] | [illegible] | [illegible] |

| | | | | | | |
|---|---|---|---|---|---|---|
| XP_013780769.1 | 416 | LOC106465117 | eukaryota>metazoa | Limulus | polyphemus | serine/threonine-protein kinase H1 homolog [Limulus polyphemus] |
| CEM36746.1 | 416 | Vbra_3415 | eukaryota>alveolata | Vitrella | brassicaformis CCMP3155 | unnamed protein product [Vitrella brassicaformis CCMP3155] |
| NP_001167827.1 | 415 | GRMZM2G422576 | eukaryota>viridiplantae | Zea | mays | putative protein kinase superfamily protein [Zea mays] |
| XP_006579812.1 | 415 | LOC100803313 | eukaryota>viridiplantae | Glycine | max | PREDICTED: CDPK-related kinase 5-like isoform X2 [Glycine max] |
| CAG08692.1 | 414 | GSTEN:00029739:G:001 | eukaryota>metazoa>chordata>vertebrata>actinopterygii | Tetraodon | nigroviridis | unnamed protein product [Tetraodon nigroviridis] |
| XP_001539950.1 | 414 | HCAG_05417 | eukaryota>fungi>ascomycota | Histoplasma | capsulatum NAm1 | calcium/calmodulin-dependent protein kinase type I [Histoplasma capsulatum NAm1] |
| XP_015155457.1 | 414 | CAMK4L | eukaryota>metazoa>chordata>vertebrata | Gallus | gallus | PREDICTED: calcium/calmodulin-dependent protein kinase type IV-like isoform X1 [Gallus gallus] |
| XP_015155458.1 | 414 | CAMK4L | eukaryota>metazoa>chordata>vertebrata | Gallus | gallus | PREDICTED: calcium/calmodulin-dependent protein kinase type IV-like isoform X1 [Gallus gallus] |
| CBN79381.1 | 414 | Esi_0202_0009 | eukaryota>stramenopiles | Ectocarpus | siliculosus | conserved unknown protein [Ectocarpus siliculosus] |
| XP_001440154.1 | 414 | GSPATT00009149001 | eukaryota>alveolata>ciliophora | Paramecium | tetraurelia strain d4-2 | hypothetical protein (macronuclear) [Paramecium tetraurelia strain d4-2] |
| XP_012897343.1 | 414 | GSBLH_T00003190001 | eukaryota>stramenopiles | Blastocystis | hominis | uncharacterized protein [Blastocystis hominis] |
| XP_003967700.1 | 413 | LOC101070748 | eukaryota>metazoa>chordata>vertebrata>actinopterygii | Takifugu | rubripes | PREDICTED: calcium/calmodulin-dependent protein kinase type 1D-like [Takifugu rubripes] |
| XP_005265573.1 | 413 | CAMKI | eukaryota>metazoa>chordata>vertebrata | Homo | sapiens | PREDICTED: calcium/calmodulin-dependent protein kinase type 1 isoform X1 [Homo sapiens] |
| XP_011393243.1 | 413 | camk-1 | eukaryota>fungi>ascomycota | Neurospora | crassa OR74A | Ca/CaM-dependent kinase-1, variant 1 [Neurospora crassa OR74A] |
| XP_011393244.1 | 413 | camk-1 | eukaryota>fungi>ascomycota | Neurospora | crassa OR74A | Ca/CaM-dependent kinase-1 [Neurospora crassa OR74A] |
| XP_011393245.1 | 413 | camk-1 | eukaryota>fungi>ascomycota | Neurospora | crassa OR74A | Ca/CaM-dependent kinase-1, variant 2 [Neurospora crassa OR74A] |
| XP_012352361.1 | 413 | CAMK2B | eukaryota>metazoa>chordata>vertebrata | Nomascus | leucogenys | PREDICTED: calcium/calmodulin-dependent protein kinase type II subunit beta isoform X6 [Nomascus leucogenys] |
| XP_016868155.1 | 413 | CAM2,CAMK2,CAMKB | eukaryota>metazoa>chordata>vertebrata | Homo | sapiens | PREDICTED: calcium/calmodulin-dependent protein kinase type II subunit beta isoform X22 [Homo sapiens] |
| XP_009300936.2 | 413 | camk1a | eukaryota>metazoa>chordata>vertebrata>actinopterygii | Danio | rerio | PREDICTED: calcium/calmodulin-dependent protein kinase type 1-like isoform X2 [Danio rerio] |
| XP_005845710.1 | 413 | CHLNCDRAFT_32147 | eukaryota>viridiplantae>chlorophyta | Chlorella | variabilis | hypothetical protein CHLNCDRAFT_32147 [Chlorella variabilis] |
| XP_004936827.1 | 412 | MKNK1 | eukaryota>metazoa>chordata>vertebrata | Gallus | gallus | PREDICTED: MAP kinase-interacting serine/threonine-protein kinase 1 isoform X3 [Gallus gallus] |
| XP_012895897.1 | 412 | GSBLH_T00001952001 | eukaryota>stramenopiles | Blastocystis | hominis | uncharacterized protein [Blastocystis hominis] |
| XP_016516068.1 | 411 | LOC107832719 | eukaryota>viridiplantae | Nicotiana | tabacum | PREDICTED: calcium-dependent protein kinase 5-like [Nicotiana tabacum] |
| XP_018184414.1 | 410 | L228DRAFT_271757 | eukaryota>fungi>ascomycota | Xylona | heveae TC161 | CMK1 Ca2+/calmodulin-dependent protein kinase [Xylona heveae TC161] |
| XP_018734691.1 | 410 | CMK2 | eukaryota>fungi>ascomycota | Sugiyamaella | lignohabitans | calmodulin-dependent protein kinase CMK2 [Sugiyamaella lignohabitans] |
| XP_001750784.1 | 410 | MONBRDRAFT_39303 | eukaryota>choanoflagellida | Monosiga | brevicollis MX1 | hypothetical protein [Monosiga brevicollis MX1] |
| EFA77412.1 | 410 | PPL_12628 | eukaryota>amoebozoa>mycetozoa>dictyosteliida | Polysphondylium | pallidum PN500 | putative protein serine/threonine kinase [Polysphondylium pallidum PN500] |
| XP_660016.1 | 409 | AN2412.2 | eukaryota>fungi>ascomycota | Aspergillus | nidulans FGSC A4 | KCC1_EMENI Calcium/calmodulin-dependent protein kinase (CMPK) [Aspergillus nidulans FGSC A4] |
| Chet1000006811 | 409 | Chet1000006811 | eukaryota>fungi>ascomycota | Cochliobolus | heterostrophus | >Chet1000006811 |
| XP_009843353.1 | 408 | H257_16578 | eukaryota>stramenopiles | Aphanomyces | astaci | CAMK/CAMK1 protein kinase [Aphanomyces astaci] |
| XP_013424048.1 | 408 | M436DRAFT_54633 | eukaryota>fungi>ascomycota | Aureobasidium | namibiae CBS 147.97 | Ca2+/calmodulin-dependent protein kinase [Aureobasidium namibiae CBS 147.97] |
| XP_001448757.1 | 408 | GSPATT00016202001 | eukaryota>alveolata>ciliophora | Paramecium | tetraurelia strain d4-2 | hypothetical protein (macronuclear) [Paramecium tetraurelia strain d4-2] |
| XP_006681018.1 | 408 | BATDEDRAFT_17461 | eukaryota>fungi>chytridiomycota | Batrachochytrium | dendrobatidis JAM81 | hypothetical protein BATDEDRAFT_17461 [Batrachochytrium dendrobatidis JAM81] |
| XP_013761964.1 | 408 | AMSG_01358 | eukaryota>apusozoa>apusomonadidae | Thecamonas | trahens ATCC 50062 | CAMK/CAMK1 protein kinase [Thecamonas trahens ATCC 50062] |
| XP_009843349.1 | 407 | H257_16578 | eukaryota>stramenopiles | Aphanomyces | astaci | CAMK/CAMK1 protein kinase, variant 1 [Aphanomyces astaci] |
| XP_008878422.1 | 407 | H310_12916 | eukaryota>stramenopiles | Aphanomyces | invadans | CAMK/CAMK1 protein kinase [Aphanomyces invadans] |
| ELR16696.1 | 407 | ACA1_089840 | eukaryota>amoebozoa>acanthamoebidae | Acanthamoeba | castellanii str. Neff | calcium/calmodulindependent protein kinase [Acanthamoeba castellanii str. Neff] |
| OHT10759.1 | 407 | TRFO_19894 | eukaryota>parabasalia | Tritrichomonas | foetus | CAMK family protein kinase [Tritrichomonas foetus] |
| XP_009843333.1 | 406 | H257_16565 | eukaryota>stramenopiles | Aphanomyces | astaci | CAMK/CAMK1 protein kinase [Aphanomyces astaci] |
| XP_014532050.1 | 406 | PDIP_78930 | eukaryota>fungi>ascomycota | Penicillium | digitatum Pd1 | Calcium/calmodulin-dependent protein kinase, putative [Penicillium digitatum Pd1] |
| XP_017306206.1 | 406 | camkv | eukaryota>metazoa>chordata>vertebrata>actinopterygii | Ictalurus | punctatus | PREDICTED: caM kinase-like vesicle-associated protein [Ictalurus punctatus] |
| XP_005703958.1 | 406 | Gasu_50310 | eukaryota>rhodophyta | Galdieria | sulphuraria | serine/threonine protein kinase [Galdieria sulphuraria] |
| NP_524622.1 | 405 | Calcium/calmodulin-dependent,protein,kinase,CaMK1,camKI,caMKI,CamKI,CG1495,dCKI,Dmel\CG1495 | eukaryota>metazoa>hexapoda | Drosophila | melanogaster | Calcium/calmodulin-dependent protein kinase I, isoform G [Drosophila melanogaster] |
| NP_726569.1 | 405 | Calcium/calmodulin-dependent,protein,kinase,CaMK1,camKI,caMKI,CamKI,CG1495,dCKI,Dmel\CG1495 | eukaryota>metazoa>hexapoda | Drosophila | melanogaster | Calcium/calmodulin-dependent protein kinase I, isoform A [Drosophila melanogaster] |
| NP_726570.1 | 405 | Calcium/calmodulin-dependent,protein,kinase,CaMK1,camKI,caMKI,CamKI,CG1495,dCKI,Dmel\CG1495 | eukaryota>metazoa>hexapoda | Drosophila | melanogaster | Calcium/calmodulin-dependent protein kinase I, isoform B [Drosophila melanogaster] |
| XP_008620433.1 | 405 | SDRG_16022 | eukaryota>stramenopiles | Saprolegnia | diclina VS20 | CAMK/CAMK1 protein kinase [Saprolegnia diclina VS20] |
| XP_012198712.1 | 405 | SPRG_04395 | eukaryota>stramenopiles | Saprolegnia | parasitica CBS 223.65 | CAMK/CAMK1 protein kinase [Saprolegnia parasitica CBS 223.65] |
| XP_009830637.1 | 405 | H257_06948 | eukaryota>stramenopiles | Aphanomyces | astaci | CAMK/CAMK1 protein kinase [Aphanomyces astaci] |
| XP_014531764.1 | 404 | PDIP_76070 | eukaryota>fungi>ascomycota | Penicillium | digitatum Pd1 | hypothetical protein PDIP_76070 [Penicillium digitatum Pd1] |
| XP_660669.1 | 404 | AN3065.2 | eukaryota>fungi>ascomycota | Aspergillus | nidulans FGSC A4 | hypothetical protein AN3065.2 [Aspergillus nidulans FGSC A4] |
| XP_019026599.1 | 404 | SAICODRAFT_64097 | eukaryota>fungi>ascomycota | Saitoella | complicata NRRL Y-17804 | Pkinase-domain-containing protein [Saitoella complicata NRRL Y-17804] |
| XP_006500894.1 | 404 | 2810011D23Rik,8030469K03Rik,CaMK,II,[d]-CaMKII | eukaryota>metazoa>chordata>vertebrata | Mus | musculus | PREDICTED: calcium/calmodulin-dependent protein kinase type II subunit delta isoform X19 [Mus musculus] |
| XP_011530594.1 | 404 | CAMKD | eukaryota>metazoa>chordata>vertebrata | Homo | sapiens | PREDICTED: calcium/calmodulin-dependent protein kinase type II subunit delta isoform X14 [Homo sapiens] |
| XP_015306010.1 | 404 | CAMK2D | eukaryota>metazoa>chordata>vertebrata | Macaca | fascicularis | PREDICTED: calcium/calmodulin-dependent protein kinase type II subunit delta isoform X22 [Macaca fascicularis] |
| XP_008896367.1 | 404 | PPTG_04245 | eukaryota>stramenopiles | Phytophthora | parasitica INRA-310 | CAMK/CAMK1 protein kinase [Phytophthora parasitica INRA-310] |
| XP_017177432.1 | 404 | 4921505G21Rik,AW061264 | eukaryota>metazoa>chordata>vertebrata | Mus | musculus | PREDICTED: serine/threonine-protein kinase 33 isoform X5 [Mus musculus] |
| XP_012208773.1 | 403 | SPRG_14268 | eukaryota>stramenopiles | Saprolegnia | parasitica CBS 223.65 | CAMK/CAMK1 protein kinase [Saprolegnia parasitica CBS 223.65] |
| XP_016513616.1 | 402 | LOC107830535 | eukaryota>viridiplantae | Nicotiana | tabacum | PREDICTED: calcium-dependent protein kinase 20-like [Nicotiana tabacum] |
| XP_006500895.1 | 401 | 2810011D23Rik,8030469K03Rik,CaMK,II,[d]-CaMKII | eukaryota>metazoa>chordata>vertebrata | Mus | musculus | PREDICTED: calcium/calmodulin-dependent protein kinase type II subunit delta isoform X20 [Mus musculus] |
| XP_015307261.1 | 401 | CAMK4 | eukaryota>metazoa>chordata>vertebrata | Macaca | fascicularis | PREDICTED: calcium/calmodulin-dependent protein kinase type IV isoform X2 [Macaca fascicularis] |
| ELR13755.1 | 400 | ACA1_014360 | eukaryota>amoebozoa>acanthamoebidae | Acanthamoeba | castellanii str. Neff | calcium/calmodulindependent protein kinase [Acanthamoeba castellanii str. Neff] |
| XP_001449552.1 | 400 | GSPATT00002284001 | eukaryota>alveolata>ciliophora | Paramecium | tetraurelia strain d4-2 | hypothetical protein (macronuclear) [Paramecium tetraurelia strain d4-2] |
| Chet1000007729 | 399 | Chet1000007729 | eukaryota>fungi>ascomycota | Cochliobolus | heterostrophus | >Chet1000007729 |
| XP_008878421.1 | 399 | H310_12916 | eukaryota>stramenopiles | Aphanomyces | invadans | CAMK/CAMK1 protein kinase, variant 1 [Aphanomyces invadans] |
| [illegible] | | | | | | |

| | | | | | | |
|---|---|---|---|---|---|---|
| XP_009830617.1 | 125 | H257_06931 | eukaryota>stramenopiles | Aphanomyces | astaci | hypothetical protein H257_06931 [Aphanomyces astaci] |
| Chet1000006047 | 125 | Chet1000006047 | eukaryota>fungi>ascomycota | Cochliobolus | heterostrophus | >Chet1000006047 |
| XP_002289889.1 | 124 | THAPSDRAFT_18469 | eukaryota>stramenopiles | Thalassiosira | pseudonana CCMP1335 | predicted protein, partial [Thalassiosira pseudonana CCMP1335] |
| XP_002508429.1 | 124 | MICPUN_85736 | eukaryota>viridiplantae>chlorophyta | Micromonas | commoda | predicted protein [Micromonas commoda] |
| XP_005763244.1 | 124 | EMIHUDRAFT_78676 | eukaryota>haptophyceae | Emiliania | huxleyi CCMP1516 | FKBP-type peptidyl-prolyl cis-trans isomerase, partial [Emiliania huxleyi CCMP1516] |
| XP_009037672.1 | 124 | AURANDRAFT_16370 | eukaryota>stramenopiles | Aureococcus | anophagefferens | hypothetical protein AURANDRAFT_16370, partial [Aureococcus anophagefferens] |
| XP_013759174.1 | 124 | AMSG_03924 | eukaryota>apusozoa>apusomonadidae | Thecamonas | trahens ATCC 50062 | peptidyl-prolyl cis-trans isomerase [Thecamonas trahens ATCC 50062] |
| XP_018191044.1 | 124 | L228DRAFT_265958 | eukaryota>fungi>ascomycota | Xylona | heveae TC161 | peptidyl-prolyl cis-trans isomerase [Xylona heveae TC161] |
| XP_710751.1 | 124 | RBP1 | eukaryota>fungi>ascomycota | Candida | albicans SC5314 | peptidylprolyl isomerase [Candida albicans SC5314] |
| Bnat1000011072 | 124 | Bnat1000011072 | eukaryota>rhizaria | Bigelowiella | natans | >Bnat1000011072 |
| Pram1000007139 | 124 | Pram1000007139 | eukaryota>stramenopiles | Phytophthora | ramorum | >Pram1000007139 |
| ABP00115.1 | 123 | FBP12 | eukaryota>viridiplantae>chlorophyta | Ostreococcus | lucimarinus CCE9901 | peptidyl-prolyl cis-trans isomerase, FKBP-type [Ostreococcus lucimarinus CCE9901] |
| XP_003055776.1 | 123 | MICPUCDRAFT_8507 | eukaryota>viridiplantae>chlorophyta | Micromonas | pusilla CCMP1545 | predicted protein, partial [Micromonas pusilla CCMP1545] |
| Bnat1000014841 | 123 | Bnat1000014841 | eukaryota>rhizaria | Bigelowiella | natans | >Bnat1000014841 |
| KPI88222.1 | 122 | ABL78_2726 | eukaryota>euglenozoa>kinetoplastida | Leptomonas | seymouri | putative peptidyl-prolyl cis-trans isomerase [Leptomonas seymouri] |
| XP_002508612.1 | 122 | MICPUN_62181 | eukaryota>viridiplantae>chlorophyta | Micromonas | commoda | predicted protein [Micromonas commoda] |
| XP_008908615.1 | 122 | PPTG_13900 | eukaryota>stramenopiles | Phytophthora | parasitica INRA-310 | hypothetical protein PPTG_13900 [Phytophthora parasitica INRA-310] |
| XP_011127622.1 | 122 | AOL_s00215g118 | eukaryota>fungi>ascomycota | Arthrobotrys | oligospora ATCC 24927 | hypothetical protein AOL_s00215g118 [Arthrobotrys oligospora ATCC 24927] |
| XP_014537689.1 | 122 | PDIP_15200 | eukaryota>fungi>ascomycota | Penicillium | digitatum Pd1 | FKBP-type peptidyl-prolyl isomerase, putative [Penicillium digitatum Pd1] |
| Bnat1000005560 | 122 | Bnat1000005560 | eukaryota>rhizaria | Bigelowiella | natans | >Bnat1000005560 |
| XP_005771612.1 | 121 | EMIHUDRAFT_46588 | eukaryota>haptophyceae | Emiliania | huxleyi CCMP1516 | FKBP-type peptidyl-prolyl cis-trans isomerase, partial [Emiliania huxleyi CCMP1516] |
| XP_005833559.1 | 121 | GUITHDRAFT_175442 | eukaryota>cryptophyta | Guillardia | theta CCMP2712 | FKBP-type peptidyl-prolyl cis-trans isomerase [Guillardia theta CCMP2712] |
| XP_009039102.1 | 121 | AURANDRAFT_6781 | eukaryota>stramenopiles | Aureococcus | anophagefferens | hypothetical protein AURANDRAFT_6781, partial [Aureococcus anophagefferens] |
| XP_014536553.1 | 121 | PDIP_30920 | eukaryota>fungi>ascomycota | Penicillium | digitatum Pd1 | FK506-binding protein 1B [Penicillium digitatum Pd1] |
| ORX94734.1 | 120 | K493DRAFT_324755 | eukaryota>fungi | Basidiobolus | meristosporus CBS 931.73 | peptidyl-prolyl cis-trans isomerase [Basidiobolus meristosporus CBS 931.73] |
| CEO95487.1 | 120 | PBRA_004213 | eukaryota>rhizaria>cercozoa>plasmodiophorida | Plasmodiophora | brassicae | hypothetical protein PBRA_004213 [Plasmodiophora brassicae] |
| EST42124.1 | 120 | SS50377_18432 | eukaryota>diplomonadida | Spironucleus | salmonicida | FKBP-type peptidyl-prolyl cis-trans isomerase [Spironucleus salmonicida] |
| XP_003383441.1 | 120 | LOC100638671 | eukaryota>metazoa | Amphimedon | queenslandica | PREDICTED: peptidyl-prolyl cis-trans isomerase FKBP12-like [Amphimedon queenslandica] |
| XP_005781001.1 | 120 | EMIHUDRAFT_58004 | eukaryota>haptophyceae | Emiliania | huxleyi CCMP1516 | FKBP-type peptidyl-prolyl cis-trans isomerase, partial [Emiliania huxleyi CCMP1516] |
| XP_006675582.1 | 120 | BATDEDRAFT_22495 | eukaryota>fungi>chytridiomycota | Batrachochytrium | dendrobatidis JAM81 | hypothetical protein BATDEDRAFT_22495 [Batrachochytrium dendrobatidis JAM81] |
| XP_014528683.1 | 120 | JH06_1212 | eukaryota>stramenopiles | Blastocystis | sp. subtype 4 | FKPB-type peptidyl-prolyl cis-trans isomerase [Blastocystis sp. subtype 4] |
| OLQ01587.1 | 119 | FKBP12 | eukaryota>alveolata | Symbiodinium | microadriaticum | Peptidyl-prolyl cis-trans isomerase FKBP12 [Symbiodinium microadriaticum] |
| XP_013428097.1 | 119 | M436DRAFT_63031 | eukaryota>fungi>ascomycota | Aureobasidium | namibiae CBS 147.97 | FKBP peptidyl-prolyl cis-trans isomerase [Aureobasidium namibiae CBS 147.97] |
| XP_013772828.1 | 119 | LOC106457927 | eukaryota>metazoa | Limulus | polyphemus | peptidyl-prolyl cis-trans isomerase FKBP12-like [Limulus polyphemus] |
| XP_014156216.1 | 119 | SARC_05406 | eukaryota>ichthyosporea | Sphaeroforma | arctica JP610 | peptidyl-prolyl isomerase FKBP12 [Sphaeroforma arctica JP610] |
| CEM25102.1 | 118 | Vbra_3277 | eukaryota>alveolata | Vitrella | brassicaformis CCMP3155 | unnamed protein product [Vitrella brassicaformis CCMP3155] |
| XP_002109901.1 | 118 | TRIADDRAFT_21313 | eukaryota>metazoa>placozoa | Trichoplax | adhaerens | hypothetical protein TRIADDRAFT_21313 [Trichoplax adhaerens] |
| XP_002287222.1 | 118 | THAPSDRAFT_32093 | eukaryota>stramenopiles | Thalassiosira | pseudonana CCMP1335 | predicted protein, partial [Thalassiosira pseudonana CCMP1335] |
| XP_002898522.1 | 118 | PITG_14620 | eukaryota>stramenopiles | Phytophthora | infestans T30-4 | FKBP-type peptidyl-prolyl cis-trans isomerase, putative [Phytophthora infestans T30-4] |
| XP_005767562.1 | 118 | EMIHUDRAFT_445648 | eukaryota>haptophyceae | Emiliania | huxleyi CCMP1516 | hypothetical protein EMIHUDRAFT_445648 [Emiliania huxleyi CCMP1516] |
| XP_009034527.1 | 118 | AURANDRAFT_22069 | eukaryota>stramenopiles | Aureococcus | anophagefferens | hypothetical protein AURANDRAFT_22069 [Aureococcus anophagefferens] |
| XP_009042283.1 | 118 | AURANDRAFT_34605 | eukaryota>stramenopiles | Aureococcus | anophagefferens | hypothetical protein AURANDRAFT_34605 [Aureococcus anophagefferens] |
| XP_009530593.1 | 118 | PHYSODRAFT_513680 | eukaryota>stramenopiles | Phytophthora | sojae | hypothetical protein PHYSODRAFT_513680 [Phytophthora sojae] |
| Smed1000010218 | 118 | Smed1000010218 | eukaryota>metazoa>platyhelminthe | Schmidtea | mediterranea | >Smed1000010218 |
| EPB90228.1 | 117 | HMPREF1544_02913 | eukaryota>fungi | Mucor | circinelloides f. circinelloides 1006PhL | FK506-binding protein 1A [Mucor circinelloides f. circinelloides 1006PhL] |
| XP_001767064.1 | 117 | PHYPADRAFT_49268 | eukaryota>viridiplantae | Physcomitrella | patens | predicted protein, partial [Physcomitrella patens] |
| XP_002668164.1 | 117 | NAEGRDRAFT_82377 | eukaryota>heterolobosea | Naegleria | gruberi strain NEG-M | predicted protein [Naegleria gruberi strain NEG-M] |
| XP_002677078.1 | 117 | NAEGRDRAFT_83142 | eukaryota>heterolobosea | Naegleria | gruberi strain NEG-M | peptidylprolyl isomerase [Naegleria gruberi strain NEG-M] |
| XP_005762909.1 | 117 | EMIHUDRAFT_57979 | eukaryota>haptophyceae | Emiliania | huxleyi CCMP1516 | FKBP-type peptidyl-prolyl cis-trans isomerase, partial [Emiliania huxleyi CCMP1516] |
| XP_008604047.1 | 117 | SDRG_00353 | eukaryota>stramenopiles | Saprolegnia | diclina VS20 | hypothetical protein SDRG_00353 [Saprolegnia diclina VS20] |
| XP_012206928.1 | 117 | SPRG_11357 | eukaryota>stramenopiles | Saprolegnia | parasitica CBS 223.65 | hypothetical protein SPRG_11357 [Saprolegnia parasitica CBS 223.65] |
| Bnat1000021493 | 117 | Bnat1000021493 | eukaryota>rhizaria | Bigelowiella | natans | >Bnat1000021493 |
| XP_006454955.1 | 116 | AGABI2DRAFT_190126 | eukaryota>fungi>basidiomycota | Agaricus | bisporus var. bisporus H97 | hypothetical protein AGABI2DRAFT_190126 [Agaricus bisporus var. bisporus H97] |
| OHS94446.1 | 116 | TRFO_39354 | eukaryota>parabasalia | Tritrichomonas | foetus | peptidyl-prolyl cis-trans isomerase [Tritrichomonas foetus] |
| OLQ13598.1 | 116 | FKBP12 | eukaryota>alveolata | Symbiodinium | microadriaticum | Peptidyl-prolyl cis-trans isomerase FKBP12 [Symbiodinium microadriaticum] |
| XP_001224999.1 | 116 | CHGG_07343 | eukaryota>fungi>ascomycota | Chaetomium | globosum CBS 148.51 | hypothetical protein CHGG_07343 [Chaetomium globosum CBS 148.51] |
| XP_001568657.1 | 116 | LBRM_35_0310 | eukaryota>euglenozoa>kinetoplastida | Leishmania | braziliensis MHOM/BR/75/M2904 | putative peptidyl-prolyl cis-trans isomerase [Leishmania braziliensis MHOM/BR/75/M2904] |
| XP_002184598.1 | 116 | PHATRDRAFT_30615 | eukaryota>stramenopiles | Phaeodactylum | tricornutum CCAP 1055/1 | predicted protein [Phaeodactylum tricornutum CCAP 1055/1] |
| XP_002289399.1 | 116 | THAPSDRAFT_33855 | eukaryota>stramenopiles | Thalassiosira | pseudonana CCMP1335 | predicted protein, partial [Thalassiosira pseudonana CCMP1335] |
| XP_002782377.1 | 116 | Pmar_PMAR029236 | eukaryota>alveolata | Perkinsus | marinus ATCC 50983 | Peptidyl-prolyl isomerase FKBP12, putative [Perkinsus marinus ATCC 50983] |
| XP_005820108.1 | 116 | GUITHDRAFT_81737 | eukaryota>cryptophyta | Guillardia | theta CCMP2712 | hypothetical protein GUITHDRAFT_81737 [Guillardia theta CCMP2712] |
| XP_007332301.1 | 116 | AGABI1DRAFT_115445 | eukaryota>fungi>basidiomycota | Agaricus | bisporus var. burnettii JB137-S8 | hypothetical protein AGABI1DRAFT_115445 [Agaricus bisporus var. burnettii JB137-S8] |
| XP_014668839.1 | 116 | LOC106810080 | eukaryota>metazoa | Priapulus | caudatus | PREDICTED: peptidyl-prolyl cis-trans isomerase FKBP12-like [Priapulus caudatus] |
| NP_001155807.1 | 115 | LOC100168864 | eukaryota>metazoa>hexapoda | Acyrthosiphon | pisum | peptidyl-prolyl cis-trans isomerase FKBP1A-like precursor [Acyrthosiphon pisum] |
| [illegible] | | | | | | |

[illegible]

| | | | | | | |
|---|---|---|---|---|---|---|
| XP_008643437.1 | 543 | GRMZM2G080083 | eukaryota>viridiplantae | Zea | mays | serine/threonine protein phosphatase 2A regulatory subunit B''alpha isoform X1 [Zea mays] |
| XP_008643438.1 | 543 | GRMZM2G080083 | eukaryota>viridiplantae | Zea | mays | serine/threonine protein phosphatase 2A regulatory subunit B''alpha isoform X1 [Zea mays] |
| XP_001438471.1 | 541 | GSPATT00000786001 | eukaryota>alveolata>ciliophora | Paramecium | tetraurelia strain d4-2 | hypothetical protein (macronuclear) [Paramecium tetraurelia strain d4-2] |
| XP_002966764.1 | 541 | SELMODRAFT_85336 | eukaryota>viridiplantae | Selaginella | moellendorffii | hypothetical protein SELMODRAFT_85336 [Selaginella moellendorffii] |
| XP_003536224.1 | 541 | LOC100793000 | eukaryota>viridiplantae | Glycine | max | PREDICTED: serine/threonine protein phosphatase 2A regulatory subunit B''beta-like [Glycine max] |
| EST46160.1 | 540 | SS50377_13752 | eukaryota>diplomonadida | Spironucleus | salmonicida | Serine/threonine-protein phosphatase 2A [Spironucleus salmonicida] |
| KPI86584.1 | 540 | ABL78_4359 | eukaryota>euglenozoa>kinetoplastida | Leptomonas | seymouri | hypothetical protein ABL78_4359 [Leptomonas seymouri] |
| XP_006345897.1 | 540 | LOC102590984 | eukaryota>viridiplantae | Solanum | tuberosum | PREDICTED: serine/threonine protein phosphatase 2A regulatory subunit B''beta-like isoform X1 [Solanum tuberosum] |
| XP_008664881.1 | 540 | LOC103643458 | eukaryota>viridiplantae | Zea | mays | probable serine/threonine protein phosphatase 2A regulatory subunit B''delta isoform X1 [Zea mays] |
| Bnat1000013915 | 540 | Bnat1000013915 | eukaryota>rhizaria | Bigelowiella | natans | >Bnat1000013915 Bnat1000013915 |
| EPY20970.1 | 539 | STCU_08752 | eukaryota>euglenozoa>kinetoplastida | Strigomonas | culicis | protein phosphatase 2 (formerly 2A), regulatory subunit B'' [Strigomonas culicis] |
| XP_002272394.1 | 539 | LOC100246385 | eukaryota>viridiplantae | Vitis | vinifera | PREDICTED: serine/threonine protein phosphatase 2A regulatory subunit B''beta isoform X3 [Vitis vinifera] |
| XP_006345898.1 | 539 | LOC102590984 | eukaryota>viridiplantae | Solanum | tuberosum | PREDICTED: serine/threonine protein phosphatase 2A regulatory subunit B''beta-like isoform X2 [Solanum tuberosum] |
| XP_006361409.1 | 539 | LOC102584799 | eukaryota>viridiplantae | Solanum | tuberosum | PREDICTED: serine/threonine protein phosphatase 2A regulatory subunit B''beta-like [Solanum tuberosum] |
| XP_010660207.1 | 539 | LOC100250118 | eukaryota>viridiplantae | Vitis | vinifera | PREDICTED: serine/threonine protein phosphatase 2A regulatory subunit B''beta [Vitis vinifera] |
| XP_016467583.1 | 539 | LOC107790195 | eukaryota>viridiplantae | Nicotiana | tabacum | PREDICTED: serine/threonine protein phosphatase 2A regulatory subunit B''beta-like [Nicotiana tabacum] |
| XP_016490243.1 | 539 | LOC107810035 | eukaryota>viridiplantae | Nicotiana | tabacum | PREDICTED: serine/threonine protein phosphatase 2A regulatory subunit B''beta-like [Nicotiana tabacum] |
| XP_016490244.1 | 539 | LOC107810035 | eukaryota>viridiplantae | Nicotiana | tabacum | PREDICTED: serine/threonine protein phosphatase 2A regulatory subunit B''beta-like [Nicotiana tabacum] |
| XP_016490245.1 | 539 | LOC107810035 | eukaryota>viridiplantae | Nicotiana | tabacum | PREDICTED: serine/threonine protein phosphatase 2A regulatory subunit B''beta-like [Nicotiana tabacum] |
| XP_016514670.1 | 539 | LOC107831426 | eukaryota>viridiplantae | Nicotiana | tabacum | PREDICTED: serine/threonine protein phosphatase 2A regulatory subunit B''beta-like [Nicotiana tabacum] |
| XP_016514671.1 | 539 | LOC107831426 | eukaryota>viridiplantae | Nicotiana | tabacum | PREDICTED: serine/threonine protein phosphatase 2A regulatory subunit B''beta-like [Nicotiana tabacum] |
| XP_016514672.1 | 539 | LOC107831426 | eukaryota>viridiplantae | Nicotiana | tabacum | PREDICTED: serine/threonine protein phosphatase 2A regulatory subunit B''beta-like [Nicotiana tabacum] |
| XP_018284469.1 | 539 | PHYBLDRAFT_70884 | eukaryota>fungi | Phycomyces | blakesleeanus NRRL 1555(-) | hypothetical protein PHYBLDRAFT_70884 [Phycomyces blakesleeanus NRRL 1555(-)] |
| CCW69367.1 | 538 | GSHART1_T00006588001 | eukaryota>euglenozoa>kinetoplastida | Phytomonas | sp. isolate Hart1 | unnamed protein product [Phytomonas sp. isolate Hart1] |
| NP_199222.1 | 538 | AT5G44090 | eukaryota>viridiplantae | Arabidopsis | thaliana | Calcium-binding EF-hand family protein [Arabidopsis thaliana] |
| XP_015774277.1 | 538 | LOC107352461 | eukaryota>metazoa>cnidaria | Acropora | digitifera | PREDICTED: serine/threonine-protein phosphatase 2A regulatory subunit B'' subunit beta-like, partial [Acropora digitifera] |
| XP_003521774.1 | 537 | LOC100807531 | eukaryota>viridiplantae | Glycine | max | PREDICTED: probable serine/threonine protein phosphatase 2A regulatory subunit B''delta [Glycine max] |
| XP_003554724.1 | 537 | LOC100815724 | eukaryota>viridiplantae | Glycine | max | PREDICTED: probable serine/threonine protein phosphatase 2A regulatory subunit B''delta [Glycine max] |
| XP_003556454.1 | 537 | LOC100799748 | eukaryota>viridiplantae | Glycine | max | PREDICTED: probable serine/threonine protein phosphatase 2A regulatory subunit B''delta isoform X1 [Glycine max] |
| XP_006589210.1 | 537 | LOC100792998 | eukaryota>viridiplantae | Glycine | max | PREDICTED: probable serine/threonine protein phosphatase 2A regulatory subunit B''delta [Glycine max] |
| XP_006589211.1 | 537 | LOC100792998 | eukaryota>viridiplantae | Glycine | max | PREDICTED: probable serine/threonine protein phosphatase 2A regulatory subunit B''delta [Glycine max] |
| XP_006589212.1 | 537 | LOC100792998 | eukaryota>viridiplantae | Glycine | max | PREDICTED: probable serine/threonine protein phosphatase 2A regulatory subunit B''delta [Glycine max] |
| XP_006606445.1 | 537 | LOC100799748 | eukaryota>viridiplantae | Glycine | max | PREDICTED: probable serine/threonine protein phosphatase 2A regulatory subunit B''delta isoform X1 [Glycine max] |
| XP_006606446.1 | 537 | LOC100799748 | eukaryota>viridiplantae | Glycine | max | PREDICTED: probable serine/threonine protein phosphatase 2A regulatory subunit B''delta isoform X1 [Glycine max] |
| XP_014629523.1 | 537 | LOC100807531 | eukaryota>viridiplantae | Glycine | max | PREDICTED: probable serine/threonine protein phosphatase 2A regulatory subunit B''delta [Glycine max] |
| EPY19001.1 | 536 | STCU_09675 | eukaryota>euglenozoa>kinetoplastida | Strigomonas | culicis | hypothetical protein STCU_09675 [Strigomonas culicis] |
| EPY33753.1 | 536 | STCU_02013 | eukaryota>euglenozoa>kinetoplastida | Strigomonas | culicis | hypothetical protein STCU_02013 [Strigomonas culicis] |
| NP_198242.1 | 536 | F7P1.80,F7P1_80 | eukaryota>viridiplantae | Arabidopsis | thaliana | Calcium-binding EF-hand family protein [Arabidopsis thaliana] |
| NP_568509.1 | 536 | F7P1.30,F7P1_30 | eukaryota>viridiplantae | Arabidopsis | thaliana | Calcium-binding EF-hand family protein [Arabidopsis thaliana] |
| XP_001568114.1 | 536 | LBRM_33_3500 | eukaryota>euglenozoa>kinetoplastida | Leishmania | braziliensis MHOM/BR/75/M2904 | conserved hypothetical protein [Leishmania braziliensis MHOM/BR/75/M2904] |
| XP_008917473.1 | 536 | PPTG_20421 | eukaryota>stramenopiles | Phytophthora | parasitica INRA-310 | hypothetical protein PPTG_20421, partial [Phytophthora parasitica INRA-310] |
| XP_010702423.1 | 536 | LPMP_333350 | eukaryota>euglenozoa>kinetoplastida | Leishmania | panamensis | hypothetical protein LPMP_333350 [Leishmania panamensis] |
| EPY25465.1 | 535 | AGDE_11742 | eukaryota>euglenozoa>kinetoplastida | Angomonas | deanei | hypothetical protein AGDE_11742 [Angomonas deanei] |
| NP_175847.1 | 535 | F20D21.27,F20D21_27 | eukaryota>viridiplantae | Arabidopsis | thaliana | Calcium-binding EF-hand family protein [Arabidopsis thaliana] |
| XP_001468382.1 | 534 | LINJ_33_3370 | eukaryota>euglenozoa>kinetoplastida | Leishmania | infantum JPCM5 | conserved hypothetical protein [Leishmania infantum JPCM5] |
| XP_001686084.1 | 534 | LMJF_33_3220 | eukaryota>euglenozoa>kinetoplastida | Leishmania | major strain Friedlin | conserved hypothetical protein [Leishmania major strain Friedlin] |
| XP_003864160.1 | 534 | LDBPK_333370 | eukaryota>euglenozoa>kinetoplastida | Leishmania | donovani | hypothetical protein, conserved [Leishmania donovani] |
| XP_003878578.1 | 534 | LMXM_32_3220 | eukaryota>euglenozoa>kinetoplastida | Leishmania | mexicana MHOM/GT/2001/U1103 | conserved hypothetical protein [Leishmania mexicana MHOM/GT/2001/U1103] |
| ERG81877.1 | 533 | ASU_09497 | eukaryota>metazoa>nematoda | Ascaris | suum | serine threonine-protein phosphatase 2a regulatory subunit b subunit alpha [Ascaris suum] |
| XP_017324900.1 | 533 | LOC108266308 | eukaryota>metazoa>chordata>vertebrata>actinopterygii | Ictalurus | punctatus | PREDICTED: serine/threonine-protein phosphatase 2A regulatory subunit B'' subunit beta-like isoform X2 [Ictalurus punctatus] |
| ABO97244.1 | 531 | OSTLU_32696 | eukaryota>viridiplantae>chlorophyta | Ostreococcus | lucimarinus CCE9901 | predicted protein [Ostreococcus lucimarinus CCE9901] |
| NP_742156.2 | 530 | 3222402P14Rik,A730042E07 | eukaryota>metazoa>chordata>vertebrata | Mus | musculus | serine/threonine-protein phosphatase 2A regulatory subunit B'' subunit alpha isoform 2 [Mus musculus] |
| NP_171892.2 | 529 | F21M11.11,F21M11_11 | eukaryota>viridiplantae | Arabidopsis | thaliana | Calcium-binding EF hand family protein [Arabidopsis thaliana] |
| NP_871626.1 | 529 | PPP2R3,PR130,PR72 | eukaryota>metazoa>chordata>vertebrata | Homo | sapiens | serine/threonine-protein phosphatase 2A regulatory subunit B'' subunit alpha isoform 2 [Homo sapiens] |
| OMJ71500.1 | 529 | SteCoe_30282 | eukaryota>alveolata>ciliophora | Stentor | coeruleus | hypothetical protein SteCoe_30282 [Stentor coeruleus] |
| OMJ93943.1 | 529 | SteCoe_2999 | eukaryota>alveolata>ciliophora | Stentor | coeruleus | hypothetical protein SteCoe_2999 [Stentor coeruleus] |
| XP_003265305.1 | 529 | PPP2R3A | eukaryota>metazoa>chordata>vertebrata | Nomascus | leucogenys | PREDICTED: serine/threonine-protein phosphatase 2A regulatory subunit B'' subunit alpha isoform X2 [Nomascus leucogenys] |
| XP_005545890.1 | 529 | PPP2R3A | eukaryota>metazoa>chordata>vertebrata | Macaca | fascicularis | PREDICTED: serine/threonine-protein phosphatase 2A regulatory subunit B'' subunit alpha isoform X2 [Macaca fascicularis] |
| XP_009312624.1 | 528 | DQ04_05721050 | eukaryota>euglenozoa>kinetoplastida | Trypanosoma | grayi | hypothetical protein DQ04_05721050 [Trypanosoma grayi] |
| XP_806034.1 | 528 | Tc00.1047053506815.50 | eukaryota>euglenozoa>kinetoplastida | Trypanosoma | cruzi strain CL Brener | hypothetical protein [Trypanosoma cruzi strain CL Brener] |
| XP_811419.1 | 528 | Tc00.1047053507105.30 | eukaryota>euglenozoa>kinetoplastida | Trypanosoma | cruzi strain CL Brener | hypothetical protein [Trypanosoma cruzi strain CL Brener] |
| OMJ79973.1 | 526 | SteCoe_19892 | eukaryota>alveolata>ciliophora | Stentor | coeruleus | hypothetical protein SteCoe_19892 [Stentor coeruleus] |
| OMJ92953.1 | 526 | SteCoe_4171 | eukaryota>alveolata>ciliophora | Stentor | coeruleus | hypothetical protein SteCoe_4171 [Stentor coeruleus] |
| XP_011613645.1 | 526 | LOC101070572 | eukaryota>metazoa>chordata>vertebrata>actinopterygii | Takifugu | rubripes | PREDICTED: serine/threonine-protein phosphatase 2A regulatory subunit B'' subunit alpha-like [Takifugu rubripes] |

[illegible]

| | | | | | | |
|---|---|---|---|---|---|---|
| NP_001124163.1 | 511 | CALN,CALNA,CALNA1,CCN1,CNA1,IECEE,IECEE1,PPP2B | eukaryota>metazoa>chordata>vertebrata | Homo | sapiens | serine/threonine-protein phosphatase 2B catalytic subunit alpha isoform isoform 2 [Homo sapiens] |
| NP_001087372.1 | 511 | calcineurin,caln,calna,calna1,ccn1,cna1,ppp2b,ppp3ca | eukaryota>metazoa>chordata>vertebrata | Xenopus | laevis | protein phosphatase 3, catalytic subunit, alpha isozyme L homeolog [Xenopus laevis] |
| NP_001017856.1 | 511 | wu:fq19c08,zgc:110141 | eukaryota>metazoa>chordata>vertebrata>actinopterygii | Danio | rerio | serine/threonine-protein phosphatase 2B catalytic subunit alpha [Danio rerio] |
| XP_005155370.1 | 511 | im:7163784,zgc:158685 | eukaryota>metazoa>chordata>vertebrata>actinopterygii | Danio | rerio | protein phosphatase 3, catalytic subunit, gamma isoform X1 [Danio rerio] |
| XP_017327655.1 | 511 | ppp3ca | eukaryota>metazoa>chordata>vertebrata>actinopterygii | Ictalurus | punctatus | PREDICTED: serine/threonine-protein phosphatase 2B catalytic subunit alpha isoform isoform X1 [Ictalurus punctatus] |
| EMH74210.1 | 511 | EHI8A_054460 | eukaryota>amoebozoa>entamoebidae | Entamoeba | histolytica HM-1:IMSS-B | calcineurin catalytic subunit A, putative [Entamoeba histolytica HM-1:IMSS-B] |
| EMD46097.1 | 511 | EHI5A_107660 | eukaryota>amoebozoa>entamoebidae | Entamoeba | histolytica KU27 | serine/threonine-protein phosphatase catalytic subunit gamma, putative [Entamoeba histolytica KU27] |
| EMH75764.1 | 511 | EHI8A_075050 | eukaryota>amoebozoa>entamoebidae | Entamoeba | histolytica HM-1:IMSS-B | calcineurin catalytic subunit A, putative [Entamoeba histolytica HM-1:IMSS-B] |
| EMS16518.1 | 511 | KM1_134500 | eukaryota>amoebozoa>entamoebidae | Entamoeba | histolytica HM-3:IMSS | serine/threonine protein phosphatase 2B catalytic subunit gamma isoform, putative [Entamoeba histolytica HM-3:IMSS] |
| ENY62767.1 | 511 | EHI7A_072900 | eukaryota>amoebozoa>entamoebidae | Entamoeba | histolytica HM-1:IMSS-A | serine/threonine protein phosphatase 2B catalytic subunit gamma isoform, putative [Entamoeba histolytica HM-1:IMSS-A] |
| GAT92021.1 | 511 | CL6EHI_050920 | eukaryota>amoebozoa>entamoebidae | Entamoeba | histolytica | calcineurin catalytic subunit A putative [Entamoeba histolytica] |
| XP_654180.2 | 511 | EHI_050920 | eukaryota>amoebozoa>entamoebidae | Entamoeba | histolytica HM-1:IMSS | calcineurin catalytic subunit A, putative [Entamoeba histolytica HM-1:IMSS] |
| ORY48385.1 | 510 | LY90DRAFT_523579 | eukaryota>fungi>chytridiomycota | Neocallimastix | californiae | Serine/threonine-protein phosphatase 2B catalytic subunit [Neocallimastix californiae] |
| XP_018080855.1 | 510 | ppp3cc.L | eukaryota>metazoa>chordata>vertebrata | Xenopus | laevis | PREDICTED: serine/threonine-protein phosphatase 2B catalytic subunit beta isoform isoform X5 [Xenopus laevis] |
| XP_015141109.1 | 510 | PPP3CA | eukaryota>metazoa>chordata>vertebrata | Gallus | gallus | PREDICTED: serine/threonine-protein phosphatase 2B catalytic subunit alpha isoform isoform X6 [Gallus gallus] |
| XP_005562855.1 | 510 | PPP3CC | eukaryota>metazoa>chordata>vertebrata | Macaca | fascicularis | PREDICTED: serine/threonine-protein phosphatase 2B catalytic subunit gamma isoform isoform X3 [Macaca fascicularis] |
| NP_001025511.1 | 510 | PPP3CB | eukaryota>metazoa>chordata>vertebrata | Gallus | gallus | serine/threonine-protein phosphatase 2B catalytic subunit beta isoform [Gallus gallus] |
| XP_021336694.1 | 510 | wu:fq19c08,zgc:110141 | eukaryota>metazoa>chordata>vertebrata>actinopterygii | Danio | rerio | serine/threonine-protein phosphatase 2B catalytic subunit alpha isoform X2 [Danio rerio] |
| XP_015833482.1 | 510 | LOC657135 | eukaryota>metazoa>hexapoda | Tribolium | castaneum | PREDICTED: serine/threonine-protein phosphatase 2B catalytic subunit 2 isoform X1 [Tribolium castaneum] |
| XP_015833483.1 | 510 | LOC657135 | eukaryota>metazoa>hexapoda | Tribolium | castaneum | PREDICTED: serine/threonine-protein phosphatase 2B catalytic subunit 2 isoform X1 [Tribolium castaneum] |
| GAT93915.1 | 510 | CL6EHI_118600 | eukaryota>amoebozoa>entamoebidae | Entamoeba | histolytica | calcineurin catalytic subunit A putative [Entamoeba histolytica] |
| XP_001913593.1 | 510 | EHI_118600 | eukaryota>amoebozoa>entamoebidae | Entamoeba | histolytica HM-1:IMSS | calcineurin catalytic subunit A, putative [Entamoeba histolytica HM-1:IMSS] |
| XP_008856053.1 | 510 | ENU1_050070 | eukaryota>amoebozoa>entamoebidae | Entamoeba | nuttalli P19 | calcineurin catalytic subunit A, putative [Entamoeba nuttalli P19] |
| XP_001448076.1 | 509 | GSPATT00015557001 | eukaryota>alveolata>ciliophora | Paramecium | tetraurelia strain d4-2 | hypothetical protein (macronuclear) [Paramecium tetraurelia strain d4-2] |
| XP_022245394.1 | 509 | LOC106465252 | eukaryota>metazoa | Limulus | polyphemus | serine/threonine-protein phosphatase 2B catalytic subunit 2-like isoform X4 [Limulus polyphemus] |
| XP_022240920.1 | 509 | LOC106458858 | eukaryota>metazoa | Limulus | polyphemus | serine/threonine-protein phosphatase 2B catalytic subunit 2-like isoform X1 [Limulus polyphemus] |
| XP_016863854.1 | 509 | CALN,CALNA,CALNA1,CCN1,CNA1,IECEE,IECEE1,PPP2B | eukaryota>metazoa>chordata>vertebrata | Homo | sapiens | serine/threonine-protein phosphatase 2B catalytic subunit alpha isoform isoform X1 [Homo sapiens] |
| XP_008199522.1 | 509 | LOC657135 | eukaryota>metazoa>hexapoda | Tribolium | castaneum | PREDICTED: serine/threonine-protein phosphatase 2B catalytic subunit 2 isoform X2 [Tribolium castaneum] |
| XP_015833484.1 | 509 | LOC657135 | eukaryota>metazoa>hexapoda | Tribolium | castaneum | PREDICTED: serine/threonine-protein phosphatase 2B catalytic subunit 2 isoform X3 [Tribolium castaneum] |
| XP_011445770.2 | 509 | LOC105341116 | eukaryota>metazoa>mollusca | Crassostrea | gigas | PREDICTED: serine/threonine-protein phosphatase 2B catalytic subunit 3 isoform X1 [Crassostrea gigas] |
| XP_001443973.1 | 508 | GSPATT00012128001 | eukaryota>alveolata>ciliophora | Paramecium | tetraurelia strain d4-2 | hypothetical protein (macronuclear) [Paramecium tetraurelia strain d4-2] |
| XP_018089403.1 | 508 | ppp3ca | eukaryota>metazoa>chordata>vertebrata | Xenopus | laevis | PREDICTED: protein phosphatase 3, catalytic subunit, alpha isozyme isoform X1 [Xenopus laevis] |
| XP_005171933.1 | 508 | ppp3cc,sb:cb767,si:dkey-21k4.5 | eukaryota>metazoa>chordata>vertebrata>actinopterygii | Danio | rerio | protein phosphatase 3, catalytic subunit, gamma isozyme-like isoform X1 [Danio rerio] |
| XP_015833485.1 | 508 | LOC657135 | eukaryota>metazoa>hexapoda | Tribolium | castaneum | PREDICTED: serine/threonine-protein phosphatase 2B catalytic subunit 2 isoform X4 [Tribolium castaneum] |
| XP_001450977.1 | 507 | GSPATT00017910001 | eukaryota>alveolata>ciliophora | Paramecium | tetraurelia strain d4-2 | hypothetical protein (macronuclear) [Paramecium tetraurelia strain d4-2] |
| XP_001431776.1 | 507 | GSPATT00033854001 | eukaryota>alveolata>ciliophora | Paramecium | tetraurelia strain d4-2 | hypothetical protein (macronuclear) [Paramecium tetraurelia strain d4-2] |
| XP_001430129.1 | 507 | GSPATT00032583001 | eukaryota>alveolata>ciliophora | Paramecium | tetraurelia strain d4-2 | hypothetical protein (macronuclear) [Paramecium tetraurelia strain d4-2] |
| EJY65152.1 | 507 | OXYTRI_14698 | eukaryota>alveolata>ciliophora | Oxytricha | trifallax | Serine/threonine-protein phosphatase (macronuclear) [Oxytricha trifallax] |
| NP_001085039.1 | 507 | calna2,calnb,cna2,pp2bbeta,ppp3cb | eukaryota>metazoa>chordata>vertebrata | Xenopus | laevis | protein phosphatase 3, catalytic subunit, beta isozyme L homeolog [Xenopus laevis] |
| XP_001456473.1 | 506 | GSPATT00022245001 | eukaryota>alveolata>ciliophora | Paramecium | tetraurelia strain d4-2 | hypothetical protein (macronuclear) [Paramecium tetraurelia strain d4-2] |
| XP_001012424.2 | 506 | TTHERM_00804750 | eukaryota>alveolata>ciliophora | Tetrahymena | thermophila SB210 | serine/threonine-protein phosphatase PP1 [Tetrahymena thermophila SB210] |
| XP_022240921.1 | 506 | LOC106458858 | eukaryota>metazoa | Limulus | polyphemus | serine/threonine-protein phosphatase 2B catalytic subunit 2-like isoform X2 [Limulus polyphemus] |
| NP_001074063.2 | 506 | im:7163784,zgc:158685 | eukaryota>metazoa>chordata>vertebrata>actinopterygii | Danio | rerio | protein phosphatase 3, catalytic subunit, gamma isoform [Danio rerio] |
| XP_009305999.1 | 506 | wu:fq19c08,zgc:110141 | eukaryota>metazoa>chordata>vertebrata>actinopterygii | Danio | rerio | serine/threonine-protein phosphatase 2B catalytic subunit alpha isoform X3 [Danio rerio] |
| XP_011678779.1 | 506 | LOC588695 | eukaryota>metazoa>echinodermata | Strongylocentrotus | purpuratus | PREDICTED: serine/threonine-protein phosphatase 2B catalytic subunit alpha isoform isoform X1 [Strongylocentrotus purpuratus] |
| CDW79014.1 | 505 | Contig16048.g787 | eukaryota>alveolata>ciliophora | Stylonychia | lemnae | UNKNOWN [Stylonychia lemnae] |
| XP_005161416.1 | 505 | fj46e08,si:dkeyp-79c2.2,wu:fj46e08 | eukaryota>metazoa>chordata>vertebrata>actinopterygii | Danio | rerio | protein phosphatase 3, catalytic subunit, alpha isozyme isoform X1 [Danio rerio] |
| XP_017307366.1 | 505 | LOC108255714 | eukaryota>metazoa>chordata>vertebrata>actinopterygii | Ictalurus | punctatus | PREDICTED: serine/threonine-protein phosphatase 2B catalytic subunit gamma isoform-like isoform X1 [Ictalurus punctatus] |
| XP_017340418.1 | 505 | LOC108274683 | eukaryota>metazoa>chordata>vertebrata>actinopterygii | Ictalurus | punctatus | PREDICTED: serine/threonine-protein phosphatase 2B catalytic subunit gamma isoform-like isoform X3 [Ictalurus punctatus] |
| XP_011678780.1 | 505 | LOC588695 | eukaryota>metazoa>echinodermata | Strongylocentrotus | purpuratus | PREDICTED: serine/threonine-protein phosphatase 2B catalytic subunit alpha isoform isoform X2 [Strongylocentrotus purpuratus] |
| XP_011445772.2 | 505 | LOC105341116 | eukaryota>metazoa>mollusca | Crassostrea | gigas | PREDICTED: serine/threonine-protein phosphatase 2B catalytic subunit 3 isoform X2 [Crassostrea gigas] |
| XP_627753.1 | 504 | cgd6_4200 | eukaryota>alveolata>apicomplexa | Cryptosporidium | parvum Iowa II | protein phosphatase 2B catalytic subunit, calcineurin like phosphatase superfamily [Cryptosporidium parvum Iowa II] |
| XP_001452806.1 | 504 | GSPATT00002809001 | eukaryota>alveolata>ciliophora | Paramecium | tetraurelia strain d4-2 | hypothetical protein (macronuclear) [Paramecium tetraurelia strain d4-2] |
| XP_001442415.1 | 504 | GSPATT00001220001 | eukaryota>alveolata>ciliophora | Paramecium | tetraurelia strain d4-2 | hypothetical protein (macronuclear) [Paramecium tetraurelia strain d4-2] |
| XP_013379079.1 | 504 | LOC106150691 | eukaryota>metazoa | Lingula | anatina | serine/threonine-protein phosphatase 2B catalytic subunit 2-like isoform X1 [Lingula anatina] |
| XP_003876556.1 | 503 | LMXM_26_2530 | eukaryota>euglenozoa>kinetoplastida | Leishmania | mexicana MHOM/GT/2001/U1103 | putative serine/threonine protein phosphatase [Leishmania mexicana MHOM/GT/2001/U1103] |
| XP_006518777.1 | 503 | Calnc,PP2BA,gamma | eukaryota>metazoa>chordata>vertebrata | Mus | musculus | PREDICTED: serine/threonine-protein phosphatase 2B catalytic subunit gamma isoform isoform X1 [Mus musculus] |
| XP_017347413.1 | 503 | LOC108278499 | eukaryota>metazoa>chordata>vertebrata>actinopterygii | Ictalurus | punctatus | PREDICTED: serine/threonine-protein phosphatase 2B catalytic subunit gamma isoform-like isoform X2 [Ictalurus punctatus] |
| XP_002142661.1 | 502 | CMU_020560 | eukaryota>alveolata>apicomplexa | Cryptosporidium | muris RN66 | serine/threonine protein phosphatase [Cryptosporidium muris RN66] |
| CDW87430.1 | 502 | Contig15030.g16014 | eukaryota>alveolata>ciliophora | Stylonychia | lemnae | serine threonine-protein phosphatase [Stylonychia lemnae] |
| XP_001456389.1 | 502 | GSPATT00022161001 | eukaryota>alveolata>ciliophora | Paramecium | tetraurelia strain d4-2 | hypothetical protein (macronuclear) [Paramecium tetraurelia strain d4-2] |
| XP_018080856.1 | 502 | ppp3cc.L | eukaryota>metazoa>chordata>vertebrata | Xenopus | laevis | PREDICTED: serine/threonine-protein phosphatase 2B catalytic subunit beta isoform isoform X6 [Xenopus laevis] |
| NP_001230904.1 | 502 | CALNA3,CNA3,PP2Bgamma | eukaryota>metazoa>chordata>vertebrata | Homo | sapiens | serine/threonine-protein phosphatase 2B catalytic subunit gamma isoform isoform 3 [Homo sapiens] |
| XP_005155371.1 | 502 | im:7163784,zgc:158685 | eukaryota>metazoa>chordata>vertebrata>actinopterygii | Danio | rerio | protein phosphatase 3, catalytic subunit, gamma isoform X2 [Danio rerio] |
| XP_011678781.1 | 502 | LOC588695 | eukaryota>metazoa>echinodermata | Strongylocentrotus | purpuratus | PREDICTED: serine/threonine-protein phosphatase 2B catalytic subunit alpha isoform isoform X3 [Strongylocentrotus purpuratus] |
| CEL93926.1 | 501 | Vbra_20271 | eukaryota>alveolata | Vitrella | brassicaformis CCMP3155 | unnamed protein product [Vitrella brassicaformis CCMP3155] |
| XP_008885873.1 | 501 | HHA_311310 | eukaryota>alveolata>apicomplexa | Hammondia | hammondi | protein phosphatase 2B catalytic subunit, calcineurin family phosphatase superfamily protein [Hammondia hammondi] |
| XP_018635527.1 | 501 | TGME49_311310 | eukaryota>alveolata>apicomplexa | Toxoplasma | gondii ME49 | protein phosphatase 2B catalytic subunit, calcineurin family phosphatase superfamily protein [Toxoplasma gondii ME49] |
| CDW84965.1 | 501 | Contig3485.g3724 | eukaryota>alveolata>ciliophora | Stylonychia | lemnae | serine threonine-protein phosphatase [Stylonychia lemnae] |
| XP_018080857.1 | 501 | ppp3cc.L | eukaryota>metazoa>chordata>vertebrata | Xenopus | laevis | PREDICTED: serine/threonine-protein phosphatase 2B catalytic subunit beta isoform isoform X7 [Xenopus laevis] |
| XP_005562856.1 | 501 | PPP3CC | eukaryota>metazoa>chordata>vertebrata | Macaca | fascicularis | PREDICTED: serine/threonine-protein phosphatase 2B catalytic subunit gamma isoform isoform X4 [Macaca fascicularis] |
| XP_015833486.1 | 501 | LOC657135 | eukaryota>metazoa>hexapoda | Tribolium | castaneum | PREDICTED: serine/threonine-protein phosphatase 2B catalytic subunit 2 isoform X5 [Tribolium castaneum] |
| XP_012650119.1 | 500 | BmR1_04g07610 | eukaryota>alveolata>apicomplexa | Babesia | microti strain RI | protein phosphatase 3, catalytic subunit [Babesia microti strain RI] |
| XP_013379080.1 | 500 | LOC106150691 | eukaryota>metazoa | Lingula | anatina | serine/threonine-protein phosphatase 2B catalytic subunit 2-like isoform X2 [Lingula anatina] |
| XP_011613631.1 | 500 | LOC101072650 | eukaryota>metazoa>chordata>vertebrata>actinopterygii | Takifugu | rubripes | PREDICTED: serine/threonine-protein phosphatase 2B catalytic subunit gamma isoform-like [Takifugu rubripes] |
| XP_001684260.1 | 499 | LMJF_26_2530 | eukaryota>euglenozoa>kinetoplastida | Leishmania | major strain Friedlin | putative serine/threonine protein phosphatase [Leishmania major strain Friedlin] |
| NP_001160100.1 | 499 | ppp3cc,sb:cb767,si:dkey-21k4.5 | eukaryota>metazoa>chordata>vertebrata>actinopterygii | Danio | rerio | protein phosphatase 3, catalytic subunit, gamma isozyme-like [Danio rerio] |
| XP_015833487.1 | 499 | LOC657135 | eukaryota>metazoa>hexapoda | Tribolium | castaneum | PREDICTED: serine/threonine-protein phosphatase 2B catalytic subunit 2 isoform X6 [Tribolium castaneum] |
| XP_001027523.2 | 498 | TTHERM_01055570 | eukaryota>alveolata>ciliophora | Tetrahymena | thermophila SB210 | serine/threonine-protein phosphatase PP1 [Tetrahymena thermophila SB210] |
| XP_003257535.1 | 498 | PPP3CA | eukaryota>metazoa>chordata>vertebrata | Nomascus | leucogenys | PREDICTED: serine/threonine-protein phosphatase 2B catalytic subunit alpha isoform isoform X3 [Nomascus leucogenys] |
| XP_008199524.1 | 498 | LOC657135 | eukaryota>metazoa>hexapoda | Tribolium | castaneum | PREDICTED: serine/threonine-protein phosphatase 2B catalytic subunit 2 isoform X7 [Tribolium castaneum] |
| NP_001158479.1 | 497 | cna1 | eukaryota>metazoa>hemichordata | Saccoglossus | kowalevskii | calcineurin A protein catalytic subunit [Saccoglossus kowalevskii] |
| XP_008199525.1 | 497 | LOC657135 | eukaryota>metazoa>hexapoda | Tribolium | castaneum | PREDICTED: serine/threonine-protein phosphatase 2B catalytic subunit 2 isoform X8 [Tribolium castaneum] |
| EMS14417.1 | 497 | KM1_300130 | eukaryota>amoebozoa>entamoebidae | Entamoeba | histolytica HM-3:IMSS | serine/threonine protein phosphatase 2B catalytic subunit, putative, partial [Entamoeba histolytica HM-3:IMSS] |
| XP_017307367.1 | 496 | LOC108255714 | eukaryota>metazoa>chordata>vertebrata>actinopterygii | Ictalurus | punctatus | PREDICTED: serine/threonine-protein phosphatase 2B catalytic subunit gamma isoform-like isoform X2 [Ictalurus punctatus] |
| XP_001868819.1 | 496 | CpipJ_CPIJ018692 | eukaryota>metazoa>hexapoda | Culex | quinquefasciatus | serine/threonine-protein phosphatase 2B catalytic subunit 2 [Culex quinquefasciatus] |
| XP_014677387.1 | 495 | LOC106817227 | eukaryota>metazoa | Priapulus | caudatus | PREDICTED: serine/threonine-protein phosphatase 2B catalytic subunit 3-like isoform X2 [Priapulus caudatus] |
| XP_014677380.1 | 495 | LOC106817227 | eukaryota>metazoa | Priapulus | caudatus | PREDICTED: serine/threonine-protein phosphatase 2B catalytic subunit 3-like isoform X1 [Priapulus caudatus] |
| XP_011608701.1 | 495 | LOC101070452 | eukaryota>metazoa>chordata>vertebrata>actinopterygii | Takifugu | rubripes | PREDICTED: LOW QUALITY PROTEIN: serine/threonine-protein phosphatase 2B catalytic subunit alpha isoform-like [Takifugu rubripes] |
| NP_001037025.1 | 495 | Cna | eukaryota>metazoa>hexapoda | Bombyx | mori | calcineurin A [Bombyx mori] |
| XP_013379081.1 | 493 | LOC106150691 | eukaryota>metazoa | Lingula | anatina | serine/threonine-protein phosphatase 2B catalytic subunit 2-like isoform X3 [Lingula anatina] |
| EMD46682.1 | 493 | EHI5A_230430 | eukaryota>amoebozoa>entamoebidae | Entamoeba | histolytica KU27 | serine/threonine protein phosphatase catalytic subunit alpha, putative, partial [Entamoeba histolytica KU27] |
| EJY85240.1 | 492 | OXYTRI_16901 | eukaryota>alveolata>ciliophora | Oxytricha | trifallax | Serine/threonine-protein phosphatase (macronuclear) [Oxytricha trifallax] |
| EJY83446.1 | 492 | OXYTRI_18825 | eukaryota>alveolata>ciliophora | Oxytricha | trifallax | Serine/threonine-protein phosphatase (macronuclear) [Oxytricha trifallax] |
| XP_001739119.1 | 492 | EDI_229070 | eukaryota>amoebozoa>entamoebidae | Entamoeba | dispar SAW760 | serine/threonine protein phosphatase 2B catalytic subunit [Entamoeba dispar SAW760] |
| CBY22738.1 | 491 | GSOID_T00013510001 | eukaryota>metazoa>chordata | Oikopleura | dioica | unnamed protein product [Oikopleura dioica] |
| XP_011678782.1 | 491 | LOC588695 | eukaryota>metazoa>echinodermata | Strongylocentrotus | purpuratus | PREDICTED: serine/threonine-protein phosphatase 2B catalytic subunit alpha isoform isoform X4 [Strongylocentrotus purpuratus] |
| XP_011130022.1 | 490 | GNI_063120 | eukaryota>alveolata>apicomplexa | Gregarina | niphandrodes | serine/threonine protein phosphatase [Gregarina niphandrodes] |
| CDW78640.1 | 490 | Contig17343.g18460 | eukaryota>alveolata>ciliophora | Stylonychia | lemnae | serine threonine-protein phosphatase [Stylonychia lemnae] |
| CCW70762.1 | 490 | GSHART1_T00008471001 | eukaryota>euglenozoa>kinetoplastida | Phytomonas | sp. isolate Hart1 | unnamed protein product [Phytomonas sp. isolate Hart1] |
| XP_003727335.1 | 490 | LOC588695 | eukaryota>metazoa>echinodermata | Strongylocentrotus | purpuratus | PREDICTED: serine/threonine-protein phosphatase 2B catalytic subunit alpha isoform isoform X5 [Strongylocentrotus purpuratus] |
| XP_008856724.1 | 490 | ENU1_074160 | eukaryota>amoebozoa>entamoebidae | Entamoeba | nuttalli P19 | Ser/Thr protein phosphatase, putative [Entamoeba nuttalli P19] |
| XP_013379082.1 | 489 | LOC106150691 | eukaryota>metazoa | Lingula | anatina | serine/threonine-protein phosphatase 2B catalytic subunit alpha isoform-like isoform X4 [Lingula anatina] |
| XP_013379083.1 | 488 | LOC106150691 | eukaryota>metazoa | Lingula | anatina | serine/threonine-protein phosphatase 2B catalytic subunit alpha isoform-like isoform X5 [Lingula anatina] |
| CCW65030.1 | 487 | GSEM1_T00005189001 | eukaryota>euglenozoa>kinetoplastida | Phytomonas | sp. isolate EM1 | unnamed protein product [Phytomonas sp. isolate EM1] |
| XP_018110019.1 | 487 | LOC108712283 | eukaryota>metazoa>chordata>vertebrata | Xenopus | laevis | PREDICTED: serine/threonine-protein phosphatase 2B catalytic subunit gamma isoform-like [Xenopus laevis] |
| XP_011678783.1 | 487 | LOC588695 | eukaryota>metazoa>echinodermata | Strongylocentrotus | purpuratus | PREDICTED: serine/threonine-protein phosphatase 2B catalytic subunit alpha isoform isoform X6 [Strongylocentrotus purpuratus] |
| XP_006819681.1 | 487 | cna1 | eukaryota>metazoa>hemichordata | Saccoglossus | kowalevskii | PREDICTED: calcineurin A protein catalytic subunit isoform X2 [Saccoglossus kowalevskii] |
| XP_006819680.1 | 487 | cna1 | eukaryota>metazoa>hemichordata | Saccoglossus | kowalevskii | PREDICTED: calcineurin A protein catalytic subunit isoform X1 [Saccoglossus kowalevskii] |
| XP_004184153.1 | 487 | EIN_283530 | eukaryota>amoebozoa>entamoebidae | Entamoeba | invadens IP1 | serine/threonine protein phosphatase 2B catalytic subunit, putative, partial [Entamoeba invadens IP1] |
| XP_002674263.1 | 486 | NAEGRDRAFT_60801 | eukaryota>heterolobosea | Naegleria | gruberi strain NEG-M | predicted protein [Naegleria gruberi strain NEG-M] |
| CBY23867.1 | 486 | GSOID_T00001193001 | eukaryota>metazoa>chordata | Oikopleura | dioica | unnamed protein product [Oikopleura dioica] |
| XP_009062091.1 | 486 | LOTGIDRAFT_183550 | eukaryota>metazoa>mollusca | Lottia | gigantea | hypothetical protein LOTGIDRAFT_183550 [Lottia gigantea] |
| XP_006518778.1 | 485 | Calnc,PP2BA,gamma | eukaryota>metazoa>chordata>vertebrata | Mus | musculus | PREDICTED: serine/threonine-protein phosphatase 2B catalytic subunit gamma isoform isoform X2 [Mus musculus] |
| CDW84032.1 | 483 | Contig471.g515 | eukaryota>alveolata>ciliophora | Stylonychia | lemnae | serine threonine-protein phosphatase [Stylonychia lemnae] |
| XP_006819682.1 | 483 | cna1 | eukaryota>metazoa>hemichordata | Saccoglossus | kowalevskii | PREDICTED: calcineurin A protein catalytic subunit isoform X3 [Saccoglossus kowalevskii] |
| EJY83969.1 | 482 | OXYTRI_18295 | eukaryota>alveolata>ciliophora | Oxytricha | trifallax | Serine/threonine-protein phosphatase (macronuclear) [Oxytricha trifallax] |
| EJY68315.1 | 482 | OXYTRI_11170 | eukaryota>alveolata>ciliophora | Oxytricha | trifallax | Serine/threonine-protein phosphatase (macronuclear) [Oxytricha trifallax] |
| XP_009313051.1 | 482 | DQ04_06571060 | eukaryota>euglenozoa>kinetoplastida | Trypanosoma | grayi | protein phosphatase 3, catalytic subunit [Trypanosoma grayi] |
| XP_002157885.2 | 482 | LOC100205420 | eukaryota>metazoa>cnidaria | Hydra | vulgaris | PREDICTED: serine/threonine-protein phosphatase 2B catalytic subunit alpha isoform [Hydra vulgaris] |
| XP_003861840.1 | 480 | LDBPK_262560 | eukaryota>euglenozoa>kinetoplastida | Leishmania | donovani | serine/threonine protein phosphatase, putative [Leishmania donovani] |
| XP_001470601.1 | 480 | LINJ_26_2560 | eukaryota>euglenozoa>kinetoplastida | Leishmania | infantum JPCM5 | putative serine/threonine protein phosphatase [Leishmania infantum JPCM5] |
| XP_011609907.1 | 480 | LOC101073844 | eukaryota>metazoa>chordata>vertebrata>actinopterygii | Takifugu | rubripes | PREDICTED: serine/threonine-protein phosphatase 2B catalytic subunit alpha isoform-like [Takifugu rubripes] |
| KXJ14923.1 | 480 | PPP3CA | eukaryota>metazoa>cnidaria | Exaiptasia | pallida | Serine/threonine-protein phosphatase 2B catalytic subunit alpha isoform [Exaiptasia pallida] |
| XP_011678784.1 | 480 | LOC588695 | eukaryota>metazoa>echinodermata | Strongylocentrotus | purpuratus | PREDICTED: serine/threonine-protein phosphatase 2B catalytic subunit alpha isoform isoform X7 [Strongylocentrotus purpuratus] |
| XP_004039734.1 | 478 | IMG5_011970 | eukaryota>alveolata>ciliophora | Ichthyophthirius | multifiliis | ser thr protein phosphatase family protein, putative [Ichthyophthirius multifiliis] |
| EJY67052.1 | 478 | OXYTRI_12653 | eukaryota>alveolata>ciliophora | Oxytricha | trifallax | Serine/threonine-protein phosphatase (macronuclear) [Oxytricha trifallax] |
| XP_014777764.1 | 478 | LOC106874522 | eukaryota>metazoa>mollusca | Octopus | bimaculoides | PREDICTED: serine/threonine-protein phosphatase 2B catalytic subunit alpha isoform-like isoform X1 [Octopus bimaculoides] |
| XP_004035302.1 | 477 | IMG5_101200 | eukaryota>alveolata>ciliophora | Ichthyophthirius | multifiliis | ser thr protein phosphatase family protein, putative, partial [Ichthyophthirius multifiliis] |
| CBY34065.1 | 477 | GSOID_T00022046001 | eukaryota>metazoa>chordata | Oikopleura | dioica | unnamed protein product [Oikopleura dioica] |
| EJY87836.1 | 476 | OXYTRI_23597 | eukaryota>alveolata>ciliophora | Oxytricha | trifallax | Serine/threonine-protein phosphatase (macronuclear) [Oxytricha trifallax] |
| XP_003885126.1 | 475 | NCLIV_055230 | eukaryota>alveolata>apicomplexa | Neospora | caninum Liverpool | putative serine/threonine protein phosphatase [Neospora caninum Liverpool] |
| XP_013755964.1 | 475 | AMSG_07462 | eukaryota>apusozoa>apusomonadidae | Thecamonas | trahens ATCC 50062 | hypothetical protein AMSG_07462 [Thecamonas trahens ATCC 50062] |
| XP_004365567.2 | 475 | CAOG_00696 | eukaryota>ichthyosporea | Capsaspora | owczarzaki ATCC 30864 | calcineurin A alpha [Capsaspora owczarzaki ATCC 30864] |
| XP_002111637.1 | 473 | TRIADDRAFT_24726 | eukaryota>placozoa | Trichoplax | adhaerens | conserved hypothetical protein, partial [Trichoplax adhaerens] |
| XP_004997803.1 | 472 | PTSG_11785 | eukaryota>choanoflagellida | Salpingoeca | rosetta | serine/threonine-protein phosphatase [Salpingoeca rosetta] |
| XP_012213090.1 | 472 | SPRG_18262 | eukaryota>stramenopiles | Saprolegnia | parasitica CBS 223.65 | hypothetical protein SPRG_18262, partial [Saprolegnia parasitica CBS 223.65] |
| EJY68175.1 | 471 | OXYTRI_11310 | eukaryota>alveolata>ciliophora | Oxytricha | trifallax | Serine/threonine-protein phosphatase (macronuclear) [Oxytricha trifallax] |
| CDW80026.1 | 471 | Contig11242.g12005 | eukaryota>alveolata>ciliophora | Stylonychia | lemnae | serine threonine-protein phosphatase [Stylonychia lemnae] |
| XP_012355006.1 | 471 | PPP3CA | eukaryota>metazoa>chordata>vertebrata | Nomascus | leucogenys | PREDICTED: serine/threonine-protein phosphatase 2B catalytic subunit alpha isoform isoform X7 [Nomascus leucogenys] |
| XP_015306174.1 | 471 | PPP3CA | eukaryota>metazoa>chordata>vertebrata | Macaca | fascicularis | PREDICTED: serine/threonine-protein phosphatase 2B catalytic subunit alpha isoform isoform X3 [Macaca fascicularis] |
| XP_015306175.1 | 471 | PPP3CA | eukaryota>metazoa>chordata>vertebrata | Macaca | fascicularis | PREDICTED: serine/threonine-protein phosphatase 2B catalytic subunit alpha isoform isoform X3 [Macaca fascicularis] |
| XP_015306176.1 | 471 | PPP3CA | eukaryota>metazoa>chordata>vertebrata | Macaca | fascicularis | PREDICTED: serine/threonine-protein phosphatase 2B catalytic subunit alpha isoform isoform X3 [Macaca fascicularis] |
| XP_015306177.1 | 471 | PPP3CA | eukaryota>metazoa>chordata>vertebrata | Macaca | fascicularis | PREDICTED: serine/threonine-protein phosphatase 2B catalytic subunit alpha isoform isoform X3 [Macaca fascicularis] |
| XP_015306178.1 | 471 | PPP3CA | eukaryota>metazoa>chordata>vertebrata | Macaca | fascicularis | PREDICTED: serine/threonine-protein phosphatase 2B catalytic subunit alpha isoform isoform X3 [Macaca fascicularis] |
| XP_017174981.1 | 471 | 2900074D19Rik,Caln,Calna,CN,CnA | eukaryota>metazoa>chordata>vertebrata | Mus | musculus | PREDICTED: serine/threonine-protein phosphatase 2B catalytic subunit alpha isoform isoform X1 [Mus musculus] |
| XP_001453246.1 | 470 | GSPATT00019553001 | eukaryota>alveolata>ciliophora | Paramecium | tetraurelia strain d4-2 | hypothetical protein (macronuclear) [Paramecium tetraurelia strain d4-2] |
| XP_001441867.1 | 470 | GSPATT00010670001 | eukaryota>alveolata>ciliophora | Paramecium | tetraurelia strain d4-2 | hypothetical protein (macronuclear) [Paramecium tetraurelia strain d4-2] |
| XP_001749964.1 | 470 | MONBRDRAFT_29480 | eukaryota>choanoflagellida | Monosiga | brevicollis MX1 | hypothetical protein [Monosiga brevicollis MX1] |
| XP_808861.1 | 470 | Tc00.1047053508413.40 | eukaryota>euglenozoa>kinetoplastida | Trypanosoma | cruzi strain CL Brener | serine/threonine protein phosphatase [Trypanosoma cruzi strain CL Brener] |
| OMJ85969.1 | 469 | SteCoe_12602 | eukaryota>alveolata>ciliophora | Stentor | coeruleus | hypothetical protein SteCoe_12602 [Stentor coeruleus] |
| OMJ76388.1 | 469 | SteCoe_24256 | eukaryota>alveolata>ciliophora | Stentor | coeruleus | hypothetical protein SteCoe_24256 [Stentor coeruleus] |
| NP_001124164.1 | 469 | CALN,CALNA,CALNA1,CCN1,CNA1,IECEE,IECEE1,PPP2B | eukaryota>metazoa>chordata>vertebrata | Homo | sapiens | serine/threonine-protein phosphatase 2B catalytic subunit alpha isoform isoform 3 [Homo sapiens] |
| XP_003257536.1 | 469 | PPP3CA | eukaryota>metazoa>chordata>vertebrata | Nomascus | leucogenys | PREDICTED: serine/threonine-protein phosphatase 2B catalytic subunit alpha isoform isoform X4 [Nomascus leucogenys] |
| OMJ92782.1 | 468 | SteCoe_4416 | eukaryota>alveolata>ciliophora | Stentor | coeruleus | hypothetical protein SteCoe_4416 [Stentor coeruleus] |
| OMJ72382.1 | 468 | SteCoe_29197 | eukaryota>alveolata>ciliophora | Stentor | coeruleus | hypothetical protein SteCoe_29197 [Stentor coeruleus] |
| XP_001897709.1 | 468 | Bm1_31320 | eukaryota>metazoa>nematoda | Brugia | malayi | Serine/threonine-protein phosphatase 2B catalytic subunit 2, putative [Brugia malayi] |
| XP_004030479.1 | 467 | IMG5_160420 | eukaryota>alveolata>ciliophora | Ichthyophthirius | multifiliis | ser thr protein phosphatase family protein, putative [Ichthyophthirius multifiliis] |
| OMJ93859.1 | 467 | SteCoe_3048 | eukaryota>alveolata>ciliophora | Stentor | coeruleus | hypothetical protein SteCoe_3048 [Stentor coeruleus] |
| OMJ88575.1 | 467 | SteCoe_9437 | eukaryota>alveolata>ciliophora | Stentor | coeruleus | hypothetical protein SteCoe_9437 [Stentor coeruleus] |
| OMJ79408.1 | 467 | SteCoe_20563 | eukaryota>alveolata>ciliophora | Stentor | coeruleus | hypothetical protein SteCoe_20563 [Stentor coeruleus] |
| OMJ75859.1 | 467 | SteCoe_24923 | eukaryota>alveolata>ciliophora | Stentor | coeruleus | hypothetical protein SteCoe_24923 [Stentor coeruleus] |
| OMJ71377.1 | 467 | SteCoe_30439 | eukaryota>alveolata>ciliophora | Stentor | coeruleus | hypothetical protein SteCoe_30439 [Stentor coeruleus] |
| EJY73574.1 | 467 | OXYTRI_05294 | eukaryota>alveolata>ciliophora | Oxytricha | trifallax | Serine/threonine-protein phosphatase (macronuclear) [Oxytricha trifallax] |
| EJY69162.1 | 467 | OXYTRI_10219 | eukaryota>alveolata>ciliophora | Oxytricha | trifallax | Serine/threonine-protein phosphatase (macronuclear) [Oxytricha trifallax] |
| OMJ70037.1 | 467 | SteCoe_32075 | eukaryota>alveolata>ciliophora | Stentor | coeruleus | hypothetical protein SteCoe_32075 [Stentor coeruleus] |
| OMJ82487.1 | 466 | SteCoe_16839 | eukaryota>alveolata>ciliophora | Stentor | coeruleus | hypothetical protein SteCoe_16839 [Stentor coeruleus] |
| OMJ82460.1 | 466 | SteCoe_16869 | eukaryota>alveolata>ciliophora | Stentor | coeruleus | hypothetical protein SteCoe_16869 [Stentor coeruleus] |
| OMJ71024.1 | 466 | SteCoe_30870 | eukaryota>alveolata>ciliophora | Stentor | coeruleus | hypothetical protein SteCoe_30870 [Stentor coeruleus] |
| XP_009022549.1 | 466 | HELRODRAFT_67240 | eukaryota>metazoa>annelida | Helobdella | robusta | hypothetical protein HELRODRAFT_67240, partial [Helobdella robusta] |
| XP_014777765.1 | 466 | LOC106874522 | eukaryota>metazoa>mollusca | Octopus | bimaculoides | PREDICTED: serine/threonine-protein phosphatase 2B catalytic subunit 3-like isoform X2 [Octopus bimaculoides] |
| OMJ91329.1 | 465 | SteCoe_6122 | eukaryota>alveolata>ciliophora | Stentor | coeruleus | hypothetical protein SteCoe_6122 [Stentor coeruleus] |
| OMJ84516.1 | 465 | SteCoe_14321 | eukaryota>alveolata>ciliophora | Stentor | coeruleus | hypothetical protein SteCoe_14321 [Stentor coeruleus] |
| OMJ83404.1 | 465 | SteCoe_15672 | eukaryota>alveolata>ciliophora | Stentor | coeruleus | hypothetical protein SteCoe_15672 [Stentor coeruleus] |
| OMJ93795.1 | 464 | SteCoe_3116 | eukaryota>alveolata>ciliophora | Stentor | coeruleus | hypothetical protein SteCoe_3116 [Stentor coeruleus] |
| OMJ87850.1 | 464 | SteCoe_10331 | eukaryota>alveolata>ciliophora | Stentor | coeruleus | hypothetical protein SteCoe_10331 [Stentor coeruleus] |
| OMJ69203.1 | 464 | SteCoe_33127 | eukaryota>alveolata>ciliophora | Stentor | coeruleus | hypothetical protein SteCoe_33127 [Stentor coeruleus] |
| OMJ93604.1 | 462 | SteCoe_3443 | eukaryota>alveolata>ciliophora | Stentor | coeruleus | hypothetical protein SteCoe_3443 [Stentor coeruleus] |
| OMJ76290.1 | 461 | SteCoe_24386 | eukaryota>alveolata>ciliophora | Stentor | coeruleus | hypothetical protein SteCoe_24386 [Stentor coeruleus] |
| OMJ72513.1 | 461 | SteCoe_29033 | eukaryota>alveolata>ciliophora | Stentor | coeruleus | hypothetical protein SteCoe_29033 [Stentor coeruleus] |
| OMJ68518.1 | 461 | SteCoe_33996 | eukaryota>alveolata>ciliophora | Stentor | coeruleus | hypothetical protein SteCoe_33996 [Stentor coeruleus] |
| CDW72256.1 | 461 | Contig19151.g20310 | eukaryota>alveolata>ciliophora | Stylonychia | lemnae | serine threonine-protein phosphatase [Stylonychia lemnae] |
| OMJ77764.1 | 461 | SteCoe_22554 | eukaryota>alveolata>ciliophora | Stentor | coeruleus | hypothetical protein SteCoe_22554 [Stentor coeruleus] |
| OMJ88118.1 | 461 | SteCoe_10019 | eukaryota>alveolata>ciliophora | Stentor | coeruleus | hypothetical protein SteCoe_10019 [Stentor coeruleus] |
| XP_017174982.1 | 461 | 2900074D19Rik,Caln,Calna,CN,CnA | eukaryota>metazoa>chordata>vertebrata | Mus | musculus | PREDICTED: serine/threonine-protein phosphatase 2B catalytic subunit alpha isoform isoform X2 [Mus musculus] |
| XP_001628335.1 | 461 | NEMVEDRAFT_v1g245588 | eukaryota>metazoa>cnidaria | Nematostella | vectensis | predicted protein [Nematostella vectensis] |
| XP_017347414.1 | 460 | LOC108278499 | eukaryota>metazoa>chordata>vertebrata>actinopterygii | Ictalurus | punctatus | PREDICTED: serine/threonine-protein phosphatase 2B catalytic subunit alpha isoform-like isoform X3 [Ictalurus punctatus] |
| XP_014777766.1 | 460 | LOC106874522 | eukaryota>metazoa>mollusca | Octopus | bimaculoides | PREDICTED: serine/threonine-protein phosphatase 2B catalytic subunit alpha isoform-like isoform X3 [Octopus bimaculoides] |
| OMJ67745.1 | 458 | SteCoe_35009 | eukaryota>alveolata>ciliophora | Stentor | coeruleus | hypothetical protein SteCoe_35009 [Stentor coeruleus] |
| OMJ74062.1 | 458 | SteCoe_27115 | eukaryota>alveolata>ciliophora | Stentor | coeruleus | hypothetical protein SteCoe_27115 [Stentor coeruleus] |
| OMJ81469.1 | 458 | SteCoe_18088 | eukaryota>alveolata>ciliophora | Stentor | coeruleus | hypothetical protein SteCoe_18088 [Stentor coeruleus] |
| XP_017327656.1 | 458 | ppp3ca | eukaryota>metazoa>chordata>vertebrata>actinopterygii | Ictalurus | punctatus | PREDICTED: serine/threonine-protein phosphatase 2B catalytic subunit alpha isoform isoform X2 [Ictalurus punctatus] |
| EJY73882.1 | 456 | OXYTRI_04865 | eukaryota>alveolata>ciliophora | Oxytricha | trifallax | Serine/threonine-protein phosphatase (macronuclear) [Oxytricha trifallax] |
| XP_014777767.1 | 456 | LOC106874522 | eukaryota>metazoa>mollusca | Octopus | bimaculoides | PREDICTED: serine/threonine-protein phosphatase 2B catalytic subunit alpha isoform-like isoform X4 [Octopus bimaculoides] |
| CDW76840.1 | 454 | Contig14573.g15525 | eukaryota>alveolata>ciliophora | Stylonychia | lemnae | serine threonine-protein phosphatase [Stylonychia lemnae] |
| EPY41515.1 | 454 | AGDE_02409 | eukaryota>euglenozoa>kinetoplastida | Angomonas | deanei | serine/threonine-protein phosphatase [Angomonas deanei] |
| EPY28419.1 | 454 | AGDE_10391 | eukaryota>euglenozoa>kinetoplastida | Angomonas | deanei | protein phosphatase 3, catalytic subunit [Angomonas deanei] |
| XP_003257537.1 | 454 | PPP3CA | eukaryota>metazoa>chordata>vertebrata | Nomascus | leucogenys | PREDICTED: serine/threonine-protein phosphatase 2B catalytic subunit alpha isoform isoform X5 [Nomascus leucogenys] |
| XP_014677393.1 | 453 | LOC106817227 | eukaryota>metazoa | Priapulus | caudatus | PREDICTED: serine/threonine-protein phosphatase 2B catalytic subunit 3-like isoform X3 [Priapulus caudatus] |
| XP_015157925.1 | 452 | LOC101748851 | eukaryota>metazoa>chordata>vertebrata | Gallus | gallus | PREDICTED: serine/threonine-protein phosphatase 2B catalytic subunit gamma isoform X2 [Gallus gallus] |
| XP_002119971.2 | 452 | LOC100185597 | eukaryota>metazoa>chordata | Ciona | intestinalis | PREDICTED: serine/threonine-protein phosphatase 2B catalytic subunit alpha isoform-like [Ciona intestinalis] |
| XP_008858421.1 | 451 | ENU1_136630 | eukaryota>amoebozoa>entamoebidae | Entamoeba | nuttalli P19 | Ser/Thr protein phosphatase, putative [Entamoeba nuttalli P19] |
| NP_001021293.2 | 450 | tax-6 | eukaryota>metazoa>nematoda | Caenorhabditis | elegans | Serine/threonine-protein phosphatase 2B catalytic subunit [Caenorhabditis elegans] |
| XP_015311334.1 | 448 | PPP3CB | eukaryota>metazoa>chordata>vertebrata | Macaca | fascicularis | PREDICTED: serine/threonine-protein phosphatase 2B catalytic subunit beta isoform isoform X1 [Macaca fascicularis] |
| XP_015311333.1 | 448 | PPP3CB | eukaryota>metazoa>chordata>vertebrata | Macaca | fascicularis | PREDICTED: serine/threonine-protein phosphatase 2B catalytic subunit beta isoform isoform X1 [Macaca fascicularis] |
| XP_011773901.1 | 448 | TbgDal_VI940 | eukaryota>euglenozoa>kinetoplastida | Trypanosoma | brucei gambiense DAL972 | kinetoplastid-specific phospho-protein phosphatase [Trypanosoma brucei gambiense DAL972] |
| XP_845245.1 | 448 | Tb06.3A7.760 | eukaryota>euglenozoa>kinetoplastida | Trypanosoma | brucei brucei TREU927 | hypothetical protein, conserved [Trypanosoma brucei brucei TREU927] |
| XP_015311335.1 | 447 | PPP3CB | eukaryota>metazoa>chordata>vertebrata | Macaca | fascicularis | PREDICTED: serine/threonine-protein phosphatase 2B catalytic subunit beta isoform isoform X2 [Macaca fascicularis] |
| OHS94116.1 | 447 | TRFO_11403 | eukaryota>parabasalia | Tritrichomonas | foetus | hypothetical protein TRFO_11403 [Tritrichomonas foetus] |
| XP_015775231.1 | 446 | LOC107353402 | eukaryota>metazoa>cnidaria | Acropora | digitifera | PREDICTED: serine/threonine-protein phosphatase 2B catalytic subunit alpha isoform-like [Acropora digitifera] |
| XP_810491.1 | 445 | Tc00.1047053510755.138 | eukaryota>euglenozoa>kinetoplastida | Trypanosoma | cruzi strain CL Brener | serine/threonine protein phosphatase [Trypanosoma cruzi strain CL Brener] |
| XP_005562857.1 | 445 | PPP3CC | eukaryota>metazoa>chordata>vertebrata | Macaca | fascicularis | PREDICTED: serine/threonine-protein phosphatase 2B catalytic subunit gamma isoform isoform X5 [Macaca fascicularis] |
| XP_012355005.1 | 444 | PPP3CA | eukaryota>metazoa>chordata>vertebrata | Nomascus | leucogenys | PREDICTED: serine/threonine-protein phosphatase 2B catalytic subunit alpha isoform isoform X6 [Nomascus leucogenys] |
| KPI86351.1 | 444 | ABL78_4577 | eukaryota>euglenozoa>kinetoplastida | Leptomonas | seymouri | hypothetical protein ABL78_4577 [Leptomonas seymouri] |
| XP_002471490.1 | 443 | PP2B-2 | eukaryota>fungi>basidiomycota | Postia | placenta Mad-698-R | serine/threonine-protein phosphatase 2B, partial [Postia placenta Mad-698-R] |
| XP_015157924.1 | 443 | LOC101748851 | eukaryota>metazoa>chordata>vertebrata | Gallus | gallus | PREDICTED: serine/threonine-protein phosphatase 2B catalytic subunit gamma isoform X1 [Gallus gallus] |
| OHT14346.1 | 440 | TRFO_15250 | eukaryota>parabasalia | Tritrichomonas | foetus | Serine/threonine-protein phosphatase 2B catalytic subunit alpha isoform [Tritrichomonas foetus] |
| XP_016869099.1 | 439 | CALNA3,CNA3,PP2Bgamma | eukaryota>metazoa>chordata>vertebrata | Homo | sapiens | PREDICTED: serine/threonine-protein phosphatase 2B catalytic subunit gamma isoform isoform X1 [Homo sapiens] |
| XP_015311336.1 | 439 | PPP3CB | eukaryota>metazoa>chordata>vertebrata | Macaca | fascicularis | PREDICTED: serine/threonine-protein phosphatase 2B catalytic subunit beta isoform isoform X3 [Macaca fascicularis] |
| XP_001453725.1 | 438 | GSPATT00020009001 | eukaryota>alveolata>ciliophora | Paramecium | tetraurelia strain d4-2 | hypothetical protein (macronuclear) [Paramecium tetraurelia strain d4-2] |
| XP_014677397.1 | 438 | LOC106817227 | eukaryota>metazoa | Priapulus | caudatus | PREDICTED: serine/threonine-protein phosphatase 2B catalytic subunit alpha isoform-like isoform X4 [Priapulus caudatus] |
| XP_015311338.1 | 438 | PPP3CB | eukaryota>metazoa>chordata>vertebrata | Macaca | fascicularis | PREDICTED: serine/threonine-protein phosphatase 2B catalytic subunit beta isoform isoform X5 [Macaca fascicularis] |
| XP_015311337.1 | 438 | PPP3CB | eukaryota>metazoa>chordata>vertebrata | Macaca | fascicularis | PREDICTED: serine/threonine-protein phosphatase 2B catalytic subunit beta isoform isoform X4 [Macaca fascicularis] |
| EXX61187.1 | 437 | RirG_173620 | eukaryota>fungi | Rhizophagus | irregularis DAOM 197198w | Cmp2p [Rhizophagus irregularis DAOM 197198w] |
| EXX61186.1 | 437 | RirG_173620 | eukaryota>fungi | Rhizophagus | irregularis DAOM 197198w | Cmp2p [Rhizophagus irregularis DAOM 197198w] |
| XP_009021900.1 | 437 | HELRODRAFT_157436 | eukaryota>metazoa>annelida | Helobdella | robusta | hypothetical protein HELRODRAFT_157436 [Helobdella robusta] |
| XP_005562858.1 | 435 | PPP3CC | eukaryota>metazoa>chordata>vertebrata | Macaca | fascicularis | PREDICTED: serine/threonine-protein phosphatase 2B catalytic subunit gamma isoform isoform X6 [Macaca fascicularis] |
| ETO26625.1 | 435 | RFI_10511 | eukaryota>rhizaria | Reticulomyxa | filosa | calcineurin A [Reticulomyxa filosa] |
| XP_011678785.1 | 434 | LOC588695 | eukaryota>metazoa>echinodermata | Strongylocentrotus | purpuratus | PREDICTED: serine/threonine-protein phosphatase 2B catalytic subunit alpha isoform isoform X8 [Strongylocentrotus purpuratus] |
| XP_015157926.1 | 433 | LOC101748851 | eukaryota>metazoa>chordata>vertebrata | Gallus | gallus | PREDICTED: serine/threonine-protein phosphatase 2B catalytic subunit gamma isoform X3 [Gallus gallus] |
| XP_011776228.1 | 431 | TbgDal_IX270 | eukaryota>euglenozoa>kinetoplastida | Trypanosoma | brucei gambiense DAL972 | serine/threonine protein phosphatase, putative [Trypanosoma brucei gambiense DAL972] |
| XP_803443.1 | 431 | Tb09.160.0480 | eukaryota>euglenozoa>kinetoplastida | Trypanosoma | brucei brucei TREU927 | serine/threonine protein phosphatase, putative [Trypanosoma brucei brucei TREU927] |
| XP_006456164.1 | 429 | AGABI2DRAFT_77099 | eukaryota>fungi>basidiomycota | Agaricus | bisporus var. bisporus H97 | hypothetical protein AGABI2DRAFT_77099, partial [Agaricus bisporus var. bisporus H97] |
| XP_015311339.1 | 429 | PPP3CB | eukaryota>metazoa>chordata>vertebrata | Macaca | fascicularis | PREDICTED: serine/threonine-protein phosphatase 2B catalytic subunit beta isoform isoform X6 [Macaca fascicularis] |
| XP_015311340.1 | 428 | PPP3CB | eukaryota>metazoa>chordata>vertebrata | Macaca | fascicularis | PREDICTED: serine/threonine-protein phosphatase 2B catalytic subunit beta isoform isoform X7 [Macaca fascicularis] |
| XP_018653155.1 | 427 | Smp_141950 | eukaryota>metazoa | Schistosoma | mansoni | putative protein phosphatase-2b [Schistosoma mansoni] |
| XP_016871868.1 | 426 | CALNA2,CALNB,CNA2,PP2Bbeta | eukaryota>metazoa>chordata>vertebrata | Homo | sapiens | serine/threonine-protein phosphatase 2B catalytic subunit beta isoform isoform X2 [Homo sapiens] |
| XP_015654428.1 | 426 | ABB37_08351 | eukaryota>euglenozoa>kinetoplastida | Leptomonas | pyrrhocoris | hypothetical protein ABB37_08351 [Leptomonas pyrrhocoris] |
| KPI87742.1 | 425 | ABL78_3151 | eukaryota>euglenozoa>kinetoplastida | Leptomonas | seymouri | putative serine/threonine protein phosphatase 2B catalytic subunit A2 [Leptomonas seymouri] |

[illegible]

| | | | | | | |
|---|---|---|---|---|---|---|
| XP_004261838.1 | 179 | EIN_253310 | eukaryota>amoebozoa>entamoebidae | Entamoeba | invadens IP1 | calcineurin subunit B, putative [Entamoeba invadens IP1] |
| XP_008860323.1 | 179 | ENU1_200580 | eukaryota>amoebozoa>entamoebidae | Entamoeba | nuttalli P19 | calcineurin B subunit, putative [Entamoeba nuttalli P19] |
| XP_011777215.1 | 179 | TbgDal_X300 | eukaryota>euglenozoa>kinetoplastida | Trypanosoma | brucei gambiense DAL972 | calcineurin B subunit, putative [Trypanosoma brucei gambiense DAL972] |
| NP_001004025.1 | 179 | CaNB2,CnB2 | eukaryota>metazoa>chordata>vertebrata | Mus | musculus | calcineurin subunit B type 2 [Mus musculus] |
| XP_016612902.1 | 178 | SPPG_00562 | eukaryota>fungi>chytridiomycota | Spizellomyces | punctatus DAOM BR117 | hypothetical protein SPPG_00562 [Spizellomyces punctatus DAOM BR117] |
| XP_004349371.2 | 178 | CAOG_02621 | eukaryota>ichthyosporea | Capsaspora | owczarzaki ATCC 30864 | calcium-binding protein p22 [Capsaspora owczarzaki ATCC 30864] |
| CEM08560.1 | 177 | Vbra_14626 | eukaryota>alveolata | Vitrella | brassicaformis CCMP3155 | unnamed protein product [Vitrella brassicaformis CCMP3155] |
| XP_001348666.2 | 177 | PF3D7_1451700 | eukaryota>alveolata>apicomplexa | Plasmodium | falciparum 3D7 | calcineurin subunit B [Plasmodium falciparum 3D7] |
| XP_001615872.1 | 177 | PVX_117895 | eukaryota>alveolata>apicomplexa | Plasmodium | vivax | calcineurin B subunit isoform 1, putative [Plasmodium vivax] |
| XP_002371117.1 | 177 | TGME49_213800 | eukaryota>alveolata>apicomplexa | Toxoplasma | gondii ME49 | protein phosphatase 2b regulatory subunit, putative [Toxoplasma gondii ME49] |
| XP_003881594.1 | 177 | NCLIV_013540 | eukaryota>alveolata>apicomplexa | Neospora | caninum Liverpool | hypothetical protein NCLIV_013540 [Neospora caninum Liverpool] |
| XP_008884045.1 | 177 | HHA_213800 | eukaryota>alveolata>apicomplexa | Hammondia | hammondi | protein phosphatase 2b regulatory subunit, putative [Hammondia hammondi] |
| XP_013234209.1 | 177 | ETH_00025715 | eukaryota>alveolata>apicomplexa | Eimeria | tenella | protein phosphatase 2b regulatory subunit, putative [Eimeria tenella] |
| OMJ81238.1 | 177 | SteCoe_18356 | eukaryota>alveolata>ciliophora | Stentor | coeruleus | hypothetical protein SteCoe_18356 [Stentor coeruleus] |
| XP_009310883.1 | 176 | DQ04_03191000 | eukaryota>euglenozoa>kinetoplastida | Trypanosoma | grayi | calcineurin B subunit [Trypanosoma grayi] |
| XP_808544.1 | 176 | Tc00.1047053506869.50 | eukaryota>euglenozoa>kinetoplastida | Trypanosoma | cruzi strain CL Brener | calcineurin B subunit [Trypanosoma cruzi strain CL Brener] |
| XP_812709.1 | 176 | Tc00.1047053510519.60 | eukaryota>euglenozoa>kinetoplastida | Trypanosoma | cruzi strain CL Brener | calcineurin B subunit [Trypanosoma cruzi strain CL Brener] |
| XP_019015494.1 | 176 | PICMEDRAFT_18295 | eukaryota>fungi>ascomycota | Pichia | membranifaciens NRRL Y-2026 | hypothetical protein PICMEDRAFT_18295 [Pichia membranifaciens NRRL Y-2026] |
| Pisp1000007246 | 176 | Pisp1000007246 | eukaryota>fungi>neocallimastigomycota | Piromyces | sp | >Pisp1000007246 |
| EST47010.1 | 176 | SS50377_12965 | eukaryota>diplomonadida | Spironucleus | salmonicida | Calcineurin regulatory subunit B [Spironucleus salmonicida] |
| XP_006459824.1 | 175 | AGABI2DRAFT_135141 | eukaryota>fungi>basidiomycota | Agaricus | bisporus var. bisporus H97 | hypothetical protein AGABI2DRAFT_135141 [Agaricus bisporus var. bisporus H97] |
| XP_021868731.1 | 175 | BD324DRAFT_636044 | eukaryota>fungi>basidiomycota | Kockovaella | imperatae | putative calcium-dependent protein serine/threonine phosphatase [Kockovaella imperatae] |
| ORY76768.1 | 175 | LY90DRAFT_377314 | eukaryota>fungi>chytridiomycota | Neocallimastix | californiae | EF-hand [Neocallimastix californiae] |
| ORY75892.1 | 175 | LY90DRAFT_377653 | eukaryota>fungi>chytridiomycota | Neocallimastix | californiae | EF-hand [Neocallimastix californiae] |
| CCW70383.1 | 175 | GSHART1_T00007958001 | eukaryota>euglenozoa>kinetoplastida | Phytomonas | sp. isolate Hart1 | unnamed protein product [Phytomonas sp. isolate Hart1] |
| EPY36052.1 | 175 | STCU_00779 | eukaryota>euglenozoa>kinetoplastida | Strigomonas | culicis | protein phosphatase 3, regulatory subunit [Strigomonas culicis] |
| KPI84786.1 | 175 | ABL78_6169 | eukaryota>euglenozoa>kinetoplastida | Leptomonas | seymouri | putative calcineurin B subunit [Leptomonas seymouri] |
| XP_001465459.1 | 175 | LINJ_21_1990 | eukaryota>euglenozoa>kinetoplastida | Leishmania | infantum JPCM5 | putative calcineurin B subunit [Leishmania infantum JPCM5] |
| XP_001564893.1 | 175 | LBRM_21_1920 | eukaryota>euglenozoa>kinetoplastida | Leishmania | braziliensis MHOM/BR/75/M2904 | putative calcineurin B subunit [Leishmania braziliensis MHOM/BR/75/M2904] |
| XP_001683127.1 | 175 | LMJF_21_1630 | eukaryota>euglenozoa>kinetoplastida | Leishmania | major strain Friedlin | putative calcineurin B subunit [Leishmania major strain Friedlin] |
| XP_003860716.1 | 175 | LDBPK_211990 | eukaryota>euglenozoa>kinetoplastida | Leishmania | donovani | calcineurin B subunit, putative [Leishmania donovani] |
| XP_003875430.1 | 175 | LMXM_21_1630 | eukaryota>euglenozoa>kinetoplastida | Leishmania | mexicana MHOM/GT/2001/U1103 | putative calcineurin B subunit [Leishmania mexicana MHOM/GT/2001/U1103] |
| XP_010698978.1 | 175 | CNB | eukaryota>euglenozoa>kinetoplastida | Leishmania | panamensis | calcineurin B subunit, putative [Leishmania panamensis] |
| XP_015656821.1 | 175 | ABB37_06498 | eukaryota>euglenozoa>kinetoplastida | Leptomonas | pyrrhocoris | calcineurin b subunit (CNB) [Leptomonas pyrrhocoris] |
| XP_015656822.1 | 175 | ABB37_06498 | eukaryota>euglenozoa>kinetoplastida | Leptomonas | pyrrhocoris | calcineurin b subunit (CNB) [Leptomonas pyrrhocoris] |
| XP_015656823.1 | 175 | ABB37_06498 | eukaryota>euglenozoa>kinetoplastida | Leptomonas | pyrrhocoris | calcineurin b subunit (CNB) [Leptomonas pyrrhocoris] |
| KNE65487.1 | 175 | AMAG_11106 | eukaryota>fungi | Allomyces | macrogynus ATCC 38327 | calcineurin subunit B [Allomyces macrogynus ATCC 38327] |
| KNE72653.1 | 175 | AMAG_16414 | eukaryota>fungi | Allomyces | macrogynus ATCC 38327 | calcineurin subunit B [Allomyces macrogynus ATCC 38327] |
| NP_012731.1 | 175 | CRV1,YCN2 | eukaryota>fungi>ascomycota | Saccharomyces | cerevisiae S288C | calcineurin regulatory subunit B [Saccharomyces cerevisiae S288C] |
| XP_002837238.1 | 175 | GSTUM_00005170001 | eukaryota>fungi>ascomycota | Tuber | melanosporum Mel28 | hypothetical protein [Tuber melanosporum Mel28] |
| XP_453201.1 | 175 | KLLA0_D02992g | eukaryota>fungi>ascomycota | Kluyveromyces | lactis | uncharacterized protein KLLA0_D02992g [Kluyveromyces lactis] |
| Chet1000004836 | 175 | Chet1000004836 | eukaryota>fungi>ascomycota | Cochliobolus | heterostrophus | >Chet1000004836 |
| EPT03113.1 | 175 | FOMPIDRAFT_1022481 | eukaryota>fungi>basidiomycota | Fomitopsis | pinicola FP-58527 SS1 | hypothetical protein FOMPIDRAFT_1022481 [Fomitopsis pinicola FP-58527 SS1] |
| XP_001839327.2 | 175 | CC1G_08194 | eukaryota>fungi>basidiomycota | Coprinopsis | cinerea okayama7#130 | calcium-dependent protein serine/threonine phosphatase [Coprinopsis cinerea okayama7#130] |
| XP_001884421.1 | 175 | LACBIDRAFT_291275 | eukaryota>fungi>basidiomycota | Laccaria | bicolor S238N-H82 | predicted protein [Laccaria bicolor S238N-H82] |
| XP_002473674.1 | 175 | CAR1 | eukaryota>fungi>basidiomycota | Postia | placenta Mad-698-R | calcium/calmodulin-dependent protein phosphatase [Postia placenta Mad-698-R] |
| XP_003322771.1 | 175 | PGTG_04308 | eukaryota>fungi>basidiomycota | Puccinia | graminis f. sp. tritici CRL 75-36-700-3 | calcineurin subunit B [Puccinia graminis f. sp. tritici CRL 75-36-700-3] |
| XP_006958264.1 | 175 | WALSEDRAFT_60317 | eukaryota>fungi>basidiomycota | Wallemia | mellicola CBS 633.66 | calcium-dependent protein serine/threonine phosphatase [Wallemia mellicola CBS 633.66] |
| XP_007333662.1 | 175 | AGABI1DRAFT_116228 | eukaryota>fungi>basidiomycota | Agaricus | bisporus var. burnettii JB137-S8 | hypothetical protein AGABI1DRAFT_116228 [Agaricus bisporus var. burnettii JB137-S8] |
| XP_011387312.1 | 175 | UMAG_10226 | eukaryota>fungi>basidiomycota | Ustilago | maydis 521 | putative calcineurin regulatory subunit B [Ustilago maydis 521] |
| XP_012049171.1 | 175 | CNAG_00888 | eukaryota>fungi>basidiomycota | Cryptococcus | neoformans var. grubii H99 | calcineurin subunit B [Cryptococcus neoformans var. grubii H99] |
| XP_013240503.1 | 175 | K437DRAFT_259661 | eukaryota>fungi>basidiomycota | Tilletiaria | anomala UBC 951 | putative CNB1-calcineurin B, regulatory subunit [Tilletiaria anomala UBC 951] |
| XP_014571409.1 | 175 | L969DRAFT_101264 | eukaryota>fungi>basidiomycota | Mixia | osmundae IAM 14324 | hypothetical protein L969DRAFT_101264 [Mixia osmundae IAM 14324] |
| XP_017992920.1 | 175 | Malapachy_2482 | eukaryota>fungi>basidiomycota | Malassezia | pachydermatis | calcium calmodulin-dependent protein phosphatase [Malassezia pachydermatis] |
| XP_018273257.1 | 175 | RHOBADRAFT_64678 | eukaryota>fungi>basidiomycota | Rhodotorula | graminis WP1 | hypothetical protein RHOBADRAFT_64678 [Rhodotorula graminis WP1] |
| XP_018280614.1 | 175 | CC85DRAFT_311339 | eukaryota>fungi>basidiomycota | Cutaneotrichosporon | oleaginosum | calcium-dependent protein serine/threonine phosphatase [Cutaneotrichosporon oleaginosum] |
| XP_006677028.1 | 175 | BATDEDRAFT_86479 | eukaryota>fungi>chytridiomycota | Batrachochytrium | dendrobatidis JAM81 | hypothetical protein BATDEDRAFT_86479 [Batrachochytrium dendrobatidis JAM81] |
| XP_016609494.1 | 175 | SPPG_03259 | eukaryota>fungi>chytridiomycota | Spizellomyces | punctatus DAOM BR117 | calcineurin subunit B, variant 2 [Spizellomyces punctatus DAOM BR117] |
| XP_016609495.1 | 175 | SPPG_03259 | eukaryota>fungi>chytridiomycota | Spizellomyces | punctatus DAOM BR117 | calcineurin subunit B, variant 1 [Spizellomyces punctatus DAOM BR117] |
| XP_016609496.1 | 175 | SPPG_03259 | eukaryota>fungi>chytridiomycota | Spizellomyces | punctatus DAOM BR117 | calcineurin subunit B [Spizellomyces punctatus DAOM BR117] |
| XP_009305321.1 | 175 | canb1,ppp3r1,si:ch211-274c22.8 | eukaryota>metazoa>chordata>vertebrata>actinopterygii | Danio | rerio | calcineurin subunit B type 1 isoform X1 [Danio rerio] |
| OZJ06015.1 | 174 | BZG36_01073 | eukaryota>fungi | Bifiguratus | adelaidae | Calcineurin subunit B [Bifiguratus adelaidae] |
| ESA14576.1 | 174 | GLOINDRAFT_78398 | eukaryota>fungi | Rhizophagus | irregularis DAOM 181602 | hypothetical protein GLOINDRAFT_78398 [Rhizophagus irregularis DAOM 181602] |
| EPY40284.1 | 174 | AGDE_03644 | eukaryota>euglenozoa>kinetoplastida | Angomonas | deanei | protein phosphatase 3, regulatory subunit [Angomonas deanei] |

[illegible]

supplement_text_new.txt

| | | | | | | |
|---|---|---|---|---|---|---|
| OMJ88964.1 | 384 | SteCoe_8973 | eukaryota>alveolata>ciliophora | Stentor | coeruleus | hypothetical protein SteCoe_8973 [Stentor coeruleus] |
| XP_001744647.1 | 384 | MONBRDRAFT_20407 | eukaryota>choanoflagellida | Monosiga | brevicollis MX1 | hypothetical protein [Monosiga brevicollis MX1] |
| XP_003275837.1 | 384 | CALR3 | eukaryota>metazoa>chordata>vertebrata | Nomascus | leucogenys | PREDICTED: calreticulin-3 [Nomascus leucogenys] |
| XP_016495583.1 | 381 | LOC107814655 | eukaryota>viridiplantae | Nicotiana | tabacum | PREDICTED: calreticulin isoform X2 [Nicotiana tabacum] |
| NP_082776.2 | 380 | 1700031L01Rik,6330586I20Rik,Crt2,cspn | eukaryota>metazoa>chordata>vertebrata | Mus | musculus | calreticulin-3 isoform 1 precursor [Mus musculus] |
| XP_013397065.1 | 380 | LOC106163893 | eukaryota>metazoa | Lingula | anatina | calreticulin-like [Lingula anatina] |
| XP_001312065.1 | 376 | TVAG_115390 | eukaryota>parabasalia | Trichomonas | vaginalis G3 | Calreticulin family protein [Trichomonas vaginalis G3] |
| XP_002291609.1 | 376 | THAPSDRAFT_34810 | eukaryota>stramenopiles | Thalassiosira | pseudonana CCMP1335 | calreticulin-like protein, partial [Thalassiosira pseudonana CCMP1335] |
| XP_001300977.1 | 375 | TVAG_207320 | eukaryota>parabasalia | Trichomonas | vaginalis G3 | Calreticulin family protein [Trichomonas vaginalis G3] |
| EPZ32318.1 | 370 | O9G_002159 | eukaryota>fungi | Rozella | allomycis CSF55 | Calreticulin domain-containing protein [Rozella allomycis CSF55] |
| NP_973793.1 | 370 | A.,thaliana,calreticulin,3,AtCRT3,calreticulin,3,CALRETICULIN,3,EBS2,EMS-MUTAGENIZED,BRI1,SUPPRESSOR,2,PRIORITY,IN,SWEET,LIFE,1,PSL1,T27G7.13,T27G7_13 | eukaryota>viridiplantae | Arabidopsis | thaliana | calreticulin 3 [Arabidopsis thaliana] |
| OLP93827.1 | 369 | AK812_SmicGene24221 | eukaryota>alveolata | Symbiodinium | microadriaticum | Calreticulin [Symbiodinium microadriaticum] |
| XP_001640172.1 | 366 | NEMVEDRAFT_v1g61368 | eukaryota>metazoa>cnidaria | Nematostella | vectensis | predicted protein, partial [Nematostella vectensis] |
| KXJ26714.1 | 363 | Crc | eukaryota>metazoa>cnidaria | Exaiptasia | pallida | Calreticulin [Exaiptasia pallida] |
| XP_006824896.1 | 360 | LOC100375401 | eukaryota>metazoa>hemichordata | Saccoglossus | kowalevskii | PREDICTED: calreticulin-like isoform X2 [Saccoglossus kowalevskii] |
| Pram1000014320 | 352 | Pram1000014320 | eukaryota>stramenopiles | Phytophthora | ramorum | >Pram1000014320 |
| XP_016611705.1 | 350 | SPPG_01136 | eukaryota>fungi>chytridiomycota | Spizellomyces | punctatus DAOM BR117 | hypothetical protein SPPG_01136 [Spizellomyces punctatus DAOM BR117] |
| EPY31781.1 | 343 | STCU_03244 | eukaryota>euglenozoa>kinetoplastida | Strigomonas | culicis | calreticulin [Strigomonas culicis] |
| XP_006509824.1 | 336 | 1700031L01Rik,6330586I20Rik,Crt2,cspn | eukaryota>metazoa>chordata>vertebrata | Mus | musculus | PREDICTED: calreticulin-3 isoform X1 [Mus musculus] |
| Smed1000027960 | 331 | Smed1000027960 | eukaryota>metazoa>platyhelminthe | Schmidtea | mediterranea | >Smed1000027960 |
| XP_001734007.1 | 318 | EDI_348940 | eukaryota>amoebozoa>entamoebidae | Entamoeba | dispar SAW760 | Calreticulin precursor [Entamoeba dispar SAW760] |
| NP_001272825.1 | 313 | 4933403L16Rik,AU017583 | eukaryota>metazoa>chordata>vertebrata | Mus | musculus | calreticulin 4 isoform c [Mus musculus] |
| NP_001272826.1 | 313 | 4933403L16Rik,AU017583 | eukaryota>metazoa>chordata>vertebrata | Mus | musculus | calreticulin 4 isoform c [Mus musculus] |
| NP_001272827.1 | 313 | 4933403L16Rik,AU017583 | eukaryota>metazoa>chordata>vertebrata | Mus | musculus | calreticulin 4 isoform c [Mus musculus] |
| XP_017175400.1 | 313 | 4933403L16Rik,AU017583 | eukaryota>metazoa>chordata>vertebrata | Mus | musculus | PREDICTED: calreticulin 4 isoform X1 [Mus musculus] |
| XP_009038526.1 | 311 | AURANDRAFT_65433 | eukaryota>stramenopiles | Aureococcus | anophagefferens | expressed protein [Aureococcus anophagefferens] |
| CDW86807.1 | 298 | Contig8228.g8773 | eukaryota>alveolata>ciliophora | Stylonychia | lemnae | calreticulin precursor [Stylonychia lemnae] |
| XP_018115337.1 | 275 | LOC108714990 | eukaryota>metazoa>chordata>vertebrata | Xenopus | laevis | PREDICTED: calreticulin-like [Xenopus laevis] |
| NP_084058.2 | 272 | 1700031L01Rik,6330586I20Rik,Crt2,cspn | eukaryota>metazoa>chordata>vertebrata | Mus | musculus | calreticulin-3 isoform 2 [Mus musculus] |
| XP_013892903.1 | 254 | MNEG_14079 | eukaryota>viridiplantae>chlorophyta | Monoraphidium | neglectum | Calreticulin Flags: Precursor [Monoraphidium neglectum] |
| XP_002900130.1 | 235 | PITG_12738 | eukaryota>stramenopiles | Phytophthora | infestans T30-4 | conserved hypothetical protein [Phytophthora infestans T30-4] |
| XP_002769367.1 | 224 | Pmar_PMAR004240 | eukaryota>alveolata | Perkinsus | marinus ATCC 50983 | Calreticulin precursor, putative, partial [Perkinsus marinus ATCC 50983] |
| ETO20325.1 | 215 | RFI_16892 | eukaryota>rhizaria | Reticulomyxa | filosa | Calreticulin precursor, partial [Reticulomyxa filosa] |
| Smed1000013190 | 159 | Smed1000013190 | eukaryota>metazoa>platyhelminthe | Schmidtea | mediterranea | >Smed1000013190 |

K. Calnexin

| | | | | | | |
|---|---|---|---|---|---|---|
| XP_006453987.1 | 1485 | AGABI2DRAFT_181983 | eukaryota>fungi>basidiomycota | Agaricus | bisporus var. bisporus H97 | hypothetical protein AGABI2DRAFT_181983 [Agaricus bisporus var. bisporus H97] |
| OWA53125.1 | 1248 | BV898_17558 | eukaryota>metazoa | Hypsibius | dujardini | Calnexin [Hypsibius dujardini] |
| XP_002674608.1 | 1118 | NAEGRDRAFT_70209 | eukaryota>heterolobosea | Naegleria | gruberi strain NEG-M | predicted protein [Naegleria gruberi strain NEG-M] |
| XP_011618359.1 | 718 | clgn | eukaryota>metazoa>chordata>vertebrata>actinopterygii | Takifugu | rubripes | PREDICTED: calmegin [Takifugu rubripes] |
| XP_005536371.1 | 660 | CYME_CMI213C | eukaryota>rhodophyta | Cyanidioschyzon | merolae strain 10D | calnexin [Cyanidioschyzon merolae strain 10D] |
| XP_004940998.1 | 655 | CLGN | eukaryota>metazoa>chordata>vertebrata | Gallus | gallus | PREDICTED: calmegin isoform X1 [Gallus gallus] |
| Chet1000006749 | 655 | Chet1000006749 | eukaryota>fungi>ascomycota | Cochliobolus | heterostrophus | >Chet1000006749 |
| XP_004940999.1 | 652 | CLGN | eukaryota>metazoa>chordata>vertebrata | Gallus | gallus | calmegin isoform X2 [Gallus gallus] |
| XP_018087171.1 | 647 | LOC108699598 | eukaryota>metazoa>chordata>vertebrata | Xenopus | laevis | PREDICTED: calmegin-like isoform X1 [Xenopus laevis] |
| XP_018087179.1 | 647 | LOC108699598 | eukaryota>metazoa>chordata>vertebrata | Xenopus | laevis | PREDICTED: calmegin-like isoform X1 [Xenopus laevis] |
| XP_011532966.1 | 646 | CNX,IP90,P90 | eukaryota>metazoa>chordata>vertebrata | Homo | sapiens | calnexin isoform X3 [Homo sapiens] |
| XP_004941000.1 | 644 | CLGN | eukaryota>metazoa>chordata>vertebrata | Gallus | gallus | calmegin isoform X3 [Gallus gallus] |
| XP_011132126.1 | 643 | GNI_129870 | eukaryota>alveolata>apicomplexa | Gregarina | niphandrodes | calreticulin family protein [Gregarina niphandrodes] |
| XP_013232333.1 | 640 | ETH_00022665 | eukaryota>alveolata>apicomplexa | Eimeria | tenella | calnexin, putative [Eimeria tenella] |
| NP_573131.1 | 639 | CG9906,Cnx,Dmel\CG9906,Xcbp1 | eukaryota>metazoa>hexapoda | Drosophila | melanogaster | calnexin 14D [Drosophila melanogaster] |
| XP_017319913.1 | 634 | clgn | eukaryota>metazoa>chordata>vertebrata>actinopterygii | Ictalurus | punctatus | PREDICTED: calmegin isoform X1 [Ictalurus punctatus] |
| XP_017319914.1 | 634 | clgn | eukaryota>metazoa>chordata>vertebrata>actinopterygii | Ictalurus | punctatus | PREDICTED: calmegin isoform X1 [Ictalurus punctatus] |
| XP_017319915.1 | 634 | clgn | eukaryota>metazoa>chordata>vertebrata>actinopterygii | Ictalurus | punctatus | PREDICTED: calmegin isoform X1 [Ictalurus punctatus] |
| XP_017319916.1 | 633 | clgn | eukaryota>metazoa>chordata>vertebrata>actinopterygii | Ictalurus | punctatus | PREDICTED: calmegin isoform X2 [Ictalurus punctatus] |
| NP_001071009.1 | 630 | canx,wu:fj24b04,zgc:153946 | eukaryota>metazoa>chordata>vertebrata>actinopterygii | Danio | rerio | calmegin precursor [Danio rerio] |
| XP_004941001.1 | 629 | CLGN | eukaryota>metazoa>chordata>vertebrata | Gallus | gallus | calmegin isoform X5 [Gallus gallus] |
| XP_005160060.1 | 629 | canx,fj49d10,wu:fj24b04,wu:fj49d10,zgc:153946 | eukaryota>metazoa>chordata>vertebrata>actinopterygii | Danio | rerio | calmegin isoform X1 [Danio rerio] |
| XP_455100.1 | 628 | KLLA0_F00462g | eukaryota>fungi>ascomycota | Kluyveromyces | lactis | uncharacterized protein KLLA0_F00462g [Kluyveromyces lactis] |
| XP_004941002.1 | 626 | CLGN | eukaryota>metazoa>chordata>vertebrata | Gallus | gallus | calmegin isoform X6 [Gallus gallus] |
| KNE60018.1 | 623 | AMAG_05457 | eukaryota>fungi | Allomyces | macrogynus ATCC 38327 | hypothetical protein AMAG_05457 [Allomyces macrogynus ATCC 38327] |
| KNE55367.1 | 622 | AMAG_01267 | eukaryota>fungi | Allomyces | macrogynus ATCC 38327 | hypothetical protein AMAG_01267 [Allomyces macrogynus ATCC 38327] |
| NP_499176.1 | 619 | cnx-1 | eukaryota>metazoa>nematoda | Caenorhabditis | elegans | Calnexin [Caenorhabditis elegans] |
| XP_001900533.1 | 615 | Bm1_45355 | eukaryota>metazoa>nematoda | Brugia | malayi | Calreticulin family protein [Brugia malayi] |
| XP_011246598.1 | 615 | 4930459O04Rik,AI528775,Cln | eukaryota>metazoa>chordata>vertebrata | Mus | musculus | PREDICTED: calmegin isoform X1 [Mus musculus] |
| XP_011246599.1 | 615 | 4930459O04Rik,AI528775,Cln | eukaryota>metazoa>chordata>vertebrata | Mus | musculus | PREDICTED: calmegin isoform X1 [Mus musculus] |
| Capi1000013491 | 614 | Capi1000013491 | eukaryota>metazoa>annelida | Capitella | spI | >Capi1000013491 |
| NP_001079415.1 | 614 | canx,canx-a,canx-b,clnx,cnx | eukaryota>metazoa>chordata>vertebrata | Xenopus | laevis | calnexin S homeolog precursor [Xenopus laevis] |
| XP_018106103.1 | 613 | canx,canx-a,canx-b,clnx,cnx | eukaryota>metazoa>chordata>vertebrata | Xenopus | laevis | PREDICTED: calnexin S homeolog isoform X1 [Xenopus laevis] |
| XP_002956983.1 | 612 | VOLCADRAFT_77379 | eukaryota>viridiplantae>chlorophyta | Volvox | carteri f. nagariensis | hypothetical protein VOLCADRAFT_77379 [Volvox carteri f. nagariensis] |
| XP_018098777.1 | 612 | LOC108706675 | eukaryota>metazoa>chordata>vertebrata | Xenopus | laevis | PREDICTED: calnexin-like isoform X1 [Xenopus laevis] |
| XP_018272653.1 | 612 | RHOBADRAFT_35252 | eukaryota>fungi>basidiomycota | Rhodotorula | graminis WP1 | hypothetical protein RHOBADRAFT_35252 [Rhodotorula graminis WP1] |
| NP_001080326.1 | 611 | canx,canx-a,canx-b,clgn,clnx,cnx | eukaryota>metazoa>chordata>vertebrata | Xenopus | laevis | calnexin L homeolog precursor [Xenopus laevis] |
| NP_034034.2 | 611 | 4930459O04Rik,A2/6,AI528775,Cln | eukaryota>metazoa>chordata>vertebrata | Mus | musculus | calmegin isoform 1 precursor [Mus musculus] |
| OMH82761.1 | 611 | AX774_g3762 | eukaryota>fungi | Zancudomyces | culisetae | Calnexin-like protein [Zancudomyces culisetae] |
| OMH85345.1 | 611 | AX774_g1100 | eukaryota>fungi | Zancudomyces | culisetae | Calnexin-like protein [Zancudomyces culisetae] |
| XP_006530698.1 | 611 | 4930459O04Rik,AI528775,Cln | eukaryota>metazoa>chordata>vertebrata | Mus | musculus | PREDICTED: calmegin isoform X2 [Mus musculus] |
| XP_011246600.1 | 611 | 4930459O04Rik,AI528775,Cln | eukaryota>metazoa>chordata>vertebrata | Mus | musculus | PREDICTED: calmegin isoform X2 [Mus musculus] |
| XP_011124147.1 | 610 | CLGN | eukaryota>metazoa>chordata>vertebrata | Homo | sapiens | calmegin precursor [Homo sapiens] |
| NP_004353.1 | 610 | CLGN | eukaryota>metazoa>chordata>vertebrata | Homo | sapiens | calmegin precursor [Homo sapiens] |
| XP_005556017.1 | 610 | CLGN | eukaryota>metazoa>chordata>vertebrata | Macaca | fascicularis | PREDICTED: calmegin [Macaca fascicularis] |
| XP_005556018.1 | 610 | CLGN | eukaryota>metazoa>chordata>vertebrata | Macaca | fascicularis | PREDICTED: calmegin [Macaca fascicularis] |
| XP_005556019.1 | 610 | CLGN | eukaryota>metazoa>chordata>vertebrata | Macaca | fascicularis | PREDICTED: calmegin [Macaca fascicularis] |
| NP_001187109.1 | 607 | canx | eukaryota>metazoa>chordata>vertebrata>actinopterygii | Ictalurus | punctatus | calnexin precursor [Ictalurus punctatus] |
| XP_018098778.1 | 607 | LOC108706675 | eukaryota>metazoa>chordata>vertebrata | Xenopus | laevis | PREDICTED: calnexin-like isoform X2 [Xenopus laevis] |
| NP_001036767.1 | 605 | CG11958,cnx,Cnx,cnx99A,Cnx99a,Dmel\CG11958 | eukaryota>metazoa>hexapoda | Drosophila | melanogaster | calnexin 99A, isoform D [Drosophila melanogaster] |
| NP_477157.1 | 605 | CG11958,cnx,Cnx,cnx99A,Cnx99a,Dmel\CG11958 | eukaryota>metazoa>hexapoda | Drosophila | melanogaster | calnexin 99A, isoform B [Drosophila melanogaster] |
| NP_733286.1 | 605 | CG11958,cnx,Cnx,cnx99A,Cnx99a,Dmel\CG11958 | eukaryota>metazoa>hexapoda | Drosophila | melanogaster | calnexin 99A, isoform A [Drosophila melanogaster] |
| NP_001036766.2 | 604 | CG11958,cnx,Cnx,cnx99A,Cnx99a,Dmel\CG11958 | eukaryota>metazoa>hexapoda | Drosophila | melanogaster | calnexin 99A, isoform E [Drosophila melanogaster] |
| XP_015773741.1 | 603 | LOC107351954 | eukaryota>metazoa>cnidaria | Acropora | digitifera | PREDICTED: calnexin-like isoform X1 [Acropora digitifera] |
| XP_006558922.1 | 602 | LOC552528 | eukaryota>metazoa>hexapoda | Apis | mellifera | PREDICTED: calnexin isoform X1 [Apis mellifera] |
| XP_016769249.1 | 602 | LOC552528 | eukaryota>metazoa>hexapoda | Apis | mellifera | PREDICTED: calnexin isoform X1 [Apis mellifera] |
| NP_001025791.1 | 601 | CANX | eukaryota>metazoa>chordata>vertebrata | Gallus | gallus | calnexin precursor [Gallus gallus] |
| XP_015149278.1 | 601 | CANX | eukaryota>metazoa>chordata>vertebrata | Gallus | gallus | calnexin isoform X1 [Gallus gallus] |
| XP_018098779.1 | 601 | LOC108706675 | eukaryota>metazoa>chordata>vertebrata | Xenopus | laevis | PREDICTED: calnexin-like isoform X3 [Xenopus laevis] |
| CEM22137.1 | 600 | Vbra_9634 | eukaryota>alveolata | Vitrella | brassicaformis CCMP3155 | unnamed protein product [Vitrella brassicaformis CCMP3155] |
| NP_998613.1 | 600 | wu:fe06b12,zgc:63524 | eukaryota>metazoa>chordata>vertebrata>actinopterygii | Danio | rerio | calnexin precursor [Danio rerio] |
| Smar1000009914 | 600 | Smar1000009914 | eukaryota>arthropoda>myriapoda | Strigamia | maritima | >Smar1000009914 |
| XP_009306088.1 | 600 | wu:fe06b12,zgc:63524 | eukaryota>metazoa>chordata>vertebrata>actinopterygii | Danio | rerio | calnexin isoform X1 [Danio rerio] |
| XP_011617847.1 | 600 | LOC101070859 | eukaryota>metazoa>chordata>vertebrata>actinopterygii | Takifugu | rubripes | PREDICTED: calnexin-like [Takifugu rubripes] |
| ERG83915.1 | 599 | ASU_06225 | eukaryota>metazoa>nematoda | Ascaris | suum | calnexin-like protein [Ascaris suum] |
| XP_003885042.1 | 599 | NCLIV_054410 | eukaryota>alveolata>apicomplexa | Neospora | caninum Liverpool | Calr protein, related [Neospora caninum Liverpool] |
| XP_014567205.1 | 597 | L969DRAFT_24275 | eukaryota>fungi>basidiomycota | Mixia | osmundae IAM 14324 | hypothetical protein L969DRAFT_24275 [Mixia osmundae IAM 14324] |
| XP_014669601.1 | 595 | LOC106810686 | eukaryota>metazoa | Priapulus | caudatus | PREDICTED: calnexin-like [Priapulus caudatus] |
| XP_014669602.1 | 595 | LOC106810686 | eukaryota>metazoa | Priapulus | caudatus | PREDICTED: calnexin-like [Priapulus caudatus] |
| XP_014669603.1 | 595 | LOC106810686 | eukaryota>metazoa | Priapulus | caudatus | PREDICTED: calnexin-like [Priapulus caudatus] |
| XP_014669604.1 | 595 | LOC106810686 | eukaryota>metazoa | Priapulus | caudatus | PREDICTED: calnexin-like [Priapulus caudatus] |
| XP_009177069.1 | 594 | T265_15627 | eukaryota>metazoa | Opisthorchis | viverrini | hypothetical protein T265_15627 [Opisthorchis viverrini] |
| CBJ28991.1 | 593 | Esi_0130_0015 | eukaryota>stramenopiles | Ectocarpus | siliculosus | Calnexin [Ectocarpus siliculosus] |
| XP_003258188.2 | 593 | CLGN | eukaryota>metazoa>chordata>vertebrata | Nomascus | leucogenys | PREDICTED: calmegin [Nomascus leucogenys] |
| XP_015773742.1 | 593 | LOC107351954 | eukaryota>metazoa>cnidaria | Acropora | digitifera | PREDICTED: calnexin-like isoform X2 [Acropora digitifera] |
| XP_018087187.1 | 593 | LOC108699598 | eukaryota>metazoa>chordata>vertebrata | Xenopus | laevis | PREDICTED: calmegin-like isoform X2 [Xenopus laevis] |
| NP_001019820.1 | 592 | CNX,IP90,P90 | eukaryota>metazoa>chordata>vertebrata | Homo | sapiens | calnexin isoform d precursor [Homo sapiens] |
| NP_001737.1 | 592 | CNX,IP90,P90 | eukaryota>metazoa>chordata>vertebrata | Homo | sapiens | calnexin isoform d precursor [Homo sapiens] |
| XP_003279601.1 | 592 | CANX | eukaryota>metazoa>chordata>vertebrata | Nomascus | leucogenys | PREDICTED: calnexin isoform X1 [Nomascus leucogenys] |
| XP_003279602.1 | 592 | CANX | eukaryota>metazoa>chordata>vertebrata | Nomascus | leucogenys | PREDICTED: calnexin isoform X1 [Nomascus leucogenys] |
| XP_005558828.1 | 592 | CANX | eukaryota>metazoa>chordata>vertebrata | Macaca | fascicularis | PREDICTED: calnexin [Macaca fascicularis] |
| XP_005558830.1 | 592 | CANX | eukaryota>metazoa>chordata>vertebrata | Macaca | fascicularis | PREDICTED: calnexin [Macaca fascicularis] |
| XP_011532967.1 | 592 | CNX,IP90,P90 | eukaryota>metazoa>chordata>vertebrata | Homo | sapiens | calnexin isoform X2 [Homo sapiens] |
| XP_012360185.1 | 592 | CANX | eukaryota>metazoa>chordata>vertebrata | Nomascus | leucogenys | PREDICTED: calnexin isoform X1 [Nomascus leucogenys] |
| XP_015307854.1 | 592 | CANX | eukaryota>metazoa>chordata>vertebrata | Macaca | fascicularis | PREDICTED: calnexin [Macaca fascicularis] |
| XP_015307855.1 | 592 | CANX | eukaryota>metazoa>chordata>vertebrata | Macaca | fascicularis | PREDICTED: calnexin [Macaca fascicularis] |
| XP_015307856.1 | 592 | CANX | eukaryota>metazoa>chordata>vertebrata | Macaca | fascicularis | PREDICTED: calnexin [Macaca fascicularis] |
| XP_019022135.1 | 592 | SAICODRAFT_9347 | eukaryota>fungi>ascomycota | Saitoella | complicata NRRL Y-17804 | calmegin [Saitoella complicata NRRL Y-17804] |
| NP_001103969.1 | 591 | 1110069N15Rik,AI988026,Cnx,D11Ertd153e | eukaryota>metazoa>chordata>vertebrata | Mus | musculus | calnexin precursor [Mus musculus] |
| NP_001103970.1 | 591 | 1110069N15Rik,AI988026,Cnx,D11Ertd153e | eukaryota>metazoa>chordata>vertebrata | Mus | musculus | calnexin precursor [Mus musculus] |
| NP_031623.1 | 591 | 1110069N15Rik,AI988026,Cnx,D11Ertd153e | eukaryota>metazoa>chordata>vertebrata | Mus | musculus | calnexin precursor [Mus musculus] |
| XP_791226.2 | 591 | LOC586348 | eukaryota>metazoa>echinodermata | Strongylocentrotus | purpuratus | PREDICTED: calnexin [Strongylocentrotus purpuratus] |
| XP_013389410.1 | 590 | LOC106158092 | eukaryota>metazoa | Lingula | anatina | calnexin isoform X1 [Lingula anatina] |
| XP_013414188.1 | 590 | LOC106176375 | eukaryota>metazoa | Lingula | anatina | calnexin isoform X1 [Lingula anatina] |

[illegible]

| | | | | | | |
|---|---|---|---|---|---|---|
| XP_014771603.1 | 679 | LOC106870127 | eukaryota>metazoa>mollusca | Octopus | bimaculoides | PREDICTED: putative uncharacterized protein DDB_G0282499, partial [Octopus bimaculoides] |
| XP_019700160.1 | 678 | LOC105190090 | eukaryota>metazoa>hexapoda | Harpegnathos | saltator | PREDICTED: protein enabled [Harpegnathos saltator] |
| XP_012552943.1 | 672 | LOC101735452 | eukaryota>metazoa>hexapoda | Bombyx | mori | protein enabled isoform X1 [Bombyx mori] |
| XP_021209110.1 | 672 | LOC101735452 | eukaryota>metazoa>hexapoda | Bombyx | mori | protein enabled isoform X1 [Bombyx mori] |
| XP_021209111.1 | 672 | LOC101735452 | eukaryota>metazoa>hexapoda | Bombyx | mori | protein enabled isoform X1 [Bombyx mori] |
| XP_021209112.1 | 671 | LOC101735452 | eukaryota>metazoa>hexapoda | Bombyx | mori | protein enabled isoform X2 [Bombyx mori] |
| XP_021209113.1 | 671 | LOC101735452 | eukaryota>metazoa>hexapoda | Bombyx | mori | protein enabled isoform X3 [Bombyx mori] |
| XP_021209114.1 | 670 | LOC101735452 | eukaryota>metazoa>hexapoda | Bombyx | mori | protein enabled isoform X4 [Bombyx mori] |
| XP_014533405.1 | 666 | PDIP_61790 | eukaryota>fungi>ascomycota | Penicillium | digitatum Pd1 | Proline-rich protein LAS17 [Penicillium digitatum Pd1] |
| NP_001286602.1 | 664 | CG15112,Dmel\CG15112,Ena,ENA/VASP,enb,Enb,l(2)02029,Vasp,VASP | eukaryota>metazoa>hexapoda | Drosophila | melanogaster | enabled, isoform G [Drosophila melanogaster] |
| XP_005158849.1 | 663 | Evl,id:ibd2260,zgc:65883,zgc:77388 | eukaryota>metazoa>chordata>vertebrata>actinopterygii | Danio | rerio | ena/VASP-like protein isoform X1 [Danio rerio] |
| ERG80807.1 | 663 | ASU_11206 | eukaryota>metazoa>nematoda | Ascaris | suum | sprouty- evh1 domain-containing protein 2 [Ascaris suum] |
| XP_021322898.1 | 662 | Evl,id:ibd2260,zgc:65883,zgc:77388 | eukaryota>metazoa>chordata>vertebrata>actinopterygii | Danio | rerio | ena/VASP-like protein isoform X2 [Danio rerio] |
| XP_005158850.1 | 653 | Evl,id:ibd2260,zgc:65883,zgc:77388 | eukaryota>metazoa>chordata>vertebrata>actinopterygii | Danio | rerio | ena/VASP-like protein isoform X3 [Danio rerio] |
| XP_452412.1 | 653 | KLLA0_C04774g | eukaryota>fungi>ascomycota | Kluyveromyces | lactis | uncharacterized protein KLLA0_C04774g [Kluyveromyces lactis] |
| XP_012749022.1 | 651 | SAMD00019534_112390 | eukaryota>amoebozoa>mycetozoa>dictyosteliida | Acytostelium | subglobosum LB1 | hypothetical protein SAMD00019534_112390 [Acytostelium subglobosum LB1] |
| XP_015306083.1 | 650 | ENAH | eukaryota>metazoa>chordata>vertebrata | Macaca | fascicularis | PREDICTED: protein enabled homolog isoform X11 [Macaca fascicularis] |
| EFA85844.1 | 650 | wasA | eukaryota>amoebozoa>mycetozoa | Heterostelium | album PN500 | Wiscott-Aldrich syndrome protein [Heterostelium album PN500] |
| XP_018655478.1 | 650 | Smp_039870.1 | eukaryota>metazoa | Schistosoma | mansoni | putative vasodilator-stimulated phosphoprotein [Schistosoma mansoni] |
| XP_019019119.1 | 644 | PICMEDRAFT_15858 | eukaryota>fungi>ascomycota | Pichia | membranifaciens NRRL Y-2026 | hypothetical protein PICMEDRAFT_15858 [Pichia membranifaciens NRRL Y-2026] |
| XP_005158851.1 | 641 | Evl,id:ibd2260,zgc:65883,zgc:77388 | eukaryota>metazoa>chordata>vertebrata>actinopterygii | Danio | rerio | ena/VASP-like protein isoform X4 [Danio rerio] |
| EPZ34321.1 | 641 | O9G_000508 | eukaryota>fungi | Rozella | allomycis CSF55 | Chitin synthase III catalytic subunit domain-containing protein [Rozella allomycis CSF55] |
| XP_005158852.1 | 638 | Evl,id:ibd2260,zgc:65883,zgc:77388 | eukaryota>metazoa>chordata>vertebrata>actinopterygii | Danio | rerio | ena/VASP-like protein isoform X5 [Danio rerio] |
| XP_964625.1 | 636 | NCU07438 | eukaryota>fungi>ascomycota | Neurospora | crassa OR74A | WASP-like protein las17p [Neurospora crassa OR74A] |
| XP_017323611.1 | 635 | LOC108265595 | eukaryota>metazoa>chordata>vertebrata>actinopterygii | Ictalurus | punctatus | PREDICTED: wiskott-Aldrich syndrome protein-like isoform X1 [Ictalurus punctatus] |
| NP_014824.1 | 633 | BEE1 | eukaryota>fungi>ascomycota | Saccharomyces | cerevisiae S288C | actin-binding protein LAS17 [Saccharomyces cerevisiae S288C] |
| XP_005541041.1 | 631 | ENAH | eukaryota>metazoa>chordata>vertebrata | Macaca | fascicularis | PREDICTED: protein enabled homolog isoform X12 [Macaca fascicularis] |
| XP_699919.5 | 630 | wasa | eukaryota>metazoa>chordata>vertebrata>actinopterygii | Danio | rerio | wiskott-Aldrich syndrome protein [Danio rerio] |
| XP_015306085.1 | 629 | ENAH | eukaryota>metazoa>chordata>vertebrata | Macaca | fascicularis | PREDICTED: protein enabled homolog isoform X13 [Macaca fascicularis] |
| XP_501207.2 | 629 | YALI0_B22110g | eukaryota>fungi>ascomycota | Yarrowia | lipolytica CLIB122 | YALI0B22110p [Yarrowia lipolytica CLIB122] |
| XP_016857242.1 | 628 | ENA,MENA,NDPP1 | eukaryota>metazoa>chordata>vertebrata | Homo | sapiens | PREDICTED: protein enabled homolog isoform X9 [Homo sapiens] |
| ORY25113.1 | 628 | LY90DRAFT_674938 | eukaryota>fungi>chytridiomycota | Neocallimastix | californiae | hypothetical protein LY90DRAFT_674938 [Neocallimastix californiae] |
| XP_018186371.1 | 620 | L228DRAFT_249631 | eukaryota>fungi>ascomycota | Xylona | heveae TC161 | WH1-domain-containing protein [Xylona heveae TC161] |
| XP_002841323.1 | 618 | GSTUM_00010426001 | eukaryota>fungi>ascomycota | Tuber | melanosporum Mel28 | hypothetical protein [Tuber melanosporum Mel28] |
| XP_021324089.1 | 616 | id:ibd2798,si:ch211-215p14.1,zgc:103638 | eukaryota>metazoa>chordata>vertebrata>actinopterygii | Danio | rerio | protein enabled homolog isoform X5 [Danio rerio] |
| XP_016857243.1 | 614 | ENA,MENA,NDPP1 | eukaryota>metazoa>chordata>vertebrata | Homo | sapiens | PREDICTED: protein enabled homolog isoform X10 [Homo sapiens] |
| XP_002111223.1 | 611 | TRIADDRAFT_54962 | eukaryota>metazoa>placozoa | Trichoplax | adhaerens | hypothetical protein TRIADDRAFT_54962 [Trichoplax adhaerens] |
| XP_005541042.1 | 610 | ENAH | eukaryota>metazoa>chordata>vertebrata | Macaca | fascicularis | PREDICTED: protein enabled homolog isoform X14 [Macaca fascicularis] |
| XP_017332339.1 | 610 | LOC108270303 | eukaryota>metazoa>chordata>vertebrata>actinopterygii | Ictalurus | punctatus | PREDICTED: ena/VASP-like protein isoform X1 [Ictalurus punctatus] |
| XP_011125450.1 | 610 | AOL_s00112g118 | eukaryota>fungi>ascomycota | Arthrobotrys | oligospora ATCC 24927 | hypothetical protein AOL_s00112g118 [Arthrobotrys oligospora ATCC 24927] |
| XP_017332340.1 | 609 | LOC108270303 | eukaryota>metazoa>chordata>vertebrata>actinopterygii | Ictalurus | punctatus | PREDICTED: ena/VASP-like protein isoform X2 [Ictalurus punctatus] |
| XP_017332341.1 | 609 | LOC108270303 | eukaryota>metazoa>chordata>vertebrata>actinopterygii | Ictalurus | punctatus | PREDICTED: ena/VASP-like protein isoform X3 [Ictalurus punctatus] |
| EUB60407.1 | 607 | EGR_04789 | eukaryota>metazoa | Echinococcus | granulosus | Protein enabled [Echinococcus granulosus] |
| XP_005273244.1 | 605 | ENA,MENA,NDPP1 | eukaryota>metazoa>chordata>vertebrata | Homo | sapiens | PREDICTED: protein enabled homolog isoform X11 [Homo sapiens] |
| XP_018119925.1 | 602 | Xena | eukaryota>metazoa>chordata>vertebrata | Xenopus | laevis | PREDICTED: enabled protein isoform X2 [Xenopus laevis] |
| XP_016857244.1 | 601 | ENA,MENA,NDPP1 | eukaryota>metazoa>chordata>vertebrata | Homo | sapiens | PREDICTED: protein enabled homolog isoform X12 [Homo sapiens] |
| XP_015139086.1 | 598 | ENAH | eukaryota>metazoa>chordata>vertebrata | Gallus | gallus | PREDICTED: protein enabled homolog isoform X11 [Gallus gallus] |
| XP_021324090.1 | 596 | id:ibd2798,si:ch211-215p14.1,zgc:103638 | eukaryota>metazoa>chordata>vertebrata>actinopterygii | Danio | rerio | protein enabled homolog isoform X6 [Danio rerio] |
| XP_015139087.1 | 594 | ENAH | eukaryota>metazoa>chordata>vertebrata | Gallus | gallus | protein enabled homolog isoform X8 [Gallus gallus] |
| XP_006496713.1 | 593 | Mena,NDPP-1,Ndpp1,WBP8 | eukaryota>metazoa>chordata>vertebrata | Mus | musculus | PREDICTED: protein enabled homolog isoform X8 [Mus musculus] |
| XP_001222599.1 | 592 | CHGG_06504 | eukaryota>fungi>ascomycota | Chaetomium | globosum CBS 148.51 | hypothetical protein CHGG_06504 [Chaetomium globosum CBS 148.51] |
| NP_001008493.1 | 591 | ENA,MENA,NDPP1 | eukaryota>metazoa>chordata>vertebrata | Homo | sapiens | protein enabled homolog isoform a [Homo sapiens] |
| XP_016885275.1 | 590 | IMD2,SCNX,THC,THC1,WASP,WASPA | eukaryota>metazoa>chordata>vertebrata | Homo | sapiens | wiskott-Aldrich syndrome protein isoform X2 [Homo sapiens] |
| XP_017323612.1 | 590 | LOC108265595 | eukaryota>metazoa>chordata>vertebrata>actinopterygii | Ictalurus | punctatus | PREDICTED: wiskott-Aldrich syndrome protein-like isoform X2 [Ictalurus punctatus] |
| XP_015306089.1 | 589 | ENAH | eukaryota>metazoa>chordata>vertebrata | Macaca | fascicularis | PREDICTED: protein enabled homolog isoform X15 [Macaca fascicularis] |
| XP_015306090.1 | 589 | ENAH | eukaryota>metazoa>chordata>vertebrata | Macaca | fascicularis | PREDICTED: protein enabled homolog isoform X15 [Macaca fascicularis] |
| XP_009013906.1 | 586 | HELRODRAFT_98248 | eukaryota>metazoa>annelida | Helobdella | robusta | hypothetical protein HELRODRAFT_98248 [Helobdella robusta] |
| XP_011419448.1 | 586 | LOC105322431 | eukaryota>metazoa>mollusca | Crassostrea | gigas | PREDICTED: homer protein homolog 2 isoform X1 [Crassostrea gigas] |
| CDS37344.1 | 586 | EmuJ_000458800 | eukaryota>metazoa | Echinococcus | multilocularis | vasodilator stimulated phosphoprotein [Echinococcus multilocularis] |
| XP_018655479.1 | 585 | Smp_039870.1 | eukaryota>metazoa | Schistosoma | mansoni | LOW QUALITY PROTEIN: putative vasodilator-stimulated phosphoprotein [Schistosoma mansoni] |
| XP_002668568.1 | 582 | NAEGRDRAFT_82342 | eukaryota>heterolobosea | Naegleria | gruberi strain NEG-M | predicted protein [Naegleria gruberi strain NEG-M] |
| XP_009292990.1 | 581 | id:ibd2798,si:ch211-215p14.1,zgc:103638 | eukaryota>metazoa>chordata>vertebrata>actinopterygii | Danio | rerio | protein enabled homolog isoform X7 [Danio rerio] |
| XP_011419449.1 | 581 | LOC105322431 | eukaryota>metazoa>mollusca | Crassostrea | gigas | PREDICTED: homer protein homolog 2 isoform X2 [Crassostrea gigas] |
| XP_003275128.1 | 579 | ENAH | eukaryota>metazoa>chordata>vertebrata | Nomascus | leucogenys | PREDICTED: protein enabled homolog [Nomascus leucogenys] |
| XP_016857245.1 | 578 | ENA,MENA,NDPP1 | eukaryota>metazoa>chordata>vertebrata | Homo | sapiens | PREDICTED: protein enabled homolog isoform X13 [Homo sapiens] |
| XP_016857246.1 | 578 | ENA,MENA,NDPP1 | eukaryota>metazoa>chordata>vertebrata | Homo | sapiens | PREDICTED: protein enabled homolog isoform X13 [Homo sapiens] |
| EFX90289.1 | 578 | DAPPUDRAFT_300115 | eukaryota>metazoa>crustacea | Daphnia | pulex | hypothetical protein DAPPUDRAFT_300115 [Daphnia pulex] |
| XP_013427246.1 | 577 | M436DRAFT_73051 | eukaryota>fungi>ascomycota | Aureobasidium | namibiae CBS 147.97 | WH1-domain-containing protein [Aureobasidium namibiae CBS 147.97] |
| XP_001897341.1 | 575 | Bm1_29495 | eukaryota>metazoa>nematoda | Brugia | malayi | WH1 domain containing protein [Brugia malayi] |
| XP_001631290.1 | 575 | NEMVEDRAFT_v1g244007 | eukaryota>metazoa>cnidaria | Nematostella | vectensis | predicted protein [Nematostella vectensis] |
| XP_015139088.1 | 574 | ENAH | eukaryota>metazoa>chordata>vertebrata | Gallus | gallus | protein enabled homolog isoform X9 [Gallus gallus] |
| NP_594758.1 | 574 | wsp1 | eukaryota>fungi>ascomycota | Schizosaccharomyces | pombe | actin assembly factor [Schizosaccharomyces pombe] |
| XP_006496714.1 | 572 | Mena,NDPP-1,Ndpp1,WBP8 | eukaryota>metazoa>chordata>vertebrata | Mus | musculus | PREDICTED: protein enabled homolog isoform X9 [Mus musculus] |
| XP_018119926.1 | 572 | Xena | eukaryota>metazoa>chordata>vertebrata | Xenopus | laevis | PREDICTED: enabled protein isoform X3 [Xenopus laevis] |
| NP_060682.2 | 570 | ENA,MENA,NDPP1 | eukaryota>metazoa>chordata>vertebrata | Homo | sapiens | protein enabled homolog isoform b [Homo sapiens] |
| XP_017311958.1 | 569 | enah | eukaryota>metazoa>chordata>vertebrata>actinopterygii | Ictalurus | punctatus | PREDICTED: protein enabled homolog isoform X6 [Ictalurus punctatus] |
| CEO99970.1 | 569 | PBRA_007704 | eukaryota>rhizaria>cercozoa>plasmodiophorida | Plasmodiophora | brassicae | hypothetical protein PBRA_007704 [Plasmodiophora brassicae] |
| CAG10529.1 | 567 | GSTEN:00032207:G:001 | eukaryota>metazoa>chordata>vertebrata>actinopterygii | Tetraodon | nigroviridis | unnamed protein product, partial [Tetraodon nigroviridis] |
| XP_015306091.1 | 566 | ENAH | eukaryota>metazoa>chordata>vertebrata | Macaca | fascicularis | PREDICTED: protein enabled homolog isoform X16 [Macaca fascicularis] |
| XP_014662305.1 | 566 | LOC106805282 | eukaryota>metazoa | Priapulus | caudatus | PREDICTED: neural Wiskott-Aldrich syndrome protein-like [Priapulus caudatus] |
| XP_019027289.1 | 565 | SAICODRAFT_4370 | eukaryota>fungi>ascomycota | Saitoella | complicata NRRL Y-17804 | WH1-domain-containing protein [Saitoella complicata NRRL Y-17804] |
| NP_001085660.1 | 562 | Ena,enah,Xena | eukaryota>metazoa>chordata>vertebrata | Xenopus | laevis | ENAH, actin regulator L homeolog [Xenopus laevis] |
| XP_002609962.1 | 561 | BRAFLDRAFT_85919 | eukaryota>metazoa>chordata | Branchiostoma | floridae | hypothetical protein BRAFLDRAFT_85919 [Branchiostoma floridae] |
| XP_015139090.1 | 561 | ENAH | eukaryota>metazoa>chordata>vertebrata | Gallus | gallus | PREDICTED: protein enabled homolog isoform X14 [Gallus gallus] |
| XP_009292991.1 | 561 | id:ibd2798,si:ch211-215p14.1,zgc:103638 | eukaryota>metazoa>chordata>vertebrata>actinopterygii | Danio | rerio | protein enabled homolog isoform X8 [Danio rerio] |
| OQV12129.1 | 559 | BV898_13608 | eukaryota>metazoa | Hypsibius | dujardini | putative Protein enabled-like protein [Hypsibius dujardini] |
| XP_003380270.1 | 557 | Tsp_06005 | eukaryota>metazoa>nematoda | Trichinella | spiralis | putative neural Wiskott-Aldrich syndrome protein [Trichinella spiralis] |
| ORZ38628.1 | 553 | BCR44DRAFT_1428628 | eukaryota>fungi | Catenaria | anguillulae PL171 | hypothetical protein BCR44DRAFT_1428628, partial [Catenaria anguillulae PL171] |
| XP_011419450.1 | 551 | LOC105322431 | eukaryota>metazoa>mollusca | Crassostrea | gigas | PREDICTED: homer protein homolog 2 isoform X3 [Crassostrea gigas] |
| XP_015139091.1 | 550 | ENAH | eukaryota>metazoa>chordata>vertebrata | Gallus | gallus | protein enabled homolog isoform X11 [Gallus gallus] |
| XP_001835070.1 | 550 | CC1G_06473 | eukaryota>fungi>basidiomycota | Coprinopsis | cinerea okayama7#130 | hypothetical protein CC1G_06473 [Coprinopsis cinerea okayama7#130] |
| OWA55093.1 | 550 | BV898_19478 | eukaryota>metazoa | Hypsibius | dujardini | putative Sprouty-related, EVH1 domain-containing protein 1 [Hypsibius dujardini] |
| XP_017311959.1 | 549 | enah | eukaryota>metazoa>chordata>vertebrata>actinopterygii | Ictalurus | punctatus | PREDICTED: protein enabled homolog isoform X7 [Ictalurus punctatus] |
| KFH74147.1 | 549 | MVEG_01360 | eukaryota>fungi | Mortierella | verticillata NRRL 6337 | hypothetical protein MVEG_01360 [Mortierella verticillata NRRL 6337] |
| ORY53338.1 | 548 | LY90DRAFT_285267 | eukaryota>fungi>chytridiomycota | Neocallimastix | californiae | hypothetical protein LY90DRAFT_285267 [Neocallimastix californiae] |
| CBY09705.1 | 545 | GSOID_T00010515001 | eukaryota>metazoa>chordata | Oikopleura | dioica | unnamed protein product [Oikopleura dioica] |
| XP_011601021.1 | 545 | was | eukaryota>metazoa>chordata>vertebrata>actinopterygii | Takifugu | rubripes | PREDICTED: wiskott-Aldrich syndrome protein [Takifugu rubripes] |
| CBY22289.1 | 544 | GSOID_T00018446001 | eukaryota>metazoa>chordata | Oikopleura | dioica | unnamed protein product [Oikopleura dioica] |
| CBY31400.1 | 544 | GSOID_T00025303001 | eukaryota>metazoa>chordata | Oikopleura | dioica | unnamed protein product [Oikopleura dioica] |
| NP_001076590.1 | 541 | Mena,NDPP-1,Ndpp1,WBP8 | eukaryota>metazoa>chordata>vertebrata | Mus | musculus | protein enabled homolog 4 [Mus musculus] |
| XP_021876053.1 | 541 | BCR41DRAFT_426279 | eukaryota>fungi | Lobosporangium | transversale | hypothetical protein BCR41DRAFT_426279 [Lobosporangium transversale] |
| XP_021323362.1 | 540 | si:ch211-255112.2 | eukaryota>metazoa>chordata>vertebrata>actinopterygii | Danio | rerio | sprouty-related, EVH1 domain containing 3 isoform X1 [Danio rerio] |
| XP_006569628.1 | 540 | LOC409218 | eukaryota>metazoa>hexapoda | Apis | mellifera | PREDICTED: neural Wiskott-Aldrich syndrome protein [Apis mellifera] |
| XP_392742.4 | 540 | LOC409218 | eukaryota>metazoa>hexapoda | Apis | mellifera | PREDICTED: neural Wiskott-Aldrich syndrome protein [Apis mellifera] |
| XP_006957313.1 | 538 | WALSEDRAFT_56819 | eukaryota>fungi>basidiomycota | Wallemia | mellicola CBS 633.66 | hypothetical protein WALSEDRAFT_56819 [Wallemia mellicola CBS 633.66] |
| KXJ29816.1 | 538 | WAS | eukaryota>metazoa>cnidaria | Exaiptasia | pallida | Wiskott-Aldrich syndrome protein [Exaiptasia pallida] |
| XP_015778704.1 | 538 | LOC107356613 | eukaryota>metazoa>cnidaria | Acropora | digitifera | PREDICTED: neural Wiskott-Aldrich syndrome protein-like [Acropora digitifera] |
| XP_015763982.1 | 538 | LOC107342971 | eukaryota>metazoa>cnidaria | Acropora | digitifera | PREDICTED: wiskott-Aldrich syndrome protein homolog [Acropora digitifera] |
| XP_015139092.1 | 537 | ENAH | eukaryota>metazoa>chordata>vertebrata | Gallus | gallus | protein enabled homolog isoform X12 [Gallus gallus] |
| XP_002610296.1 | 536 | BRAFLDRAFT_126860 | eukaryota>metazoa>chordata | Branchiostoma | floridae | hypothetical protein BRAFLDRAFT_126860 [Branchiostoma floridae] |
| XP_006496715.1 | 535 | Mena,NDPP-1,Ndpp1,WBP8 | eukaryota>metazoa>chordata>vertebrata | Mus | musculus | PREDICTED: protein enabled homolog isoform X10 [Mus musculus] |
| NP_001086441.1 | 535 | ena,Xena | eukaryota>metazoa>chordata>vertebrata | Xenopus | laevis | ENAH, actin regulator S homeolog [Xenopus laevis] |
| XP_002127752.1 | 535 | LOC100183769 | eukaryota>metazoa>chordata | Ciona | intestinalis | PREDICTED: sprouty-related, EVH1 domain-containing protein 2-like [Ciona intestinalis] |
| XP_001635928.1 | 534 | NEMVEDRAFT_v1g241376 | eukaryota>metazoa>cnidaria | Nematostella | vectensis | predicted protein [Nematostella vectensis] |
| NP_001013539.2 | 533 | id:ibd2798,si:ch211-215p14.1,zgc:103638 | eukaryota>metazoa>chordata>vertebrata>actinopterygii | Danio | rerio | protein enabled homolog 1 [Danio rerio] |
| NP_001036341.2 | 531 | BEST:LD03829,CG11324,dhom,Dhom,Dhomer,Dm,Homer,Dmel\CG11324,Dvh,Hom,Homer,LD03829 | eukaryota>metazoa>hexapoda | Drosophila | melanogaster | homer, isoform E [Drosophila melanogaster] |
| KXN66161.1 | 529 | CONCODRAFT_80577 | eukaryota>fungi | Conidiobolus | coronatus NRRL 28638 | hypothetical protein CONCODRAFT_80577 [Conidiobolus coronatus NRRL 28638] |
| OQV18372.1 | 528 | BV898_07575 | eukaryota>metazoa | Hypsibius | dujardini | putative Wiskott-Aldrich syndrome protein [Hypsibius dujardini] |
| XP_001868297.1 | 528 | CpipJ_CPIJ018200 | eukaryota>metazoa>hexapoda | Culex | quinquefasciatus | conserved hypothetical protein [Culex quinquefasciatus] |
| XP_007331884.1 | 527 | AGABI1DRAFT_101672 | eukaryota>fungi>basidiomycota | Agaricus | bisporus var. burnettii JB137-S8 | hypothetical protein AGABI1DRAFT_101672 [Agaricus bisporus var. burnettii JB137-S8] |
| XP_014149294.1 | 527 | SARC_12081 | eukaryota>ichthyosporea | Sphaeroforma | arctica JP610 | hypothetical protein SARC_12081, partial [Sphaeroforma arctica JP610] |

[illegible]

| | | | | | | |
|---|---|---|---|---|---|---|
| CDS40850.1 | 331 | EmuJ_000845000 | eukaryota>metazoa | Echinococcus | multilocularis | wiskott Aldrich syndrome protein [Echinococcus multilocularis] |
| XP_011520536.1 | 330 | ACPD,CPD,DFNA68,HOMER-2,VESL-2 | eukaryota>metazoa>chordata>vertebrata | Homo | sapiens | homer protein homolog 2 isoform X6 [Homo sapiens] |
| NP_001139196.1 | 325 | HOMER-3,VESL3 | eukaryota>metazoa>chordata>vertebrata | Homo | sapiens | homer protein homolog 3 isoform 3 [Homo sapiens] |
| XP_015295838.1 | 325 | HOMER3 | eukaryota>metazoa>chordata>vertebrata | Macaca | fascicularis | PREDICTED: homer protein homolog 3 isoform X2 [Macaca fascicularis] |
| NP_001300273.1 | 324 | unc-34 | eukaryota>metazoa>nematoda | Caenorhabditis | elegans | Uncharacterized protein CELE_Y50D4C.1 [Caenorhabditis elegans] |
| XP_014156175.1 | 324 | SARC_05445 | eukaryota>ichthyosporea | Sphaeroforma | arctica JP610 | hypothetical protein SARC_05445 [Sphaeroforma arctica JP610] |
| XP_012353371.1 | 320 | HOMER2 | eukaryota>metazoa>chordata>vertebrata | Nomascus | leucogenys | PREDICTED: LOW QUALITY PROTEIN: homer protein homolog 2 [Nomascus leucogenys] |
| CBY32835.1 | 318 | GSOID_T00032191001 | eukaryota>metazoa>chordata | Oikopleura | dioica | unnamed protein product [Oikopleura dioica] |
| XP_001850671.1 | 317 | CpipJ_CPIJ009183 | eukaryota>metazoa>hexapoda | Culex | quinquefasciatus | homer [Culex quinquefasciatus] |
| NP_001018470.1 | 314 | zgc:111960,zmp:0000000677 | eukaryota>metazoa>chordata>vertebrata>actinopterygii | Danio | rerio | homer protein homolog 2 [Danio rerio] |
| CAG01507.1 | 312 | GSTEN:00020297:G:001 | eukaryota>metazoa>chordata>vertebrata>actinopterygii | Tetraodon | nigroviridis | unnamed protein product, partial [Tetraodon nigroviridis] |
| NP_001080709.1 | 310 | homer2 | eukaryota>metazoa>chordata>vertebrata | Xenopus | laevis | homer 2 [Xenopus laevis] |
| NP_001024254.1 | 310 | unc-34 | eukaryota>metazoa>nematoda | Caenorhabditis | elegans | Uncharacterized protein CELE_Y50D4C.1 [Caenorhabditis elegans] |
| XP_013382185.1 | 306 | LOC106152976 | eukaryota>metazoa | Lingula | anatina | homer protein homolog 2-like [Lingula anatina] |
| XP_001749347.1 | 304 | MONBRDRAFT_28813 | eukaryota>choanoflagellida | Monosiga | brevicollis MX1 | hypothetical protein [Monosiga brevicollis MX1] |
| CBY23156.1 | 301 | GSOID_T00015088001 | eukaryota>metazoa>chordata | Oikopleura | dioica | unnamed protein product [Oikopleura dioica] |
| XP_001919116.3 | 299 | si:dkey-197j19.6 | eukaryota>metazoa>chordata>vertebrata>actinopterygii | Danio | rerio | proline-rich protein LAS17-like [Danio rerio] |
| XP_017335405.1 | 296 | LOC108271935 | eukaryota>metazoa>chordata>vertebrata>actinopterygii | Ictalurus | punctatus | PREDICTED: wiskott-Aldrich syndrome protein-like isoform X1 [Ictalurus punctatus] |
| XP_013759887.1 | 293 | AMSG_03539 | eukaryota>apusozoa>apusomonadidae | Thecamonas | trahens ATCC 50062 | hypothetical protein AMSG_03539 [Thecamonas trahens ATCC 50062] |
| NP_001157559.1 | 290 | 9330120H11Rik,AW539445,CPD,Vesl-2 | eukaryota>metazoa>chordata>vertebrata | Mus | musculus | homer protein homolog 2 isoform 3 [Mus musculus] |
| XP_006540972.1 | 290 | 9330120H11Rik,AW539445,CPD,Vesl-2 | eukaryota>metazoa>chordata>vertebrata | Mus | musculus | PREDICTED: homer protein homolog 2 isoform X1 [Mus musculus] |
| XP_017177403.1 | 290 | AW047143,D130060H24Rik,Spred-3 | eukaryota>metazoa>chordata>vertebrata | Mus | musculus | PREDICTED: sprouty-related, EVH1 domain-containing protein 3 isoform X5 [Mus musculus] |
| XP_021329015.1 | 279 | vaspb | eukaryota>metazoa>chordata>vertebrata>actinopterygii | Danio | rerio | protein enabled homolog isoform X3 [Danio rerio] |
| XP_001848257.1 | 279 | CpipJ_CPIJ006699 | eukaryota>metazoa>hexapoda | Culex | quinquefasciatus | wiskott-aldrich syndrome protein [Culex quinquefasciatus] |
| XP_022238305.1 | 277 | LOC106478900 | eukaryota>metazoa | Limulus | polyphemus | neural Wiskott-Aldrich syndrome protein-like isoform X5 [Limulus polyphemus] |
| XP_001846730.1 | 271 | CpipJ_CPIJ004706 | eukaryota>metazoa>hexapoda | Culex | quinquefasciatus | vasodilator-stimulated phosphoprotein [Culex quinquefasciatus] |
| XP_022250318.1 | 267 | LOC106466578 | eukaryota>metazoa | Limulus | polyphemus | homer protein homolog 2-like, partial [Limulus polyphemus] |
| XP_009497368.1 | 266 | H696_05232 | eukaryota | Fonticula | alba | hypothetical protein H696_05232 [Fonticula alba] |
| XP_015770466.1 | 262 | LOC107348895 | eukaryota>metazoa>cnidaria | Acropora | digitifera | PREDICTED: uncharacterized protein LOC107348895 isoform X1 [Acropora digitifera] |
| XP_017177404.1 | 261 | AW047143,D130060H24Rik,Spred-3 | eukaryota>metazoa>chordata>vertebrata | Mus | musculus | PREDICTED: sprouty-related, EVH1 domain-containing protein 3 isoform X6 [Mus musculus] |
| XP_017177405.1 | 261 | AW047143,D130060H24Rik,Spred-3 | eukaryota>metazoa>chordata>vertebrata | Mus | musculus | PREDICTED: sprouty-related, EVH1 domain-containing protein 3 isoform X7 [Mus musculus] |
| ORX84632.1 | 254 | K493DRAFT_239192 | eukaryota>fungi | Basidiobolus | meristosporus CBS 931.73 | WH1-domain-containing protein [Basidiobolus meristosporus CBS 931.73] |
| EPT04783.1 | 253 | FOMPIDRAFT_1096116 | eukaryota>fungi>basidiomycota | Fomitopsis | pinicola FP-58527 SS1 | hypothetical protein FOMPIDRAFT_1096116, partial [Fomitopsis pinicola FP-58527 SS1] |
| XP_014775879.1 | 250 | LOC106873164 | eukaryota>metazoa>mollusca | Octopus | bimaculoides | PREDICTED: vasodilator-stimulated phosphoprotein-like, partial [Octopus bimaculoides] |
| XP_005264257.2 | 248 | Spred-2 | eukaryota>metazoa>chordata>vertebrata | Homo | sapiens | sprouty-related, EVH1 domain-containing protein 2 isoform X2 [Homo sapiens] |
| XP_021324005.1 | 246 | LOC100330028 | eukaryota>metazoa>chordata>vertebrata>actinopterygii | Danio | rerio | homer protein homolog 3-like isoform X1 [Danio rerio] |
| XP_017207954.1 | 243 | LOC100330028 | eukaryota>metazoa>chordata>vertebrata>actinopterygii | Danio | rerio | homer protein homolog 3-like isoform X2 [Danio rerio] |
| XP_004365930.1 | 243 | CAOG_01059 | eukaryota>ichthyosporea | Capsaspora | owczarzaki ATCC 30864 | hypothetical protein CAOG_01059, partial [Capsaspora owczarzaki ATCC 30864] |
| ORE11625.1 | 239 | BCV72DRAFT_324576 | eukaryota>fungi | Rhizopus | microsporus var. microsporus | WH1-domain-containing protein, partial [Rhizopus microsporus var. microsporus] |
| CAG07864.1 | 237 | GSTEN:00028670:G:001 | eukaryota>metazoa>chordata>vertebrata>actinopterygii | Tetraodon | nigroviridis | unnamed protein product [Tetraodon nigroviridis] |
| XP_009010952.1 | 236 | HELRODRAFT_96961 | eukaryota>metazoa>annelida | Helobdella | robusta | hypothetical protein HELRODRAFT_96961, partial [Helobdella robusta] |
| XP_015770467.1 | 232 | LOC107348895 | eukaryota>metazoa>cnidaria | Acropora | digitifera | PREDICTED: uncharacterized protein LOC107348895 isoform X2 [Acropora digitifera] |
| XP_021331746.1 | 231 | homer1,zgc:100854,zgc:92844 | eukaryota>metazoa>chordata>vertebrata>actinopterygii | Danio | rerio | homer protein homolog 1 isoform X1 [Danio rerio] |
| XP_018274847.1 | 231 | CC85DRAFT_294450 | eukaryota>fungi>basidiomycota | Cutaneotrichosporon | oleaginosum | PH domain-like protein, partial [Cutaneotrichosporon oleaginosum] |
| NP_001264185.1 | 229 | 5730461F13Rik,AL024345,AW047647 | eukaryota>metazoa>chordata>vertebrata | Mus | musculus | sprouty-related, EVH1 domain-containing protein 1 isoform 2 [Mus musculus] |
| XP_009061757.1 | 229 | LOTGIDRAFT_127578 | eukaryota>metazoa>mollusca | Lottia | gigantea | hypothetical protein LOTGIDRAFT_127578, partial [Lottia gigantea] |
| NP_001264006.1 | 224 | HOMER,HOMER1A,HOMER1B,HOMER1C,SYN47,Ves-1 | eukaryota>metazoa>chordata>vertebrata | Homo | sapiens | homer protein homolog 1 isoform 2 [Homo sapiens] |
| XP_012361038.1 | 224 | HOMER1 | eukaryota>metazoa>chordata>vertebrata | Nomascus | leucogenys | PREDICTED: homer protein homolog 1 isoform X2 [Nomascus leucogenys] |
| XP_015770468.1 | 223 | LOC107348895 | eukaryota>metazoa>cnidaria | Acropora | digitifera | PREDICTED: uncharacterized protein LOC107348895 isoform X3 [Acropora digitifera] |
| XP_015770469.1 | 223 | LOC107348895 | eukaryota>metazoa>cnidaria | Acropora | digitifera | PREDICTED: uncharacterized protein LOC107348895 isoform X3 [Acropora digitifera] |
| XP_019856528.1 | 219 | LOC105314031 | eukaryota>metazoa | Amphimedon | queenslandica | PREDICTED: neural Wiskott-Aldrich syndrome protein-like isoform X1 [Amphimedon queenslandica] |
| XP_019856529.1 | 219 | LOC105314031 | eukaryota>metazoa | Amphimedon | queenslandica | PREDICTED: neural Wiskott-Aldrich syndrome protein-like isoform X1 [Amphimedon queenslandica] |
| XP_005264259.1 | 218 | Spred-2 | eukaryota>metazoa>chordata>vertebrata | Homo | sapiens | sprouty-related, EVH1 domain-containing protein 2 isoform X4 [Homo sapiens] |
| XP_017335406.1 | 215 | LOC108271935 | eukaryota>metazoa>chordata>vertebrata>actinopterygii | Ictalurus | punctatus | PREDICTED: wiskott-Aldrich syndrome protein-like isoform X2 [Ictalurus punctatus] |
| ESA14394.1 | 210 | GLOINDRAFT_171470 | eukaryota>fungi | Rhizophagus | irregularis DAOM 181602 | hypothetical protein GLOINDRAFT_171470 [Rhizophagus irregularis DAOM 181602] |
| XP_001901591.1 | 205 | Bm1_50615 | eukaryota>metazoa>nematoda | Brugia | malayi | WH1 domain containing protein [Brugia malayi] |
| XP_017171007.1 | 203 | homer-1,PSD-Zip45,SYN47,Ves-1,vesl-1 | eukaryota>metazoa>chordata>vertebrata | Mus | musculus | PREDICTED: homer protein homolog 1 isoform X1 [Mus musculus] |
| XP_018292777.1 | 203 | PHYBLDRAFT_124118 | eukaryota>fungi | Phycomyces | blakesleeanus NRRL 1555(-) | hypothetical protein PHYBLDRAFT_124118, partial [Phycomyces blakesleeanus NRRL 1555(-)] |
| XP_012567082.1 | 199 | LOC100215288 | eukaryota>metazoa>cnidaria | Hydra | vulgaris | PREDICTED: homer protein homolog 2-like [Hydra vulgaris] |
| XP_018089156.1 | 196 | LOC108700459 | eukaryota>metazoa>chordata>vertebrata | Xenopus | laevis | PREDICTED: sprouty-related, EVH1 domain-containing protein 1-like [Xenopus laevis] |
| KNE72913.1 | 196 | AMAG_20633 | eukaryota>fungi | Allomyces | macrogynus ATCC 38327 | hypothetical protein AMAG_20633 [Allomyces macrogynus ATCC 38327] |
| ELR25045.1 | 194 | ACA1_287240 | eukaryota>amoebozoa>acanthamoebidae | Acanthamoeba | castellanii str. Neff | WH1 domain containing protein [Acanthamoeba castellanii str. Neff] |
| XP_005157353.1 | 193 | homer1,zgc:100854,zgc:92844 | eukaryota>metazoa>chordata>vertebrata>actinopterygii | Danio | rerio | homer protein homolog 1 isoform X2 [Danio rerio] |
| NP_001161217.1 | 193 | 2900021I12Rik,3110031I02Rik,N-WASP | eukaryota>metazoa>chordata>vertebrata | Mus | musculus | neural Wiskott-Aldrich syndrome protein isoform 2 [Mus musculus] |
| CAG12079.1 | 191 | GSTEN:00034222:G:001 | eukaryota>metazoa>chordata>vertebrata>actinopterygii | Tetraodon | nigroviridis | unnamed protein product, partial [Tetraodon nigroviridis] |
| XP_001636870.1 | 190 | NEMVEDRAFT_v1g202473 | eukaryota>metazoa>cnidaria | Nematostella | vectensis | predicted protein [Nematostella vectensis] |
| XP_016865548.1 | 188 | HOMER,HOMER1A,HOMER1B,HOMER1C,SYN47,Ves-1 | eukaryota>metazoa>chordata>vertebrata | Homo | sapiens | homer protein homolog 1 isoform X1 [Homo sapiens] |
| NP_036112.1 | 186 | homer-1,PSD-Zip45,SYN47,Ves-1,vesl-1 | eukaryota>metazoa>chordata>vertebrata | Mus | musculus | homer protein homolog 1 isoform S [Mus musculus] |
| XP_005557308.1 | 184 | HOMER1 | eukaryota>metazoa>chordata>vertebrata | Macaca | fascicularis | PREDICTED: homer protein homolog 1 isoform X2 [Macaca fascicularis] |
| XP_013753353.1 | 184 | AMSG_10645 | eukaryota>apusozoa>apusomonadidae | Thecamonas | trahens ATCC 50062 | hypothetical protein AMSG_10645 [Thecamonas trahens ATCC 50062] |
| XP_013389218.1 | 182 | LOC106157956 | eukaryota>metazoa | Lingula | anatina | protein enabled homolog [Lingula anatina] |
| XP_011606554.1 | 181 | LOC105417035 | eukaryota>metazoa>chordata>vertebrata>actinopterygii | Takifugu | rubripes | PREDICTED: vasodilator-stimulated phosphoprotein-like [Takifugu rubripes] |
| XP_019856530.1 | 179 | LOC105314031 | eukaryota>metazoa | Amphimedon | queenslandica | PREDICTED: neural Wiskott-Aldrich syndrome protein-like isoform X2 [Amphimedon queenslandica] |
| NP_001264007.1 | 177 | HOMER,HOMER1A,HOMER1B,HOMER1C,SYN47,Ves-1 | eukaryota>metazoa>chordata>vertebrata | Homo | sapiens | homer protein homolog 1 isoform 3 [Homo sapiens] |
| XP_012361039.1 | 177 | HOMER1 | eukaryota>metazoa>chordata>vertebrata | Nomascus | leucogenys | PREDICTED: homer protein homolog 1 isoform X3 [Nomascus leucogenys] |
| CBY34382.1 | 177 | GSOID_T00024405001 | eukaryota>metazoa>chordata | Oikopleura | dioica | unnamed protein product [Oikopleura dioica] |
| XP_021323364.1 | 176 | si:ch211-255112.2 | eukaryota>metazoa>chordata>vertebrata>actinopterygii | Danio | rerio | sprouty-related, EVH1 domain containing 3 isoform X4 [Danio rerio] |
| XP_009044835.1 | 173 | LOTGIDRAFT_170749 | eukaryota>metazoa>mollusca | Lottia | gigantea | hypothetical protein LOTGIDRAFT_170749 [Lottia gigantea] |
| EFX81577.1 | 161 | DAPPUDRAFT_242162 | eukaryota>metazoa>crustacea | Daphnia | pulex | hypothetical protein DAPPUDRAFT_242162 [Daphnia pulex] |
| ETO21130.1 | 156 | RFI_16075 | eukaryota>rhizaria | Reticulomyxa | filosa | hypothetical protein RFI_16075 [Reticulomyxa filosa] |
| XP_019040119.1 | 154 | WICANDRAFT_29494 | eukaryota>fungi>ascomycota | Wickerhamomyces | anomalus NRRL Y-366-8 | hypothetical protein WICANDRAFT_29494, partial [Wickerhamomyces anomalus NRRL Y-366-8] |
| XP_013389219.1 | 152 | LOC106157956 | eukaryota>metazoa | Lingula | anatina | PREDICTED: protein enabled homolog isoform X2 [Lingula anatina] |
| EFX77865.1 | 151 | DAPPUDRAFT_213242 | eukaryota>metazoa>crustacea | Daphnia | pulex | hypothetical protein DAPPUDRAFT_213242, partial [Daphnia pulex] |
| XP_022249327.1 | 145 | LOC111087333 | eukaryota>metazoa | Limulus | polyphemus | homer protein homolog 2-like [Limulus polyphemus] |
| XP_004212513.1 | 145 | LOC101238251 | eukaryota>metazoa>cnidaria | Hydra | vulgaris | PREDICTED: wiskott-Aldrich syndrome protein-like [Hydra vulgaris] |
| XP_006822068.1 | 138 | LOC100367435 | eukaryota>metazoa>hemichordata | Saccoglossus | kowalevskii | PREDICTED: protein still life, isoform SIF type 1-like [Saccoglossus kowalevskii] |
| XP_009018259.1 | 134 | HELRODRAFT_79834 | eukaryota>metazoa>annelida | Helobdella | robusta | hypothetical protein HELRODRAFT_79834, partial [Helobdella robusta] |
| XP_009055943.1 | 134 | LOTGIDRAFT_119577 | eukaryota>metazoa>mollusca | Lottia | gigantea | hypothetical protein LOTGIDRAFT_119577 [Lottia gigantea] |
| XP_001633271.1 | 128 | NEMVEDRAFT_v1g104827 | eukaryota>metazoa>cnidaria | Nematostella | vectensis | predicted protein, partial [Nematostella vectensis] |
| XP_009166784.1 | 128 | T265_03919 | eukaryota>metazoa | Opisthorchis | viverrini | hypothetical protein T265_03919 [Opisthorchis viverrini] |
| XP_018668300.1 | 127 | LOC100179781 | eukaryota>metazoa>chordata | Ciona | intestinalis | PREDICTED: neural Wiskott-Aldrich syndrome protein-like [Ciona intestinalis] |
| XP_002585829.1 | 126 | BRAFLDRAFT_256955 | eukaryota>metazoa>chordata | Branchiostoma | floridae | hypothetical protein BRAFLDRAFT_256955 [Branchiostoma floridae] |
| XP_009016952.1 | 125 | HELRODRAFT_78249 | eukaryota>metazoa>annelida | Helobdella | robusta | hypothetical protein HELRODRAFT_78249, partial [Helobdella robusta] |
| XP_001743073.1 | 119 | MONBRDRAFT_14345 | eukaryota>choanoflagellida | Monosiga | brevicollis MX1 | hypothetical protein, partial [Monosiga brevicollis MX1] |
| XP_009021879.1 | 118 | HELRODRAFT_127531 | eukaryota>metazoa>annelida | Helobdella | robusta | hypothetical protein HELRODRAFT_127531, partial [Helobdella robusta] |
| XP_009052802.1 | 118 | LOTGIDRAFT_214569 | eukaryota>metazoa>mollusca | Lottia | gigantea | hypothetical protein LOTGIDRAFT_214569 [Lottia gigantea] |
| XP_681984.1 | 118 | AN8715.2 | eukaryota>fungi>ascomycota | Aspergillus | nidulans FGSC A4 | hypothetical protein AN8715.2 [Aspergillus nidulans FGSC A4] |
| XP_001619064.1 | 117 | NEMVEDRAFT_v1g224549 | eukaryota>metazoa>cnidaria | Nematostella | vectensis | hypothetical protein NEMVEDRAFT_v1g224549 [Nematostella vectensis] |
| OAD07496.1 | 106 | MUCCIDRAFT_123792 | eukaryota>fungi | Mucor | circinelloides f. lusitanicus CBS 277.49 | hypothetical protein MUCCIDRAFT_123792, partial [Mucor circinelloides f. lusitanicus CBS 277.49] |
| OAC98115.1 | 104 | MUCCIDRAFT_123755 | eukaryota>fungi | Mucor | circinelloides f. lusitanicus CBS 277.49 | hypothetical protein MUCCIDRAFT_123755, partial [Mucor circinelloides f. lusitanicus CBS 277.49] |
| XP_009022206.1 | 101 | HELRODRAFT_127019 | eukaryota>metazoa>annelida | Helobdella | robusta | hypothetical protein HELRODRAFT_127019, partial [Helobdella robusta] |
| XP_002111045.1 | 98 | TRIADDRAFT_22944 | eukaryota>metazoa>placozoa | Trichoplax | adhaerens | hypothetical protein TRIADDRAFT_22944, partial [Trichoplax adhaerens] |

N. ERdj5

| | | | | | | |
|---|---|---|---|---|---|---|
| OLP80009.1 | 1597 | DNAJB11 | eukaryota>alveolata | Symbiodinium | microadriaticum | DnaJ-like subfamily B member 11, partial [Symbiodinium microadriaticum] |
| CEO95860.1 | 918 | PBRA_004573 | eukaryota>rhizaria>cercozoa>plasmodiophorida | Plasmodiophora | brassicae | hypothetical protein PBRA_004573 [Plasmodiophora brassicae] |
| KXJ25509.1 | 875 | dnajc10 | eukaryota>metazoa>cnidaria | Exaiptasia | pallida | DnaJ-like subfamily C member 10 [Exaiptasia pallida] |
| XP_009026746.1 | 872 | HELRODRAFT_193633 | eukaryota>metazoa>annelida | Helobdella | robusta | hypothetical protein HELRODRAFT_193633 [Helobdella robusta] |
| XP_001635821.1 | 844 | NEMVEDRAFT_v1g163820 | eukaryota>metazoa>cnidaria | Nematostella | vectensis | predicted protein [Nematostella vectensis] |
| XP_001894589.1 | 839 | Bm1_15660 | eukaryota>metazoa>nematoda | Brugia | malayi | DnaJ domain containing protein [Brugia malayi] |
| CAG01766.1 | 831 | GSTEN:00020649:G:001 | eukaryota>metazoa>chordata>vertebrata>actinopterygii | Tetraodon | nigroviridis | unnamed protein product, partial [Tetraodon nigroviridis] |
| XP_001862421.1 | 831 | CpipJ_CPIJ012337 | eukaryota>metazoa>hexapoda | Culex | quinquefasciatus | conserved hypothetical protein [Culex quinquefasciatus] |
| XP_017325246.1 | 830 | dnajc10 | eukaryota>metazoa>chordata>vertebrata>actinopterygii | Ictalurus | punctatus | PREDICTED: dnaJ homolog subfamily C member 10 isoform X1 [Ictalurus punctatus] |
| ERG81725.1 | 830 | ASU_09741 | eukaryota>metazoa>nematoda | Ascaris | suum | hypothetical protein ASU_09741 [Ascaris suum] |
| Capa1000007577 | 826 | Caps1000007577 | eukaryota>metazoa>annelida | Capitella | spI | >Caps1000007577 Capa1000007577 |
| XP_395584.4 | 815 | LOC412118 | eukaryota>metazoa>hexapoda | Apis | mellifera | PREDICTED: dnaJ homolog subfamily C member 16 [Apis mellifera] |
| NP_001040753.1 | 813 | dnj-8 | eukaryota>metazoa>nematoda | Caenorhabditis | elegans | DNaJ domain (prokaryotic heat shock protein) [Caenorhabditis elegans] |
| XP_011146374.1 | 812 | LOC105187332 | eukaryota>metazoa>hexapoda | Harpegnathos | saltator | dnaJ homolog subfamily C member 16 [Harpegnathos saltator] |
| XP_002122728.1 | 811 | LOC100179175 | eukaryota>metazoa>chordata | Ciona | intestinalis | PREDICTED: dnaJ homolog subfamily C member 10 [Ciona intestinalis] |
| XP_019700629.1 | 810 | LOC105191156 | eukaryota>metazoa>hexapoda | Harpegnathos | saltator | dnaJ homolog subfamily C member 10 isoform X1 [Harpegnathos saltator] |
| CEL95543.1 | 810 | Vbra_11973 | eukaryota>alveolata | Vitrella | brassicaformis CCMP3155 | unnamed protein product [Vitrella brassicaformis CCMP3155] |
| XP_003372029.1 | 809 | Tsp_05680 | eukaryota>metazoa>nematoda | Trichinella | spiralis | DnaJ protein subfamily C member 10 [Trichinella spiralis] |
| CBY20690.1 | 807 | GSOID_T00000693001 | eukaryota>metazoa>chordata | Oikopleura | dioica | unnamed protein product [Oikopleura dioica] |
| EFX65186.1 | 807 | DAPPUDRAFT_333446 | eukaryota>metazoa>crustacea | Daphnia | pulex | hypothetical protein DAPPUDRAFT_333446 [Daphnia pulex] |
| XP_019700630.1 | 807 | LOC105191156 | eukaryota>metazoa>hexapoda | Harpegnathos | saltator | dnaJ homolog subfamily C member 10 isoform X3 [Harpegnathos saltator] |
| XP_782974.2 | 807 | LOC577666 | eukaryota>metazoa>echinodermata | Strongylocentrotus | purpuratus | PREDICTED: dnaJ homolog subfamily C member 10 [Strongylocentrotus purpuratus] |
| XP_003967687.1 | 807 | LOC101061594 | eukaryota>metazoa>chordata>vertebrata>actinopterygii | Takifugu | rubripes | PREDICTED: dnaJ homolog subfamily C member 16-like [Takifugu rubripes] |
| XP_011152658.1 | 806 | LOC105191156 | eukaryota>metazoa>hexapoda | Harpegnathos | saltator | dnaJ homolog subfamily C member 10 isoform X4 [Harpegnathos saltator] |
| XP_015287942.1 | 802 | DNAJC10 | eukaryota>metazoa>chordata>vertebrata | Macaca | fascicularis | PREDICTED: dnaJ homolog subfamily C member 10 isoform X1 [Macaca fascicularis] |
| XP_015287943.1 | 802 | DNAJC10 | eukaryota>metazoa>chordata>vertebrata | Macaca | fascicularis | PREDICTED: dnaJ homolog subfamily C member 10 isoform X1 [Macaca fascicularis] |
| XP_015287944.1 | 802 | DNAJC10 | eukaryota>metazoa>chordata>vertebrata | Macaca | fascicularis | PREDICTED: dnaJ homolog subfamily C member 10 isoform X1 [Macaca fascicularis] |
| XP_017325247.1 | 801 | dnajc10 | eukaryota>metazoa>chordata>vertebrata>actinopterygii | Ictalurus | punctatus | PREDICTED: dnaJ homolog subfamily C member 10 isoform X2 [Ictalurus punctatus] |
| XP_017325248.1 | 801 | dnajc10 | eukaryota>metazoa>chordata>vertebrata>actinopterygii | Ictalurus | punctatus | PREDICTED: dnaJ homolog subfamily C member 10 isoform X2 [Ictalurus punctatus] |
| XP_002170794.2 | 798 | LOC100206669 | eukaryota>metazoa>cnidaria | Hydra | vulgaris | PREDICTED: dnaJ homolog subfamily C member 10 [Hydra vulgaris] |
| XP_003961647.2 | 798 | dnajc10 | eukaryota>metazoa>chordata>vertebrata>actinopterygii | Takifugu | rubripes | PREDICTED: dnaJ homolog subfamily C member 10 [Takifugu rubripes] |
| XP_018091505.1 | 797 | LOC108701402 | eukaryota>metazoa>chordata>vertebrata | Xenopus | laevis | PREDICTED: dnaJ homolog subfamily C member 10-like [Xenopus laevis] |
| XP_018091506.1 | 797 | LOC108701402 | eukaryota>metazoa>chordata>vertebrata | Xenopus | laevis | PREDICTED: dnaJ homolog subfamily C member 10-like [Xenopus laevis] |
| XP_018091507.1 | 797 | LOC108701402 | eukaryota>metazoa>chordata>vertebrata | Xenopus | laevis | PREDICTED: dnaJ homolog subfamily C member 10-like [Xenopus laevis] |
| XP_421968.2 | 797 | DNAJC10 | eukaryota>metazoa>chordata>vertebrata | Gallus | gallus | dnaJ homolog subfamily C member 10 [Gallus gallus] |
| NP_001084933.1 | 796 | dnajc10 | eukaryota>metazoa>chordata>vertebrata | Xenopus | laevis | dnaJ homolog subfamily C member 10 precursor [Xenopus laevis] |
| EFX69369.1 | 794 | DAPPUDRAFT_113685 | eukaryota>metazoa>crustacea | Daphnia | pulex | hypothetical protein DAPPUDRAFT_113685 [Daphnia pulex] |
| NP_061854.1 | 793 | ERdj5,JPDI,MTHr,PDIA19 | eukaryota>metazoa>chordata>vertebrata | Homo | sapiens | dnaJ homolog subfamily C member 10 isoform 1 precursor [Homo sapiens] |
| NP_077143.2 | 793 | 1200006L06Rik,D2Ertd706e,ERdj5,JPDI | eukaryota>metazoa>chordata>vertebrata | Mus | musculus | dnaJ homolog subfamily C member 10 precursor [Mus musculus] |
| XP_005573707.1 | 793 | DNAJC10 | eukaryota>metazoa>chordata>vertebrata | Macaca | fascicularis | PREDICTED: dnaJ homolog subfamily C member 10 isoform X2 [Macaca fascicularis] |
| XP_005573709.1 | 793 | DNAJC10 | eukaryota>metazoa>chordata>vertebrata | Macaca | fascicularis | PREDICTED: dnaJ homolog subfamily C member 10 isoform X2 [Macaca fascicularis] |
| XP_013395942.1 | 793 | LOC106163023 | eukaryota>metazoa | Lingula | anatina | dnaJ homolog subfamily C member 10 [Lingula anatina] |
| XP_013774797.1 | 793 | LOC106459696 | eukaryota>metazoa | Limulus | polyphemus | dnaJ homolog subfamily C member 16-like [Limulus polyphemus] |
| XP_013773201.1 | 792 | LOC106458262 | eukaryota>metazoa | Limulus | polyphemus | dnaJ homolog subfamily C member 10-like [Limulus polyphemus] |
| NP_001077016.1 | 791 | zgc:162218 | eukaryota>metazoa>chordata>vertebrata>actinopterygii | Danio | rerio | dnaJ homolog subfamily C member 10 [Danio rerio] |
| XP_011449285.1 | 791 | LOC105343574 | eukaryota>metazoa>mollusca | Crassostrea | gigas | PREDICTED: dnaJ homolog subfamily C member 10 isoform X1 [Crassostrea gigas] |
| XP_012356558.1 | 791 | DNAJC10 | eukaryota>metazoa>chordata>vertebrata | Nomascus | leucogenys | PREDICTED: LOW QUALITY PROTEIN: dnaJ homolog subfamily C member 10 [Nomascus leucogenys] |
| XP_019929032.1 | 791 | LOC105343574 | eukaryota>metazoa>mollusca | Crassostrea | gigas | PREDICTED: dnaJ homolog subfamily C member 10 isoform X1 [Crassostrea gigas] |
| XP_002410093.1 | 791 | IscW_ISCW020369 | eukaryota>metazoa | Ixodes | scapularis | conserved hypothetical protein [Ixodes scapularis] |
| XP_013775072.1 | 791 | LOC106459946 | eukaryota>metazoa | Limulus | polyphemus | dnaJ homolog subfamily C member 16-like isoform X2 [Limulus polyphemus] |
| XP_011449286.1 | 790 | LOC105343574 | eukaryota>metazoa>mollusca | Crassostrea | gigas | PREDICTED: dnaJ homolog subfamily C member 10 isoform X2 [Crassostrea gigas] |
| XP_002663742.2 | 789 | dnajc16 | eukaryota>metazoa>chordata>vertebrata>actinopterygii | Danio | rerio | PREDICTED: dnaJ homolog subfamily C member 16 [Danio rerio] |
| NP_001040704.1 | 788 | dnj-27 | eukaryota>metazoa>nematoda | Caenorhabditis | elegans | DNaJ domain (prokaryotic heat shock protein) [Caenorhabditis elegans] |
| XP_002119344.1 | 787 | LOC100176543 | eukaryota>metazoa>chordata | Ciona | intestinalis | PREDICTED: dnaJ homolog subfamily C member 16-like [Ciona intestinalis] |
| XP_017342364.1 | 787 | dnajc16 | eukaryota>metazoa>chordata>vertebrata>actinopterygii | Ictalurus | punctatus | PREDICTED: dnaJ homolog subfamily C member 16 [Ictalurus punctatus] |
| XP_017342365.1 | 787 | dnajc16 | eukaryota>metazoa>chordata>vertebrata>actinopterygii | Ictalurus | punctatus | PREDICTED: dnaJ homolog subfamily C member 16 [Ictalurus punctatus] |
| XP_003978530.1 | 786 | dnajc16 | eukaryota>metazoa>chordata>vertebrata>actinopterygii | Takifugu | rubripes | PREDICTED: dnaJ homolog subfamily C member 16 [Takifugu rubripes] |
| XP_012552871.1 | 786 | LOC101737705 | eukaryota>metazoa>hexapoda | Bombyx | mori | PREDICTED: dnaJ homolog subfamily C member 16 [Bombyx mori] |
| OAF69206.1 | 785 | A3Q56_03049 | eukaryota>metazoa | Intoshia | linei | DnaJ subfamily C member 10 [Intoshia linei] |
| XP_008190269.1 | 785 | LOC660466 | eukaryota>metazoa>hexapoda | Tribolium | castaneum | PREDICTED: dnaJ homolog subfamily C member 10-like [Tribolium castaneum] |
| Capa1000028443 | 785 | Caps1000028443 | eukaryota>metazoa>annelida | Capitella | spI | >Caps1000028443 Capa1000028443 |
| XP_002435441.1 | 782 | IscW_ISCW018779 | eukaryota>metazoa | Ixodes | scapularis | DnaJ domain and thioredoxin-containing protein, putative [Ixodes scapularis] |
| NP_056106.1 | 782 | DNAJC16 | eukaryota>metazoa>chordata>vertebrata | Homo | sapiens | dnaJ homolog subfamily C member 16 isoform 1 precursor [Homo sapiens] |
| XP_012354025.1 | 782 | DNAJC16 | eukaryota>metazoa>chordata>vertebrata | Nomascus | leucogenys | PREDICTED: dnaJ homolog subfamily C member 16 isoform X1 [Nomascus leucogenys] |
| XP_015297931.1 | 782 | DNAJC16 | eukaryota>metazoa>chordata>vertebrata | Macaca | fascicularis | PREDICTED: dnaJ homolog subfamily C member 16 [Macaca fascicularis] |
| XP_015297936.1 | 782 | DNAJC16 | eukaryota>metazoa>chordata>vertebrata | Macaca | fascicularis | PREDICTED: dnaJ homolog subfamily C member 16 [Macaca fascicularis] |
| XP_969135.2 | 781 | LOC657591 | eukaryota>metazoa>hexapoda | Tribolium | castaneum | PREDICTED: dnaJ homolog subfamily C member 16 isoform X1 [Tribolium castaneum] |
| NP_001015187.2 | 780 | CG40178/CG41283,Dmel\CG40178 | eukaryota>metazoa>hexapoda | Drosophila | melanogaster | uncharacterized protein Dmel_CG40178, isoform A [Drosophila melanogaster] |
| NP_001015188.2 | 780 | CG40178/CG41283,Dmel\CG40178 | eukaryota>metazoa>hexapoda | Drosophila | melanogaster | uncharacterized protein Dmel_CG40178, isoform B [Drosophila melanogaster] |
| NP_001263164.1 | 780 | CG40178/CG41283,Dmel\CG40178 | eukaryota>metazoa>hexapoda | Drosophila | melanogaster | uncharacterized protein Dmel_CG40178, isoform C [Drosophila melanogaster] |
| XP_001901387.1 | 779 | Bm1_49600 | eukaryota>metazoa>nematoda | Brugia | malayi | Thioredoxin family protein [Brugia malayi] |
| XP_002605314.1 | 779 | BRAFLDRAFT_125408 | eukaryota>metazoa>chordata | Branchiostoma | floridae | hypothetical protein BRAFLDRAFT_125408 [Branchiostoma floridae] |
| XP_019700631.1 | 778 | LOC105191156 | eukaryota>metazoa>hexapoda | Harpegnathos | saltator | dnaJ homolog subfamily C member 10 isoform X6 [Harpegnathos saltator] |
| XP_012546160.1 | 778 | LOC101737299 | eukaryota>metazoa>hexapoda | Bombyx | mori | dnaJ homolog subfamily C member 10 isoform X1 [Bombyx mori] |
| NP_001034419.1 | 777 | DNAJC16 | eukaryota>metazoa>chordata>vertebrata | Gallus | gallus | dnaJ homolog subfamily C member 16 precursor [Gallus gallus] |
| XP_005166271.1 | 777 | dnajc16 | eukaryota>metazoa>chordata>vertebrata>actinopterygii | Danio | rerio | dnaJ homolog subfamily C member 16 isoform X1 [Danio rerio] |
| XP_018082048.1 | 777 | LOC108696869 | eukaryota>metazoa>chordata>vertebrata | Xenopus | laevis | PREDICTED: dnaJ homolog subfamily C member 16-like [Xenopus laevis] |
| XP_018083834.1 | 777 | LOC108697879 | eukaryota>metazoa>chordata>vertebrata | Xenopus | laevis | PREDICTED: dnaJ homolog subfamily C member 16-like [Xenopus laevis] |
| XP_009046138.1 | 776 | LOTGIDRAFT_171701 | eukaryota>metazoa>mollusca | Lottia | gigantea | hypothetical protein LOTGIDRAFT_171701 [Lottia gigantea] |
| XP_015287945.1 | 776 | DNAJC10 | eukaryota>metazoa>chordata>vertebrata | Macaca | fascicularis | PREDICTED: dnaJ homolog subfamily C member 10 isoform X3 [Macaca fascicularis] |
| XP_015152426.1 | 776 | DNAJC16 | eukaryota>metazoa>chordata>vertebrata | Gallus | gallus | PREDICTED: dnaJ homolog subfamily C member 16 isoform X1 [Gallus gallus] |
| XP_688223.4 | 776 | dnajc16 | eukaryota>metazoa>chordata>vertebrata>actinopterygii | Danio | rerio | dnaJ homolog subfamily C member 16 isoform X2 [Danio rerio] |
| XP_011442421.1 | 774 | LOC105338837 | eukaryota>metazoa>mollusca | Crassostrea | gigas | PREDICTED: dnaJ homolog subfamily C member 16 [Crassostrea gigas] |
| XP_009026128.1 | 773 | HELRODRAFT_102482 | eukaryota>metazoa>annelida | Helobdella | robusta | hypothetical protein HELRODRAFT_102482 [Helobdella robusta] |
| Smar1000011868 | 772 | Smar1000011868 | eukaryota>metazoa>arthropoda>myriapoda | Strigamia | maritima | >Smar1000011868 Smar1000011868 |
| NP_758841.1 | 772 | 2900037O03Rik,4732437J24Rik,D030060M24,mKIAA0962 | eukaryota>metazoa>chordata>vertebrata | Mus | musculus | dnaJ homolog subfamily C member 16 precursor [Mus musculus] |
| XP_001010992.2 | 768 | TTHERM_00709590 | eukaryota>alveolata>ciliophora | Tetrahymena | thermophila SB210 | DnaJ domain protein [Tetrahymena thermophila SB210] |
| XP_008197271.1 | 767 | LOC657591 | eukaryota>metazoa>hexapoda | Tribolium | castaneum | PREDICTED: dnaJ homolog subfamily C member 16 isoform X2 [Tribolium castaneum] |
| XP_006815097.1 | 766 | LOC102808049 | eukaryota>metazoa>hemichordata | Saccoglossus | kowalevskii | PREDICTED: dnaJ homolog subfamily C member 16-like [Saccoglossus kowalevskii] |
| XP_008197276.1 | 766 | LOC657591 | eukaryota>metazoa>hexapoda | Tribolium | castaneum | PREDICTED: dnaJ homolog subfamily C member 16 isoform X3 [Tribolium castaneum] |
| Smar1000014991 | 765 | Smar1000014991 | eukaryota>metazoa>arthropoda>myriapoda | Strigamia | maritima | >Smar1000014991 Smar1000014991 |
| XP_002740489.1 | 763 | LOC100374701 | eukaryota>metazoa>hemichordata | Saccoglossus | kowalevskii | PREDICTED: dnaJ homolog subfamily C member 10-like [Saccoglossus kowalevskii] |
| XP_014787454.1 | 763 | LOC106881543 | eukaryota>metazoa>mollusca | Octopus | bimaculoides | PREDICTED: dnaJ homolog subfamily C member 16-like [Octopus bimaculoides] |
| XP_017306360.1 | 761 | LOC108255153 | eukaryota>metazoa>chordata>vertebrata>actinopterygii | Ictalurus | punctatus | PREDICTED: dnaJ homolog subfamily C member 16-like [Ictalurus punctatus] |
| XP_017306361.1 | 761 | LOC108255153 | eukaryota>metazoa>chordata>vertebrata>actinopterygii | Ictalurus | punctatus | PREDICTED: dnaJ homolog subfamily C member 16-like [Ictalurus punctatus] |
| XP_014771843.1 | 757 | LOC106870330 | eukaryota>metazoa>mollusca | Octopus | bimaculoides | PREDICTED: dnaJ homolog subfamily C member 16-like [Octopus bimaculoides] |
| XP_012556009.1 | 756 | LOC100199979 | eukaryota>metazoa>cnidaria | Hydra | vulgaris | PREDICTED: dnaJ homolog subfamily C member 16-like [Hydra vulgaris] |
| XP_019700632.1 | 755 | LOC105191156 | eukaryota>metazoa>hexapoda | Harpegnathos | saltator | PREDICTED: dnaJ homolog subfamily C member 10 isoform X5 [Harpegnathos saltator] |
| XP_003372041.1 | 753 | Tsp_05693 | eukaryota>metazoa>nematoda | Trichinella | spiralis | putative DnaJ domain protein [Trichinella spiralis] |
| XP_001742547.1 | 753 | MONBRDRAFT_21911 | eukaryota>choanoflagellida | Monosiga | brevicollis MX1 | hypothetical protein [Monosiga brevicollis MX1] |
| NP_001258510.1 | 747 | ERdj5,JPDI,MTHr,PDIA19 | eukaryota>metazoa>chordata>vertebrata | Homo | sapiens | dnaJ homolog subfamily C member 10 isoform 2 precursor [Homo sapiens] |
| XP_002591687.1 | 734 | BRAFLDRAFT_223511 | eukaryota>metazoa>chordata | Branchiostoma | floridae | hypothetical protein BRAFLDRAFT_223511 [Branchiostoma floridae] |
| XP_019700633.1 | 731 | LOC105191156 | eukaryota>metazoa>hexapoda | Harpegnathos | saltator | PREDICTED: dnaJ homolog subfamily C member 10 isoform X6 [Harpegnathos saltator] |
| XP_011664449.1 | 707 | LOC582790 | eukaryota>metazoa>echinodermata | Strongylocentrotus | purpuratus | PREDICTED: dnaJ homolog subfamily C member 16, partial [Strongylocentrotus purpuratus] |
| XP_019700634.1 | 689 | LOC105191156 | eukaryota>metazoa>hexapoda | Harpegnathos | saltator | dnaJ homolog subfamily C member 10 isoform X7 [Harpegnathos saltator] |
| XP_004993461.1 | 683 | PTSG_05593 | eukaryota>choanoflagellida | Salpingoeca | rosetta | molecular chaperone DnaJ [Salpingoeca rosetta] |
| XP_002141552.1 | 678 | CMU_000730 | eukaryota>alveolata>apicomplexa | Cryptosporidium | muris RN66 | DnaJ domain-containing protein [Cryptosporidium muris RN66] |
| XP_008887413.1 | 659 | HHA_204480 | eukaryota>alveolata>apicomplexa | Hammondia | hammondi | thioredoxin domain-containing protein [Hammondia hammondi] |
| XP_018637303.1 | 659 | TGME49_204480 | eukaryota>alveolata>apicomplexa | Toxoplasma | gondii ME49 | thioredoxin domain-containing protein [Toxoplasma gondii ME49] |
| XP_015751062.1 | 658 | LOC107331046 | eukaryota>metazoa>cnidaria | Acropora | digitifera | PREDICTED: dnaJ homolog subfamily C member 10-like [Acropora digitifera] |
| XP_012354026.1 | 655 | DNAJC16 | eukaryota>metazoa>chordata>vertebrata | Nomascus | leucogenys | PREDICTED: dnaJ homolog subfamily C member 16 isoform X2 [Nomascus leucogenys] |
| XP_009032704.1 | 635 | AURANDRAFT_60606 | eukaryota>stramenopiles | Aureococcus | anophagefferens | hypothetical protein AURANDRAFT_60606 [Aureococcus anophagefferens] |
| XP_626829.1 | 631 | cgd3_2690 | eukaryota>alveolata>apicomplexa | Cryptosporidium | parvum Iowa II | DNAj domain, possible transmembrane domain [Cryptosporidium parvum Iowa II] |
| XP_013401458.1 | 624 | LOC106167272 | eukaryota>metazoa | Lingula | anatina | dnaJ homolog subfamily C member 10-like, partial [Lingula anatina] |
| XP_002270607.2 | 618 | LOC100260777 | eukaryota>viridiplantae | Vitis | vinifera | PREDICTED: dnaJ protein ERDJ3A isoform X1 [Vitis vinifera] |
| CEP00246.1 | 601 | PBRA_007980 | eukaryota>rhizaria>cercozoa>plasmodiophorida | Plasmodiophora | brassicae | hypothetical protein PBRA_007980 [Plasmodiophora brassicae] |
| XP_001388233.1 | 601 | cgd2_3230 | eukaryota>alveolata>apicomplexa | Cryptosporidium | parvum Iowa II | heat shock protein DnaJ Pfj2 [Cryptosporidium parvum Iowa II] |
| XP_012546161.1 | 597 | LOC101737299 | eukaryota>metazoa>hexapoda | Bombyx | mori | dnaJ homolog subfamily C member 10 isoform X2 [Bombyx mori] |
| XP_010653325.1 | 596 | LOC100260777 | eukaryota>viridiplantae | Vitis | vinifera | PREDICTED: dnaJ protein ERDJ3A isoform X2 [Vitis vinifera] |
| XP_014769377.1 | 586 | LOC106868569 | eukaryota>metazoa>mollusca | Octopus | bimaculoides | PREDICTED: dnaJ homolog subfamily C member 10-like [Octopus bimaculoides] |
| XP_006362374.1 | 585 | LOC102579268 | eukaryota>viridiplantae | Solanum | tuberosum | PREDICTED: dnaJ protein ERDJ3A [Solanum tuberosum] |
| XP_015158438.1 | 585 | LOC102579268 | eukaryota>viridiplantae | Solanum | tuberosum | PREDICTED: dnaJ protein ERDJ3A [Solanum tuberosum] |
| NP_001169161.1 | 583 | GRMZM5G898471 | eukaryota>viridiplantae | Zea | mays | uncharacterized LOC100383011 precursor [Zea mays] |
| GAQ91638.1 | 582 | KFL_008230030 | eukaryota>viridiplantae | Klebsormidium | nitens | DnaJ heat shock family protein [Klebsormidium nitens] |
| XP_003530294.1 | 580 | LOC100785691 | eukaryota>viridiplantae | Glycine | max | PREDICTED: dnaJ protein ERDJ3A-like [Glycine max] |
| XP_013386220.1 | 580 | LOC106155772 | eukaryota>metazoa | Lingula | anatina | dnaJ homolog subfamily C member 16, partial [Lingula anatina] |
| XP_002142846.1 | 579 | CMU_003270 | eukaryota>alveolata>apicomplexa | Cryptosporidium | muris RN66 | DnaJ / Thioredoxin domain-containing protein [Cryptosporidium muris RN66] |
| XP_003551564.1 | 577 | LOC100796871 | eukaryota>viridiplantae | Glycine | max | PREDICTED: dnaJ protein ERDJ3A [Glycine max] |
| XP_001610190.1 | 577 | 18.m06550 | eukaryota>alveolata>apicomplexa | Babesia | bovis T2Bo | DnaJ domain containing protein [Babesia bovis T2Bo] |
| XP_016447176.1 | 574 | LOC107772193 | eukaryota>viridiplantae | Nicotiana | tabacum | PREDICTED: dnaJ protein ERDJ3A-like [Nicotiana tabacum] |
| XP_016467250.1 | 573 | LOC107789888 | eukaryota>viridiplantae | Nicotiana | tabacum | PREDICTED: dnaJ protein ERDJ3A-like [Nicotiana tabacum] |
| NP_001319504.1 | 572 | THERMOSENSITIVE,MALE,STERILE,1,TMS1 | eukaryota>viridiplantae | Arabidopsis | thaliana | DNAJ heat shock N-terminal domain-containing protein [Arabidopsis thaliana] |
| NP_187509.1 | 572 | THERMOSENSITIVE,MALE,STERILE,1,TMS1 | eukaryota>viridiplantae | Arabidopsis | thaliana | DNAJ heat shock N-terminal domain-containing protein [Arabidopsis thaliana] |
| XP_003078103.1 | 565 | Ot03g01910 | eukaryota>viridiplantae>chlorophyta | Ostreococcus | tauri | CG4599-PB, isoform B (ISS) [Ostreococcus tauri] |
| XP_012766217.1 | 563 | BBBOND_0103450 | eukaryota>alveolata>apicomplexa | Babesia | bigemina | DnaJ domain containing protein, putative [Babesia bigemina] |
| EWM30375.1 | 559 | Naga_100026g6 | eukaryota>stramenopiles | Nannochloropsis | gaditana | disulfide isomerase [Nannochloropsis gaditana] |
| XP_001580079.1 | 559 | TVAG_189840 | eukaryota>parabasalia | Trichomonas | vaginalis G3 | DnaJ domain containing protein [Trichomonas vaginalis G3] |
| XP_009690664.1 | 557 | TOT_020000621 | eukaryota>alveolata>apicomplexa | Theileria | orientalis strain Shintoku | molecular chaperone protein DnaJ [Theileria orientalis strain Shintoku] |
| XP_727981.1 | 553 | PY17X_0940800 | eukaryota>alveolata>apicomplexa | Plasmodium | yoelii | heat shock protein DnaJ [Plasmodium yoelii] |
| OHT15350.1 | 551 | TRFO_42559 | eukaryota>parabasalia | Tritrichomonas | foetus | hypothetical protein TRFO_42559 [Tritrichomonas foetus] |
| XP_002778681.1 | 550 | Pmar_PMAR005811 | eukaryota>alveolata | Perkinsus | marinus ATCC 50983 | hypothetical protein Pmar_PMAR005811 [Perkinsus marinus ATCC 50983] |
| XP_002778682.1 | 550 | Pmar_PMAR005811 | eukaryota>alveolata | Perkinsus | marinus ATCC 50983 | hypothetical protein Pmar_PMAR005811 [Perkinsus marinus ATCC 50983] |
| XP_001615215.1 | 549 | PVX_091110 | eukaryota>alveolata>apicomplexa | Plasmodium | vivax | DnaJ domain containing protein [Plasmodium vivax] |
| XP_012768962.1 | 547 | BBBOND_0306800 | eukaryota>alveolata>apicomplexa | Babesia | bigemina | DnaJ domain containing protein, putative [Babesia bigemina] |
| XP_952078.1 | 542 | TA14295 | eukaryota>alveolata>apicomplexa | Theileria | annulata | hypothetical protein, conserved [Theileria annulata] |
| OHT05368.1 | 542 | TRFO_05951 | eukaryota>parabasalia | Tritrichomonas | foetus | hypothetical protein TRFO_05951 [Tritrichomonas foetus] |
| XP_001347774.1 | 540 | PF3D7_1108700 | eukaryota>alveolata>apicomplexa | Plasmodium | falciparum 3D7 | heat shock protein DnaJ homologue Pfj2 [Plasmodium falciparum 3D7] |
| XP_001783528.1 | 540 | PHYPADRAFT_226536 | eukaryota>viridiplantae | Physcomitrella | patens | predicted protein [Physcomitrella patens] |
| XP_004829278.1 | 539 | BEWA_024610 | eukaryota>alveolata>apicomplexa | Theileria | equi strain WA | chaperone protein DNAj, putative [Theileria equi strain WA] |
| CEL99775.1 | 537 | Vbra_5321 | eukaryota>alveolata | Vitrella | brassicaformis CCMP3155 | unnamed protein product [Vitrella brassicaformis CCMP3155] |
| XP_002966705.1 | 537 | SELMODRAFT_407813 | eukaryota>viridiplantae | Selaginella | moellendorffii | hypothetical protein SELMODRAFT_407813 [Selaginella moellendorffii] |
| XP_002978034.1 | 537 | SELMODRAFT_417930 | eukaryota>viridiplantae | Selaginella | moellendorffii | hypothetical protein SELMODRAFT_417930 [Selaginella moellendorffii] |
| EJY87165.1 | 535 | OXYTRI_06273 | eukaryota>alveolata>ciliophora | Oxytricha | trifallax | DnaJ domain containing protein [Oxytricha trifallax] |
| XP_001696952.1 | 535 | DNJ3 | eukaryota>viridiplantae>chlorophyta | Chlamydomonas | reinhardtii | DnaJ-like protein [Chlamydomonas reinhardtii] |
| XP_012650419.1 | 535 | BBM_III09175 | eukaryota>alveolata>apicomplexa | Babesia | microti strain RI | uncharacterized protein [Babesia microti strain RI] |
| ELR20284.1 | 533 | ACA1_307490 | eukaryota>amoebozoa>acanthamoebidae | Acanthamoeba | castellanii str. Neff | DnaJ domain containing protein [Acanthamoeba castellanii str. Neff] |
| XP_005827189.1 | 530 | GUITHDRAFT_113690 | eukaryota>cryptophyta | Guillardia | theta CCMP2712 | hypothetical protein GUITHDRAFT_113690 [Guillardia theta CCMP2712] |
| XP_002783521.1 | 521 | Pmar_PMAR001367 | eukaryota>alveolata | Perkinsus | marinus ATCC 50983 | conserved hypothetical protein [Perkinsus marinus ATCC 50983] |
| XP_001017243.1 | 519 | TTHERM_00195890 | eukaryota>alveolata>ciliophora | Tetrahymena | thermophila SB210 | DnaJ domain protein [Tetrahymena thermophila SB210] |
| XP_004031789.1 | 518 | IMG5_138460 | eukaryota>alveolata>ciliophora | Ichthyophthirius | multifiliis | hypothetical protein IMG5_138460 [Ichthyophthirius multifiliis] |
| XP_764324.1 | 509 | TP04_0688 | eukaryota>alveolata>apicomplexa | Theileria | parva strain Muguga | heat shock protein DnaJ [Theileria parva strain Muguga] |
| XP_009692153.1 | 508 | TOT_040000232 | eukaryota>alveolata>apicomplexa | Theileria | orientalis strain Shintoku | Pbj2 [Theileria orientalis strain Shintoku] |
| OMJ76060.1 | 504 | SteCoe_24666 | eukaryota>alveolata>ciliophora | Stentor | coeruleus | hypothetical protein SteCoe_24666 [Stentor coeruleus] |
| XP_004831200.1 | 504 | BEWA_009480 | eukaryota>alveolata>apicomplexa | Theileria | equi strain WA | DnaJ domain-containing protein [Theileria equi strain WA] |
| XP_001423972.1 | 503 | GSPATT00027602001 | eukaryota>alveolata>ciliophora | Paramecium | tetraurelia strain d4-2 | hypothetical protein (macronuclear) [Paramecium tetraurelia strain d4-2] |
| XP_009060482.1 | 503 | LOTGIDRAFT_218980 | eukaryota>metazoa>mollusca | Lottia | gigantea | hypothetical protein LOTGIDRAFT_218980 [Lottia gigantea] |
| XP_005822954.1 | 503 | GUITHDRAFT_146120 | eukaryota>cryptophyta | Guillardia | theta CCMP2712 | hypothetical protein GUITHDRAFT_146120 [Guillardia theta CCMP2712] |
| XP_001430366.1 | 499 | GSPATT00032820001 | eukaryota>alveolata>ciliophora | Paramecium | tetraurelia strain d4-2 | hypothetical protein (macronuclear) [Paramecium tetraurelia strain d4-2] |
| ABO95202.1 | 495 | OSTLU_14554 | eukaryota>viridiplantae>chlorophyta | Ostreococcus | lucimarinus CCE9901 | predicted protein [Ostreococcus lucimarinus CCE9901] |
| OHT09988.1 | 494 | TRFO_20990 | eukaryota>parabasalia | Tritrichomonas | foetus | hypothetical protein TRFO_20990 [Tritrichomonas foetus] |
| XP_013900451.1 | 493 | MNEG_6529 | eukaryota>viridiplantae>chlorophyta | Monoraphidium | neglectum | protein disulfide isomerase family A, member 6 [Monoraphidium neglectum] |
| CDW76835.1 | 492 | Contig950.g1040 | eukaryota>alveolata>ciliophora | Stylonychia | lemnae | UNKNOWN [Stylonychia lemnae] |
| CBJ26316.1 | 487 | Esi_0029_0127 | eukaryota>stramenopiles | Ectocarpus | siliculosus | Heat shock protein 40 like protein [Ectocarpus siliculosus] |
| XP_002503525.1 | 487 | MICPUN_59923 | eukaryota>viridiplantae>chlorophyta | Micromonas | commoda | predicted protein [Micromonas commoda] |
| XP_002948079.1 | 485 | dnj3 | eukaryota>viridiplantae>chlorophyta | Volvox | carteri f. nagariensis | molecular chaperone [Volvox carteri f. nagariensis] |
| XP_003081576.1 | 484 | Ot10g01430 | eukaryota>viridiplantae>chlorophyta | Ostreococcus | tauri | Molecular chaperone (DnaJ superfamily) (ISS) [Ostreococcus tauri] |
| XP_004029747.1 | 481 | IMG5_173670 | eukaryota>alveolata>ciliophora | Ichthyophthirius | multifiliis | hypothetical protein IMG5_173670 [Ichthyophthirius multifiliis] |
| XP_001461982.1 | 478 | GSPATT00026933001 | eukaryota>alveolata>ciliophora | Paramecium | tetraurelia strain d4-2 | hypothetical protein (macronuclear) [Paramecium tetraurelia strain d4-2] |
| XP_005648989.1 | 476 | COCSUDRAFT_40840 | eukaryota>viridiplantae>chlorophyta | Coccomyxa | subellipsoidea C-169 | DnaJ-domain-containing protein [Coccomyxa subellipsoidea C-169] |
| XP_002908020.1 | 468 | PITG_01349 | eukaryota>stramenopiles | Phytophthora | infestans T30-4 | Type II (General) Secretory Pathway (IISP) Family [Phytophthora infestans T30-4] |
| XP_005852011.1 | 466 | CHLNCDRAFT_132941 | eukaryota>viridiplantae>chlorophyta | Chlorella | variabilis | hypothetical protein CHLNCDRAFT_132941 [Chlorella variabilis] |
| XP_012649810.1 | 462 | BBM_III06020 | eukaryota>alveolata>apicomplexa | Babesia | microti strain RI | uncharacterized protein [Babesia microti strain RI] |
| OMJ88336.1 | 456 | SteCoe_9749 | eukaryota>alveolata>ciliophora | Stentor | coeruleus | hypothetical protein SteCoe_9749 [Stentor coeruleus] |
| XP_009526141.1 | 448 | PHYSODRAFT_544780 | eukaryota>stramenopiles | Phytophthora | sojae | hypothetical protein PHYSODRAFT_544780 [Phytophthora sojae] |
| CAG02636.1 | 444 | GSTEN:00021791:G:001 | eukaryota>metazoa>chordata>vertebrata>actinopterygii | Tetraodon | nigroviridis | unnamed protein product, partial [Tetraodon nigroviridis] |
| XP_011398221.1 | 442 | F751_0866 | eukaryota>viridiplantae>chlorophyta | Auxenochlorella | protothecoides | Putative protein disulfide-isomerase [Auxenochlorella protothecoides] |
| XP_008866253.1 | 440 | H310_03946 | eukaryota>stramenopiles | Aphanomyces | invadans | hypothetical protein H310_03946 [Aphanomyces invadans] |
| XP_009823351.1 | 440 | H257_01713 | eukaryota>stramenopiles | Aphanomyces | astaci | hypothetical protein H257_01713 [Aphanomyces astaci] |
| Smed1000007201 | 430 | Smed1000007201 | eukaryota>metazoa>platyhelminthe | Schmidtea | mediterranea | >Smed1000007201 Smed1000007201 |
| XP_012203900.1 | 423 | SPRG_09418 | eukaryota>stramenopiles | Saprolegnia | parasitica CBS 223.65 | hypothetical protein SPRG_09418 [Saprolegnia parasitica CBS 223.65] |
| XP_008612210.1 | 417 | SDRG_08120 | eukaryota>stramenopiles | Saprolegnia | diclina VS20 | DnaJ like subfamily C member 7 [Saprolegnia diclina VS20] |
| XP_013233440.1 | 408 | ETH_00006695 | eukaryota>alveolata>apicomplexa | Eimeria | tenella | DnaJ domain-containing protein, putative [Eimeria tenella] |
| CEM27509.1 | 400 | Vbra_17492 | eukaryota>alveolata | Vitrella | brassicaformis CCMP3155 | unnamed protein product [Vitrella brassicaformis CCMP3155] |
| XP_001611912.1 | 387 | 17.m07688 | eukaryota>alveolata>apicomplexa | Babesia | bovis T2Bo | dnaJ domain containing protein [Babesia bovis T2Bo] |
| XP_002371338.1 | 360 | TGME49_217710 | eukaryota>alveolata>apicomplexa | Toxoplasma | gondii ME49 | DnaJ domain-containing protein [Toxoplasma gondii ME49] |
| XP_015754256.1 | 347 | LOC107333902 | eukaryota>metazoa>cnidaria | Acropora | digitifera | PREDICTED: dnaJ homolog subfamily C member 16-like [Acropora digitifera] |
| XP_003885850.1 | 340 | NCLIV_062500 | eukaryota>alveolata>apicomplexa | Neospora | caninum Liverpool | putative DnaJ domain-containing protein [Neospora caninum Liverpool] |
| XP_008889274.1 | 335 | HHA_217710 | eukaryota>alveolata>apicomplexa | Hammondia | hammondi | DnaJ domain-containing protein [Hammondia hammondi] |
| OHT01240.1 | 308 | TRFO_31983 | eukaryota>parabasalia | Tritrichomonas | foetus | hypothetical protein TRFO_31983 [Tritrichomonas foetus] |
| XP_953331.1 | 284 | TA10480 | eukaryota>alveolata>apicomplexa | Theileria | annulata | molecular chaperone (DnaJ family), putative [Theileria annulata] |
| XP_005788416.1 | 281 | EMIHUDRAFT_252473 | eukaryota>haptophyceae | Emiliania | huxleyi CCMP1516 | hypothetical protein EMIHUDRAFT_252473 [Emiliania huxleyi CCMP1516] |
| XP_001625279.1 | 249 | NEMVEDRAFT_v1g130443 | eukaryota>metazoa>cnidaria | Nematostella | vectensis | predicted protein [Nematostella vectensis] |
| ABO98241.1 | 184 | OSTLU_5385 | eukaryota>viridiplantae>chlorophyta | Ostreococcus | lucimarinus CCE9901 | predicted protein, partial [Ostreococcus lucimarinus CCE9901] |

O. Sarcalumenin/EHD

| | | | | | | |
|---|---|---|---|---|---|---|
| XP_001952603.2 | 1078 | LOC100160121 | eukaryota>metazoa>hexapoda | Acyrthosiphon | pisum | PREDICTED: sarcalumenin isoform X1 [Acyrthosiphon pisum] |
| XP_008185565.1 | 1045 | LOC100160121 | eukaryota>metazoa>hexapoda | Acyrthosiphon | pisum | PREDICTED: sarcalumenin isoform X2 [Acyrthosiphon pisum] |
| XP_008185566.1 | 1039 | LOC100160121 | eukaryota>metazoa>hexapoda | Acyrthosiphon | pisum | PREDICTED: sarcalumenin isoform X3 [Acyrthosiphon pisum] |
| XP_012352596.1 | 1033 | SRL | eukaryota>metazoa>chordata>vertebrata | Nomascus | leucogenys | PREDICTED: LOW QUALITY PROTEIN: sarcalumenin [Nomascus leucogenys] |
| EPZ35530.1 | 983 | O9G_003963 | eukaryota>fungi | Rozella | allomycis CSF55 | Phosphoacetylglucosamine mutase-like protein [Rozella allomycis CSF55] |
| XP_001863387.1 | 958 | CpipJ_CPIJ013085 | eukaryota>metazoa>hexapoda | Culex | quinquefasciatus | sarcalumenin [Culex quinquefasciatus] |
| XP_011153590.1 | 958 | LOC105191705 | eukaryota>metazoa>hexapoda | Harpegnathos | saltator | sarcalumenin isoform X1 [Harpegnathos saltator] |
| XP_011153599.1 | 956 | LOC105191705 | eukaryota>metazoa>hexapoda | Harpegnathos | saltator | PREDICTED: sarcalumenin isoform X2 [Harpegnathos saltator] |
| XP_006521750.1 | 955 | 9830004M20Rik,AW108387,sar | eukaryota>metazoa>chordata>vertebrata | Mus | musculus | PREDICTED: sarcalumenin isoform X1 [Mus musculus] |
| XP_006521751.1 | 954 | 9830004M20Rik,AW108387,sar | eukaryota>metazoa>chordata>vertebrata | Mus | musculus | PREDICTED: sarcalumenin isoform X2 [Mus musculus] |
| NP_001262553.1 | 952 | Dmel\CG9297 | eukaryota>metazoa>hexapoda | Drosophila | melanogaster | uncharacterized protein Dmel_CG9297, isoform C [Drosophila melanogaster] |
| NP_001262554.1 | 952 | Dmel\CG9297 | eukaryota>metazoa>hexapoda | Drosophila | melanogaster | uncharacterized protein Dmel_CG9297, isoform D [Drosophila melanogaster] |
| NP_001262555.1 | 952 | Dmel\CG9297 | eukaryota>metazoa>hexapoda | Drosophila | melanogaster | uncharacterized protein Dmel_CG9297, isoform E [Drosophila melanogaster] |
| NP_788674.1 | 952 | Dmel\CG9297 | eukaryota>metazoa>hexapoda | Drosophila | melanogaster | uncharacterized protein Dmel_CG9297, isoform B [Drosophila melanogaster] |
| XP_011153610.1 | 951 | LOC105191705 | eukaryota>metazoa>hexapoda | Harpegnathos | saltator | sarcalumenin isoform X2 [Harpegnathos saltator] |
| XP_005591158.1 | 932 | SRL | eukaryota>metazoa>chordata>vertebrata | Macaca | fascicularis | PREDICTED: sarcalumenin isoform X2 [Macaca fascicularis] |
| NP_788675.1 | 930 | Dmel\CG9297 | eukaryota>metazoa>hexapoda | Drosophila | melanogaster | uncharacterized protein Dmel_CG9297, isoform A [Drosophila melanogaster] |
| XP_016767580.1 | 926 | LOC413012 | eukaryota>metazoa>hexapoda | Apis | mellifera | PREDICTED: EH domain-containing protein 3 [Apis mellifera] |
| NP_780556.2 | 910 | 9830004M20Rik,AW108387,sar | eukaryota>metazoa>chordata>vertebrata | Mus | musculus | sarcalumenin isoform 1 precursor [Mus musculus] |
| XP_013787331.1 | 894 | LOC106471282 | eukaryota>metazoa | Limulus | polyphemus | sarcalumenin-like isoform X1 [Limulus polyphemus] |
| XP_016768229.1 | 889 | LOC100576703 | eukaryota>metazoa>hexapoda | Apis | mellifera | PREDICTED: sarcalumenin [Apis mellifera] |
| XP_018092020.1 | 886 | srl.L | eukaryota>metazoa>chordata>vertebrata | Xenopus | laevis | PREDICTED: sarcalumenin isoform X1 [Xenopus laevis] |
| XP_009042966.1 | 880 | AURANDRAFT_68976 | eukaryota>stramenopiles | Aureococcus | anophagefferens | hypothetical protein AURANDRAFT_68976 [Aureococcus anophagefferens] |
| XP_018092021.1 | 874 | srl.L | eukaryota>metazoa>chordata>vertebrata | Xenopus | laevis | PREDICTED: sarcalumenin isoform X2 [Xenopus laevis] |
| XP_005591160.1 | 857 | SRL | eukaryota>metazoa>chordata>vertebrata | Macaca | fascicularis | PREDICTED: sarcalumenin isoform X1 [Macaca fascicularis] |
| XP_016879016.1 | 857 | SRL | eukaryota>metazoa>chordata>vertebrata | Homo | sapiens | sarcalumenin isoform X1 [Homo sapiens] |
| XP_013787660.1 | 842 | LOC106471590 | eukaryota>metazoa | Limulus | polyphemus | sarcalumenin-like isoform X1 [Limulus polyphemus] |
| XP_969014.1 | 841 | LOC657463 | eukaryota>metazoa>hexapoda | Tribolium | castaneum | PREDICTED: sarcalumenin isoform X1 [Tribolium castaneum] |
| XP_013787667.1 | 840 | LOC106471590 | eukaryota>metazoa | Limulus | polyphemus | sarcalumenin-like isoform X2 [Limulus polyphemus] |
| NP_001024193.1 | 835 | rme-1 | eukaryota>metazoa>nematoda | Caenorhabditis | elegans | Receptor Mediated Endocytosis [Caenorhabditis elegans] |
| XP_013777210.1 | 826 | LOC106461889 | eukaryota>metazoa | Limulus | polyphemus | sarcalumenin-like [Limulus polyphemus] |
| XP_012550298.1 | 816 | LOC101744044 | eukaryota>metazoa>hexapoda | Bombyx | mori | LOW QUALITY PROTEIN: sarcalumenin [Bombyx mori] |
| XP_015833583.1 | 811 | LOC657463 | eukaryota>metazoa>hexapoda | Tribolium | castaneum | PREDICTED: sarcalumenin isoform X2 [Tribolium castaneum] |
| XP_003377409.1 | 809 | Tsp_01593 | eukaryota>metazoa>nematoda | Trichinella | spiralis | putative EH domain-containing protein 1 [Trichinella spiralis] |
| NP_001024192.1 | 786 | rme-1 | eukaryota>metazoa>nematoda | Caenorhabditis | elegans | Receptor Mediated Endocytosis [Caenorhabditis elegans] |
| XP_011610829.1 | 768 | LOC101074425 | eukaryota>metazoa>chordata>vertebrata>actinopterygii | Takifugu | rubripes | PREDICTED: sarcalumenin-like isoform X1 [Takifugu rubripes] |
| CBJ31714.1 | 762 | RME1L3 | eukaryota>stramenopiles | Ectocarpus | siliculosus | RME1L3, RME1-like GTPase/ATPase without a C-terminal EH domain [Ectocarpus siliculosus] |
| XP_011610830.1 | 748 | LOC101074425 | eukaryota>metazoa>chordata>vertebrata>actinopterygii | Takifugu | rubripes | PREDICTED: sarcalumenin-like isoform X2 [Takifugu rubripes] |
| XP_013787332.1 | 743 | LOC106471282 | eukaryota>metazoa | Limulus | polyphemus | sarcalumenin-like [Limulus polyphemus] |
| EJK64730.1 | 734 | THAOC_14506 | eukaryota>stramenopiles | Thalassiosira | oceanica | hypothetical protein THAOC_14506 [Thalassiosira oceanica] |
| EJK66267.1 | 718 | THAOC_12827 | eukaryota>stramenopiles | Thalassiosira | oceanica | hypothetical protein THAOC_12827 [Thalassiosira oceanica] |
| CBJ27330.1 | 703 | RME1L2 | eukaryota>stramenopiles | Ectocarpus | siliculosus | RME1L2, RME1-like GTPase/ATPase without a C-terminal EH domain [Ectocarpus siliculosus] |
| OLP92823.1 | 700 | EHD4 | eukaryota>alveolata | Symbiodinium | microadriaticum | EH domain-containing protein 4 [Symbiodinium microadriaticum] |
| XP_017314466.1 | 700 | LOC108259451 | eukaryota>metazoa>chordata>vertebrata>actinopterygii | Ictalurus | punctatus | PREDICTED: sarcalumenin-like [Ictalurus punctatus] |
| OLP96537.1 | 698 | Ehd4 | eukaryota>alveolata | Symbiodinium | microadriaticum | EH domain-containing protein 4 [Symbiodinium microadriaticum] |
| XP_011131356.1 | 681 | GNI_105920 | eukaryota>alveolata>apicomplexa | Gregarina | niphandrodes | putative EH-domain protein [Gregarina niphandrodes] |
| XP_813710.1 | 673 | Tc00.1047053509207.70 | eukaryota>euglenozoa>kinetoplastida | Trypanosoma | cruzi strain CL Brener | hypothetical protein [Trypanosoma cruzi strain CL Brener] |
| XP_002610295.1 | 669 | BRAFLDRAFT_126859 | eukaryota>metazoa>chordata | Branchiostoma | floridae | hypothetical protein BRAFLDRAFT_126859 [Branchiostoma floridae] |
| XP_005782411.1 | 667 | EMIHUDRAFT_449725 | eukaryota>haptophyceae | Emiliania | huxleyi CCMP1516 | hypothetical protein EMIHUDRAFT_449725 [Emiliania huxleyi CCMP1516] |
| CAF93029.1 | 648 | GSTEN:00008495:G:001 | eukaryota>metazoa>chordata>vertebrata>actinopterygii | Tetraodon | nigroviridis | unnamed protein product, partial [Tetraodon nigroviridis] |
| XP_807180.1 | 646 | Tc00.1047053508321.60 | eukaryota>euglenozoa>kinetoplastida | Trypanosoma | cruzi strain CL Brener | hypothetical protein [Trypanosoma cruzi strain CL Brener] |
| XP_005784921.1 | 641 | EMIHUDRAFT_456074 | eukaryota>haptophyceae | Emiliania | huxleyi CCMP1516 | hypothetical protein EMIHUDRAFT_456074 [Emiliania huxleyi CCMP1516] |
| KPI83560.1 | 637 | ABL78_7399 | eukaryota>euglenozoa>kinetoplastida | Leptomonas | seymouri | hypothetical protein ABL78_7399 [Leptomonas seymouri] |
| KFH65415.1 | 636 | MVEG_08893 | eukaryota>fungi | Mortierella | verticillata NRRL 6337 | hypothetical protein MVEG_08893 [Mortierella verticillata NRRL 6337] |
| XP_001682605.1 | 633 | LMJF_19_0280 | eukaryota>euglenozoa>kinetoplastida | Leishmania | major strain Friedlin | putative sarcoplasmic reticulum glycoprotein [Leishmania major strain Friedlin] |
| XP_009313934.1 | 632 | DQ04_08691020 | eukaryota>euglenozoa>kinetoplastida | Trypanosoma | grayi | hypothetical protein DQ04_08691020 [Trypanosoma grayi] |
| XP_013234175.1 | 632 | ETH_00031045 | eukaryota>alveolata>apicomplexa | Eimeria | tenella | EH protein, putative [Eimeria tenella] |
| XP_001464991.1 | 631 | LINJ_19_0270 | eukaryota>euglenozoa>kinetoplastida | Leishmania | infantum JPCM5 | putative sarcalumenin precursor [Leishmania infantum JPCM5] |
| XP_003860206.1 | 631 | LDBPK_190270 | eukaryota>euglenozoa>kinetoplastida | Leishmania | donovani | sarcoplasmic reticulum glycoprotein, putative [Leishmania donovani] |
| EWM27210.1 | 629 | RME1L4 | eukaryota>stramenopiles | Nannochloropsis | gaditana | rme1-like gtpase atpase without a c-terminal eh domain protein [Nannochloropsis gaditana] |
| XP_003874206.1 | 628 | LMXM_19_0280 | eukaryota>euglenozoa>kinetoplastida | Leishmania | mexicana MHOM/GT/2001/U1103 | putative sarcalumenin precursor [Leishmania mexicana MHOM/GT/2001/U1103] |
| XP_001564155.1 | 626 | LBRM_19_0590 | eukaryota>euglenozoa>kinetoplastida | Leishmania | braziliensis MHOM/BR/75/M2904 | putative sarcoplasmic reticulum glycoprotein [Leishmania braziliensis MHOM/BR/75/M2904] |
| XP_010698020.1 | 626 | LPMP_190250 | eukaryota>euglenozoa>kinetoplastida | Leishmania | panamensis | sarcoplasmic reticulum glycoprotein-like protein [Leishmania panamensis] |
| XP_011772821.1 | 624 | TbgDal_IV2340 | eukaryota>euglenozoa>kinetoplastida | Trypanosoma | brucei gambiense DAL972 | sarcoplasmic reticulum glycoprotein, putative [Trypanosoma brucei gambiense DAL972] |
| KFH73719.1 | 623 | MVEG_02014 | eukaryota>fungi | Mortierella | verticillata NRRL 6337 | hypothetical protein MVEG_02014 [Mortierella verticillata NRRL 6337] |
| XP_005762092.1 | 623 | RME1B | eukaryota>haptophyceae | Emiliania | huxleyi CCMP1516 | RME1B, RME-1/EHD family protein [Emiliania huxleyi CCMP1516] |
| ERG83767.1 | 614 | ASU_06468 | eukaryota>metazoa>nematoda | Ascaris | suum | eh domain-containing protein 3 [Ascaris suum] |
| NP_001024196.1 | 613 | rme-1 | eukaryota>metazoa>nematoda | Caenorhabditis | elegans | Receptor Mediated Endocytosis [Caenorhabditis elegans] |
| EJK48101.1 | 611 | THAOC_33126 | eukaryota>stramenopiles | Thalassiosira | oceanica | hypothetical protein THAOC_33126 [Thalassiosira oceanica] |
| EPY23687.1 | 611 | STCU_07553 | eukaryota>euglenozoa>kinetoplastida | Strigomonas | culicis | sarcoplasmic reticulum glycoprotein [Strigomonas culicis] |
| XP_819453.1 | 610 | Tc00.1047053506211.100 | eukaryota>euglenozoa>kinetoplastida | Trypanosoma | cruzi strain CL Brener | sarcoplasmic reticulum glycoprotein [Trypanosoma cruzi strain CL Brener] |
| XP_009009818.1 | 609 | HELRODRAFT_62236 | eukaryota>metazoa>annelida | Helobdella | robusta | hypothetical protein HELRODRAFT_62236 [Helobdella robusta] |
| XP_002286661.1 | 608 | THAPSDRAFT_20698 | eukaryota>stramenopiles | Thalassiosira | pseudonana CCMP1335 | predicted protein [Thalassiosira pseudonana CCMP1335] |
| XP_011774589.1 | 600 | TbgDal_VII2680 | eukaryota>euglenozoa>kinetoplastida | Trypanosoma | brucei gambiense DAL972 | hypothetical protein, conserved [Trypanosoma brucei gambiense DAL972] |
| XP_011402336.1 | 598 | F751_1659 | eukaryota>viridiplantae>chlorophyta | Auxenochlorella | protothecoides | EH domain-containing protein 1 [Auxenochlorella protothecoides] |
| OQV26138.1 | 597 | BV898_00260 | eukaryota>metazoa | Hypsibius | dujardini | Sarcalumenin [Hypsibius dujardini] |
| CAG09748.1 | 597 | GSTEN:00031164:G:001 | eukaryota>metazoa>chordata>vertebrata>actinopterygii | Tetraodon | nigroviridis | unnamed protein product, partial [Tetraodon nigroviridis] |
| KPI90049.1 | 597 | ABL78_0802 | eukaryota>euglenozoa>kinetoplastida | Leptomonas | seymouri | putative sarcoplasmic reticulum glycoprotein putative sarcalumenin precursor [Leptomonas seymouri] |
| XP_015661079.1 | 597 | ABB37_02479 | eukaryota>euglenozoa>kinetoplastida | Leptomonas | pyrrhocoris | putative sarcoplasmic reticulum glycoprotein putativesarcalumenin precursor [Leptomonas pyrrhocoris] |
| XP_002368022.1 | 594 | TGME49_231210 | eukaryota>alveolata>apicomplexa | Toxoplasma | gondii ME49 | sarcalumenin/eps15 family protein [Toxoplasma gondii ME49] |
| XP_008885711.1 | 594 | HHA_231210 | eukaryota>alveolata>apicomplexa | Hammondia | hammondi | sarcalumenin/eps15 family protein [Hammondia hammondi] |
| OLP95718.1 | 592 | Ehd1 | eukaryota>alveolata | Symbiodinium | microadriaticum | EH domain-containing protein 1 [Symbiodinium microadriaticum] |
| NP_001024194.1 | 589 | rme-1 | eukaryota>metazoa>nematoda | Caenorhabditis | elegans | Receptor Mediated Endocytosis [Caenorhabditis elegans] |
| CBJ27036.1 | 587 | RME1 | eukaryota>stramenopiles | Ectocarpus | siliculosus | RME1, RME-1/EHD family ATPase with a C-terminal EH domain [Ectocarpus siliculosus] |
| XP_009034850.1 | 583 | AURANDRAFT_52816 | eukaryota>stramenopiles | Aureococcus | anophagefferens | hypothetical protein AURANDRAFT_52816 [Aureococcus anophagefferens] |
| XP_003883403.1 | 582 | NCLIV_031580 | eukaryota>alveolata>apicomplexa | Neospora | caninum Liverpool | GA19392, related [Neospora caninum Liverpool] |
| XP_003382701.2 | 581 | LOC100636011 | eukaryota>metazoa | Amphimedon | queenslandica | PREDICTED: EH domain-containing protein 1-like [Amphimedon queenslandica] |
| XP_009537814.1 | 578 | PHYSODRAFT_529936 | eukaryota>stramenopiles | Phytophthora | sojae | hypothetical protein PHYSODRAFT_529936 [Phytophthora sojae] |
| XP_002112862.1 | 577 | TRIADDRAFT_56433 | eukaryota>metazoa>placozoa | Trichoplax | adhaerens | hypothetical protein TRIADDRAFT_56433 [Trichoplax adhaerens] |
| XP_002899341.1 | 577 | PITG_14253 | eukaryota>stramenopiles | Phytophthora | infestans T30-4 | conserved hypothetical protein [Phytophthora infestans T30-4] |
| NP_001024195.1 | 576 | rme-1 | eukaryota>metazoa>nematoda | Caenorhabditis | elegans | Receptor Mediated Endocytosis [Caenorhabditis elegans] |
| XP_008916119.1 | 575 | PPTG_19423 | eukaryota>stramenopiles | Phytophthora | parasitica INRA-310 | hypothetical protein PPTG_19423 [Phytophthora parasitica INRA-310] |
| Pram1000000693 | 575 | Pram1000000693 | eukaryota>stramenopiles | Phytophthora | ramorum | >Pram1000000693 Pram1000000693 |
| XP_002408077.1 | 573 | IscW_ISCW009690 | eukaryota>metazoa | Ixodes | scapularis | endocytosis/signaling protein EHD1, putative, partial [Ixodes scapularis] |
| XP_002112876.1 | 572 | TRIADDRAFT_25492 | eukaryota>metazoa>placozoa | Trichoplax | adhaerens | hypothetical protein TRIADDRAFT_25492 [Trichoplax adhaerens] |
| XP_005717519.1 | 572 | CHC_T00005843001 | eukaryota>rhodophyta | Chondrus | crispus | unnamed protein product [Chondrus crispus] |
| EJK62928.1 | 571 | THAOC_16443 | eukaryota>stramenopiles | Thalassiosira | oceanica | hypothetical protein THAOC_16443 [Thalassiosira oceanica] |
| EPY37221.1 | 570 | STCU_00076 | eukaryota>euglenozoa>kinetoplastida | Strigomonas | culicis | hypothetical protein STCU_00076 [Strigomonas culicis] |
| XP_723685.1 | 570 | PY03528 | eukaryota>alveolata>apicomplexa | Plasmodium | yoelii yoelii 17XNL | hypothetical protein PY03528 [Plasmodium yoelii yoelii 17XNL] |
| XP_012897311.1 | 569 | GSBLH_T00003164001 | eukaryota>stramenopiles | Blastocystis | hominis | uncharacterized protein [Blastocystis hominis] |
| XP_005851211.1 | 568 | CHLNCDRAFT_33853 | eukaryota>viridiplantae>chlorophyta | Chlorella | variabilis | hypothetical protein CHLNCDRAFT_33853 [Chlorella variabilis] |
| XP_014529158.1 | 568 | JH06_0753 | eukaryota>stramenopiles | Blastocystis | sp. subtype 4 | hypothetical protein JH06_0753 [Blastocystis sp. subtype 4] |
| XP_640300.1 | 568 | DDB_G0282233 | eukaryota>amoebozoa>mycetozoa>dictyosteliida | Dictyostelium | discoideum AX4 | hypothetical protein DDB_G0282233 [Dictyostelium discoideum AX4] |
| OEU13291.1 | 567 | FRACYDRAFT_226667 | eukaryota>stramenopiles | Fragilariopsis | cylindrus CCMP1102 | putative EH-domain-containing protein [Fragilariopsis cylindrus CCMP1102] |
| CDS40913.1 | 563 | EmuJ_000851900 | eukaryota>metazoa | Echinococcus | multilocularis | Receptor Mediated Endocytosis family member [Echinococcus multilocularis] |
| EUB63467.1 | 563 | EGR_01549 | eukaryota>metazoa | Echinococcus | granulosus | EH domain-containing protein [Echinococcus granulosus] |
| XP_009516092.1 | 563 | PHYSODRAFT_475086 | eukaryota>stramenopiles | Phytophthora | sojae | hypothetical protein PHYSODRAFT_475086 [Phytophthora sojae] |
| XP_005780580.1 | 562 | RME1A-1 | eukaryota>haptophyceae | Emiliania | huxleyi CCMP1516 | RME1A, RME-1/EHD family protein [Emiliania huxleyi CCMP1516] |
| XP_008895591.1 | 562 | PPTG_04974 | eukaryota>stramenopiles | Phytophthora | parasitica INRA-310 | hypothetical protein PPTG_04974 [Phytophthora parasitica INRA-310] |
| XP_015660715.1 | 561 | ABB37_03384 | eukaryota>euglenozoa>kinetoplastida | Leptomonas | pyrrhocoris | hypothetical protein ABB37_03384 [Leptomonas pyrrhocoris] |
| XP_009168139.1 | 560 | T265_04985 | eukaryota>metazoa | Opisthorchis | viverrini | hypothetical protein T265_04985 [Opisthorchis viverrini] |
| XP_001945062.1 | 559 | LOC100165475 | eukaryota>metazoa>hexapoda | Acyrthosiphon | pisum | PREDICTED: EH domain-containing protein 3 [Acyrthosiphon pisum] |
| XP_002184252.1 | 559 | PHATRDRAFT_30394 | eukaryota>stramenopiles | Phaeodactylum | tricornutum CCAP 1055/1 | predicted protein [Phaeodactylum tricornutum CCAP 1055/1] |
| XP_002906916.1 | 559 | PITG_03877 | eukaryota>stramenopiles | Phytophthora | infestans T30-4 | conserved hypothetical protein [Phytophthora infestans T30-4] |
| XP_002985697.1 | 558 | SELMODRAFT_122705 | eukaryota>viridiplantae | Selaginella | moellendorffii | hypothetical protein SELMODRAFT_122705 [Selaginella moellendorffii] |
| XP_009032070.1 | 557 | HELRODRAFT_91367 | eukaryota>metazoa>annelida | Helobdella | robusta | hypothetical protein HELRODRAFT_91367 [Helobdella robusta] |
| OMH81055.1 | 556 | AX774_g5496 | eukaryota>fungi | Zancudomyces | culisetae | EH domain-containing protein 1 [Zancudomyces culisetae] |
| XP_009492794.1 | 556 | H696_00638 | eukaryota | Fonticula | alba | hypothetical protein H696_00638 [Fonticula alba] |
| NP_001024197.1 | 555 | rme-1 | eukaryota>metazoa>nematoda | Caenorhabditis | elegans | Receptor Mediated Endocytosis [Caenorhabditis elegans] |
| XP_002131017.1 | 555 | LOC100176986 | eukaryota>metazoa>chordata | Ciona | intestinalis | PREDICTED: EH domain-containing protein 3-like [Ciona intestinalis] |
| XP_002293462.1 | 554 | THAPSDRAFT_24736 | eukaryota>stramenopiles | Thalassiosira | pseudonana CCMP1335 | receptor mediated endocytosis protein-like protein [Thalassiosira pseudonana CCMP1335] |
| XP_014767553.1 | 554 | LOC106867255 | eukaryota>metazoa>mollusca | Octopus | bimaculoides | PREDICTED: EH domain-containing protein 3-like isoform X1 [Octopus bimaculoides] |
| GAQ88779.1 | 553 | KFL_004590010 | eukaryota>viridiplantae | Klebsormidium | nitens | Endocytosis/signaling protein [Klebsormidium nitens] |
| XP_002985283.1 | 552 | SELMODRAFT_121714 | eukaryota>viridiplantae | Selaginella | moellendorffii | hypothetical protein SELMODRAFT_121714 [Selaginella moellendorffii] |
| XP_017315836.1 | 552 | LOC108260244 | eukaryota>metazoa>chordata>vertebrata>actinopterygii | Ictalurus | punctatus | PREDICTED: EH domain-containing protein 2-like [Ictalurus punctatus] |
| KPI90477.1 | 551 | ABL78_0407 | eukaryota>euglenozoa>kinetoplastida | Leptomonas | seymouri | hypothetical protein ABL78_0407 [Leptomonas seymouri] |
| XP_016510443.1 | 551 | LOC107827761 | eukaryota>viridiplantae | Nicotiana | tabacum | PREDICTED: EH domain-containing protein 1-like [Nicotiana tabacum] |
| XP_016604678.1 | 551 | SPPG_07851 | eukaryota>fungi>chytridiomycota | Spizellomyces | punctatus DAOM BR117 | hypothetical protein SPPG_07851 [Spizellomyces punctatus DAOM BR117] |
| EPY24407.1 | 550 | STCU_07196 | eukaryota>euglenozoa>kinetoplastida | Strigomonas | culicis | sarcoplasmic reticulum glycoprotein [Strigomonas culicis] |
| EXX52865.1 | 550 | RirG_249160 | eukaryota>fungi | Rhizophagus | irregularis DAOM 197198w | Pan1p [Rhizophagus irregularis DAOM 197198w] |
| OQV16329.1 | 549 | BV898_09476 | eukaryota>metazoa | Hypsibius | dujardini | EH domain-containing protein 1 [Hypsibius dujardini] |
| EPY23867.1 | 549 | AGDE_12675 | eukaryota>euglenozoa>kinetoplastida | Angomonas | deanei | sarcoplasmic reticulum glycoprotein [Angomonas deanei] |
| XP_001748021.1 | 549 | MONBRDRAFT_38103 | eukaryota>choanoflagellida | Monosiga | brevicollis MX1 | hypothetical protein [Monosiga brevicollis MX1] |
| XP_003379210.1 | 549 | Tsp_02879 | eukaryota>metazoa>nematoda | Trichinella | spiralis | EH domain-containing protein [Trichinella spiralis] |
| XP_014626129.1 | 549 | LOC100804468 | eukaryota>viridiplantae | Glycine | max | PREDICTED: EH domain-containing protein 1 isoform X1 [Glycine max] |
| XP_016435237.1 | 549 | LOC107761527 | eukaryota>viridiplantae | Nicotiana | tabacum | PREDICTED: EH domain-containing protein 1-like isoform X1 [Nicotiana tabacum] |
| XP_017321328.1 | 549 | LOC101154661 | eukaryota>metazoa>chordata>vertebrata>actinopterygii | Ictalurus | punctatus | PREDICTED: EH domain-containing protein 2 isoform X1 [Ictalurus punctatus] |
| NP_001269374.1 | 548 | H-PAST,HPAST1,PAST,PAST1 | eukaryota>metazoa>chordata>vertebrata | Homo | sapiens | EH domain-containing protein 1 isoform 2 [Homo sapiens] |
| XP_001733332.1 | 548 | EDI_092910 | eukaryota>amoebozoa>entamoebidae | Entamoeba | dispar SAW760 | EH domain-containing protein [Entamoeba dispar SAW760] |
| XP_005577417.1 | 548 | EHD1 | eukaryota>metazoa>chordata>vertebrata | Macaca | fascicularis | PREDICTED: EH domain-containing protein 1 isoform X1 [Macaca fascicularis] |
| XP_009310476.1 | 548 | DQ04_02771000 | eukaryota>euglenozoa>kinetoplastida | Trypanosoma | grayi | sarcoplasmic reticulum glycoprotein [Trypanosoma grayi] |
| XP_012353246.1 | 548 | EHD1 | eukaryota>metazoa>chordata>vertebrata | Nomascus | leucogenys | PREDICTED: EH domain-containing protein 1 isoform X1 [Nomascus leucogenys] |
| XP_016435238.1 | 548 | LOC107761527 | eukaryota>viridiplantae | Nicotiana | tabacum | PREDICTED: EH domain-containing protein 1-like isoform X2 [Nicotiana tabacum] |
| XP_016477437.1 | 548 | LOC107798924 | eukaryota>viridiplantae | Nicotiana | tabacum | PREDICTED: EH domain-containing protein 1-like isoform X1 [Nicotiana tabacum] |
| XP_016505166.1 | 548 | LOC107823084 | eukaryota>viridiplantae | Nicotiana | tabacum | PREDICTED: EH domain-containing protein 1-like isoform X1 [Nicotiana tabacum] |
| XP_018644902.1 | 548 | Smp_019100 | eukaryota>metazoa | Schistosoma | mansoni | eh domain containing/past-1-related [Schistosoma mansoni] |
| EFA79769.1 | 547 | PPL_06588 | eukaryota>amoebozoa>mycetozoa | Heterostelium | album PN500 | hypothetical protein PPL_06588 [Heterostelium album PN500] |
| NP_001140588.1 | 547 | GRMZM2G052740 | eukaryota>viridiplantae | Zea | mays | uncharacterized protein LOC100272658 [Zea mays] |
| XP_001772546.1 | 547 | PHYPADRAFT_167618 | eukaryota>viridiplantae | Physcomitrella | patens | predicted protein [Physcomitrella patens] |
| XP_006352380.1 | 547 | LOC102604422 | eukaryota>viridiplantae | Solanum | tuberosum | PREDICTED: EH domain-containing protein 1-like [Solanum tuberosum] |
| XP_012546894.1 | 547 | LOC101737204 | eukaryota>metazoa>hexapoda | Bombyx | mori | EH domain-containing protein 3 isoform X1 [Bombyx mori] |
| NP_567299.2 | 546 | ATEHD2,EPS15,homology,domain,2,T1J24.10,T1J24_10 | eukaryota>viridiplantae | Arabidopsis | thaliana | EPS15 homology domain 2 [Arabidopsis thaliana] |
| XP_001641640.1 | 546 | NEMVEDRAFT_v1g158412 | eukaryota>metazoa>cnidaria | Nematostella | vectensis | predicted protein [Nematostella vectensis] |
| XP_001951167.1 | 546 | LOC100166389 | eukaryota>metazoa>hexapoda | Acyrthosiphon | pisum | PREDICTED: EH domain-containing protein 1 [Acyrthosiphon pisum] |
| XP_003962855.1 | 546 | LOC101073507 | eukaryota>metazoa>chordata>vertebrata>actinopterygii | Takifugu | rubripes | PREDICTED: EH domain-containing protein 4-like [Takifugu rubripes] |
| XP_009174734.1 | 546 | T265_15072 | eukaryota>metazoa | Opisthorchis | viverrini | hypothetical protein T265_15072 [Opisthorchis viverrini] |
| XP_016435239.1 | 546 | LOC107761527 | eukaryota>viridiplantae | Nicotiana | tabacum | PREDICTED: EH domain-containing protein 1-like isoform X3 [Nicotiana tabacum] |
| XP_016477438.1 | 546 | LOC107798924 | eukaryota>viridiplantae | Nicotiana | tabacum | PREDICTED: EH domain-containing protein 1-like isoform X2 [Nicotiana tabacum] |
| XP_016505167.1 | 546 | LOC107823084 | eukaryota>viridiplantae | Nicotiana | tabacum | PREDICTED: EH domain-containing protein 1-like isoform X2 [Nicotiana tabacum] |
| XP_017211861.1 | 546 | ehd2a | eukaryota>metazoa>chordata>vertebrata>actinopterygii | Danio | rerio | EH domain-containing protein 2-like [Danio rerio] |
| XP_017332379.1 | 546 | LOC108270320 | eukaryota>metazoa>chordata>vertebrata>actinopterygii | Ictalurus | punctatus | PREDICTED: EH domain-containing protein 4-like [Ictalurus punctatus] |
| XP_698033.4 | 546 | ehd4 | eukaryota>metazoa>chordata>vertebrata>actinopterygii | Danio | rerio | EH domain-containing protein 4 [Danio rerio] |
| NP_001030731.1 | 545 | ATEHD1,EPS15,homology,domain,1 | eukaryota>viridiplantae | Arabidopsis | thaliana | EPS15 homology domain 1 [Arabidopsis thaliana] |
| NP_001327849.1 | 545 | ATEHD1,EPS15,homology,domain,1 | eukaryota>viridiplantae | Arabidopsis | thaliana | EPS15 homology domain 1 [Arabidopsis thaliana] |
| NP_566657.1 | 545 | ATEHD1,EPS15,homology,domain,1 | eukaryota>viridiplantae | Arabidopsis | thaliana | EPS15 homology domain 1 [Arabidopsis thaliana] |
| NP_974517.1 | 545 | ATEHD2,EPS15,homology,domain,2,T1J24.10,T1J24_10 | eukaryota>viridiplantae | Arabidopsis | thaliana | EPS15 homology domain 2 [Arabidopsis thaliana] |
| XP_002264320.1 | 545 | LOC100262940 | eukaryota>viridiplantae | Vitis | vinifera | PREDICTED: EH domain-containing protein 1 [Vitis vinifera] |
| XP_002277794.1 | 545 | LOC100241890 | eukaryota>viridiplantae | Vitis | vinifera | PREDICTED: EH domain-containing protein 1 [Vitis vinifera] |
| XP_003529656.1 | 545 | LOC100786047 | eukaryota>viridiplantae | Glycine | max | PREDICTED: EH domain-containing protein 1-like [Glycine max] |
| XP_003550477.1 | 545 | LOC100803757 | eukaryota>viridiplantae | Glycine | max | PREDICTED: EH domain-containing protein 1 [Glycine max] |
| XP_006344871.1 | 545 | LOC102589096 | eukaryota>viridiplantae | Solanum | tuberosum | PREDICTED: EH domain-containing protein 1 [Solanum tuberosum] |
| XP_006346193.1 | 545 | LOC102601806 | eukaryota>viridiplantae | Solanum | tuberosum | PREDICTED: EH domain-containing protein 1 [Solanum tuberosum] |
| XP_008618010.1 | 545 | SDRG_13692 | eukaryota>stramenopiles | Saprolegnia | diclina VS20 | hypothetical protein SDRG_13692 [Saprolegnia diclina VS20] |
| XP_015166444.1 | 545 | LOC102598398 | eukaryota>viridiplantae | Solanum | tuberosum | PREDICTED: EH domain-containing protein 1-like [Solanum tuberosum] |
| XP_016435240.1 | 545 | LOC107761527 | eukaryota>viridiplantae | Nicotiana | tabacum | PREDICTED: EH domain-containing protein 1-like isoform X4 [Nicotiana tabacum] |
| XP_016435242.1 | 545 | LOC107761527 | eukaryota>viridiplantae | Nicotiana | tabacum | PREDICTED: EH domain-containing protein 1-like isoform X5 [Nicotiana tabacum] |
| XP_821779.1 | 545 | Tc00.1047053506559.270 | eukaryota>euglenozoa>kinetoplastida | Trypanosoma | cruzi strain CL Brener | sarcoplasmic reticulum glycoprotein [Trypanosoma cruzi strain CL Brener] |
| CBY21336.1 | 544 | GSOID_T00009696001 | eukaryota>metazoa>chordata | Oikopleura | dioica | unnamed protein product [Oikopleura dioica] |
| XP_001564459.1 | 544 | LBRM_20_1770 | eukaryota>euglenozoa>kinetoplastida | Leishmania | braziliensis MHOM/BR/75/M2904 | conserved hypothetical protein [Leishmania braziliensis MHOM/BR/75/M2904] |
| XP_002970092.1 | 544 | LOC9645670 | eukaryota>viridiplantae | Selaginella | moellendorffii | EH domain-containing protein 1 isoform X1 [Selaginella moellendorffii] |
| XP_008644157.1 | 544 | c137059_1 | eukaryota>viridiplantae | Zea | mays | PREDICTED: uncharacterized protein LOC100384394 isoform X1 [Zea mays] |
| XP_008644158.1 | 544 | GRMZM2G113613 | eukaryota>viridiplantae | Zea | mays | EH domain-containing protein 1 isoform X1 [Zea mays] |
| XP_010698396.1 | 544 | LPMP_201950 | eukaryota>euglenozoa>kinetoplastida | Leishmania | panamensis | hypothetical protein LPMP_201950 [Leishmania panamensis] |
| KXJ17775.1 | 543 | EHD1 | eukaryota>metazoa>cnidaria | Exaiptasia | pallida | EH domain-containing protein 1 [Exaiptasia pallida] |
| NP_001167939.1 | 543 | GRMZM2G065012 | eukaryota>viridiplantae | Zea | mays | uncharacterized protein LOC100381653 [Zea mays] |
| XP_001270364.1 | 543 | EHD2 | eukaryota>metazoa>chordata>vertebrata | Macaca | fascicularis | EH domain-containing protein 2 [Macaca fascicularis] |
| NP_055416.2 | 543 | PAST2 | eukaryota>metazoa>chordata>vertebrata | Homo | sapiens | EH domain-containing protein 2 [Homo sapiens] |
| NP_694708.2 | 543 | BC027084,C130052H20Rik | eukaryota>metazoa>chordata>vertebrata | Mus | musculus | EH domain-containing protein 2 [Mus musculus] |
| NP_956357.1 | 543 | ehd2,fj66d02,wu:fj66d02,zgc:77671 | eukaryota>metazoa>chordata>vertebrata>actinopterygii | Danio | rerio | EH domain-containing protein 2 [Danio rerio] |
| XP_003519595.1 | 543 | LOC100804468 | eukaryota>viridiplantae | Glycine | max | PREDICTED: EH domain-containing protein 1 isoform X2 [Glycine max] |
| XP_011248884.1 | 543 | BC027084,C130052H20Rik | eukaryota>metazoa>chordata>vertebrata | Mus | musculus | PREDICTED: EH domain-containing protein 2 isoform X1 [Mus musculus] |
| XP_016452172.1 | 543 | LOC107776767 | eukaryota>viridiplantae | Nicotiana | tabacum | PREDICTED: EH domain-containing protein 1-like isoform X1 [Nicotiana tabacum] |
| XP_016452173.1 | 543 | LOC107776767 | eukaryota>viridiplantae | Nicotiana | tabacum | PREDICTED: EH domain-containing protein 1-like isoform X1 [Nicotiana tabacum] |
| XP_016452174.1 | 543 | LOC107776767 | eukaryota>viridiplantae | Nicotiana | tabacum | PREDICTED: EH domain-containing protein 1-like isoform X1 [Nicotiana tabacum] |
| XP_016452175.1 | 543 | LOC107776767 | eukaryota>viridiplantae | Nicotiana | tabacum | PREDICTED: EH domain-containing protein 1-like isoform X1 [Nicotiana tabacum] |
| XP_017321329.1 | 543 | LOC101154661 | eukaryota>metazoa>chordata>vertebrata>actinopterygii | Ictalurus | punctatus | PREDICTED: EH domain-containing protein 2 isoform X2 [Ictalurus punctatus] |
| XP_009011174.1 | 542 | HELRODRAFT_109211 | eukaryota>metazoa>annelida | Helobdella | robusta | hypothetical protein HELRODRAFT_109211 [Helobdella robusta] |
| XP_015142603.1 | 542 | LOC100859860 | eukaryota>metazoa>chordata>vertebrata | Gallus | gallus | PREDICTED: EH domain-containing protein 4 isoform X2 [Gallus gallus] |
| EFX80375.1 | 541 | DAPPUDRAFT_304007 | eukaryota>metazoa>crustacea | Daphnia | pulex | hypothetical protein DAPPUDRAFT_304007 [Daphnia pulex] |
| EPB81232.1 | 541 | HMPREF1544_12055 | eukaryota>fungi | Mucor | circinelloides f. circinelloides 1006PhL | hypothetical protein HMPREF1544_12055 [Mucor circinelloides f. circinelloides 1006PhL] |
| NP_598599.2 | 541 | 2210022F10Rik,AI197390,AI846352,AV006278,Past2 | eukaryota>metazoa>chordata>vertebrata | Mus | musculus | EH domain-containing protein 4 [Mus musculus] |
| NP_644670.1 | 541 | PAST4 | eukaryota>metazoa>chordata>vertebrata | Homo | sapiens | EH domain-containing protein 4 [Homo sapiens] |
| XP_001468576.1 | 541 | LINJ_34_2040 | eukaryota>euglenozoa>kinetoplastida | Leishmania | infantum JPCM5 | conserved hypothetical protein [Leishmania infantum JPCM5] |
| XP_001686325.1 | 541 | LMJF_34_2270 | eukaryota>euglenozoa>kinetoplastida | Leishmania | major strain Friedlin | conserved hypothetical protein [Leishmania major strain Friedlin] |
| XP_003266806.1 | 541 | EHD4 | eukaryota>metazoa>chordata>vertebrata | Nomascus | leucogenys | PREDICTED: EH domain-containing protein 4 [Nomascus leucogenys] |
| XP_003864363.1 | 541 | LDBPK_342040 | eukaryota>euglenozoa>kinetoplastida | Leishmania | donovani | hypothetical protein, conserved [Leishmania donovani] |
| XP_005559332.1 | 541 | EHD4 | eukaryota>metazoa>chordata>vertebrata | Macaca | fascicularis | PREDICTED: EH domain-containing protein 4 [Macaca fascicularis] |
| XP_006602183.1 | 541 | LOC100816958 | eukaryota>viridiplantae | Glycine | max | PREDICTED: EH domain-containing protein 1-like [Glycine max] |
| NP_731737.1 | 540 | CG6148,CG6148,dEHD,dEHD1,Dmel\CG6148,Ehd,PAST,Past-1,PAST-1,PAST1 | eukaryota>metazoa>hexapoda | Drosophila | melanogaster | putative achaete scute target 1, isoform A [Drosophila melanogaster] |
| XP_002974220.1 | 540 | LOC9658060 | eukaryota>viridiplantae | Selaginella | moellendorffii | EH domain-containing protein 1 [Selaginella moellendorffii] |
| XP_003286160.1 | 540 | DICPUDRAFT_30504 | eukaryota>amoebozoa>mycetozoa>dictyosteliida | Dictyostelium | purpureum | hypothetical protein DICPUDRAFT_30504 [Dictyostelium purpureum] |
| XP_005643284.1 | 540 | COCSUDRAFT_38543 | eukaryota>viridiplantae>chlorophyta | Coccomyxa | subellipsoidea C-169 | hypothetical protein COCSUDRAFT_38543 [Coccomyxa subellipsoidea C-169] |
| XP_790578.1 | 540 | LOC585665 | eukaryota>metazoa>echinodermata | Strongylocentrotus | purpuratus | PREDICTED: EH domain-containing protein 1 [Strongylocentrotus purpuratus] |
| XP_003547274.1 | 539 | LOC100805498 | eukaryota>viridiplantae | Glycine | max | PREDICTED: EH domain-containing protein 1-like [Glycine max] |
| XP_003878780.1 | 539 | LMXM_33_2270 | eukaryota>euglenozoa>kinetoplastida | Leishmania | mexicana MHOM/GT/2001/U1103 | conserved hypothetical protein [Leishmania mexicana MHOM/GT/2001/U1103] |
| XP_003968629.1 | 539 | ehd2 | eukaryota>metazoa>chordata>vertebrata>actinopterygii | Takifugu | rubripes | PREDICTED: EH domain-containing protein 2 [Takifugu rubripes] |
| XP_006586764.1 | 539 | LOC100809091 | eukaryota>viridiplantae | Glycine | max | PREDICTED: EH domain-containing protein 1-like [Glycine max] |
| XP_009826739.1 | 539 | H257_04064 | eukaryota>stramenopiles | Aphanomyces | astaci | hypothetical protein H257_04064 [Aphanomyces astaci] |
| XP_011606911.1 | 539 | ehd2 | eukaryota>metazoa>chordata>vertebrata>actinopterygii | Takifugu | rubripes | PREDICTED: EH domain-containing protein 2 [Takifugu rubripes] |
| Capa1000009297 | 539 | Caps1000009297 | eukaryota>metazoa>annelida | Capitella | spI | >Caps1000009297 Capa1000009297 |
| CEM32347.1 | 538 | Vbra_1035 | eukaryota>alveolata | Vitrella | brassicaformis CCMP3155 | unnamed protein product [Vitrella brassicaformis CCMP3155] |
| NP_001079813.1 | 538 | ehd2 | eukaryota>metazoa>chordata>vertebrata | Xenopus | laevis | EH domain containing 2 L homeolog [Xenopus laevis] |
| NP_001082536.1 | 538 | LOC398546 | eukaryota>metazoa>chordata>vertebrata | Xenopus | laevis | EH domain-containing protein 2-like [Xenopus laevis] |
| XP_002107982.1 | 538 | TRIADDRAFT_18691 | eukaryota>metazoa>placozoa | Trichoplax | adhaerens | hypothetical protein TRIADDRAFT_18691 [Trichoplax adhaerens] |
| XP_002403083.1 | 538 | IscW_ISCW024633 | eukaryota>metazoa | Ixodes | scapularis | endocytosis/signaling protein EHD1, putative [Ixodes scapularis] |
| XP_018295848.1 | 538 | PHYBLDRAFT_122594 | eukaryota>fungi | Phycomyces | blakesleeanus NRRL 1555(-) | hypothetical protein PHYBLDRAFT_122594 [Phycomyces blakesleeanus NRRL 1555(-)] |
| Uram1000008116 | 538 | Uram1000008116 | eukaryota>fungi>mucoromycotina | Umbelopsis | ramanniana | >Uram1000008116 Uram1000008116 |
| NP_001080707.1 | 537 | ehd4 | eukaryota>metazoa>chordata>vertebrata | Xenopus | laevis | EH domain containing 4 L homeolog [Xenopus laevis] |
| XP_013773832.1 | 537 | LOC106458828 | eukaryota>metazoa | Limulus | polyphemus | EH domain-containing protein 3-like [Limulus polyphemus] |
| XP_013776303.1 | 537 | LOC106461062 | eukaryota>metazoa | Limulus | polyphemus | EH domain-containing protein 3-like [Limulus polyphemus] |
| XP_013781142.1 | 537 | LOC106465459 | eukaryota>metazoa | Limulus | polyphemus | EH domain-containing protein 3-like [Limulus polyphemus] |
| XP_018087753.1 | 537 | LOC108699742 | eukaryota>metazoa>chordata>vertebrata | Xenopus | laevis | PREDICTED: EH domain-containing protein 4 [Xenopus laevis] |
| Lhya1000000989 | 537 | Lhya1000000989 | eukaryota>fungi>mucoromycotina | Lichtheimia | hyalospora | >Lhya1000000989 Lhya1000000989 |
| Smar1000010913 | 537 | Smar1000010913 | eukaryota>metazoa>arthropoda>myriapoda | Strigamia | maritima | >Smar1000010913 Smar1000010913 |
| EIE85679.1 | 536 | RO3G_10389 | eukaryota>fungi | Rhizopus | delemar RA 99-880 | hypothetical protein RO3G_10389 [Rhizopus delemar RA 99-880] |
| XP_001850207.1 | 536 | CpipJ_CPIJ008639 | eukaryota>metazoa>hexapoda | Culex | quinquefasciatus | past-1 [Culex quinquefasciatus] |
| XP_002736717.1 | 536 | LOC100366667 | eukaryota>metazoa>hemichordata | Saccoglossus | kowalevskii | PREDICTED: EH domain-containing protein 1-like [Saccoglossus kowalevskii] |
| XP_011448866.1 | 536 | LOC105343266 | eukaryota>metazoa>mollusca | Crassostrea | gigas | PREDICTED: EH domain-containing protein 3 [Crassostrea gigas] |
| XP_014767554.1 | 536 | LOC106867255 | eukaryota>metazoa>mollusca | Octopus | bimaculoides | PREDICTED: EH domain-containing protein 3-like isoform X2 [Octopus bimaculoides] |
| Bcir1000011263 | 536 | Bcir1000011263 | eukaryota>fungi>mucoromycotina | Backusella | circina | >Bcir1000011263 Bcir1000011263 |
| NP_001026211.1 | 535 | EHD3 | eukaryota>metazoa>chordata>vertebrata | Gallus | gallus | EH domain-containing protein 3 [Gallus gallus] |
| NP_001038469.1 | 535 | si:ch211-147c1.2 | eukaryota>metazoa>chordata>vertebrata>actinopterygii | Danio | rerio | EH domain-containing protein 3 [Danio rerio] |
| NP_055415.1 | 535 | PAST3 | eukaryota>metazoa>chordata>vertebrata | Homo | sapiens | EH domain-containing protein 3 [Homo sapiens] |
| NP_065603.2 | 535 | Ehd2 | eukaryota>metazoa>chordata>vertebrata | Mus | musculus | EH domain-containing protein 3 [Mus musculus] |
| XP_002112863.1 | 535 | TRIADDRAFT_26061 | eukaryota>metazoa>placozoa | Trichoplax | adhaerens | expressed hypothetical protein [Trichoplax adhaerens] |
| XP_003262753.1 | 535 | EHD3 | eukaryota>metazoa>chordata>vertebrata | Nomascus | leucogenys | PREDICTED: LOW QUALITY PROTEIN: EH domain-containing protein 3 [Nomascus leucogenys] |
| XP_003971602.1 | 535 | LOC101071455 | eukaryota>metazoa>chordata>vertebrata>actinopterygii | Takifugu | rubripes | PREDICTED: EH domain-containing protein 3 [Takifugu rubripes] |
| XP_005576248.1 | 535 | EHD3 | eukaryota>metazoa>chordata>vertebrata | Macaca | fascicularis | PREDICTED: EH domain-containing protein 3 [Macaca fascicularis] |
| XP_011610269.1 | 535 | LOC101071455 | eukaryota>metazoa>chordata>vertebrata>actinopterygii | Takifugu | rubripes | PREDICTED: EH domain-containing protein 3 [Takifugu rubripes] |
| XP_013905406.1 | 535 | MNEG_1562 | eukaryota>viridiplantae>chlorophyta | Monoraphidium | neglectum | EH domain-containing protein 4 [Monoraphidium neglectum] |
| XP_014680692.1 | 535 | LOC106820696 | eukaryota>metazoa | Priapulus | caudatus | PREDICTED: EH domain-containing protein 3-like [Priapulus caudatus] |
| XP_017313237.1 | 535 | LOC108258807 | eukaryota>metazoa>chordata>vertebrata>actinopterygii | Ictalurus | punctatus | PREDICTED: EH domain-containing protein 3 [Ictalurus punctatus] |
| XP_017313246.1 | 535 | LOC108258807 | eukaryota>metazoa>chordata>vertebrata>actinopterygii | Ictalurus | punctatus | PREDICTED: EH domain-containing protein 3 [Ictalurus punctatus] |
| XP_017344465.1 | 535 | LOC108276895 | eukaryota>metazoa>chordata>vertebrata>actinopterygii | Ictalurus | punctatus | PREDICTED: EH domain-containing protein 1-like [Ictalurus punctatus] |
| XP_018120414.1 | 535 | LOC108717680 | eukaryota>metazoa>chordata>vertebrata | Xenopus | laevis | PREDICTED: EH domain-containing protein 3 [Xenopus laevis] |
| ELR14590.1 | 534 | ACA1_271250 | eukaryota>amoebozoa>acanthamoebidae | Acanthamoeba | castellanii str. Neff | hypothetical protein ACA1_271250 [Acanthamoeba castellanii str. Neff] |
| EPB88290.1 | 534 | HMPREF1544_04874 | eukaryota>fungi | Mucor | circinelloides f. circinelloides 1006PhL | hypothetical protein HMPREF1544_04874 [Mucor circinelloides f. circinelloides 1006PhL] |
| NP_001088212.1 | 534 | ehd1,ehd1.S | eukaryota>metazoa>chordata>vertebrata | Xenopus | laevis | EH domain containing 1 L homeolog [Xenopus laevis] |
| NP_001269373.1 | 534 | H-PAST,HPAST1,PAST,PAST1 | eukaryota>metazoa>chordata>vertebrata | Homo | sapiens | EH domain-containing protein 1 isoform 1 [Homo sapiens] |
| NP_006786.2 | 534 | H-PAST,HPAST1,PAST,PAST1 | eukaryota>metazoa>chordata>vertebrata | Homo | sapiens | EH domain-containing protein 1 isoform 1 [Homo sapiens] |
| NP_034249.1 | 534 | AA409636,Past1,RME-1 | eukaryota>metazoa>chordata>vertebrata | Mus | musculus | EH domain-containing protein 1 [Mus musculus] |
| NP_524332.2 | 534 | CG6148,CG6148,dEHD,dEHD1,Dmel\CG6148,Ehd,PAST,Past-1,PAST-1,PAST1 | eukaryota>metazoa>hexapoda | Drosophila | melanogaster | putative achaete scute target 1, isoform B [Drosophila melanogaster] |
| XP_003274216.1 | 534 | EHD1 | eukaryota>metazoa>chordata>vertebrata | Nomascus | leucogenys | PREDICTED: EH domain-containing protein 1 isoform X2 [Nomascus leucogenys] |
| XP_004365433.1 | 534 | CAOG_00562 | eukaryota>ichthyosporea | Capsaspora | owczarzaki ATCC 30864 | hypothetical protein CAOG_00562 [Capsaspora owczarzaki ATCC 30864] |
| XP_009064815.1 | 534 | LOTGIDRAFT_196647 | eukaryota>metazoa>mollusca | Lottia | gigantea | hypothetical protein LOTGIDRAFT_196647 [Lottia gigantea] |
| XP_011152664.1 | 534 | LOC105191158 | eukaryota>metazoa>hexapoda | Harpegnathos | saltator | EH domain-containing protein 3 [Harpegnathos saltator] |
| XP_011152665.1 | 534 | LOC105191158 | eukaryota>metazoa>hexapoda | Harpegnathos | saltator | EH domain-containing protein 3 [Harpegnathos saltator] |
| XP_011152666.1 | 534 | LOC105191158 | eukaryota>metazoa>hexapoda | Harpegnathos | saltator | EH domain-containing protein 3 [Harpegnathos saltator] |
| XP_011152667.1 | 534 | LOC105191158 | eukaryota>metazoa>hexapoda | Harpegnathos | saltator | EH domain-containing protein 3 [Harpegnathos saltator] |
| XP_011543041.1 | 534 | H-PAST,HPAST1,PAST,PAST1 | eukaryota>metazoa>chordata>vertebrata | Homo | sapiens | EH domain-containing protein 1 isoform X1 [Homo sapiens] |
| XP_012756732.1 | 534 | SAMD00019534_031450 | eukaryota>amoebozoa>mycetozoa>dictyosteliida | Acytostelium | subglobosum LB1 | hypothetical protein SAMD00019534_031450, partial [Acytostelium subglobosum LB1] |
| XP_015289987.1 | 534 | EHD1 | eukaryota>metazoa>chordata>vertebrata | Macaca | fascicularis | PREDICTED: EH domain-containing protein 1 isoform X2 [Macaca fascicularis] |
| XP_017329712.1 | 534 | LOC108268895 | eukaryota>metazoa>chordata>vertebrata>actinopterygii | Ictalurus | punctatus | PREDICTED: EH domain-containing protein 1-like [Ictalurus punctatus] |
| XP_018115558.1 | 534 | LOC495107 | eukaryota>metazoa>chordata>vertebrata | Xenopus | laevis | PREDICTED: EH domain-containing protein 1 [Xenopus laevis] |
| XP_018118164.1 | 534 | ehd3.S | eukaryota>metazoa>chordata>vertebrata | Xenopus | laevis | PREDICTED: EH domain-containing protein 3 [Xenopus laevis] |
| XP_975287.1 | 534 | LOC664181 | eukaryota>metazoa>hexapoda | Tribolium | castaneum | PREDICTED: EH domain-containing protein 1 [Tribolium castaneum] |
| Lhya1000004186 | 534 | Lhya1000004186 | eukaryota>fungi>mucoromycotina | Lichtheimia | hyalospora | >Lhya1000004186 Lhya1000004186 |
| Smed1000006299 | 534 | Smed1000006299 | eukaryota>metazoa>platyhelminthe | Schmidtea | mediterranea | >Smed1000006299 Smed1000006299 |
| CBJ31873.1 | 533 | RME1L4 | eukaryota>stramenopiles | Ectocarpus | siliculosus | RME1L4, RME1-like GTPase/ATPase without a C-terminal EH domain [Ectocarpus siliculosus] |
| EIE86499.1 | 533 | RO3G_11210 | eukaryota>fungi | Rhizopus | delemar RA 99-880 | hypothetical protein RO3G_11210 [Rhizopus delemar RA 99-880] |
| EPY39116.1 | 533 | AGDE_04813 | eukaryota>euglenozoa>kinetoplastida | Angomonas | deanei | sarcoplasmic reticulum glycoprotein [Angomonas deanei] |
| NP_998039.1 | 533 | zgc:77049 | eukaryota>metazoa>chordata>vertebrata>actinopterygii | Danio | rerio | EH-domain containing 1a [Danio rerio] |
| XP_001351117.1 | 533 | PF3D7_0304200 | eukaryota>alveolata>apicomplexa | Plasmodium | falciparum 3D7 | EH domain-containing protein [Plasmodium falciparum 3D7] |
| XP_003970287.1 | 533 | LOC101062556 | eukaryota>metazoa>chordata>vertebrata>actinopterygii | Takifugu | rubripes | PREDICTED: EH domain-containing protein 1-like [Takifugu rubripes] |
| XP_009289526.1 | 533 | zgc:77049 | eukaryota>metazoa>chordata>vertebrata>actinopterygii | Danio | rerio | EH-domain containing 1a isoform X1 [Danio rerio] |
| XP_009313222.1 | 533 | DQ04_06941000 | eukaryota>euglenozoa>kinetoplastida | Trypanosoma | grayi | sarcoplasmic reticulum glycoprotein [Trypanosoma grayi] |
| XP_812028.1 | 533 | Tc00.1047053508895.30 | eukaryota>euglenozoa>kinetoplastida | Trypanosoma | cruzi strain CL Brener | sarcoplasmic reticulum glycoprotein [Trypanosoma cruzi strain CL Brener] |
| NP_001004578.1 | 532 | cb862,ehd1,ehd3,ehd3l,wu:fb95h05,zgc:92331 | eukaryota>metazoa>chordata>vertebrata>actinopterygii | Danio | rerio | EH domain-containing protein 1 [Danio rerio] |
| XP_004927463.1 | 532 | LOC101737204 | eukaryota>metazoa>hexapoda | Bombyx | mori | EH domain-containing protein 3 isoform X2 [Bombyx mori] |
| XP_008869163.1 | 532 | H310_06040 | eukaryota>stramenopiles | Aphanomyces | invadans | hypothetical protein H310_06040 [Aphanomyces invadans] |
| XP_013406368.1 | 532 | LOC106170884 | eukaryota>metazoa | Lingula | anatina | EH domain-containing protein 3 [Lingula anatina] |
| XP_966867.1 | 532 | LOC660971 | eukaryota>metazoa>hexapoda | Tribolium | castaneum | PREDICTED: EH domain-containing protein 3-like [Tribolium castaneum] |
| Crev1000004822 | 532 | Crev1000004822 | eukaryota>fungi>kickxellomycotina | Coemansia | reversa | >Crev1000004822 Crev1000004822 |
| XP_009534169.1 | 531 | PHYSODRAFT_304910 | eukaryota>stramenopiles | Phytophthora | sojae | hypothetical protein PHYSODRAFT_304910 [Phytophthora sojae] |
| Bcir1000010377 | 530 | Bcir1000010377 | eukaryota>fungi>mucoromycotina | Backusella | circina | >Bcir1000010377 Bcir1000010377 |
| XP_001613064.1 | 529 | PVX_119335 | eukaryota>alveolata>apicomplexa | Plasmodium | vivax | EH (for Eps15 Homology) domain containing protein [Plasmodium vivax] |

| | | | | | | |
|---|---|---|---|---|---|---|
| Uram1000008023 | 529 | Uram1000008023 | eukaryota>fungi>mucoromycotina | Umbelopsis | ramanniana | >Uram1000008023 Uram1000008023 |
| XP_008620887.1 | 528 | SDRG_16454 | eukaryota>stramenopiles | Saprolegnia | diclina VS20 | hypothetical protein SDRG_16454 [Saprolegnia diclina VS20] |
| XP_012205107.1 | 528 | SPRG_10591 | eukaryota>stramenopiles | Saprolegnia | parasitica CBS 223.65 | hypothetical protein SPRG_10591 [Saprolegnia parasitica CBS 223.65] |
| Pram1000000294 | 528 | Pram1000000294 | eukaryota>stramenopiles | Phytophthora | ramorum | >Pram1000000294 Pram1000000294 |
| CAG09040.1 | 527 | GSTEN:00030205:G:001 | eukaryota>metazoa>chordata>vertebrata>actinopterygii | Tetraodon | nigroviridis | unnamed protein product, partial [Tetraodon nigroviridis] |
| XP_002897239.1 | 527 | PITG_17035 | eukaryota>stramenopiles | Phytophthora | infestans T30-4 | conserved hypothetical protein [Phytophthora infestans T30-4] |
| XP_008914002.1 | 527 | PPTG_17882 | eukaryota>stramenopiles | Phytophthora | parasitica INRA-310 | hypothetical protein PPTG_17882 [Phytophthora parasitica INRA-310] |
| XP_003055188.1 | 526 | MICPUCDRAFT_31096 | eukaryota>viridiplantae>chlorophyta | Micromonas | pusilla CCMP1545 | predicted protein [Micromonas pusilla CCMP1545] |
| XP_002785185.1 | 525 | Pmar_PMAR008147 | eukaryota>alveolata | Perkinsus | marinus ATCC 50983 | EH-domain-containing protein, putative [Perkinsus marinus ATCC 50983] |
| XP_013758078.1 | 525 | AMSG_05001 | eukaryota>apusozoa>apusomonadidae | Thecamonas | trahens ATCC 50062 | EH domain-containing protein 3 [Thecamonas trahens ATCC 50062] |
| XP_018299378.1 | 525 | PHYBLDRAFT_120878 | eukaryota>fungi | Phycomyces | blakesleeanus NRRL 1555(-) | hypothetical protein PHYBLDRAFT_120878 [Phycomyces blakesleeanus NRRL 1555(-)] |
| Bcir1000003721 | 524 | Bcir1000003721 | eukaryota>fungi>mucoromycotina | Backusella | circina | >Bcir1000003721 Bcir1000003721 |
| Pram1000015057 | 524 | Pram1000015057 | eukaryota>stramenopiles | Phytophthora | ramorum | >Pram1000015057 Pram1000015057 |
| XP_003369002.1 | 523 | Tsp_04432 | eukaryota>metazoa>nematoda | Trichinella | spiralis | putative EH domain-containing protein 3, partial [Trichinella spiralis] |
| XP_001784382.1 | 522 | PHYPADRAFT_153849 | eukaryota>viridiplantae | Physcomitrella | patens | predicted protein [Physcomitrella patens] |
| XP_012758189.1 | 520 | SAMD00019534_021440 | eukaryota>amoebozoa>mycetozoa>dictyosteliida | Acytostelium | subglobosum LB1 | hypothetical protein SAMD00019534_021440, partial [Acytostelium subglobosum LB1] |
| OQV22217.1 | 519 | BV898_03719 | eukaryota>metazoa | Hypsibius | dujardini | hypothetical protein BV898_03719 [Hypsibius dujardini] |
| XP_008860576.1 | 517 | ENU1_210180 | eukaryota>amoebozoa>entamoebidae | Entamoeba | nuttalli P19 | ENTH domain containing protein, putative [Entamoeba nuttalli P19] |
| XP_008861213.1 | 517 | H310_00281 | eukaryota>stramenopiles | Aphanomyces | invadans | hypothetical protein H310_00281 [Aphanomyces invadans] |
| XP_011778973.1 | 517 | TbgDal_X18120 | eukaryota>euglenozoa>kinetoplastida | Trypanosoma | brucei gambiense DAL972 | sarcalumenin precursor, putative [Trypanosoma brucei gambiense DAL972] |
| XP_012203446.1 | 517 | SPRG_08827 | eukaryota>stramenopiles | Saprolegnia | parasitica CBS 223.65 | hypothetical protein SPRG_08827 [Saprolegnia parasitica CBS 223.65] |
| XP_013239309.1 | 516 | DI09_12p50 | eukaryota>fungi>microsporidia | Mitosporidium | daphniae | hypothetical protein DI09_12p50 [Mitosporidium daphniae] |
| XP_015664083.1 | 516 | ABB37_00043 | eukaryota>euglenozoa>kinetoplastida | Leptomonas | pyrrhocoris | putative mitochondrial hypothetical protein [Leptomonas pyrrhocoris] |
| XP_015664084.1 | 516 | ABB37_00043 | eukaryota>euglenozoa>kinetoplastida | Leptomonas | pyrrhocoris | putative mitochondrial hypothetical protein [Leptomonas pyrrhocoris] |
| Smar1000012255 | 516 | Smar1000012255 | eukaryota>metazoa>arthropoda>myriapoda | Strigamia | maritima | >Smar1000012255 Smar1000012255 |
| CAG01795.1 | 515 | GSTEN:00020682:G:001 | eukaryota>metazoa>chordata>vertebrata>actinopterygii | Tetraodon | nigroviridis | unnamed protein product, partial [Tetraodon nigroviridis] |
| XP_009040221.1 | 514 | AURANDRAFT_31432 | eukaryota>stramenopiles | Aureococcus | anophagefferens | hypothetical protein AURANDRAFT_31432 [Aureococcus anophagefferens] |
| KFH65080.1 | 511 | MVEG_08561 | eukaryota>fungi | Mortierella | verticillata NRRL 6337 | hypothetical protein MVEG_08561 [Mortierella verticillata NRRL 6337] |
| XP_008870165.1 | 509 | H310_06767 | eukaryota>stramenopiles | Aphanomyces | invadans | hypothetical protein H310_06767 [Aphanomyces invadans] |
| XP_009822927.1 | 509 | H257_01428 | eukaryota>stramenopiles | Aphanomyces | astaci | hypothetical protein H257_01428 [Aphanomyces astaci] |
| GAT91521.1 | 508 | CL6EHI_152680 | eukaryota>amoebozoa>entamoebidae | Entamoeba | histolytica | EH-domain containing protein putative [Entamoeba histolytica] |
| XP_006683041.1 | 508 | BATDEDRAFT_14976 | eukaryota>fungi>chytridiomycota | Batrachochytrium | dendrobatidis JAM81 | hypothetical protein BATDEDRAFT_14976 [Batrachochytrium dendrobatidis JAM81] |
| XP_018650317.1 | 508 | Smp_065180 | eukaryota>metazoa | Schistosoma | mansoni | eh domain containing/past-1-related [Schistosoma mansoni] |
| GAT94904.1 | 507 | CL6EHI_105270 | eukaryota>amoebozoa>entamoebidae | Entamoeba | histolytica | receptor mediated endocytosis protein putative [Entamoeba histolytica] |
| GAT95987.1 | 507 | CL6EHI_052870 | eukaryota>amoebozoa>entamoebidae | Entamoeba | histolytica | enth domain protein putative [Entamoeba histolytica] |
| XP_001734264.1 | 507 | EDI_210720 | eukaryota>amoebozoa>entamoebidae | Entamoeba | dispar SAW760 | EH domain-containing protein [Entamoeba dispar SAW760] |
| XP_001735333.1 | 507 | EDI_315860 | eukaryota>amoebozoa>entamoebidae | Entamoeba | dispar SAW760 | EH domain-containing protein [Entamoeba dispar SAW760] |
| XP_004183208.1 | 507 | EIN_198150 | eukaryota>amoebozoa>entamoebidae | Entamoeba | invadens IP1 | EH domain containing protein, putative [Entamoeba invadens IP1] |
| XP_004261439.1 | 507 | EIN_499380 | eukaryota>amoebozoa>entamoebidae | Entamoeba | invadens IP1 | EH domain containing protein, putative [Entamoeba invadens IP1] |
| XP_004261078.1 | 505 | EIN_130360 | eukaryota>amoebozoa>entamoebidae | Entamoeba | invadens IP1 | EH domain containing protein, putative [Entamoeba invadens IP1] |
| XP_005841179.1 | 503 | GUITHDRAFT_160834 | eukaryota>cryptophyta | Guillardia | theta CCMP2712 | hypothetical protein GUITHDRAFT_160834 [Guillardia theta CCMP2712] |
| ELR12122.1 | 502 | ACA1_139840 | eukaryota>amoebozoa>acanthamoebidae | Acanthamoeba | castellanii str. Neff | receptor mediated endocytosis family member (rme1), putative [Acanthamoeba castellanii str. Neff] |
| XP_009830450.1 | 501 | H257_06788 | eukaryota>stramenopiles | Aphanomyces | astaci | hypothetical protein H257_06788 [Aphanomyces astaci] |
| XP_008861214.1 | 500 | H310_00281 | eukaryota>stramenopiles | Aphanomyces | invadans | hypothetical protein, variant [Aphanomyces invadans] |
| XP_008615202.1 | 498 | SDRG_10963 | eukaryota>stramenopiles | Saprolegnia | diclina VS20 | hypothetical protein SDRG_10963 [Saprolegnia diclina VS20] |
| XP_012196138.1 | 498 | SPRG_02197 | eukaryota>stramenopiles | Saprolegnia | parasitica CBS 223.65 | hypothetical protein SPRG_02197 [Saprolegnia parasitica CBS 223.65] |
| XP_003641397.2 | 494 | EHD4 | eukaryota>metazoa>chordata>vertebrata | Gallus | gallus | EH domain-containing protein 4 isoform X2 [Gallus gallus] |
| XP_001564962.1 | 493 | LBRM_22_0430 | eukaryota>euglenozoa>kinetoplastida | Leishmania | braziliensis MHOM/BR/75/M2904 | conserved hypothetical protein [Leishmania braziliensis MHOM/BR/75/M2904] |
| XP_010699047.1 | 493 | LPMP_220460 | eukaryota>euglenozoa>kinetoplastida | Leishmania | panamensis | hypothetical protein LPMP_220460 [Leishmania panamensis] |
| XP_017317733.1 | 490 | LOC108261333 | eukaryota>metazoa>chordata>vertebrata>actinopterygii | Ictalurus | punctatus | PREDICTED: sarcalumenin-like [Ictalurus punctatus] |
| XP_014668558.1 | 489 | LOC106809844 | eukaryota>metazoa | Priapulus | caudatus | PREDICTED: LOW QUALITY PROTEIN: EH domain-containing protein 3-like [Priapulus caudatus] |
| EPY29863.1 | 487 | AGDE_09748 | eukaryota>euglenozoa>kinetoplastida | Angomonas | deanei | hypothetical protein AGDE_09748 [Angomonas deanei] |
| EPY41112.1 | 487 | AGDE_02813 | eukaryota>euglenozoa>kinetoplastida | Angomonas | deanei | hypothetical protein AGDE_02813 [Angomonas deanei] |
| XP_018094740.1 | 484 | LOC446267 | eukaryota>metazoa>chordata>vertebrata | Xenopus | laevis | PREDICTED: sarcalumenin isoform X1 [Xenopus laevis] |
| CBJ30463.1 | 483 | RME1L1 | eukaryota>stramenopiles | Ectocarpus | siliculosus | RME1L1, RME1-like GTPase/ATPase without a C-terminal EH domain [Ectocarpus siliculosus] |
| CAG00163.1 | 482 | GSTEN:00018510:G:001 | eukaryota>metazoa>chordata>vertebrata>actinopterygii | Tetraodon | nigroviridis | unnamed protein product, partial [Tetraodon nigroviridis] |
| NP_001070999.1 | 482 | im:6899248,wu:fb94b10,zgc:153844 | eukaryota>metazoa>chordata>vertebrata>actinopterygii | Danio | rerio | sarcalumenin precursor [Danio rerio] |
| XP_003964691.1 | 482 | srl | eukaryota>metazoa>chordata>vertebrata>actinopterygii | Takifugu | rubripes | PREDICTED: sarcalumenin isoform X1 [Takifugu rubripes] |
| XP_013413281.1 | 482 | LOC106175703 | eukaryota>metazoa | Lingula | anatina | sarcalumenin [Lingula anatina] |
| XP_006823804.1 | 477 | LOC102801464 | eukaryota>metazoa>hemichordata | Saccoglossus | kowalevskii | PREDICTED: sarcalumenin-like [Saccoglossus kowalevskii] |
| XP_002946165.1 | 476 | VOLCADRAFT_55400 | eukaryota>viridiplantae>chlorophyta | Volvox | carteri f. nagariensis | hypothetical protein VOLCADRAFT_55400 [Volvox carteri f. nagariensis] |
| NP_001092284.1 | 473 | SAR | eukaryota>metazoa>chordata>vertebrata | Homo | sapiens | sarcalumenin isoform 1 precursor [Homo sapiens] |
| XP_002772566.1 | 473 | Pmar_PMAR021539 | eukaryota>alveolata | Perkinsus | marinus ATCC 50983 | EH-domain-containing protein, putative [Perkinsus marinus ATCC 50983] |
| XP_017172333.1 | 473 | 9830004M20Rik,AW108387,sar | eukaryota>metazoa>chordata>vertebrata | Mus | musculus | PREDICTED: sarcalumenin isoform X3 [Mus musculus] |
| XP_017172334.1 | 472 | 9830004M20Rik,AW108387,sar | eukaryota>metazoa>chordata>vertebrata | Mus | musculus | PREDICTED: sarcalumenin isoform X4 [Mus musculus] |
| XP_018092022.1 | 472 | srl.L | eukaryota>metazoa>chordata>vertebrata | Xenopus | laevis | PREDICTED: sarcalumenin isoform X3 [Xenopus laevis] |
| XP_018094741.1 | 472 | LOC446267 | eukaryota>metazoa>chordata>vertebrata | Xenopus | laevis | PREDICTED: sarcalumenin isoform X2 [Xenopus laevis] |
| NP_990660.1 | 471 | SRL | eukaryota>metazoa>chordata>vertebrata | Gallus | gallus | sarcalumenin precursor [Gallus gallus] |
| OEU18895.1 | 471 | FRACYDRAFT_183357 | eukaryota>stramenopiles | Fragilariopsis | cylindrus CCMP1102 | hypothetical protein FRACYDRAFT_183357 [Fragilariopsis cylindrus CCMP1102] |
| XP_003964690.1 | 471 | srl | eukaryota>metazoa>chordata>vertebrata>actinopterygii | Takifugu | rubripes | PREDICTED: sarcalumenin isoform X2 [Takifugu rubripes] |
| XP_009298079.1 | 470 | im:6899248,wu:fb94b10,zgc:153844 | eukaryota>metazoa>chordata>vertebrata>actinopterygii | Danio | rerio | sarcalumenin isoform X1 [Danio rerio] |
| XP_016452176.1 | 467 | LOC107776767 | eukaryota>viridiplantae | Nicotiana | tabacum | PREDICTED: EH domain-containing protein 1-like isoform X2 [Nicotiana tabacum] |
| XP_017208913.1 | 466 | wu:fb80c09 | eukaryota>metazoa>chordata>vertebrata>actinopterygii | Danio | rerio | PREDICTED: sarcalumenin, partial [Danio rerio] |
| XP_009857606.1 | 460 | LOC104265388 | eukaryota>metazoa>chordata | Ciona | intestinalis | PREDICTED: sarcalumenin-like [Ciona intestinalis] |
| XP_014526523.1 | 458 | JH06_3112 | eukaryota>stramenopiles | Blastocystis | sp. subtype 4 | hypothetical protein JH06_3112 [Blastocystis sp. subtype 4] |
| XP_016877589.1 | 453 | PAST4 | eukaryota>metazoa>chordata>vertebrata | Homo | sapiens | PREDICTED: EH domain-containing protein 4 isoform X1 [Homo sapiens] |
| XP_017174841.1 | 453 | 2210022F10Rik,AI197390,AI846352,AV006278,Past2 | eukaryota>metazoa>chordata>vertebrata | Mus | musculus | PREDICTED: EH domain-containing protein 4 isoform X1 [Mus musculus] |
| XP_012895831.1 | 449 | GSBLH_T00001899001 | eukaryota>stramenopiles | Blastocystis | hominis | uncharacterized protein [Blastocystis hominis] |
| XP_009014569.1 | 446 | HELRODRAFT_76627 | eukaryota>metazoa>annelida | Helobdella | robusta | hypothetical protein HELRODRAFT_76627, partial [Helobdella robusta] |
| XP_001465567.1 | 444 | LINJ_22_0360 | eukaryota>euglenozoa>kinetoplastida | Leishmania | infantum JPCM5 | conserved hypothetical protein [Leishmania infantum JPCM5] |
| XP_001683200.1 | 444 | LMJF_22_0480 | eukaryota>euglenozoa>kinetoplastida | Leishmania | major strain Friedlin | conserved hypothetical protein [Leishmania major strain Friedlin] |
| XP_003860786.1 | 444 | LDBPK_220360 | eukaryota>euglenozoa>kinetoplastida | Leishmania | donovani | hypothetical protein, conserved [Leishmania donovani] |
| XP_003875503.1 | 444 | LMXM_22_0480 | eukaryota>euglenozoa>kinetoplastida | Leishmania | mexicana MHOM/GT/2001/U1103 | conserved hypothetical protein [Leishmania mexicana MHOM/GT/2001/U1103] |
| CAG06078.1 | 441 | GSTEN:00026313:G:001 | eukaryota>metazoa>chordata>vertebrata>actinopterygii | Tetraodon | nigroviridis | unnamed protein product, partial [Tetraodon nigroviridis] |
| OLP82663.1 | 439 | Ehd1 | eukaryota>alveolata | Symbiodinium | microadriaticum | EH domain-containing protein 1 [Symbiodinium microadriaticum] |
| XP_002765121.1 | 439 | Pmar_PMAR003862 | eukaryota>alveolata | Perkinsus | marinus ATCC 50983 | EH-domain-containing protein, putative [Perkinsus marinus ATCC 50983] |
| XP_009039421.1 | 437 | AURANDRAFT_30262 | eukaryota>stramenopiles | Aureococcus | anophagefferens | hypothetical protein AURANDRAFT_30262 [Aureococcus anophagefferens] |
| NP_001310596.1 | 431 | SAR | eukaryota>metazoa>chordata>vertebrata | Homo | sapiens | sarcalumenin isoform 2 [Homo sapiens] |
| NP_001310597.1 | 431 | SAR | eukaryota>metazoa>chordata>vertebrata | Homo | sapiens | sarcalumenin isoform 2 [Homo sapiens] |
| XP_016879017.1 | 431 | SRL | eukaryota>metazoa>chordata>vertebrata | Homo | sapiens | sarcalumenin isoform X2 [Homo sapiens] |
| XP_016879018.1 | 431 | SRL | eukaryota>metazoa>chordata>vertebrata | Homo | sapiens | sarcalumenin isoform X2 [Homo sapiens] |
| XP_017172335.1 | 431 | 9830004M20Rik,AW108387,sar | eukaryota>metazoa>chordata>vertebrata | Mus | musculus | PREDICTED: sarcalumenin isoform X5 [Mus musculus] |
| XP_017172336.1 | 431 | 9830004M20Rik,AW108387,sar | eukaryota>metazoa>chordata>vertebrata | Mus | musculus | PREDICTED: sarcalumenin isoform X5 [Mus musculus] |
| XP_002499973.1 | 427 | MICPUN_107789 | eukaryota>viridiplantae>chlorophyta | Micromonas | commoda | predicted protein [Micromonas commoda] |
| XP_016486423.1 | 427 | LOC107806716 | eukaryota>viridiplantae | Nicotiana | tabacum | PREDICTED: EH domain-containing protein 1-like isoform X1 [Nicotiana tabacum] |
| EWM25448.1 | 420 | RME1L1 | eukaryota>stramenopiles | Nannochloropsis | gaditana | receptor mediated endocytosis family member (rme-1) [Nannochloropsis gaditana] |
| Pisp1000006440 | 420 | Pisp1000006440 | eukaryota>fungi>neocallimastigomycota | Piromyces | sp | >Pisp1000006440 Pisp1000006440 |
| XP_005835236.1 | 414 | GUITHDRAFT_157516 | eukaryota>cryptophyta | Guillardia | theta CCMP2712 | hypothetical protein GUITHDRAFT_157516 [Guillardia theta CCMP2712] |
| XP_012352130.1 | 413 | EHD2 | eukaryota>metazoa>chordata>vertebrata | Nomascus | leucogenys | PREDICTED: EH domain-containing protein 2 [Nomascus leucogenys] |
| EFX74164.1 | 412 | DAPPUDRAFT_307451 | eukaryota>metazoa>crustacea | Daphnia | pulex | hypothetical protein DAPPUDRAFT_307451 [Daphnia pulex] |
| ETO05470.1 | 412 | RFI_31925 | eukaryota>rhizaria | Reticulomyxa | filosa | hypothetical protein RFI_31925 [Reticulomyxa filosa] |
| XP_009040272.1 | 405 | AURANDRAFT_1674 | eukaryota>stramenopiles | Aureococcus | anophagefferens | hypothetical protein AURANDRAFT_1674, partial [Aureococcus anophagefferens] |
| Smed1000013153 | 398 | Smed1000013153 | eukaryota>metazoa>platyhelminthe | Schmidtea | mediterranea | >Smed1000013153 Smed1000013153 |
| XP_016484342.1 | 397 | LOC107804905 | eukaryota>viridiplantae | Nicotiana | tabacum | PREDICTED: EH domain-containing protein 1-like, partial [Nicotiana tabacum] |
| XP_016487400.1 | 397 | LOC107807506 | eukaryota>viridiplantae | Nicotiana | tabacum | PREDICTED: EH domain-containing protein 1-like, partial [Nicotiana tabacum] |
| OEU22607.1 | 391 | FRACYDRAFT_166535 | eukaryota>stramenopiles | Fragilariopsis | cylindrus CCMP1102 | Dynamin_N-domain-containing protein [Fragilariopsis cylindrus CCMP1102] |
| XP_002177526.1 | 390 | PHATRDRAFT_8770 | eukaryota>stramenopiles | Phaeodactylum | tricornutum CCAP 1055/1 | predicted protein [Phaeodactylum tricornutum CCAP 1055/1] |
| XP_002291221.1 | 385 | THAPSDRAFT_13909 | eukaryota>stramenopiles | Thalassiosira | pseudonana CCMP1335 | eh domain-containing protein, partial [Thalassiosira pseudonana CCMP1335] |
| XP_002966377.1 | 385 | SELMODRAFT_85478 | eukaryota>viridiplantae | Selaginella | moellendorffii | hypothetical protein SELMODRAFT_85478, partial [Selaginella moellendorffii] |
| XP_002978188.1 | 385 | SELMODRAFT_108442 | eukaryota>viridiplantae | Selaginella | moellendorffii | hypothetical protein SELMODRAFT_108442, partial [Selaginella moellendorffii] |
| XP_009036435.1 | 383 | AURANDRAFT_1666 | eukaryota>stramenopiles | Aureococcus | anophagefferens | hypothetical protein AURANDRAFT_1666, partial [Aureococcus anophagefferens] |
| XP_002289284.1 | 382 | THAPSDRAFT_33868 | eukaryota>stramenopiles | Thalassiosira | pseudonana CCMP1335 | probable eh domain-containing protein, partial [Thalassiosira pseudonana CCMP1335] |
| XP_013898083.1 | 381 | MNEG_8899 | eukaryota>viridiplantae>chlorophyta | Monoraphidium | neglectum | EH domain-containing protein 4 [Monoraphidium neglectum] |
| ETO26663.1 | 374 | RFI_10471 | eukaryota>rhizaria | Reticulomyxa | filosa | EH-domain containing 4, partial [Reticulomyxa filosa] |
| XP_009293166.1 | 374 | si:ch211-147c1.2 | eukaryota>metazoa>chordata>vertebrata>actinopterygii | Danio | rerio | PREDICTED: EH domain-containing protein 3 isoform X1 [Danio rerio] |
| Bnat1000005148 | 374 | Bnat1000005148 | eukaryota>rhizaria | Bigelowiella | natans | >Bnat1000005148 Bnat1000005148 |
| XP_015129283.1 | 372 | LOC107049991 | eukaryota>metazoa>chordata>vertebrata | Gallus | gallus | PREDICTED: LOW QUALITY PROTEIN: EH domain-containing protein 2-like, partial [Gallus gallus] |
| XP_001896504.1 | 361 | Bm1_25225 | eukaryota>metazoa>nematoda | Brugia | malayi | EH-domain containing protein 3 [Brugia malayi] |
| ERG82150.1 | 353 | ASU_09059 | eukaryota>metazoa>nematoda | Ascaris | suum | eh domain-containing protein 1 [Ascaris suum] |
| ETO26762.1 | 351 | RFI_10373 | eukaryota>rhizaria | Reticulomyxa | filosa | hypothetical protein RFI_10373, partial [Reticulomyxa filosa] |
| XP_002779611.1 | 331 | Pmar_PMAR004572 | eukaryota>alveolata | Perkinsus | marinus ATCC 50983 | conserved hypothetical protein, partial [Perkinsus marinus ATCC 50983] |
| Bnat1000007215 | 329 | Bnat1000007215 | eukaryota>rhizaria | Bigelowiella | natans | >Bnat1000007215 Bnat1000007215 |
| XP_002411757.1 | 324 | IscW_ISCW021581 | eukaryota>metazoa | Ixodes | scapularis | endocytosis/signaling protein EHD1, putative [Ixodes scapularis] |
| XP_011531108.1 | 322 | PAST3 | eukaryota>metazoa>chordata>vertebrata | Homo | sapiens | EH domain-containing protein 3 isoform X1 [Homo sapiens] |
| ETO08814.1 | 320 | RFI_28574 | eukaryota>rhizaria | Reticulomyxa | filosa | EH domain-containing protein 4, partial [Reticulomyxa filosa] |
| XP_002769901.1 | 320 | Pmar_PMAR018864 | eukaryota>alveolata | Perkinsus | marinus ATCC 50983 | EH-domain-containing protein, putative, partial [Perkinsus marinus ATCC 50983] |
| ETN97991.1 | 312 | RFI_39531 | eukaryota>rhizaria | Reticulomyxa | filosa | hypothetical protein RFI_39531, partial [Reticulomyxa filosa] |
| XP_005837811.1 | 310 | GUITHDRAFT_157255 | eukaryota>cryptophyta | Guillardia | theta CCMP2712 | hypothetical protein GUITHDRAFT_157255 [Guillardia theta CCMP2712] |
| XP_012557383.1 | 301 | LOC100211925 | eukaryota>metazoa>cnidaria | Hydra | vulgaris | PREDICTED: EH domain-containing protein 3-like, partial [Hydra vulgaris] |
| Smed1000027697 | 296 | Smed1000027697 | eukaryota>metazoa>platyhelminthe | Schmidtea | mediterranea | >Smed1000027697 Smed1000027697 |
| Bnat1000019652 | 282 | Bnat1000019652 | eukaryota>rhizaria | Bigelowiella | natans | >Bnat1000019652 Bnat1000019652 |
| EPY43845.1 | 276 | AGDE_00076 | eukaryota>euglenozoa>kinetoplastida | Angomonas | deanei | hypothetical protein AGDE_00076 [Angomonas deanei] |
| OLP76986.1 | 271 | Ehd1 | eukaryota>alveolata | Symbiodinium | microadriaticum | EH domain-containing protein 1 [Symbiodinium microadriaticum] |
| XP_005833601.1 | 265 | GUITHDRAFT_52547 | eukaryota>cryptophyta | Guillardia | theta CCMP2712 | hypothetical protein GUITHDRAFT_52547, partial [Guillardia theta CCMP2712] |
| XP_006585912.1 | 248 | LOC100794877 | eukaryota>viridiplantae | Glycine | max | PREDICTED: EH domain-containing protein 2-like isoform X1 [Glycine max] |
| XP_006585913.1 | 246 | LOC100794877 | eukaryota>viridiplantae | Glycine | max | PREDICTED: EH domain-containing protein 2-like isoform X2 [Glycine max] |
| CAF87806.1 | 245 | GSTEN:00000798:G:001 | eukaryota>metazoa>chordata>vertebrata>actinopterygii | Tetraodon | nigroviridis | unnamed protein product, partial [Tetraodon nigroviridis] |
| XP_013406319.1 | 242 | LOC106170870 | eukaryota>metazoa | Lingula | anatina | EH domain-containing protein 1-like [Lingula anatina] |
| XP_016486424.1 | 242 | LOC107806716 | eukaryota>viridiplantae | Nicotiana | tabacum | PREDICTED: EH domain-containing protein 1-like isoform X2 [Nicotiana tabacum] |
| XP_006585914.1 | 241 | LOC100794877 | eukaryota>viridiplantae | Glycine | max | PREDICTED: EH domain-containing protein 2-like isoform X3 [Glycine max] |
| XP_006585915.1 | 241 | LOC100794877 | eukaryota>viridiplantae | Glycine | max | PREDICTED: EH domain-containing protein 2-like isoform X3 [Glycine max] |
| XP_006585916.1 | 236 | LOC100794877 | eukaryota>viridiplantae | Glycine | max | PREDICTED: EH domain-containing protein 2-like isoform X4 [Glycine max] |
| XP_006585917.1 | 236 | LOC100794877 | eukaryota>viridiplantae | Glycine | max | PREDICTED: EH domain-containing protein 2-like isoform X4 [Glycine max] |
| XP_004995955.1 | 230 | PTSG_03370 | eukaryota>choanoflagellida | Salpingoeca | rosetta | hypothetical protein PTSG_03370 [Salpingoeca rosetta] |
| XP_002411758.1 | 210 | IscW_ISCW021582 | eukaryota>metazoa | Ixodes | scapularis | hypothetical protein IscW_ISCW021582, partial [Ixodes scapularis] |
| XP_002182541.1 | 204 | PHATRDRAFT_14787 | eukaryota>stramenopiles | Phaeodactylum | tricornutum CCAP 1055/1 | predicted protein, partial [Phaeodactylum tricornutum CCAP 1055/1] |
| XP_016489207.1 | 204 | LOC107809121 | eukaryota>viridiplantae | Nicotiana | tabacum | PREDICTED: EH domain-containing protein 1-like [Nicotiana tabacum] |
| EWM24454.1 | 203 | RME1 | eukaryota>stramenopiles | Nannochloropsis | gaditana | dynamin GTPase family protein [Nannochloropsis gaditana] |
| ETO13471.1 | 196 | RFI_23901 | eukaryota>rhizaria | Reticulomyxa | filosa | EH domain-containing protein 4, partial [Reticulomyxa filosa] |
| XP_015129697.1 | 185 | LOC107050461 | eukaryota>metazoa>chordata>vertebrata | Gallus | gallus | PREDICTED: EH domain-containing protein 2-like, partial [Gallus gallus] |
| NP_001170407.1 | 177 | GRMZM2G113613 | eukaryota>viridiplantae | Zea | mays | uncharacterized protein LOC100384394 isoform 2 [Zea mays] |
| XP_008859898.1 | 162 | ENU1_187130 | eukaryota>amoebozoa>entamoebidae | Entamoeba | nuttalli P19 | receptor mediated endocytosis protein, putative, partial [Entamoeba nuttalli P19] |
| XP_005792863.1 | 148 | RME1A-2 | eukaryota>haptophyceae | Emiliania | huxleyi CCMP1516 | RME1A, RME-1/EHD family protein, partial [Emiliania huxleyi CCMP1516] |
| ENY62486.1 | 125 | EHI7A_091100 | eukaryota>amoebozoa>entamoebidae | Entamoeba | histolytica HM-1:IMSS-A | receptor mediated endocytosis protein, putative, partial [Entamoeba histolytica HM-1:IMSS-A] |
| EWM24449.1 | 116 | RME1 | eukaryota>stramenopiles | Nannochloropsis | gaditana | receptor mediated endocytosis protein, partial [Nannochloropsis gaditana] |
| ETO08813.1 | 106 | RFI_28573 | eukaryota>rhizaria | Reticulomyxa | filosa | receptor mediated endocytosis protein-like protein, partial [Reticulomyxa filosa] |
| EIE88190.1 | 88 | RO3G_12901 | eukaryota>fungi | Rhizopus | delemar RA 99-880 | hypothetical protein RO3G_12901 [Rhizopus delemar RA 99-880] |

P. IP3R

| | | | | | | |
|---|---|---|---|---|---|---|
| Bnat1000003017 | 4130 | Bnat1000003017 | eukaryota>rhizaria | Bigelowiella | natans | >Bnat1000003017 |
| EJY82846.1 | 3898 | OXYTRI_19537 | eukaryota>alveolata>ciliophora | Oxytricha | trifallax | Type 3 inositol 1,4,5-trisphosphate receptor, putative (macronuclear) [Oxytricha trifallax] |
| XP_001947344.2 | 3831 | LOC100163622 | eukaryota>metazoa>hexapoda | Acyrthosiphon | pisum | PREDICTED: inositol 1,4,5-trisphosphate receptor [Acyrthosiphon pisum] |
| XP_008185766.1 | 3831 | LOC100163622 | eukaryota>metazoa>hexapoda | Acyrthosiphon | pisum | PREDICTED: inositol 1,4,5-trisphosphate receptor [Acyrthosiphon pisum] |
| KPI87904.1 | 3823 | ABL78_3014 | eukaryota>euglenozoa>kinetoplastida | Leptomonas | seymouri | hypothetical protein ABL78_3014 [Leptomonas seymouri] |
| XP_001011602.2 | 3659 | TTHERM_00885700 | eukaryota>alveolata>ciliophora | Tetrahymena | thermophila SB210 | MIR domain protein [Tetrahymena thermophila SB210] |
| CDW75428.1 | 3530 | Contig1440.g1580 | eukaryota>alveolata>ciliophora | Stylonychia | lemnae | cation channel family protein [Stylonychia lemnae] |
| XP_001013405.2 | 3525 | TTHERM_00454060 | eukaryota>alveolata>ciliophora | Tetrahymena | thermophila SB210 | cation channel family protein [Tetrahymena thermophila SB210] |
| XP_976640.2 | 3489 | TTHERM_00545830 | eukaryota>alveolata>ciliophora | Tetrahymena | thermophila SB210 | MIR domain protein [Tetrahymena thermophila SB210] |
| XP_001009504.2 | 3449 | TTHERM_00370740 | eukaryota>alveolata>ciliophora | Tetrahymena | thermophila SB210 | cation channel family protein [Tetrahymena thermophila SB210] |
| XP_009493566.1 | 3443 | H696_01395 | eukaryota | Fonticula | alba | hypothetical protein H696_01395 [Fonticula alba] |
| XP_001029961.2 | 3412 | TTHERM_01205340 | eukaryota>alveolata>ciliophora | Tetrahymena | thermophila SB210 | IP3 receptor calcium ion channel protein [Tetrahymena thermophila SB210] |
| XP_001030533.2 | 3401 | TTHERM_01076970 | eukaryota>alveolata>ciliophora | Tetrahymena | thermophila SB210 | MIR domain protein [Tetrahymena thermophila SB210] |
| XP_017334767.1 | 3389 | LOC108271603 | eukaryota>metazoa>chordata>vertebrata>actinopterygii | Ictalurus | punctatus | PREDICTED: inositol 1,4,5-trisphosphate receptor type 1-like [Ictalurus punctatus] |
| CAG03532.1 | 3383 | GSTEN:00022986:G:001 | eukaryota>metazoa>chordata>vertebrata>actinopterygii | Tetraodon | nigroviridis | unnamed protein product [Tetraodon nigroviridis] |
| XP_001007910.2 | 3380 | TTHERM_00538860 | eukaryota>alveolata>ciliophora | Tetrahymena | thermophila SB210 | MIR domain protein [Tetrahymena thermophila SB210] |
| XP_001007911.2 | 3344 | TTHERM_00538870 | eukaryota>alveolata>ciliophora | Tetrahymena | thermophila SB210 | MIR domain protein [Tetrahymena thermophila SB210] |
| ELR11529.1 | 3344 | ACA1_256780 | eukaryota>amoebozoa>acanthamoebidae | Acanthamoeba | castellanii str. Neff | Inositol 1,4,5trisphosphate receptor type 2, putative [Acanthamoeba castellanii str. Neff] |
| XP_001010363.2 | 3260 | TTHERM_00354640 | eukaryota>alveolata>ciliophora | Tetrahymena | thermophila SB210 | IP3 receptor calcium ion channel protein [Tetrahymena thermophila SB210] |
| XP_001024368.2 | 3243 | TTHERM_00532470 | eukaryota>alveolata>ciliophora | Tetrahymena | thermophila SB210 | MIR domain protein [Tetrahymena thermophila SB210] |
| XP_004347577.1 | 3232 | CAOG_04826 | eukaryota>ichthyosporea | Capsaspora | owczarzaki ATCC 30864 | inositol 1,4,5-triphosphate receptor [Capsaspora owczarzaki ATCC 30864] |
| XP_009839519.1 | 3199 | H257_13636 | eukaryota>stramenopiles | Aphanomyces | astaci | hypothetical protein H257_13636 [Aphanomyces astaci] |
| XP_013762131.1 | 3179 | AMSG_00953 | eukaryota>apusozoa>apusomonadidae | Thecamonas | trahens ATCC 50062 | hypothetical protein AMSG_00953 [Thecamonas trahens ATCC 50062] |
| XP_629509.1 | 3177 | iplA | eukaryota>amoebozoa>mycetozoa>dictyosteliida | Dictyostelium | discoideum AX4 | inositol 1,4,5-trisphosphate receptor-like protein [Dictyostelium discoideum AX4] |
| XP_002959187.1 | 3167 | VOLCADRAFT_121781 | eukaryota>viridiplantae>chlorophyta | Volvox | carteri f. nagariensis | hypothetical protein VOLCADRAFT_121781 [Volvox carteri f. nagariensis] |
| XP_001019190.2 | 3160 | TTHERM_00997500 | eukaryota>alveolata>ciliophora | Tetrahymena | thermophila SB210 | MIR domain protein [Tetrahymena thermophila SB210] |
| XP_001701472.1 | 3140 | RYR1 | eukaryota>viridiplantae>chlorophyta | Chlamydomonas | reinhardtii | ryanodine-inositol 1,4,5-triphosphate receptor Ca2+ channel [Chlamydomonas reinhardtii] |
| XP_001442703.1 | 3130 | GSPATT00015090001 | eukaryota>alveolata>ciliophora | Paramecium | tetraurelia strain d4-2 | hypothetical protein (macronuclear) [Paramecium tetraurelia strain d4-2] |
| XP_001452546.1 | 3125 | GSPATT00025490001 | eukaryota>alveolata>ciliophora | Paramecium | tetraurelia strain d4-2 | hypothetical protein (macronuclear) [Paramecium tetraurelia strain d4-2] |
| XP_001440436.1 | 3121 | GSPATT00009387001 | eukaryota>alveolata>ciliophora | Paramecium | tetraurelia strain d4-2 | hypothetical protein (macronuclear) [Paramecium tetraurelia strain d4-2] |
| XP_001453569.1 | 3118 | GSPATT00019865001 | eukaryota>alveolata>ciliophora | Paramecium | tetraurelia strain d4-2 | hypothetical protein (macronuclear) [Paramecium tetraurelia strain d4-2] |
| XP_001019838.3 | 3116 | TTHERM_00138560 | eukaryota>alveolata>ciliophora | Tetrahymena | thermophila SB210 | inositol 1,4,5-trisphosphate receptor type 3 [Tetrahymena thermophila SB210] |
| XP_011775562.1 | 3099 | TbgDal_VIII2330 | eukaryota>euglenozoa>kinetoplastida | Trypanosoma | brucei gambiense DAL972 | hypothetical protein, conserved [Trypanosoma brucei gambiense DAL972] |
| XP_003283834.1 | 3089 | DICPUDRAFT_45096 | eukaryota>amoebozoa>mycetozoa>dictyosteliida | Dictyostelium | purpureum | hypothetical protein DICPUDRAFT_45096 [Dictyostelium purpureum] |
| XP_001013154.2 | 3056 | TTHERM_00295220 | eukaryota>alveolata>ciliophora | Tetrahymena | thermophila SB210 | zinc finger, C2H2 type family protein [Tetrahymena thermophila SB210] |
| XP_008878887.1 | 3037 | H310_13238 | eukaryota>stramenopiles | Aphanomyces | invadans | hypothetical protein H310_13238 [Aphanomyces invadans] |
| XP_001462377.1 | 3036 | GSPATT00027332001 | eukaryota>alveolata>ciliophora | Paramecium | tetraurelia strain d4-2 | hypothetical protein (macronuclear) [Paramecium tetraurelia strain d4-2] |
| XP_018668305.1 | 3019 | LOC101243127 | eukaryota>metazoa>chordata | Ciona | intestinalis | PREDICTED: inositol 1,4,5-trisphosphate receptor type 1 [Ciona intestinalis] |
| [illegible] | | | | | | |
| XP_019850149.1 | 2261 | LOC100633240 | eukaryota>metazoa | Amphimedon | queenslandica | PREDICTED: uncharacterized protein LOC100633240 [Amphimedon queenslandica] |
| XP_017305879.1 | 2244 | itpr1 | eukaryota>metazoa>chordata>vertebrata>actinopterygii | Ictalurus | punctatus | PREDICTED: inositol 1,4,5-trisphosphate receptor type 1 isoform X11 [Ictalurus punctatus] |
| Caps1000017277 | 2239 | Caps1000017277 | eukaryota>metazoa>annelida | Capitella | spI | >Caps1000017277 |
| ERG81215.1 | 2230 | ASU_10556 | eukaryota>metazoa>nematoda | Ascaris | suum | inositol -trisphosphate receptor type 1 [Ascaris suum] |
| XP_008613183.1 | 2216 | SDRG_09050 | eukaryota>stramenopiles | Saprolegnia | diclina VS20 | hypothetical protein SDRG_09050 [Saprolegnia diclina VS20] |
| XP_018104755.1 | 2206 | LOC108709441 | eukaryota>metazoa>chordata>vertebrata | Xenopus | laevis | PREDICTED: inositol 1,4,5-trisphosphate receptor type 3-like isoform X7 [Xenopus laevis] |
| XP_012208812.1 | 2168 | SPRG_14004 | eukaryota>stramenopiles | Saprolegnia | parasitica CBS 223.65 | hypothetical protein SPRG_14004 [Saprolegnia parasitica CBS 223.65] |
| XP_018104756.1 | 2160 | LOC108709441 | eukaryota>metazoa>chordata>vertebrata | Xenopus | laevis | PREDICTED: inositol 1,4,5-trisphosphate receptor type 3-like isoform X8 [Xenopus laevis] |
| XP_009827764.1 | 1933 | H257_04824 | eukaryota>stramenopiles | Aphanomyces | astaci | hypothetical protein, variant 4 [Aphanomyces astaci] |
| XP_009827765.1 | 1933 | H257_04824 | eukaryota>stramenopiles | Aphanomyces | astaci | hypothetical protein, variant 5 [Aphanomyces astaci] |
| XP_011531993.1 | 1910 | ACV,CLA4,INSP3R1,IP3R,IP3R1,PPP1R94,SCA15,SCA16,SCA29 | eukaryota>metazoa>chordata>vertebrata | Homo | sapiens | PREDICTED: inositol 1,4,5-trisphosphate receptor type 1 isoform X13 [Homo sapiens] |
| XP_012199467.1 | 1905 | SPRG_04939 | eukaryota>stramenopiles | Saprolegnia | parasitica CBS 223.65 | hypothetical protein SPRG_04939 [Saprolegnia parasitica CBS 223.65] |
| XP_011531994.1 | 1884 | ACV,CLA4,INSP3R1,IP3R,IP3R1,PPP1R94,SCA15,SCA16,SCA29 | eukaryota>metazoa>chordata>vertebrata | Homo | sapiens | PREDICTED: inositol 1,4,5-trisphosphate receptor type 1 isoform X14 [Homo sapiens] |
| XP_016861847.1 | 1883 | ACV,CLA4,INSP3R1,IP3R,IP3R1,PPP1R94,SCA15,SCA16,SCA29 | eukaryota>metazoa>chordata>vertebrata | Homo | sapiens | PREDICTED: inositol 1,4,5-trisphosphate receptor type 1 isoform X15 [Homo sapiens] |
| XP_016874758.1 | 1733 | ANHD,CFAP48,INSP3R2,IP3R2 | eukaryota>metazoa>chordata>vertebrata | Homo | sapiens | PREDICTED: inositol 1,4,5-trisphosphate receptor type 2 isoform X4 [Homo sapiens] |
| XP_016866321.1 | 1312 | IP3R,IP3R3 | eukaryota>metazoa>chordata>vertebrata | Homo | sapiens | PREDICTED: inositol 1,4,5-trisphosphate receptor type 3 isoform X2 [Homo sapiens] |
| OAF66980.1 | 1190 | A3Q56_05306 | eukaryota>metazoa | Intoshia | linei | hypothetical protein A3Q56_05306, partial [Intoshia linei] |
| OAF68200.1 | 928 | A3Q56_04067 | eukaryota>metazoa | Intoshia | linei | hypothetical protein A3Q56_04067 [Intoshia linei] |
| XP_014679742.1 | 367 | LOC106819653 | eukaryota>metazoa | Priapulus | caudatus | PREDICTED: inositol 1,4,5-trisphosphate receptor-like, partial [Priapulus caudatus] |

Q. ORAI

| | | | | | | |
|---|---|---|---|---|---|---|
| XP_001697902.1 | 2249 | CHLREDRAFT_193029 | eukaryota>viridiplantae>chlorophyta | Chlamydomonas | reinhardtii | predicted protein [Chlamydomonas reinhardtii] |
| OAF69866.1 | 1128 | A3Q56_02405 | eukaryota>metazoa | Intoshia | linei | 40S ribosomal protein S7 [Intoshia linei] |
| XP_005775441.1 | 1008 | EMIHUDRAFT_95611 | eukaryota>haptophyceae | Emiliania | huxleyi CCMP1516 | hypothetical protein EMIHUDRAFT_95611 [Emiliania huxleyi CCMP1516] |
| EUB64231.1 | 811 | EGR_00775 | eukaryota>metazoa | Echinococcus | granulosus | Pyridine nucleotide-disulfide oxidoreductase domain-containing protein [Echinococcus granulosus] |
| XP_009038390.1 | 777 | AURANDRAFT_65073 | eukaryota>stramenopiles | Aureococcus | anophagefferens | hypothetical protein AURANDRAFT_65073 [Aureococcus anophagefferens] |
| OLQ02130.1 | 762 | GTF3C5 | eukaryota>alveolata | Symbiodinium | microadriaticum | General transcription factor 3C polypeptide 5 [Symbiodinium microadriaticum] |
| XP_009174231.1 | 753 | T265_09793 | eukaryota>metazoa | Opisthorchis | viverrini | hypothetical protein T265_09793 [Opisthorchis viverrini] |
| XP_001769112.1 | 710 | PHYPADRAFT_166295 | eukaryota>viridiplantae | Physcomitrella | patens | predicted protein [Physcomitrella patens] |
| OLQ11388.1 | 561 | AK812_SmicGene4819 | eukaryota>alveolata | Symbiodinium | microadriaticum | hypothetical protein AK812_SmicGene4819 [Symbiodinium microadriaticum] |
| XP_001759550.1 | 535 | PHYPADRAFT_72636 | eukaryota>viridiplantae | Physcomitrella | patens | predicted protein [Physcomitrella patens] |
| XP_003056729.1 | 529 | MICPUCDRAFT_56086 | eukaryota>viridiplantae>chlorophyta | Micromonas | pusilla CCMP1545 | predicted protein [Micromonas pusilla CCMP1545] |
| XP_002972992.1 | 519 | SELMODRAFT_441966 | eukaryota>viridiplantae | Selaginella | moellendorffii | hypothetical protein SELMODRAFT_441966 [Selaginella moellendorffii] |
| NP_001137696.1 | 507 | BcDNA:LD03241,CG11430,CRACM1,Dmel\CG11430,dorai,dOrai,DOrai,dorai1,l(2)k11505,Olf186-F,orai,Orai,Orai1,SPH206 | eukaryota>metazoa>hexapoda | Drosophila | melanogaster | olf186-F, isoform E [Drosophila melanogaster] |
| NP_001261063.1 | 507 | BcDNA:LD03241,CG11430,CRACM1,Dmel\CG11430,dorai,dOrai,DOrai,dorai1,l(2)k11505,Olf186-F,orai,Orai,Orai1,SPH206 | eukaryota>metazoa>hexapoda | Drosophila | melanogaster | olf186-F, isoform G [Drosophila melanogaster] |
| XP_005788454.1 | 498 | EMIHUDRAFT_440744 | eukaryota>haptophyceae | Emiliania | huxleyi CCMP1516 | hypothetical protein EMIHUDRAFT_440744 [Emiliania huxleyi CCMP1516] |
| XP_005790774.1 | 498 | EMIHUDRAFT_440173 | eukaryota>haptophyceae | Emiliania | huxleyi CCMP1516 | hypothetical protein EMIHUDRAFT_440173 [Emiliania huxleyi CCMP1516] |
| XP_003081046.1 | 494 | Ot09g00560 | eukaryota>viridiplantae>chlorophyta | Ostreococcus | tauri | unnamed protein product [Ostreococcus tauri] |
| XP_001952386.2 | 493 | LOC100167625 | eukaryota>metazoa>hexapoda | Acyrthosiphon | pisum | PREDICTED: calcium release-activated calcium channel protein 1 [Acyrthosiphon pisum] |
| XP_008184407.1 | 493 | LOC100167625 | eukaryota>metazoa>hexapoda | Acyrthosiphon | pisum | PREDICTED: calcium release-activated calcium channel protein 1 [Acyrthosiphon pisum] |
| XP_016661006.1 | 493 | LOC100167625 | eukaryota>metazoa>hexapoda | Acyrthosiphon | pisum | PREDICTED: calcium release-activated calcium channel protein 1 [Acyrthosiphon pisum] |
| XP_002506166.1 | 473 | MICPUN_64064 | eukaryota>viridiplantae>chlorophyta | Micromonas | commoda | predicted protein [Micromonas commoda] |
| XP_005764484.1 | 464 | EMIHUDRAFT_213663 | eukaryota>haptophyceae | Emiliania | huxleyi CCMP1516 | hypothetical protein EMIHUDRAFT_213663 [Emiliania huxleyi CCMP1516] |
| XP_004365003.1 | 462 | CAOG_00132 | eukaryota>ichthyosporea | Capsaspora | owczarzaki ATCC 30864 | hypothetical protein CAOG_00132 [Capsaspora owczarzaki ATCC 30864] |
| XP_005761869.1 | 452 | EMIHUDRAFT_448361 | eukaryota>haptophyceae | Emiliania | huxleyi CCMP1516 | hypothetical protein EMIHUDRAFT_448361 [Emiliania huxleyi CCMP1516] |
| XP_002954571.1 | 444 | VOLCADRAFT_121327 | eukaryota>viridiplantae>chlorophyta | Volvox | carteri f. nagariensis | hypothetical protein VOLCADRAFT_121327 [Volvox carteri f. nagariensis] |
| ABO97995.1 | 442 | OSTLU_33356 | eukaryota>viridiplantae>chlorophyta | Ostreococcus | lucimarinus CCE9901 | predicted protein [Ostreococcus lucimarinus CCE9901] |
| OLP92260.1 | 438 | AK812_SmicGene25957 | eukaryota>alveolata | Symbiodinium | microadriaticum | hypothetical protein AK812_SmicGene25957 [Symbiodinium microadriaticum] |
| XP_001893964.1 | 433 | Bm1_12470 | eukaryota>metazoa>nematoda | Brugia | malayi | hypothetical protein Bm1_12470 [Brugia malayi] |
| XP_005788899.1 | 427 | EMIHUDRAFT_440534 | eukaryota>haptophyceae | Emiliania | huxleyi CCMP1516 | hypothetical protein EMIHUDRAFT_440534 [Emiliania huxleyi CCMP1516] |
| XP_005791795.1 | 427 | EMIHUDRAFT_439862 | eukaryota>haptophyceae | Emiliania | huxleyi CCMP1516 | hypothetical protein EMIHUDRAFT_439862 [Emiliania huxleyi CCMP1516] |
| XP_005757607.1 | 424 | EMIHUDRAFT_107392 | eukaryota>haptophyceae | Emiliania | huxleyi CCMP1516 | hypothetical protein EMIHUDRAFT_107392 [Emiliania huxleyi CCMP1516] |
| XP_005763784.1 | 411 | EMIHUDRAFT_452442 | eukaryota>haptophyceae | Emiliania | huxleyi CCMP1516 | hypothetical protein EMIHUDRAFT_452442 [Emiliania huxleyi CCMP1516] |
| XP_001689548.1 | 386 | SSA12 | eukaryota>viridiplantae>chlorophyta | Chlamydomonas | reinhardtii | predicted protein [Chlamydomonas reinhardtii] |
| CBJ26914.1 | 385 | Esi_0050_0029 | eukaryota>stramenopiles | Ectocarpus | siliculosus | expressed unknown protein [Ectocarpus siliculosus] |

| | | | | | | |
|---|---|---|---|---|---|---|
| XP_005773691.1 | 382 | EMIHUDRAFT_444540 | eukaryota>haptophyceae | Emiliania | huxleyi CCMP1516 | hypothetical protein EMIHUDRAFT_444540, partial [Emiliania huxleyi CCMP1516] |
| XP_005763709.1 | 381 | EMIHUDRAFT_247965 | eukaryota>haptophyceae | Emiliania | huxleyi CCMP1516 | hypothetical protein EMIHUDRAFT_247965 [Emiliania huxleyi CCMP1516] |
| XP_006559876.1 | 377 | LOC102655332 | eukaryota>metazoa>hexapoda | Apis | mellifera | PREDICTED: protein orai-2-like [Apis mellifera] |
| XP_005775253.1 | 374 | EMIHUDRAFT_95445 | eukaryota>haptophyceae | Emiliania | huxleyi CCMP1516 | hypothetical protein EMIHUDRAFT_95445 [Emiliania huxleyi CCMP1516] |
| XP_021205517.1 | 374 | LOC101739927 | eukaryota>metazoa>hexapoda | Bombyx | mori | calcium release-activated calcium channel protein 1 isoform X1 [Bombyx mori] |
| XP_012548235.1 | 371 | LOC101739927 | eukaryota>metazoa>hexapoda | Bombyx | mori | calcium release-activated calcium channel protein 1 isoform X2 [Bombyx mori] |
| XP_012548236.1 | 370 | LOC101739927 | eukaryota>metazoa>hexapoda | Bombyx | mori | calcium release-activated calcium channel protein 1 isoform X3 [Bombyx mori] |
| XP_012548237.1 | 366 | LOC101739927 | eukaryota>metazoa>hexapoda | Bombyx | mori | calcium release-activated calcium channel protein 1 isoform X4 [Bombyx mori] |
| XP_011144695.1 | 359 | LOC105186300 | eukaryota>metazoa>hexapoda | Harpegnathos | saltator | calcium release-activated calcium channel protein 1 [Harpegnathos saltator] |
| XP_001758939.1 | 353 | PHYPADRAFT_161647 | eukaryota>viridiplantae | Physcomitrella | patens | predicted protein [Physcomitrella patens] |
| NP_611273.1 | 351 | BcDNA:LD03241,CG11430,CRACM1,Dmel\CG11430,dorai,dOrai,DOrai,dOrai1,l(2)k11505,Olf186-F,orai,Orai,Orai1,SPH206 | eukaryota>metazoa>hexapoda | Drosophila | melanogaster | olf186-F, isoform B [Drosophila melanogaster] |
| NP_725727.1 | 351 | BcDNA:LD03241,CG11430,CRACM1,Dmel\CG11430,dorai,dOrai,DOrai,dOrai1,l(2)k11505,Olf186-F,orai,Orai,Orai1,SPH206 | eukaryota>metazoa>hexapoda | Drosophila | melanogaster | olf186-F, isoform A [Drosophila melanogaster] |
| NP_995881.2 | 351 | BcDNA:LD03241,CG11430,CRACM1,Dmel\CG11430,dorai,dOrai,DOrai,dOrai1,l(2)k11505,Olf186-F,orai,Orai,Orai1,SPH206 | eukaryota>metazoa>hexapoda | Drosophila | melanogaster | olf186-F, isoform F [Drosophila melanogaster] |
| XP_013227949.1 | 350 | ETH_00012845 | eukaryota>alveolata>apicomplexa | Eimeria | tenella | hypothetical protein, conserved [Eimeria tenella] |
| XP_002953952.1 | 340 | ssa12 | eukaryota>viridiplantae>chlorophyta | Volvox | carteri f. nagariensis | hypothetical protein VOLCADRAFT_127367 [Volvox carteri f. nagariensis] |
| XP_004993864.1 | 339 | PTSG_05291 | eukaryota>choanoflagellida | Salpingoeca | rosetta | hypothetical protein PTSG_05291 [Salpingoeca rosetta] |
| XP_021205518.1 | 339 | LOC101739927 | eukaryota>metazoa>hexapoda | Bombyx | mori | calcium release-activated calcium channel protein 1 isoform X5 [Bombyx mori] |
| OLP89363.1 | 333 | AK812_SmicGene29195 | eukaryota>alveolata | Symbiodinium | microadriaticum | hypothetical protein AK812_SmicGene29195 [Symbiodinium microadriaticum] |
| XP_002366727.1 | 329 | TGME49_241180 | eukaryota>alveolata>apicomplexa | Toxoplasma | gondii ME49 | hypothetical protein TGME49_241180 [Toxoplasma gondii ME49] |
| XP_005777770.1 | 329 | EMIHUDRAFT_471701 | eukaryota>haptophyceae | Emiliania | huxleyi CCMP1516 | hypothetical protein EMIHUDRAFT_471701 [Emiliania huxleyi CCMP1516] |
| XP_008887798.1 | 329 | HHA_241180 | eukaryota>alveolata>apicomplexa | Hammondia | hammondi | hypothetical protein HHA_241180 [Hammondia hammondi] |
| XP_008194887.1 | 323 | LOC657890 | eukaryota>metazoa>hexapoda | Tribolium | castaneum | PREDICTED: calcium release-activated calcium channel protein 1 isoform X1 [Tribolium castaneum] |
| XP_018670031.1 | 323 | LOC100180182 | eukaryota>metazoa>chordata | Ciona | intestinalis | PREDICTED: calcium release-activated calcium channel protein 1 isoform X1 [Ciona intestinalis] |
| XP_001850020.1 | 322 | CpipJ_CPIJ007873 | eukaryota>metazoa>hexapoda | Culex | quinquefasciatus | conserved hypothetical protein [Culex quinquefasciatus] |
| XP_013904039.1 | 320 | MNEG_2934 | eukaryota>viridiplantae>chlorophyta | Monoraphidium | neglectum | hypothetical protein MNEG_2934 [Monoraphidium neglectum] |
| XP_969411.1 | 320 | LOC657890 | eukaryota>metazoa>hexapoda | Tribolium | castaneum | PREDICTED: calcium release-activated calcium channel protein 1 isoform X2 [Tribolium castaneum] |
| XP_005782447.1 | 317 | EMIHUDRAFT_462903 | eukaryota>haptophyceae | Emiliania | huxleyi CCMP1516 | hypothetical protein EMIHUDRAFT_462903 [Emiliania huxleyi CCMP1516] |
| XP_002970256.1 | 316 | SELMODRAFT_441115 | eukaryota>viridiplantae | Selaginella | moellendorffii | hypothetical protein SELMODRAFT_441115 [Selaginella moellendorffii] |
| XP_002978414.1 | 316 | SELMODRAFT_418284 | eukaryota>viridiplantae | Selaginella | moellendorffii | hypothetical protein SELMODRAFT_418284 [Selaginella moellendorffii] |
| CDW80608.1 | 314 | Contig5165.g5534 | eukaryota>alveolata>ciliophora | Stylonychia | lemnae | superoxide dismutase [Stylonychia lemnae] |
| XP_005768431.1 | 313 | EMIHUDRAFT_210980 | eukaryota>haptophyceae | Emiliania | huxleyi CCMP1516 | hypothetical protein EMIHUDRAFT_210980 [Emiliania huxleyi CCMP1516] |
| XP_005770971.1 | 313 | EMIHUDRAFT_243748 | eukaryota>haptophyceae | Emiliania | huxleyi CCMP1516 | hypothetical protein EMIHUDRAFT_243748 [Emiliania huxleyi CCMP1516] |
| XP_002125402.1 | 312 | LOC100180182 | eukaryota>metazoa>chordata | Ciona | intestinalis | PREDICTED: calcium release-activated calcium channel protein 1 isoform X2 [Ciona intestinalis] |
| XP_005763455.1 | 312 | EMIHUDRAFT_446144 | eukaryota>haptophyceae | Emiliania | huxleyi CCMP1516 | hypothetical protein EMIHUDRAFT_446144 [Emiliania huxleyi CCMP1516] |
| XP_005767816.1 | 312 | EMIHUDRAFT_436817 | eukaryota>haptophyceae | Emiliania | huxleyi CCMP1516 | hypothetical protein EMIHUDRAFT_436817 [Emiliania huxleyi CCMP1516] |
| XP_012196003.1 | 311 | SPRG_02056 | eukaryota>stramenopiles | Saprolegnia | parasitica CBS 223.65 | hypothetical protein SPRG_02056 [Saprolegnia parasitica CBS 223.65] |
| XP_003249558.1 | 310 | LOC412524 | eukaryota>metazoa>hexapoda | Apis | mellifera | PREDICTED: calcium release-activated calcium channel protein 1 isoform X1 [Apis mellifera] |
| XP_009518362.1 | 309 | PHYSODRAFT_486152 | eukaryota>stramenopiles | Phytophthora | sojae | hypothetical protein PHYSODRAFT_486152 [Phytophthora sojae] |
| OEU19244.1 | 307 | FRACYDRAFT_235292 | eukaryota>stramenopiles | Fragilariopsis | cylindrus CCMP1102 | hypothetical protein FRACYDRAFT_235292 [Fragilariopsis cylindrus CCMP1102] |
| XP_008612529.1 | 306 | SDRG_08420 | eukaryota>stramenopiles | Saprolegnia | diclina VS20 | hypothetical protein SDRG_08420 [Saprolegnia diclina VS20] |
| NP_780632.1 | 304 | D730049H07Rik,orai-1,Tmem142a | eukaryota>metazoa>chordata>vertebrata | Mus | musculus | calcium release-activated calcium channel protein 1 [Mus musculus] |
| XP_005645699.1 | 304 | COCSUDRAFT_17677 | eukaryota>viridiplantae>chlorophyta | Coccomyxa | subellipsoidea C-169 | hypothetical protein COCSUDRAFT_17677, partial [Coccomyxa subellipsoidea C-169] |
| XP_008612530.1 | 304 | SDRG_08420 | eukaryota>stramenopiles | Saprolegnia | diclina VS20 | hypothetical protein, variant [Saprolegnia diclina VS20] |
| XP_009518359.1 | 304 | PHYSODRAFT_295628 | eukaryota>stramenopiles | Phytophthora | sojae | hypothetical protein PHYSODRAFT_295628 [Phytophthora sojae] |
| NP_116179.2 | 301 | CRACM1,IMD9,ORAT1,TAM2,TMEM142A | eukaryota>metazoa>chordata>vertebrata | Homo | sapiens | calcium release-activated calcium channel protein 1 [Homo sapiens] |
| XP_008861378.1 | 301 | H310_00383 | eukaryota>stramenopiles | Aphanomyces | invadans | hypothetical protein H310_00383 [Aphanomyces invadans] |
| XP_011142627.1 | 301 | LOC105185086 | eukaryota>metazoa>hexapoda | Harpegnathos | saltator | PREDICTED: calcium release-activated calcium channel protein 1 isoform X1 [Harpegnathos saltator] |
| EJY70280.1 | 300 | OXYTRI_08975 | eukaryota>alveolata>ciliophora | Oxytricha | trifallax | Superoxide dismutase [Cu-Zn] (macronuclear) [Oxytricha trifallax] |
| XP_005591748.1 | 300 | ORAI3 | eukaryota>metazoa>chordata>vertebrata | Macaca | fascicularis | PREDICTED: protein orai-3 isoform X1 [Macaca fascicularis] |
| XP_004928907.1 | 299 | LOC101739927 | eukaryota>metazoa>hexapoda | Bombyx | mori | calcium release-activated calcium channel protein 1 isoform X6 [Bombyx mori] |
| NP_001254835.1 | 297 | orai-1 | eukaryota>metazoa>nematoda | Caenorhabditis | elegans | ORAI (store-operated calcium channel) homolog [Caenorhabditis elegans] |
| XP_002984309.1 | 297 | SELMODRAFT_423451 | eukaryota>viridiplantae | Selaginella | moellendorffii | hypothetical protein SELMODRAFT_423451 [Selaginella moellendorffii] |
| EJK58875.1 | 296 | THAOC_20967 | eukaryota>stramenopiles | Thalassiosira | oceanica | hypothetical protein THAOC_20967 [Thalassiosira oceanica] |
| NP_689501.1 | 295 | TMEM142C | eukaryota>metazoa>chordata>vertebrata | Homo | sapiens | protein orai-3 [Homo sapiens] |
| XP_003280496.1 | 295 | ORAI3 | eukaryota>metazoa>chordata>vertebrata | Nomascus | leucogenys | PREDICTED: protein orai-3 [Nomascus leucogenys] |
| XP_005591749.1 | 295 | ORAI3 | eukaryota>metazoa>chordata>vertebrata | Macaca | fascicularis | PREDICTED: protein orai-3 isoform X2 [Macaca fascicularis] |
| NP_001254834.1 | 293 | orai-1 | eukaryota>metazoa>nematoda | Caenorhabditis | elegans | Protein orai [Caenorhabditis elegans] |
| XP_009822282.1 | 292 | H257_01002 | eukaryota>stramenopiles | Aphanomyces | astaci | hypothetical protein H257_01002 [Aphanomyces astaci] |
| XP_002898394.1 | 291 | PITG_15443 | eukaryota>stramenopiles | Phytophthora | infestans T30-4 | conserved hypothetical protein [Phytophthora infestans T30-4] |
| NP_940816.1 | 290 | 9930124N15,BC061259,Tmem142c | eukaryota>metazoa>chordata>vertebrata | Mus | musculus | protein orai-3 [Mus musculus] |
| XP_003881962.1 | 289 | NCLIV_017210 | eukaryota>alveolata>apicomplexa | Neospora | caninum Liverpool | conserved hypothetical protein [Neospora caninum Liverpool] |
| XP_003882987.1 | 288 | NCLIV_027440 | eukaryota>alveolata>apicomplexa | Neospora | caninum Liverpool | conserved hypothetical protein [Neospora caninum Liverpool] |
| XP_018637036.1 | 286 | TGME49_259050 | eukaryota>alveolata>apicomplexa | Toxoplasma | gondii ME49 | hypothetical protein TGME49_259050 [Toxoplasma gondii ME49] |
| XP_008883102.1 | 283 | HHA_259050 | eukaryota>alveolata>apicomplexa | Hammondia | hammondi | hypothetical protein HHA_259050 [Hammondia hammondi] |

[illegible]

| | | | | | | |
|---|---|---|---|---|---|---|
| XP_004092266.1 | 116 | RCAN3 | eukaryota>metazoa>chordata>vertebrata | Nomascus | leucogenys | PREDICTED: calcipressin-3 isoform X3 [Nomascus leucogenys] |
| XP_012353797.1 | 116 | RCAN3 | eukaryota>metazoa>chordata>vertebrata | Nomascus | leucogenys | PREDICTED: calcipressin-3 isoform X3 [Nomascus leucogenys] |
| XP_006825325.1 | 102 | LOC102806619 | eukaryota>metazoa>hemichordata | Saccoglossus | kowalevskii | PREDICTED: protein sarah-like, partial [Saccoglossus kowalevskii] |
| XP_001745353.1 | 53 | MONBRDRAFT_3061 | eukaryota>choanoflagellida | Monosiga | brevicollis MX1 | hypothetical protein, partial [Monosiga brevicollis MX1] |

T. STIM

| | | | | | | |
|---|---|---|---|---|---|---|
| XP_017317088.1 | 1028 | LOC108260897 | eukaryota>metazoa>chordata>vertebrata>actinopterygii | Ictalurus | punctatus | PREDICTED: stromal interaction molecule 2-like [Ictalurus punctatus] |
| XP_017317089.1 | 1028 | LOC108260897 | eukaryota>metazoa>chordata>vertebrata>actinopterygii | Ictalurus | punctatus | PREDICTED: stromal interaction molecule 2-like [Ictalurus punctatus] |
| XP_017317090.1 | 1028 | LOC108260897 | eukaryota>metazoa>chordata>vertebrata>actinopterygii | Ictalurus | punctatus | PREDICTED: stromal interaction molecule 2-like [Ictalurus punctatus] |
| XP_017317091.1 | 1028 | LOC108260897 | eukaryota>metazoa>chordata>vertebrata>actinopterygii | Ictalurus | punctatus | PREDICTED: stromal interaction molecule 2-like [Ictalurus punctatus] |
| XP_017317092.1 | 1028 | LOC108260897 | eukaryota>metazoa>chordata>vertebrata>actinopterygii | Ictalurus | punctatus | PREDICTED: stromal interaction molecule 2-like [Ictalurus punctatus] |
| XP_002660454.3 | 992 | stim2a | eukaryota>metazoa>chordata>vertebrata>actinopterygii | Danio | rerio | stromal interaction molecule 2 isoform X1 [Danio rerio] |
| XP_002154218.2 | 858 | LOC100198735 | eukaryota>metazoa>cnidaria | Hydra | vulgaris | PREDICTED: uncharacterized protein LOC100198735, partial [Hydra vulgaris] |
| XP_003258607.1 | 841 | STIM2 | eukaryota>metazoa>chordata>vertebrata | Nomascus | leucogenys | PREDICTED: stromal interaction molecule 2 isoform X1 [Nomascus leucogenys] |
| XP_005554693.1 | 840 | STIM2 | eukaryota>metazoa>chordata>vertebrata | Macaca | fascicularis | PREDICTED: stromal interaction molecule 2 isoform X1 [Macaca fascicularis] |
| XP_003258605.1 | 833 | STIM2 | eukaryota>metazoa>chordata>vertebrata | Nomascus | leucogenys | PREDICTED: stromal interaction molecule 2 isoform X2 [Nomascus leucogenys] |
| XP_005554694.1 | 832 | STIM2 | eukaryota>metazoa>chordata>vertebrata | Macaca | fascicularis | PREDICTED: stromal interaction molecule 2 isoform X2 [Macaca fascicularis] |
| XP_006503745.1 | 820 | Stim2 | eukaryota>metazoa>chordata>vertebrata | Mus | musculus | PREDICTED: stromal interaction molecule 2 isoform X1 [Mus musculus] |
| CAG04573.1 | 820 | GSTEN:00024339:G:001 | eukaryota>metazoa>chordata>vertebrata>actinopterygii | Tetraodon | nigroviridis | unnamed protein product, partial [Tetraodon nigroviridis] |
| XP_013773068.1 | 809 | LOC106458154 | eukaryota>metazoa | Limulus | polyphemus | PREDICTED: uncharacterized protein LOC106458154 [Limulus polyphemus] |
| XP_005173449.1 | 808 | si:dkey-24p1.5 | eukaryota>metazoa>chordata>vertebrata>actinopterygii | Danio | rerio | stromal interaction molecule 1 isoform X1 [Danio rerio] |
| XP_006507596.1 | 793 | SIM | eukaryota>metazoa>chordata>vertebrata | Mus | musculus | PREDICTED: stromal interaction molecule 1 isoform X1 [Mus musculus] |
| XP_004090689.1 | 792 | STIM1 | eukaryota>metazoa>chordata>vertebrata | Nomascus | leucogenys | PREDICTED: stromal interaction molecule 1 isoform X1 [Nomascus leucogenys] |
| NP_001264890.1 | 791 | D11S4896E,GOK,IMD10,STRMK,TAM,TAM1 | eukaryota>metazoa>chordata>vertebrata | Homo | sapiens | stromal interaction molecule 1 isoform 1 precursor [Homo sapiens] |
| XP_005579046.1 | 784 | STIM1 | eukaryota>metazoa>chordata>vertebrata | Macaca | fascicularis | PREDICTED: stromal interaction molecule 1 isoform X1 [Macaca fascicularis] |
| XP_005173450.1 | 781 | si:dkey-24p1.5 | eukaryota>metazoa>chordata>vertebrata>actinopterygii | Danio | rerio | stromal interaction molecule 1 isoform X2 [Danio rerio] |
| NP_001162589.1 | 754 | STIM2 | eukaryota>metazoa>chordata>vertebrata | Homo | sapiens | stromal interaction molecule 2 isoform 1 precursor [Homo sapiens] |
| XP_009162000.1 | 753 | T265_00079 | eukaryota>metazoa | Opisthorchis | viverrini | hypothetical protein T265_00079 [Opisthorchis viverrini] |
| XP_017327752.1 | 753 | stim2 | eukaryota>metazoa>chordata>vertebrata>actinopterygii | Ictalurus | punctatus | PREDICTED: stromal interaction molecule 2 isoform X1 [Ictalurus punctatus] |
| NP_001074572.2 | 746 | Stim2 | eukaryota>metazoa>chordata>vertebrata | Mus | musculus | stromal interaction molecule 2 precursor [Mus musculus] |
| NP_065911.3 | 746 | STIM2 | eukaryota>metazoa>chordata>vertebrata | Homo | sapiens | stromal interaction molecule 2 isoform 2 precursor [Homo sapiens] |
| XP_018098572.1 | 746 | LOC108706560 | eukaryota>metazoa>chordata>vertebrata | Xenopus | laevis | PREDICTED: stromal interaction molecule 2-like [Xenopus laevis] |
| XP_420749.5 | 740 | STIM2 | eukaryota>metazoa>chordata>vertebrata | Gallus | gallus | PREDICTED: stromal interaction molecule 2 isoform X1 [Gallus gallus] |
| XP_011605056.1 | 736 | stim2 | eukaryota>metazoa>chordata>vertebrata>actinopterygii | Takifugu | rubripes | PREDICTED: stromal interaction molecule 2 [Takifugu rubripes] |
| XP_692356.4 | 727 | stim2b | eukaryota>metazoa>chordata>vertebrata>actinopterygii | Danio | rerio | stromal interaction molecule 2 [Danio rerio] |
| XP_018124030.1 | 722 | LOC108719591 | eukaryota>metazoa>chordata>vertebrata | Xenopus | laevis | PREDICTED: stromal interaction molecule 2-like [Xenopus laevis] |
| XP_017327753.1 | 722 | stim2 | eukaryota>metazoa>chordata>vertebrata>actinopterygii | Ictalurus | punctatus | PREDICTED: stromal interaction molecule 2 isoform X2 [Ictalurus punctatus] |
| XP_011439185.1 | 720 | LOC105336527 | eukaryota>metazoa>mollusca | Crassostrea | gigas | PREDICTED: stromal interaction molecule 1 isoform X1 [Crassostrea gigas] |
| XP_006507597.1 | 716 | SIM | eukaryota>metazoa>chordata>vertebrata | Mus | musculus | PREDICTED: stromal interaction molecule 1 isoform X2 [Mus musculus] |
| XP_005579047.1 | 715 | STIM1 | eukaryota>metazoa>chordata>vertebrata | Macaca | fascicularis | PREDICTED: stromal interaction molecule 1 isoform X2 [Macaca fascicularis] |
| XP_011439186.1 | 712 | LOC105336527 | eukaryota>metazoa>mollusca | Crassostrea | gigas | PREDICTED: stromal interaction molecule 1 isoform X2 [Crassostrea gigas] |
| XP_009290146.1 | 703 | si:dkey-24p1.5 | eukaryota>metazoa>chordata>vertebrata>actinopterygii | Danio | rerio | stromal interaction molecule 1 isoform X3 [Danio rerio] |
| XP_013386698.1 | 701 | LOC106156136 | eukaryota>metazoa | Lingula | anatina | PREDICTED: stromal interaction molecule homolog [Lingula anatina] |
| XP_011439187.1 | 701 | LOC105336527 | eukaryota>metazoa>mollusca | Crassostrea | gigas | PREDICTED: stromal interaction molecule 1 isoform X3 [Crassostrea gigas] |
| XP_002126383.1 | 700 | LOC100178893 | eukaryota>metazoa>chordata | Ciona | intestinalis | PREDICTED: stromal interaction molecule homolog isoform X1 [Ciona intestinalis] |
| XP_002598264.1 | 698 | BRAFLDRAFT_119144 | eukaryota>metazoa>chordata | Branchiostoma | floridae | hypothetical protein BRAFLDRAFT_119144 [Branchiostoma floridae] |
| XP_009857576.1 | 698 | LOC100178893 | eukaryota>metazoa>chordata | Ciona | intestinalis | PREDICTED: stromal interaction molecule homolog isoform X2 [Ciona intestinalis] |
| XP_012551138.1 | 695 | Stim1 | eukaryota>metazoa>hexapoda | Bombyx | mori | stromal interaction molecule 1 isoform X1 [Bombyx mori] |
| XP_012551147.1 | 694 | Stim1 | eukaryota>metazoa>hexapoda | Bombyx | mori | stromal interaction molecule 1 isoform X2 [Bombyx mori] |
| XP_011439188.1 | 693 | LOC105336527 | eukaryota>metazoa>mollusca | Crassostrea | gigas | PREDICTED: stromal interaction molecule 1 isoform X4 [Crassostrea gigas] |
| Smar1000002138 | 692 | Smar1000002138 | eukaryota>metazoa>arthropoda>myriapoda | Strigamia | maritima | >Smar1000002138 |
| XP_005554695.1 | 692 | STIM2 | eukaryota>metazoa>chordata>vertebrata | Macaca | fascicularis | PREDICTED: stromal interaction molecule 2 isoform X3 [Macaca fascicularis] |
| XP_011609388.1 | 692 | LOC101077194 | eukaryota>metazoa>chordata>vertebrata>actinopterygii | Takifugu | rubripes | PREDICTED: stromal interaction molecule 1-like isoform X1 [Takifugu rubripes] |
| XP_011609389.1 | 692 | LOC101077194 | eukaryota>metazoa>chordata>vertebrata>actinopterygii | Takifugu | rubripes | PREDICTED: stromal interaction molecule 1-like isoform X1 [Takifugu rubripes] |
| XP_011609390.1 | 691 | LOC101077194 | eukaryota>metazoa>chordata>vertebrata>actinopterygii | Takifugu | rubripes | PREDICTED: stromal interaction molecule 1-like isoform X2 [Takifugu rubripes] |
| XP_011609391.1 | 690 | LOC101077194 | eukaryota>metazoa>chordata>vertebrata>actinopterygii | Takifugu | rubripes | PREDICTED: stromal interaction molecule 1-like isoform X3 [Takifugu rubripes] |
| XP_395207.4 | 687 | LOC411739 | eukaryota>metazoa>hexapoda | Apis | mellifera | PREDICTED: stromal interaction molecule homolog isoform X1 [Apis mellifera] |
| XP_003258606.1 | 686 | STIM2 | eukaryota>metazoa>chordata>vertebrata | Nomascus | leucogenys | PREDICTED: stromal interaction molecule 2 isoform X3 [Nomascus leucogenys] |
| NP_003147.2 | 685 | D11S4896E,GOK,IMD10,STRMK,TAM,TAM1 | eukaryota>metazoa>chordata>vertebrata | Homo | sapiens | stromal interaction molecule 1 isoform 2 precursor [Homo sapiens] |
| NP_033313.2 | 685 | SIM | eukaryota>metazoa>chordata>vertebrata | Mus | musculus | stromal interaction molecule 1 precursor [Mus musculus] |
| XP_003254842.1 | 685 | STIM1 | eukaryota>metazoa>chordata>vertebrata | Nomascus | leucogenys | PREDICTED: stromal interaction molecule 1 isoform X2 [Nomascus leucogenys] |
| XP_005579048.1 | 684 | STIM1 | eukaryota>metazoa>chordata>vertebrata | Macaca | fascicularis | PREDICTED: stromal interaction molecule 1 isoform X3 [Macaca fascicularis] |
| XP_018673097.1 | 677 | LOC100178893 | eukaryota>metazoa>chordata | Ciona | intestinalis | PREDICTED: stromal interaction molecule homolog isoform X3 [Ciona intestinalis] |
| NP_001038264.1 | 676 | si:dkey-24p1.5 | eukaryota>metazoa>chordata>vertebrata>actinopterygii | Danio | rerio | stromal interaction molecule 1 precursor [Danio rerio] |
| XP_011142898.1 | 676 | LOC105185257 | eukaryota>metazoa>hexapoda | Harpegnathos | saltator | PREDICTED: stromal interaction molecule homolog [Harpegnathos saltator] |
| XP_011606881.1 | 671 | LOC101077340 | eukaryota>metazoa>chordata>vertebrata>actinopterygii | Takifugu | rubripes | PREDICTED: stromal interaction molecule 1-like [Takifugu rubripes] |
| XP_013776985.1 | 669 | LOC106461685 | eukaryota>metazoa | Limulus | polyphemus | PREDICTED: stromal interaction molecule homolog isoform X1 [Limulus polyphemus] |
| XP_018105965.1 | 669 | LOC108710082 | eukaryota>metazoa>chordata>vertebrata | Xenopus | laevis | PREDICTED: stromal interaction molecule 1-like isoform X1 [Xenopus laevis] |
| XP_017346476.1 | 669 | LOC108277929 | eukaryota>metazoa>chordata>vertebrata>actinopterygii | Ictalurus | punctatus | PREDICTED: stromal interaction molecule 1-like [Ictalurus punctatus] |
| NP_001090506.1 | 668 | stim1,stim1.S | eukaryota>metazoa>chordata>vertebrata | Xenopus | laevis | stromal interaction molecule 1 L homeolog precursor [Xenopus laevis] |
| XP_018105966.1 | 668 | LOC108710082 | eukaryota>metazoa>chordata>vertebrata | Xenopus | laevis | PREDICTED: stromal interaction molecule 1-like isoform X2 [Xenopus laevis] |
| XP_005161345.1 | 668 | si:ch211-112117.2,stim1 | eukaryota>metazoa>chordata>vertebrata>actinopterygii | Danio | rerio | stromal interaction molecule 1 isoform X1 [Danio rerio] |
| XP_015778612.1 | 667 | LOC107356518 | eukaryota>metazoa>cnidaria | Acropora | digitifera | PREDICTED: stromal interaction molecule 1-like [Acropora digitifera] |
| XP_017348463.1 | 666 | stim1 | eukaryota>metazoa>chordata>vertebrata>actinopterygii | Ictalurus | punctatus | PREDICTED: stromal interaction molecule 1 isoform X1 [Ictalurus punctatus] |
| XP_003970828.1 | 661 | LOC101077194 | eukaryota>metazoa>chordata>vertebrata>actinopterygii | Takifugu | rubripes | PREDICTED: stromal interaction molecule 1-like isoform X4 [Takifugu rubripes] |
| XP_001951196.2 | 657 | LOC100166274 | eukaryota>metazoa>hexapoda | Acyrthosiphon | pisum | PREDICTED: stromal interaction molecule homolog isoform X1 [Acyrthosiphon pisum] |
| CAF95381.1 | 656 | GSTEN:00012009:G:001 | eukaryota>metazoa>chordata>vertebrata>actinopterygii | Tetraodon | nigroviridis | unnamed protein product, partial [Tetraodon nigroviridis] |
| XP_008192163.1 | 652 | LOC664047 | eukaryota>metazoa>hexapoda | Tribolium | castaneum | PREDICTED: stromal interaction molecule homolog isoform X1 [Tribolium castaneum] |
| XP_019853545.1 | 650 | LOC100633433 | eukaryota>metazoa | Amphimedon | queenslandica | PREDICTED: stromal interaction molecule 1-like isoform X1 [Amphimedon queenslandica] |
| XP_019853546.1 | 650 | LOC100633433 | eukaryota>metazoa | Amphimedon | queenslandica | PREDICTED: stromal interaction molecule 1-like isoform X1 [Amphimedon queenslandica] |
| XP_003387480.1 | 649 | LOC100633433 | eukaryota>metazoa | Amphimedon | queenslandica | PREDICTED: stromal interaction molecule 1-like isoform X2 [Amphimedon queenslandica] |
| CDS35874.1 | 646 | EmuJ_001183000 | eukaryota>metazoa | Echinococcus | multilocularis | stromal interaction molecule 1 [Echinococcus multilocularis] |
| XP_975159.2 | 646 | LOC664047 | eukaryota>metazoa>hexapoda | Tribolium | castaneum | PREDICTED: stromal interaction molecule homolog isoform X2 [Tribolium castaneum] |
| XP_004936189.2 | 642 | STIM2 | eukaryota>metazoa>chordata>vertebrata | Gallus | gallus | PREDICTED: stromal interaction molecule 2 isoform X2 [Gallus gallus] |
| XP_006507598.1 | 642 | SIM | eukaryota>metazoa>chordata>vertebrata | Mus | musculus | PREDICTED: stromal interaction molecule 1 isoform X3 [Mus musculus] |
| XP_017177567.1 | 642 | SIM | eukaryota>metazoa>chordata>vertebrata | Mus | musculus | PREDICTED: stromal interaction molecule 1 isoform X3 [Mus musculus] |
| NP_001189470.1 | 642 | si:ch211-112117.2,stim1 | eukaryota>metazoa>chordata>vertebrata>actinopterygii | Danio | rerio | stromal interaction molecule 1 precursor [Danio rerio] |
| XP_009293832.1 | 641 | si:ch211-112117.2,stim1 | eukaryota>metazoa>chordata>vertebrata>actinopterygii | Danio | rerio | stromal interaction molecule 1 isoform X2 [Danio rerio] |
| XP_017348464.1 | 641 | stim1 | eukaryota>metazoa>chordata>vertebrata>actinopterygii | Ictalurus | punctatus | PREDICTED: stromal interaction molecule 1 isoform X2 [Ictalurus punctatus] |
| XP_006818739.1 | 639 | LOC100366516 | eukaryota>metazoa>hemichordata | Saccoglossus | kowalevskii | PREDICTED: stromal interaction molecule 1-like [Saccoglossus kowalevskii] |
| XP_008186734.1 | 635 | LOC100166274 | eukaryota>metazoa>hexapoda | Acyrthosiphon | pisum | PREDICTED: stromal interaction molecule homolog isoform X2 [Acyrthosiphon pisum] |
| XP_003976755.2 | 631 | LOC101080161 | eukaryota>metazoa>chordata>vertebrata>actinopterygii | Takifugu | rubripes | PREDICTED: LOW QUALITY PROTEIN: stromal interaction molecule 2-like, partial [Takifugu rubripes] |
| XP_008192165.1 | 631 | LOC664047 | eukaryota>metazoa>hexapoda | Tribolium | castaneum | PREDICTED: stromal interaction molecule homolog isoform X3 [Tribolium castaneum] |
| XP_013407301.1 | 627 | LOC106171477 | eukaryota>metazoa | Lingula | anatina | PREDICTED: stromal interaction molecule 1-like [Lingula anatina] |
| XP_008192166.1 | 625 | LOC664047 | eukaryota>metazoa>hexapoda | Tribolium | castaneum | PREDICTED: stromal interaction molecule homolog isoform X4 [Tribolium castaneum] |
| XP_013776228.1 | 616 | LOC106461002 | eukaryota>metazoa | Limulus | polyphemus | PREDICTED: stromal interaction molecule homolog, partial [Limulus polyphemus] |
| NP_001026009.1 | 612 | STIM1 | eukaryota>metazoa>chordata>vertebrata | Gallus | gallus | stromal interaction molecule 1 [Gallus gallus] |
| XP_009031508.1 | 609 | HELRODRAFT_189915 | eukaryota>metazoa>annelida | Helobdella | robusta | hypothetical protein HELRODRAFT_189915 [Helobdella robusta] |
| XP_017327754.1 | 607 | stim2 | eukaryota>metazoa>chordata>vertebrata>actinopterygii | Ictalurus | punctatus | PREDICTED: stromal interaction molecule 2 isoform X3 [Ictalurus punctatus] |
| XP_018105967.1 | 606 | LOC108710082 | eukaryota>metazoa>chordata>vertebrata | Xenopus | laevis | PREDICTED: stromal interaction molecule 1-like isoform X3 [Xenopus laevis] |
| KXJ22905.1 | 605 | STIM1 | eukaryota>metazoa>cnidaria | Exaiptasia | pallida | Stromal interaction molecule 1 [Exaiptasia pallida] |
| XP_016767778.1 | 604 | LOC411739 | eukaryota>metazoa>hexapoda | Apis | mellifera | PREDICTED: stromal interaction molecule homolog isoform X2 [Apis mellifera] |
| XP_014788351.1 | 600 | LOC106882253 | eukaryota>metazoa>mollusca | Octopus | bimaculoides | PREDICTED: stromal interaction molecule 1-like [Octopus bimaculoides] |
| NP_001162588.1 | 599 | STIM2 | eukaryota>metazoa>chordata>vertebrata | Homo | sapiens | stromal interaction molecule 2 isoform 3 precursor [Homo sapiens] |
| CAG08997.1 | 598 | GSTEN:00030156:G:001 | eukaryota>metazoa>chordata>vertebrata>actinopterygii | Tetraodon | nigroviridis | unnamed protein product, partial [Tetraodon nigroviridis] |
| XP_001900500.1 | 590 | Bm1_45190 | eukaryota>metazoa>nematoda | Brugia | malayi | hypothetical protein Bm1_45190 [Brugia malayi] |
| XP_001850572.1 | 589 | CpipJ_CPIJ009010 | eukaryota>metazoa>hexapoda | Culex | quinquefasciatus | conserved hypothetical protein [Culex quinquefasciatus] |
| OQV20132.1 | 584 | BV898_05924 | eukaryota>metazoa | Hypsibius | dujardini | Stromal interaction molecule 1 [Hypsibius dujardini] |
| XP_011666877.1 | 578 | LOC581356 | eukaryota>metazoa>echinodermata | Strongylocentrotus | purpuratus | PREDICTED: stromal interaction molecule 1, partial [Strongylocentrotus purpuratus] |
| NP_001128674.1 | 577 | Stim1 | eukaryota>metazoa>hexapoda | Bombyx | mori | stromal interaction molecule 1 precursor [Bombyx mori] |
| XP_003382288.1 | 575 | Tsp_09128 | eukaryota>metazoa>nematoda | Trichinella | spiralis | stromal interaction molecule-like protein [Trichinella spiralis] |
| XP_006507599.1 | 571 | SIM | eukaryota>metazoa>chordata>vertebrata | Mus | musculus | PREDICTED: stromal interaction molecule 1 isoform X4 [Mus musculus] |
| NP_523357.2 | 570 | C-Stim,CG9126,D-Stim,D-STIM,Dmel\CG9126,dstim,dStim,dSTIM,STIM,stim1,STIM1 | eukaryota>metazoa>hexapoda | Drosophila | melanogaster | stromal interaction molecule, isoform A [Drosophila melanogaster] |
| XP_013776986.1 | 569 | LOC106461685 | eukaryota>metazoa | Limulus | polyphemus | PREDICTED: stromal interaction molecule homolog isoform X2 [Limulus polyphemus] |
| XP_004365004.1 | 563 | CAOG_00133 | eukaryota>ichthyosporea | Capsaspora | owczarzaki ATCC 30864 | hypothetical protein CAOG_00133 [Capsaspora owczarzaki ATCC 30864] |
| NP_996470.2 | 561 | C-Stim,CG9126,D-Stim,D-STIM,Dmel\CG9126,dstim,dStim,dSTIM,STIM,stim1,STIM1 | eukaryota>metazoa>hexapoda | Drosophila | melanogaster | stromal interaction molecule, isoform D [Drosophila melanogaster] |
| XP_011609392.1 | 559 | LOC101077194 | eukaryota>metazoa>chordata>vertebrata>actinopterygii | Takifugu | rubripes | PREDICTED: stromal interaction molecule 1-like isoform X5 [Takifugu rubripes] |
| CBY11086.1 | 545 | GSOID_T00014829001 | eukaryota>metazoa>chordata | Oikopleura | dioica | unnamed protein product [Oikopleura dioica] |
| NP_001264891.1 | 540 | D11S4896E,GOK,IMD10,STRMK,TAM,TAM1 | eukaryota>metazoa>chordata>vertebrata | Homo | sapiens | stromal interaction molecule 1 isoform 3 precursor [Homo sapiens] |
| XP_005579050.1 | 540 | STIM1 | eukaryota>metazoa>chordata>vertebrata | Macaca | fascicularis | PREDICTED: stromal interaction molecule 1 isoform X4 [Macaca fascicularis] |
| XP_006507600.1 | 540 | SIM | eukaryota>metazoa>chordata>vertebrata | Mus | musculus | PREDICTED: stromal interaction molecule 1 isoform X5 [Mus musculus] |
| XP_012359606.1 | 540 | STIM1 | eukaryota>metazoa>chordata>vertebrata | Nomascus | leucogenys | PREDICTED: stromal interaction molecule 1 isoform X3 [Nomascus leucogenys] |
| NP_741074.1 | 530 | stim-1 | eukaryota>metazoa>nematoda | Caenorhabditis | elegans | Stromal interaction molecule 1 [Caenorhabditis elegans] |
| XP_004994296.1 | 524 | PTSG_04978 | eukaryota>choanoflagellida | Salpingoeca | rosetta | hypothetical protein PTSG_04978 [Salpingoeca rosetta] |
| XP_006570264.1 | 520 | LOC411739 | eukaryota>metazoa>hexapoda | Apis | mellifera | PREDICTED: stromal interaction molecule homolog isoform X3 [Apis mellifera] |
| XP_018105968.1 | 516 | LOC108710082 | eukaryota>metazoa>chordata>vertebrata | Xenopus | laevis | PREDICTED: stromal interaction molecule 1-like isoform X4 [Xenopus laevis] |
| XP_018105969.1 | 515 | LOC108710082 | eukaryota>metazoa>chordata>vertebrata | Xenopus | laevis | PREDICTED: stromal interaction molecule 1-like isoform X5 [Xenopus laevis] |
| NP_741073.1 | 510 | stim-1 | eukaryota>metazoa>nematoda | Caenorhabditis | elegans | Stromal interaction molecule 1 [Caenorhabditis elegans] |
| EUB61676.1 | 506 | EGR_03490 | eukaryota>metazoa | Echinococcus | granulosus | Stromal interaction molecule [Echinococcus granulosus] |
| XP_014667541.1 | 501 | LOC106809096 | eukaryota>metazoa | Priapulus | caudatus | PREDICTED: stromal interaction molecule homolog [Priapulus caudatus] |
| XP_008186735.1 | 499 | LOC100166274 | eukaryota>metazoa>hexapoda | Acyrthosiphon | pisum | PREDICTED: stromal interaction molecule homolog isoform X3 [Acyrthosiphon pisum] |
| EFX72167.1 | 465 | DAPPUDRAFT_59440 | eukaryota>metazoa>crustacea | Daphnia | pulex | hypothetical protein DAPPUDRAFT_59440 [Daphnia pulex] |
| Caps1000012710 | 425 | Caps1000012710 | eukaryota>metazoa>annelida | Capitella | spI | >Caps1000012710 |
| XP_002407630.1 | 410 | IscW_ISCW017522 | eukaryota>metazoa | Ixodes | scapularis | conserved hypothetical protein, partial [Ixodes scapularis] |
| OAF68587.1 | 401 | A3Q56_03679 | eukaryota>metazoa | Intoshia | linei | hypothetical protein A3Q56_03679 [Intoshia linei] |
| XP_009055385.1 | 396 | LOTGIDRAFT_104732 | eukaryota>metazoa>mollusca | Lottia | gigantea | hypothetical protein LOTGIDRAFT_104732, partial [Lottia gigantea] |
| NP_001022916.1 | 395 | stim-1 | eukaryota>metazoa>nematoda | Caenorhabditis | elegans | Stromal interaction molecule 1 [Caenorhabditis elegans] |
| XP_001742986.1 | 386 | MONBRDRAFT_35825 | eukaryota>choanoflagellida | Monosiga | brevicollis MX1 | hypothetical protein [Monosiga brevicollis MX1] |
| XP_001641745.1 | 383 | NEMVEDRAFT_v1g31023 | eukaryota>metazoa>cnidaria | Nematostella | vectensis | predicted protein, partial [Nematostella vectensis] |
| XP_013792951.1 | 328 | LOC106476880 | eukaryota>metazoa | Limulus | polyphemus | PREDICTED: stromal interaction molecule 2-like, partial [Limulus polyphemus] |

U. ERp44

| | | | | | | |
|---|---|---|---|---|---|---|
| XP_004347444.1 | 541 | CAOG_04697 | eukaryota>ichthyosporea | Capsaspora | owczarzaki ATCC 30864 | thioredoxin domain-containing protein [Capsaspora owczarzaki ATCC 30864] |
| CBY18544.1 | 503 | GSOID_T00002411001 | eukaryota>metazoa>chordata | Oikopleura | dioica | unnamed protein product [Oikopleura dioica] |
| XP_001750772.1 | 499 | MONBRDRAFT_39286 | eukaryota>choanoflagellida | Monosiga | brevicollis MX1 | hypothetical protein [Monosiga brevicollis MX1] |
| ERG86510.1 | 473 | ASU_02091 | eukaryota>metazoa>nematoda | Ascaris | suum | thioredoxin domain-containing protein [Ascaris suum] |
| OQV11721.1 | 473 | BV898_13992 | eukaryota>metazoa | Hypsibius | dujardini | Endoplasmic reticulum resident protein 44 [Hypsibius dujardini] |
| OQV11722.1 | 468 | BV898_13992 | eukaryota>metazoa | Hypsibius | dujardini | Endoplasmic reticulum resident protein 44 [Hypsibius dujardini] |
| XP_004990284.1 | 452 | PTSG_08898 | eukaryota>choanoflagellida | Salpingoeca | rosetta | hypothetical protein PTSG_08898 [Salpingoeca rosetta] |
| XP_001893038.1 | 449 | Bm1_07840 | eukaryota>metazoa>nematoda | Brugia | malayi | Thioredoxin family protein [Brugia malayi] |
| EUB56965.1 | 442 | EGR_08202 | eukaryota>metazoa | Echinococcus | granulosus | Endoplasmic reticulum resident protein 44 [Echinococcus granulosus] |
| XP_003372150.1 | 440 | Tsp_11162 | eukaryota>metazoa>nematoda | Trichinella | spiralis | thioredoxin domain-containing protein 4 [Trichinella spiralis] |
| XP_002155688.1 | 439 | LOC100203097 | eukaryota>metazoa>cnidaria | Hydra | vulgaris | PREDICTED: endoplasmic reticulum resident protein 44-like [Hydra vulgaris] |
| XP_008179159.1 | 437 | LOC100163514 | eukaryota>metazoa>hexapoda | Acyrthosiphon | pisum | PREDICTED: endoplasmic reticulum resident protein 44 isoform X1 [Acyrthosiphon pisum] |
| NP_501357.3 | 434 | erp-44.3 | eukaryota>metazoa>nematoda | Caenorhabditis | elegans | Endoplasmic Reticulum Protein homolog [Caenorhabditis elegans] |
| XP_011455351.1 | 429 | LOC105347815 | eukaryota>metazoa>mollusca | Crassostrea | gigas | PREDICTED: endoplasmic reticulum resident protein 44 [Crassostrea gigas] |
| XP_001629332.1 | 428 | NEMVEDRAFT_v1g237233 | eukaryota>metazoa>cnidaria | Nematostella | vectensis | predicted protein [Nematostella vectensis] |
| XP_624571.2 | 427 | LOC552191 | eukaryota>metazoa>hexapoda | Apis | mellifera | PREDICTED: endoplasmic reticulum resident protein 44 isoform X1 [Apis mellifera] |
| XP_011150169.1 | 425 | LOC105189633 | eukaryota>metazoa>hexapoda | Harpegnathos | saltator | PREDICTED: endoplasmic reticulum resident protein 44 isoform X1 [Harpegnathos saltator] |
| XP_009860646.1 | 424 | LOC100176147 | eukaryota>metazoa>chordata | Ciona | intestinalis | PREDICTED: endoplasmic reticulum resident protein 44-like [Ciona intestinalis] |
| XP_018670204.1 | 424 | LOC100176147 | eukaryota>metazoa>chordata | Ciona | intestinalis | PREDICTED: endoplasmic reticulum resident protein 44-like [Ciona intestinalis] |
| EFX75490.1 | 421 | DAPPUDRAFT_306744 | eukaryota>metazoa>crustacea | Daphnia | pulex | hypothetical protein DAPPUDRAFT_306744 [Daphnia pulex] |
| XP_009016078.1 | 421 | HELRODRAFT_106306 | eukaryota>metazoa>annelida | Helobdella | robusta | hypothetical protein HELRODRAFT_106306 [Helobdella robusta] |
| XP_002736479.1 | 419 | LOC100366713 | eukaryota>metazoa>hemichordata | Saccoglossus | kowalevskii | PREDICTED: endoplasmic reticulum resident protein 44-like [Saccoglossus kowalevskii] |
| OAF69394.1 | 418 | A3Q56_02845 | eukaryota>metazoa | Intoshia | linei | hypothetical protein A3Q56_02845 [Intoshia linei] |
| XP_002605179.1 | 418 | BRAFLDRAFT_122706 | eukaryota>metazoa>chordata | Branchiostoma | floridae | hypothetical protein BRAFLDRAFT_122706 [Branchiostoma floridae] |
| CAG11934.1 | 416 | GSTEN:00034033:G:001 | eukaryota>metazoa>chordata>vertebrata>actinopterygii | Tetraodon | nigroviridis | unnamed protein product [Tetraodon nigroviridis] |
| CDS36497.1 | 413 | EmuJ_000359800 | eukaryota>metazoa | Echinococcus | multilocularis | Thioredoxin fold [Echinococcus multilocularis] |
| NP_501303.1 | 413 | erp-44.2 | eukaryota>metazoa>nematoda | Caenorhabditis | elegans | Thioredoxin domain-containing protein C06A6.5 [Caenorhabditis elegans] |
| XP_019855521.1 | 413 | LOC100640805 | eukaryota>metazoa | Amphimedon | queenslandica | PREDICTED: endoplasmic reticulum resident protein 44-like isoform X1 [Amphimedon queenslandica] |
| ERG79435.1 | 412 | ASU_13634 | eukaryota>metazoa>nematoda | Ascaris | suum | thioredoxin domain-containing protein [Ascaris suum] |
| NP_573111.1 | 412 | CG9912,Dmel\CG9911,LD41494p | eukaryota>metazoa>hexapoda | Drosophila | melanogaster | CG9911, isoform A [Drosophila melanogaster] |
| NP_727947.1 | 412 | CG9912,Dmel\CG9911,LD41494p | eukaryota>metazoa>hexapoda | Drosophila | melanogaster | CG9911, isoform B [Drosophila melanogaster] |
| NP_727948.1 | 412 | CG9912,Dmel\CG9911,LD41494p | eukaryota>metazoa>hexapoda | Drosophila | melanogaster | CG9911, isoform D [Drosophila melanogaster] |
| XP_001899062.1 | 412 | Bm1_38045 | eukaryota>metazoa>nematoda | Brugia | malayi | Thioredoxin family protein [Brugia malayi] |
| NP_001040641.1 | 411 | erp-44.1 | eukaryota>metazoa>nematoda | Caenorhabditis | elegans | Endoplasmic Reticulum Protein homolog [Caenorhabditis elegans] |
| XP_006558595.1 | 411 | LOC552191 | eukaryota>metazoa>hexapoda | Apis | mellifera | PREDICTED: endoplasmic reticulum resident protein 44 isoform X2 [Apis mellifera] |
| XP_011150170.1 | 411 | LOC105189633 | eukaryota>metazoa>hexapoda | Harpegnathos | saltator | PREDICTED: endoplasmic reticulum resident protein 44 isoform X2 [Harpegnathos saltator] |
| NP_649716.1 | 410 | Dmel\CG10029 | eukaryota>metazoa>hexapoda | Drosophila | melanogaster | uncharacterized protein Dmel_CG10029 [Drosophila melanogaster] |
| XP_004923220.1 | 410 | LOC101741420 | eukaryota>metazoa>hexapoda | Bombyx | mori | endoplasmic reticulum resident protein 44 isoform X2 [Bombyx mori] |
| XP_001951494.1 | 409 | LOC100158679 | eukaryota>metazoa>hexapoda | Acyrthosiphon | pisum | PREDICTED: endoplasmic reticulum resident protein 44 [Acyrthosiphon pisum] |
| XP_011150171.1 | 409 | LOC105189633 | eukaryota>metazoa>hexapoda | Harpegnathos | saltator | PREDICTED: endoplasmic reticulum resident protein 44 isoform X3 [Harpegnathos saltator] |
| XP_011150172.1 | 409 | LOC105189633 | eukaryota>metazoa>hexapoda | Harpegnathos | saltator | PREDICTED: endoplasmic reticulum resident protein 44 isoform X4 [Harpegnathos saltator] |
| XP_791945.2 | 409 | LOC587099 | eukaryota>metazoa>echinodermata | Strongylocentrotus | purpuratus | PREDICTED: endoplasmic reticulum resident protein 44 [Strongylocentrotus purpuratus] |
| NP_957186.1 | 408 | txndc4,zgc:77883 | eukaryota>metazoa>chordata>vertebrata>actinopterygii | Danio | rerio | endoplasmic reticulum resident protein 44 [Danio rerio] |
| XP_008179161.1 | 408 | LOC100163514 | eukaryota>metazoa>hexapoda | Acyrthosiphon | pisum | PREDICTED: endoplasmic reticulum resident protein 44 isoform X2 [Acyrthosiphon pisum] |
| XP_011603971.1 | 408 | erp44 | eukaryota>metazoa>chordata>vertebrata>actinopterygii | Takifugu | rubripes | PREDICTED: endoplasmic reticulum resident protein 44 [Takifugu rubripes] |
| XP_017308738.1 | 408 | erp44 | eukaryota>metazoa>chordata>vertebrata>actinopterygii | Ictalurus | punctatus | PREDICTED: endoplasmic reticulum resident protein 44 isoform X1 [Ictalurus punctatus] |
| XP_014663701.1 | 407 | LOC106806318 | eukaryota>metazoa | Priapulus | caudatus | PREDICTED: endoplasmic reticulum resident protein 44-like isoform X1 [Priapulus caudatus] |
| XP_017308739.1 | 407 | erp44 | eukaryota>metazoa>chordata>vertebrata>actinopterygii | Ictalurus | punctatus | PREDICTED: endoplasmic reticulum resident protein 44 isoform X2 [Ictalurus punctatus] |
| NP_001180332.1 | 406 | erp44,txndc4 | eukaryota>metazoa>chordata>vertebrata | Xenopus | laevis | endoplasmic reticulum protein 44 L homeolog precursor [Xenopus laevis] |
| NP_001271680.1 | 406 | ERP44 | eukaryota>metazoa>chordata>vertebrata | Macaca | fascicularis | endoplasmic reticulum resident protein 44 precursor [Macaca fascicularis] |
| NP_055866.1 | 406 | PDIA10,TXNDC4 | eukaryota>metazoa>chordata>vertebrata | Homo | sapiens | endoplasmic reticulum resident protein 44 precursor [Homo sapiens] |
| NP_083848.1 | 406 | 1110001E24Rik,AI849526,AL033348,Txndc4 | eukaryota>metazoa>chordata>vertebrata | Mus | musculus | endoplasmic reticulum resident protein 44 precursor [Mus musculus] |
| XP_003260331.1 | 406 | ERP44 | eukaryota>metazoa>chordata>vertebrata | Nomascus | leucogenys | PREDICTED: endoplasmic reticulum resident protein 44 [Nomascus leucogenys] |
| XP_013783081.1 | 406 | LOC106467291 | eukaryota>metazoa | Limulus | polyphemus | PREDICTED: endoplasmic reticulum resident protein 44-like [Limulus polyphemus] |
| XP_419079.1 | 406 | ERP44 | eukaryota>metazoa>chordata>vertebrata | Gallus | gallus | PREDICTED: endoplasmic reticulum resident protein 44 [Gallus gallus] |
| XP_971182.1 | 406 | LOC659817 | eukaryota>metazoa>hexapoda | Tribolium | castaneum | PREDICTED: endoplasmic reticulum resident protein 44 [Tribolium castaneum] |
| XP_014663709.1 | 405 | LOC106806318 | eukaryota>metazoa | Priapulus | caudatus | PREDICTED: endoplasmic reticulum resident protein 44-like isoform X2 [Priapulus caudatus] |
| XP_014663719.1 | 405 | LOC106806318 | eukaryota>metazoa | Priapulus | caudatus | PREDICTED: endoplasmic reticulum resident protein 44-like isoform X2 [Priapulus caudatus] |
| XP_013791647.1 | 403 | LOC106475511 | eukaryota>metazoa | Limulus | polyphemus | PREDICTED: endoplasmic reticulum resident protein 44-like [Limulus polyphemus] |
| Caps1000012460 | 403 | Caps1000012460 | eukaryota>metazoa>annelida | Capitella | spI | >Caps1000012460 |
| XP_013387938.1 | 402 | LOC106157020 | eukaryota>metazoa | Lingula | anatina | PREDICTED: endoplasmic reticulum resident protein 44-like [Lingula anatina] |
| XP_019855522.1 | 394 | LOC100640805 | eukaryota>metazoa | Amphimedon | queenslandica | PREDICTED: endoplasmic reticulum resident protein 44-like isoform X2 [Amphimedon queenslandica] |
| NP_001040640.1 | 393 | erp-44.1 | eukaryota>metazoa>nematoda | Caenorhabditis | elegans | Endoplasmic Reticulum Protein homolog [Caenorhabditis elegans] |
| XP_008178361.1 | 392 | LOC100572906 | eukaryota>metazoa>hexapoda | Acyrthosiphon | pisum | PREDICTED: endoplasmic reticulum resident protein 44-like isoform X1 [Acyrthosiphon pisum] |
| Smar1000006250 | 391 | Smar1000006250 | eukaryota>metazoa>arthropoda>myriapoda | Strigamia | maritima | >Smar1000006250 |
| XP_013787956.1 | 390 | LOC106471878 | eukaryota>metazoa | Limulus | polyphemus | PREDICTED: endoplasmic reticulum resident protein 44-like, partial [Limulus polyphemus] |
| XP_002110325.1 | 382 | TRIADDRAFT_37324 | eukaryota>metazoa>placozoa | Trichoplax | adhaerens | hypothetical protein TRIADDRAFT_37324 [Trichoplax adhaerens] |
| XP_013387936.1 | 377 | LOC106157019 | eukaryota>metazoa | Lingula | anatina | PREDICTED: endoplasmic reticulum resident protein 44-like [Lingula anatina] |
| XP_015761302.1 | 376 | LOC107340462 | eukaryota>metazoa>cnidaria | Acropora | digitifera | PREDICTED: endoplasmic reticulum resident protein 44-like [Acropora digitifera] |
| NP_727949.1 | 375 | CG9912,Dmel\CG9911,LD41494p | eukaryota>metazoa>hexapoda | Drosophila | melanogaster | CG9911, isoform C [Drosophila melanogaster] |
| XP_009064776.1 | 366 | LOTGIDRAFT_131935 | eukaryota>metazoa>mollusca | Lottia | gigantea | hypothetical protein LOTGIDRAFT_131935, partial [Lottia gigantea] |
| XP_014159730.1 | 355 | SARC_02011 | eukaryota>ichthyosporea | Sphaeroforma | arctica JP610 | hypothetical protein SARC_02011 [Sphaeroforma arctica JP610] |
| XP_016656180.1 | 345 | LOC100572906 | eukaryota>metazoa>hexapoda | Acyrthosiphon | pisum | PREDICTED: endoplasmic reticulum resident protein 44-like isoform X2 [Acyrthosiphon pisum] |
| XP_005581253.1 | 344 | ERP44 | eukaryota>metazoa>chordata>vertebrata | Macaca | fascicularis | PREDICTED: endoplasmic reticulum resident protein 44 isoform X1 [Macaca fascicularis] |
| XP_015291957.1 | 344 | ERP44 | eukaryota>metazoa>chordata>vertebrata | Macaca | fascicularis | PREDICTED: endoplasmic reticulum resident protein 44 isoform X1 [Macaca fascicularis] |
| Smed1000009550 | 332 | Smed1000009550 | eukaryota>metazoa>platyhelminthe | Schmidtea | mediterranea | >Smed1000009550 |
| XP_006538404.1 | 314 | 1110001E24Rik,AI849526,AL033348,Txndc4 | eukaryota>metazoa>chordata>vertebrata | Mus | musculus | PREDICTED: endoplasmic reticulum resident protein 44 isoform X1 [Mus musculus] |
| XP_014787973.1 | 289 | LOC106881956 | eukaryota>metazoa>mollusca | Octopus | bimaculoides | PREDICTED: endoplasmic reticulum resident protein 44-like, partial [Octopus bimaculoides] |
| XP_018653631.1 | 230 | Smp_144840 | eukaryota>metazoa | Schistosoma | mansoni | putative thioredoxin-like protein [Schistosoma mansoni] |
| Smed1000010608 | 205 | Smed1000010608 | eukaryota>metazoa>platyhelminthe | Schmidtea | mediterranea | >Smed1000010608 |

V. RyR

| | | | | | | |
|---|---|---|---|---|---|---|
| XP_004342590.1 | 6625 | CAOG_07989 | eukaryota>ichthyosporea | Capsaspora | owczarzaki ATCC 30864 | ryanodine receptor [Capsaspora owczarzaki ATCC 30864] |
| XP_011670306.1 | 5731 | LOC580223 | eukaryota>metazoa>echinodermata | Strongylocentrotus | purpuratus | PREDICTED: LOW QUALITY PROTEIN: ryanodine receptor 2 [Strongylocentrotus purpuratus] |
| XP_009173549.1 | 5727 | T265_09254 | eukaryota>metazoa | Opisthorchis | viverrini | hypothetical protein T265_09254 [Opisthorchis viverrini] |
| NP_001256077.1 | 5387 | unc-68 | eukaryota>metazoa>nematoda | Caenorhabditis | elegans | Uncharacterized protein CELE_K11C4.5 [Caenorhabditis elegans] |
| NP_001256076.1 | 5363 | unc-68 | eukaryota>metazoa>nematoda | Caenorhabditis | elegans | Uncharacterized protein CELE_K11C4.5 [Caenorhabditis elegans] |
| XP_004993045.1 | 5340 | PTSG_06153 | eukaryota>choanoflagellida | Salpingoeca | rosetta | ryanodine receptor type 1 [Salpingoeca rosetta] |
| NP_001256074.1 | 5333 | unc-68 | eukaryota>metazoa>nematoda | Caenorhabditis | elegans | Uncharacterized protein CELE_K11C4.5 [Caenorhabditis elegans] |
| NP_001256075.2 | 5273 | unc-68 | eukaryota>metazoa>nematoda | Caenorhabditis | elegans | Uncharacterized protein CELE_K11C4.5 [Caenorhabditis elegans] |
| XP_014787737.1 | 5242 | LOC106881762 | eukaryota>metazoa>mollusca | Octopus | bimaculoides | PREDICTED: ryanodine receptor-like isoform X1 [Octopus bimaculoides] |
| XP_014787738.1 | 5242 | LOC106881762 | eukaryota>metazoa>mollusca | Octopus | bimaculoides | PREDICTED: ryanodine receptor-like isoform X2 [Octopus bimaculoides] |
| XP_014787740.1 | 5241 | LOC106881762 | eukaryota>metazoa>mollusca | Octopus | bimaculoides | PREDICTED: ryanodine receptor-like isoform X3 [Octopus bimaculoides] |
| XP_014787741.1 | 5241 | LOC106881762 | eukaryota>metazoa>mollusca | Octopus | bimaculoides | PREDICTED: ryanodine receptor-like isoform X4 [Octopus bimaculoides] |
| XP_014787742.1 | 5236 | LOC106881762 | eukaryota>metazoa>mollusca | Octopus | bimaculoides | PREDICTED: ryanodine receptor-like isoform X5 [Octopus bimaculoides] |
| XP_014787743.1 | 5235 | LOC106881762 | eukaryota>metazoa>mollusca | Octopus | bimaculoides | PREDICTED: ryanodine receptor-like isoform X6 [Octopus bimaculoides] |
| XP_014787744.1 | 5235 | LOC106881762 | eukaryota>metazoa>mollusca | Octopus | bimaculoides | PREDICTED: ryanodine receptor-like isoform X7 [Octopus bimaculoides] |
| XP_014787745.1 | 5235 | LOC106881762 | eukaryota>metazoa>mollusca | Octopus | bimaculoides | PREDICTED: ryanodine receptor-like isoform X8 [Octopus bimaculoides] |
| XP_014787746.1 | 5233 | LOC106881762 | eukaryota>metazoa>mollusca | Octopus | bimaculoides | PREDICTED: ryanodine receptor 2-like isoform X9 [Octopus bimaculoides] |
| XP_014787753.1 | 5229 | LOC106881762 | eukaryota>metazoa>mollusca | Octopus | bimaculoides | PREDICTED: ryanodine receptor-like isoform X16 [Octopus bimaculoides] |
| XP_014787747.1 | 5226 | LOC106881762 | eukaryota>metazoa>mollusca | Octopus | bimaculoides | PREDICTED: ryanodine receptor 2-like isoform X10 [Octopus bimaculoides] |
| XP_014787748.1 | 5223 | LOC106881762 | eukaryota>metazoa>mollusca | Octopus | bimaculoides | PREDICTED: ryanodine receptor-like isoform X11 [Octopus bimaculoides] |
| XP_014787754.1 | 5222 | LOC106881762 | eukaryota>metazoa>mollusca | Octopus | bimaculoides | PREDICTED: ryanodine receptor-like isoform X17 [Octopus bimaculoides] |
| XP_014787751.1 | 5213 | LOC106881762 | eukaryota>metazoa>mollusca | Octopus | bimaculoides | PREDICTED: ryanodine receptor-like isoform X14 [Octopus bimaculoides] |
| XP_014787752.1 | 5206 | LOC106881762 | eukaryota>metazoa>mollusca | Octopus | bimaculoides | PREDICTED: ryanodine receptor-like isoform X15 [Octopus bimaculoides] |
| XP_014787749.1 | 5187 | LOC106881762 | eukaryota>metazoa>mollusca | Octopus | bimaculoides | PREDICTED: ryanodine receptor-like isoform X12 [Octopus bimaculoides] |
| XP_006569086.1 | 5152 | LOC408680 | eukaryota>metazoa>hexapoda | Apis | mellifera | PREDICTED: ryanodine receptor isoform X1 [Apis mellifera] |
| XP_006569087.1 | 5152 | LOC408680 | eukaryota>metazoa>hexapoda | Apis | mellifera | PREDICTED: ryanodine receptor isoform X2 [Apis mellifera] |
| XP_016766251.1 | 5152 | LOC408680 | eukaryota>metazoa>hexapoda | Apis | mellifera | PREDICTED: ryanodine receptor isoform X1 [Apis mellifera] |
| XP_006569088.1 | 5151 | LOC408680 | eukaryota>metazoa>hexapoda | Apis | mellifera | PREDICTED: ryanodine receptor isoform X4 [Apis mellifera] |
| XP_006569089.1 | 5151 | LOC408680 | eukaryota>metazoa>hexapoda | Apis | mellifera | PREDICTED: ryanodine receptor isoform X3 [Apis mellifera] |
| XP_006569090.1 | 5149 | LOC408680 | eukaryota>metazoa>hexapoda | Apis | mellifera | PREDICTED: ryanodine receptor isoform X5 [Apis mellifera] |
| XP_013775910.1 | 5149 | LOC106460719 | eukaryota>metazoa | Limulus | polyphemus | PREDICTED: ryanodine receptor 44F-like [Limulus polyphemus] |
| XP_006569091.1 | 5147 | LOC408680 | eukaryota>metazoa>hexapoda | Apis | mellifera | PREDICTED: ryanodine receptor isoform X6 [Apis mellifera] |
| XP_006569092.1 | 5147 | LOC408680 | eukaryota>metazoa>hexapoda | Apis | mellifera | PREDICTED: ryanodine receptor isoform X7 [Apis mellifera] |
| XP_006569093.1 | 5147 | LOC408680 | eukaryota>metazoa>hexapoda | Apis | mellifera | PREDICTED: ryanodine receptor isoform X8 [Apis mellifera] |
| XP_006569094.1 | 5145 | LOC408680 | eukaryota>metazoa>hexapoda | Apis | mellifera | PREDICTED: ryanodine receptor isoform X9 [Apis mellifera] |
| XP_002735182.2 | 5145 | RYR | eukaryota>metazoa>hemichordata | Saccoglossus | kowalevskii | PREDICTED: LOW QUALITY PROTEIN: ryanodine receptor 2 [Saccoglossus kowalevskii] |
| XP_006569095.1 | 5144 | LOC408680 | eukaryota>metazoa>hexapoda | Apis | mellifera | PREDICTED: ryanodine receptor isoform X10 [Apis mellifera] |
| XP_006569096.1 | 5144 | LOC408680 | eukaryota>metazoa>hexapoda | Apis | mellifera | PREDICTED: ryanodine receptor isoform X11 [Apis mellifera] |
| XP_006569097.1 | 5143 | LOC408680 | eukaryota>metazoa>hexapoda | Apis | mellifera | PREDICTED: ryanodine receptor isoform X12 [Apis mellifera] |
| XP_006569098.1 | 5142 | LOC408680 | eukaryota>metazoa>hexapoda | Apis | mellifera | PREDICTED: ryanodine receptor isoform X13 [Apis mellifera] |
| XP_012544748.1 | 5142 | LOC101739070 | eukaryota>metazoa>hexapoda | Bombyx | mori | ryanodine receptor isoform X1 [Bombyx mori] |
| XP_012544749.1 | 5142 | LOC101739070 | eukaryota>metazoa>hexapoda | Bombyx | mori | PREDICTED: ryanodine receptor 44F isoform X1 [Bombyx mori] |
| XP_012544750.1 | 5142 | LOC101739070 | eukaryota>metazoa>hexapoda | Bombyx | mori | PREDICTED: ryanodine receptor 44F isoform X2 [Bombyx mori] |
| XP_006569099.1 | 5140 | LOC408680 | eukaryota>metazoa>hexapoda | Apis | mellifera | PREDICTED: ryanodine receptor isoform X14 [Apis mellifera] |
| XP_012544751.1 | 5140 | LOC101739070 | eukaryota>metazoa>hexapoda | Bombyx | mori | ryanodine receptor isoform X2 [Bombyx mori] |
| XP_006569100.1 | 5137 | LOC408680 | eukaryota>metazoa>hexapoda | Apis | mellifera | PREDICTED: ryanodine receptor isoform X17 [Apis mellifera] |
| XP_006569101.1 | 5137 | LOC408680 | eukaryota>metazoa>hexapoda | Apis | mellifera | PREDICTED: ryanodine receptor isoform X16 [Apis mellifera] |
| XP_012544752.1 | 5137 | LOC101739070 | eukaryota>metazoa>hexapoda | Bombyx | mori | PREDICTED: ryanodine receptor 44F isoform X4 [Bombyx mori] |
| XP_012544753.1 | 5137 | LOC101739070 | eukaryota>metazoa>hexapoda | Bombyx | mori | PREDICTED: ryanodine receptor 44F isoform X5 [Bombyx mori] |
| XP_016766252.1 | 5137 | LOC408680 | eukaryota>metazoa>hexapoda | Apis | mellifera | PREDICTED: ryanodine receptor isoform X15 [Apis mellifera] |
| XP_012544754.1 | 5136 | LOC101739070 | eukaryota>metazoa>hexapoda | Bombyx | mori | PREDICTED: ryanodine receptor 44F isoform X6 [Bombyx mori] |
| XP_012544771.1 | 5136 | LOC101739070 | eukaryota>metazoa>hexapoda | Bombyx | mori | ryanodine receptor isoform X15 [Bombyx mori] |
| NP_001246211.1 | 5134 | CG10844,D-RyR,Dmel\CG10844,DRR,dry,DRY,dRyr,dRyR,dya,l(2)k00424,l(2)k04913,Rya-44F,rya-r44F,Rya-r44F,Rya-R44F,Rya-r76CD,ryr,Ryr,RYR,RyRs | eukaryota>metazoa>hexapoda | Drosophila | melanogaster | ryanodine receptor, isoform I [Drosophila melanogaster] |
| XP_012544756.1 | 5134 | LOC101739070 | eukaryota>metazoa>hexapoda | Bombyx | mori | PREDICTED: ryanodine receptor 44F isoform X7 [Bombyx mori] |
| XP_012544757.1 | 5132 | LOC101739070 | eukaryota>metazoa>hexapoda | Bombyx | mori | ryanodine receptor isoform X9 [Bombyx mori] |
| XP_016766253.1 | 5132 | LOC408680 | eukaryota>metazoa>hexapoda | Apis | mellifera | PREDICTED: ryanodine receptor isoform X18 [Apis mellifera] |
| XP_012544758.1 | 5131 | LOC101739070 | eukaryota>metazoa>hexapoda | Bombyx | mori | PREDICTED: ryanodine receptor 44F isoform X9 [Bombyx mori] |
| XP_012544769.1 | 5131 | LOC101739070 | eukaryota>metazoa>hexapoda | Bombyx | mori | ryanodine receptor isoform X13 [Bombyx mori] |
| NP_001246210.1 | 5130 | CG10844,D-RyR,Dmel\CG10844,DRR,dry,DRY,dRyr,dRyR,dya,l(2)k00424,l(2)k04913,Rya-44F,rya-r44F,Rya-r44F,Rya-R44F,Rya-r76CD,ryr,Ryr,RYR,RyRs | eukaryota>metazoa>hexapoda | Drosophila | melanogaster | ryanodine receptor, isoform H [Drosophila melanogaster] |
| XP_012544764.1 | 5130 | LOC101739070 | eukaryota>metazoa>hexapoda | Bombyx | mori | ryanodine receptor isoform X5 [Bombyx mori] |
| XP_012544759.1 | 5129 | LOC101739070 | eukaryota>metazoa>hexapoda | Bombyx | mori | ryanodine receptor isoform X3 [Bombyx mori] |
| XP_012544772.1 | 5129 | LOC101739070 | eukaryota>metazoa>hexapoda | Bombyx | mori | PREDICTED: ryanodine receptor 44F isoform X23 [Bombyx mori] |
| XP_019699201.1 | 5129 | LOC105188290 | eukaryota>metazoa>hexapoda | Harpegnathos | saltator | PREDICTED: LOW QUALITY PROTEIN: ryanodine receptor [Harpegnathos saltator] |
| NP_476991.1 | 5127 | CG10844,D-RyR,Dmel\CG10844,DRR,dry,DRY,dRyr,dRyR,dya,l(2)k00424,l(2)k04913,Rya-44F,rya-r44F,Rya-r44F,Rya-R44F,Rya-r76CD,ryr,Ryr,RYR,RyRs | eukaryota>metazoa>hexapoda | Drosophila | melanogaster | ryanodine receptor, isoform A [Drosophila melanogaster] |
| NP_476993.1 | 5127 | CG10844,D-RyR,Dmel\CG10844,DRR,dry,DRY,dRyr,dRyR,dya,l(2)k00424,l(2)k04913,Rya-44F,rya-r44F,Rya-r44F,Rya-R44F,Rya-r76CD,ryr,Ryr,RYR,RyRs | eukaryota>metazoa>hexapoda | Drosophila | melanogaster | ryanodine receptor, isoform C [Drosophila melanogaster] |
| XP_012544770.1 | 5127 | LOC101739070 | eukaryota>metazoa>hexapoda | Bombyx | mori | ryanodine receptor isoform X14 [Bombyx mori] |
| XP_012544765.1 | 5125 | LOC101739070 | eukaryota>metazoa>hexapoda | Bombyx | mori | ryanodine receptor isoform X6 [Bombyx mori] |
| XP_012544760.1 | 5124 | LOC101739070 | eukaryota>metazoa>hexapoda | Bombyx | mori | PREDICTED: ryanodine receptor 44F isoform X11 [Bombyx mori] |
| XP_012544768.1 | 5124 | LOC101739070 | eukaryota>metazoa>hexapoda | Bombyx | mori | ryanodine receptor isoform X12 [Bombyx mori] |
| NP_001246209.1 | 5123 | CG10844,D-RyR,Dmel\CG10844,DRR,dry,DRY,dRyr,dRyR,dya,l(2)k00424,l(2)k04913,Rya-44F,rya-r44F,Rya-r44F,Rya-R44F,Rya-r76CD,ryr,Ryr,RYR,RyRs | eukaryota>metazoa>hexapoda | Drosophila | melanogaster | ryanodine receptor, isoform G [Drosophila melanogaster] |
| XP_012544761.1 | 5123 | LOC101739070 | eukaryota>metazoa>hexapoda | Bombyx | mori | ryanodine receptor isoform X4 [Bombyx mori] |
| XP_006569104.1 | 5122 | LOC408680 | eukaryota>metazoa>hexapoda | Apis | mellifera | PREDICTED: ryanodine receptor isoform X19 [Apis mellifera] |
| EFX89429.1 | 5119 | DAPPUDRAFT_310496 | eukaryota>metazoa>crustacea | Daphnia | pulex | hypothetical protein DAPPUDRAFT_310496 [Daphnia pulex] |
| NP_001246208.1 | 5119 | CG10844,D-RyR,Dmel\CG10844,DRR,dry,DRY,dRyr,dRyR,dya,l(2)k00424,l(2)k04913,Rya-44F,rya-r44F,Rya-r44F,Rya-R44F,Rya-r76CD,ryr,Ryr,RYR,RyRs | eukaryota>metazoa>hexapoda | Drosophila | melanogaster | ryanodine receptor, isoform F [Drosophila melanogaster] |
| XP_006569110.1 | 5119 | LOC408680 | eukaryota>metazoa>hexapoda | Apis | mellifera | PREDICTED: ryanodine receptor isoform X26 [Apis mellifera] |
| NP_001246207.1 | 5117 | CG10844,D-RyR,Dmel\CG10844,DRR,dry,DRY,dRyr,dRyR,dya,l(2)k00424,l(2)k04913,Rya-44F,rya-r44F,Rya-r44F,Rya-R44F,Rya-r76CD,ryr,Ryr,RYR,RyRs | eukaryota>metazoa>hexapoda | Drosophila | melanogaster | ryanodine receptor, isoform E [Drosophila melanogaster] |
| XP_006569105.1 | 5117 | LOC408680 | eukaryota>metazoa>hexapoda | Apis | mellifera | PREDICTED: ryanodine receptor isoform X20 [Apis mellifera] |
| XP_006569108.1 | 5117 | LOC408680 | eukaryota>metazoa>hexapoda | Apis | mellifera | PREDICTED: ryanodine receptor isoform X24 [Apis mellifera] |
| XP_006569109.1 | 5117 | LOC408680 | eukaryota>metazoa>hexapoda | Apis | mellifera | PREDICTED: ryanodine receptor isoform X25 [Apis mellifera] |
| XP_012544762.1 | 5116 | LOC101739070 | eukaryota>metazoa>hexapoda | Bombyx | mori | PREDICTED: ryanodine receptor 44F isoform X13 [Bombyx mori] |
| NP_001246206.1 | 5115 | CG10844,D-RyR,Dmel\CG10844,DRR,dry,DRY,dRyr,dRyR,dya,l(2)k00424,l(2)k04913,Rya-44F,rya-r44F,Rya-r44F,Rya-R44F,Rya-r76CD,ryr,Ryr,RYR,RyRs | eukaryota>metazoa>hexapoda | Drosophila | melanogaster | ryanodine receptor, isoform J [Drosophila melanogaster] |
| XP_012544763.1 | 5115 | LOC101739070 | eukaryota>metazoa>hexapoda | Bombyx | mori | ryanodine receptor isoform X7 [Bombyx mori] |
| NP_476992.1 | 5113 | CG10844,D-RyR,Dmel\CG10844,DRR,dry,DRY,dRyr,dRyR,dya,l(2)k00424,l(2)k04913,Rya-44F,rya-r44F,Rya-r44F,Rya-R44F,Rya-r76CD,ryr,Ryr,RYR,RyRs | eukaryota>metazoa>hexapoda | Drosophila | melanogaster | ryanodine receptor, isoform B [Drosophila melanogaster] |
| NP_476994.1 | 5113 | CG10844,D-RyR,Dmel\CG10844,DRR,dry,DRY,dRyr,dRyR,dya,l(2)k00424,l(2)k04913,Rya-44F,rya-r44F,Rya-r44F,Rya-R44F,Rya-r76CD,ryr,Ryr,RYR,RyRs | eukaryota>metazoa>hexapoda | Drosophila | melanogaster | ryanodine receptor, isoform D [Drosophila melanogaster] |
| XP_012544766.1 | 5110 | LOC101739070 | eukaryota>metazoa>hexapoda | Bombyx | mori | ryanodine receptor isoform X8 [Bombyx mori] |
| XP_017207446.1 | 5109 | im:7137738,si:dkey-250i19.1 | eukaryota>metazoa>chordata>vertebrata>actinopterygii | Danio | rerio | ryanodine receptor 1 isoform X1 [Danio rerio] |
| XP_012544767.1 | 5108 | LOC101739070 | eukaryota>metazoa>hexapoda | Bombyx | mori | ryanodine receptor isoform X11 [Bombyx mori] |
| XP_006569106.1 | 5107 | LOC408680 | eukaryota>metazoa>hexapoda | Apis | mellifera | PREDICTED: ryanodine receptor isoform X22 [Apis mellifera] |
| XP_017207447.1 | 5104 | im:7137738,si:dkey-250i19.1 | eukaryota>metazoa>chordata>vertebrata>actinopterygii | Danio | rerio | ryanodine receptor 1 isoform X2 [Danio rerio] |
| XP_006569107.1 | 5102 | LOC408680 | eukaryota>metazoa>hexapoda | Apis | mellifera | PREDICTED: ryanodine receptor isoform X23 [Apis mellifera] |
| XP_392217.5 | 5102 | LOC408680 | eukaryota>metazoa>hexapoda | Apis | mellifera | PREDICTED: ryanodine receptor isoform X21 [Apis mellifera] |
| XP_016661950.1 | 5101 | LOC100167965 | eukaryota>metazoa>hexapoda | Acyrthosiphon | pisum | PREDICTED: ryanodine receptor [Acyrthosiphon pisum] |
| XP_017207448.1 | 5098 | im:7137738,si:dkey-250i19.1 | eukaryota>metazoa>chordata>vertebrata>actinopterygii | Danio | rerio | ryanodine receptor 1 isoform X6 [Danio rerio] |
| XP_017322026.1 | 5093 | LOC108264731 | eukaryota>metazoa>chordata>vertebrata>actinopterygii | Ictalurus | punctatus | PREDICTED: ryanodine receptor 1-like isoform X1 [Ictalurus punctatus] |
| XP_017207449.1 | 5089 | im:7137738,si:dkey-250i19.1 | eukaryota>metazoa>chordata>vertebrata>actinopterygii | Danio | rerio | ryanodine receptor 1 isoform X8 [Danio rerio] |
| XP_017322029.1 | 5088 | LOC108264731 | eukaryota>metazoa>chordata>vertebrata>actinopterygii | Ictalurus | punctatus | PREDICTED: ryanodine receptor 1-like isoform X2 [Ictalurus punctatus] |
| XP_017322030.1 | 5087 | LOC108264731 | eukaryota>metazoa>chordata>vertebrata>actinopterygii | Ictalurus | punctatus | PREDICTED: ryanodine receptor 1-like isoform X3 [Ictalurus punctatus] |
| XP_017322031.1 | 5087 | LOC108264731 | eukaryota>metazoa>chordata>vertebrata>actinopterygii | Ictalurus | punctatus | PREDICTED: ryanodine receptor 1-like isoform X4 [Ictalurus punctatus] |
| XP_017322032.1 | 5086 | LOC108264731 | eukaryota>metazoa>chordata>vertebrata>actinopterygii | Ictalurus | punctatus | PREDICTED: ryanodine receptor 1-like isoform X5 [Ictalurus punctatus] |
| XP_017322033.1 | 5083 | LOC108264731 | eukaryota>metazoa>chordata>vertebrata>actinopterygii | Ictalurus | punctatus | PREDICTED: ryanodine receptor 1-like isoform X6 [Ictalurus punctatus] |
| XP_017322034.1 | 5082 | LOC108264731 | eukaryota>metazoa>chordata>vertebrata>actinopterygii | Ictalurus | punctatus | PREDICTED: ryanodine receptor 1-like isoform X7 [Ictalurus punctatus] |
| XP_017322035.1 | 5081 | LOC108264731 | eukaryota>metazoa>chordata>vertebrata>actinopterygii | Ictalurus | punctatus | PREDICTED: ryanodine receptor 1-like isoform X8 [Ictalurus punctatus] |
| XP_017322036.1 | 5081 | LOC108264731 | eukaryota>metazoa>chordata>vertebrata>actinopterygii | Ictalurus | punctatus | PREDICTED: ryanodine receptor 1-like isoform X9 [Ictalurus punctatus] |
| XP_011609906.1 | 5080 | LOC101072939 | eukaryota>metazoa>chordata>vertebrata>actinopterygii | Takifugu | rubripes | PREDICTED: ryanodine receptor 1-like [Takifugu rubripes] |
| XP_017344262.1 | 5078 | LOC108276801 | eukaryota>metazoa>chordata>vertebrata>actinopterygii | Ictalurus | punctatus | PREDICTED: ryanodine receptor 1-like isoform X1 [Ictalurus punctatus] |
| XP_017344263.1 | 5077 | LOC108276801 | eukaryota>metazoa>chordata>vertebrata>actinopterygii | Ictalurus | punctatus | PREDICTED: ryanodine receptor 1-like isoform X2 [Ictalurus punctatus] |
| NP_001096041.1 | 5076 | im:7137738,si:dkey-250i19.1 | eukaryota>metazoa>chordata>vertebrata>actinopterygii | Danio | rerio | ryanodine receptor 1 [Danio rerio] |
| XP_017207451.1 | 5076 | im:7137738,si:dkey-250i19.1 | eukaryota>metazoa>chordata>vertebrata>actinopterygii | Danio | rerio | ryanodine receptor 1 isoform X10 [Danio rerio] |
| XP_017344264.1 | 5073 | LOC108276801 | eukaryota>metazoa>chordata>vertebrata>actinopterygii | Ictalurus | punctatus | PREDICTED: ryanodine receptor 1-like isoform X3 [Ictalurus punctatus] |
| XP_015833833.1 | 5073 | LOC655265 | eukaryota>metazoa>hexapoda | Tribolium | castaneum | PREDICTED: LOW QUALITY PROTEIN: ryanodine receptor [Tribolium castaneum] |
| XP_017207450.1 | 5072 | im:7137738,si:dkey-250i19.1 | eukaryota>metazoa>chordata>vertebrata>actinopterygii | Danio | rerio | ryanodine receptor 1 isoform X9 [Danio rerio] |
| XP_017344265.1 | 5072 | LOC108276801 | eukaryota>metazoa>chordata>vertebrata>actinopterygii | Ictalurus | punctatus | PREDICTED: ryanodine receptor 1-like isoform X4 [Ictalurus punctatus] |
| XP_017344266.1 | 5072 | LOC108276801 | eukaryota>metazoa>chordata>vertebrata>actinopterygii | Ictalurus | punctatus | PREDICTED: ryanodine receptor 1-like isoform X5 [Ictalurus punctatus] |
| XP_001900491.1 | 5072 | Bm1_45145 | eukaryota>metazoa>nematoda | Brugia | malayi | Ryanodine Receptor TM 4-6 family protein [Brugia malayi] |
| XP_017207452.1 | 5071 | im:7137738,si:dkey-250i19.1 | eukaryota>metazoa>chordata>vertebrata>actinopterygii | Danio | rerio | ryanodine receptor 1 isoform X11 [Danio rerio] |
| XP_017322038.1 | 5070 | LOC108264731 | eukaryota>metazoa>chordata>vertebrata>actinopterygii | Ictalurus | punctatus | PREDICTED: ryanodine receptor 1-like isoform X11 [Ictalurus punctatus] |
| XP_017322037.1 | 5068 | LOC108264731 | eukaryota>metazoa>chordata>vertebrata>actinopterygii | Ictalurus | punctatus | PREDICTED: ryanodine receptor 1-like isoform X10 [Ictalurus punctatus] |
| XP_017344267.1 | 5067 | LOC108276801 | eukaryota>metazoa>chordata>vertebrata>actinopterygii | Ictalurus | punctatus | PREDICTED: ryanodine receptor 1-like isoform X6 [Ictalurus punctatus] |
| XP_017344268.1 | 5066 | LOC108276801 | eukaryota>metazoa>chordata>vertebrata>actinopterygii | Ictalurus | punctatus | PREDICTED: ryanodine receptor 1-like isoform X7 [Ictalurus punctatus] |
| XP_009303777.1 | 5063 | ryr1a | eukaryota>metazoa>chordata>vertebrata>actinopterygii | Danio | rerio | ryanodine receptor 1 isoform X2 [Danio rerio] |
| XP_011606962.1 | 5063 | LOC101073831 | eukaryota>metazoa>chordata>vertebrata>actinopterygii | Takifugu | rubripes | PREDICTED: ryanodine receptor 1-like [Takifugu rubripes] |
| XP_006539751.1 | 5061 | AI528790,Ryr,RYR-1,skrr | eukaryota>metazoa>chordata>vertebrata | Mus | musculus | PREDICTED: ryanodine receptor 1 isoform X1 [Mus musculus] |
| XP_017177540.1 | 5056 | AI528790,Ryr,RYR-1,skrr | eukaryota>metazoa>chordata>vertebrata | Mus | musculus | PREDICTED: ryanodine receptor 1 isoform X2 [Mus musculus] |
| XP_017177541.1 | 5055 | AI528790,Ryr,RYR-1,skrr | eukaryota>metazoa>chordata>vertebrata | Mus | musculus | PREDICTED: ryanodine receptor 1 isoform X3 [Mus musculus] |
| XP_017177542.1 | 5054 | AI528790,Ryr,RYR-1,skrr | eukaryota>metazoa>chordata>vertebrata | Mus | musculus | PREDICTED: ryanodine receptor 1 isoform X4 [Mus musculus] |
| XP_009059256.1 | 5045 | LOTGIDRAFT_123762 | eukaryota>metazoa>mollusca | Lottia | gigantea | hypothetical protein LOTGIDRAFT_123762 [Lottia gigantea] |
| XP_017177543.1 | 5042 | AI528790,Ryr,RYR-1,skrr | eukaryota>metazoa>chordata>vertebrata | Mus | musculus | PREDICTED: ryanodine receptor 1 isoform X5 [Mus musculus] |
| Caps1000000006 | 5039 | Caps1000000006 | eukaryota>metazoa>annelida | Capitella | spI | >Caps1000000006 |
| NP_000531.2 | 5038 | CCO,MHS,MHS1,PPP1R137,RYDR,RYR,RYR-1,SKRR | eukaryota>metazoa>chordata>vertebrata | Homo | sapiens | ryanodine receptor 1 isoform 1 [Homo sapiens] |
| XP_017177544.1 | 5036 | AI528790,Ryr,RYR-1,skrr | eukaryota>metazoa>chordata>vertebrata | Mus | musculus | PREDICTED: ryanodine receptor 1 isoform X6 [Mus musculus] |
| NP_033135.2 | 5035 | AI528790,Ryr,RYR-1,skrr | eukaryota>metazoa>chordata>vertebrata | Mus | musculus | ryanodine receptor 1 [Mus musculus] |
| NP_001036188.1 | 5033 | CCO,MHS,MHS1,PPP1R137,RYDR,RYR,RYR-1,SKRR | eukaryota>metazoa>chordata>vertebrata | Homo | sapiens | ryanodine receptor 1 isoform 2 [Homo sapiens] |
| XP_006723380.1 | 5032 | CCO,MHS,MHS1,PPP1R137,RYDR,RYR,RYR-1,SKRR | eukaryota>metazoa>chordata>vertebrata | Homo | sapiens | PREDICTED: ryanodine receptor 1 isoform X1 [Homo sapiens] |
| XP_006539755.1 | 5031 | AI528790,Ryr,RYR-1,skrr | eukaryota>metazoa>chordata>vertebrata | Mus | musculus | PREDICTED: ryanodine receptor 1 isoform X11 [Mus musculus] |
| XP_006539752.1 | 5030 | AI528790,Ryr,RYR-1,skrr | eukaryota>metazoa>chordata>vertebrata | Mus | musculus | PREDICTED: ryanodine receptor 1 isoform X7 [Mus musculus] |
| XP_018085767.1 | 5030 | LOC108698642 | eukaryota>metazoa>chordata>vertebrata | Xenopus | laevis | PREDICTED: ryanodine receptor 1-like isoform X1 [Xenopus laevis] |
| XP_015292237.1 | 5028 | RYR2 | eukaryota>metazoa>chordata>vertebrata | Macaca | fascicularis | PREDICTED: ryanodine receptor 2 isoform X1 [Macaca fascicularis] |
| XP_006723382.1 | 5027 | CCO,MHS,MHS1,PPP1R137,RYDR,RYR,RYR-1,SKRR | eukaryota>metazoa>chordata>vertebrata | Homo | sapiens | PREDICTED: ryanodine receptor 1 isoform X2 [Homo sapiens] |
| XP_018085768.1 | 5027 | LOC108698642 | eukaryota>metazoa>chordata>vertebrata | Xenopus | laevis | PREDICTED: ryanodine receptor 1-like isoform X2 [Xenopus laevis] |
| XP_018085769.1 | 5025 | LOC108698642 | eukaryota>metazoa>chordata>vertebrata | Xenopus | laevis | PREDICTED: ryanodine receptor 1-like isoform X3 [Xenopus laevis] |
| XP_018085770.1 | 5024 | LOC108698642 | eukaryota>metazoa>chordata>vertebrata | Xenopus | laevis | PREDICTED: ryanodine receptor 1-like isoform X4 [Xenopus laevis] |
| XP_018085771.1 | 5023 | LOC108698642 | eukaryota>metazoa>chordata>vertebrata | Xenopus | laevis | PREDICTED: ryanodine receptor 1-like isoform X5 [Xenopus laevis] |
| XP_018085772.1 | 5022 | LOC108698642 | eukaryota>metazoa>chordata>vertebrata | Xenopus | laevis | PREDICTED: ryanodine receptor 1-like isoform X6 [Xenopus laevis] |
| XP_018085773.1 | 5020 | LOC108698642 | eukaryota>metazoa>chordata>vertebrata | Xenopus | laevis | PREDICTED: ryanodine receptor 1-like isoform X7 [Xenopus laevis] |
| XP_015139906.1 | 5017 | RYR2 | eukaryota>metazoa>chordata>vertebrata | Gallus | gallus | PREDICTED: ryanodine receptor 2 isoform X1 [Gallus gallus] |
| XP_015292256.1 | 5017 | RYR2 | eukaryota>metazoa>chordata>vertebrata | Macaca | fascicularis | PREDICTED: ryanodine receptor 2 isoform X2 [Macaca fascicularis] |
| XP_015292289.1 | 5016 | RYR2 | eukaryota>metazoa>chordata>vertebrata | Macaca | fascicularis | PREDICTED: ryanodine receptor 2 isoform X3 [Macaca fascicularis] |
| XP_018085776.1 | 5016 | LOC108698642 | eukaryota>metazoa>chordata>vertebrata | Xenopus | laevis | PREDICTED: ryanodine receptor 1-like isoform X10 [Xenopus laevis] |
| XP_018085774.1 | 5015 | LOC108698642 | eukaryota>metazoa>chordata>vertebrata | Xenopus | laevis | PREDICTED: ryanodine receptor 1-like isoform X8 [Xenopus laevis] |
| XP_018085775.1 | 5014 | LOC108698642 | eukaryota>metazoa>chordata>vertebrata | Xenopus | laevis | PREDICTED: ryanodine receptor 1-like isoform X9 [Xenopus laevis] |
| XP_011525507.1 | 5009 | CCO,MHS,MHS1,PPP1R137,RYDR,RYR,RYR-1,SKRR | eukaryota>metazoa>chordata>vertebrata | Homo | sapiens | PREDICTED: ryanodine receptor 1 isoform X3 [Homo sapiens] |
| XP_015292339.1 | 5009 | RYR2 | eukaryota>metazoa>chordata>vertebrata | Macaca | fascicularis | PREDICTED: ryanodine receptor 2 isoform X4 [Macaca fascicularis] |
| XP_018669433.1 | 5002 | LOC100178148 | eukaryota>metazoa>chordata | Ciona | intestinalis | PREDICTED: ryanodine receptor 3 [Ciona intestinalis] |
| XP_018645937.1 | 4998 | Smp_163570 | eukaryota>metazoa | Schistosoma | mansoni | ryanodine receptor related [Schistosoma mansoni] |
| XP_015292368.1 | 4997 | RYR2 | eukaryota>metazoa>chordata>vertebrata | Macaca | fascicularis | PREDICTED: ryanodine receptor 2 isoform X5 [Macaca fascicularis] |
| XP_015292403.1 | 4996 | RYR2 | eukaryota>metazoa>chordata>vertebrata | Macaca | fascicularis | PREDICTED: ryanodine receptor 2 isoform X6 [Macaca fascicularis] |
| XP_017170934.1 | 4995 | 9330127I20Rik,RYR-2 | eukaryota>metazoa>chordata>vertebrata | Mus | musculus | PREDICTED: ryanodine receptor 2 isoform X1 [Mus musculus] |
| XP_017170935.1 | 4993 | 9330127I20Rik,RYR-2 | eukaryota>metazoa>chordata>vertebrata | Mus | musculus | PREDICTED: ryanodine receptor 2 isoform X2 [Mus musculus] |
| XP_015139907.1 | 4992 | RYR2 | eukaryota>metazoa>chordata>vertebrata | Gallus | gallus | PREDICTED: ryanodine receptor 2 isoform X2 [Gallus gallus] |
| XP_017170936.1 | 4989 | 9330127I20Rik,RYR-2 | eukaryota>metazoa>chordata>vertebrata | Mus | musculus | PREDICTED: ryanodine receptor 2 isoform X3 [Mus musculus] |
| XP_017170937.1 | 4988 | 9330127I20Rik,RYR-2 | eukaryota>metazoa>chordata>vertebrata | Mus | musculus | PREDICTED: ryanodine receptor 2 isoform X4 [Mus musculus] |
| XP_017170938.1 | 4987 | 9330127I20Rik,RYR-2 | eukaryota>metazoa>chordata>vertebrata | Mus | musculus | PREDICTED: ryanodine receptor 2 isoform X5 [Mus musculus] |
| XP_015292439.1 | 4986 | RYR2 | eukaryota>metazoa>chordata>vertebrata | Macaca | fascicularis | PREDICTED: ryanodine receptor 2 isoform X7 [Macaca fascicularis] |
| XP_006516673.1 | 4985 | 9330127I20Rik,RYR-2 | eukaryota>metazoa>chordata>vertebrata | Mus | musculus | PREDICTED: ryanodine receptor 2 isoform X6 [Mus musculus] |
| XP_006711865.1 | 4985 | ARVC2,ARVD2,RyR,RYR-2,VTSIP | eukaryota>metazoa>chordata>vertebrata | Homo | sapiens | PREDICTED: ryanodine receptor 2 isoform X1 [Homo sapiens] |
| XP_006711866.1 | 4984 | ARVC2,ARVD2,RyR,RYR-2,VTSIP | eukaryota>metazoa>chordata>vertebrata | Homo | sapiens | PREDICTED: ryanodine receptor 2 isoform X2 [Homo sapiens] |
| XP_015139908.1 | 4984 | RYR2 | eukaryota>metazoa>chordata>vertebrata | Gallus | gallus | PREDICTED: ryanodine receptor 2 isoform X3 [Gallus gallus] |
| XP_017170939.1 | 4984 | 9330127I20Rik,RYR-2 | eukaryota>metazoa>chordata>vertebrata | Mus | musculus | PREDICTED: ryanodine receptor 2 isoform X7 [Mus musculus] |
| XP_017170940.1 | 4983 | 9330127I20Rik,RYR-2 | eukaryota>metazoa>chordata>vertebrata | Mus | musculus | PREDICTED: ryanodine receptor 2 isoform X8 [Mus musculus] |
| XP_017170941.1 | 4983 | 9330127I20Rik,RYR-2 | eukaryota>metazoa>chordata>vertebrata | Mus | musculus | PREDICTED: ryanodine receptor 2 isoform X9 [Mus musculus] |
| XP_016857517.1 | 4978 | ARVC2,ARVD2,RyR,RYR-2,VTSIP | eukaryota>metazoa>chordata>vertebrata | Homo | sapiens | PREDICTED: ryanodine receptor 2 isoform X3 [Homo sapiens] |
| XP_006711867.1 | 4977 | ARVC2,ARVD2,RyR,RYR-2,VTSIP | eukaryota>metazoa>chordata>vertebrata | Homo | sapiens | PREDICTED: ryanodine receptor 2 isoform X4 [Homo sapiens] |
| XP_017170942.1 | 4976 | 9330127I20Rik,RYR-2 | eukaryota>metazoa>chordata>vertebrata | Mus | musculus | PREDICTED: ryanodine receptor 2 isoform X10 [Mus musculus] |
| XP_006711868.1 | 4975 | ARVC2,ARVD2,RyR,RYR-2,VTSIP | eukaryota>metazoa>chordata>vertebrata | Homo | sapiens | PREDICTED: ryanodine receptor 2 isoform X5 [Homo sapiens] |
| XP_015292504.1 | 4975 | RYR2 | eukaryota>metazoa>chordata>vertebrata | Macaca | fascicularis | PREDICTED: ryanodine receptor 2 isoform X9 [Macaca fascicularis] |
| XP_017170943.1 | 4975 | 9330127I20Rik,RYR-2 | eukaryota>metazoa>chordata>vertebrata | Mus | musculus | PREDICTED: ryanodine receptor 2 isoform X11 [Mus musculus] |
| XP_017170949.1 | 4974 | 9330127I20Rik,RYR-2 | eukaryota>metazoa>chordata>vertebrata | Mus | musculus | PREDICTED: ryanodine receptor 2 isoform X18 [Mus musculus] |
| XP_006711869.1 | 4973 | ARVC2,ARVD2,RyR,RYR-2,VTSIP | eukaryota>metazoa>chordata>vertebrata | Homo | sapiens | PREDICTED: ryanodine receptor 2 isoform X6 [Homo sapiens] |
| XP_017339435.1 | 4971 | ryr2 | eukaryota>metazoa>chordata>vertebrata>actinopterygii | Ictalurus | punctatus | PREDICTED: LOW QUALITY PROTEIN: ryanodine receptor 2 [Ictalurus punctatus] |
| CBY11776.1 | 4968 | GSOID_T00016960001 | eukaryota>metazoa>chordata | Oikopleura | dioica | unnamed protein product [Oikopleura dioica] |
| NP_001026.2 | 4967 | ARVC2,ARVD2,RyR,RYR-2,VTSIP | eukaryota>metazoa>chordata>vertebrata | Homo | sapiens | ryanodine receptor 2 [Homo sapiens] |
| XP_005539757.1 | 4967 | RYR2 | eukaryota>metazoa>chordata>vertebrata | Macaca | fascicularis | PREDICTED: ryanodine receptor 2 isoform X10 [Macaca fascicularis] |
| NP_076357.2 | 4966 | 9330127I20Rik,RYR-2 | eukaryota>metazoa>chordata>vertebrata | Mus | musculus | ryanodine receptor 2 [Mus musculus] |
| XP_006711870.1 | 4965 | ARVC2,ARVD2,RyR,RYR-2,VTSIP | eukaryota>metazoa>chordata>vertebrata | Homo | sapiens | PREDICTED: ryanodine receptor 2 isoform X7 [Homo sapiens] |
| XP_017170944.1 | 4955 | 9330127I20Rik,RYR-2 | eukaryota>metazoa>chordata>vertebrata | Mus | musculus | PREDICTED: ryanodine receptor 2 isoform X12 [Mus musculus] |
| XP_006711873.1 | 4954 | ARVC2,ARVD2,RyR,RYR-2,VTSIP | eukaryota>metazoa>chordata>vertebrata | Homo | sapiens | PREDICTED: ryanodine receptor 2 isoform X10 [Homo sapiens] |
| XP_017170950.1 | 4953 | 9330127I20Rik,RYR-2 | eukaryota>metazoa>chordata>vertebrata | Mus | musculus | PREDICTED: ryanodine receptor 2 isoform X19 [Mus musculus] |
| XP_002109077.1 | 4949 | TRIADDRAFT_52741 | eukaryota>metazoa>placozoa | Trichoplax | adhaerens | hypothetical protein TRIADDRAFT_52741 [Trichoplax adhaerens] |
| XP_018089264.1 | 4949 | LOC108700581 | eukaryota>metazoa>chordata>vertebrata | Xenopus | laevis | PREDICTED: ryanodine receptor 1-like [Xenopus laevis] |
| XP_011242598.1 | 4947 | 9330127I20Rik,RYR-2 | eukaryota>metazoa>chordata>vertebrata | Mus | musculus | PREDICTED: ryanodine receptor 2 isoform X13 [Mus musculus] |
| XP_011616342.1 | 4947 | LOC101062804 | eukaryota>metazoa>chordata>vertebrata>actinopterygii | Takifugu | rubripes | PREDICTED: LOW QUALITY PROTEIN: ryanodine receptor 2-like [Takifugu rubripes] |
| XP_012361884.1 | 4933 | RYR2 | eukaryota>metazoa>chordata>vertebrata | Nomascus | leucogenys | PREDICTED: LOW QUALITY PROTEIN: ryanodine receptor 2 [Nomascus leucogenys] |
| XP_018089157.1 | 4927 | LOC108700460 | eukaryota>metazoa>chordata>vertebrata | Xenopus | laevis | PREDICTED: ryanodine receptor 3-like [Xenopus laevis] |
| XP_017177545.1 | 4917 | AI528790,Ryr,RYR-1,skrr | eukaryota>metazoa>chordata>vertebrata | Mus | musculus | PREDICTED: ryanodine receptor 1 isoform X8 [Mus musculus] |
| XP_015292467.1 | 4917 | RYR2 | eukaryota>metazoa>chordata>vertebrata | Macaca | fascicularis | PREDICTED: ryanodine receptor 2 isoform X8 [Macaca fascicularis] |
| CAG00547.1 | 4910 | GSTEN:00019016:G:001 | eukaryota>metazoa>chordata>vertebrata>actinopterygii | Tetraodon | nigroviridis | unnamed protein product, partial [Tetraodon nigroviridis] |
| XP_006711871.1 | 4906 | ARVC2,ARVD2,RyR,RYR-2,VTSIP | eukaryota>metazoa>chordata>vertebrata | Homo | sapiens | PREDICTED: ryanodine receptor 2 isoform X8 [Homo sapiens] |
| XP_017172189.1 | 4895 | AI851294,C230090H21,RYR-3 | eukaryota>metazoa>chordata>vertebrata | Mus | musculus | PREDICTED: ryanodine receptor 3 isoform X1 [Mus musculus] |
| XP_017172191.1 | 4890 | AI851294,C230090H21,RYR-3 | eukaryota>metazoa>chordata>vertebrata | Mus | musculus | PREDICTED: ryanodine receptor 3 isoform X2 [Mus musculus] |
| NP_808320.2 | 4888 | AI851294,C230090H21,RYR-3 | eukaryota>metazoa>chordata>vertebrata | Mus | musculus | ryanodine receptor 3 isoform 1 [Mus musculus] |
| XP_012362990.1 | 4883 | RYR3 | eukaryota>metazoa>chordata>vertebrata | Nomascus | leucogenys | PREDICTED: LOW QUALITY PROTEIN: ryanodine receptor 3 [Nomascus leucogenys] |
| XP_017331022.1 | 4882 | LOC108269589 | eukaryota>metazoa>chordata>vertebrata>actinopterygii | Ictalurus | punctatus | PREDICTED: ryanodine receptor 2-like [Ictalurus punctatus] |
| XP_015142778.1 | 4882 | RYR3 | eukaryota>metazoa>chordata>vertebrata | Gallus | gallus | PREDICTED: ryanodine receptor 3 isoform X1 [Gallus gallus] |
| XP_015142779.1 | 4881 | RYR3 | eukaryota>metazoa>chordata>vertebrata | Gallus | gallus | PREDICTED: ryanodine receptor 3 isoform X2 [Gallus gallus] |
| XP_015142780.1 | 4881 | RYR3 | eukaryota>metazoa>chordata>vertebrata | Gallus | gallus | PREDICTED: ryanodine receptor 3 isoform X3 [Gallus gallus] |
| XP_015142781.1 | 4878 | RYR3 | eukaryota>metazoa>chordata>vertebrata | Gallus | gallus | PREDICTED: ryanodine receptor 3 isoform X4 [Gallus gallus] |
| XP_015142782.1 | 4877 | RYR3 | eukaryota>metazoa>chordata>vertebrata | Gallus | gallus | PREDICTED: ryanodine receptor 3 isoform X5 [Gallus gallus] |
| XP_015142783.1 | 4877 | RYR3 | eukaryota>metazoa>chordata>vertebrata | Gallus | gallus | PREDICTED: ryanodine receptor 3 isoform X6 [Gallus gallus] |
| XP_015142784.1 | 4876 | RYR3 | eukaryota>metazoa>chordata>vertebrata | Gallus | gallus | PREDICTED: ryanodine receptor 3 isoform X7 [Gallus gallus] |
| XP_015142785.1 | 4876 | RYR3 | eukaryota>metazoa>chordata>vertebrata | Gallus | gallus | PREDICTED: ryanodine receptor 3 isoform X8 [Gallus gallus] |
| XP_015142786.1 | 4875 | RYR3 | eukaryota>metazoa>chordata>vertebrata | Gallus | gallus | PREDICTED: ryanodine receptor 3 isoform X9 [Gallus gallus] |
| XP_017172201.1 | 4875 | AI851294,C230090H21,RYR-3 | eukaryota>metazoa>chordata>vertebrata | Mus | musculus | PREDICTED: ryanodine receptor 3 isoform X8 [Mus musculus] |
| XP_015308076.1 | 4874 | RYR3 | eukaryota>metazoa>chordata>vertebrata | Macaca | fascicularis | PREDICTED: ryanodine receptor 3 isoform X1 [Macaca fascicularis] |
| XP_017332742.1 | 4874 | LOC108270491 | eukaryota>metazoa>chordata>vertebrata>actinopterygii | Ictalurus | punctatus | PREDICTED: ryanodine receptor 3-like [Ictalurus punctatus] |
| XP_015308077.1 | 4873 | RYR3 | eukaryota>metazoa>chordata>vertebrata | Macaca | fascicularis | PREDICTED: ryanodine receptor 3 isoform X2 [Macaca fascicularis] |
| XP_011617561.1 | 4872 | LOC101061096 | eukaryota>metazoa>chordata>vertebrata>actinopterygii | Takifugu | rubripes | PREDICTED: ryanodine receptor 3-like [Takifugu rubripes] |
| NP_001027.3 | 4870 | RYR-3 | eukaryota>metazoa>chordata>vertebrata | Homo | sapiens | ryanodine receptor 3 isoform 1 [Homo sapiens] |
| NP_996757.2 | 4870 | RYR3 | eukaryota>metazoa>chordata>vertebrata | Gallus | gallus | ryanodine receptor 3 [Gallus gallus] |
| XP_005559133.1 | 4870 | RYR3 | eukaryota>metazoa>chordata>vertebrata | Macaca | fascicularis | PREDICTED: ryanodine receptor 3 isoform X3 [Macaca fascicularis] |
| XP_015142787.1 | 4870 | RYR3 | eukaryota>metazoa>chordata>vertebrata | Gallus | gallus | PREDICTED: ryanodine receptor 3 isoform X10 [Gallus gallus] |
| XP_018086652.1 | 4870 | LOC108699223 | eukaryota>metazoa>chordata>vertebrata | Xenopus | laevis | PREDICTED: ryanodine receptor 3 isoform X1 [Xenopus laevis] |
| XP_015308078.1 | 4869 | RYR3 | eukaryota>metazoa>chordata>vertebrata | Macaca | fascicularis | PREDICTED: ryanodine receptor 3 isoform X4 [Macaca fascicularis] |
| XP_015308079.1 | 4869 | RYR3 | eukaryota>metazoa>chordata>vertebrata | Macaca | fascicularis | PREDICTED: ryanodine receptor 3 isoform X5 [Macaca fascicularis] |
| XP_018086654.1 | 4869 | LOC108699223 | eukaryota>metazoa>chordata>vertebrata | Xenopus | laevis | PREDICTED: ryanodine receptor 3 isoform X2 [Xenopus laevis] |
| NP_001306085.1 | 4868 | AI851294,C230090H21,RYR-3 | eukaryota>metazoa>chordata>vertebrata | Mus | musculus | ryanodine receptor 3 [Mus musculus] |
| XP_011610427.1 | 4867 | LOC101061364 | eukaryota>metazoa>chordata>vertebrata>actinopterygii | Takifugu | rubripes | PREDICTED: LOW QUALITY PROTEIN: ryanodine receptor 3-like [Takifugu rubripes] |
| XP_015142789.1 | 4866 | RYR3 | eukaryota>metazoa>chordata>vertebrata | Gallus | gallus | PREDICTED: ryanodine receptor 3 isoform X12 [Gallus gallus] |
| XP_016877957.1 | 4866 | RYR-3 | eukaryota>metazoa>chordata>vertebrata | Homo | sapiens | PREDICTED: ryanodine receptor 3 isoform X1 [Homo sapiens] |
| NP_001230925.1 | 4865 | RYR-3 | eukaryota>metazoa>chordata>vertebrata | Homo | sapiens | ryanodine receptor 3 isoform 2 [Homo sapiens] |
| XP_005559134.1 | 4865 | RYR3 | eukaryota>metazoa>chordata>vertebrata | Macaca | fascicularis | PREDICTED: ryanodine receptor 3 isoform X6 [Macaca fascicularis] |
| XP_015142788.1 | 4865 | RYR3 | eukaryota>metazoa>chordata>vertebrata | Gallus | gallus | PREDICTED: ryanodine receptor 3 isoform X11 [Gallus gallus] |
| XP_016877958.1 | 4865 | RYR-3 | eukaryota>metazoa>chordata>vertebrata | Homo | sapiens | PREDICTED: ryanodine receptor 3 isoform X2 [Homo sapiens] |
| XP_016877959.1 | 4865 | RYR-3 | eukaryota>metazoa>chordata>vertebrata | Homo | sapiens | PREDICTED: ryanodine receptor 3 isoform X3 [Homo sapiens] |
| XP_016877960.1 | 4865 | RYR-3 | eukaryota>metazoa>chordata>vertebrata | Homo | sapiens | PREDICTED: ryanodine receptor 3 isoform X4 [Homo sapiens] |
| XP_015308080.1 | 4864 | RYR3 | eukaryota>metazoa>chordata>vertebrata | Macaca | fascicularis | PREDICTED: ryanodine receptor 3 isoform X7 [Macaca fascicularis] |
| XP_016877961.1 | 4864 | RYR-3 | eukaryota>metazoa>chordata>vertebrata | Homo | sapiens | PREDICTED: ryanodine receptor 3 isoform X5 [Homo sapiens] |
| XP_017172200.1 | 4864 | AI851294,C230090H21,RYR-3 | eukaryota>metazoa>chordata>vertebrata | Mus | musculus | PREDICTED: ryanodine receptor 3 isoform X7 [Mus musculus] |
| XP_018086655.1 | 4864 | LOC108699223 | eukaryota>metazoa>chordata>vertebrata | Xenopus | laevis | PREDICTED: ryanodine receptor 3 isoform X3 [Xenopus laevis] |
| XP_009293048.1 | 4863 | ryr3 | eukaryota>metazoa>chordata>vertebrata>actinopterygii | Danio | rerio | ryanodine receptor 3 isoform X1 [Danio rerio] |
| XP_017172194.1 | 4863 | AI851294,C230090H21,RYR-3 | eukaryota>metazoa>chordata>vertebrata | Mus | musculus | PREDICTED: ryanodine receptor 3 isoform X3 [Mus musculus] |
| XP_018086656.1 | 4863 | LOC108699223 | eukaryota>metazoa>chordata>vertebrata | Xenopus | laevis | PREDICTED: ryanodine receptor 3 isoform X4 [Xenopus laevis] |
| XP_016877962.1 | 4860 | RYR-3 | eukaryota>metazoa>chordata>vertebrata | Homo | sapiens | PREDICTED: ryanodine receptor 3 isoform X6 [Homo sapiens] |
| XP_011520182.1 | 4859 | RYR-3 | eukaryota>metazoa>chordata>vertebrata | Homo | sapiens | PREDICTED: ryanodine receptor 3 isoform X8 [Homo sapiens] |
| XP_016877963.1 | 4859 | RYR-3 | eukaryota>metazoa>chordata>vertebrata | Homo | sapiens | PREDICTED: ryanodine receptor 3 isoform X7 [Homo sapiens] |
| XP_017311641.1 | 4859 | ryr3 | eukaryota>metazoa>chordata>vertebrata>actinopterygii | Ictalurus | punctatus | PREDICTED: ryanodine receptor 3 isoform X1 [Ictalurus punctatus] |
| XP_009293049.1 | 4857 | ryr3 | eukaryota>metazoa>chordata>vertebrata>actinopterygii | Danio | rerio | ryanodine receptor 3 isoform X2 [Danio rerio] |
| XP_009293050.1 | 4856 | ryr3 | eukaryota>metazoa>chordata>vertebrata>actinopterygii | Danio | rerio | ryanodine receptor 3 isoform X3 [Danio rerio] |
| XP_017311642.1 | 4853 | ryr3 | eukaryota>metazoa>chordata>vertebrata>actinopterygii | Ictalurus | punctatus | PREDICTED: ryanodine receptor 3 isoform X2 [Ictalurus punctatus] |
| XP_017311643.1 | 4852 | ryr3 | eukaryota>metazoa>chordata>vertebrata>actinopterygii | Ictalurus | punctatus | PREDICTED: ryanodine receptor 3 isoform X3 [Ictalurus punctatus] |
| XP_017311645.1 | 4846 | ryr3 | eukaryota>metazoa>chordata>vertebrata>actinopterygii | Ictalurus | punctatus | PREDICTED: ryanodine receptor 3 isoform X5 [Ictalurus punctatus] |
| XP_015308081.1 | 4845 | RYR3 | eukaryota>metazoa>chordata>vertebrata | Macaca | fascicularis | PREDICTED: ryanodine receptor 3 isoform X8 [Macaca fascicularis] |
| XP_011601311.1 | 4841 | LOC101066256 | eukaryota>metazoa>chordata>vertebrata>actinopterygii | Takifugu | rubripes | PREDICTED: ryanodine receptor 2-like [Takifugu rubripes] |
| XP_016877964.1 | 4837 | RYR-3 | eukaryota>metazoa>chordata>vertebrata | Homo | sapiens | PREDICTED: ryanodine receptor 3 isoform X9 [Homo sapiens] |
| CAG10072.1 | 4833 | GSTEN:00031574:G:001 | eukaryota>metazoa>chordata>vertebrata>actinopterygii | Tetraodon | nigroviridis | unnamed protein product, partial [Tetraodon nigroviridis] |
| XP_016877965.1 | 4830 | RYR-3 | eukaryota>metazoa>chordata>vertebrata | Homo | sapiens | PREDICTED: ryanodine receptor 3 isoform X10 [Homo sapiens] |
| XP_017207197.1 | 4799 | ryr2b | eukaryota>metazoa>chordata>vertebrata>actinopterygii | Danio | rerio | PREDICTED: ryanodine receptor 2 [Danio rerio] |
| XP_017311644.1 | 4779 | ryr3 | eukaryota>metazoa>chordata>vertebrata>actinopterygii | Ictalurus | punctatus | PREDICTED: ryanodine receptor 3 isoform X4 [Ictalurus punctatus] |
| XP_015308082.1 | 4778 | RYR3 | eukaryota>metazoa>chordata>vertebrata | Macaca | fascicularis | PREDICTED: ryanodine receptor 3 isoform X9 [Macaca fascicularis] |
| XP_019855347.1 | 4753 | LOC100637092 | eukaryota>metazoa | Amphimedon | queenslandica | PREDICTED: ryanodine receptor 2 [Amphimedon queenslandica] |
| XP_013393030.1 | 4715 | LOC106160817 | eukaryota>metazoa | Lingula | anatina | PREDICTED: ryanodine receptor 44F-like [Lingula anatina] |
| XP_003378799.1 | 4543 | Tsp_02477 | eukaryota>metazoa>nematoda | Trichinella | spiralis | putative RIH domain protein [Trichinella spiralis] |
| XP_003199712.4 | 4534 | ryr2a | eukaryota>metazoa>chordata>vertebrata>actinopterygii | Danio | rerio | PREDICTED: ryanodine receptor 2 [Danio rerio] |
| XP_015296463.1 | 4457 | RYR1 | eukaryota>metazoa>chordata>vertebrata | Macaca | fascicularis | PREDICTED: ryanodine receptor 1, partial [Macaca fascicularis] |
| XP_012365616.1 | 4405 | RYR1 | eukaryota>metazoa>chordata>vertebrata | Nomascus | leucogenys | PREDICTED: LOW QUALITY PROTEIN: ryanodine receptor 1 [Nomascus leucogenys] |
| XP_014787750.1 | 4339 | LOC106881762 | eukaryota>metazoa>mollusca | Octopus | bimaculoides | PREDICTED: ryanodine receptor-like isoform X13 [Octopus bimaculoides] |
| CAG12758.1 | 4251 | GSTEN:00035194:G:001 | eukaryota>metazoa>chordata>vertebrata>actinopterygii | Tetraodon | nigroviridis | unnamed protein product [Tetraodon nigroviridis] |
| XP_015308083.1 | 3863 | RYR3 | eukaryota>metazoa>chordata>vertebrata | Macaca | fascicularis | PREDICTED: ryanodine receptor 3 isoform X10 [Macaca fascicularis] |
| XP_016877966.1 | 3855 | RYR-3 | eukaryota>metazoa>chordata>vertebrata | Homo | sapiens | PREDICTED: ryanodine receptor 3 isoform X11 [Homo sapiens] |
| XP_017172197.1 | 3575 | AI851294,C230090H21,RYR-3 | eukaryota>metazoa>chordata>vertebrata | Mus | musculus | PREDICTED: ryanodine receptor 3 isoform X4 [Mus musculus] |
| XP_017208123.1 | 3508 | ryr3 | eukaryota>metazoa>chordata>vertebrata>actinopterygii | Danio | rerio | PREDICTED: ryanodine receptor 3 isoform X4 [Danio rerio] |
| XP_017172198.1 | 3367 | AI851294,C230090H21,RYR-3 | eukaryota>metazoa>chordata>vertebrata | Mus | musculus | PREDICTED: ryanodine receptor 3 isoform X5 [Mus musculus] |
| XP_014664675.1 | 3274 | LOC106806993 | eukaryota>metazoa | Priapulus | caudatus | PREDICTED: ryanodine receptor-like [Priapulus caudatus] |
| XP_017172199.1 | 3083 | AI851294,C230090H21,RYR-3 | eukaryota>metazoa>chordata>vertebrata | Mus | musculus | PREDICTED: ryanodine receptor 3 isoform X6 [Mus musculus] |
| XP_017177547.1 | 3076 | AI528790,Ryr,RYR-1,skrr | eukaryota>metazoa>chordata>vertebrata | Mus | musculus | PREDICTED: ryanodine receptor 1 isoform X10 [Mus musculus] |
| OQV16826.1 | 3045 | BV898_08999 | eukaryota>metazoa | Hypsibius | dujardini | Ryanodine receptor [Hypsibius dujardini] |
| OQV16827.1 | 3033 | BV898_08999 | eukaryota>metazoa | Hypsibius | dujardini | Ryanodine receptor [Hypsibius dujardini] |
| XP_017170946.1 | 3033 | 9330127I20Rik,RYR-2 | eukaryota>metazoa>chordata>vertebrata | Mus | musculus | PREDICTED: ryanodine receptor 2 isoform X15 [Mus musculus] |
| XP_017170947.1 | 3016 | 9330127I20Rik,RYR-2 | eukaryota>metazoa>chordata>vertebrata | Mus | musculus | PREDICTED: ryanodine receptor 2 isoform X16 [Mus musculus] |
| XP_006711872.1 | 2735 | ARVC2,ARVD2,RyR,RYR-2,VTSIP | eukaryota>metazoa>chordata>vertebrata | Homo | sapiens | PREDICTED: ryanodine receptor 2 isoform X9 [Homo sapiens] |
| XP_002597150.1 | 2638 | BRAFLDRAFT_215612 | eukaryota>metazoa>chordata | Branchiostoma | floridae | hypothetical protein BRAFLDRAFT_215612, partial [Branchiostoma floridae] |
| OQV16829.1 | 2143 | BV898_09001 | eukaryota>metazoa | Hypsibius | dujardini | Ryanodine receptor [Hypsibius dujardini] |

W. Sorcin

| | | | | | | |
|---|---|---|---|---|---|---|
| XP_006712461.1 | 259 | GCL | eukaryota>metazoa>chordata>vertebrata | Homo | sapiens | PREDICTED: grancalcin isoform X1 [Homo sapiens] |
| XP_013404159.1 | 258 | LOC106169292 | eukaryota>metazoa | Lingula | anatina | PREDICTED: sorcin-like [Lingula anatina] |
| XP_003727947.1 | 253 | LOC582258 | eukaryota>metazoa>echinodermata | Strongylocentrotus | purpuratus | PREDICTED: sorcin isoform X2 [Strongylocentrotus purpuratus] |
| XP_003966584.1 | 252 | LOC101070744 | eukaryota>metazoa>chordata>vertebrata>actinopterygii | Takifugu | rubripes | PREDICTED: grancalcin-like [Takifugu rubripes] |
| XP_006712463.1 | 248 | GCL | eukaryota>metazoa>chordata>vertebrata | Homo | sapiens | PREDICTED: grancalcin isoform X2 [Homo sapiens] |
| XP_011435979.1 | 246 | LOC105334287 | eukaryota>metazoa>mollusca | Crassostrea | gigas | PREDICTED: sorcin isoform X1 [Crassostrea gigas] |
| XP_011435980.1 | 246 | LOC105334287 | eukaryota>metazoa>mollusca | Crassostrea | gigas | PREDICTED: sorcin isoform X2 [Crassostrea gigas] |
| XP_006712464.1 | 243 | GCL | eukaryota>metazoa>chordata>vertebrata | Homo | sapiens | PREDICTED: grancalcin isoform X3 [Homo sapiens] |
| CAG04972.1 | 237 | GSTEN:00024843:G:001 | eukaryota>metazoa>chordata>vertebrata>actinopterygii | Tetraodon | nigroviridis | unnamed protein product [Tetraodon nigroviridis] |
| XP_005246503.1 | 233 | GCL | eukaryota>metazoa>chordata>vertebrata | Homo | sapiens | PREDICTED: grancalcin isoform X4 [Homo sapiens] |
| XP_016859253.1 | 232 | GCL | eukaryota>metazoa>chordata>vertebrata | Homo | sapiens | PREDICTED: grancalcin isoform X5 [Homo sapiens] |
| XP_016859254.1 | 232 | GCL | eukaryota>metazoa>chordata>vertebrata | Homo | sapiens | PREDICTED: grancalcin isoform X5 [Homo sapiens] |
| XP_016859255.1 | 232 | GCL | eukaryota>metazoa>chordata>vertebrata | Homo | sapiens | PREDICTED: grancalcin isoform X5 [Homo sapiens] |
| XP_002605900.1 | 230 | BRAFLDRAFT_124895 | eukaryota>metazoa>chordata | Branchiostoma | floridae | hypothetical protein BRAFLDRAFT_124895 [Branchiostoma floridae] |
| XP_017336976.1 | 228 | LOC108272774 | eukaryota>metazoa>chordata>vertebrata>actinopterygii | Ictalurus | punctatus | PREDICTED: grancalcin-like isoform X1 [Ictalurus punctatus] |
| NP_663498.1 | 220 | 5133401E04Rik,AI573844 | eukaryota>metazoa>chordata>vertebrata | Mus | musculus | grancalcin [Mus musculus] |
| XP_001748649.1 | 220 | MONBRDRAFT_35370 | eukaryota>choanoflagellida | Monosiga | brevicollis MX1 | hypothetical protein [Monosiga brevicollis MX1] |
| XP_017350789.1 | 220 | LOC108280385 | eukaryota>metazoa>chordata>vertebrata>actinopterygii | Ictalurus | punctatus | PREDICTED: sorcin-like isoform X1 [Ictalurus punctatus] |
| XP_005573380.1 | 218 | GCA | eukaryota>metazoa>chordata>vertebrata | Macaca | fascicularis | PREDICTED: grancalcin isoform X1 [Macaca fascicularis] |
| XP_012351979.1 | 218 | GCA | eukaryota>metazoa>chordata>vertebrata | Nomascus | leucogenys | PREDICTED: grancalcin isoform X1 [Nomascus leucogenys] |
| NP_036330.1 | 217 | GCL | eukaryota>metazoa>chordata>vertebrata | Homo | sapiens | grancalcin isoform b [Homo sapiens] |
| XP_003383149.1 | 217 | LOC100633073 | eukaryota>metazoa | Amphimedon | queenslandica | PREDICTED: sorcin-like [Amphimedon queenslandica] |
| XP_012351980.1 | 215 | GCA | eukaryota>metazoa>chordata>vertebrata | Nomascus | leucogenys | PREDICTED: grancalcin isoform X2 [Nomascus leucogenys] |
| XP_011509228.1 | 214 | GCL | eukaryota>metazoa>chordata>vertebrata | Homo | sapiens | PREDICTED: grancalcin isoform X6 [Homo sapiens] |
| XP_016859256.1 | 214 | GCL | eukaryota>metazoa>chordata>vertebrata | Homo | sapiens | PREDICTED: grancalcin isoform X6 [Homo sapiens] |
| XP_004998129.1 | 210 | PTSG_00660 | eukaryota>choanoflagellida | Salpingoeca | rosetta | grancalcin [Salpingoeca rosetta] |
| XP_017336977.1 | 208 | LOC108272774 | eukaryota>metazoa>chordata>vertebrata>actinopterygii | Ictalurus | punctatus | PREDICTED: grancalcin-like isoform X2 [Ictalurus punctatus] |
| NP_001083493.1 | 207 | gca.S | eukaryota>metazoa>chordata>vertebrata | Xenopus | laevis | uncharacterized protein LOC398957 [Xenopus laevis] |
| NP_001085654.1 | 207 | gca | eukaryota>metazoa>chordata>vertebrata | Xenopus | laevis | grancalcin L homeolog [Xenopus laevis] |
| NP_001005585.1 | 205 | zgc:92027 | eukaryota>metazoa>chordata>vertebrata>actinopterygii | Danio | rerio | grancalcin [Danio rerio] |
| XP_004343120.1 | 205 | CAOG_07261 | eukaryota>ichthyosporea | Capsaspora | owczarzaki ATCC 30864 | sorcin [Capsaspora owczarzaki ATCC 30864] |
| XP_012556640.1 | 201 | LOC105844501 | eukaryota>metazoa>cnidaria | Hydra | vulgaris | PREDICTED: sorcin-like [Hydra vulgaris] |
| XP_003266225.1 | 199 | GCA | eukaryota>metazoa>chordata>vertebrata | Nomascus | leucogenys | PREDICTED: grancalcin isoform X3 [Nomascus leucogenys] |
| NP_001074334.1 | 198 | SRI | eukaryota>metazoa>chordata>vertebrata | Gallus | gallus | sorcin [Gallus gallus] |
| NP_001074443.1 | 198 | 2210417O06Rik,2900070H08Rik,Sor | eukaryota>metazoa>chordata>vertebrata | Mus | musculus | sorcin isoform 1 [Mus musculus] |
| NP_001274256.1 | 198 | SRI | eukaryota>metazoa>chordata>vertebrata | Macaca | fascicularis | sorcin [Macaca fascicularis] |
| NP_003121.1 | 198 | CP-22,CP22,SCN,V19 | eukaryota>metazoa>chordata>vertebrata | Homo | sapiens | sorcin isoform A [Homo sapiens] |
| XP_003252404.1 | 198 | SRI | eukaryota>metazoa>chordata>vertebrata | Nomascus | leucogenys | PREDICTED: sorcin isoform X1 [Nomascus leucogenys] |
| XP_011509229.1 | 198 | GCL | eukaryota>metazoa>chordata>vertebrata | Homo | sapiens | PREDICTED: grancalcin isoform X7 [Homo sapiens] |
| XP_016859257.1 | 198 | GCL | eukaryota>metazoa>chordata>vertebrata | Homo | sapiens | PREDICTED: grancalcin isoform X7 [Homo sapiens] |
| XP_015287784.1 | 197 | GCA | eukaryota>metazoa>chordata>vertebrata | Macaca | fascicularis | PREDICTED: grancalcin isoform X2 [Macaca fascicularis] |
| XP_015287785.1 | 197 | GCA | eukaryota>metazoa>chordata>vertebrata | Macaca | fascicularis | PREDICTED: grancalcin isoform X2 [Macaca fascicularis] |
| XP_017336978.1 | 197 | LOC108272774 | eukaryota>metazoa>chordata>vertebrata>actinopterygii | Ictalurus | punctatus | PREDICTED: grancalcin-like isoform X3 [Ictalurus punctatus] |
| NP_001089357.1 | 196 | scn,sri | eukaryota>metazoa>chordata>vertebrata | Xenopus | laevis | sorcin S homeolog [Xenopus laevis] |
| NP_001091359.1 | 196 | scn,sri | eukaryota>metazoa>chordata>vertebrata | Xenopus | laevis | sorcin L homeolog [Xenopus laevis] |
| XP_018092870.1 | 196 | MGC68559 | eukaryota>metazoa>chordata>vertebrata | Xenopus | laevis | PREDICTED: uncharacterized protein LOC398957 isoform X1 [Xenopus laevis] |
| NP_001243820.1 | 195 | CP-22,CP22,SCN,V19 | eukaryota>metazoa>chordata>vertebrata | Homo | sapiens | sorcin isoform C [Homo sapiens] |
| XP_002599500.1 | 192 | RAB9A.1 | eukaryota>metazoa>chordata | Branchiostoma | floridae | hypothetical protein BRAFLDRAFT_58969 [Branchiostoma floridae] |
| XP_011669403.1 | 192 | LOC578745 | eukaryota>metazoa>echinodermata | Strongylocentrotus | purpuratus | PREDICTED: sorcin [Strongylocentrotus purpuratus] |
| XP_005157918.1 | 190 | zgc:64070 | eukaryota>metazoa>chordata>vertebrata>actinopterygii | Danio | rerio | sorcin isoform X1 [Danio rerio] |
| XP_017350790.1 | 188 | LOC108280385 | eukaryota>metazoa>chordata>vertebrata>actinopterygii | Ictalurus | punctatus | PREDICTED: sorcin-like isoform X2 [Ictalurus punctatus] |
| XP_002741663.1 | 187 | LOC100368657 | eukaryota>metazoa>hemichordata | Saccoglossus | kowalevskii | PREDICTED: sorcin-like [Saccoglossus kowalevskii] |
| XP_014160674.1 | 187 | SARC_01106 | eukaryota>ichthyosporea | Sphaeroforma | arctica JP610 | hypothetical protein SARC_01106 [Sphaeroforma arctica JP610] |
| XP_014160675.1 | 187 | SARC_01106 | eukaryota>ichthyosporea | Sphaeroforma | arctica JP610 | hypothetical protein, variant [Sphaeroforma arctica JP610] |
| NP_079894.2 | 183 | 2210417O06Rik,2900070H08Rik,Sor | eukaryota>metazoa>chordata>vertebrata | Mus | musculus | sorcin isoform 2 [Mus musculus] |
| NP_944490.1 | 183 | CP-22,CP22,SCN,V19 | eukaryota>metazoa>chordata>vertebrata | Homo | sapiens | sorcin isoform B [Homo sapiens] |
| XP_003252405.1 | 183 | SRI | eukaryota>metazoa>chordata>vertebrata | Nomascus | leucogenys | PREDICTED: sorcin isoform X2 [Nomascus leucogenys] |
| XP_005550355.1 | 183 | SRI | eukaryota>metazoa>chordata>vertebrata | Macaca | fascicularis | PREDICTED: sorcin isoform X1 [Macaca fascicularis] |
| NP_001243821.1 | 180 | CP-22,CP22,SCN,V19 | eukaryota>metazoa>chordata>vertebrata | Homo | sapiens | sorcin isoform D [Homo sapiens] |
| XP_011270782.1 | 180 | CAOG_07261 | eukaryota>ichthyosporea | Capsaspora | owczarzaki ATCC 30864 | sorcin, variant [Capsaspora owczarzaki ATCC 30864] |
| XP_002585553.1 | 179 | BRAFLDRAFT_290057 | eukaryota>metazoa>chordata | Branchiostoma | floridae | hypothetical protein BRAFLDRAFT_290057 [Branchiostoma floridae] |
| XP_003965936.2 | 173 | LOC101066788 | eukaryota>metazoa>chordata>vertebrata>actinopterygii | Takifugu | rubripes | PREDICTED: sorcin-like [Takifugu rubripes] |
| NP_956667.1 | 169 | zgc:64070 | eukaryota>metazoa>chordata>vertebrata>actinopterygii | Danio | rerio | sorcin [Danio rerio] |
| XP_002110749.1 | 167 | TRIADDRAFT_22435 | eukaryota>metazoa>placozoa | Trichoplax | adhaerens | hypothetical protein TRIADDRAFT_22435, partial [Trichoplax adhaerens] |
| CAF95586.1 | 165 | GSTEN:00012303:G:001 | eukaryota>metazoa>chordata>vertebrata>actinopterygii | Tetraodon | nigroviridis | unnamed protein product, partial [Tetraodon nigroviridis] |
| XP_001641156.1 | 161 | NEMVEDRAFT_v1g81972 | eukaryota>metazoa>cnidaria | Nematostella | vectensis | predicted protein, partial [Nematostella vectensis] |
| XP_011514830.1 | 155 | CP-22,CP22,SCN,V19 | eukaryota>metazoa>chordata>vertebrata | Homo | sapiens | PREDICTED: sorcin isoform X1 [Homo sapiens] |
| XP_011509230.1 | 137 | GCL | eukaryota>metazoa>chordata>vertebrata | Homo | sapiens | PREDICTED: grancalcin isoform X8 [Homo sapiens] |
| KXJ21168.1 | 135 | SRI | eukaryota>metazoa>cnidaria | Exaiptasia | pallida | Sorcin [Exaiptasia pallida] |
| XP_015750990.1 | 121 | LOC107330972 | eukaryota>metazoa>cnidaria | Acropora | digitifera | PREDICTED: sorcin-like [Acropora digitifera] |
| XP_015750991.1 | 121 | LOC107330973 | eukaryota>metazoa>cnidaria | Acropora | digitifera | PREDICTED: sorcin-like [Acropora digitifera] |
| XP_006503590.1 | 113 | 2210417O06Rik,2900070H08Rik,Sor | eukaryota>metazoa>chordata>vertebrata | Mus | musculus | PREDICTED: sorcin isoform X1 [Mus musculus] |
| XP_017176089.1 | 113 | 2210417O06Rik,2900070H08Rik,Sor | eukaryota>metazoa>chordata>vertebrata | Mus | musculus | PREDICTED: sorcin isoform X1 [Mus musculus] |

X. L-type VGCC (pore)

| | | | | | | |
|---|---|---|---|---|---|---|
| NP_723952.1 | 2552 | alpha1,alpha1D,BG:DS02795.1,Ca-a1D,Ca-alpha1,ca-alpha1d,ca-alpha1D,Ca-alpha[[1]]D,Ca-_1D,Caalpha1D,cad,caID,Ca_alpha1D,CG4894,DmCa1alpha1,Dmca1D,DmCa1D,Dmel\CG4894,DroCa1,l(2)35Fa,l35Fa | eukaryota>metazoa>hexapoda | Drosophila | melanogaster | Ca[2+]-channel protein alpha[[1]] subunit D, isoform C [Drosophila melanogaster] |
| NP_001260480.1 | 2546 | alpha1,alpha1D,BG:DS02795.1,Ca-a1D,Ca-alpha1,ca-alpha1d,ca-alpha1D,Ca-alpha[[1]]D,Ca-_1D,Caalpha1D,cad,caID,Ca_alpha1D,CG4894,DmCa1alpha1,Dmca1D,DmCa1D,Dmel\CG4894,DroCa1,l(2)35Fa,l35Fa | eukaryota>metazoa>hexapoda | Drosophila | melanogaster | Ca[2+]-channel protein alpha[[1]] subunit D, isoform E [Drosophila melanogaster] |
| XP_013414688.1 | 2539 | LOC106176729 | eukaryota>metazoa | Lingula | anatina | PREDICTED: muscle calcium channel subunit alpha-1-like [Lingula anatina] |
| NP_525104.2 | 2519 | alpha1,alpha1D,BG:DS02795.1,Ca-a1D,Ca-alpha1,ca-alpha1d,ca-alpha1D,Ca-alpha[[1]]D,Ca-_1D,Caalpha1D,cad,caID,Ca_alpha1D,CG4894,DmCa1alpha1,Dmca1D,DmCa1D,Dmel\CG4894,DroCa1,l(2)35Fa,l35Fa | eukaryota>metazoa>hexapoda | Drosophila | melanogaster | Ca[2+]-channel protein alpha[[1]] subunit D, isoform B [Drosophila melanogaster] |
| XP_011419158.1 | 2519 | LOC105322251 | eukaryota>metazoa>mollusca | Crassostrea | gigas | PREDICTED: voltage-dependent calcium channel type A subunit alpha-1 isoform X1 [Crassostrea gigas] |
| NP_602305.1 | 2516 | alpha1,alpha1D,BG:DS02795.1,Ca-a1D,Ca-alpha1,ca-alpha1d,ca-alpha1D,Ca-alpha[[1]]D,Ca-_1D,Caalpha1D,cad,caID,Ca_alpha1D,CG4894,DmCa1alpha1,Dmca1D,DmCa1D,Dmel\CG4894,DroCa1,l(2)35Fa,l35Fa | eukaryota>metazoa>hexapoda | Drosophila | melanogaster | Ca[2+]-channel protein alpha[[1]] subunit D, isoform A [Drosophila melanogaster] |
| XP_011419159.1 | 2515 | LOC105322251 | eukaryota>metazoa>mollusca | Crassostrea | gigas | PREDICTED: voltage-dependent calcium channel type A subunit alpha-1 isoform X2 [Crassostrea gigas] |
| XP_011419160.1 | 2514 | LOC105322251 | eukaryota>metazoa>mollusca | Crassostrea | gigas | PREDICTED: voltage-dependent calcium channel type A subunit alpha-1 isoform X3 [Crassostrea gigas] |
| XP_011419161.1 | 2513 | LOC105322251 | eukaryota>metazoa>mollusca | Crassostrea | gigas | PREDICTED: voltage-dependent calcium channel type A subunit alpha-1 isoform X4 [Crassostrea gigas] |
| XP_011419163.1 | 2513 | LOC105322251 | eukaryota>metazoa>mollusca | Crassostrea | gigas | PREDICTED: voltage-dependent calcium channel type A subunit alpha-1 isoform X5 [Crassostrea gigas] |
| XP_011419164.1 | 2509 | LOC105322251 | eukaryota>metazoa>mollusca | Crassostrea | gigas | PREDICTED: voltage-dependent calcium channel type A subunit alpha-1 isoform X6 [Crassostrea gigas] |
| XP_011419165.1 | 2507 | LOC105322251 | eukaryota>metazoa>mollusca | Crassostrea | gigas | PREDICTED: voltage-dependent calcium channel type A subunit alpha-1 isoform X7 [Crassostrea gigas] |
| XP_019920406.1 | 2503 | LOC105322251 | eukaryota>metazoa>mollusca | Crassostrea | gigas | PREDICTED: voltage-dependent calcium channel type A subunit alpha-1 isoform X19 [Crassostrea gigas] |
| XP_011419166.1 | 2502 | LOC105322251 | eukaryota>metazoa>mollusca | Crassostrea | gigas | PREDICTED: voltage-dependent calcium channel type A subunit alpha-1 isoform X8 [Crassostrea gigas] |
| XP_011419167.1 | 2501 | LOC105322251 | eukaryota>metazoa>mollusca | Crassostrea | gigas | PREDICTED: voltage-dependent calcium channel type A subunit alpha-1 isoform X9 [Crassostrea gigas] |
| XP_011419168.1 | 2492 | LOC105322251 | eukaryota>metazoa>mollusca | Crassostrea | gigas | PREDICTED: voltage-dependent calcium channel type A subunit alpha-1 isoform X10 [Crassostrea gigas] |
| XP_011419169.1 | 2486 | LOC105322251 | eukaryota>metazoa>mollusca | Crassostrea | gigas | PREDICTED: voltage-dependent calcium channel type A subunit alpha-1 isoform X11 [Crassostrea gigas] |
| XP_011419170.1 | 2486 | LOC105322251 | eukaryota>metazoa>mollusca | Crassostrea | gigas | PREDICTED: voltage-dependent calcium channel type A subunit alpha-1 isoform X12 [Crassostrea gigas] |
| XP_011419180.1 | 2481 | LOC105322251 | eukaryota>metazoa>mollusca | Crassostrea | gigas | PREDICTED: voltage-dependent calcium channel type A subunit alpha-1 isoform X21 [Crassostrea gigas] |
| XP_019920407.1 | 2481 | LOC105322251 | eukaryota>metazoa>mollusca | Crassostrea | gigas | PREDICTED: voltage-dependent calcium channel type A subunit alpha-1 isoform X20 [Crassostrea gigas] |
| XP_011419171.1 | 2470 | LOC105322251 | eukaryota>metazoa>mollusca | Crassostrea | gigas | PREDICTED: voltage-dependent calcium channel type A subunit alpha-1 isoform X13 [Crassostrea gigas] |

| | | | | | | |
|---|---|---|---|---|---|---|
| EUB63035.1 | 2470 | EGR 02129 | eukaryota>metazoa | Echinococcus | granulosus | Voltage-dependent calcium channel type D subunit alpha-1 [Echinococcus granulosus] |
| XP_016875415.1 | 2412 | CACH2,CACN2,CACNL1A1,CaV1.2,CCHL1A1,LQT8,TS | eukaryota>metazoa>chordata>vertebrata | Homo | sapiens | PREDICTED: voltage-dependent L-type calcium channel subunit alpha-1C isoform X1 [Homo sapiens] |
| XP_016875416.1 | 2377 | CACH2,CACN2,CACNL1A1,CaV1.2,CCHL1A1,LQT8,TS | eukaryota>metazoa>chordata>vertebrata | Homo | sapiens | PREDICTED: voltage-dependent L-type calcium channel subunit alpha-1C isoform X2 [Homo sapiens] |
| CAG03249.1 | 2340 | GSTEN:00022583:G:001 | eukaryota>metazoa>chordata>vertebrata>actinopterygii | Tetraodon | nigroviridis | unnamed protein product, partial [Tetraodon nigroviridis] |
| CDS41920.2 | 2337 | EmuJ 000961000 | eukaryota>metazoa | Echinococcus | multilocularis | voltage dependent calcium channel subunit [Echinococcus multilocularis] |
| XP_011419172.1 | 2314 | LOC105322251 | eukaryota>metazoa>mollusca | Crassostrea | gigas | PREDICTED: voltage-dependent calcium channel type A subunit alpha-1 isoform X14 [Crassostrea gigas] |
| XP_016875417.1 | 2306 | CACH2,CACN2,CACNL1A1,CaV1.2,CCHL1A1,LQT8,TS | eukaryota>metazoa>chordata>vertebrata | Homo | sapiens | PREDICTED: voltage-dependent L-type calcium channel subunit alpha-1C isoform X3 [Homo sapiens] |
| XP_011670127.1 | 2304 | LOC578128 | eukaryota>metazoa>echinodermata | Strongylocentrotus | purpuratus | PREDICTED: voltage dependent calcium channel L-type isoform X1 [Strongylocentrotus purpuratus] |
| XP_011670128.1 | 2303 | LOC578128 | eukaryota>metazoa>echinodermata | Strongylocentrotus | purpuratus | PREDICTED: voltage dependent calcium channel L-type isoform X2 [Strongylocentrotus purpuratus] |
| XP_011662956.1 | 2301 | LOC574743 | eukaryota>metazoa>echinodermata | Strongylocentrotus | purpuratus | PREDICTED: voltage-dependent calcium channel type A subunit alpha-1 [Strongylocentrotus purpuratus] |
| NP_723953.1 | 2300 | alpha1,alpha1D,BG:DS02795.1,Ca-a1D,Ca-alpha1,ca-alpha1d,ca-alpha1D,Ca-alpha[[1]]D,Ca-_1D,Caalpha1D,cad,caID,Ca_alpha1D,CG4894,DmCa1alpha1,Dmca1D,DmCa1D,Dmel\CG4894,DroCa1,l(2)35Fa,l35Fa | eukaryota>metazoa>hexapoda | Drosophila | melanogaster | Ca[2+]-channel protein alpha[[1]] subunit D, isoform D [Drosophila melanogaster] |
| XP_011670130.1 | 2300 | LOC578128 | eukaryota>metazoa>echinodermata | Strongylocentrotus | purpuratus | PREDICTED: voltage dependent calcium channel L-type isoform X3 [Strongylocentrotus purpuratus] |
| XP_011670131.1 | 2299 | LOC578128 | eukaryota>metazoa>echinodermata | Strongylocentrotus | purpuratus | PREDICTED: voltage dependent calcium channel L-type isoform X4 [Strongylocentrotus purpuratus] |
| XP_011670132.1 | 2298 | LOC578128 | eukaryota>metazoa>echinodermata | Strongylocentrotus | purpuratus | PREDICTED: voltage dependent calcium channel L-type isoform X5 [Strongylocentrotus purpuratus] |
| XP_011670133.1 | 2293 | LOC578128 | eukaryota>metazoa>echinodermata | Strongylocentrotus | purpuratus | PREDICTED: voltage dependent calcium channel L-type isoform X6 [Strongylocentrotus purpuratus] |
| XP_011670134.1 | 2289 | LOC578128 | eukaryota>metazoa>echinodermata | Strongylocentrotus | purpuratus | PREDICTED: voltage dependent calcium channel L-type isoform X7 [Strongylocentrotus purpuratus] |
| XP_016875418.1 | 2287 | CACH2,CACN2,CACNL1A1,CaV1.2,CCHL1A1,LQT8,TS | eukaryota>metazoa>chordata>vertebrata | Homo | sapiens | PREDICTED: voltage-dependent L-type calcium channel subunit alpha-1C isoform X4 [Homo sapiens] |
| XP_016875419.1 | 2287 | CACH2,CACN2,CACNL1A1,CaV1.2,CCHL1A1,LQT8,TS | eukaryota>metazoa>chordata>vertebrata | Homo | sapiens | PREDICTED: voltage-dependent L-type calcium channel subunit alpha-1C isoform X5 [Homo sapiens] |
| XP_011419175.1 | 2282 | LOC105322251 | eukaryota>metazoa>mollusca | Crassostrea | gigas | PREDICTED: voltage-dependent calcium channel type A subunit alpha-1 isoform X15 [Crassostrea gigas] |
| XP_012365582.1 | 2264 | CACNA1D | eukaryota>metazoa>chordata>vertebrata | Nomascus | leucogenys | PREDICTED: LOW QUALITY PROTEIN: voltage-dependent L-type calcium channel subunit alpha-1D [Nomascus leucogenys] |
| XP_016875420.1 | 2259 | CACH2,CACN2,CACNL1A1,CaV1.2,CCHL1A1,LQT8,TS | eukaryota>metazoa>chordata>vertebrata | Homo | sapiens | PREDICTED: voltage-dependent L-type calcium channel subunit alpha-1C isoform X6 [Homo sapiens] |
| XP_016875421.1 | 2254 | CACH2,CACN2,CACNL1A1,CaV1.2,CCHL1A1,LQT8,TS | eukaryota>metazoa>chordata>vertebrata | Homo | sapiens | PREDICTED: voltage-dependent L-type calcium channel subunit alpha-1C isoform X7 [Homo sapiens] |
| XP_011532396.1 | 2246 | CACH3,CACN4,CACNL1A2,Cav1.3,CCHL1A2,PASNA,SANDD | eukaryota>metazoa>chordata>vertebrata | Homo | sapiens | PREDICTED: voltage-dependent L-type calcium channel subunit alpha-1D isoform X2 [Homo sapiens] |
| XP_016862626.1 | 2246 | CACH3,CACN4,CACNL1A2,Cav1.3,CCHL1A2,PASNA,SANDD | eukaryota>metazoa>chordata>vertebrata | Homo | sapiens | PREDICTED: voltage-dependent L-type calcium channel subunit alpha-1D isoform X1 [Homo sapiens] |
| XP_016862627.1 | 2245 | CACH3,CACN4,CACNL1A2,Cav1.3,CCHL1A2,PASNA,SANDD | eukaryota>metazoa>chordata>vertebrata | Homo | sapiens | PREDICTED: voltage-dependent L-type calcium channel subunit alpha-1D isoform X3 [Homo sapiens] |
| XP_015148461.1 | 2238 | CACNA1D | eukaryota>metazoa>chordata>vertebrata | Gallus | gallus | PREDICTED: voltage-dependent L-type calcium channel subunit alpha-1D isoform X1 [Gallus gallus] |
| XP_015148462.1 | 2238 | CACNA1D | eukaryota>metazoa>chordata>vertebrata | Gallus | gallus | PREDICTED: voltage-dependent L-type calcium channel subunit alpha-1D isoform X2 [Gallus gallus] |
| XP_016862628.1 | 2237 | CACH3,CACN4,CACNL1A2,Cav1.3,CCHL1A2,PASNA,SANDD | eukaryota>metazoa>chordata>vertebrata | Homo | sapiens | PREDICTED: voltage-dependent L-type calcium channel subunit alpha-1D isoform X4 [Homo sapiens] |
| XP_011419176.1 | 2226 | LOC105322251 | eukaryota>metazoa>mollusca | Crassostrea | gigas | PREDICTED: voltage-dependent calcium channel type A subunit alpha-1 isoform X16 [Crassostrea gigas] |
| XP_016862629.1 | 2226 | CACH3,CACN4,CACNL1A2,Cav1.3,CCHL1A2,PASNA,SANDD | eukaryota>metazoa>chordata>vertebrata | Homo | sapiens | PREDICTED: voltage-dependent L-type calcium channel subunit alpha-1D isoform X5 [Homo sapiens] |
| XP_016862630.1 | 2226 | CACH3,CACN4,CACNL1A2,Cav1.3,CCHL1A2,PASNA,SANDD | eukaryota>metazoa>chordata>vertebrata | Homo | sapiens | PREDICTED: voltage-dependent L-type calcium channel subunit alpha-1D isoform X6 [Homo sapiens] |
| XP_011239477.1 | 2225 | Cav1.2,Cchl1a1,D930026N18Rik,MBC,MELC-CC | eukaryota>metazoa>chordata>vertebrata | Mus | musculus | PREDICTED: voltage-dependent L-type calcium channel subunit alpha-1C isoform X1 [Mus musculus] |
| XP_011239478.1 | 2225 | Cav1.2,Cchl1a1,D930026N18Rik,MBC,MELC-CC | eukaryota>metazoa>chordata>vertebrata | Mus | musculus | PREDICTED: voltage-dependent L-type calcium channel subunit alpha-1C isoform X3 [Mus musculus] |
| XP_017176845.1 | 2225 | Cav1.2,Cchl1a1,D930026N18Rik,MBC,MELC-CC | eukaryota>metazoa>chordata>vertebrata | Mus | musculus | PREDICTED: voltage-dependent L-type calcium channel subunit alpha-1C isoform X2 [Mus musculus] |
| XP_017176846.1 | 2223 | Cav1.2,Cchl1a1,D930026N18Rik,MBC,MELC-CC | eukaryota>metazoa>chordata>vertebrata | Mus | musculus | PREDICTED: voltage-dependent L-type calcium channel subunit alpha-1C isoform X4 [Mus musculus] |
| NP_001242931.1 | 2222 | Cav1.2,Cchl1a1,D930026N18Rik,MBC,MELC-CC | eukaryota>metazoa>chordata>vertebrata | Mus | musculus | voltage-dependent L-type calcium channel subunit alpha-1C isoform 10 [Mus musculus] |
| XP_006505478.1 | 2222 | Cav1.2,Cchl1a1,D930026N18Rik,MBC,MELC-CC | eukaryota>metazoa>chordata>vertebrata | Mus | musculus | PREDICTED: voltage-dependent L-type calcium channel subunit alpha-1C isoform X6 [Mus musculus] |
| XP_016875422.1 | 2222 | CACH2,CACN2,CACNL1A1,CaV1.2,CCHL1A1,LQT8,TS | eukaryota>metazoa>chordata>vertebrata | Homo | sapiens | PREDICTED: voltage-dependent L-type calcium channel subunit alpha-1C isoform X8 [Homo sapiens] |
| XP_017176847.1 | 2222 | Cav1.2,Cchl1a1,D930026N18Rik,MBC,MELC-CC | eukaryota>metazoa>chordata>vertebrata | Mus | musculus | PREDICTED: voltage-dependent L-type calcium channel subunit alpha-1C isoform X5 [Mus musculus] |
| NP_955630.3 | 2221 | CACH2,CACN2,CACNL1A1,CaV1.2,CCHL1A1,LQT8,TS | eukaryota>metazoa>chordata>vertebrata | Homo | sapiens | voltage-dependent L-type calcium channel subunit alpha-1C isoform 1 [Homo sapiens] |
| XP_019697108.1 | 2221 | LOC105183370 | eukaryota>metazoa>hexapoda | Harpegnathos | saltator | PREDICTED: muscle calcium channel subunit alpha-1 isoform X6 [Harpegnathos saltator] |
| XP_015142138.1 | 2220 | CACNA1C | eukaryota>metazoa>chordata>vertebrata | Gallus | gallus | PREDICTED: voltage-dependent L-type calcium channel subunit alpha-1C isoform X31 [Gallus gallus] |
| XP_016875423.1 | 2219 | CACH2,CACN2,CACNL1A1,CaV1.2,CCHL1A1,LQT8,TS | eukaryota>metazoa>chordata>vertebrata | Homo | sapiens | PREDICTED: voltage-dependent L-type calcium channel subunit alpha-1C isoform X9 [Homo sapiens] |
| XP_016875424.1 | 2219 | CACH2,CACN2,CACNL1A1,CaV1.2,CCHL1A1,LQT8,TS | eukaryota>metazoa>chordata>vertebrata | Homo | sapiens | PREDICTED: voltage-dependent L-type calcium channel subunit alpha-1C isoform X10 [Homo sapiens] |
| XP_016875444.1 | 2219 | CACH2,CACN2,CACNL1A1,CaV1.2,CCHL1A1,LQT8,TS | eukaryota>metazoa>chordata>vertebrata | Homo | sapiens | PREDICTED: voltage-dependent L-type calcium channel subunit alpha-1C isoform X32 [Homo sapiens] |
| XP_015148463.1 | 2218 | CACNA1D | eukaryota>metazoa>chordata>vertebrata | Gallus | gallus | PREDICTED: voltage-dependent L-type calcium channel subunit alpha-1D isoform X3 [Gallus gallus] |
| XP_015148464.1 | 2218 | CACNA1D | eukaryota>metazoa>chordata>vertebrata | Gallus | gallus | PREDICTED: voltage-dependent L-type calcium channel subunit alpha-1D isoform X4 [Gallus gallus] |
| XP_016862631.1 | 2218 | CACH3,CACN4,CACNL1A2,Cav1.3,CCHL1A2,PASNA,SANDD | eukaryota>metazoa>chordata>vertebrata | Homo | sapiens | PREDICTED: voltage-dependent L-type calcium channel subunit alpha-1D isoform X7 [Homo sapiens] |
| XP_016862632.1 | 2218 | CACH3,CACN4,CACNL1A2,Cav1.3,CCHL1A2,PASNA,SANDD | eukaryota>metazoa>chordata>vertebrata | Homo | sapiens | PREDICTED: voltage-dependent L-type calcium channel subunit alpha-1D isoform X8 [Homo sapiens] |
| XP_016862633.1 | 2218 | CACH3,CACN4,CACNL1A2,Cav1.3,CCHL1A2,PASNA,SANDD | eukaryota>metazoa>chordata>vertebrata | Homo | sapiens | PREDICTED: voltage-dependent L-type calcium channel subunit alpha-1D isoform X9 [Homo sapiens] |
| XP_006505481.1 | 2217 | Cav1.2,Cchl1a1,D930026N18Rik,MBC,MELC-CC | eukaryota>metazoa>chordata>vertebrata | Mus | musculus | PREDICTED: voltage-dependent L-type calcium channel subunit alpha-1C isoform X35 [Mus musculus] |
| XP_017176848.1 | 2214 | Cav1.2,Cchl1a1,D930026N18Rik,MBC,MELC-CC | eukaryota>metazoa>chordata>vertebrata | Mus | musculus | PREDICTED: voltage-dependent L-type calcium channel subunit alpha-1C isoform X7 [Mus musculus] |
| XP_016875425.1 | 2213 | CACH2,CACN2,CACNL1A1,CaV1.2,CCHL1A1,LQT8,TS | eukaryota>metazoa>chordata>vertebrata | Homo | sapiens | PREDICTED: voltage-dependent L-type calcium channel subunit alpha-1C isoform X11 [Homo sapiens] |
| XP_016875426.1 | 2213 | CACH2,CACN2,CACNL1A1,CaV1.2,CCHL1A1,LQT8,TS | eukaryota>metazoa>chordata>vertebrata | Homo | sapiens | PREDICTED: voltage-dependent L-type calcium channel subunit alpha-1C isoform X12 [Homo sapiens] |
| XP_016875427.1 | 2213 | CACH2,CACN2,CACNL1A1,CaV1.2,CCHL1A1,LQT8,TS | eukaryota>metazoa>chordata>vertebrata | Homo | sapiens | PREDICTED: voltage-dependent L-type calcium channel subunit alpha-1C isoform X13 [Homo sapiens] |
| NP_001118235.1 | 2212 | LOC578128 | eukaryota>metazoa>echinodermata | Strongylocentrotus | purpuratus | voltage dependent calcium channel L-type [Strongylocentrotus purpuratus] |
| XP_015148465.1 | 2212 | CACNA1D | eukaryota>metazoa>chordata>vertebrata | Gallus | gallus | PREDICTED: voltage-dependent L-type calcium channel subunit alpha-1D isoform X5 [Gallus gallus] |
| XP_015148466.1 | 2212 | CACNA1D | eukaryota>metazoa>chordata>vertebrata | Gallus | gallus | PREDICTED: voltage-dependent L-type calcium channel subunit alpha-1D isoform X6 [Gallus gallus] |
| XP_011532402.1 | 2211 | CACH3,CACN4,CACNL1A2,Cav1.3,CCHL1A2,PASNA,SANDD | eukaryota>metazoa>chordata>vertebrata | Homo | sapiens | PREDICTED: voltage-dependent L-type calcium channel subunit alpha-1D isoform X15 [Homo sapiens] |
| XP_016875428.1 | 2211 | CACH2,CACN2,CACNL1A1,CaV1.2,CCHL1A1,LQT8,TS | eukaryota>metazoa>chordata>vertebrata | Homo | sapiens | PREDICTED: voltage-dependent L-type calcium channel subunit alpha-1C isoform X14 [Homo sapiens] |
| XP_015148467.1 | 2210 | CACNA1D | eukaryota>metazoa>chordata>vertebrata | Gallus | gallus | PREDICTED: voltage-dependent L-type calcium channel subunit alpha-1D isoform X7 [Gallus gallus] |
| XP_011419177.1 | 2208 | LOC105322251 | eukaryota>metazoa>mollusca | Crassostrea | gigas | PREDICTED: voltage-dependent calcium channel type A subunit alpha-1 isoform X17 [Crassostrea gigas] |
| XP_015148468.1 | 2208 | CACNA1D | eukaryota>metazoa>chordata>vertebrata | Gallus | gallus | PREDICTED: voltage-dependent L-type calcium channel subunit alpha-1D isoform X8 [Gallus gallus] |
| XP_015148469.1 | 2208 | CACNA1D | eukaryota>metazoa>chordata>vertebrata | Gallus | gallus | PREDICTED: voltage-dependent L-type calcium channel subunit alpha-1D isoform X9 [Gallus gallus] |
| [illegible] | [illegible] | [illegible] | [illegible] | [illegible] | [illegible] | [illegible] |

| | | | | | | |
|---|---|---|---|---|---|---|
| XP_003971275.1 | 1016 | LOC101070073 | eukaryota>metazoa>chordata>vertebrata>actinopterygii | Takifugu | rubripes | PREDICTED: short transient receptor potential channel 2-like [Takifugu rubripes] |
| XP_013384065.1 | 1014 | LOC106154271 | eukaryota>metazoa | Lingula | anatina | PREDICTED: short transient receptor potential channel 7-like [Lingula anatina] |
| XP_018669082.1 | 1013 | LOC100184754 | eukaryota>metazoa>chordata | Ciona | intestinalis | PREDICTED: short transient receptor potential channel 4-like isoform X1 [Ciona intestinalis] |
| XP_018669083.1 | 1010 | LOC100184754 | eukaryota>metazoa>chordata | Ciona | intestinalis | PREDICTED: short transient receptor potential channel 4-like isoform X2 [Ciona intestinalis] |
| ERG81205.1 | 1007 | ASU_10569 | eukaryota>metazoa>nematoda | Ascaris | suum | transient receptor potential-gamma protein [Ascaris suum] |
| CDS41436.1 | 989 | EmuJ_000908800 | eukaryota>metazoa | Echinococcus | multilocularis | transient receptor potential gamma protein [Echinococcus multilocularis] |
| XP_420310.2 | 989 | TRPC5 | eukaryota>metazoa>chordata>vertebrata | Gallus | gallus | PREDICTED: short transient receptor potential channel 5 [Gallus gallus] |
| NP_003297.1 | 982 | HTRP-4,HTRP4,TRP4 | eukaryota>metazoa>chordata>vertebrata | Homo | sapiens | short transient receptor potential channel 4 isoform epsilon [Homo sapiens] |
| XP_012357404.1 | 982 | TRPC4 | eukaryota>metazoa>chordata>vertebrata | Nomascus | leucogenys | PREDICTED: short transient receptor potential channel 4 isoform X1 [Nomascus leucogenys] |
| XP_018085351.1 | 982 | LOC108698404 | eukaryota>metazoa>chordata>vertebrata | Xenopus | laevis | PREDICTED: short transient receptor potential channel 5-like [Xenopus laevis] |
| XP_018088468.1 | 982 | LOC108700107 | eukaryota>metazoa>chordata>vertebrata | Xenopus | laevis | PREDICTED: short transient receptor potential channel 5-like [Xenopus laevis] |
| EUB64483.1 | 980 | EGR_00433 | eukaryota>metazoa | Echinococcus | granulosus | Transient receptor potential-gamma protein [Echinococcus granulosus] |
| NP_057263.1 | 977 | HTRP-4,HTRP4,TRP4 | eukaryota>metazoa>chordata>vertebrata | Homo | sapiens | short transient receptor potential channel 4 isoform alpha [Homo sapiens] |
| XP_003270343.1 | 977 | TRPC4 | eukaryota>metazoa>chordata>vertebrata | Nomascus | leucogenys | PREDICTED: short transient receptor potential channel 4 isoform X2 [Nomascus leucogenys] |
| XP_005585748.1 | 977 | TRPC4 | eukaryota>metazoa>chordata>vertebrata | Macaca | fascicularis | PREDICTED: short transient receptor potential channel 4 [Macaca fascicularis] |
| NP_033454.1 | 975 | CCE2,TRP-5,TRP5,Trrp5 | eukaryota>metazoa>chordata>vertebrata | Mus | musculus | short transient receptor potential channel 5 [Mus musculus] |
| NP_058680.1 | 974 | CCE1,STRPC4,Trp4,Trrp4 | eukaryota>metazoa>chordata>vertebrata | Mus | musculus | short transient receptor potential channel 4 isoform 1 [Mus musculus] |
| XP_003262260.1 | 974 | TRPC5 | eukaryota>metazoa>chordata>vertebrata | Nomascus | leucogenys | PREDICTED: short transient receptor potential channel 5 [Nomascus leucogenys] |
| XP_005594455.1 | 974 | TRPC5 | eukaryota>metazoa>chordata>vertebrata | Macaca | fascicularis | PREDICTED: short transient receptor potential channel 5 [Macaca fascicularis] |
| XP_006501359.1 | 974 | CCE1,STRPC4,Trp4,Trrp4 | eukaryota>metazoa>chordata>vertebrata | Mus | musculus | PREDICTED: short transient receptor potential channel 4 isoform X1 [Mus musculus] |
| XP_417089.2 | 974 | TRPC4 | eukaryota>metazoa>chordata>vertebrata | Gallus | gallus | PREDICTED: short transient receptor potential channel 4 [Gallus gallus] |
| NP_036603.1 | 973 | PPP1R159,TRP5 | eukaryota>metazoa>chordata>vertebrata | Homo | sapiens | short transient receptor potential channel 5 [Homo sapiens] |
| XP_016885263.1 | 973 | PPP1R159,TRP5 | eukaryota>metazoa>chordata>vertebrata | Homo | sapiens | PREDICTED: short transient receptor potential channel 5 isoform X1 [Homo sapiens] |
| XP_011666677.1 | 970 | LOC586844 | eukaryota>metazoa>echinodermata | Strongylocentrotus | purpuratus | PREDICTED: short transient receptor potential channel 3 isoform X1 [Strongylocentrotus purpuratus] |
| XP_011666683.1 | 970 | LOC586844 | eukaryota>metazoa>echinodermata | Strongylocentrotus | purpuratus | PREDICTED: short transient receptor potential channel 3 isoform X1 [Strongylocentrotus purpuratus] |
| XP_009009132.1 | 967 | HELRODRAFT_63393 | eukaryota>metazoa>annelida | Helobdella | robusta | hypothetical protein HELRODRAFT_63393 [Helobdella robusta] |
| XP_018103124.1 | 966 | LOC108708683 | eukaryota>metazoa>chordata>vertebrata | Xenopus | laevis | PREDICTED: short transient receptor potential channel 4-like [Xenopus laevis] |
| XP_003376781.1 | 957 | Tsp_00950 | eukaryota>metazoa>nematoda | Trichinella | spiralis | transient receptor potential-gamma protein [Trichinella spiralis] |
| XP_791700.3 | 956 | LOC586844 | eukaryota>metazoa>echinodermata | Strongylocentrotus | purpuratus | PREDICTED: short transient receptor potential channel 3 isoform X2 [Strongylocentrotus purpuratus] |
| XP_013386796.1 | 954 | LOC106156216 | eukaryota>metazoa | Lingula | anatina | PREDICTED: transient-receptor-potential-like protein [Lingula anatina] |
| XP_002733765.1 | 953 | LOC100378311 | eukaryota>metazoa>hemichordata | Saccoglossus | kowalevskii | PREDICTED: transient-receptor-potential-like protein-like [Saccoglossus kowalevskii] |
| XP_004938897.2 | 952 | TRPC6 | eukaryota>metazoa>chordata>vertebrata | Gallus | gallus | PREDICTED: short transient receptor potential channel 6 isoform X1 [Gallus gallus] |
| XP_006509912.1 | 949 | AV025995,LLHWJM002,LLHWJM003,LLHWJM004,mtrp6,TRP-6,Trrp6 | eukaryota>metazoa>chordata>vertebrata | Mus | musculus | PREDICTED: short transient receptor potential channel 6 isoform X3 [Mus musculus] |
| XP_013398015.1 | 942 | LOC106164593 | eukaryota>metazoa | Lingula | anatina | PREDICTED: short transient receptor potential channel 3-like [Lingula anatina] |
| OAF71995.1 | 941 | A3Q56_00219 | eukaryota>metazoa | Intoshia | linei | Transient receptor protein 2 [Intoshia linei] |
| XP_017343908.1 | 941 | trpc4 | eukaryota>metazoa>chordata>vertebrata>actinopterygii | Ictalurus | punctatus | PREDICTED: short transient receptor potential channel 4 [Ictalurus punctatus] |
| XP_018124202.1 | 941 | LOC108719617 | eukaryota>metazoa>chordata>vertebrata | Xenopus | laevis | PREDICTED: short transient receptor potential channel 3-like [Xenopus laevis] |
| XP_011530519.1 | 937 | SCA41,TRP3 | eukaryota>metazoa>chordata>vertebrata | Homo | sapiens | PREDICTED: short transient receptor potential channel 3 isoform X1 [Homo sapiens] |
| XP_015306276.1 | 937 | TRPC3 | eukaryota>metazoa>chordata>vertebrata | Macaca | fascicularis | PREDICTED: short transient receptor potential channel 3 isoform X1 [Macaca fascicularis] |
| XP_011667283.1 | 936 | LOC580692 | eukaryota>metazoa>echinodermata | Strongylocentrotus | purpuratus | PREDICTED: short transient receptor potential channel 3 [Strongylocentrotus purpuratus] |
| XP_016864067.1 | 936 | SCA41,TRP3 | eukaryota>metazoa>chordata>vertebrata | Homo | sapiens | PREDICTED: short transient receptor potential channel 3 isoform X2 [Homo sapiens] |
| XP_785826.2 | 936 | LOC580692 | eukaryota>metazoa>echinodermata | Strongylocentrotus | purpuratus | PREDICTED: short transient receptor potential channel 3 [Strongylocentrotus purpuratus] |
| NP_001276811.1 | 934 | si:dkey-109i21.3,trpc4b | eukaryota>metazoa>chordata>vertebrata>actinopterygii | Danio | rerio | short transient receptor potential channel 4 [Danio rerio] |
| XP_683649.2 | 934 | si:dkey-109i21.3,trpc4b | eukaryota>metazoa>chordata>vertebrata>actinopterygii | Danio | rerio | short transient receptor potential channel 4 isoform X1 [Danio rerio] |
| NP_001244208.1 | 933 | TRPC3 | eukaryota>metazoa>chordata>vertebrata | Gallus | gallus | short transient receptor potential channel 3 [Gallus gallus] |
| NP_004612.2 | 931 | FSGS2,TRP6 | eukaryota>metazoa>chordata>vertebrata | Homo | sapiens | short transient receptor potential channel 6 [Homo sapiens] |
| XP_005579492.1 | 931 | TRPC6 | eukaryota>metazoa>chordata>vertebrata | Macaca | fascicularis | PREDICTED: short transient receptor potential channel 6 [Macaca fascicularis] |
| NP_038866.2 | 930 | AV025995,LLHWJM002,LLHWJM003,LLHWJM004,mtrp6,TRP-6,Trrp6 | eukaryota>metazoa>chordata>vertebrata | Mus | musculus | short transient receptor potential channel 6 isoform 1 [Mus musculus] |
| XP_006509913.1 | 930 | AV025995,LLHWJM002,LLHWJM003,LLHWJM004,mtrp6,TRP-6,Trrp6 | eukaryota>metazoa>chordata>vertebrata | Mus | musculus | PREDICTED: short transient receptor potential channel 6 isoform X1 [Mus musculus] |
| XP_003253082.1 | 928 | TRPC6 | eukaryota>metazoa>chordata>vertebrata | Nomascus | leucogenys | PREDICTED: LOW QUALITY PROTEIN: short transient receptor potential channel 6 [Nomascus leucogenys] |
| XP_011455889.1 | 926 | LOC105348246 | eukaryota>metazoa>mollusca | Crassostrea | gigas | PREDICTED: short transient receptor potential channel 3 [Crassostrea gigas] |
| XP_011420089.1 | 924 | LOC105322886 | eukaryota>metazoa>mollusca | Crassostrea | gigas | PREDICTED: short transient receptor potential channel 3 [Crassostrea gigas] |
| XP_011428523.1 | 924 | LOC105329068 | eukaryota>metazoa>mollusca | Crassostrea | gigas | PREDICTED: short transient receptor potential channel 3 isoform X1 [Crassostrea gigas] |
| XP_002607434.1 | 923 | BRAFLDRAFT_69859 | eukaryota>metazoa>chordata | Branchiostoma | floridae | hypothetical protein BRAFLDRAFT_69859 [Branchiostoma floridae] |
| XP_011428524.1 | 923 | LOC105329068 | eukaryota>metazoa>mollusca | Crassostrea | gigas | PREDICTED: short transient receptor potential channel 3 isoform X2 [Crassostrea gigas] |
| XP_013780206.1 | 923 | LOC106464597 | eukaryota>metazoa | Limulus | polyphemus | PREDICTED: transient-receptor-potential-like protein isoform X1 [Limulus polyphemus] |
| XP_013780207.1 | 923 | LOC106464597 | eukaryota>metazoa | Limulus | polyphemus | PREDICTED: transient-receptor-potential-like protein isoform X2 [Limulus polyphemus] |
| NP_001124170.1 | 921 | SCA41,TRP3 | eukaryota>metazoa>chordata>vertebrata | Homo | sapiens | short transient receptor potential channel 3 isoform a [Homo sapiens] |
| XP_003271391.1 | 921 | TRPC3 | eukaryota>metazoa>chordata>vertebrata | Nomascus | leucogenys | PREDICTED: short transient receptor potential channel 3 isoform X1 [Nomascus leucogenys] |
| XP_005555901.1 | 921 | TRPC3 | eukaryota>metazoa>chordata>vertebrata | Macaca | fascicularis | PREDICTED: short transient receptor potential channel 3 isoform X2 [Macaca fascicularis] |
| XP_011530520.1 | 920 | SCA41,TRP3 | eukaryota>metazoa>chordata>vertebrata | Homo | sapiens | PREDICTED: short transient receptor potential channel 3 isoform X3 [Homo sapiens] |
| XP_003971385.1 | 918 | trpc4 | eukaryota>metazoa>chordata>vertebrata>actinopterygii | Takifugu | rubripes | PREDICTED: short transient receptor potential channel 4 [Takifugu rubripes] |
| XP_013775532.1 | 917 | LOC106460388 | eukaryota>metazoa | Limulus | polyphemus | PREDICTED: transient-receptor-potential-like protein [Limulus polyphemus] |
| XP_003968682.1 | 916 | LOC101062185 | eukaryota>metazoa>chordata>vertebrata>actinopterygii | Takifugu | rubripes | PREDICTED: short transient receptor potential channel 4-like [Takifugu rubripes] |
| [illegible] | | | | | | |

| | | | | | | |
|---|---|---|---|---|---|---|
| EUB65028.1 | 313 | EGR 00297 | eukaryota>metazoa | Echinococcus | granulosus | Calumenin-B [Echinococcus granulosus] |
| NP_001085102.1 | 313 | MGC78878 | eukaryota>metazoa>chordata>vertebrata | Xenopus | laevis | uncharacterized protein LOC432173 precursor [Xenopus laevis] |
| NP 001087007.1 | 313 | rcn2,rcn2.L | eukaryota>metazoa>chordata>vertebrata | Xenopus | laevis | reticulocalbin 2 S homeolog precursor [Xenopus laevis] |
| NP 001187699.1 | 313 | calu | eukaryota>metazoa>chordata>vertebrata>actinopterygii | Ictalurus | punctatus | reticulocalbin-3 precursor [Ictalurus punctatus] |
| XP_009043912.1 | 313 | LOTGIDRAFT_103075 | eukaryota>metazoa>mollusca | Lottia | gigantea | hypothetical protein LOTGIDRAFT_103075 [Lottia gigantea] |
| XP 017338425.1 | 313 | rcn3 | eukaryota>metazoa>chordata>vertebrata>actinopterygii | Ictalurus | punctatus | PREDICTED: reticulocalbin-3 isoform X1 [Ictalurus punctatus] |
| ERG83583.1 | 312 | ASU 06763 | eukaryota>metazoa>nematoda | Ascaris | suum | calumenin-a [Ascaris suum] |
| XP_002129310.1 | 312 | LOC100182984 | eukaryota>metazoa>chordata | Ciona | intestinalis | PREDICTED: reticulocalbin-2-like [Ciona intestinalis] |
| XP 002167284.1 | 312 | LOC100206632 | eukaryota>metazoa>cnidaria | Hydra | vulgaris | PREDICTED: calumenin-B-like [Hydra vulgaris] |
| OQV20397.1 | 312 | BV898 05683 | eukaryota>metazoa | Hypsibius | dujardini | putative Calumenin [Hypsibius dujardini] |
| XP_003378780.1 | 312 | Tsp_12034 | eukaryota>metazoa>nematoda | Trichinella | spiralis | putative calumenin [Trichinella spiralis] |
| XP 002123414.1 | 311 | LOC100184283 | eukaryota>metazoa>chordata | Ciona | intestinalis | PREDICTED: calumenin-B-like [Ciona intestinalis] |
| XP 011605652.1 | 310 | rcn2 | eukaryota>metazoa>chordata>vertebrata>actinopterygii | Takifugu | rubripes | PREDICTED: reticulocalbin-2 isoform X2 [Takifugu rubripes] |
| CBY10162.1 | 309 | GSOID_T00010993001 | eukaryota>metazoa>chordata | Oikopleura | dioica | unnamed protein product [Oikopleura dioica] |
| XP 008184569.1 | 309 | LOC100159618 | eukaryota>metazoa>hexapoda | Acyrthosiphon | pisum | PREDICTED: calumenin-A [Acyrthosiphon pisum] |
| XP 009031860.1 | 309 | HELRODRAFT_186226 | eukaryota>metazoa>annelida | Helobdella | robusta | hypothetical protein HELRODRAFT 186226 [Helobdella robusta] |
| KXJ19679.1 | 308 | calua | eukaryota>metazoa>cnidaria | Exaiptasia | pallida | Calumenin-A [Exaiptasia pallida] |
| KXJ19691.1 | 308 | calub | eukaryota>metazoa>cnidaria | Exaiptasia | pallida | Calumenin-B [Exaiptasia pallida] |
| XP 001627907.1 | 308 | NEMVEDRAFT v1g190767 | eukaryota>metazoa>cnidaria | Nematostella | vectensis | predicted protein [Nematostella vectensis] |
| Smar1000001370 | 308 | Smar1000001370 | eukaryota>metazoa>arthropoda>myriapoda | Strigamia | maritima | >Smar1000001370 |
| NP 001027627.1 | 307 | calumenin | eukaryota>metazoa>chordata | Ciona | intestinalis | calumenin homologue precursor [Ciona intestinalis] |
| NP 001280137.1 | 307 | RCN2 | eukaryota>metazoa>chordata>vertebrata | Gallus | gallus | reticulocalbin-2 isoform 1 precursor [Gallus gallus] |
| XP_002122894.2 | 307 | LOC100176707 | eukaryota>metazoa>chordata | Ciona | intestinalis | PREDICTED: reticulocalbin-2-like [Ciona intestinalis] |
| XP_009029673.1 | 307 | HELRODRAFT 89801 | eukaryota>metazoa>annelida | Helobdella | robusta | hypothetical protein HELRODRAFT_89801 [Helobdella robusta] |
| CAF94652.1 | 306 | GSTEN:00010846:G:001 | eukaryota>metazoa>chordata>vertebrata>actinopterygii | Tetraodon | nigroviridis | unnamed protein product [Tetraodon nigroviridis] |
| XP 002590942.1 | 304 | BRAFLDRAFT_101089 | eukaryota>metazoa>chordata | Branchiostoma | floridae | hypothetical protein BRAFLDRAFT 101089 [Branchiostoma floridae] |
| NP 001280159.1 | 303 | RCN2 | eukaryota>metazoa>chordata>vertebrata | Gallus | gallus | reticulocalbin-2 isoform 2 precursor [Gallus gallus] |
| XP_001633892.1 | 303 | NEMVEDRAFT_v1g102868 | eukaryota>metazoa>cnidaria | Nematostella | vectensis | predicted protein [Nematostella vectensis] |
| XP 016440728.1 | 303 | LOC107766464 | eukaryota>viridiplantae | Nicotiana | tabacum | PREDICTED: calumenin-A-like, partial [Nicotiana tabacum] |
| XP 002162407.1 | 302 | LOC100213173 | eukaryota>metazoa>cnidaria | Hydra | vulgaris | PREDICTED: calumenin-A-like [Hydra vulgaris] |
| XP_002109885.1 | 301 | TRIADDRAFT_20674 | eukaryota>metazoa>placozoa | Trichoplax | adhaerens | hypothetical protein TRIADDRAFT_20674 [Trichoplax adhaerens] |
| XP 003376324.1 | 299 | Tsp 00516 | eukaryota>metazoa>nematoda | Trichinella | spiralis | calumenin [Trichinella spiralis] |
| XP 015758332.1 | 299 | LOC107337629 | eukaryota>metazoa>cnidaria | Acropora | digitifera | PREDICTED: calumenin-like isoform X3 [Acropora digitifera] |
| XP_001639736.1 | 297 | NEMVEDRAFT_v1g86027 | eukaryota>metazoa>cnidaria | Nematostella | vectensis | predicted protein, partial [Nematostella vectensis] |
| XP 002118306.1 | 297 | TRIADDRAFT 33976 | eukaryota>metazoa>placozoa | Trichoplax | adhaerens | hypothetical protein TRIADDRAFT_33976 [Trichoplax adhaerens] |
| XP 015760930.1 | 297 | LOC107340092 | eukaryota>metazoa>cnidaria | Acropora | digitifera | PREDICTED: calumenin-like [Acropora digitifera] |
| KXJ10711.1 | 294 | calua | eukaryota>metazoa>cnidaria | Exaiptasia | pallida | Calumenin-A [Exaiptasia pallida] |
| Smed1000002375 | 294 | Smed1000002375 | eukaryota>metazoa>platyhelminthe | Schmidtea | mediterranea | >Smed1000002375 |
| XP 002155513.3 | 293 | LOC100198810 | eukaryota>metazoa>cnidaria | Hydra | vulgaris | PREDICTED: calumenin-B-like [Hydra vulgaris] |
| XP_018648564.1 | 293 | Smp_005230.1 | eukaryota>metazoa | Schistosoma | mansoni | putative ef hand containing protein [Schistosoma mansoni] |
| CBY20674.1 | 291 | GSOID T00000677001 | eukaryota>metazoa>chordata | Oikopleura | dioica | unnamed protein product [Oikopleura dioica] |
| CDI98589.1 | 288 | EmuJ 000245300 | eukaryota>metazoa | Echinococcus | multilocularis | calumenin [Echinococcus multilocularis] |
| CDS41602.1 | 286 | EmuJ_000926900 | eukaryota>metazoa | Echinococcus | multilocularis | reticulocalbin [Echinococcus multilocularis] |
| NP 491936.3 | 286 | calu-2 | eukaryota>metazoa>nematoda | Caenorhabditis | elegans | CALUmenin (calcium-binding protein) homolog [Caenorhabditis elegans] |
| XP_005589982.1 | 281 | RCN3 | eukaryota>metazoa>chordata>vertebrata | Macaca | fascicularis | PREDICTED: reticulocalbin-3 [Macaca fascicularis] |
| XP_018654312.1 | 281 | Smp_103740.1 | eukaryota>metazoa | Schistosoma | mansoni | putative ef hand containing protein [Schistosoma mansoni] |
| Smed1000012997 | 281 | Smed1000012997 | eukaryota>metazoa>platyhelminthe | Schmidtea | mediterranea | >Smed1000012997 |
| XP_012360521.1 | 280 | RCN1 | eukaryota>metazoa>chordata>vertebrata | Nomascus | leucogenys | PREDICTED: reticulocalbin-1 isoform X1 [Nomascus leucogenys] |
| Caps1000012157 | 272 | Caps1000012157 | eukaryota>metazoa>annelida | Capitella | spI | >Caps1000012157 |
| XP 001623269.1 | 269 | NEMVEDRAFT v1g138173 | eukaryota>metazoa>cnidaria | Nematostella | vectensis | predicted protein, partial [Nematostella vectensis] |
| XP_001634852.1 | 267 | NEMVEDRAFT_v1g184260 | eukaryota>metazoa>cnidaria | Nematostella | vectensis | predicted protein [Nematostella vectensis] |
| XP 003279772.2 | 266 | SDF4 | eukaryota>metazoa>chordata>vertebrata | Nomascus | leucogenys | PREDICTED: 45 kDa calcium-binding protein [Nomascus leucogenys] |
| XP 001626862.1 | 264 | NEMVEDRAFT v1g235717 | eukaryota>metazoa>cnidaria | Nematostella | vectensis | predicted protein [Nematostella vectensis] |
| XP_009016089.1 | 261 | HELRODRAFT_64364 | eukaryota>metazoa>annelida | Helobdella | robusta | hypothetical protein HELRODRAFT_64364 [Helobdella robusta] |
| XP_002410091.1 | 251 | IscW ISCW020366 | eukaryota>metazoa | Ixodes | scapularis | reticulocalbin, putative [Ixodes scapularis] |
| XP 001641982.1 | 250 | NEMVEDRAFT v1g79348 | eukaryota>metazoa>cnidaria | Nematostella | vectensis | predicted protein, partial [Nematostella vectensis] |
| CBY14654.1 | 250 | GSOID T00009701001 | eukaryota>metazoa>chordata | Oikopleura | dioica | unnamed protein product [Oikopleura dioica] |

[illegible]

| | | | | | | |
|---|---|---|---|---|---|---|
| XP_011438715.1 | 1663 | LOC105336190 | eukaryota>metazoa>mollusca | Crassostrea | gigas | PREDICTED: neurabin-1 isoform X5 [Crassostrea gigas] |
| XP_011438716.1 | 1662 | LOC105336190 | eukaryota>metazoa>mollusca | Crassostrea | gigas | PREDICTED: neurabin-1 isoform X6 [Crassostrea gigas] |
| XP_013422095.1 | 1656 | LOC106182023 | eukaryota>metazoa | Lingula | anatina | PREDICTED: uncharacterized protein LOC106182023 isoform X10 [Lingula anatina] |
| XP_011438717.1 | 1650 | LOC105336190 | eukaryota>metazoa>mollusca | Crassostrea | gigas | PREDICTED: neurabin-1 isoform X7 [Crassostrea gigas] |
| XP_013422096.1 | 1626 | LOC106182023 | eukaryota>metazoa | Lingula | anatina | PREDICTED: uncharacterized protein LOC106182023 isoform X11 [Lingula anatina] |
| XP_011667300.1 | 1613 | LOC577610 | eukaryota>metazoa>echinodermata | Strongylocentrotus | purpuratus | PREDICTED: uncharacterized protein LOC577610 isoform X1 [Strongylocentrotus purpuratus] |
| XP_011667301.1 | 1603 | LOC577610 | eukaryota>metazoa>echinodermata | Strongylocentrotus | purpuratus | PREDICTED: uncharacterized protein LOC577610 isoform X2 [Strongylocentrotus purpuratus] |
| XP_011667302.1 | 1600 | LOC577610 | eukaryota>metazoa>echinodermata | Strongylocentrotus | purpuratus | PREDICTED: uncharacterized protein LOC577610 isoform X3 [Strongylocentrotus purpuratus] |
| XP_011667303.1 | 1598 | LOC577610 | eukaryota>metazoa>echinodermata | Strongylocentrotus | purpuratus | PREDICTED: uncharacterized protein LOC577610 isoform X4 [Strongylocentrotus purpuratus] |
| XP_011667304.1 | 1575 | LOC577610 | eukaryota>metazoa>echinodermata | Strongylocentrotus | purpuratus | PREDICTED: uncharacterized protein LOC577610 isoform X5 [Strongylocentrotus purpuratus] |
| XP_011667305.1 | 1567 | LOC577610 | eukaryota>metazoa>echinodermata | Strongylocentrotus | purpuratus | PREDICTED: neurabin-1 isoform X6 [Strongylocentrotus purpuratus] |
| XP_004210881.2 | 1560 | LOC101236396 | eukaryota>metazoa>cnidaria | Hydra | vulgaris | PREDICTED: merozoite surface protein 1-like [Hydra vulgaris] |
| XP_017211649.1 | 1552 | si:ch211-269i23.2 | eukaryota>metazoa>chordata>vertebrata>actinopterygii | Danio | rerio | PREDICTED: neurabin-1 isoform X1 [Danio rerio] |
| XP_011667306.1 | 1543 | LOC577610 | eukaryota>metazoa>echinodermata | Strongylocentrotus | purpuratus | PREDICTED: uncharacterized protein LOC577610 isoform X7 [Strongylocentrotus purpuratus] |
| XP_002122996.1 | 1539 | LOC100179135 | eukaryota>metazoa>chordata | Ciona | intestinalis | PREDICTED: neurabin-1 isoform X1 [Ciona intestinalis] |
| XP_018668829.1 | 1538 | LOC100179135 | eukaryota>metazoa>chordata | Ciona | intestinalis | PREDICTED: neurabin-1 isoform X2 [Ciona intestinalis] |
| XP_008191010.1 | 1531 | LOC664255 | eukaryota>metazoa>hexapoda | Tribolium | castaneum | PREDICTED: uncharacterized protein LOC664255 isoform X1 [Tribolium castaneum] |
| XP_008191012.1 | 1530 | LOC664255 | eukaryota>metazoa>hexapoda | Tribolium | castaneum | PREDICTED: uncharacterized protein LOC664255 isoform X2 [Tribolium castaneum] |
| XP_008191013.1 | 1519 | LOC664255 | eukaryota>metazoa>hexapoda | Tribolium | castaneum | PREDICTED: uncharacterized protein LOC664255 isoform X3 [Tribolium castaneum] |
| XP_008191014.1 | 1519 | LOC664255 | eukaryota>metazoa>hexapoda | Tribolium | castaneum | PREDICTED: uncharacterized protein LOC664255 isoform X4 [Tribolium castaneum] |
| XP_013422097.1 | 1515 | LOC106182023 | eukaryota>metazoa | Lingula | anatina | PREDICTED: neurabin-1-like isoform X12 [Lingula anatina] |
| XP_009290272.1 | 1498 | si:ch211-269i23.2 | eukaryota>metazoa>chordata>vertebrata>actinopterygii | Danio | rerio | PREDICTED: neurabin-1 isoform X2 [Danio rerio] |
| XP_018668830.1 | 1498 | LOC100179135 | eukaryota>metazoa>chordata | Ciona | intestinalis | PREDICTED: neurabin-1 isoform X3 [Ciona intestinalis] |
| XP_017312924.1 | 1495 | LOC108258654 | eukaryota>metazoa>chordata>vertebrata>actinopterygii | Ictalurus | punctatus | PREDICTED: neurabin-1-like isoform X1 [Ictalurus punctatus] |
| XP_009172250.1 | 1487 | T265_08239 | eukaryota>metazoa | Opisthorchis | viverrini | hypothetical protein T265_08239 [Opisthorchis viverrini] |
| XP_009290277.1 | 1482 | si:ch211-269i23.2 | eukaryota>metazoa>chordata>vertebrata>actinopterygii | Danio | rerio | neurabin-1 isoform X1 [Danio rerio] |
| XP_014782972.1 | 1473 | LOC106878301 | eukaryota>metazoa>mollusca | Octopus | bimaculoides | PREDICTED: neurabin-1-like [Octopus bimaculoides] |
| XP_013422098.1 | 1470 | LOC106182023 | eukaryota>metazoa | Lingula | anatina | PREDICTED: neurabin-1-like isoform X13 [Lingula anatina] |
| XP_017312926.1 | 1460 | LOC108258654 | eukaryota>metazoa>chordata>vertebrata>actinopterygii | Ictalurus | punctatus | PREDICTED: neurabin-1-like isoform X2 [Ictalurus punctatus] |
| XP_001849160.1 | 1449 | CpipJ_CPIJ007358 | eukaryota>metazoa>hexapoda | Culex | quinquefasciatus | conserved hypothetical protein [Culex quinquefasciatus] |
| XP_018668831.1 | 1443 | LOC100179135 | eukaryota>metazoa>chordata | Ciona | intestinalis | PREDICTED: neurabin-2 isoform X4 [Ciona intestinalis] |
| XP_011607982.1 | 1435 | LOC101065336 | eukaryota>metazoa>chordata>vertebrata>actinopterygii | Takifugu | rubripes | PREDICTED: neurabin-1-like isoform X1 [Takifugu rubripes] |
| XP_011607987.1 | 1431 | LOC101065336 | eukaryota>metazoa>chordata>vertebrata>actinopterygii | Takifugu | rubripes | PREDICTED: neurabin-1-like isoform X2 [Takifugu rubripes] |
| XP_011607991.1 | 1420 | LOC101065336 | eukaryota>metazoa>chordata>vertebrata>actinopterygii | Takifugu | rubripes | PREDICTED: neurabin-1-like isoform X3 [Takifugu rubripes] |
| KXJ17399.1 | 1420 | PPP1R9A | eukaryota>metazoa>cnidaria | Exaiptasia | pallida | Neurabin-1 [Exaiptasia pallida] |
| XP_011608000.1 | 1416 | LOC101065336 | eukaryota>metazoa>chordata>vertebrata>actinopterygii | Takifugu | rubripes | PREDICTED: neurabin-1-like isoform X5 [Takifugu rubripes] |
| XP_011514682.1 | 1395 | Neurabin-I,NRB1,NRBI | eukaryota>metazoa>chordata>vertebrata | Homo | sapiens | PREDICTED: neurabin-1 isoform X1 [Homo sapiens] |
| XP_011514683.1 | 1395 | Neurabin-I,NRB1,NRBI | eukaryota>metazoa>chordata>vertebrata | Homo | sapiens | PREDICTED: neurabin-1 isoform X1 [Homo sapiens] |
| XP_011514684.1 | 1395 | Neurabin-I,NRB1,NRBI | eukaryota>metazoa>chordata>vertebrata | Homo | sapiens | PREDICTED: neurabin-1 isoform X1 [Homo sapiens] |
| XP_011514685.1 | 1395 | Neurabin-I,NRB1,NRBI | eukaryota>metazoa>chordata>vertebrata | Homo | sapiens | PREDICTED: neurabin-1 isoform X1 [Homo sapiens] |
| XP_017324930.1 | 1389 | LOC108266327 | eukaryota>metazoa>chordata>vertebrata>actinopterygii | Ictalurus | punctatus | PREDICTED: neurabin-1-like isoform X1 [Ictalurus punctatus] |
| XP_017324931.1 | 1389 | LOC108266327 | eukaryota>metazoa>chordata>vertebrata>actinopterygii | Ictalurus | punctatus | PREDICTED: neurabin-1-like isoform X1 [Ictalurus punctatus] |
| XP_003241912.1 | 1381 | LOC100162635 | eukaryota>metazoa>hexapoda | Acyrthosiphon | pisum | PREDICTED: neurabin-1 isoform X1 [Acyrthosiphon pisum] |
| XP_011607996.1 | 1381 | LOC101065336 | eukaryota>metazoa>chordata>vertebrata>actinopterygii | Takifugu | rubripes | PREDICTED: neurabin-1-like isoform X4 [Takifugu rubripes] |
| XP_015303203.1 | 1379 | PPP1R9A | eukaryota>metazoa>chordata>vertebrata | Macaca | fascicularis | PREDICTED: neurabin-1 isoform X1 [Macaca fascicularis] |
| XP_015303204.1 | 1379 | PPP1R9A | eukaryota>metazoa>chordata>vertebrata | Macaca | fascicularis | PREDICTED: neurabin-1 isoform X1 [Macaca fascicularis] |
| XP_015303205.1 | 1379 | PPP1R9A | eukaryota>metazoa>chordata>vertebrata | Macaca | fascicularis | PREDICTED: neurabin-1 isoform X1 [Macaca fascicularis] |
| XP_015303206.1 | 1379 | PPP1R9A | eukaryota>metazoa>chordata>vertebrata | Macaca | fascicularis | PREDICTED: neurabin-1 isoform X1 [Macaca fascicularis] |
| XP_015303207.1 | 1379 | PPP1R9A | eukaryota>metazoa>chordata>vertebrata | Macaca | fascicularis | PREDICTED: neurabin-1 isoform X1 [Macaca fascicularis] |
| XP_016867883.1 | 1377 | Neurabin-I,NRB1,NRBI | eukaryota>metazoa>chordata>vertebrata | Homo | sapiens | PREDICTED: neurabin-1 isoform X2 [Homo sapiens] |
| NP_001159632.1 | 1374 | Neurabin-I,NRB1,NRBI | eukaryota>metazoa>chordata>vertebrata | Homo | sapiens | neurabin-1 isoform 1 [Homo sapiens] |
| XP_003241913.1 | 1374 | LOC100162635 | eukaryota>metazoa>hexapoda | Acyrthosiphon | pisum | PREDICTED: neurabin-1 isoform X2 [Acyrthosiphon pisum] |
| XP_003252562.1 | 1374 | PPP1R9A | eukaryota>metazoa>chordata>vertebrata | Nomascus | leucogenys | PREDICTED: neurabin-1 [Nomascus leucogenys] |
| XP_012365960.1 | 1374 | PPP1R9A | eukaryota>metazoa>chordata>vertebrata | Nomascus | leucogenys | PREDICTED: neurabin-1 [Nomascus leucogenys] |
| XP_017324932.1 | 1374 | LOC108266327 | eukaryota>metazoa>chordata>vertebrata>actinopterygii | Ictalurus | punctatus | PREDICTED: neurabin-1-like isoform X2 [Ictalurus punctatus] |
| XP_016656811.1 | 1373 | LOC100162635 | eukaryota>metazoa>hexapoda | Acyrthosiphon | pisum | PREDICTED: neurabin-1 isoform X3 [Acyrthosiphon pisum] |
| XP_005550235.1 | 1371 | PPP1R9A | eukaryota>metazoa>chordata>vertebrata | Macaca | fascicularis | PREDICTED: neurabin-1 isoform X2 [Macaca fascicularis] |
| [illegible] | [illegible] | [illegible] | [illegible] | [illegible] | [illegible] | [illegible] |

[illegible]

XP 003261542.1 99 S100Z eukaryota>metazoa>chordata>vertebrata Nomascus leucogenys PREDICTED: protein S100 [illegible]
XP_011541541.1 99 Gm625,S100-zeta eukaryota>metazoa>chordata>vertebrata Homo sapiens PREDICTED: protein S100 [illegible]
XP_011541542.1 99 Gm625,S100-zeta eukaryota>metazoa>chordata>vertebrata Homo sapiens PREDICTED: protein S100 [illegible]
XP_011541543.1 99 Gm625,S100-zeta eukaryota>metazoa>chordata>vertebrata Homo sapiens PREDICTED: protein S100 [illegible]
XP_011541546.1 99 Gm625,S100-zeta eukaryota>metazoa>chordata>vertebrata Homo sapiens PREDICTED: protein S100 [illegible]
XP_011541547.1 99 Gm625,S100-zeta eukaryota>metazoa>chordata>vertebrata Homo sapiens PREDICTED: protein S100 [illegible]
XP 015154048.1 99 S100A14 eukaryota>metazoa>chordata>vertebrata Gallus gallus PREDICTED: protein S100 [illegible]
XP 016864660.1 99 Gm625,S100-zeta eukaryota>metazoa>chordata>vertebrata Homo sapiens PREDICTED: protein S100 [illegible]
XP 016864661.1 99 Gm625,S100-zeta eukaryota>metazoa>chordata>vertebrata Homo sapiens PREDICTED: protein S100 [illegible]
XP 016864662.1 99 Gm625,S100-zeta eukaryota>metazoa>chordata>vertebrata Homo sapiens PREDICTED: protein S100 [illegible]
XP 018098399.1 99 LOC108706452 eukaryota>metazoa>chordata>vertebrata Xenopus laevis PREDICTED: protein S100 [illegible]
XP 018115390.1 99 LOC108715051 eukaryota>metazoa>chordata>vertebrata Xenopus laevis PREDICTED: protein S100 [illegible]
XP 018123499.1 99 gm625,s100-zeta,s100z eukaryota>metazoa>chordata>vertebrata Xenopus laevis PREDICTED: S100 calcium [illegible]
CAF96543.1 98 GSTEN:00013622:G:001 eukaryota>metazoa>chordata>vertebrata>actinopterygii Tetraodon nigroviridis unnamed protein product [illegible]
CAG11861.1 98 GSTEN:00033945:G:001 eukaryota>metazoa>chordata>vertebrata>actinopterygii Tetraodon nigroviridis unnamed protein product [illegible]
NP 001019381.1 98 S100A13 eukaryota>metazoa>chordata>vertebrata Homo sapiens protein S100-A13 [Homo [illegible]
NP_001019382.1 98 S100A13 eukaryota>metazoa>chordata>vertebrata Homo sapiens protein S100-A13 [Homo [illegible]
NP 001019383.1 98 S100A13 eukaryota>metazoa>chordata>vertebrata Homo sapiens protein S100-A13 [Homo [illegible]
NP 001019384.1 98 S100A13 eukaryota>metazoa>chordata>vertebrata Homo sapiens protein S100-A13 [Homo [illegible]
NP 001187547.1 98 s10a1 eukaryota>metazoa>chordata>vertebrata>actinopterygii Ictalurus punctatus s100-a1 [Ictalurus punc[illegible]
NP 001289127.1 98 S100Z eukaryota>metazoa>chordata>vertebrata Gallus gallus protein S100-Z [Gallus [illegible]
NP 005970.1 98 S100A13 eukaryota>metazoa>chordata>vertebrata Homo sapiens protein S100-A13 [Homo [illegible]
NP_058020.1 98 cal,Emap1,EMAPI,S100a14,S100c eukaryota>metazoa>chordata>vertebrata Mus musculus protein S100-A11 [Mus m[illegible]
XP 005245491.1 98 S100A13 eukaryota>metazoa>chordata>vertebrata Homo sapiens PREDICTED: protein S100 [illegible]
XP 005541821.1 98 S100A13 eukaryota>metazoa>chordata>vertebrata Macaca fascicularis PREDICTED: protein S100 [illegible]
XP 005541822.1 98 S100A13 eukaryota>metazoa>chordata>vertebrata Macaca fascicularis PREDICTED: protein S100 [illegible]
XP 005541829.1 98 S100A2 eukaryota>metazoa>chordata>vertebrata Macaca fascicularis PREDICTED: protein S100 [illegible]
XP 005541830.1 98 S100A2 eukaryota>metazoa>chordata>vertebrata Macaca fascicularis PREDICTED: protein S100 [illegible]
XP 011508164.1 98 S100A13 eukaryota>metazoa>chordata>vertebrata Homo sapiens PREDICTED: protein S100 [illegible]
XP 011508165.1 98 S100A13 eukaryota>metazoa>chordata>vertebrata Homo sapiens PREDICTED: protein S100 [illegible]
XP 011508166.1 98 S100A13 eukaryota>metazoa>chordata>vertebrata Homo sapiens PREDICTED: protein S100 [illegible]
XP 015132839.1 98 S100Z eukaryota>metazoa>chordata>vertebrata Gallus gallus PREDICTED: protein S100 [illegible]
XP 016857523.1 98 S100A13 eukaryota>metazoa>chordata>vertebrata Homo sapiens PREDICTED: protein S100 [illegible]
XP 016857524.1 98 S100A13 eukaryota>metazoa>chordata>vertebrata Homo sapiens PREDICTED: protein S100 [illegible]
XP 016857525.1 98 S100A13 eukaryota>metazoa>chordata>vertebrata Homo sapiens PREDICTED: protein S100 [illegible]
CAG05262.1 97 GSTEN:00025230:G:001 eukaryota>metazoa>chordata>vertebrata>actinopterygii Tetraodon nigroviridis unnamed protein product [illegible]
NP 001076426.1 97 zgc:171263 eukaryota>metazoa>chordata>vertebrata>actinopterygii Danio rerio ictacalcin 2 [Danio rer[illegible]
NP 002957.1 97 42C,ANX2L,ANX2LG,CAL1L,Ca[1],CLP11,GP11,p10,P11 eukaryota>metazoa>chordata>vertebrata Homo sapiens protein S100-A10 [Homo [illegible]
NP 005969.1 97 CAN19,S100L eukaryota>metazoa>chordata>vertebrata Homo sapiens protein S100-A2 [Homo s[illegible]
NP_033138.1 97 42C,AA409961,AL024248,CAL12,Cal11,CLP11,p10,p11 eukaryota>metazoa>chordata>vertebrata Mus musculus protein S100-A10 [Mus m[illegible]
NP 990837.1 97 P11 eukaryota>metazoa>chordata>vertebrata Gallus gallus protein S100-A10 [Gallu[illegible]
XP 003259351.1 97 S100A2 eukaryota>metazoa>chordata>vertebrata Nomascus leucogenys PREDICTED: protein S100 [illegible]
XP_003259352.1 97 S100A2 eukaryota>metazoa>chordata>vertebrata Nomascus leucogenys PREDICTED: protein S100 [illegible]
XP 003282254.1 97 S100A10 eukaryota>metazoa>chordata>vertebrata Nomascus leucogenys PREDICTED: protein S100 [illegible]
XP 003973567.1 97 s100b eukaryota>metazoa>chordata>vertebrata>actinopterygii Takifugu rubripes PREDICTED: protein S100 [illegible]
XP_005541900.1 97 S100A10 eukaryota>metazoa>chordata>vertebrata Macaca fascicularis PREDICTED: protein S100 [illegible]
XP 005541901.1 97 S100A10 eukaryota>metazoa>chordata>vertebrata Macaca fascicularis PREDICTED: protein S100 [illegible]
XP 012366634.1 97 S100A2 eukaryota>metazoa>chordata>vertebrata Nomascus leucogenys PREDICTED: protein S100 [illegible]
XP 012366636.1 97 S100A2 eukaryota>metazoa>chordata>vertebrata Nomascus leucogenys PREDICTED: protein S100 [illegible]
XP 015135488.1 97 P11 eukaryota>metazoa>chordata>vertebrata Gallus gallus PREDICTED: protein S100 [illegible]
XP 015312662.1 97 S100A10 eukaryota>metazoa>chordata>vertebrata Macaca fascicularis PREDICTED: protein S100 [illegible]
XP 017340854.1 97 LOC108274911 eukaryota>metazoa>chordata>vertebrata>actinopterygii Ictalurus punctatus PREDICTED: protein S100 [illegible]
XP 017347915.1 97 s100z eukaryota>metazoa>chordata>vertebrata>actinopterygii Ictalurus punctatus PREDICTED: protein S100 [illegible]
XP 017347916.1 97 s100z eukaryota>metazoa>chordata>vertebrata>actinopterygii Ictalurus punctatus PREDICTED: protein S100 [illegible]
XP 017347917.1 97 s100z eukaryota>metazoa>chordata>vertebrata>actinopterygii Ictalurus punctatus PREDICTED: protein S100 [illegible]
XP 017347918.1 97 s100z eukaryota>metazoa>chordata>vertebrata>actinopterygii Ictalurus punctatus PREDICTED: protein S100 [illegible]
NP 001013513.1 96 im:7139587,zgc:110771 eukaryota>metazoa>chordata>vertebrata>actinopterygii Danio rerio S100 calcium binding pr[illegible]

[illegible]

XP 006501243.1 93 S100D9 eukaryota>metazoa>chordata>vertebrata [illegible]
XP 006501244.1 93 S100D9 eukaryota>metazoa>chordata>vertebrata [illegible]
XP 011607737.1 93 LOC101065443 eukaryota>metazoa>chordata>vertebrata>actinopterygii [illegible]
XP 015312549.1 93 S100A8 eukaryota>metazoa>chordata>vertebrata [illegible]
XP 017351975.1 93 LOC108280979 eukaryota>metazoa>chordata>vertebrata>actinopterygii [illegible]
NP 001187203.1 92 S10I eukaryota>metazoa>chordata>vertebrata>actinopterygii [illegible]
NP_001270589.1 92 S100B eukaryota>metazoa>chordata>vertebrata [illegible]
NP_001289071.1 92 S100B eukaryota>metazoa>chordata>vertebrata [illegible]
NP_002953.2 92 S100D eukaryota>metazoa>chordata>vertebrata [illegible]
NP_005612.1 92 CAAF1,CAGC,CGRP,ENRAGE,MRP-6,MRP6,p6 eukaryota>metazoa>chordata>vertebrata [illegible]
NP_006263.1 92 NEF,S100,S100-B,S100beta eukaryota>metazoa>chordata>vertebrata [illegible]
NP 033141.1 92 AI850290,Bpb eukaryota>metazoa>chordata>vertebrata [illegible]
NP 989479.1 92 S100A6 eukaryota>metazoa>chordata>vertebrata [illegible]
XP 003259366.1 92 S100A5 eukaryota>metazoa>chordata>vertebrata [illegible]
XP 003277452.1 92 S100B eukaryota>metazoa>chordata>vertebrata [illegible]
XP 003277454.1 92 S100B eukaryota>metazoa>chordata>vertebrata [illegible]
XP 003966278.2 92 LOC101071509 eukaryota>metazoa>chordata>vertebrata>actinopterygii [illegible]
XP_005541836.1 92 S100A5 eukaryota>metazoa>chordata>vertebrata [illegible]
XP_005548529.1 92 S100B eukaryota>metazoa>chordata>vertebrata [illegible]
XP_012357257.1 92 S100B eukaryota>metazoa>chordata>vertebrata [illegible]
XP_015312559.1 92 S100A12 eukaryota>metazoa>chordata>vertebrata [illegible]
XP_016857520.1 92 S100D eukaryota>metazoa>chordata>vertebrata [illegible]
XP 016857521.1 92 S100D eukaryota>metazoa>chordata>vertebrata [illegible]
XP 016883913.1 92 NEF,S100,S100-B,S100beta eukaryota>metazoa>chordata>vertebrata [illegible]
XP 017323558.1 92 LOC108265566 eukaryota>metazoa>chordata>vertebrata>actinopterygii [illegible]
XP 017323574.1 92 LOC108265573 eukaryota>metazoa>chordata>vertebrata>actinopterygii [illegible]
XP 017323586.1 92 LOC108265578 eukaryota>metazoa>chordata>vertebrata>actinopterygii [illegible]
XP 017323594.1 92 LOC108265578 eukaryota>metazoa>chordata>vertebrata>actinopterygii [illegible]
XP_017323603.1 92 LOC108265578 eukaryota>metazoa>chordata>vertebrata>actinopterygii [illegible]
XP_018087243.1 92 LOC108699649 eukaryota>metazoa>chordata>vertebrata [illegible]
XP_018089084.1 92 LOC108700425 eukaryota>metazoa>chordata>vertebrata [illegible]
XP_421894.1 92 S100B eukaryota>metazoa>chordata>vertebrata [illegible]
NP 001313411.1 91 fd10a10,wu:fd10a10 eukaryota>metazoa>chordata>vertebrata>actinopterygii [illegible]
XP 003259339.1 91 S100A12 eukaryota>metazoa>chordata>vertebrata [illegible]
XP 017324242.1 91 LOC108265907 eukaryota>metazoa>chordata>vertebrata>actinopterygii [illegible]
CAG10828.1 90 GSTEN:00032574:G:001 eukaryota>metazoa>chordata>vertebrata>actinopterygii [illegible]
NP 055439.1 90 2A9,5B10,CABP,CACY,PRA eukaryota>metazoa>chordata>vertebrata [illegible]
XP 003259360.1 90 S100A6 eukaryota>metazoa>chordata>vertebrata [illegible]
XP_003966262.1 90 LOC101067460 eukaryota>metazoa>chordata>vertebrata>actinopterygii [illegible]
XP_005541837.1 90 S100A6 eukaryota>metazoa>chordata>vertebrata [illegible]
XP_012366875.1 90 S100A6 eukaryota>metazoa>chordata>vertebrata [illegible]
XP_012366876.1 90 S100A6 eukaryota>metazoa>chordata>vertebrata [illegible]
XP_012366877.1 90 S100A6 eukaryota>metazoa>chordata>vertebrata [illegible]
XP 012366878.1 90 S100A6 eukaryota>metazoa>chordata>vertebrata [illegible]
XP 012366879.1 90 S100A6 eukaryota>metazoa>chordata>vertebrata [illegible]
XP 012366880.1 90 S100A6 eukaryota>metazoa>chordata>vertebrata [illegible]
XP 012366881.1 90 S100A6 eukaryota>metazoa>chordata>vertebrata [illegible]
XP 016857522.1 90 2A9,5B10,CABP,CACY,PRA eukaryota>metazoa>chordata>vertebrata [illegible]
NP 035443.1 89 2A9,5B10,Cacy,CALCYCLIN,PRA eukaryota>metazoa>chordata>vertebrata [illegible]
NP_038678.1 89 60B8Ag,AI323541,B8Ag,Caga,CFAg,CP-10,MRP8,p8 eukaryota>metazoa>chordata>vertebrata [illegible]
CAG10836.1 87 GSTEN:00032585:G:001 eukaryota>metazoa>chordata>vertebrata>actinopterygii [illegible]
NP_001188085.1 87 s10ab eukaryota>metazoa>chordata>vertebrata>actinopterygii [illegible]
XP_003969336.1 86 LOC101064776 eukaryota>metazoa>chordata>vertebrata>actinopterygii [illegible]
CAF87358.1 85 GSTEN:00038360:G:001 eukaryota>metazoa>chordata>vertebrata>actinopterygii [illegible]
NP 001269112.1 85 si:dkey-105n5.1 eukaryota>metazoa>chordata>vertebrata>actinopterygii [illegible]
XP 017207662.1 85 si:dkey-105n5.1 eukaryota>metazoa>chordata>vertebrata>actinopterygii [illegible]
CBY12809.1 81 GSOID T00002868001 eukaryota>metazoa>chordata [illegible]
CBY12423.1 80 GSOID T00001792001 eukaryota>metazoa>chordata [illegible]
NP 004048.1 79 CABP,CABP1,CABP9K,CALB3 eukaryota>metazoa>chordata>vertebrata [illegible]
NP 033919.1 79 AU042539,CaBP-D9K,CABP1,Cabp9k,Calb3 eukaryota>metazoa>chordata>vertebrata [illegible]
XP_003261140.1 79 S100G eukaryota>metazoa>chordata>vertebrata [illegible]
XP_005593108.1 79 S100G eukaryota>metazoa>chordata>vertebrata [illegible]
XP_016885330.1 79 CABP,CABP1,CABP9K,CALB3 eukaryota>metazoa>chordata>vertebrata [illegible]

III. Accession numbers for prokaryotic relatives of proteins and protein families key to calcium storage and signaling, with their phylogenies

A. Accession numbers for selected calmodulin-like proteins with domain fusions

Accession numbers of proteins containing a calmoudlin-like EF-hand fused to some additional domains are provided below in clusters which correspond to the different fused domains discussed. Accession numbers are further annotated by the following columns in order from left to right: length, GenBank gene name, phylogeny, organism, and GenBank defline.

i. Fusions to Tic110-like domains

ADB95776.1 797 UCYN_11030 bacteria>cyanobacteria Candidatus Atelocyanobacterium thalassa isolate ALOHA Ca2+-binding protein [illegible]
KFF41613.1 812 ucyna2 00487 bacteria>cyanobacteria Candidatus Atelocyanobacterium thalassa isolate SIO64986 Ca2+-binding protein [illegible]
ACB51194.1 815 cce 1844 bacteria>cyanobacteria Cyanothece sp. ATCC 51142 unknown [Cyanothece sp[illegible]
ACK71329.1 802 PCC7424 2925 bacteria>cyanobacteria Cyanothece sp. PCC 7424 putative signal transd[illegible]
ACU99706.1 815 Cyan8802 0768 bacteria>cyanobacteria Cyanothece sp. PCC 8802 putative signal transd[illegible]
AFY37740.1 727 Lepto7376 1390 bacteria>cyanobacteria Leptolyngbya sp. PCC 7376 EF hand repeat-contain[illegible]
AOC54009.1 723 amyaer 3304 bacteria>cyanobacteria Microcystis aeruginosa NIES-2481 cAMP-binding protein [illegible]
AKE65596.1 723 MYAER_3258 bacteria>cyanobacteria Microcystis aeruginosa NIES-2549 cAMP-binding protein [illegible]
BAG02189.1 723 MAE_23670 bacteria>cyanobacteria Microcystis aeruginosa NIES-843 hypothetical protein [illegible]
ARI81737.1 723 BH695_2457 bacteria>cyanobacteria Microcystis aeruginosa PCC 7806SL hypothetical protein [illegible]
AKV68896.1 723 VL20_3943 bacteria>cyanobacteria Microcystis panniformis FACHB-1757 cAMP-binding protein [illegible]
ABG51286.1 716 Tery_2045 bacteria>cyanobacteria Trichodesmium erythraeum IMS101 putative signal transd[illegible]
XP_011613778.1 196 cabp1 eukaryota>metazoa>chordata>vertebrata>actinopterygii Takifugu rubripes PREDICTED: calcium-bi[illegible]

ii. Fusions to Heme-Oxygenase domains

ARF79381.1 455 B6264 11010 bacteria>actinobacteria Kitasatospora aureofaciens calcium-binding protein [Kitasatospora aureofaciens]
BAJ33050.1 471 KSE 72950 bacteria>actinobacteria Kitasatospora setae KM-6054 hypothetical protein KSE 72950 [Kitasatospora setae KM-6054]
CCH31773.1 55 BN6_44930 bacteria>actinobacteria Saccharothrix espanaensis DSM 44229 hypothetical protein BN6_44930 [Saccharothrix espanaensis DSM [illegible]

[illegible]

[illegible]

D. ORAI and Jiraiya

[illegible]

V. Alignments

A. Alignment of representative Jiraya and ORAI family members

```
Secondary Structure                   ------HHHHHHHHHHHHHHHHHHHHHHHHHHHHH----H-HHHH-----------------HHH---------HHHH-HHH-HH--HHHHH-HHHHHHHHHHHHHHHH-------------------------------------------H-HHH---HHHHHHHHHHHHHHHHHH-----HHHHHHHHHHHHHHHHHHHHH-H-------------------------------------HHHHHHHHHHHHHHHHHHHHHHHHH-HHHHHHHHH
Hsapiens__NP_001177773.1           1  MARSY-GGRVLAAMTLLGIPAAVLVALAAQLLFQ-----LQAGRAELRGLRGELQGLGAGPGLPEDAAGT------LLPLA-TAL-AA--LVLVLG-LSCLLLAALCGHLGAELAR-------------------------------GP--GPGRSDW-FLYD-CRLLRHVALGLFCCG-----ISVYLAALSIYALLLFEIET------------------GAAAASLSGSGTLVLVAVLTHT-LLRAARARR---GLHE---LSPPSFED------DIARPA---EVSH---ASPRAQPQQGIH  215  Jiraiya
Mmusculus__XP_006509768.1          1  MGRSY-GGRVLAAMTLLGIPAAVLVALAAQLLFQ-----LQAGRAELRRVRTDGLHPE------LDPDAGLPEAAAGA------LLPLA-TAL-AA--LAQVLG-LSCLLLAALCGHLGAELAR-------------------------------GP--GPGRSDW-FLYD-CRLLRHSALGLFCCG-----VSVYLAALAIYALLLFEIEA------------------GAAAASLLGSSGALVLVAVMTHT-LFRAVQATRR---GLHE---LSPPSFED------EPARPS---EDSH---SGCRAQPQPQGTH  215  Jiraiya
Ggallus__XP_015155607.2            1  MP-SY-PQRTLVTLLLFGTLSAAMALLASSLIFQ-----LPXGADLSGSGPGAVRGA------LPESAAV-------------LLPVS-AVL-AA--LCLVLN-VSCLLLCLLHGYFSTELCR-------------------------------GQP--GPERADW-FLLD-SRTVRHAAIGLFCCG-----VSYLTALAIYMLLLFELEA------------------GAAAASLISSGIIVLLVTVTHA-LLRASQLSRR---SRSE---ISHTLYEN------DSAQHS---ETPG---SNLNNKSTAVPR  208  Jiraiya
Drerio__XP_003201087.1             1  MTHSY-SQRSLVILAFLGILSSIMAVLSVILIFQ-----LQTQQAVIKDSPTF----------VPKDITAV------------LLPVS-TVL-SA--LSLTLN-LSSVVVCLLHSYFATEICR-------------------------------GEP--DTQRADW-FLMD-SRAVRHVTIGLFCCG-----VSVYLAAMSIYMVLVFEVET------------------GAAAASITGSSVLLLIIIHS-LLRAARAKRH---HLHTMYHNNP---EQIRTS---NLSIM-ENPRRQHSLANL  208  Jiraiya
Tcastaneum__XP_971722.1          104  SKQVG-VKEMLVSLGLLCLVSLLLALLSLIFLLK-----ISPV-------------------TASDIRELLRAEQLTVITAEEYVV--VYEVT-LAL-CA--LTLSLN-LCCLLVCAIQFLFAVKLVKS--------------------PHG--GNDRTNK-YLQK-SSVSRVCAVGGFFIS----AIVTSVFIGLGIIFCCAMVHN-VFINQR------EKTN---AVKELALA------QHEQSY---HTT---QLNHSHVSHVNH  311  Jiraiya
Lgigantea__XP_009061052.1         76  PADNS-AVKFLYSLAVLCLASLILSALSFQVLIS-----LES--------------------LEDGLKNTSGDGMLNSSAMYSI----VYEAT-TSL-AT--LVIGLD-LSCLMVCSMQCFFTAKILR-------------------------TYE--GSERAFK-YLKE-CSSSRFIAVFSFFFS----IPVYMVALALYLIMKFRSTA------------------ALSCIIICLCIVLFCVLSVMQN-IYHWRV------EKSR---SDEGLPV------DGHVSS---GRTV---SMAVNRHELSTL  279  Jiraiya
Celegans__NP_491975.1             73  MDRKT-VHWNMLLLLFMSVCSALFTVVALQFHFSNAERFLR------------------TAKSPASAYRGV-------------IDVIR-DVI-ASFDVAIVILSILAFFVSALQLLFLMKIVKQT---------------------QQP--DDIDVALRYIKS-TSFLRIVAFSFWFFS----IIVFVIVLMLTVSID----PSRSLAS------------------KCVSITLGSSAFLATCFAVKT-LYFWAN------KSLR---HSCGHVYR------EDCYNY---LSTL---V--------  263  Jiraiya
Hsapiens__NP_116179.2             74  LSWRK-LYLSRAKLKASSRTSALLSGFAMVAMVE-----VQ---------------------LDADHDYPPG------------LLIAF-SAC-TT--VLVAVH-LFALMISTCILPNIEAVSNV------------------------HNL--NSVKESP-HERM-HRHIELAWAFSTVIG-----TLLFLAEVVLLCWVKFLPLKKQPGQPRPTSKPPASGAAANVSTSGITPG-QAAAIASTTIMVPFGLIFIVFA-VHFYRSLVSH---KTDR---QFQELNEL------AEFARL---QDQL---DHRGDHPLTPGS  298  ORAI
Mmusculus__NP_780632.1            76  LSWRK-LYLSRAKLKASSRTSALLSGFAMVAMVE-----VQ---------------------LDTDHDYPPG------------LLIVF-SAC-TT--VLVAVH-LFALMISTCILPNIEAVSNV------------------------HNL--NSVKESP-HERM-HRHIELAWAFSTVIG-----TLLFLAEVVLLCWVKFLPLKRQAGQPSPTKPPAESVIVANHSDSSGITPG-EAAAIASTAIMVPCGLVFIVFA-VHFYRSLVSH---KTDR---QFQELNEL------AEFARL---QDQL---DHRGDHSLTPGT  301  ORAI
Ggallus__NP_001025829.1           11  LSWRK-LYLSRAKLKASSRTSALLSGFAMVAMVE-----VQ---------------------LDAEHDYPPG------------LLIAF-SAC-TT--VLVAVH-LFALMISTCILPNIEAVSNV------------------------HNL--NSVKESP-HERM-HRHIELAWAFSTVIG-----TLLFLAEVVLLCWVKFLPLKKQPGQPTPTKPPAENVIVANHSDSSGITPG-EAAAIASTAIMVPFGLIFIVFA-VHFYRSLVSH---KTDR---QFQELNEL------AEFARL---QDQL---DHRGDSAVPAVT  223  ORAI
Drerio__NP_991163.1               13  LSWRK-LYLSRAKLKASSRTSALLSGFAMVAMVE-----VQ---------------------LDTNHDYPPG------------LLIAF-SAC-TT--VLVAVH-LFALMISTCILPNIEAVSNV------------------------HNL--NSVKESP-HERM-HRHIELAWAFSTVIG-----TLLFLAEVVLLCWVKFLPIKPKDQKNGTVSAG-----------------VAAAITSTSIMVPFGLVFIVFA-VHFYRSLVSH---KTDR---QFQELEEL------EDLQNE---LDHR---EEVSTLQSPGSL  220  ORAI
Tcastaneum__XP_008194887.1       136  LSWRR-LHMSRAKLKATATTSELLSGFAMVAMVE-----LQ---------------------INDDTAVPEW------------LFVIF-AVC-TT--VLVAVH-IFALMISTYILPNIDAVAK------------------------LEIC--EMVAESP-HERM-RGFIELAWAFSTVIG-----LFLFLEVALLCWVKFDVPRHKKDNN--------------------------FTAATAATVIVPVLIIFVLFA-VHFYRSLVSH---KCNA---SVNDMEKL------ENMKKH---LDEV---T--------SPV  323  ORAI
Lgigantea__XP_009053888.1         14  LSWRH-LLLSRTKLKRSSRTSALLSGFAMMAIVE-----LE---------------------LDDDTEVPDN------------LLICF-GVC-TT--LLVFVH-LLALLISTCILPYIEAVSGT------------------------HDI--SMVHDSP-HESM-KWCVELAWISSTGIG-----ILFLAQGLGLLCWVKFHDVS-------------------------------INTAIATCIVIPGAIMFVAFA-LLFYKSLVKH---KNTR---EDELRAIYYNNK--KLLDSSQSLSDN  212  ORAI
Celegans__NP_001254835.1         107  LSEKYNYDLSRAQLKASSRTSALLAGFAMVCLVE-----LQ---------------------YDQSTPKP--------------LLIVL-GVV-TS--LLVSVH-LLALMMSTCILPYMEAT----------------------------GCTQDSP-HIKL-KFYIDLSWLFSTCIG-----LLLFLVEIGVIFYVKFTAVGY-------------------------------PTAGYITTAMLVPVGVVFVVFS-YLIHKNRVSH---SLGR---FKHKVDTM------KQFLDV---EANL---QKSTLAPSTIRD  296  ORAI
Epallida__KXJ12979.1              18  LSWRR-LFLTRGKLKASSRTSGLMSGFAMIAMVE-----IQ---------------------LEKGIPEG--------------LLIAF-SVI-TT--ILISVH-VFALMISVCILPNIESIASLH----------------------HNDFDHSVQDSP-HDKM-HSYIELAWIFSTGIG-----ILFLLAEVLLLCWVKFYNYS-------------------------------HPAAIASMVVLPVLFIVVFVFA-VHFYRSLVSH---KYER---SYQGIEEL------QQIADE---LQTD---NQPSQGPSPSHL  214  ORAI
Tadhaerens__XP_002108336.1         4  IDIAP-VQSTNNIIGSNTTT-----------------N------------------------CPPGHKCVPLY-----------LLVCF-AVL-TT--ILISVH-LFAVMISTCILPHIDAAES-----------------------EGP--SDGMESP-HFEL-HKYIELAWGFSTVIG-----IFLFLCEIIVLCWVKFITID-------------------------------EYAALGTCTMIPVLVVFTVFA-CHFYRRLSRH---EYRR---HSRQINEL------KRMAHA---NLKG---S--GFNAGENTV  182  ORAI
Aqueenslandica__XP_019848729.1     8  SNQLN-CILPVVSLLQVGLV-------------E-----IE---------------------VSPTTPVP--------------LLIIF-AMH-SA--VLVFIH-LLSLMLATFLLPELEAA----------------------------GT--GDISSSL-DIAK-SFTVQLCWVLSNVVG-----ILLLITEIVAFVKFYPTETSEHES-------------------------IHAATATLTFLLFVLCVMAVPLI-IYQSRVVAKH---KLKF---HERGLGRA------RQMLEDM---NFQI---GEGGESSSRDAT  191  ORAI
Srosetta__XP_004993864.1           6  ARRRQ-LALFTHKLEQTGGTSALLSGFAVAMVE-----LN----------------------LPEGRTHVEEFDNNP-------VFYIF-SAV-TS--VLIALH-LFALMVSTSVLPYLKSAIEDIDHHLAQRALTARAKGQALHNLIGPQDTLFDDTTDTSHDN--GLLERVK-HLDQFSSYISKAWFFANGVG-----IAFLLQLALISIYKFLAHT-------------------------------TSSAVAAAVSGYTLSPAFVLFMVFI-YRFYRKIVGSQSTVLKA---TAEQVEQA------ANDYDQ---TNTVAFQLPRIESNSL  249  ORAI
Smoellendorffii__XP_002970256.1   86  KNRRD-VEEKSEQLKAISNLAALFAGFAVVTLTQ-----FT---------------------IAPDPVLT--------------VWIAFYGVL-TA--IAVGLM-TISMVTCTLILGSILKNGKSYVNEEAEEEFMFKCRDFVKSYRTGDKP-----------PSP--RRTFEAF-WEIR-CEDDWKRAFQFFAFGRFLNSRVLSFLFSTLIGWIKYNSE---------------LTAGLFMGVIAVSILVWGFVQ-------------YLWGGYLT---KTFRYN---RTSI---AESGSSHAASIS  298  ORAI
Vcarteri__XP_002953952.1          81  RNRAE-VQEKAEQLKALSALAALLAGFALVRYVEHRVRVLGTHAVHDPC--------------ADDNRRYPKCSFLEFQFVVDANYNRALLPLF-ALT-TS--ITVGVE-VAAVVLCSLMLASIMRIGKMYVNEQEEAEFMHRCQIFITQYRP------------PAP--SRTFSAY-WAHR-CEKEWSTAFRLFSVGALGTLLSFLLAIWAFAFTNKRD--------------LLSEQEAA----WNLT-HRKWGR--------------------CVRR---RMFYQFPLPVAP  320  ORAI
Tpseudonana__XP_002287927.1       38  DARRA-VDEKAQQLKVLANQSALVAGFSMVVLVE-----SN---------------------IPPDINGA--------------LLTLF-GGV-TA--CVIALM-LVSTLNATYMLVAILRYDC------------------------VNR--EVPFDEF-WRKR-CEPDWKMALRTFSVGIPNLLLLFAVIAVFAVVEHKLAAG----------------------------YYSASVVTAISVMVLILWFGST-ERKWGEFLRM---SDAR---ILTRGMSF------EGHGDP---TKQS---QHDGNGGTAMRV  232  ORAI
Pinfestans__XP_002898394.1        62  SEHLV-CEARAEQLTSVSEQATLLCGFIVSAMCN-----VG---------------------VPDNVHAY--------------VLFLY-TVSGTA--VCVSML-ICAVMCTNLQLAVTRYAAHTLEDSVRTMDMTQ----------------LEW--GSPFSSW-WLTR-CEKEQMTAYKFMLFG-----VASFFFYLGAVSWIQFYMST--------------------------GTSLSISSLSFVGFLI-WQLRI-ASKWRYLLTPPSLGTSA---NNLGASHLSHSYLSAPRTPA---SLTL---RKSALEPTAAPV  276  ORAI
Tgondii__XP_002366727.1           20  VQHME-VEYLVMVMNKITSLASLLSGFAASSFLQATG--LQ---------------------RGESWIKL--------------IYVLF-TSS-A-----LGFM-LLVLLICTLCTMWGPGLAL-------------------------RGSKDESLQTAV-NIMD-SS--VQATFRLFSQ-----LACYMLSSILSMVLL---Q-------------------------------PTLSSLVSVTIISVGGVFICRAGLHVVEQLSPKY---VTPA----TICGDPLLFA------SRRVET---EKTA---EMEGSGSALPPA  212  ORAI
Slemnae__CDW77297.1               20  IREKE-LNYWTTTFQTMASQSALLAG------------------------------------NQILLFNL-------------IGFCY-GGL-S-----VPIN-EK---ISSFCVMFGPGLALR------------------------GGE--GAVSMHK-AVEN-MKKESNQCFIFFLQ-----LLFFHVSSFLLMWLV---YP-------------------------------FKVALIININILLVFLVFVKNG-YEIYDKLYKV---EDEAVTGQFQDFGMY----ENMHDL---DRKR---SLPSSPSKRIRN  197  ORAI
Ehuxleyi__XP_005777770.1          20  LRQKE-LDYYVERYSNIATQASIVAGFAFDSLVE-----LE---------------------ISDDMRAALAPGGRWKAELRW-----METVY-YGS-AS--LTMALA-LYTVCVASFATVYGHRLAL-------------------QGPT--GSVERAV-AVMM-KQ--RHSIFVMYGLS----LSVLVAAITMSWIK----M-------------------------------APSWGETIVSGIFLALLL-GLVRK-HQQMKALFRI---PQEH---MVQGDVRLFAG----AEHVDV---SRLE---AGFGEAASAPPG  222  ORAI
consensus/100%                        ............h...s......................................................h..h.....s.....h.......s.............................................p.......hh....hl.............................ss.......h...........................................................
consensus/95%                         ............h...s......................................................s...........l..h.....s....h.hh....s.h.s.h.........................................p.......hh....hl.............................ss.......h...........................................................
consensus/90%                         ........b....h...s..u.hhs.......p.........................s............lb.h..ss..ss..h.lsl..h.shhhssh........h..........................................p.......hh....ah..h.hh.hl........................................................................................
consensus/85%                         ........b....h...s..u.lhs.h.......p.......h...............s.s..........lh.h..ss..ss..l.lsl..l.shhhssh.......h..........................s...............p.......hh.....hahh.h.lh.hlb...............................su...s..h.l.hh...h.h.....................................
consensus/80%                         .pb..h.b.....l...u..uslhushuh..hhp.....lp.................hs.s...s...............lh.hh.ssh.ss..l.lsl..l.shlhssh........hs.........................ss..abb..pp..b.shh....shG.........hahh.l.lh.hlbF...............uu..h..s.h.l.hhhh...s.h.bh.p............p..p.......p.........
consensus/75%                         .sbp..h.b....L...u..uulhuuhuh..hhp.....lp.................hs.s...s...............lh.hh.ssh.ss..lhlsl..lhslhlsshhh.......hsp......................ss.bps..abb..pp..b.shh....shG.........hahh.l.lh.hlbF...............uu..h..s.hl.hhhhhh.s.h.bh.ph.........p....bh.......p..p......
consensus/70%                         hsbp..h.b....pLp..u..uulluuhuh.hhhp.....lp.................ls.s...s...............lh.hh.ssh.ss..lhlsl..lhslhlsshhh.....lsp.................ss.bps..abbb.pp..b.shh....shG.........lalh.lhlhsalbFb..............uu..h..s.hl.hhhhhhbs.h.bhpph.........p.......p..bh.......p...ppp...p......
```

VI. List of completely sequenced curated eukaryotic genomes used for the analysis.

| TaxID | Species.name | Taxonomy |
|---|---|---|
| 5855 | Plasmodium vivax | Alveolata; Apicomplexa; Aconoidasida; Haemosporida; Plasmodiidae; Plasmodium; Plasmodium (Plasmodium) |
| 73239 | Plasmodium yoelii yoelii | Alveolata; Apicomplexa; Aconoidasida; Haemosporida; Plasmodiidae; Plasmodium; Plasmodium (Vinckeia); Plasmodium yoelii |
| 36329 | Plasmodium falciparum 3D7 | Alveolata; Apicomplexa; Aconoidasida; Haemosporida; Plasmodiidae; Plasmodium; Plasmodium (Laverania); Plasmodium falciparum |
| 5865 | Babesia bovis | Alveolata; Apicomplexa; Aconoidasida; Piroplasmida; Babesiidae; Babesia |
| 5866 | Babesia bigemina | Alveolata; Apicomplexa; Aconoidasida; Piroplasmida; Babesiidae; Babesia |
| 1133968 | Babesia microti strain RI | Alveolata; Apicomplexa; Aconoidasida; Piroplasmida; Babesiidae; Babesia; Babesia microti |
| 5875 | Theileria parva | Alveolata; Apicomplexa; Aconoidasida; Piroplasmida; Theileriidae; Theileria |
| 353154 | Theileria annulata strain Ankara | Alveolata; Apicomplexa; Aconoidasida; Piroplasmida; Theileriidae; Theileria; Theileria annulata |
| 1537102 | Theileria equi strain WA | Alveolata; Apicomplexa; Aconoidasida; Piroplasmida; Theileriidae; Theileria; Theileria equi |
| 869250 | Theileria orientalis strain Shintoku | Alveolata; Apicomplexa; Aconoidasida; Piroplasmida; Theileriidae; Theileria; Theileria orientalis |
| 353152 | Cryptosporidium parvum Iowa II | Alveolata; Apicomplexa; Conoidasida; Coccidia; Eucoccidiorida; Eimeriorina; Cryptosporidiidae; Cryptosporidium; Cryptosporidium parvum |
| 5802 | Eimeria tenella | Alveolata; Apicomplexa; Conoidasida; Coccidia; Eucoccidiorida; Eimeriorina; Eimeriidae; Eimeria |
| 99158 | Hammondia hammondi | Alveolata; Apicomplexa; Conoidasida; Coccidia; Eucoccidiorida; Eimeriorina; Sarcocystidae; Hammondia |
| 572307 | Neospora caninum Liverpool | Alveolata; Apicomplexa; Conoidasida; Coccidia; Eucoccidiorida; Eimeriorina; Sarcocystidae; Neospora; Neospora caninum |
| 508771 | Toxoplasma gondii ME49 | Alveolata; Apicomplexa; Conoidasida; Coccidia; Eucoccidiorida; Eimeriorina; Sarcocystidae; Toxoplasma; Toxoplasma gondii |
| 441375 | Cryptosporidium muris RN66 | Alveolata; Apicomplexa; Conoidasida; Coccidia; Eucoccidiorida; Eimeriorina; Cryptosporidiidae; Cryptosporidium; Cryptosporidium muris |
| 110365 | Gregarina niphandrodes | Alveolata; Apicomplexa; Conoidasida; Gregarinasina; Eugregarinorida; Gregarinidae; Gregarina |
| 5888 | Paramecium tetraurelia | Alveolata; Ciliophora; Intramacronucleata; Oligohymenophorea; Peniculida; Parameciidae; Paramecium |
| 5932 | Ichthyophthirius multifiliis | Alveolata; Ciliophora; Intramacronucleata; Oligohymenophorea; Hymenostomatida; Ophryoglenina; Ichthyophthirius |
| 312017 | Tetrahymena thermophila SB210 | Alveolata; Ciliophora; Intramacronucleata; Oligohymenophorea; Hymenostomatida; Tetrahymenina; Tetrahymenidae; Tetrahymena; Tetrahymena thermophila |
| 423536 | Perkinsus marinus ATCC 50983 | Alveolata; Perkinsea; Perkinsida; Perkinsidae; Perkinsus; Perkinsus marinus |
| 370355 | Entamoeba invadens IP1 | Amoebozoa; Archamoebae; Entamoebidae; Entamoeba; Entamoeba invadens |
| 370354 | Entamoeba dispar SAW760 | Amoebozoa; Archamoebae; Entamoebidae; Entamoeba; Entamoeba dispar |
| 1076696 | Entamoeba nuttalli P19 | Amoebozoa; Archamoebae; Entamoebidae; Entamoeba; Entamoeba nuttalli |
| 294381 | Entamoeba histolytica HM-1:IMSS | Amoebozoa; Archamoebae; Entamoebidae; Entamoeba; Entamoeba histolytica |
| 1410327 | Acytostelium subglobosum LB1 | Amoebozoa; Mycetozoa; Dictyosteliida; Acytostelium; Acytostelium subglobosum |
| 5786 | Dictyostelium purpureum | Amoebozoa; Mycetozoa; Dictyosteliida; Dictyostelium |
| 44689 | Dictyostelium discoideum | Amoebozoa; Mycetozoa; Dictyosteliida; Dictyostelium |
| 461836 | Thecamonas trahens ATCC 50062 | Apusozoa; Apusomonadidae; Thecamonas; Thecamonas trahens |
| 2898 | Cryptomonas paramecium | Cryptophyta; Cryptomonadales; Cryptomonadaceae; Cryptomonas |
| 464988 | Hemiselmis andersenii | Cryptophyta; Cryptomonadales; Hemiselmidaceae; Hemiselmis |
| 905079 | Guillardia theta CCMP2712 | Cryptophyta; Pyrenomonadales; Geminigeraceae; Guillardia; Guillardia theta |
| 55529 | Guillardia theta | Cryptophyta; Pyrenomonadales; Geminigeraceae; Guillardia |
| 157538 | Leptomonas pyrrhocoris | Euglenozoa; Kinetoplastida; Trypanosomatidae; Leishmaniinae; Leptomonas |
| 420245 | Leishmania braziliensis MHOM/BR/75/M2904 | Euglenozoa; Kinetoplastida; Trypanosomatidae; Leishmaniinae; Leishmania; Viannia; Leishmania braziliensis species complex; Leishmania braziliensis |
| 71804 | Trypanosoma grayi | Euglenozoa; Kinetoplastida; Trypanosomatidae; Trypanosoma; Trypanosoma with unspecified subgenus |
| 185431 | Trypanosoma brucei brucei TREU927 | Euglenozoa; Kinetoplastida; Trypanosomatidae; Trypanosoma; Trypanozoon; Trypanosoma brucei; Trypanosoma brucei brucei |
| 679716 | Trypanosoma brucei gambiense DAL972 | Euglenozoa; Kinetoplastida; Trypanosomatidae; Trypanosoma; Trypanozoon; Trypanosoma brucei; Trypanosoma brucei gambiense |
| 435258 | Leishmania infantum JPCM5 | Euglenozoa; Kinetoplastida; Trypanosomatidae; Leishmaniinae; Leishmania; Leishmania; Leishmania donovani species complex; Leishmania infantum |
| 5693 | Trypanosoma cruzi | Euglenozoa; Kinetoplastida; Trypanosomatidae; Trypanosoma; Schizotrypanum |
| 5679 | Leishmania panamensis | Euglenozoa; Kinetoplastida; Trypanosomatidae; Leishmaniinae; Leishmania; Viannia; Leishmania guyanensis species complex |
| 929439 | Leishmania mexicana MHOM/GT/2001/U1103 | Euglenozoa; Kinetoplastida; Trypanosomatidae; Leishmaniinae; Leishmania; Leishmania; Leishmania mexicana species complex; Leishmania mexicana |
| 5661 | Leishmania donovani | Euglenozoa; Kinetoplastida; Trypanosomatidae; Leishmaniinae; Leishmania; Leishmania; Leishmania donovani species complex |
| 347515 | Leishmania major strain Friedlin | Euglenozoa; Kinetoplastida; Trypanosomatidae; Leishmaniinae; Leishmania; Leishmania; Leishmania major species complex; Leishmania major |
| 184922 | Giardia lamblia ATCC 50803 | Fornicata; Diplomonadida; Hexamitidae; Giardiinae; Giardia; Giardia intestinalis |
| 280463 | Emiliania huxleyi CCMP1516 | Haptophyceae; Isochrysidales; Noelaerhabdaceae; Emiliania; Emiliania huxleyi |
| 5762 | Naegleria gruberi | Heterolobosea; Schizopyrenida; Vahlkampfiidae; Naegleria |
| 431895 | Monosiga brevicollis MX1 | Opisthokonta; Choanoflagellida; Craspedida; Salpingoecidae; Monosiga; Monosiga brevicollis |
| 946362 | Salpingoeca rosetta | Opisthokonta; Choanoflagellida; Craspedida; Salpingoecidae; Salpingoeca |
| 645134 | Spizellomyces punctatus DAOM BR117 | Opisthokonta; Fungi; Chytridiomycota; Chytridiomycetes; Spizellomycetales; Spizellomycetaceae; Spizellomyces; Spizellomyces punctatus |
| 684364 | Batrachochytrium dendrobatidis JAM81 | Opisthokonta; Fungi; Chytridiomycota; Chytridiomycetes; Rhizophydiales; Rhizophydiales incertae sedis; Batrachochytrium; Batrachochytrium dendrobatidis |
| 1328760 | Xylona heveae TC161 | Opisthokonta; Fungi; Dikarya; Ascomycota; saccharomyceta; Pezizomycotina; leotiomyceta; Xylonomycetes; Xylonomycetales; Xylonomycetaceae; Xylona; Xylona heveae |
| 756982 | Arthrobotrys oligospora ATCC 24927 | Opisthokonta; Fungi; Dikarya; Ascomycota; saccharomyceta; Pezizomycotina; Orbiliomycetes; Orbiliales; Orbiliaceae; Arthrobotrys; Arthrobotrys oligospora |
| 39416 | Tuber melanosporum | Opisthokonta; Fungi; Dikarya; Ascomycota; saccharomyceta; Pezizomycotina; Pezizomycetes; Pezizales; Tuberaceae; Tuber |
| 237561 | Candida albicans SC5314 | Opisthokonta; Fungi; Dikarya; Ascomycota; saccharomyceta; Saccharomycotina; Saccharomycetes; Saccharomycetales; Debaryomycetaceae; Candida/Lodderomyces clade; Candida; Candida albicans |
| 367110 | Neurospora crassa OR74A | Opisthokonta; Fungi; Dikarya; Ascomycota; saccharomyceta; Pezizomycotina; leotiomyceta; sordariomyceta; Sordariomycetes; Sordariomycetidae; Sordariales; Sordariaceae; Neurospora; Neurospora crassa |
| 306902 | Clavispora lusitaniae ATCC 42720 | Opisthokonta; Fungi; Dikarya; Ascomycota; saccharomyceta; Saccharomycotina; Saccharomycetes; Saccharomycetales; Metschnikowiaceae; Clavispora; Clavispora lusitaniae |
| 683960 | Wickerhamomyces anomalus NRRL Y-366-8 | Opisthokonta; Fungi; Dikarya; Ascomycota; saccharomyceta; Saccharomycotina; Saccharomycetes; Saccharomycetales; Phaffomycetaceae; Wickerhamomyces; Wickerhamomyces anomalus |
| 763406 | Pichia membranifaciens NRRL Y-2026 | Opisthokonta; Fungi; Dikarya; Ascomycota; saccharomyceta; Saccharomycotina; Saccharomycetes; Saccharomycetales; Pichiaceae; Pichia; Pichia membranifaciens |
| 559292 | Saccharomyces cerevisiae S288c | Opisthokonta; Fungi; Dikarya; Ascomycota; saccharomyceta; Saccharomycotina; Saccharomycetes; Saccharomycetales; Saccharomycetaceae; Saccharomyces; Saccharomyces cerevisiae |
| 796027 | Sugiyamaella lignohabitans | Opisthokonta; Fungi; Dikarya; Ascomycota; saccharomyceta; Saccharomycotina; Saccharomycetes; Saccharomycetales; Trichomonascaceae; Sugiyamaella |
| 28985 | Kluyveromyces lactis | Opisthokonta; Fungi; Dikarya; Ascomycota; saccharomyceta; Saccharomycotina; Saccharomycetes; Saccharomycetales; Saccharomycetaceae; Kluyveromyces |
| 1170230 | Penicillium digitatum Pd1 | Opisthokonta; Fungi; Dikarya; Ascomycota; saccharomyceta; Pezizomycotina; leotiomyceta; Eurotiomycetes; Eurotiomycetidae; Eurotiales; Aspergillaceae; Penicillium; Penicillium digitatum |
| 306901 | Chaetomium globosum CBS 148.51 | Opisthokonta; Fungi; Dikarya; Ascomycota; saccharomyceta; Pezizomycotina; leotiomyceta; sordariomyceta; Sordariomycetes; Sordariomycetidae; Sordariales; Chaetomiaceae; Chaetomium; Chaetomium globosum |
| 339724 | Histoplasma capsulatum NAm1 | Opisthokonta; Fungi; Dikarya; Ascomycota; saccharomyceta; Pezizomycotina; leotiomyceta; Eurotiomycetes; Eurotiomycetidae; Onygenales; Ajellomycetaceae; Histoplasma; Histoplasma capsulatum |
| 1043004 | Aureobasidium namibiae CBS 147.97 | Opisthokonta; Fungi; Dikarya; Ascomycota; saccharomyceta; Pezizomycotina; leotiomyceta; dothideomyceta; Dothideomycetes; Dothideomycetidae; Dothideales; Aureobasidiaceae; Aureobasidium; Aureobasidium namibiae |
| 284591 | Yarrowia lipolytica CLIB122 | Opisthokonta; Fungi; Dikarya; Ascomycota; saccharomyceta; Saccharomycotina; Saccharomycetes; Saccharomycetales; Dipodascaceae; Yarrowia; Yarrowia lipolytica |
| 227321 | Aspergillus nidulans FGSC A4 | Opisthokonta; Fungi; Dikarya; Ascomycota; saccharomyceta; Pezizomycotina; leotiomyceta; Eurotiomycetes; Eurotiomycetidae; Eurotiales; Aspergillaceae; Aspergillus; Aspergillus nidulans |
| 4896 | Schizosaccharomyces pombe | Opisthokonta; Fungi; Dikarya; Ascomycota; Taphrinomycotina; Schizosaccharomycetes; Schizosaccharomycetales; Schizosaccharomycetaceae; Schizosaccharomyces |
| 402676 | Schizosaccharomyces japonicus yFS275 | Opisthokonta; Fungi; Dikarya; Ascomycota; Taphrinomycotina; Schizosaccharomycetes; Schizosaccharomycetales; Schizosaccharomycetaceae; Schizosaccharomyces; Schizosaccharomyces japonicus |
| 698492 | Saitoella complicata NRRL Y-17804 | Opisthokonta; Fungi; Dikarya; Ascomycota; Taphrinomycotina; Taphrinomycotina incertae sedis; Saitoella; Saitoella complicata |
| 671144 | Wallemia mellicola CBS 633.66 | Opisthokonta; Fungi; Dikarya; Basidiomycota; Agaricomycotina; Wallemiomycetes; Wallemiales; Wallemiales incertae sedis; Wallemia; Wallemia mellicola |
| 879819 | Cutaneotrichosporon oleaginosus | Opisthokonta; Fungi; Dikarya; Basidiomycota; Agaricomycotina; Tremellomycetes; Trichosporonales; Trichosporonaceae; Cutaneotrichosporon |
| 235443 | Cryptococcus neoformans var. grubii H99 | Opisthokonta; Fungi; Dikarya; Basidiomycota; Agaricomycotina; Tremellomycetes; Tremellales; Cryptococcaceae; Cryptococcus; Cryptococcus neoformans species complex; Cryptococcus neoformans; Cryptococcus neoformans var. grubii |
| 597362 | Agaricus bisporus var. burnettii JB137-S8 | Opisthokonta; Fungi; Dikarya; Basidiomycota; Agaricomycotina; Agaricomycetes; Agaricomycetidae; Agaricales; Agaricaceae; Agaricus; Agaricus bisporus; Agaricus bisporus var. burnettii |
| 240176 | Coprinopsis cinerea okayama7#130 | Opisthokonta; Fungi; Dikarya; Basidiomycota; Agaricomycotina; Agaricomycetes; Agaricomycetidae; Agaricales; Psathyrellaceae; Coprinopsis; Coprinopsis cinerea |
| 4999 | Kockovaella imperatae | Opisthokonta; Fungi; Dikarya; Basidiomycota; Agaricomycotina; Tremellomycetes; Tremellales; Cuniculitremaceae; Kockovaella |
| 936046 | Agaricus bisporus var. bisporus H97 | Opisthokonta; Fungi; Dikarya; Basidiomycota; Agaricomycotina; Agaricomycetes; Agaricomycetidae; Agaricales; Agaricaceae; Agaricus; Agaricus bisporus; Agaricus bisporus var. bisporus |
| 486041 | Laccaria bicolor S238N-H82 | Opisthokonta; Fungi; Dikarya; Basidiomycota; Agaricomycotina; Agaricomycetes; Agaricomycetidae; Agaricales; Tricholomataceae; Laccaria; Laccaria bicolor |
| 561896 | Postia placenta Mad-698-R | Opisthokonta; Fungi; Dikarya; Basidiomycota; Agaricomycotina; Agaricomycetes; Agaricomycetes incertae sedis; Polyporales; Coriolaceae; Postia; Postia placenta |
| 764103 | Mixia osmundae IAM 14324 | Opisthokonta; Fungi; Dikarya; Basidiomycota; Pucciniomycotina; Mixiomycetes; Mixiales; Mixiaceae; Mixia; Mixia osmundae |
| 578459 | Rhodotorula graminis WP1 | Opisthokonta; Fungi; Dikarya; Basidiomycota; Pucciniomycotina; Microbotryomycetes; Sporidiobolales; Sporidiobolaceae; Rhodotorula; Rhodotorula graminis |
| 418459 | Puccinia graminis f. sp. tritici CRL 75-36-700-3 | Opisthokonta; Fungi; Dikarya; Basidiomycota; Pucciniomycotina; Pucciniomycetes; Pucciniales; Pucciniaceae; Puccinia; Puccinia graminis; Puccinia graminis f. sp. tritici |
| 77020 | Malassezia pachydermatis | Opisthokonta; Fungi; Dikarya; Basidiomycota; Ustilaginomycotina; Malasseziomycetes; Malasseziales; Malasseziaceae; Malassezia |
| 237631 | Ustilago maydis 521 | Opisthokonta; Fungi; Dikarya; Basidiomycota; Ustilaginomycotina; Ustilaginomycetes; Ustilaginales; Ustilaginaceae; Ustilago; Ustilago maydis |
| 1037660 | Tilletiaria anomala UBC 951 | Opisthokonta; Fungi; Dikarya; Basidiomycota; Ustilaginomycotina; Exobasidiomycetes; Georgefischeriales; Tilletiariaceae; Tilletiaria; Tilletiaria anomala |
| 284813 | Encephalitozoon cuniculi GB-M1 | Opisthokonta; Fungi; Microsporidia; Apansporoblastina; Unikaryonidae; Encephalitozoon; Encephalitozoon cuniculi |
| 578460 | Nosema ceranae BRL01 | Opisthokonta; Fungi; Microsporidia; Apansporoblastina; Nosematidae; Nosema; Nosema ceranae |
| 481877 | Enterocytozoon bieneusi H348 | Opisthokonta; Fungi; Microsporidia; Apansporoblastina; Enterocytozoonidae; Enterocytozoon; Enterocytozoon bieneusi |
| 993615 | Vittaforma corneae ATCC 50505 | Opisthokonta; Fungi; Microsporidia; Apansporoblastina; Nosematidae; Vittaforma; Vittaforma corneae |
| 1485682 | Mitosporidium daphniae | Opisthokonta; Fungi; Microsporidia; Microsporidia incertae sedis; Mitosporidium |
| 881290 | Nematocida parisii ERTm1 | Opisthokonta; Fungi; Microsporidia; Microsporidia incertae sedis; Nematocida; Nematocida parisii |
| 1354746 | Ordospora colligata OC4 | Opisthokonta; Fungi; Microsporidia; Microsporidia incertae sedis; Ordosporidae; Ordospora; Ordospora colligata |
| 948595 | Vavraia culicis subsp. floridensis | Opisthokonta; Fungi; Microsporidia; Pansporoblastina; Pleistophoridae; Vavraia; Vavraia culicis |
| 64571 | Lobosporangium transversale | Opisthokonta; Fungi; Mucoromycota; Mortierellomycotina; Mortierellales; Mortierellaceae; Lobosporangium |
| 763407 | Phycomyces blakesleeanus NRRL 1555(-) | Opisthokonta; Fungi; Mucoromycota; Mucoromycotina; Mucorales; Phycomycetaceae; Phycomyces; Phycomyces blakesleeanus |
| 7955 | Danio rerio | Opisthokonta; Metazoa; Eumetazoa; Bilateria; Deuterostomia; Chordata; Craniata; Vertebrata; Gnathostomata; Teleostomi; Euteleostomi; Actinopterygii; Actinopteri; Neopterygii; Teleostei; Osteoglossocephalai; Clupeocephala; Otomorpha; Ostariophysi; Otophysi; Cypriniphysae; Cypriniformes; Cyprinoidea; Cyprinidae; Danio |
| 7719 | Ciona intestinalis | Opisthokonta; Metazoa; Eumetazoa; Bilateria; Deuterostomia; Chordata; Tunicata; Ascidiacea; Enterogona; Phlebobranchia; Cionidae; Ciona |
| 7668 | Strongylocentrotus purpuratus | Opisthokonta; Metazoa; Eumetazoa; Bilateria; Deuterostomia; Echinodermata; Eleutherozoa; Echinozoa; Echinoidea; Euechinoidea; Echinacea; Echinoida; Strongylocentrotidae; Strongylocentrotus |
| 7998 | Ictalurus punctatus | Opisthokonta; Metazoa; Eumetazoa; Bilateria; Deuterostomia; Chordata; Craniata; Vertebrata; Gnathostomata; Teleostomi; Euteleostomi; Actinopterygii; Actinopteri; Neopterygii; Teleostei; Osteoglossocephalai; Clupeocephala; Otomorpha; Ostariophysi; Otophysi; Characiphysae; Siluriformes; Siluroidei; Ictaluridae; Ictalurus |
| 9031 | Gallus gallus | Opisthokonta; Metazoa; Eumetazoa; Bilateria; Deuterostomia; Chordata; Craniata; Vertebrata; Gnathostomata; Teleostomi; Euteleostomi; Sarcopterygii; Dipnotetrapodomorpha; Tetrapoda; Amniota; Sauropsida; Sauria; Archelosauria; Archosauria; Dinosauria; Saurischia; Theropoda; Coelurosauria; Aves; Neognathae; Galloanserae; Galliformes; Phasianidae; Phasianinae; Gallus |
| 61853 | Nomascus leucogenys | Opisthokonta; Metazoa; Eumetazoa; Bilateria; Deuterostomia; Chordata; Craniata; Vertebrata; Gnathostomata; Teleostomi; Euteleostomi; Sarcopterygii; Dipnotetrapodomorpha; Tetrapoda; Amniota; Mammalia; Theria; Eutheria; Boreoeutheria; Euarchontoglires; Primates; Haplorrhini; Simiiformes; Catarrhini; Hominoidea; Hylobatidae; Nomascus |
| 10090 | Mus musculus | Opisthokonta; Metazoa; Eumetazoa; Bilateria; Deuterostomia; Chordata; Craniata; Vertebrata; Gnathostomata; Teleostomi; Euteleostomi; Sarcopterygii; Dipnotetrapodomorpha; Tetrapoda; Amniota; Mammalia; Theria; Eutheria; Boreoeutheria; Euarchontoglires; Glires; Rodentia; Sciurognathi; Muroidea; Muridae; Murinae; Mus; Mus |
| 9541 | Macaca fascicularis | Opisthokonta; Metazoa; Eumetazoa; Bilateria; Deuterostomia; Chordata; Craniata; Vertebrata; Gnathostomata; Teleostomi; Euteleostomi; Sarcopterygii; Dipnotetrapodomorpha; Tetrapoda; Amniota; Mammalia; Theria; Eutheria; Boreoeutheria; Euarchontoglires; Primates; Haplorrhini; Simiiformes; Catarrhini; Cercopithecoidea; Cercopithecidae; Cercopithecinae; Macaca |
| 10224 | Saccoglossus kowalevskii | Opisthokonta; Metazoa; Eumetazoa; Bilateria; Deuterostomia; Hemichordata; Enteropneusta; Harrimaniidae; Saccoglossus |
| 9606 | Homo sapiens | Opisthokonta; Metazoa; Eumetazoa; Bilateria; Deuterostomia; Chordata; Craniata; Vertebrata; Gnathostomata; Teleostomi; Euteleostomi; Sarcopterygii; Dipnotetrapodomorpha; Tetrapoda; Amniota; Mammalia; Theria; Eutheria; Boreoeutheria; Euarchontoglires; Primates; Haplorrhini; Simiiformes; Catarrhini; Hominoidea; Hominidae; Homininae; Homo |
| 7739 | Branchiostoma floridae | Opisthokonta; Metazoa; Eumetazoa; Bilateria; Deuterostomia; Chordata; Cephalochordata; Branchiostomidae; Branchiostoma |
| 8355 | Xenopus laevis | Opisthokonta; Metazoa; Eumetazoa; Bilateria; Deuterostomia; Chordata; Craniata; Vertebrata; Gnathostomata; Teleostomi; Euteleostomi; Sarcopterygii; Dipnotetrapodomorpha; Tetrapoda; Amphibia; Batrachia; Anura; Pipoidea; Pipidae; Xenopodinae; Xenopus; Xenopus |
| 31033 | Takifugu rubripes | Opisthokonta; Metazoa; Eumetazoa; Bilateria; Deuterostomia; Chordata; Craniata; Vertebrata; Gnathostomata; Teleostomi; Euteleostomi; Actinopterygii; Actinopteri; Neopterygii; Teleostei; Osteoglossocephalai; Clupeocephala; Euteleosteomorpha; Neoteleostei; Eurypterygia; Ctenosquamata; Acanthomorphata; Euacanthomorphacea; Percomorphaceae; Eupercaria; Tetraodontiformes; Tetraodontoidei; Tetraodontoidea; Tetraodontidae; Takifugu |
| 7070 | Tribolium castaneum | Opisthokonta; Metazoa; Eumetazoa; Bilateria; Protostomia; Ecdysozoa; Panarthropoda; Arthropoda; Mandibulata; Pancrustacea; Hexapoda; Insecta; Dicondylia; Pterygota; Neoptera; Endopterygota; Coleoptera; Polyphaga; Cucujiformia; Tenebrionoidea; Tenebrionidae; Tenebrionidae incertae sedis; Tribolium |
| 7091 | Bombyx mori | Opisthokonta; Metazoa; Eumetazoa; Bilateria; Protostomia; Ecdysozoa; Panarthropoda; Arthropoda; Mandibulata; Pancrustacea; Hexapoda; Insecta; Dicondylia; Pterygota; Neoptera; Endopterygota; Amphiesmenoptera; Lepidoptera; Glossata; Neolepidoptera; Heteroneura; Ditrysia; Obtectomera; Bombycoidea; Bombycidae; Bombycinae; Bombyx |
| 7227 | Drosophila melanogaster | Opisthokonta; Metazoa; Eumetazoa; Bilateria; Protostomia; Ecdysozoa; Panarthropoda; Arthropoda; Mandibulata; Pancrustacea; Hexapoda; Insecta; Dicondylia; Pterygota; Neoptera; Endopterygota; Diptera; Brachycera; Muscomorpha; Eremoneura; Cyclorrhapha; Schizophora; Acalyptratae; Ephydroidea; Drosophilidae; Drosophilinae; Drosophilini; Drosophila; Sophophora; melanogaster group; melanogaster subgroup |
| 6850 | Limulus polyphemus | Opisthokonta; Metazoa; Eumetazoa; Bilateria; Protostomia; Ecdysozoa; Panarthropoda; Arthropoda; Chelicerata; Merostomata; Xiphosura; Limulidae; Limulus |
| 37621 | Priapulus caudatus | Opisthokonta; Metazoa; Eumetazoa; Bilateria; Protostomia; Ecdysozoa; Scalidophora; Priapulida; Priapulidae; Priapulus |
| 6412 | Helobdella robusta | Opisthokonta; Metazoa; Eumetazoa; Bilateria; Protostomia; Lophotrochozoa; Annelida; Clitellata; Hirudinida; Hirudinea; Rhynchobdellida; Glossiphoniidae; Helobdella |
| 7574 | Lingula anatina | Opisthokonta; Metazoa; Eumetazoa; Bilateria; Protostomia; Lophotrochozoa; Brachiopoda; Linguliformea; Lingulata; Lingulida; Linguloidea; Lingulidae; Lingula |
| 29159 | Crassostrea gigas | Opisthokonta; Metazoa; Eumetazoa; Bilateria; Protostomia; Lophotrochozoa; Mollusca; Bivalvia; Pteriomorphia; Ostreoida; Ostreoidea; Ostreidae; Crassostrea |
| 37653 | Octopus bimaculoides | Opisthokonta; Metazoa; Eumetazoa; Bilateria; Protostomia; Lophotrochozoa; Mollusca; Cephalopoda; Coleoidea; Neocoleoidea; Octopodiformes; Octopoda; Incirrata; Octopodidae; Octopus |
| 225164 | Lottia gigantea | Opisthokonta; Metazoa; Eumetazoa; Bilateria; Protostomia; Lophotrochozoa; Mollusca; Gastropoda; Patellogastropoda; Lottioidea; Lottiidae; Lottia |
| 7176 | Culex quinquefasciatus | Opisthokonta; Metazoa; Eumetazoa; Bilateria; Protostomia; Ecdysozoa; Panarthropoda; Arthropoda; Mandibulata; Pancrustacea; Hexapoda; Insecta; Dicondylia; Pterygota; Neoptera; Endopterygota; Diptera; Nematocera; Culicomorpha; Culicoidea; Culicidae; Culicinae; Culicini; Culex; Culex; Culex pipiens complex |
| 7460 | Apis mellifera | Opisthokonta; Metazoa; Eumetazoa; Bilateria; Protostomia; Ecdysozoa; Panarthropoda; Arthropoda; Mandibulata; Pancrustacea; Hexapoda; Insecta; Dicondylia; Pterygota; Neoptera; Endopterygota; Hymenoptera; Apocrita; Aculeata; Apoidea; Apidae; Apinae; Apini; Apis |
| 7029 | Acyrthosiphon pisum | Opisthokonta; Metazoa; Eumetazoa; Bilateria; Protostomia; Ecdysozoa; Panarthropoda; Arthropoda; Mandibulata; Pancrustacea; Hexapoda; Insecta; Dicondylia; Pterygota; Neoptera; Paraneoptera; Hemiptera; Sternorrhyncha; Aphidiformes; Aphidomorpha; Aphidoidea; Aphididae; Aphidinae; Macrosiphini; Acyrthosiphon |
| 6334 | Trichinella spiralis | Opisthokonta; Metazoa; Eumetazoa; Bilateria; Protostomia; Ecdysozoa; Nematoda; Enoplea; Dorylaimia; Trichocephalida; Trichinellidae; Trichinella |
| 6279 | Brugia malayi | Opisthokonta; Metazoa; Eumetazoa; Bilateria; Protostomia; Ecdysozoa; Nematoda; Chromadorea; Spirurida; Filarioidea; Onchocercidae; Brugia |
| 610380 | Harpegnathos saltator | Opisthokonta; Metazoa; Eumetazoa; Bilateria; Protostomia; Ecdysozoa; Panarthropoda; Arthropoda; Mandibulata; Pancrustacea; Hexapoda; Insecta; Dicondylia; Pterygota; Neoptera; Holometabola; Hymenoptera; Apocrita; Aculeata; Vespoidea; Formicidae; Ponerinae; Ponerini; Harpegnathos |
| 6239 | Caenorhabditis elegans | Opisthokonta; Metazoa; Eumetazoa; Bilateria; Protostomia; Ecdysozoa; Nematoda; Chromadorea; Rhabditida; Rhabditoidea; Rhabditidae; Peloderinae; Caenorhabditis |
| 6183 | Schistosoma mansoni | Opisthokonta; Metazoa; Eumetazoa; Bilateria; Protostomia; Platyhelminthes; Trematoda; Digenea; Strigeidida; Schistosomatoidea; Schistosomatidae; Schistosoma |
| 6945 | Ixodes scapularis | Opisthokonta; Metazoa; Eumetazoa; Bilateria; Protostomia; Ecdysozoa; Panarthropoda; Arthropoda; Chelicerata; Arachnida; Acari; Parasitiformes; Ixodida; Ixodoidea; Ixodidae; Ixodinae; Ixodes |
| 6198 | Opisthorchis viverrini | Opisthokonta; Metazoa; Eumetazoa; Bilateria; Protostomia; Platyhelminthes; Trematoda; Digenea; Opisthorchiida; Opisthorchiata; Opisthorchiidae; Opisthorchis |
| 45351 | Nematostella vectensis | Opisthokonta; Metazoa; Eumetazoa; Cnidaria; Anthozoa; Hexacorallia; Actiniaria; Edwardsiidae; Nematostella |
| 70779 | Acropora digitifera | Opisthokonta; Metazoa; Eumetazoa; Cnidaria; Anthozoa; Hexacorallia; Scleractinia; Astrocoeniina; Acroporidae; Acropora |
| 6087 | Hydra vulgaris | Opisthokonta; Metazoa; Eumetazoa; Cnidaria; Hydrozoa; Hydroidolina; Anthoathecata; Aplanulata; Hydridae; Hydra |
| 10228 | Trichoplax adhaerens | Opisthokonta; Metazoa; Placozoa; Trichoplax |
| 400682 | Amphimedon queenslandica | Opisthokonta; Metazoa; Porifera; Demospongiae; Heteroscleromorpha; Haplosclerida; Niphatidae; Amphimedon |
| 691883 | Fonticula alba | Opisthokonta; Nucleariidae and Fonticula group; Fonticula |
| 595528 | Capsaspora owczarzaki ATCC 30864 | Opisthokonta; Opisthokonta incertae sedis; Ichthyosporea; Capsaspora; Capsaspora owczarzaki |
| 667725 | Sphaeroforma arctica JP610 | Opisthokonta; Opisthokonta incertae sedis; Ichthyosporea; Ichthyophonida; Sphaeroforma; Sphaeroforma arctica |
| 412133 | Trichomonas vaginalis G3 | Parabasalia; Trichomonadida; Trichomonadidae; Trichomonas; Trichomonas vaginalis |
| 227086 | Bigelowiella natans | Rhizaria; Cercozoa; Chlorarachniophyceae; Bigelowiella |
| 280699 | Cyanidioschyzon merolae strain 10D | Rhodophyta; Bangiophyceae; Cyanidiales; Cyanidiaceae; Cyanidioschyzon; Cyanidioschyzon merolae |
| 130081 | Galdieria sulphuraria | Rhodophyta; Bangiophyceae; Cyanidiales; Cyanidiaceae; Galdieria |
| 2769 | Chondrus crispus | Rhodophyta; Florideophyceae; Gigartinales; Gigartinaceae; Chondrus |
| 556484 | Phaeodactylum tricornutum CCAP 1055/1 | Stramenopiles; Bacillariophyta; Bacillariophyceae; Bacillariophycidae; Naviculales; Phaeodactylaceae; Phaeodactylum; Phaeodactylum tricornutum |
| 296543 | Thalassiosira pseudonana CCMP1335 | Stramenopiles; Bacillariophyta; Coscinodiscophyceae; Thalassiosirophycidae; Thalassiosirales; Thalassiosiraceae; Thalassiosira; Thalassiosira pseudonana |
| 12968 | Blastocystis hominis | Stramenopiles; Blastocystis |
| 944170 | Blastocystis sp. subtype 4 | Stramenopiles; Blastocystis; Blastocystis sp. subtypes |
| 67593 | Phytophthora sojae | Stramenopiles; Oomycetes; Peronosporales; Phytophthora |
| 403677 | Phytophthora infestans T30-4 | Stramenopiles; Oomycetes; Peronosporales; Phytophthora; Phytophthora infestans |
| 761204 | Phytophthora parasitica INRA-310 | Stramenopiles; Oomycetes; Peronosporales; Phytophthora; Phytophthora parasitica |
| 695850 | Saprolegnia parasitica CBS 223.65 | Stramenopiles; Oomycetes; Saprolegniales; Saprolegniaceae; Saprolegnia; Saprolegnia parasitica |
| 157072 | Aphanomyces invadans | Stramenopiles; Oomycetes; Saprolegniales; Saprolegniaceae; Aphanomyces |
| 112090 | Aphanomyces astaci | Stramenopiles; Oomycetes; Saprolegniales; Saprolegniaceae; Aphanomyces |
| 1156394 | Saprolegnia diclina VS20 | Stramenopiles; Oomycetes; Saprolegniales; Saprolegniaceae; Saprolegnia; Saprolegnia diclina |
| 44056 | Aureococcus anophagefferens | Stramenopiles; Pelagophyceae; Pelagomonadales; Aureococcus |
| 3055 | Chlamydomonas reinhardtii | Viridiplantae; Chlorophyta; Chlorophyceae; Chlamydomonadales; Chlamydomonadaceae; Chlamydomonas |
| 3068 | Volvox carteri f. nagariensis | Viridiplantae; Chlorophyta; Chlorophyceae; Chlamydomonadales; Volvocaceae; Volvox; Volvox carteri |
| 145388 | Monoraphidium neglectum | Viridiplantae; Chlorophyta; Chlorophyceae; Sphaeropleales; Selenastraceae; Monoraphidium |
| 70448 | Ostreococcus tauri | Viridiplantae; Chlorophyta; prasinophytes; Mamiellophyceae; Mamiellales; Bathycoccaceae; Ostreococcus |
| 564608 | Micromonas pusilla CCMP1545 | Viridiplantae; Chlorophyta; prasinophytes; Mamiellophyceae; Mamiellales; Mamiellaceae; Micromonas; Micromonas pusilla |
| 296587 | Micromonas commoda | Viridiplantae; Chlorophyta; prasinophytes; Mamiellophyceae; Mamiellales; Mamiellaceae; Micromonas |
| 554065 | Chlorella variabilis | Viridiplantae; Chlorophyta; Trebouxiophyceae; Chlorellales; Chlorellaceae; Chlorella |
| 574566 | Coccomyxa subellipsoidea C-169 | Viridiplantae; Chlorophyta; Trebouxiophyceae; Trebouxiophyceae incertae sedis; Coccomyxaceae; Coccomyxa; Coccomyxa subellipsoidea |
| 3075 | Auxenochlorella protothecoides | Viridiplantae; Chlorophyta; Trebouxiophyceae; Chlorellales; Chlorellaceae; Auxenochlorella |
| 3702 | Arabidopsis thaliana | Viridiplantae; Streptophyta; Streptophytina; Embryophyta; Tracheophyta; Euphyllophyta; Spermatophyta; Magnoliophyta; Mesangiospermae; eudicotyledons; Gunneridae; Pentapetalae; rosids; malvids; Brassicales; Brassicaceae; Camelineae; Arabidopsis |
| 29760 | Vitis vinifera | Viridiplantae; Streptophyta; Streptophytina; Embryophyta; Tracheophyta; Euphyllophyta; Spermatophyta; Magnoliophyta; Mesangiospermae; eudicotyledons; Gunneridae; Pentapetalae; rosids; rosids incertae sedis; Vitales; Vitaceae; Vitis |
| 4577 | Zea mays | Viridiplantae; Streptophyta; Streptophytina; Embryophyta; Tracheophyta; Euphyllophyta; Spermatophyta; Magnoliophyta; Mesangiospermae; Liliopsida; Petrosaviidae; commelinids; Poales; Poaceae; PACMAD clade; Panicoideae; Andropogonodae; Andropogoneae; Tripsacinae; Zea |
| 4097 | Nicotiana tabacum | Viridiplantae; Streptophyta; Streptophytina; Embryophyta; Tracheophyta; Euphyllophyta; Spermatophyta; Magnoliophyta; Mesangiospermae; eudicotyledons; Gunneridae; Pentapetalae; asterids; lamiids; Solanales; Solanaceae; Nicotianoideae; Nicotianeae; Nicotiana |
| 3847 | Glycine max | Viridiplantae; Streptophyta; Streptophytina; Embryophyta; Tracheophyta; Euphyllophyta; Spermatophyta; Magnoliophyta; Mesangiospermae; eudicotyledons; Gunneridae; Pentapetalae; rosids; fabids; Fabales; Fabaceae; Papilionoideae; Phaseoleae; Glycine; Soja |
| 3218 | Physcomitrella patens | Viridiplantae; Streptophyta; Streptophytina; Embryophyta; Bryophyta; Bryophytina; Bryopsida; Funariidae; Funariales; Funariaceae; Physcomitrella |
| 88036 | Selaginella moellendorffii | Viridiplantae; Streptophyta; Streptophytina; Embryophyta; Tracheophyta; Lycopodiidae; Selaginellales; Selaginellaceae; Selaginella |
| 4113 | Solanum tuberosum | Viridiplantae; Streptophyta; Streptophytina; Embryophyta; Tracheophyta; Euphyllophyta; Spermatophyta; Magnoliophyta; Mesangiospermae; eudicotyledons; Gunneridae; Pentapetalae; asterids; lamiids; Solanales; Solanaceae; Solanoideae; Solaneae; Solanum |
| 1169540 | Vitrella brassicaformis CCMP3155 | Alveolata; Chromerida; Vitrella; Vitrella brassicaformis |
| 47911 | Entodinium caudatum | Alveolata; Ciliophora; Intramacronucleata; Litostomatea; Trichostomatia; Entodiniomorphida; Ophryoscolecidae; Entodinium |
| 5949 | Stylonychia lemnae | Alveolata; Ciliophora; Intramacronucleata; Spirotrichea; Stichotrichia; Sporadotrichida; Oxytrichidae; Stylonychinae; Stylonychia |
| 1172189 | Oxytricha trifallax | Alveolata; Ciliophora; Intramacronucleata; Spirotrichea; Stichotrichia; Sporadotrichida; Oxytrichidae; Oxytrichinae; Oxytricha |
| 5963 | Stentor coeruleus | Alveolata; Ciliophora; Postciliodesmatophora; Heterotrichea; Heterotrichida; Stentoridae; Stentor |
| 2951 | Symbiodinium microadriaticum | Alveolata; Dinophyceae; Suessiales; Symbiodiniaceae; Symbiodinium |
| 5759 | Entamoeba histolytica | Amoebozoa; Archamoebae; Entamoebidae; Entamoeba |
| 180228 | Paramoeba pemaquidensis | Amoebozoa; Discosea; Flabellinia; Dactylopodida; Paramoebidae; Paramoeba |
| 65659 | Acanthamoeba comandoni | Amoebozoa; Discosea; Longamoebia; Centramoebida; Acanthamoebidae; Acanthamoeba |
| 361140 | Acytostelium ellipticum | Amoebozoa; Mycetozoa; Dictyosteliida; Acytostelium |
| 133408 | Acytostelium leptosomum | Amoebozoa; Mycetozoa; Dictyosteliida; Acytostelium |
| 670386 | Polysphondylium pallidum PN500 | Amoebozoa; Mycetozoa; Dictyosteliida; Polysphondylium; Polysphondylium pallidum |
| 472931 | Protostelium mycophagum | Amoebozoa; Mycetozoa; Protosteliales; Protosteliaceae; Protostelium |
| 1195612 | Chroomonas mesostigmatica CCMP1168 | Cryptophyta; Pyrenomonadales; Chroomonadaceae; Chroomonas; Chroomonas mesostigmatica |
| 5684 | Leptomonas seymouri | Euglenozoa; Kinetoplastida; Trypanosomatidae; Leishmaniinae; Leptomonas |
| 134006 | Phytomonas sp. isolate EM1 | Euglenozoa; Kinetoplastida; Trypanosomatidae; Phytomonas; unclassified Phytomonas |
| 134013 | Phytomonas sp. isolate Hart1 | Euglenozoa; Kinetoplastida; Trypanosomatidae; Phytomonas; unclassified Phytomonas |
| 59799 | Angomonas deanei | Euglenozoa; Kinetoplastida; Trypanosomatidae; Strigomonadinae; Angomonas |
| 28005 | Strigomonas culicis | Euglenozoa; Kinetoplastida; Trypanosomatidae; Strigomonadinae; Strigomonas |
| 598745 | Giardia intestinalis ATCC 50581 | Fornicata; Diplomonadida; Hexamitidae; Giardiinae; Giardia; Giardia intestinalis |
| 5741 | Giardia intestinalis | Fornicata; Diplomonadida; Hexamitidae; Giardiinae; Giardia |
| 348837 | Spironucleus salmonicida | Fornicata; Diplomonadida; Hexamitidae; Hexamitinae; Spironucleus |
| 765915 | Catenaria anguillulae PL171 | Opisthokonta; Fungi; Blastocladiomycota; Blastocladiomycetes; Blastocladiales; Catenariaceae; Catenaria; Catenaria anguillulae |
| 578462 | Allomyces macrogynus ATCC 38327 | Opisthokonta; Fungi; Blastocladiomycota; Blastocladiomycetes; Blastocladiales; Blastocladiaceae; Allomyces; Allomyces macrogynus |
| 988480 | Rozella allomycis CSF55 | Opisthokonta; Fungi; Cryptomycota; Rozella; Rozella allomycis |
| 743788 | Fomitopsis pinicola FP-58527 SS1 | Opisthokonta; Fungi; Dikarya; Basidiomycota; Agaricomycotina; Agaricomycetes; Agaricomycetes incertae sedis; Polyporales; Fomitopsidaceae; Fomitopsis; Fomitopsis pinicola |
| 1069443 | Mortierella verticillata NRRL 6337 | Opisthokonta; Fungi; Mucoromycota; Mortierellomycotina; Mortierellales; Mortierellaceae; Mortierella; Mortierella verticillata |
| 64571 | Lobosporangium transversale | Opisthokonta; Fungi; Mucoromycota; Mortierellomycotina; Mortierellales; Mortierellaceae; Lobosporangium |
| 747725 | Mucor circinelloides f. lusitanicus CBS 277.49 | Opisthokonta; Fungi; Mucoromycota; Mucoromycotina; Mucorales; Mucorineae; Mucoraceae; Mucor; Mucor circinelloides; Mucor circinelloides f. lusitanicus |
| 246409 | Rhizopus delemar RA 99-880 | Opisthokonta; Fungi; Mucoromycota; Mucoromycotina; Mucorales; Mucorineae; Rhizopodaceae; Rhizopus; Rhizopus delemar |
| 1938954 | Bifiguratus adelaidae | Opisthokonta; Fungi; Mucoromycota; Mucoromycotina; Mucoromycotina incertae sedis; Bifiguratus |
| 1220926 | Mucor circinelloides f. circinelloides 1006PhL | Opisthokonta; Fungi; Mucoromycota; Mucoromycotina; Mucorales; Mucorineae; Mucoraceae; Mucor; Mucor circinelloides; Mucor circinelloides f. circinelloides |
| 86635 | Rhizopus microsporus var microsporus | Opisthokonta; Fungi; Mucoromycota; Mucoromycotina; Mucorales; Mucorineae; Rhizopodaceae; Rhizopus; Rhizopus microsporus |
| 1754190 | Neocallimastix californiae | Opisthokonta; Fungi; Neocallimastigomycota; Neocallimastigomycetes; Neocallimastigales; Neocallimastigaceae; Neocallimastix |
| 1987568 | Pecoramyces ruminatium | Opisthokonta; Fungi; Neocallimastigomycota; Neocallimastigomycetes; Neocallimastigales; Neocallimastigaceae; Pecoramyces |
| 796925 | Conidiobolus coronatus NRRL 28638 | Opisthokonta; Fungi; Zoopagomycota; Entomophthoromycotina; Entomophthoromycetes; Entomophthorales; Ancylistaceae; Conidiobolus; Conidiobolus coronatus |
| 1314790 | Basidiobolus meristosporus CBS 93173 | Opisthokonta; Fungi; Zoopagomycota; Entomophthoromycotina; Basidiobolomycetes; Basidiobolales; Basidiobolaceae; Basidiobolus; Basidiobolus meristosporus |
| 1213189 | Zancudomyces culisetae | Opisthokonta; Fungi; Zoopagomycota; Kickxellomycotina; Harpellales; Legeriomycetaceae; Zancudomyces |
| 34765 | Oikopleura dioica | Opisthokonta; Metazoa; Eumetazoa; Bilateria; Deuterostomia; Chordata; Tunicata; Appendicularia; Oikopleuridae; Oikopleura |
| 99883 | Tetraodon nigroviridis | Opisthokonta; Metazoa; Eumetazoa; Bilateria; Deuterostomia; Chordata; Craniata; Vertebrata; Gnathostomata; Teleostomi; Euteleostomi; Actinopterygii; Actinopteri; Neopterygii; Teleostei; Osteoglossocephalai; Clupeocephala; Euteleosteomorpha; Neoteleostei; Eurypterygia; Ctenosquamata; Acanthomorphata; Euacanthomorphacea; Percomorphaceae; Eupercaria; Tetraodontiformes; Tetraodontoidei; Tetraodontoidea; Tetraodontidae; Tetraodon |
| 6210 | Echinococcus granulosus | Opisthokonta; Metazoa; Eumetazoa; Bilateria; Protostomia; Platyhelminthes; Cestoda; Eucestoda; Cyclophyllidea; Taeniidae; Echinococcus |
| 6211 | Echinococcus multilocularis | Opisthokonta; Metazoa; Eumetazoa; Bilateria; Protostomia; Platyhelminthes; Cestoda; Eucestoda; Cyclophyllidea; Taeniidae; Echinococcus |
| 6669 | Daphnia pulex | Opisthokonta; Metazoa; Eumetazoa; Bilateria; Protostomia; Ecdysozoa; Panarthropoda; Arthropoda; Mandibulata; Pancrustacea; Crustacea; Branchiopoda; Phyllopoda; Diplostraca; Cladocera; Anomopoda; Daphniidae; Daphnia |
| 6257 | Parascaris univalens | Opisthokonta; Metazoa; Eumetazoa; Bilateria; Protostomia; Ecdysozoa; Nematoda; Chromadorea; Ascaridida; Ascaridoidea; Ascarididae; Parascaris |
| 6253 | Ascaris suum | Opisthokonta; Metazoa; Eumetazoa; Bilateria; Protostomia; Ecdysozoa; Nematoda; Chromadorea; Ascaridida; Ascaridoidea; Ascarididae; Ascaris |
| 1720309 | Exaiptasia pallida | Opisthokonta; Metazoa; Eumetazoa; Cnidaria; Anthozoa; Hexacorallia; Actiniaria; Aiptasiidae; Exaiptasia |
| 6136 | Renilla reniformis | Opisthokonta; Metazoa; Eumetazoa; Cnidaria; Anthozoa; Octocorallia; Pennatulacea; Sessiliflorae; Renillidae; Renilla |
| 1819745 | Intoshia linei | Opisthokonta; Metazoa; Mesozoa; Orthonectida; Rhopaluridae; Intoshia |
| 1660239 | Fonticula-like sp. SCN 57-25 | Opisthokonta; Nucleariidae and Fonticula group |
| 1144522 | Tritrichomonas foetus | Parabasalia; Tritrichomonadida; Tritrichomonadidae; Tritrichomonas |
| 641309 | Lotharella oceanica | Rhizaria; Cercozoa; Chlorarachniophyceae; Lotharella |
| 37360 | Plasmodiophora brassicae | Rhizaria; Cercozoa; Imbricatea; Plasmodiophorida; Plasmodiophoridae; Plasmodiophora |
| 46433 | Reticulomyxa filosa | Rhizaria; Foraminifera; Monothalamids; Reticulomyxidae; Reticulomyxa |
| 38544 | Kappaphycus alvarezii | Rhodophyta; Florideophyceae; Gigartinales; Solieriaceae; Kappaphycus |
| 635003 | Fragilariopsis cylindrus CCMP1102 | Stramenopiles; Bacillariophyta; Bacillariophyceae; Bacillariophycidae; Bacillariales; Bacillariaceae; Fragilariopsis; Fragilariopsis cylindrus |
| 183589 | Pseudonitzschia multistriata | Stramenopiles; Bacillariophyta; Bacillariophyceae; Bacillariophycidae; Bacillariales; Bacillariaceae; Pseudo-nitzschia |
| 159749 | Thalassiosira oceanica | Stramenopiles; Bacillariophyta; Coscinodiscophyceae; Thalassiosirophycidae; Thalassiosirales; Thalassiosiraceae; Thalassiosira |
| 72520 | Nannochloropsis gaditana | Stramenopiles; Eustigmatophyceae; Eustigmatales; Monodopsidaceae; Nannochloropsis |
| 672127 | Thraustochytrium sp ATCC 26185 | Stramenopiles; Labyrinthulomycetes; Thraustochytrida; Thraustochytriaceae; Thraustochytrium; unclassified Thraustochytrium |
| 2880 | Ectocarpus siliculosus | Stramenopiles; PX clade; Phaeophyceae; Ectocarpales; Ectocarpaceae; Ectocarpus |
| 42746 | Proteromonas lacertae | Stramenopiles; Slopalinida; Proteromonadidae; Proteromonas |
| 436017 | Ostreococcus lucimarinus CCE9901 | Viridiplantae; Chlorophyta; prasinophytes; Mamiellophyceae; Mamiellales; Bathycoccaceae; Ostreococcus; Ostreococcus 'lucimarinus' |
| 1917417 | Trebouxia sp TZW2008 | Viridiplantae; Chlorophyta; Trebouxiophyceae; Microthamniales; Trebouxia; unclassified Trebouxia photobionts |
| 3175 | Klebsormidium flaccidum | Viridiplantae; Streptophyta; Klebsormidiophyceae; Klebsormidiales; Klebsormidium |
| 232323 | Hypsibius dujardini | Opisthokonta; Metazoa; Eumetazoa; Bilateria; Protostomia; Ecdysozoa; Panarthropoda; Tardigrada; Eutardigrada; Parachela; Hypsibiidae; Hypsibius |
